# In situ single-cell activities of microbial populations revealed by spatial transcriptomics

**DOI:** 10.1101/2021.02.24.432792

**Authors:** Daniel Dar, Nina Dar, Long Cai, Dianne K. Newman

## Abstract

Microbial populations and communities are heterogeneous, yet capturing their diverse activities has proven challenging at the relevant spatiotemporal scales. Here we present par-seqFISH, a targeted transcriptome-imaging approach that records both gene-expression and spatial context within microscale assemblies at a single-cell and molecule resolution. We apply this approach to the opportunistic bacterial pathogen, *Pseudomonas aeruginosa*, analyzing ∼600,000 individuals across dozens of physiological conditions in planktonic and biofilm cultures. We explore the phenotypic landscape of this bacterium and identify metabolic and virulence related cell-states that emerge dynamically during growth. We chart the spatial context of biofilm-related processes including motility and kin-exclusion mechanisms and identify extensive and highly spatially-resolved metabolic heterogeneity. We find that distinct physiological states can co-exist within the same biofilm, just a few microns away, underscoring the importance of the microenvironment. Together, our results illustrate the complexity of microbial populations and present a new way of studying them at high-resolution.

## Introduction

Life exists in context. Cells within microbial populations and communities are typically closely associated with one another in multicellular biofilms, whether found within infected tissues, attached to diverse surfaces, or forming assemblages in the deep sea (Costerton et al., 1987; Flemming and Wuertz, 2019). Natural microbiota and infectious bacteria generally exist in biofilm aggregates that are on the order of several dozen microns and which can contain many interacting species (DePas et al., 2016; Mark Welch et al., 2016; Schaber et al., 2007). Despite the ubiquity of the biofilm lifestyle in both natural and manmade habitats, understanding what life is like within it for individual microbes has proven highly challenging. While single-cell level activities have been tracked at high spatial resolution using a variety of approaches in diverse contexts (Chadwick et al., 2019; Hatzenpichler et al., 2020; Jorth et al., 2019), we have been unable to resolve the hundreds if not thousands of concurrent activities that characterize microbial life at relevant spatiotemporal scales. What we understand about microbial life literally has been limited by our ability to see.

Despite this limitation, it has become clear in recent years that extreme phenotypic heterogeneity defines the microbial experience (Ackermann, 2015; Evans et al., 2020). This is as true for isogenic populations as it is for complex biofilm communities. Clonemates sampled from the same environment often display significant differences that are thought to result from stochastic gene-expression and variable environmental factors (Ackermann, 2015; Schreiber and Ackermann, 2019; Schreiber et al., 2016). The detection of phenotypic diversity even in seemingly well-mixed environments such as chemostats (Kopf et al., 2015; Schreiber et al., 2016) also serves as a powerful reminder that life at the microscale may inhabit far more diverse niches than are readily apparent. Phenotypic diversity has been rationalized as providing microbes with a fitness advantage in an unpredictable world (Ackermann, 2015; Veening et al., 2008). In addition, specialized functions have been proposed to underpin collective interactions such as division of labor (Ackermann, 2015; Armbruster et al., 2019; Diard et al., 2013; Rosenthal et al., 2018). However, little is still known about the range of possible cellular phenotypic states and their roles in most biological processes.

What triggers such phenotypic plasticity, and are there underlying “rules” that govern any patterns that may exist at the microscale? In sessile communities, both clonal or multispecies, biological activities give rise to changing chemical gradients that create a range of local microenvironments (Stewart, 2003; Stewart and Franklin, 2008). Furthermore, spatial organization enables different conflicting metabolic states or species to co-exist via physical separation, increasing the potential for diversity and allowing for new interactions to emerge (Bocci et al., 2018; Evans et al., 2020; Kotte et al., 2014; Nadell et al., 2016; Wolfsberg et al., 2018). Indeed, natural communities often contain many interacting species that assemble into intricate spatial structures. These microscale assemblies can promote interactions between species and represent a key ecosystem feature (Cordero and Datta, 2016; Nadell et al., 2016). Yet a wide gulf— limited by technology—still separates such observations from a coherent conceptual framework to explain the rules governing microbial ecology.

Recent advances in imaging methods provide a means to chart the physical associations between different species in natural environments (Mark Welch et al., 2016; Shi et al., 2020; Tropini et al., 2017; Wilbert et al., 2020). However, interpreting these maps remains challenging without additional functional information on the physiological states and activities of relevant community members. In contrast, recent adaptions of eukaryotic single-cell RNA-sequencing (scRNA-seq) approaches provide a powerful means of exploring the phenotypic landscape of planktonic bacteria (Blattman et al., 2020; Imdahl et al., 2020; Kuchina et al., 2021). However, these approaches do not preserve the spatial context of analyzed cells and are therefore limited in their capacity to address single and multispecies biofilms. Thus, a major gap exists in our ability to account for both spatial and functional complexity, limiting progression toward a high-resolution understanding of microbial life.

Single-molecule fluorescence in situ hybridization (FISH) based technologies have been used to measure gene-expression directly within native tissues, recording both spatial and functional information. However, while these methods have shed important light on single-cell heterogeneity they have been traditionally limited to measuring the expression of only a few genes at a time (Choi et al., 2014; Femino et al., 1998; Raj et al., 2008; So et al., 2011). In addition to this limited throughput, single-gene measurements do not provide a means to capture coordinated cellular responses—the molecular “fingerprint” of multiple biological activities that underpin distinct physiological states. Recent advances in combinatorial mRNA labeling and sequential FISH (seqFISH) now allow for hundreds and even thousands of genes to be analyzed within the same sample at a sub-micron resolution (Chen et al., 2015; Eng et al., 2019; Lubeck et al., 2014). Until now, seqFISH has been used in mammalian systems to expose the physical organization of cell states within tissues (Chen et al., 2015; Eng et al., 2019; Lubeck et al., 2014; Moffitt et al., 2018; Shah et al., 2016). We reasoned that the high spatial resolution of these modern transcriptome-imaging techniques also had the potential to illuminate the microscale organization of microbial populations and communities.

In this study, we adapted and further developed seqFISH for studying bacteria, measuring the expression of hundreds of genes within individual cells while also capturing their spatial context. We utilized *Pseudomonas aeruginosa* planktonic and biofilm populations to demonstrate how different cellular functions are coordinated in time and space. Our proof-of-concept work illustrates how the ability to observe transcriptional activities at the microscale permits insights into the spatiotemporal regulation and coordination of critical life processes, enabling hitherto unrecognized, transient physiological states to be identified and new hypotheses to be generated. These findings represent the tip of the iceberg and the opportunities for discovery our approach enables promise to reveal new insights about the rules governing microbial ecology.

## Results

### A sequential mRNA-FISH framework for studying bacterial gene-expression

Combinatorial mRNA labeling requires that each measured mRNA molecule be individually resolved. However, this is much more challenging in bacteria due to the small size of their cells, as many different mRNA molecules occur in close proximity and cannot be resolved using standard fluorescent microscopy. We therefore utilized a nonbarcoded seqFISH approach (Lignell et al., 2017).

In seqFISH, target mRNAs are first hybridized with a set of primary, non-fluorescent probes, which are flanked by short sequences uniquely assigned per gene (Figure 1A). Specific genes can be turned “ON” via a secondary hybridization with short fluorescently labeled “readout” probes, complementary to the gene-specific flanking sequences (Figure 1A). Several genes can be measured at once using a set of readout probes labeled with different fluorophores (Methods). Importantly, these short fluorescent readout probes can be efficiently stripped and washed away from the sample without affecting the primary probes (Shah et al., 2018) (Figure 1A). Thus, once expression is measured, fluorescence can be turned OFF and a new set of genes can be measured by introducing a new set of readout probes (Figure 1B). This 2-step design allows for potentially hundreds of genes to be measured sequentially, one after the other in the same sample, using automated microscopy (Figure 1B). The individual gene mRNA-FISH data can be combined into spatially resolved multigene profiles at the single-bacterium level (Figure 1B).

**Figure 1.**
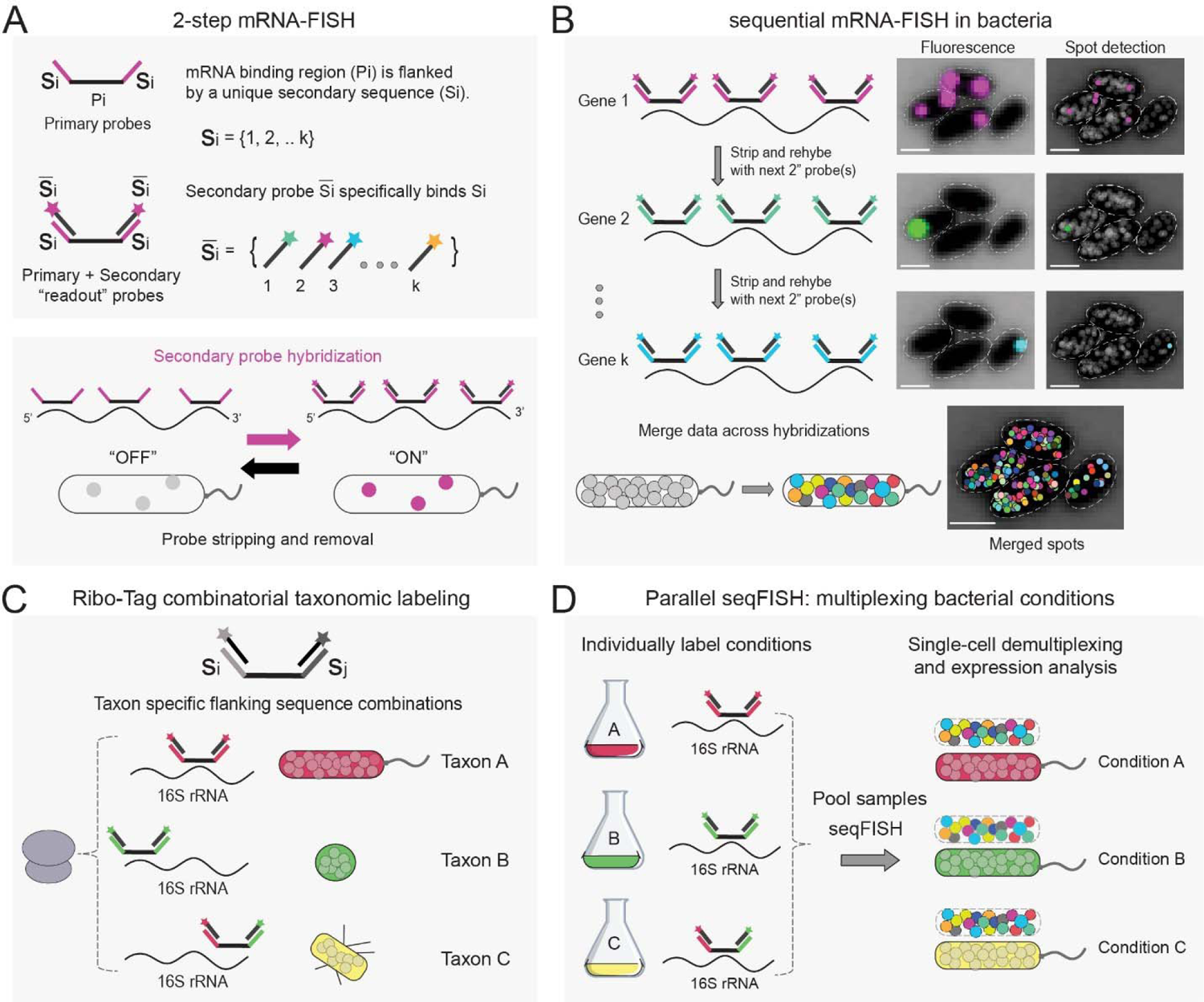
Parallel and sequential mRNA-FISH in bacteria. (A) seqFISH probe design scheme. Primary probes contain unique sequences (Si) that are read by secondary probes (colored wands). Each gene is read by a unique probe and its fluorescence can be turned “ON” or “OFF”. (B) mRNA-FISH applied sequentially to the same sample. In each cycle, a new set of secondary readout probes are introduced. Raw fluorescence data is shown on the right and the detected local spot maxima are shown in the spot detection image. Merged spots for many genes shown in shuffled colors. (C) Combinatorial labeling can be used to encode species taxonomy using 16S rRNA, or to enable the parallel study of (D) bacteria grown in different conditions.

Due to the diffraction limit and the small size of bacteria, mRNA-FISH fluorescent signals (appearing as spots within cells) can contain overlapping mRNA molecules that cannot be spatially resolved in standard microscopes. Thus, counting the number of spots within a bacterial cell severely underestimates expression levels. This problem can be overcome by integrating the fluorescent intensity per spot, which scales linearly with the number of mRNAs. Fluorescent intensity can be converted to discrete mRNA counts by measuring the characteristic intensity of a single transcript. This analog to digital conversion approach has been shown to provide a wide dynamic range in bacteria (Skinner et al., 2013; So et al., 2011).

We developed seqFISH in the study of *Pseudomonas aeruginosa*, an opportunistic human pathogen and a severe cause of morbidity and mortality in cystic fibrosis (CF) patients (Bhagirath et al., 2016; Malhotra et al., 2019). We generated a probe library targeting a set of 105 marker genes that capture many core physiological aspects of this pathogen (Tables S1-S2). These included genes involved in biosynthetic capacity (ribosome and RNA-polymerase subunits), anerobic physiology (fermentation and denitrification pathways), stress responses (oxidative and nutrient limitation), cellular signaling (c-di-GMP), biofilm matrix components, motility (flagella and T4P), all major quorum-sensing (QS) systems, as well as multiple antibiotic resistance and core virulence factors. In addition, to control for false positives, we designed probes targeting three different negative control genes that do not exist in *Pseudomonas* (Figure S1).

### Parallel and sequential mRNA-FISH in single bacterial cells

To test our bacterial seqFISH approach, we first studied *P. aeruginosa* grown in well-understood batch culture conditions. We performed a growth curve experiment in LB medium, where key parameters such as cell density, growth rate, and oxygen levels change in a predictable manner. We collected 11 time points representing the lag phase, exponential growth, and stationary phase and imaged them simultaneously (Figure 2A). Independent imaging of these samples in a serial manner would have taken ∼3 weeks of automated microscopy time.

**Figure 2.**
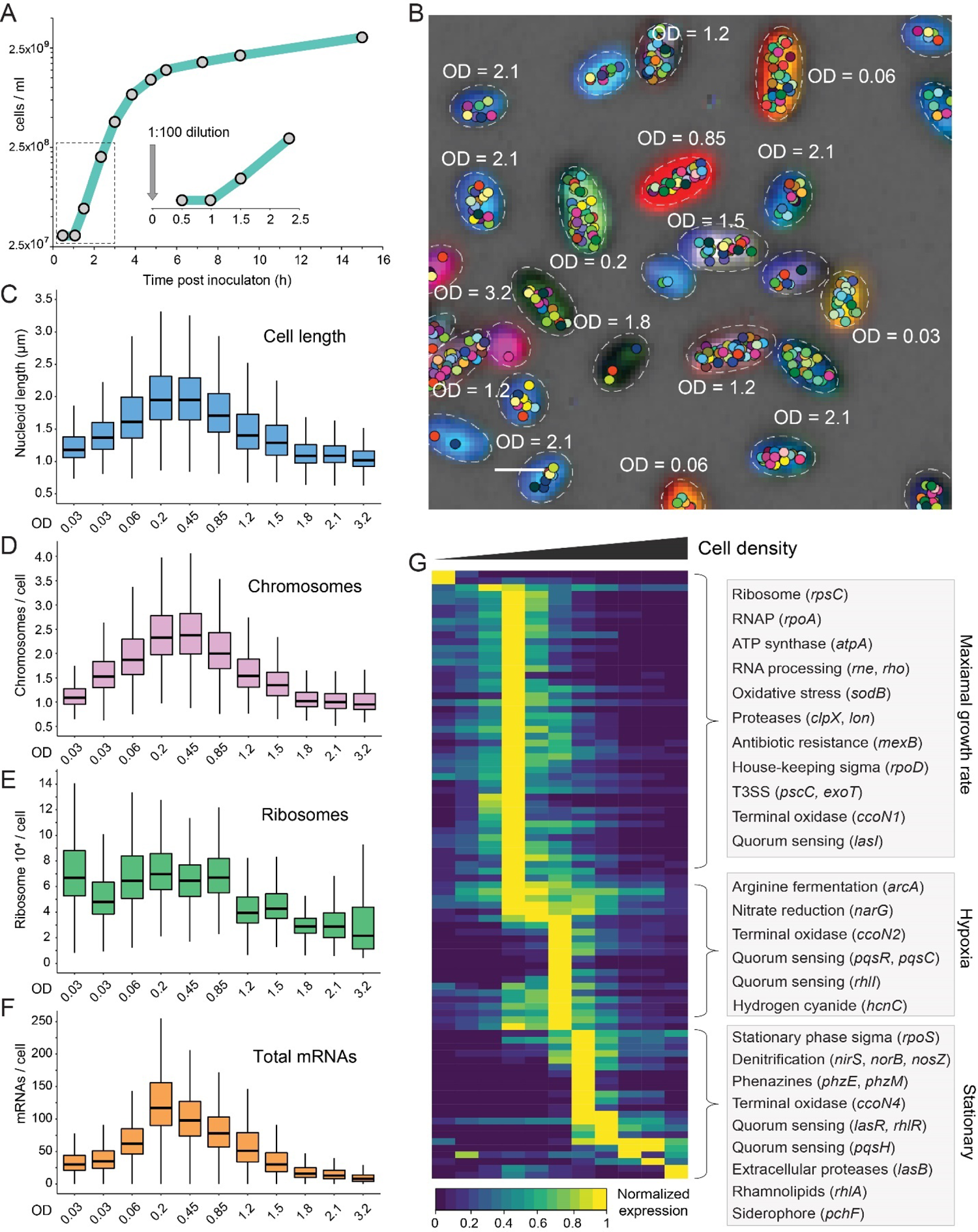
Parallel seqFISH (par-seqFISH) of an LB growth curve experiment. (A) The sampled LB growth curve. Collected time points are indicated with gray circles. A zoom-in shows the sampled lag phase. (B) Demultiplexed bacteria and their mRNAs. The merged, raw Ribo-Tag 16S rRNA fluorescence is shown for a representative region. Different barcodes (16S combinations) result in unique colors that visually report the condition of which they originated from (indicated with the corresponding OD_600_ value). Ellipses fitted to the segmented cell boundaries are shown. The mRNA spots (fitted position of maximal intensity) for all genes per cell are shown in unique colors per gene. Each spot may represent more than one mRNA copy. (C-F) Condition specific distributions of nucleoid length, chromosome copy, ribosome levels and total mRNAs detected across our gene set. (G) Heatmap showing average gene expression normalized to the maximal value for each gene across all conditions. Highlighted gene groups and their functions are indicated on the right.

To perform simultaneous imaging, we developed an efficient multiplexing method that enables parallel seqFISH experiments (par-seqFISH). We designed a set of primary probes targeting the 16S rRNA (Ribo-Tags), which contain unique combinations of flanking sequences (barcodes), that serve as the “readout” in a seqFISH run (Figure 1C-D; Table S3). In principle, this multiplexing approach can be applied to studying combinations of different species (Figure 1C) or for pooling bacteria from different growth conditions (Figure 1D). We validated the latter application by individually labeling the 16S rRNAs of each of the 11 growth curve samples with unique Ribo-Tags. The samples were pooled, collectively hybridized with the 105 gene probe library, and subjected to sequential hybridizations to measure gene-expression and to decode cell identity (Figure 2B). We acquired expression profiles for >50,000 individual *P. aeruginosa* cells, over 91.8% of which were unambiguously decoded and assigned to the condition from which they originated (Figure 2B). We estimate the false positive decoding rate at 0.04% (1 in 2500 cells) by counting the number of hits for barcodes left out of the experiment, demonstrating both high efficiency and accuracy for par-seqFISH.

In addition to acquiring mRNA expression profiles, our imaging-based platform permits concurrent tracking of key information such as cell size and shape, and can be combined with functional stains, markers and/or immunofluorescence measurements (Takei et al., 2021). This opens up the possibility of correlating particular expression profiles at the single cell level with integrative physiological or cell biological parameters. We applied a 4′ diamidino-2-phenylindole (DAPI) stain as a part of the par-, 6-seqFISH experiment and used DAPI fluorescence to estimate the nucleoid size and chromosome copy per cell. Comparing cells at different stages of growth shows that both nucleoid size (estimating cell size) and chromosome number distributions follow identical trends, in agreement with the *P. aeruginosa* literature (Vallet-Gely and Boccard, 2013) (Figure 2C-D). We also estimated ribosome abundance using 16S rRNA fluorescence. Notably, the distribution of this metric differed significantly from that of the chromosome parameters, displaying contrasting intensities at different stages of lag phase, increased variability at deep stationary and a delay in signal decline during the shift from exponential growth to stationary phase (Figure 2E). In contrast, the total number of mRNAs per cell (estimated by our 105 genes) differentiates each time point along the growth curve, reaching a maxima and minima at the fastest and slowest growth rates, respectively (Figure 2F). These data further support the accuracy of our par-seqFISH multiplexing approach and demonstrate the unique ability of this method to integrate single cell gene-expression with global parameters.

To examine whether our expression profiles faithfully capture known physiological processes that occur during culture development, we grouped the cells according to their decoded conditions and calculated their average gene expression profiles. We find a temporally resolved expression pattern associated with different stages of growth (Figure 2G). For example, genes representing high replicative/biosynthetic capacity such as those involved in RNA and protein biosynthesis reach their peak expression during maximal division rate but decreased between 90 to 250-fold in stationary phase (Figure 2G). In contrast, stress factors involved in stationary phase adaptation and nutrient limitation peak at low division rates and higher cell densities (Figure 2G). QS signal production, receptor expression and target activation reflect the known hierarchical QS regulatory network (Lee and Zhang, 2015). Notably, the expression of anaerobic metabolism genes occurred in two stages: early induction of the fermentation and nitrate/nitrite reduction genes in the entry to stationary phase, in which hypoxic conditions emerge, followed by expression of the remaining denitrification pathway at lower predicted oxygen levels (Price-Whelan et al., 2007) (Figure 2G). Furthermore, the shift from aerobic to anaerobic metabolism was accompanied by sequential exchanges in terminal oxidase identities, from *ccoN1* to *ccoN2* and finally, *ccoN4*, concomitantly with the induction of phenazine biosynthesis (Arai, 2011; Jo et al., 2017) (Figure 2G).

Notably, repeated mRNA measurements of the same genes in independent and spaced hybridization rounds were well correlated, both in average expression and at single-bacterium levels (Pearson R = 0.86, 0.89 and 0.9, for *sigX*, *rpsC* and *rpoS*, respectively). In addition, the three negative control genes had an average false positive rate of 0.002 transcripts per cell (Figure S1). Together, these results further validate the accuracy of our multiplexing method and demonstrate that our marker genes capture diverse transcriptional states across a wide range of physiological conditions.

Transient emergence of physiologically distinct sub-populations during LB growth Phenotypic diversity in clonal populations can generate distinct sub-populations that specialize in different tasks at different times, setting a fertile ground for bet-hedging behaviors and complex interactions (Ackermann, 2015; Rosenthal et al., 2018). The single-cell resolution and high sensitivity of seqFISH has the potential to shed light on this important yet largely unexplored aspect of microbial life.

We applied Uniform Manifold Approximation and Projection (UMAP) dimensionality reduction and unsupervised clustering to our single-cell expression data (McInnes et al., 2018). This analysis charted the single-cell phenotypic landscape in LB growth, from the perspective of our marker genes. Analyzing the 11 time points together, we detect 20 sub-populations with diverse predicted functional capabilities. These included among others, differential replicative capacity, exoproduct biosynthesis, and virulence factor production (Figure 3A-B). We find that the sampled populations of most of the growth conditions are partitioned into multiple co-existing sub-groups with distinct expression profiles (Figure 3; Figure S2; Table S4). Notably, our data suggest that the degree of dispersion within this expression space (estimating phenotypic diversity) varies significantly between conditions and is elevated during stationary phase (Figure S2; Table S4).

**Figure 3.**
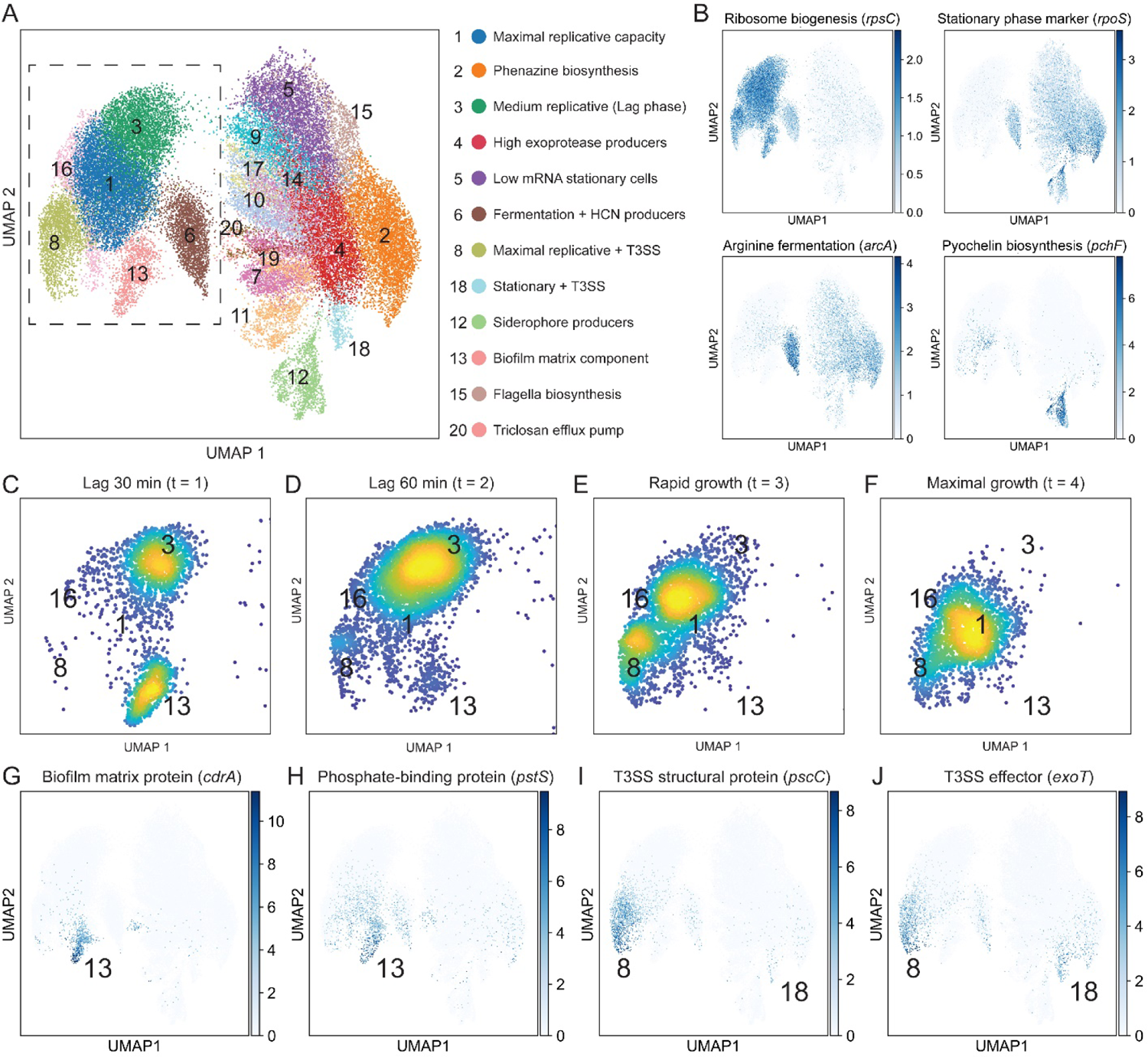
Single-bacterium analysis reveals physiologically distinct dynamic sub-populations. (A) UMAP analysis using cells from all 11 time points. Identified clusters are shown in different colors and are indexed by group size. Specific group and their enriched functions are shown to the right. (B) Gene expression overlays for four genes that report on metabolic state, stationary phase progression and exoproduct biosynthesis. (C-F) Density scatter plots of cells from individual conditions in a zoom-in of the UMAP (dashed box in panel A). The clusters are indicated by their index. (G-J) Gene expression overlays shown as in B and indicated in the figure.

Our growth condition-specific analysis revealed intriguing dynamics during lag phase progression. It could be expected that lag phase cultures will follow a steady ribosome accumulation as the cells progress toward exponential growth and maximal ribosome content (Bosdriesz et al., 2015). In contrast, we found a relative decline in the average ribosomal rRNA levels: early lag phase populations (30 min post dilution) had a higher signal than the more advanced lag culture (60 min post dilution; Figure 2E). These differences appear to be rooted in the transient emergence and disappearance of an early lag sub-population with exceptionally high levels of 16S rRNA (cluster 13; comprising 34.6% of the population in early lag; Figure 3C-F; Figure S3; Table S4). In agreement with the deviation in the rRNA signal, this sub-population also shows a proportional increase in total mRNA counts. However, its size and chromosome copy distributions were not elevated (Figure S3; cluster 13 vs. 3).

Beyond illuminating the extent of heterogeneity in seemingly well mixed cultures and classifying subpopulations into particular types, seqFISH can directly connect global cell-specific parameters such as ribosome levels or cell shape to particular gene-expression signatures. For example, a closer examination of the metabolically hyperactive sub-population revealed a 186-fold enrichment in *cdrA* expression relative to the rest of the population (Figure 3G). The *cdrA* gene encodes a major adhesive protein component of the *P. aeruginosa* biofilm matrix (Borlee et al., 2010; Reichhardt et al., 2018). Expression of *cdrA* is commonly used as a reporter for cyclic diguanylate monophosphate (c-di-GMP) levels, a key signaling molecule involved in surface attachment (Armbruster et al., 2019). In addition, this sub-population also displays a 30-fold enrichment in *pstS* expression, which encodes for the phosphate-binding component of the *pstSCAB* phosphate uptake system (Figure 3H). PstS has been previously detected in extracellular appendages of *P. aeruginosa* and has been suggested to provide an adhesion phenotype to intestinal epithelial cells (Zaborina et al., 2008). In support of this non-canonical role, *pstS* was recently suggested to confer a similar adherence phenotype in *Acinetobacter baumannii*, another human pathogenic bacterium (Gil-Marqués et al., 2020).

A second example from our dataset of the type of fine-grained information seqFISH can provide comes from the temporal expression of genes involved in virulence factor production. Single cell variation in virulence factor production has been suggested as a mechanism for division of labor during infection (Diard et al., 2013). *P. aeruginosa* employs a variety of virulence factors to overcome the host immune response (Malhotra et al., 2019), including the type 3 secretion system (T3SS) that translocate toxins (effectors) directly into host cells (Hauser, 2009). Our gene set monitors two T3SS structural genes (*pscC* and *pcrD*) and two main effectors (*exoT* and *exoY*), all of which are encoded in different operons (Wurtzel et al., 2012). We detected two different types of sub-populations with enriched T3SS related genes, suggesting a unique division of cells into virulent and avirulent states (Figure 3I-J). The first group transiently appears during exponential growth and constitutes 8-30% of the population (Figure 3C-F and 3I-J; Table S4). This group expresses both the secretion system genes (86-fold enrichment) and the effectors (28-fold). In contrast, the second group appears 3-4 divisions later, close to the replicative minima at stationary phase, and occupies only ∼2.7% of cells (Table S4). This sub-population is strongly enriched for the two effectors (average 26-fold; Figure 3I-J) but only mildly so for the secretion system (6-fold), as compared with the earlier group.

We can potentially reconcile these observations as follows: *P. aeruginosa* has been shown to contain approximately 1-3 T3SS units per cell under inducing conditions (Lombardi et al., 2019). Thus, successive divisions following T3SS expression will result in rapid dilution of the T3SS^+^ group. Assuming the inheritance of the T3SS and effectors is uncoupled, then T3SS^+^ stationary phase cells are likely to lose their effectors during division and are predicted to be “inactive”. Thus, an intriguing hypothesis is that *P. aeruginosa* invests in the costly T3SS^+^ sub-population during “times of plenty” (rapid growth) and specifically expresses the effectors at stationary to “reload” and maintain this sub-population following division-based dilution, just prior to growth arrest. Together, these examples underscore the power of seqFISH to suggest hypotheses that can be tested going forward.

### Spatial transcriptomics at a single-cell resolution in *P. aeruginosa* biofilms

Though much can be learned by applying seqFISH to planktonic cultures, in many contexts, bacteria exist in biofilms (Costerton et al., 1987; Flemming and Wuertz, 2019). Variation in local environmental conditions and the effect of spatially confined metabolic activities in biofilm populations can promote the emergence of chemically distinct microenvironments and phenotypes (Evans et al., 2020; Stewart, 2003). We reasoned that seqFISH’s capacity to record transcriptional activities with micron resolution would be particularly useful in shedding light on these processes.

The *P. aeruginosa* biofilm mode of life is particularly important in chronic infections such as those residing in the airways of individuals with CF (Bjarnsholt et al., 2009; Høiby et al., 2011). Accordingly, having used LB to validate bacterial seqFISH, we switched to synthetic cystic fibrosis sputum medium (SCFM) for our biofilm studies (Palmer et al., 2007). Briefly, bacteria were incubated in coverslip attached microwells and the medium was replaced every several hours (Methods). Using biofilms that were allowed to develop for 10 or 35 hours, we imaged hundreds of aggregates ranging in size from several bacteria to tens-of-thousands of tightly bound members (Figure 4A-B). As a reference for cellular physiological states, we also performed a planktonic growth curve experiment in SCFM. We applied par-seqFISH multiplexing to image 10 time points matching those sampled in the planktonic LB experiment (Methods). We extracted the physical coordinates of individual bacterial cells within microaggregates, acquiring a microscale spatial expression profile for ∼365,000 surface attached bacteria (Figure 4A-B). In addition, we collected single-cell expression data for ∼218,000 planktonic cells.

**Figure 4.**
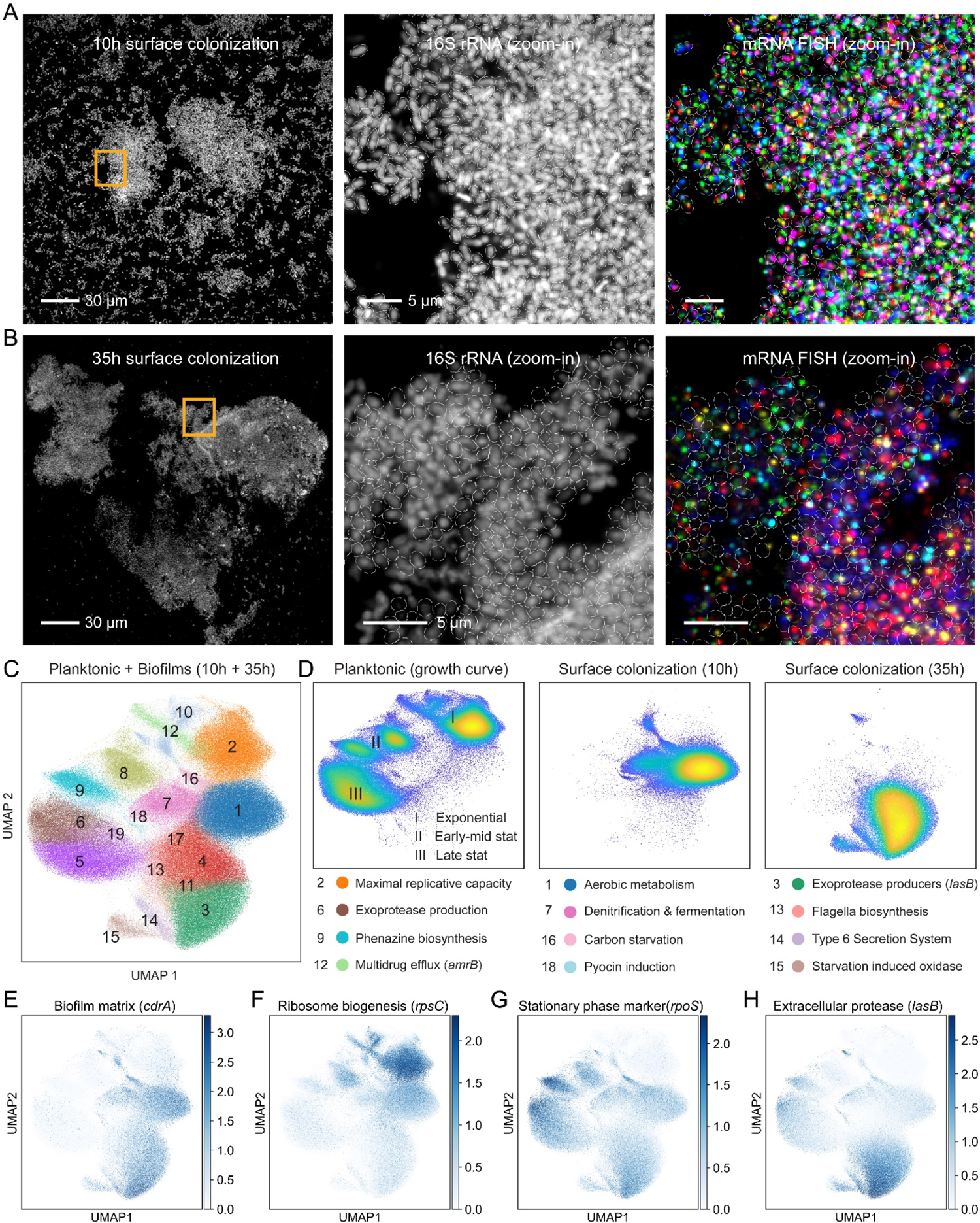
Spatial transcriptomics in *P. aeruginosa* biofilms at a single cell resolution. (A) A representative field of view collected during a 10h surface colonization experiment showing cells via 16S rRNA fluorescence (gray). A zoom-in (orange box) shows the cell segmentation masks depicted as white ellipses. The 16S rRNA signal and mRNA-FISH data for several genes are shown in different colors. (B) A 35h experiment field is shown in an identical manner to panel A. Scale bar length is annotated within the figure. (C) Joint UMAP cluster analysis of biofilm and planktonic experiments. Planktonic cells are shown for all time points collected (D) UMAP scatter plots showing cells from either planktonic or biofilm experiments as indicated. Below, a highlighted set of UMAP clusters associated with each experiment is annotated with enriched functions. (E-H) UMAP overlay with specific gene data.

A basic question we sought to answer was the extent to which transcriptional responses are unique to the biofilm lifestyle. We performed a joint UMAP analysis using both biofilm and planktonic samples (Figure 4C). These different modes of growth cluster into independent groups in expression space, reflecting their significant physiological differences (Figure 4D). Ribosome and RNAP subunit expression in the planktonic experiment correlated strongly with growth rate, as observed in LB (Figure 4D). Examining these marker genes in the biofilm-derived cells places the average replicative capacity of the 10h and 35h biofilm populations at roughly equal to those of early-mid and late stationary planktonic populations, respectively (Figure 4F). Expression of the stationary phase master regulator, *rpoS* further supports this classification (Figure 4G). However, biofilm cells also have unique expression profiles that distinguish them from liquid cultures. For example, the matrix component gene, *cdrA*, is uniformly expressed in both 10 and 35h biofilms but repressed in most planktonic cells (Figure 4E). In addition, compared with stationary liquid cells, our data indicates that early biofilms (10h) have higher expression of *sigX* (5.1-fold), a transcription factor recently implicated in biofilm formation (Gicquel et al., 2013), *mexB* (>4.5-fold), of the *mexA*-*mexB*-*oprM* antibiotic efflux system, and an increase in the 3’-5’ exonuclease, polynucleotide phosphorylase (*pnp*) (7.5-fold). Comparing the 35h biofilm to the stationary cells, we find a 3.3-fold increase in the extracellular protease, *lasB*, but reduced expression of other proteases such *lasA* (3-fold lower), as well as *aprA* and the rhamnolipid biosynthesis gene, *rhlA* (∼10-fold lower). Notably, these genes are quorum-sensing (QS) regulated and our liquid cultures expressed both *lasA* and *rhlA* at later time points than *lasB*, suggesting these differences may reflect the age of the biofilm rather than features that define the biofilm state per *se*.

### In situ analysis of biofilm specific functions

The above data demonstrate that seqFISH can capture both cell states and their physical position directly within intact biofilms, providing an opportunity to examine known and new processes that contribute to biofilm development from a quantitative and highly spatially resolved perspective. To illustrate this, we focused on the expression patterns of representative genes known to define critical stages in biofilm development such as attachment, maturation and exclusion of competitors.

Motility systems such as the flagella and the type 4 pilus (T4P) are major determinant of surface colonization subsequent biofilm formation (Belas, 2014; Burrows, 2012; O’Toole and Kolter, 1998). Recent work identified an asymmetric division process coined “Touch-Seed-and-Go”, in which flagellated mother cells first attach to a surface and then produce un-flagellated daughter cells that contain the T4P. This c-di-GMP dependent phenotypic diversification enables the mother “spreader” cell to spawn multiple adherent “seed” populations (Laventie et al., 2019). This is thought to be mainly regulated by surface sensing (Laventie et al., 2019). However, how such motility-based division of labor affects the organization of biofilms at stages beyond surface attachment remains unknown.

We examined the spatial expression patterns of the major flagellum and T4P components, *fliC* and *pilA*, respectively in the early surface colonization experiment (10h biofilm). An abundant “checkerboard” like pattern is evident, in which cells express high levels of either *fliC* or *pilA* but generally not both (Figure 5A). This pattern is apparent in both small groups (∼tens of cells) and in microaggregates that contain thousands of cells. In contrast, the older 35h biofilms showed lower expression of *pilA* but contained a sparse but uniform distribution of *fliC*^+^ cells, suggesting that biofilm associated bacteria invest in a costly motility apparatus despite being spatially confined (Figure 5B), effectively, the bacterial equivalent of purchasing a sports car during a midlife crisis. Strikingly, examining the expression of *fliC* and *pilA* in our paired planktonic experiment we find a similar mutually exclusive pattern (Figure 5C). Thus, in contrast to the current model, our planktonic control experiment suggests that the asymmetric distribution of motility systems is unlikely to be directly regulated by surface sensing (Figure 5C); such a conclusion would not be possible without the means to compare transcriptional activities at the single cell level.

**Figure 5.**
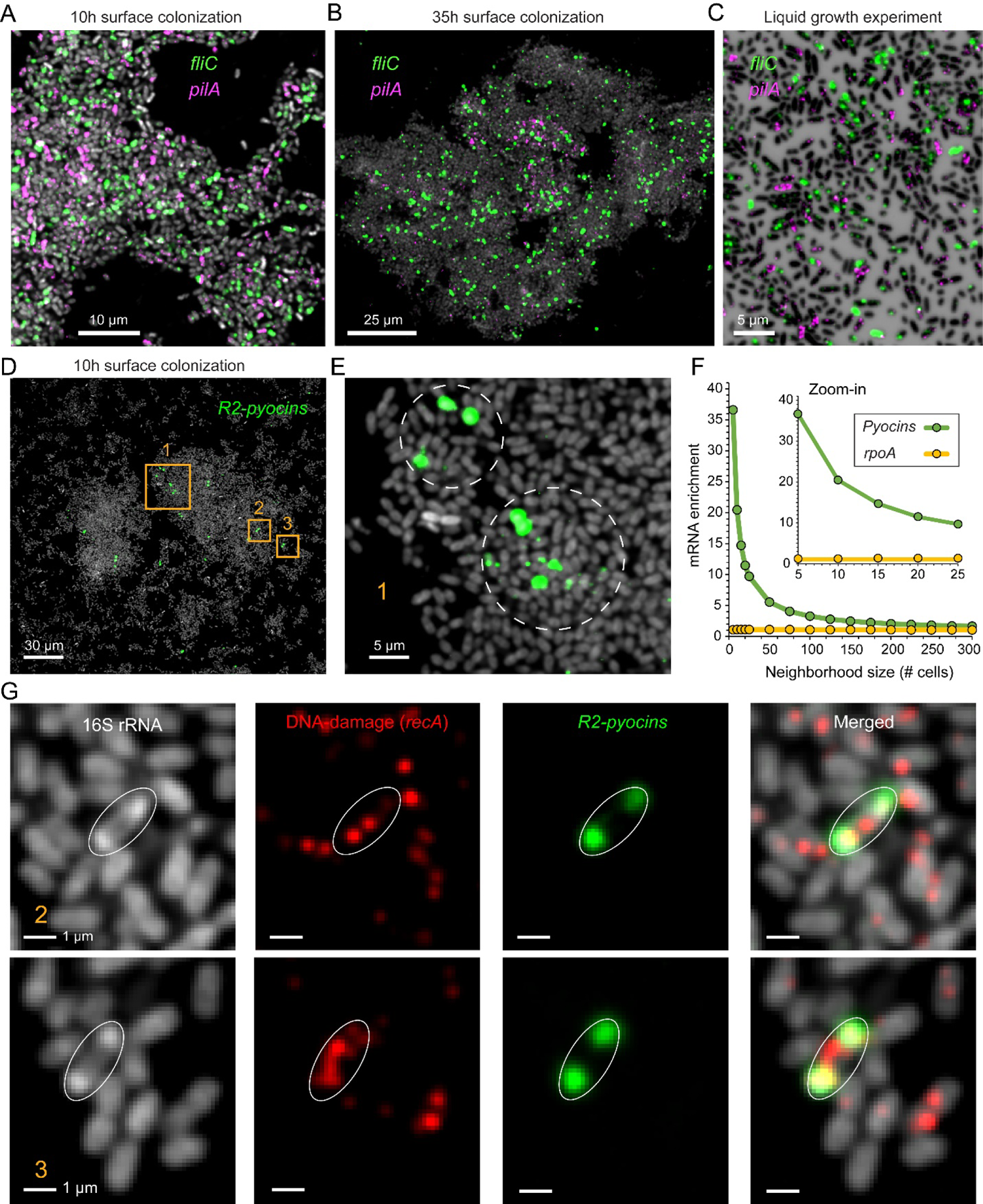
Spatial expression patterns for motility and pyocin related genes. (A-B) Representative regions from the 10h and 35h biofilm experiments, cells are shown in via 16S rRNA fluorescence (gray) and overlayed with raw mRNA-FISH fluorescence for different genes as indicated. (C) planktonic cells from the pair liquid experiments. Cells are shown via DAPI and expression as indicated (D-E) 10h aggregate showing R2-pyocin expression. (F) Enrichment of R2-Pyocin mRNA near strong induction sites (cell with 99.5^th^ percentile pyocin expression). X-axis shows the number of cells closest to an induction site that were analyzed (neighborhood size; center cell was excluded). Y-axis shows the enrichment in each neighborhood relative to the total population. A non-pyocin control gene is shown (*rpoA*). (G) Examples of mRNA R-pyocin transcript and ribosome polar localization as indicated in the legends.

Beyond initial surface attachment, bacteria must establish a strong foothold for colony development as well as outcompete resident microbes. One strategy that potentially address both needs is the utilization of phage tail-like bacteriocins, broadly called tailocins (Ghequire and De Mot, 2015). These elements are thought to be adapted from prophages and are applied as narrow-spectrum toxins for kin exclusion (Bobay et al., 2014; Ghequire and De Mot, 2015). However, in contrast with antibiotics, these phage tail-like structures are released into the environment via explosive lysis events that kill the producer and spray the toxin locally to inhibit nearby competitors (Turnbull et al., 2016; Vacheron et al., 2021). This event also releases extracellular DNA that integrates into the biofilm matrix, structurally supporting biofilm maturation (Turnbull et al., 2016; Whitchurch et al., 2002). Yet how this “sacrificial” process is regulated within developing biofilms is not well understood. Our UMAP analysis identified a sub-population (cluster 18; Figure 4C) exhibiting >1000-fold enrichment in expression of the R2-pyocin operon (*Pseudomonas* tailocin), represented by the *PA14_08150* gene.

This UMAP cluster was enriched ∼4-fold in 10h biofilm derived cells, suggesting pyocin induction is upregulated during surface attachment. Furthermore, we find an 11-fold higher expression of the DNA-repair gene, *recA*, in agreement with its role in inducing pyocin expression (Brazas and Hancock, 2005). Visualizing the expression of the pyocin producers, we find that induction events are spread across various microaggregates regions but often appear in local clusters (Figure 5D-E). Indeed, we find a ∼37-fold average spatial enrichment in pyocin expression in the immediate vicinity of strong induction sites as compared with the general population (Figure 5F). This enrichment decayed rapidly as a function of neighborhood size, suggesting a highly localized effect (Figure 5F).

Remarkably, in addition to reporting gene-expression levels, seqFISH also reports the physical position of measured mRNA molecules at a sub-micron resolution. During this analysis we noticed that R2-pyocin transcript fluorescence generally appeared as two spots. Upon closer examination, we discovered that this mRNA is strongly localized to the two cell poles (Figure 5G). The 16S rRNA fluorescent signal in these pyocin producers show identical polarization, a rare pattern not observed in neighboring non-inducing cells (Figure 5G). These data suggest that ribosomes and the R2-pyocin transcript are mobilized following induction and spatially co-localize. In contrast, the expression of *recA* did not follow this pattern, suggesting a pyocin-specific effect (Figure 5G). Notably, a recent study discovered an identical polar localization for two different *Pseudomonas protegens* R-tailocins at the protein level (Vacheron et al., 2021). Together, these data hint at a potentially evolutionary conserved RNA-dependent mechanism for R-tailocin protein polar localization. We hypothesize that the spatially correlated ribosomal enrichment may provide efficient local translation and particle accumulation prior to cell lysis.

### Temporal evolution of metabolic heterogeneity during biofilm development

Beyond resolving transcriptional activities that contribute to biofilm developmental processes, seqFISH can reveal how biofilm cells metabolically respond to subtle changes in their local microenvironment. Chemical heterogeneity is a key feature of spatially structured environments, and metabolic heterogeneity characterizes mature biofilms (Evans et al., 2020; Povolotsky et al., 2021; Stewart, 2003; Stewart and Franklin, 2008). Yet until now, it has been impossible to capture the development of fine-grained metabolic structure across multiple suites of genes at different times.

To map biofilm metabolic development, we focused on genes whose regulation and functions are well understood. In particular, we focused on catabolic genes whose gene products enable energy conservation under different oxygen concentrations. Oxygen is a central and dynamic factor that influences metabolic activity in bacterial biofilms (Dietrich et al., 2013; Evans et al., 2020; Stewart, 2003; Wessel et al., 2014). Local oxygen availability can vary significantly within structured environments and is biotically shaped within biofilms (Cowley et al., 2015; Stewart and Franklin, 2008; Wessel et al., 2014). *P. aeruginosa* can survive under anaerobic conditions by fermenting different substrates and/or denitrifying (Arai, 2011; Eschbach et al., 2004; Yoon et al., 2002). Accordingly, monitoring the expression of these catabolic genes as well as others that are co-regulated with them provides a means to track local oxygen availability and its dynamic effects on biofilm metabolic coordination.

How quickly and over what spatial scales do biofilm cells metabolically differentiate? Following the *uspL* gene, which was strongly induced during hypoxic conditions and correlated with anaerobic fermentation and denitrification genes in our planktonic growth experiments, we observed surprisingly heterogeneous responses to oxygen depletion over just a few microns in young (10h) biofilms (Figure 6A). Notably, *uspL* expression is strongly spatially correlated with multiple anaerobic markers (Figure S4), indicating that this gene reports on local anaerobic activities. A closer examination of these putative hypoxic sites showed a frequent anti-correlation of *uspL* with multiple genes that are otherwise uniformly expressed in 10h biofilms, appearing as co-localized but reversed expression patches (Figure 6B). Among the anti-correlated functions are the TCA cycle gene, *sucC*, and replicative capacity genes such as RNAP and ribosome subunits (Figure 6B; Figure S4-S5). However, exceptions to this anti-correlation were also visible (Figure S5).

**Figure 6.**
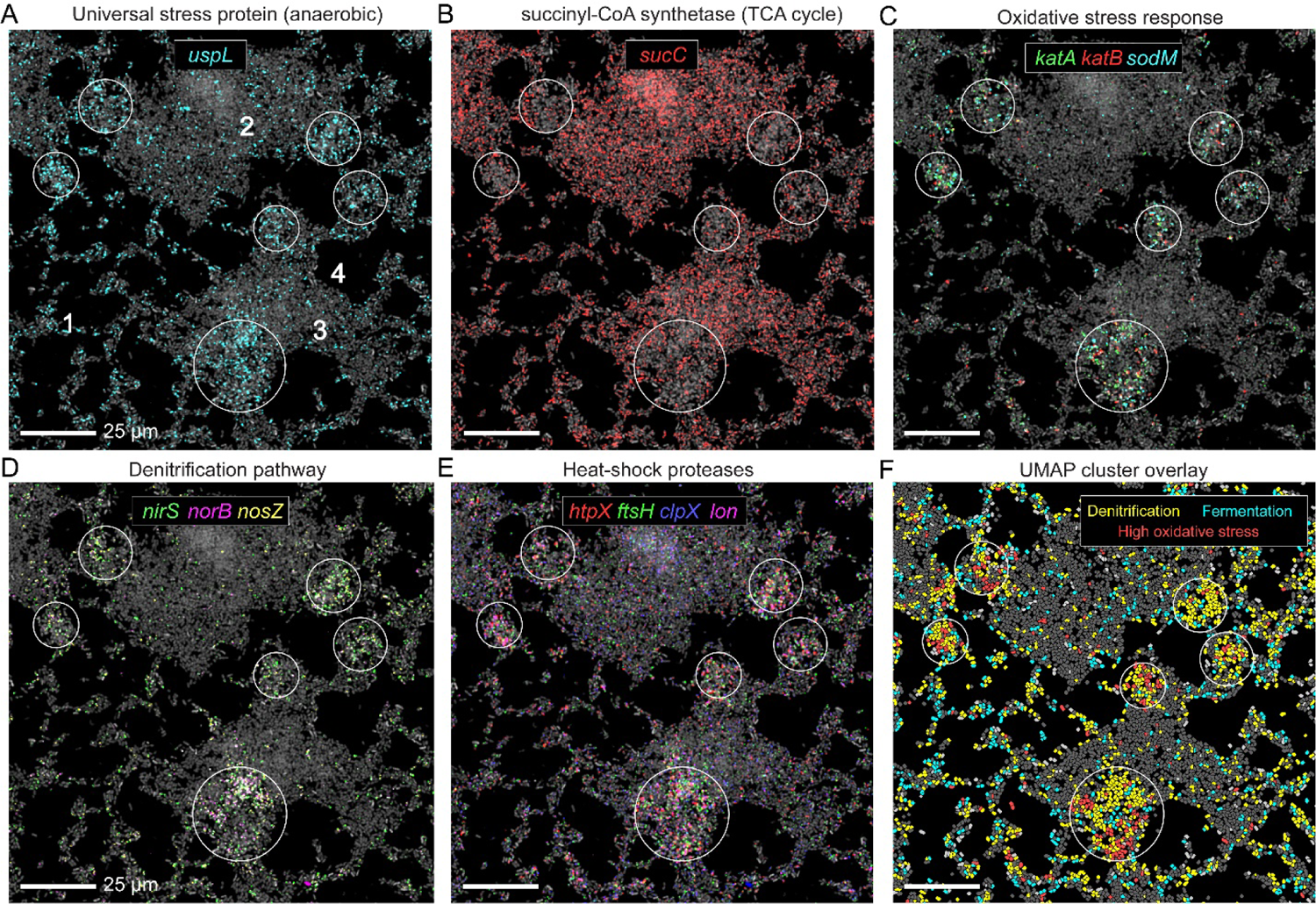
Oxygen availability shapes microscale metabolic heterogeneity in biofilms (A-E) Representative 10h biofilms. Cells are shown via 16S rRNA FISH fluorescence (gray) and overlayed with raw mRNA-FISH fluorescence for different genes as indicated in each panel. White circles highlight regions of interest. (F) Cells painted according to their UMAP derived metabolic state as indicated in the panel legends (also see Figure S6 clusters, 0, 8, 12 and 15), showing co-localization of multiple metabolic states within a given region.

Can the metabolic heterogeneity revealed by oxygen-responsive marker genes provide an entry point for the discovery of more nuanced cellular responses at the microscale? Our spatial correlation analysis revealed an intriguing association between anaerobic metabolism genes, such as the denitrification pathway (*narG*-*nirS*-*norB*-*nosZ*), and the oxidative stress response genes *katA*, *katB* and *sodM*, encoding for the inducible catalases and an Mn-dependent superoxide dismutase, respectively (Brown et al., 1995; Hassett et al., 1992; Su et al., 2014) (Figure 6C-D; Figure S4-S5). Nitrite respiring *P. aeruginosa* produce the highly toxic intermediate nitric oxide (NO) (Cutruzzolà and Frankenberg-Dinkel, 2016). Indeed, KatA was recently demonstrated to play a role in protection from NO-associated stress (Su et al., 2014), suggesting that these sub-aggregate regions correspond to microenvironments with high NO levels. In agreement with this hypothesis, we find that the stress response pattern is also spatially correlated with heat-shock protease expression, including the membrane protease, *ftsH*, which was found to play an important role in survival under anoxic conditions (Basta et al., 2017) (Figure 6E; Figure S4). These data highlight how contrasting physiological states can be established just a few microns away early in biofilm development.

These coordinated expressions patterns for particular genes led us to hypothesize that these patterns reflected the spatiometabolic distribution of distinct physiological “states” across the biofilm. To test this hypothesis we conducted a targeted UMAP analysis using only the 10h biofilm cells (Figure S6). We identified two main anaerobic sub-populations corresponding to denitrification and fermentation dominated metabolic states (Figure S6). In addition, we detected a smaller sub-population of denitrifying cells with 5.3-fold average increase in the oxidative stress factors *katB*, *sodM*, and *ahpF*, which encodes for an alkyl hydroperoxide reductase (Ochsner et al., 2000). Relative to the main denitrifying sub-group, stressed cells have lower expression of the denitrification pathway (∼4-fold) and a >2-fold reduction in replicative capacity marker levels (*rpoA*, *rpsC* and *atpA*), in support of a potentially damaged state.

Projecting these single-cell metabolic states over their respective biofilm positions showed a strong overlap with the above predicted hypoxic pockets, supporting our hypothesis and revealing that multiple metabolic states can co-exist in the same patch (Figure 6F; Figure S5).

Given the extent of transcriptional heterogeneity manifest in young biofilms, we wondered whether such heterogeneity would persist as biofilms aged. We speculated that the higher cell densities and more committed spatial structuring of mature biofilms might favor larger scale metabolic zonation. We therefore examined the spatial expression patterns in a 35h biofilm experiment.

In contrast to the spatial variation in aerobic and anaerobic metabolic processes seen in 10h biofilms, 35h biofilms have ∼50-fold lower average expression of the denitrification pathway genes *nar*-*nirs*-*norB*-*nosZ*. Indeed, these genes are known to be repressed by the *las* and *rhl* QS-systems, indicating *P. aeruginosa* is programmed to shut down denitrification at high cell densities (Toyofuku et al., 2007; Yoon et al., 2002). However, in addition to this complete and co-regulated pathway, *P. aeruginosa* also encodes an independent periplasmic nitrate reductase (*nap*) (Van Alst et al., 2009). Intriguingly, the *napA* gene is uniformly expressed at low levels across the 35h aggregates, a pattern that was closely shared with the *uspL* gene (Figure 7A; Figure S7). NapA has been implicated in maintaining redox homeostasis under oxygen limitation (Dietrich et al., 2013) and the *uspL* paralogue, *uspK*, was shown to play a role in survival under such conditions (Basta et al., 2017; Schreiber et al., 2006). At first blush, these results suggest that as aggregate cell mass grows, survival physiology dominates over growth-promoting processes on average. Yet we also find substantial and large-scale heterogeneity in certain genes, such as the replicative capacity markers (Figure 7B; Figure S7) and, *lasB*, encoding a QS-regulated extracellular protease (Figure 7C; Figure S7). These data demonstrate that while older biofilms generally comprise larger zones of particular activities than younger biofilms, a single microaggregate can still contain cell types with distinct physiological states and virulence-related activities.

**Figure 7.**
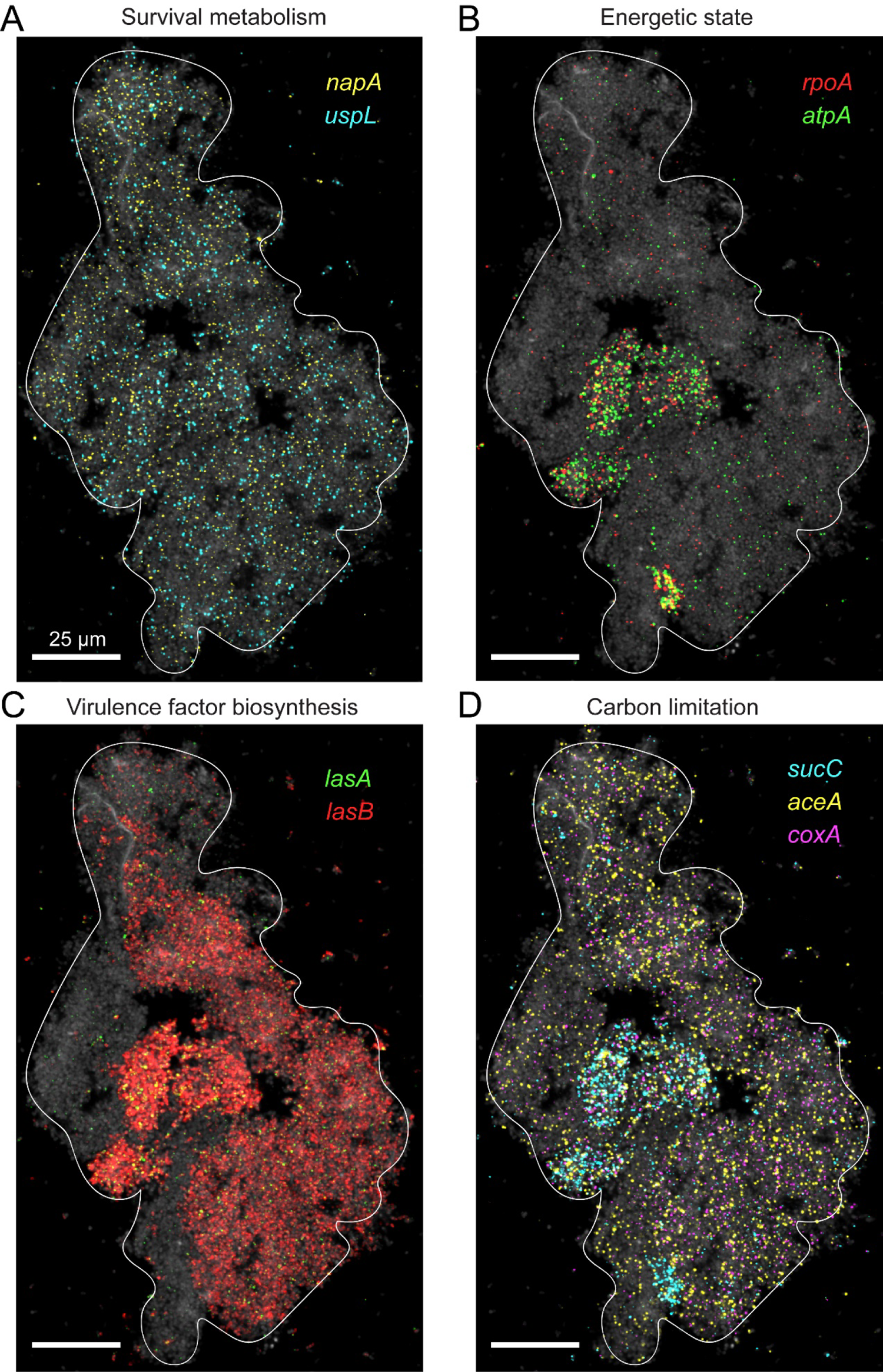
Functional zonation in a single microaggregate. (A-D) A *P. aeruginosa* 35h aggregate. Bacteria are shown via 16S rRNA FISH fluorescence (gray) and are overlaid with raw mRNA-FISH fluorescence for different genes as described in the panel legends.

Finally, that metabolism dynamically shapes the microenvironment leads to the prediction that differences in local nutrient availability will be reflected in heterogenous transcriptional activities over small spatial scales (Evans et al., 2020). We see evidence of this phenomenon in our data when focusing on carbon metabolism, for example. Where replicative capacity appears to be high and carbon is presumably replete, we see co-expression of the TCA cycle gene (*sucC*) (Fig. 7B-7D). However, when carbon is limiting, bacteria can utilize the glyoxylate shunt (GS), which bypasses the oxidative decarboxylation steps of the TCA. The GS provides an alternative metabolic pathway for utilizing acetate and fatty acids as carbon sources (Crousilles et al., 2018; Dolan and Welch, 2018). In the GS, carbon flux is redirected by isocitrate lyase (ICL) which competes with the TCA enzyme isocitrate dehydrogenase (ICD) for isocitrate.

However, since ICD has a much lower K_m_ it must be enzymatically inactivated via phosphorylation for the carbon flux to be redirected to the GS (Crousilles et al., 2018). However, little is still known about the transcriptional regulation of these pathways (Dolan et al., 2020). Our gene set contains both the GS gene, *aceA*, as well as a downstream TCA cycle gene, *sucC*. While these genes are often co-expressed, we find that only the GS marker, *aceA*, is expressed in low energetic capacity biofilm zones (Figure 7D; Figure S7), suggesting these subregions experience carbon limitation. In support of this hypothesis, these regions also express the tightly regulated terminal oxidase gene, *coxA*, which is transcriptionally induced by carbon starvation, a condition in which it promotes survival (Basta et al., 2017; Kawakami et al., 2010) (Figure 7D; Figure S7). This is just one example of the type of coherent spatiometabolic stratification pattern seqFISH can reveal at a given moment in time.

## Discussion

We adapted seqFISH to the study of bacterial planktonic and biofilm populations and showed that this approach can measure hundreds of genes within individual bacteria. Furthermore, we introduced par-seqFISH, a novel multiplexing approach that enables parallel seqFISH imaging of dozens of samples. Studying *P. aeruginosa* across a range of physiological conditions and two different modes of life (planktonic v. sessile), we analyzed the expression of ∼600,000 individual cells. We identified the dynamic emergence of planktonic subpopulations with distinct metabolic and/or virulence-related functions throughout the course of planktonic growth, effectively capturing evolving cell states. In both young and older biofilms, we observed strikingly high levels of metabolic heterogeneity and zonation, yet coherent co-expression patterns emerged from the subcellular to the microscale that gave rise to new insights. Together, the patterns this new way of “seeing” reveals provide a means to chart and understand the extraordinary diversity that defines the microbial world.

Our approach is based on non-barcoded sequential single-molecule FISH, a sensitive and accurate method for measuring mRNA expression (Chen et al., 2015; Eng et al., 2019; Lignell et al., 2017; Raj et al., 2008; So et al., 2011). As each gene is measured individually, the number of genes analyzed per experiment scales linearly with the number of hybridization cycles. Thus, from a practical point of view, this method is better suited for more targeted studies of several hundred genes. Our results nonetheless demonstrate that gene sets of this magnitude can drive explorative studies of population sub-structure and functional cell states. Considering the operonic encoding of bacterial genes, the effective gene coverage of seqFISH is in fact significantly higher (∼3-fold higher in our set). In contrast, bacterial scRNA-seq methods are unbiased and provide a genome-wide sampling. However, these methods are not spatially resolved and must deal with relatively low capture efficiencies, estimated as 2.5-10% of the total mRNAs in recent studies (Blattman et al., 2020; Kuchina et al., 2021). Thus, seqFISH and scRNA-seq provide complementary approaches for studying individual planktonic bacteria in terms of sensitivity and throughput.

In addition to mRNA profiles, our method has the unique advantage of capturing key cell biological parameters such as cell size and shape and can be further integrated with functional markers to connect between specific processes and expression within the same cell. In a recent study, seqFISH was integrated with DNA-FISH and protein abundances via immunofluorescence within the same cells (Takei et al., 2021). In addition to its integrative capacity, the spatial resolution and single-molecule nature of seqFISH provides a record of the physical location of all studied mRNAs. As just one example, our analysis exposed a strong intracellular localization signature of R-pyocin transcripts and ribosomal rRNA within inducing cells. Beyond the potential biological implications of these observations, these data highlight seqFISH as a high-throughput platform for exploring sub-cellular transcript organization in bacteria (Fei and Sharma, 2018).

To increase the throughput of seqFISH for single-cell analysis we developed the par-seqFISH multiplexing approach, which enabled the study of ∼270,000 planktonic cells grown in 21 different conditions. However, this approach can potentially be applied in various other ways, both in synthetic and natural communities. For example, since par-seqFISH is based on 16S rRNA labels (Ribo-Tags), it could in principle be used to encode bacterial taxonomy (Figure 1C). Recently, a conceptually similar and exciting method for combinatorial labeling of taxonomy was introduced in a biogeographical study of the human microbiome (Shi et al., 2020). In principle, the par-seqFISH strategy could be readily extended to capture similar or higher level of taxonomic complexity, as well as adding the currently missing feature of mRNA expression. We predict that this future application will be useful in providing a functional measure for interpreting spatial associations between microbial species.

To our knowledge, this study presents the first highly multiplexed and spatially resolved single-cell analysis of bacterial populations. Future application of this approach at greater temporal resolution or in other biofilm models could help shed light on biofilm functional organization and development.

Furthermore, extension of this approach to natural and clinical samples could provide important insights into the conditions experienced by microbes in more complex environments and the coordinated physiological responses that emerge in turn. Understanding the roles that spatial and temporal heterogeneity play in microbial populations represents an exciting and important research aim for modern microbiology. The results presented in this manuscript provide a new way of addressing these defining features on our way to understanding microbial life on its own terms.

## Materials and methods

### Bacterial strains and growth conditions

*P. aeruginosa* strain UCBPP-PA14 was grown aerobically with shaking at 250 rpm in lysogeny broth (LB) (Difco) or on LB agar plates at 37°C. SCFM was made as previously described (Palmer et al., 2007). For the growth curve experiments, an overnight LB culture was washed twice using fresh growth media (either LB or SCFM) and then diluted 1:100 into 100 ml prewarmed fresh media. The cultures were grown at 37°C with shaking at 250 rpm and collected at various time points as indicated in Figure 2A. The SCFM samples were collected cell densities identical to the LB experiment, except the OD_600_ = 3.2 sample was omitted. Collected samples were immediately fixed in ice-cold 2% paraformaldehyde (PFA) and were incubated on ice for 1.5h in the dark, and then washed twice with 1x PBS. Samples were resuspended in 70% EtOH and incubated at −20°C for at 24h to permeabilize the cells. Surface colonization was performed by washing and diluting an LB overnight culture 1:100 into fresh SCFM and dispensing 100 µl into coverslip attached open incubation chambers (Electron Microscopy Sciences, 70333-42). The coverslips were incubated in parafilm sealed sterile petri dishes at 37°C and the media was gently exchanged every 4 hours. A damp Kimwipe was placed in the petri dish to control media evaporation. During the overnight stage of the 35h experiment, the media was exchanged only once after 8h. Biofilm experiments were collected by gently exchanging the SCFM with 100 µl ice cold 2% PFA solution and incubating the sample at 4°C for 1.5h. The samples were washed twice with 1x PBS, resuspended in 70% EtOH and incubated overnight at 4°C and prepared for seqFISH as described below the following day.

### SeqFISH probe design and library generation

Primary probes were designed as 30 nt stretches in a GC range of 45-65%. Probe sequences containing more than four consecutive base repeats were removed. The remaining probes were compared to the reference genome using blast and any probe with non-specific binding of at 18nt or more was discarded. Negative control genes were selected from the P1 phage genome (NC_005856.1) with the same criteria.

Each selected gene was covered by 12-20 nonoverlapping probes randomly selected from the gene probe set. The probes were designed as a 30nt mRNA binding region flanked by overhangs composed of four repeats of the secondary hybridization sequence (complementary to a designated fluorescent readout probe; Table S2). Thus, it is estimated that during secondary hybridization, each mRNA is covered by 48-80 fluorescent readout probes (12-20 x 4), on par with previous mRNA-FISH experiments in bacteria (Skinner et al., 2013; So et al., 2011).

A library of 1,763 probes targeting 105 *P. aeruginosa* genes and three negative controls were designed (Tables S1-S2). Additional flanking sequences were added to the primary probe sequences to enable library amplification via PCR (Forward 5’-TTTCGTCCGCGAGTGACCAG-3’ and reverse 5’-CAACGTCCATGTCGGGATGC-3’). The primary probe set was purchased as oligoarray complex pool from Twist Bioscience and constructed as previously described (Eng et al., 2019) (Table S2). Briefly, a set of 9 PCR cycles were used to amplify the designated probe sequences from the oligo pool. The amplified PCR products were purified using the QIAquick PCR Purification Kit (28104; Qiagen) according to the manufacturer’s instructions. The PCR products were used as the template for in vitro transcription (E2040S; NEB) followed by reverse transcription (EP7051; Thermo Fisher). Then, the single-stranded DNA (ssDNA) probes were alkaline hydrolyzed with 1 M NaOH at 65c C for 15 min to degrade the RNA templates, followed by 1 M acetic acid neutralization. Next, to clean up the probes, we performed ethanol precipitation to remove stray nucleotides, phenol–chloroform extraction to remove protein, and used Zeba Spin Desalting Columns (7K MWCO) (89882; Thermo Fisher) to remove residual nucleotides and phenol contaminants. Readout probes were designed as previously described and ordered from Integrated DNA Technologies (IDT) (Eng et al., 2019).

Ribo-Tag probes were designed to target the same region in the 16S rRNA gene according to the criteria described above, but with a 28nt binding regions. Each probe sequence was flanked with two secondary sequences selected out a set of six that were dedicated to multiplexing (Table S3). An additional 16S rRNA probe was generated as a standard between all multiplexed samples and was hybridized to an independent region of the 16S rRNA (Table S3). This probe provided an additional reference and was used to register images from different channels (see below).

### Coverslip functionalization

Coverslips were cleaned with a plasma cleaner on a high setting (PDC-001, Harrick Plasma) for 5 min, followed by immersion in 1% bind-silane solution (GE; 17-1330-01) made in pH 3.5 10% (v/v) acidic ethanol solution for 30 min at room temperature. The coverslips were washed with 100% ethanol three times and dried in an oven at >90 °C for 30 min. The coverslips were then treated with 100 μg μl^−1^ of c poly-D-lysine (P6407; Sigma) in water for at least one hour at room temperature, followed by three rinses with water. Coverslips were air-dried and kept at −20°C for no longer than 2 weeks before use.

### Parallel seqFISH

Independent fixed samples were individually hybridized with 16S rRNA labels, washed and then pooled into a single mixture that was hybridized with the gene probe library and prepared for imaging. Approximately 10^8^ cells were collected from each sample into a microcentrifuge, pelleted via centrifugation (6,000 rpm) and then resuspended in 20 μ H_2_0 with 6 nM of the designated 16S rRNA label (sample specific) and another 6 nM of a shared reference 16S rRNA probe (Table S3). Each sample was then mixed with prewarmed 30 μ of primary hybridization buffer (50% formamide, 10% dextran sulfate and 2× SSC) via gentle pipetting, incubated at 37°C for >16 h, washed twice with 100 μ wash buffer (55% formamide and 0.1% Triton-X 100 in 2× SSC; 5 min 8,000 rpm for the viscous hybridization buffer) and then incubated at 37°C in 100 μl wash buffer for 30 min to remove non-specific probe binding. Samples were washed twice with 100 μl 2x SSC and pooled together into a new microcentrifuge μl H_2_0 and 10 μl gene probe library mixture and mixed well with prewarmed 30 μ buffer. The hybridizations were incubated for >16 h at 37°C and were washed and prepared as described above. The final mixture was resuspended in 20-25 μ 1x PBS and 5-10 μ were gently spotted at the center of the coverslip and incubated at RT for 10 min to allow the cells to sediment and bind the surface. The coverslips were centrifuged for 5 min at 1,000 rpm to create a smooth and dense cell monolayer. The cells were immobilized using a hydrogel as previously described (Eng et al., 2019) and stained with 10 μl ml^−1^ DAPI (D8417; Sigma) for 5 min before imaging so that cells could be visualized.

In biofilm experiments, the fixed and permeabilized surface attached microaggregates were air dried, covered with a hydrogel and hybridized with the gene library and a rRNA probes in one single reaction, as described above.

### seqFISH imaging

All seqFISH experiments were performed using a combined imaging and automated fluidics delivery system as previously described (Eng et al., 2019). DAPI stained samples mounted on coverslips were connected to the fluidic system. The ROIs were registered using the DAPI fluorescence and a set of sequential secondary hybridizations, washes and imaging was performed.

Each hybridization round contained three unique 15-nt readouts probes each conjugated to either Alexa Fluor 647 (A647), Cy3B and Alexa Fluor 488 (A488). All readout probes were ordered from Integrated DNA Technologies and prepared into 500 nM stock solutions. Each serial probe mixture was prepared in EC buffer (10% ethylene carbonate (E26258; Sigma), 10% dextran sulfate (D4911; Sigma), 4× SSC).

Hybridizations were incubated with the sample for 20 min to allow for secondary probe binding. The samples were then washed to remove excess readout probes and to limited non-specific binding using ∼300 μ of 10% formamide wash buffer (10% formamide and 0.1% Triton X-100 in 2× SSC). Samples were then rinsed with ∼200 l of 4× SSC and then stained with DAPI solution (10 μg ml^−1^ of DAPI, 4× SSC). Lastly, an anti-bleaching buffer solution made was flowed through the samples (10% (w/v) glucose, 1:100 diluted catalase (Sigma C3155), 0.5 mg ml^−1^ glucose oxidase (Sigma G2133) and 50 mM pH 8 Tris-HCl in 4× SSC). Imaging was performed with a Leica DMi8 microscope equipped with a confocal scanner unit (Yokogawa CSU-W1), a sCMOS camera (Andor Zyla 4.2 Plus), a 63× oil objective lens (Leica 1.40 NA) and a motorized stage (ASI MS2000). Lasers from CNI and filter sets from Semrock were used. Snapshots were acquired using 647-nm, 561-nm, 488-nm and 405-nm fluorescent channels with 0.5-μm z-steps for all experiments with the exception of the 35h biofilm experiment in which 1.0-μm z-steps were collected. After imaging, readout probes were stripped using 55% wash buffer (55% formamide and 0.1% Triton-X 100 in 2× SSC) that was flowed through for 1 min, followed by an incubation time of 15 min before rinsing with 4× SSC solution. This protocol: serial hybridizations, imaging and signal quenching steps, was repeated for ∼40 rounds to capture 16S rRNA for multiplexing, mRNA expression and background signal. The integration of automated fluidics delivery system and imaging was controlled via μ Manager (Edelstein et al., 2010).

### Image analysis demultiplexing and gene-expression measurement

Maximal projection images were generated using ImageJ (Schneider et al., 2012) for DAPI and 16S rRNA and hybridization rounds were registered using the DAPI fluorescence. Aberrations between fluorophores were corrected by alignment of 16S rRNA signals across all channels. Cells were segmented using the DAPI signal with SuperSegger using the 60XPa configuration (Stylianidou et al., 2016) and filtered using custom scripts to eliminate odd shapes, autofluorescent or low signal components.

For par-seqFISH demultiplexing, the background (no readouts) and 16S rRNA fluorescent intensity for each relevant secondary readout probe was measured within segmented cell boundaries to provide a signal-to-background score for each readout. The cells were classified according to the positive readout combinations (Table S3). The level of false positives was estimated by counting the number of cells classified into combinations left out of the experiment.

The mRNA-FISH data was analyzed using Spätzcells (Skinner et al., 2013). Briefly, spots were detected as regional maxima with intensity greater than a threshold value that was set using the negative control genes and were fit with a 2D gaussian model. The integrated intensity of the spot and the position of its estimated maxima were determined (Skinner et al., 2013). Spots were assigned to cells using the cell segmentation masks (Skinner et al., 2013). In biofilm experiments spots were assigned to cells in a z-section sensitive manner. Deviating spots maxima positions that did not overlap a cell boundary were tested against the flanking z-sections to identify their cell of origin. If no cell was detected the spots were discarded. All predicted low expression genes (defined as gene with spots in less than 30% of all cells) were identified and the distribution of their spot intensities was fit with a gaussian mixture model to identify the characteristic intensity of a single mRNA, normalized to the number of probes used for the specific gene. The median characteristic single-mRNA signal was then calculated using all low expression genes for each fluorophore (A647, A488 and cy3B). The variation between different genes labeled with the same fluorophore was low, with a coefficient of variation of 18-21%. This median characteristic value was used to transform fluorescent intensity into to discrete mRNA counts per gene within each cell. The A488 characteristic signal was corrected by a factor of 1.5 to account for its lower intensity in our system. In each cell, the total intensity of each gene was calculated by summing the intensities of all spots. The total value was normalized by the characteristic value for a single mRNA in the corresponding fluorophore.

### Single-cell expression analysis and cell biological parameter calculations

Single-cell UMAP analysis was performed using Scanpy v1.7.0 (Wolf et al., 2018). Genes detected at consistently low levels were excluded from the analysis. These included *pilY1*, *flgK*, *nasA*, *algU*, *purF*, *phzH*, *phzS* and *pslG* (Table S1). We followed the standard Scanpy normalization and scaling, dimensionality reduction, and clustering as described in the Scanpy tutorial, minus the high variance gene selection and without a library size normalization. We used 15 neighbors and 15 and 17 PCA components, for the LB and merged SCFM analyses, respectively. Clustering was performed using the Leiden method. Jupyter notebooks with chosen parameters, run lines, output files and source data are available at https://github.com/daniedar/seqFISH.

Cell nucleoid size was calculated using the segmentation mask. A chromosome score was calculated as the median DAPI intensity multiplied by the nucleoid size. The median chromosome score was calculated for the last time point in our LB experiment (deep stationary; OD_600_ = 3.2). Because most cells in this stage are in a non-dividing state, we set this value as a reference for a single chromosome copy. We then normalized the scores of all cells in the experiment using this value, as seen in Figure 2. In addition to using Ribo-Tags to label cells from different conditions, we also hybridized another region in the 16S rRNA with a probe that was shared across all samples (Table S3; described above). We used this reference signal to compare the 16S rRNA intensity between cells from different conditions. We measured the median 16S rRNA signal per cells and multiplied it by the nucleoid size (which completely overlaps the 16S signal and estimates cell size). In *E. coli*, maximal ribosome numbers appear at the maximal growth rate and have been estimated at 72,000 (Milo et al., 2010). The median rRNA score was calculated for the maximal growth (OD_600_ = 0.2) and normalized to 72,000 as in *E. coli* for a rough estimate (Figure 2).

### Image analysis in surface colonization experiments

Images were registered as described above and segmentation was performed using the pixel classification workflow in Ilastik (Berg et al., 2019). We trained the Ilastik classification model with background, cell boundaries and cell bodies, using the 16S rRNA signal. We find that Ilastik performs extremely well.

However, in high density regions, segmentation often resulted in over-connectivity due to incorrect 3D overlaps. We disconnected such cells clusters. The binary masks (segmentation output) were thinned, and all 3D connected components (CCs) were re-calculated. This reduced spurious connections. Then, all CCs traversing more than 2.5 µm were set aside for re-evaluation for potential over-connections. For each such 3D component we examined each z-slice at a time and identified all 2D CCs. We removed overly large or curved blobs, which represent segmentation artefacts that often incorrectly connect distinct cells across z-sections. In addition, for each 2D detected component we calculated its orientation and overlap with components in the previous flanking z-section. If this component exhibited a significant change in its orientation (the direction it is pointing) we disconnected it from the component below. We then continued the analysis using the newly oriented component as a seed. Cell clusters that could not be properly disentangled were removed from the analysis. At the end of the analysis the cell 3D masks were re-thickened.

We conducted bulk neighborhood analysis where we studied the immediate neighborhoods associated with high expression of a specific gene. For the gene of interest, we identified all top 99^th^ percentile cells (99.5^th^ for the pyocin specific analysis), denoted as “center cells”. Using the 3D centroid coordinates of center cells we identified their closest neighbors within a specified distance (up to 10 µm for pyocins and 3 µm for the rest). We then collected up to k closest cells (up to 5-300 neighbors in the pyocin analysis to view the enrichment decay and up to 5 for the rest of the genes). All of the neighborhood cells selected (not including the center cells) were then analyzed in bulk together and their mean gene-expression was calculated and compared to the population (minus all center cells not used). We conducted this analysis across all genes and performed a Pearson correlation analyses to identify spatially correlating genes (Figure S4).

## Acknowledgments

We thank George A. O’Toole and Marvin Whiteley for help with designing the gene set. We also thank Megan Bergkessel and Rotem Sorek and members of the Newman lab for critically reading the manuscript. We thank the Newman lab for fruitful discussions and comments. Grants to DKN from the NIH (1R01AI127850-01A1 and 1R01HL152190-01) and ARO (W911NF-17-1-0024) supported this work. LC was funded by the Allen Frontier group. DD was supported by the Rothschild foundation, EMBO Long-Term, and the Helen Hay Whitney postdoctoral fellowships, as well as a Geobiology Postdoctoral Fellowship from the Division of Geological and Planetary Sciences, Caltech.

## Author contributions

DD ND, LC and DKN designed the study. DD led the study, designed the experiments, and performed the experiments with ND. Analyses were performed by DD. DKN and LC supervised the study, and all authors wrote the manuscript.

## Data and Code Availability

Custom MATLAB scripts and single-cell source data from this study were deposited in https://github.com/daniedar/seqFISH. Imaging data obtained during this study are available from the corresponding author upon reasonable request.

## Supplemental Figures

**Figure S1.**
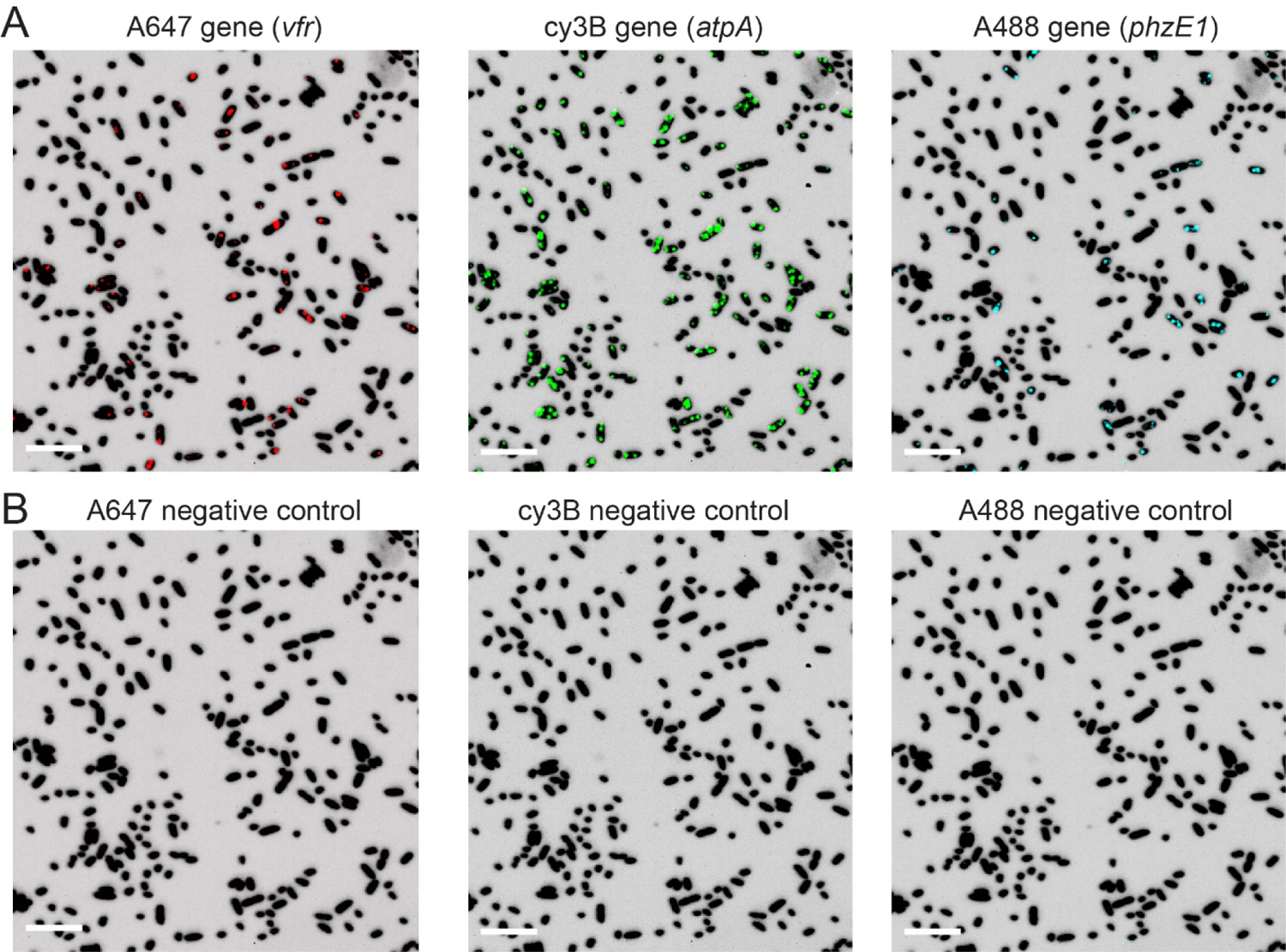
Negative control genes estimate the false positive rate. (A) Examples of positive signal for genes labeled with one of the three fluorophores used in this study A647 (red), cy3B (green), and A488 (cyan). For context, the mRNA-FISH fluorescence is shown over DAPI (dark silhouette). (B) Same regions as is in panel A but showing the raw fluorescence of the negative control genes for each fluorophore. For direct comparison, the intensity range is identical between positive and negative panels in A-B. Scale bar represents 5µm.

**Figure S2.**
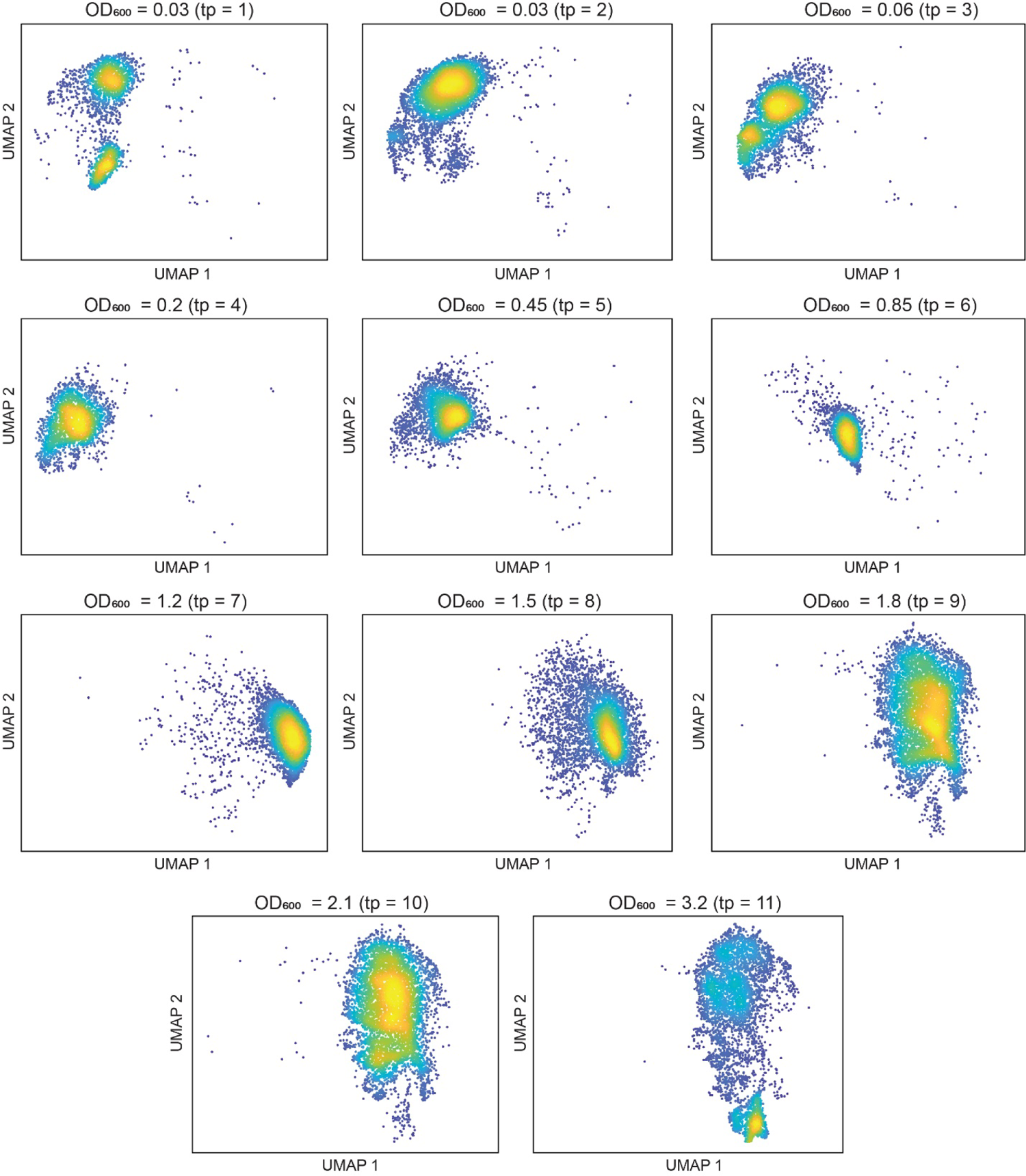
Single-cell dispersions in UMAP space for each growth curve time point. A UMAP density plot of cells belonging to specific time points. The OD_600_ values and the number of the time points are shown over each plot. Color intensity represents cell density.

**Figure S3.**
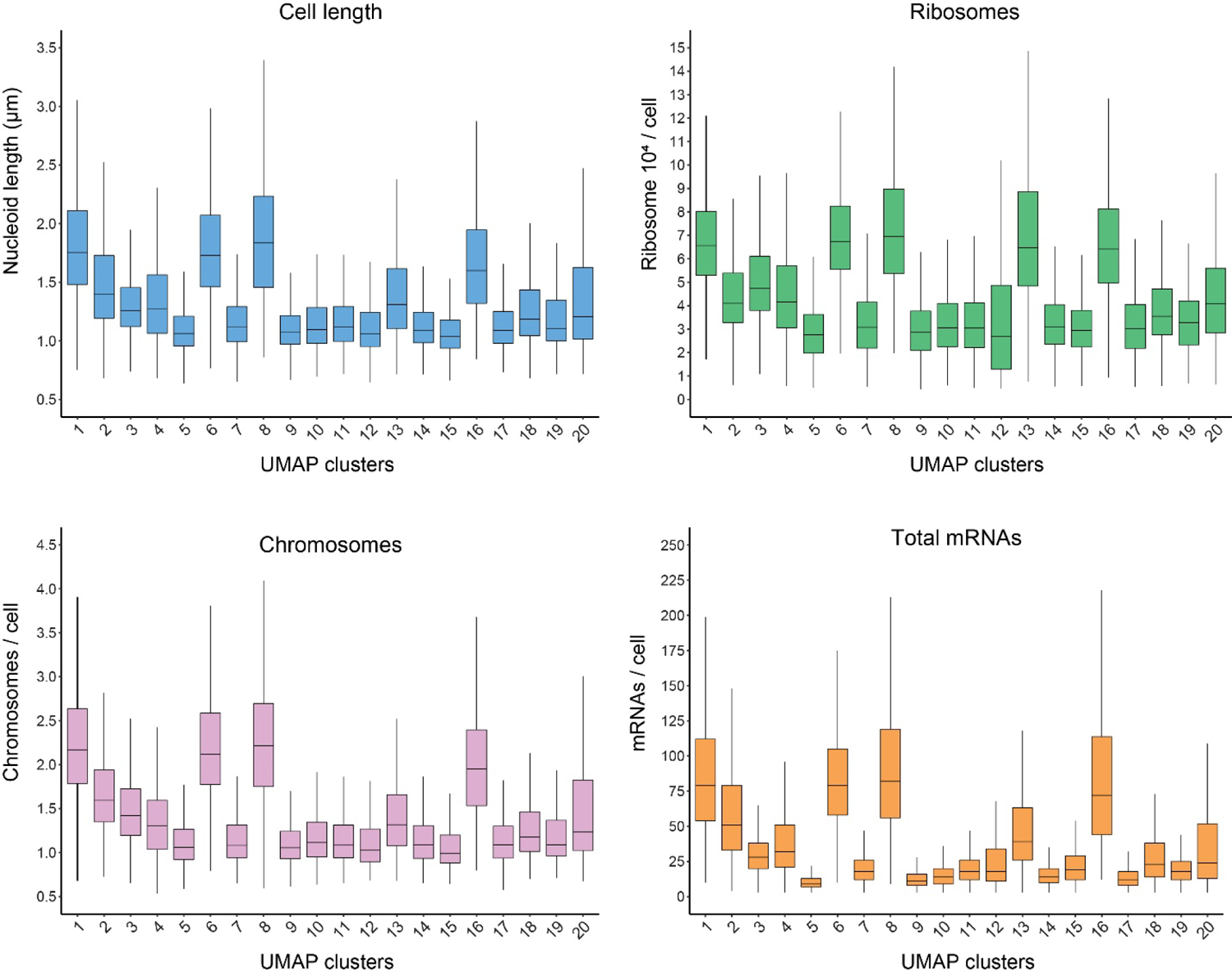
Distributions of single-cell parameters across the detected UMAP clusters Distributions of nucleoid length, chromosome copy, ribosome levels and total mRNAs for each of the UMAP clusters described in main figure 3.

**Figure S4.**
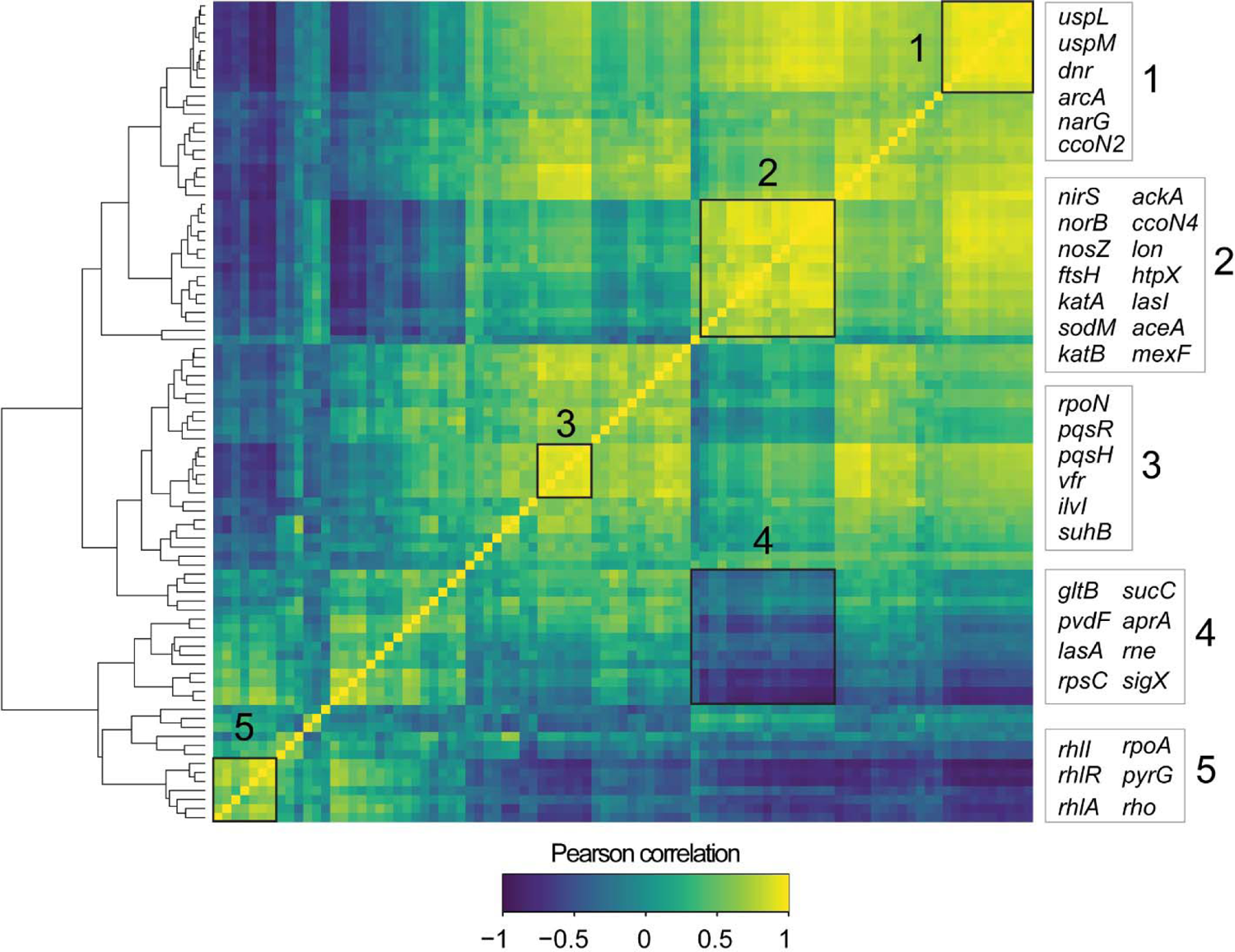
Spatial correlation analysis. Gene centered neighborhood analysis for detecting spatial correlation. For each gene, its 99^th^ percentile expressing cells were identified and their 5 immediate neighbors within 3 µm were collected (leaving out the enriched center cell). The set of all such neighbors cross the experiment was analyzed together to produce a mean expression profile that was compared with the total population to produce a local enrichment/depletion ratio. The Pearson correlation between such gene neighborhood profiles was calculated and shown above as a clustered heat map. Five selected regions are highlighted and numbered. Key genes within each cluster are described to the right.

**Figure S5.**
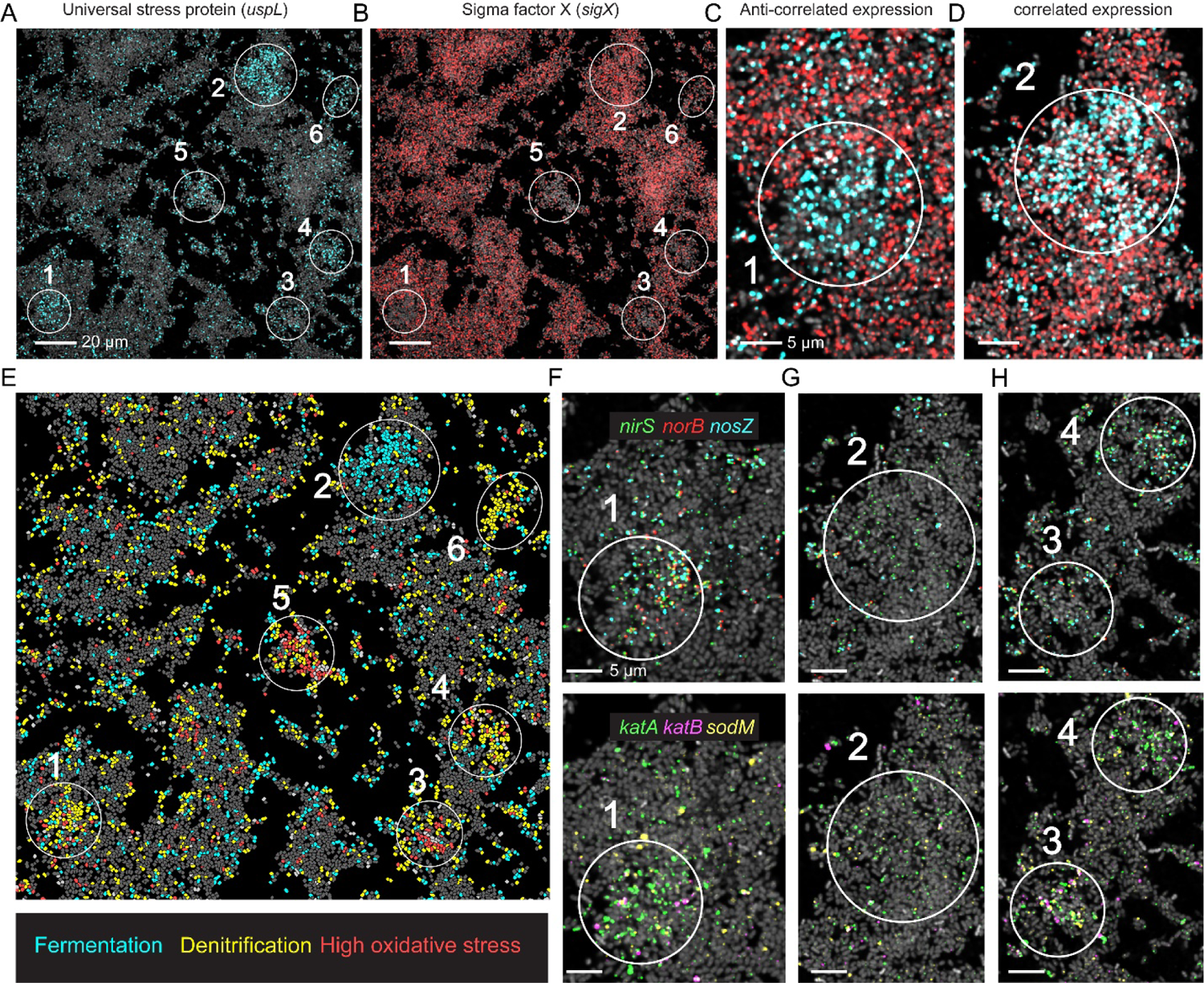
Distributions of single-cell parameters per UMAP cluster (A-B) Representative 10h microaggregates. Cells are shown via 16S rRNA FISH fluorescence (gray) and overlayed with gene-expression as indicated in each panel. White circles highlight regions of interest. (C-D) Zoom-in on region 1 and 2 showing *uspL* (cyan) and *sigX* (red). (E) Cells painted according to their neighborhood class as indicated in the panel legend. (F-H) Zoom-in highlighted regions overlaid with raw gene-expression as indicated in the panel legends.

**Figure S6.**
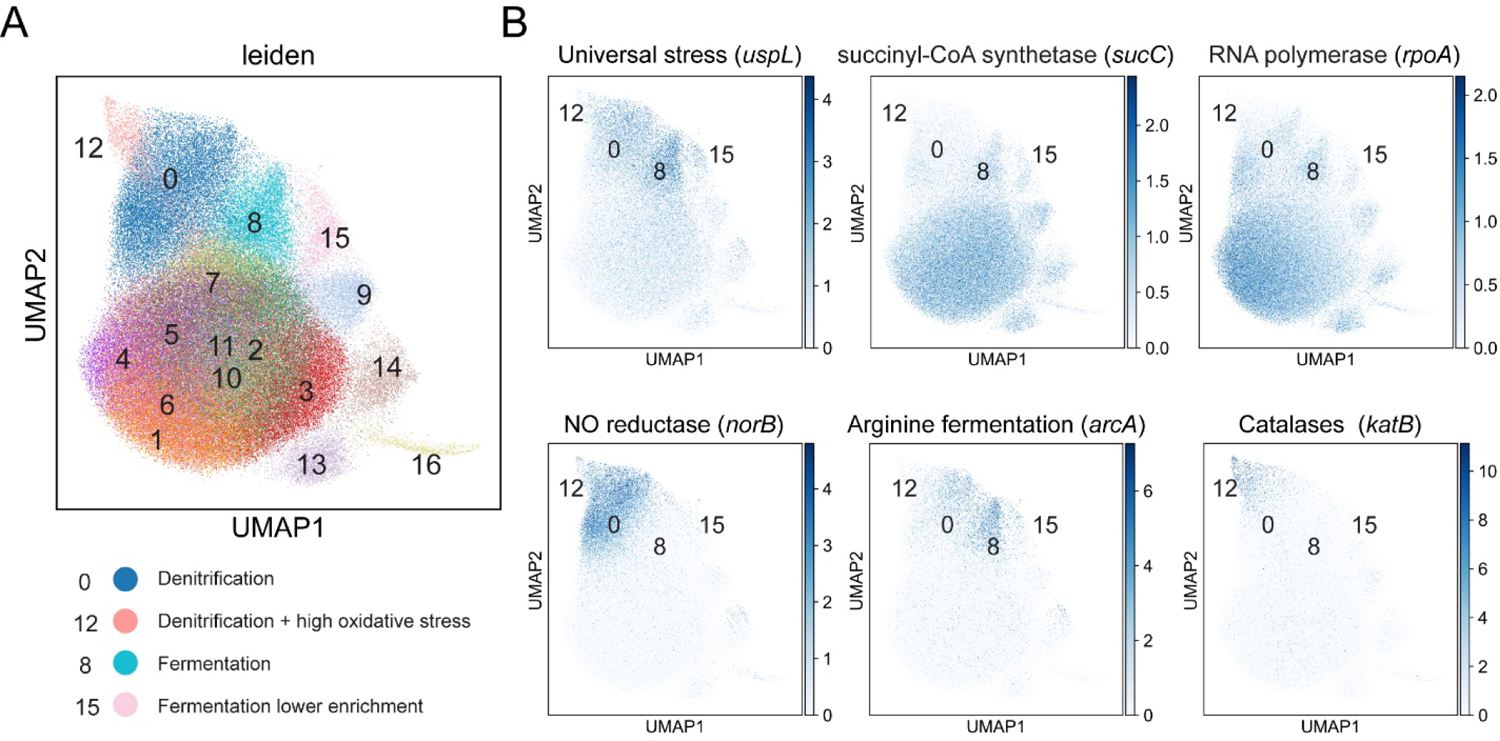
UMAP analysis of 10h biofilms (A) UMAP analysis was performed using the 10h biofilm experiment. Below, clusters are labeled and are divided into predicted anaerobic groups. (B) UMAP overlaid with specific gene data. The cluster number positions are shown in the figure.

**Figure S7.**
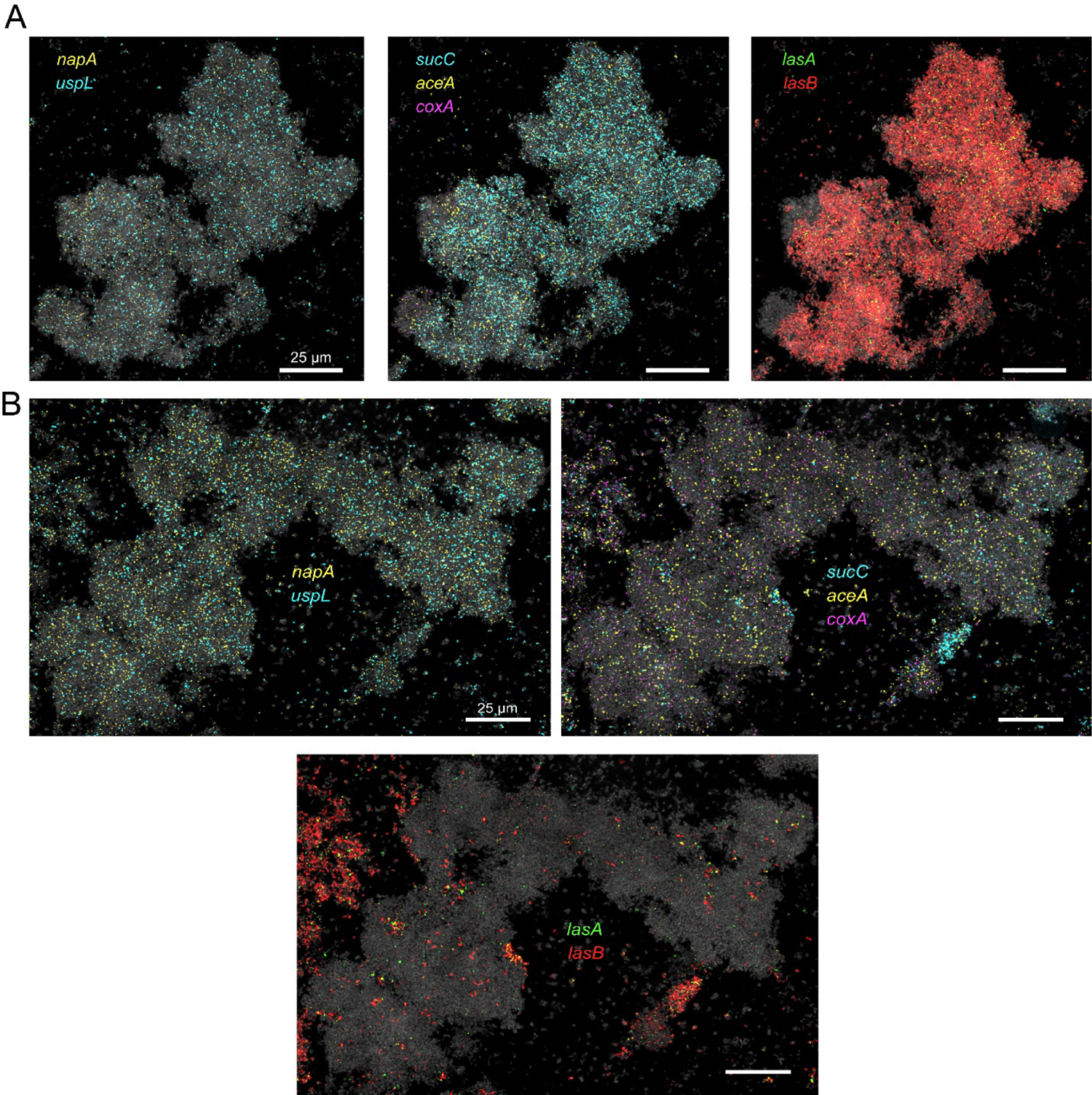
Functional zonation in 35h microaggregates (A-B) Various *P. aeruginosa* 35h aggregate. Bacteria are shown via 16S rRNA FISH fluorescence (gray) and are overlaid with raw mRNA-FISH fluorescence for several genes as described in the images.

