## Supplemental Table 1 for "In situ single-cell activities of microbial populations revealed by spatial transcriptomics"

### Table S1: gene library used in study

| Locus_tag | Gene name | Gene description |
|---|---|---|
| P1_gp003 | cre | Phage P1 negative control gene |
| P1_gp013 | pro | Phage P1 negative control gene |
| P1_gp078 | dbn | Phage P1 negative control gene |
| PA14_00010 | dnaA | chromosomal replication initiator protein DnaA |
| PA14_00560 | exoT | exoenzyme T |
| PA14_00640 | phzH | phenazine-modifying enzyme |
| PA14_01160 | vgrG | T6SS protein |
| PA14_01300 | coxA | cytochrome c oxidase, subunit I |
| PA14_01720 | ahpF | alkyl hydroperoxide reductase subunit F |
| PA14_01970 | TriC | Triclosan RND efflux transporter |
| PA14_03650 | cysA | sulfate transport protein CysA |
| PA14_03920 | spuD | polyamine transport protein |
| PA14_04410 | ptsP | phosphoenolpyruvate-protein phosphotransferase |
| PA14_04930 | rpoH | RNA polymerase sigma-32 factor |
| PA14_05310 | gshB | glutathione synthetase |
| PA14_05540 | mexB | RND multidrug efflux transporter MexB |
| PA14_06750 | nirS | nitrite reductase precursor |
| PA14_06830 | norB | nitric-oxide reductase subunit B |
| PA14_06870 | dnr | transcriptional regulator Dnr |
| PA14_07520 | rpoD | sigma factor RpoD |
| PA14_08150 | PA14_08150 | Pyocin R2 F2 operon |
| PA14_08370 | vfr | cyclic AMP receptor-like protein |
| PA14_08910 | rpsC | 30S ribosomal protein S3 |
| PA14_09115 | rpoA | DNA-directed RNA polymerase alpha chain |
| PA14_09150 | katA | catalase |
| PA14_09280 | pchF | pyochelin synthetase PchF |
| PA14_09400 | phzS | flavin-containing monooxygenase |
| PA14_09440 | phzE1 | phenazine biosynthesis protein PhzE |
| PA14_09490 | phzM | phenazine-specific methyltransferase |
| PA14_09520 | mexI | probable RND efflux transporter |
| PA14_10500 | ccoN4 | cytochrome c oxidase subunit (cbb3-type) |
| PA14_10790 | ampC | cephalosporinase |

| | | |
|---|---|---|
| PA14_13780 | narG | respiratory nitrate reductase alpha subun |
| PA14_14680 | suhB | extragenic suppressor protein SuhB |
| PA14_16250 | lasB | elastase LasB |
| PA14_16500 | wspR | two-component response regulator |
| PA14_17290 | pyrG | CTP synthase |
| PA14_17480 | rpoS | sigma factor RpoS |
| PA14_17530 | recA | RecA protein |
| PA14_18580 | algD | GDP-mannose 6-dehydrogenase AlgD |
| PA14_19100 | rhlA | rhamnosyltransferase chain A |
| PA14_19120 | rhlR | acylhomoserine lactone dependent transcriptional regulator |
| PA14_19130 | rhlI | autoinducer synthesis protein RhlI |
| PA14_20200 | nosZ | nitrous-oxide reductase precursor |
| PA14_22980 | gltB | binding protein component of ABC sugar transporter |
| PA14_23920 | purF | amidophosphoribosyltransferase |
| PA14_24480 | pelA | conserved hypothetical protein |
| PA14_25080 | fadB | fatty-acid oxidation complex alpha-subunit |
| PA14_25560 | rne | ribonuclease E |
| PA14_27480 | htpX | heat shock protein HtpX |
| PA14_30050 | aceA | isocitrate lyase |
| PA14_30630 | pqsH | FAD-dependent monooxygenase |
| PA14_32390 | mexF | RND multidrug efflux transporter MexF |
| PA14_33700 | pvdF | pyoverdine synthetase F |
| PA14_34050 | impC | conserved hypothetical protein |
| PA14_35670 | pslG | glycosyl hydrolase |
| PA14_36310 | hcnC | hydrogen cyanide synthase HcnC |
| PA14_36345 | exoY | adenylate cyclase ExoY |
| PA14_38410 | amrB | RND multidrug efflux transporter |
| PA14_39330 | rbsA | ribose ABC transporter, ATP-binding protein |
| PA14_40290 | lasA | staphylolytic protease preproenzyme LasA |
| PA14_40510 | ccoN3 | cytochrome c oxidase, cbb3-type, subunit |
| PA14_41220 | lon | Lon protease |
| PA14_41230 | clpX | ATP-dependent Clp protease ATP-binding subunit ClpX |
| PA14_41440 | uspL | universal stress protein |
| PA14_41510 | nasA | nitrate transporter |

| | | |
|---|---|---|
| PA14_41575 | sigX | ECF sigma factor SigX |
| PA14_42350 | pscC | Type III secretion outer membrane protein PscC precursor |
| PA14_42500 | pcrD | type III secretory apparatus protein PcrD |
| PA14_43950 | sucC | succinyl-CoA synthetase beta subunit |
| PA14_44340 | ccoN2 | cytochrome oxidase subunit (cbb3-type) |
| PA14_44370 | ccoN1 | cytochrome oxidase subunit (cbb3-type) |
| PA14_44490 | anr | transcriptional regulator Anr |
| PA14_45630 | fliA | motility sigma factor FliA |
| PA14_45940 | lasI | autoinducer synthesis protein LasI |
| PA14_45960 | lasR | transcriptional regulator LasR |
| PA14_48060 | aprA | alkaline metalloproteinase precursor |
| PA14_48930 | PA14_48930 | pf5 phage gene |
| PA14_49250 | napA | periplasmic nitrate reductase protein NapA |
| PA14_50290 | fliC | flagellin type B |
| PA14_50360 | flgK | flagellar hook-associated protein 1 FlgK |
| PA14_51340 | mvfR | Transcriptional regulator MvfR |
| PA14_51410 | pqsC | PqsC |
| PA14_52180 | relA | GTP pyrophosphokinase |
| PA14_52270 | ldhA | D-lactate dehydrogenase (fermentative) |
| PA14_52580 | lysC | aspartate kinase alpha and beta chain |
| PA14_53470 | ackA | probable acetate kinase |
| PA14_54430 | algU | sigma factor AlgU |
| PA14_56220 | uspM | universal stress protein |
| PA14_56780 | sodB | iron superoxide dismutase |
| PA14_57940 | rpoN | RNA polymerase sigma-54 factor |
| PA14_58000 | sodM | superoxide dismutase |
| PA14_58730 | pilA | type IV pilin structural subunit |
| PA14_60310 | pilY1 | type 4 fimbrial biogenesis protein PilY1 |
| PA14_61040 | katB | catalase |
| PA14_61060 | fnr-2 | ferredoxin--NADP+ reductase |
| PA14_61200 | cdrA | conserved hypothetical protein |
| PA14_62160 | ilvI | acetolactate synthase large subunit |
| PA14_62710 | pnp | polyribonucleotide nucleotidyltransferase |
| PA14_62730 | truB | tRNA pseudouridine 55 synthase |

| | | |
|---|---|---|
| PA14_62860 | ftsH | cell division protein FtsH |
| PA14_68330 | arcA | Arginine fermentation |
| PA14_69190 | rho | transcription termination factor Rho |
| PA14_70470 | spoT | guanosine-3',5'-bis(diphosphate) 3'-pyrophosphohydrolase |
| PA14_70560 | OxyR | transcriptional regulator, LysR family |
| PA14_70860 | pstS | phosphate ABC transporter, periplasmic phosphate-binding protein |
| PA14_72960 | kgtP | MFS dicarboxylate transporter |
| PA14_73260 | atpA | ATP synthase alpha chain |
