## Supplemental Table 2 for "In situ single-cell activities of microbial populations revealed by spatial transcriptomics"

**Table S2: gene library used in study**

| Locus | Gene name | mRNA binding region sequence | Full primary probe sequence |
|---|---|---|---|
| P1_gp013 | pro | GACGAAAATCGGAGGTTATCCAGTTCGTC | TTTCGTCCGCGAGTGACCAGTAATCAAAGGCCGCAAATAATCAAAGGCCGCAAAGACGAAAATCGGAGGTTATCCAGTTCGTCAATAATCAAAGGCCGCAAATAATCAAAGGCCGCACAACGTCCATGTCGGGATGC |
| P1_gp013 | pro | ACGCTGGAACATTGTCGATTACGACCGGGC | TTTCGTCCGCGAGTGACCAGTAATCAAAGGCCGCAAATAATCAAAGGCCGCAAAACGCTGGAACATTGTCGATTACGACCGGGCAATAATCAAAGGCCGCAAATAATCAAAGGCCGCACAACGTCCATGTCGGGATGC |
| P1_gp013 | pro | CCAGGTTCTTTGAGGCGGAATGCGTCTTTT | TTTCGTCCGCGAGTGACCAGTAATCAAAGGCCGCAAATAATCAAAGGCCGCAAACCAGGTTCTTTGAGGCGGAATGCGTCTTTTAATAATCAAAGGCCGCAAATAATCAAAGGCCGCACAACGTCCATGTCGGGATGC |
| P1_gp013 | pro | TGGTTACAGCGATTTTCCCGGACTCGCGCC | TTTCGTCCGCGAGTGACCAGTAATCAAAGGCCGCAAATAATCAAAGGCCGCAAATGGTTACAGCGATTTTCCCGGACTCGCGCCAATAATCAAAGGCCGCAAATAATCAAAGGCCGCACAACGTCCATGTCGGGATGC |
| P1_gp013 | pro | GCTATAGAGACAAAAGCGGTCAGTGACCGT | TTTCGTCCGCGAGTGACCAGTAATCAAAGGCCGCAAATAATCAAAGGCCGCAAAGCTATAGAGACAAAAGCGGTCAGTGACCGTAATAATCAAAGGCCGCAAATAATCAAAGGCCGCACAACGTCCATGTCGGGATGC |
| P1_gp013 | pro | CTTGCTTTCAAGCATGGTCTTTACCGCGCT | TTTCGTCCGCGAGTGACCAGTAATCAAAGGCCGCAAATAATCAAAGGCCGCAAACTTGCTTTCAAGCATGGTCTTTACCGCGCTAATAATCAAAGGCCGCAAATAATCAAAGGCCGCACAACGTCCATGTCGGGATGC |
| P1_gp013 | pro | CGCCAGGGACTCTGCCAGTAAAGATTTAGA | TTTCGTCCGCGAGTGACCAGTAATCAAAGGCCGCAAATAATCAAAGGCCGCAAACGCCAGGGACTCTGCCAGTAAAGATTTAGAAATAATCAAAGGCCGCAAATAATCAAAGGCCGCACAACGTCCATGTCGGGATGC |
| P1_gp013 | pro | TAACGGCCTGCACTGACTCGTCGGAGTACC | TTTCGTCCGCGAGTGACCAGTAATCAAAGGCCGCAAATAATCAAAGGCCGCAAATAACGGCCTGCACTGACTCGTCGGAGTACCAATAATCAAAGGCCGCAAATAATCAAAGGCCGCACAACGTCCATGTCGGGATGC |
| P1_gp013 | pro | GCCTCCACCATCATTTCCAGTTCAGCCATT | TTTCGTCCGCGAGTGACCAGTAATCAAAGGCCGCAAATAATCAAAGGCCGCAAAGCCTCCACCATCATTTCCAGTTCAGCCATTAATAATCAAAGGCCGCAAATAATCAAAGGCCGCACAACGTCCATGTCGGGATGC |
| P1_gp013 | pro | GTGCAGTTTCCAGTTCTGCCGTACGCTCTG | TTTCGTCCGCGAGTGACCAGTAATCAAAGGCCGCAAATAATCAAAGGCCGCAAAGTGCAGTTTCCAGTTCTGCCGTACGCTCTGAATAATCAAAGGCCGCAAATAATCAAAGGCCGCACAACGTCCATGTCGGGATGC |
| P1_gp013 | pro | GGCGACAATTTTACCCGGCTCTGTGTTAAG | TTTCGTCCGCGAGTGACCAGTAATCAAAGGCCGCAAATAATCAAAGGCCGCAAAGGCGACAATTTTACCCGGCTCTGTGTTAAGAATAATCAAAGGCCGCAAATAATCAAAGGCCGCACAACGTCCATGTCGGGATGC |
| P1_gp013 | pro | CCTTCCACCATGATCACGCTGGTTTCTGGT | TTTCGTCCGCGAGTGACCAGTAATCAAAGGCCGCAAATAATCAAAGGCCGCAAACCTTCCACCATGATCACGCTGGTTTCTGGTAATAATCAAAGGCCGCAAATAATCAAAGGCCGCACAACGTCCATGTCGGGATGC |
| P1_gp013 | pro | GATGCTTGTCTCACGCGGTGCCTGGACGTT | TTTCGTCCGCGAGTGACCAGTAATCAAAGGCCGCAAATAATCAAAGGCCGCAAAGATGCTTGTCTCACGCGGTGCCTGGACGTTAATAATCAAAGGCCGCAAATAATCAAAGGCCGCACAACGTCCATGTCGGGATGC |
| P1_gp013 | pro | GCGTTGTCACATAATCCACACCATGGAAGC | TTTCGTCCGCGAGTGACCAGTAATCAAAGGCCGCAAATAATCAAAGGCCGCAAAGCGTTGTCACATAATCCACACCATGGAAGCAATAATCAAAGGCCGCAAATAATCAAAGGCCGCACAACGTCCATGTCGGGATGC |
| P1_gp013 | pro | AGCTCACCAAGGCGTAAACCTTCCTGAGTT | TTTCGTCCGCGAGTGACCAGTAATCAAAGGCCGCAAATAATCAAAGGCCGCAAAAGCTCACCAAGGCGTAAACCTTCCTGAGTTAATAATCAAAGGCCGCAAATAATCAAAGGCCGCACAACGTCCATGTCGGGATGC |
| P1_gp078 | dbn | CGGTCAGACGCTGAAGGTTCATAAGGATAC | TTTCGTCCGCGAGTGACCAGTGACGGTTGTAACGGAATGACGGTTGTAACGGAACGGTCAGACGCTGAAGGTTCATAAGGATACAATGACGGTTGTAACGGAATGACGGTTGTAACGGCAACGTCCATGTCGGGATGC |
| P1_gp078 | dbn | GCATGCCTATTTCCCTGGGTACCGTAGTAA | TTTCGTCCGCGAGTGACCAGTGACGGTTGTAACGGAATGACGGTTGTAACGGAAGCATGCCTATTTCCCTGGGTACCGTAGTAAAATGACGGTTGTAACGGAATGACGGTTGTAACGGCAACGTCCATGTCGGGATGC |
| P1_gp078 | dbn | GTTAATCCAAGGCTTTCGGGATGCGTAACG | TTTCGTCCGCGAGTGACCAGTGACGGTTGTAACGGAATGACGGTTGTAACGGAAGTTAATCCAAGGCTTTCGGGATGCGTAACGAATGACGGTTGTAACGGAATGACGGTTGTAACGGCAACGTCCATGTCGGGATGC |
| P1_gp078 | dbn | CTGTCGCAGCAGTCAACACAGCTTCAACAT | TTTCGTCCGCGAGTGACCAGTGACGGTTGTAACGGAATGACGGTTGTAACGGAACTGTCGCAGCAGTCAACACAGCTTCAACATAATGACGGTTGTAACGGAATGACGGTTGTAACGGCAACGTCCATGTCGGGATGC |
| P1_gp078 | dbn | GTCATCAGTCAACGTTCTGGCCTCATACGG | TTTCGTCCGCGAGTGACCAGTGACGGTTGTAACGGAATGACGGTTGTAACGGAAGTCATCAGTCAACGTTCTGGCCTCATACGGAATGACGGTTGTAACGGAATGACGGTTGTAACGGCAACGTCCATGTCGGGATGC |
| P1_gp078 | dbn | ATACCAACGCCCCGCACATTGCCGCCTCAA | TTTCGTCCGCGAGTGACCAGTGACGGTTGTAACGGAATGACGGTTGTAACGGAAATACCAACGCCCCGCACATTGCCGCCTCAAAATGACGGTTGTAACGGAATGACGGTTGTAACGGCAACGTCCATGTCGGGATGC |
| P1_gp078 | dbn | GCCGAATCCAGCATAGAACCGTCATATACC | TTTCGTCCGCGAGTGACCAGTGACGGTTGTAACGGAATGACGGTTGTAACGGAAGCCGAATCCAGCATAGAACCGTCATATACCAATGACGGTTGTAACGGAATGACGGTTGTAACGGCAACGTCCATGTCGGGATGC |
| P1_gp078 | dbn | TCCTGGTGCCAAAATGGCCTTCGCAGAACA | TTTCGTCCGCGAGTGACCAGTGACGGTTGTAACGGAATGACGGTTGTAACGGAATCCTGGTGCCAAAATGGCCTTCGCAGAACAAATGACGGTTGTAACGGAATGACGGTTGTAACGGCAACGTCCATGTCGGGATGC |
| P1_gp078 | dbn | GATGGTATTGCATTGTTACCCCTGCCTCAC | TTTCGTCCGCGAGTGACCAGTGACGGTTGTAACGGAATGACGGTTGTAACGGAAGATGGTATTGCATTGTTACCCCTGCCTCACAATGACGGTTGTAACGGAATGACGGTTGTAACGGCAACGTCCATGTCGGGATGC |
| P1_gp078 | dbn | GACGTTCTAACCGACCGGGATTATCCAGGT | TTTCGTCCGCGAGTGACCAGTGACGGTTGTAACGGAATGACGGTTGTAACGGAAGACGTTCTAACCGACCGGGATTATCCAGGTAATGACGGTTGTAACGGAATGACGGTTGTAACGGCAACGTCCATGTCGGGATGC |
| P1_gp078 | dbn | GGCTACCCATCCAGAGAATCGGCGACACAG | TTTCGTCCGCGAGTGACCAGTGACGGTTGTAACGGAATGACGGTTGTAACGGAAGGCTACCCATCCAGAGAATCGGCGACACAGAATGACGGTTGTAACGGAATGACGGTTGTAACGGCAACGTCCATGTCGGGATGC |
| P1_gp078 | dbn | GACGGTTTCTGCGTGATAAACGTCTCAAGC | TTTCGTCCGCGAGTGACCAGTGACGGTTGTAACGGAATGACGGTTGTAACGGAAGACGGTTTCTGCGTGATAAACGTCTCAAGCAATGACGGTTGTAACGGAATGACGGTTGTAACGGCAACGTCCATGTCGGGATGC |
| P1_gp078 | dbn | GCAGGCCAGGTTATAAGACCACATTTCCGT | TTTCGTCCGCGAGTGACCAGTGACGGTTGTAACGGAATGACGGTTGTAACGGAAGCAGGCCAGGTTATAAGACCACATTTCCGTAATGACGGTTGTAACGGAATGACGGTTGTAACGGCAACGTCCATGTCGGGATGC |
| P1_gp078 | dbn | CGCTGCCCATCCAACATGTTTGTTAAAGCG | TTTCGTCCGCGAGTGACCAGTGACGGTTGTAACGGAATGACGGTTGTAACGGAACGCTGCCCATCCAACATGTTTGTTAAAGCGAATGACGGTTGTAACGGAATGACGGTTGTAACGGCAACGTCCATGTCGGGATGC |
| P1_gp078 | dbn | CGTTTTGGTTTCCAGATGGAACAGCTCCTG | TTTCGTCCGCGAGTGACCAGTGACGGTTGTAACGGAATGACGGTTGTAACGGAACGTTTTGGTTTCCAGATGGAACAGCTCCTGAATGACGGTTGTAACGGAATGACGGTTGTAACGGCAACGTCCATGTCGGGATGC |
| P1_gp003 | cre | CTGTTTTGCACGTTCACCGGCATCAACGTT | TTTCGTCCGCGAGTGACCAGGCGGTGATTGCACCAAAGCGGTGATTGCACCAAACTGTTTTGCACGTTCACCGGCATCAACGTTAAGCGGTGATTGCACCAAAGCGGTGATTGCACCACAACGTCCATGTCGGGATGC |
| P1_gp003 | cre | AATCCATCGCTCGACCAGTTTAGTTACCCC | TTTCGTCCGCGAGTGACCAGGCGGTGATTGCACCAAAGCGGTGATTGCACCAAAAATCCATCGCTCGACCAGTTTAGTTACCCCAAGCGGTGATTGCACCAAAGCGGTGATTGCACCACAACGTCCATGTCGGGATGC |
| P1_gp003 | cre | TGGTCCAGCCACCAGCTTGCATGATCTCCG | TTTCGTCCGCGAGTGACCAGGCGGTGATTGCACCAAAGCGGTGATTGCACCAAATGGTCCAGCCACCAGCTTGCATGATCTCCGAAGCGGTGATTGCACCAAAGCGGTGATTGCACCACAACGTCCATGTCGGGATGC |
| P1_gp003 | cre | ATTCAACTTGCACCATGCCGCCCACGACCG | TTTCGTCCGCGAGTGACCAGGCGGTGATTGCACCAAAGCGGTGATTGCACCAAAATTCAACTTGCACCATGCCGCCCACGACCGAAGCGGTGATTGCACCAAAGCGGTGATTGCACCACAACGTCCATGTCGGGATGC |
| P1_gp003 | cre | GTTCTTGCGAACCTCATCACTCGTTGCATC | TTTCGTCCGCGAGTGACCAGGCGGTGATTGCACCAAAGCGGTGATTGCACCAAAGTTCTTGCGAACCTCATCACTCGTTGCATCAAGCGGTGATTGCACCAAAGCGGTGATTGCACCACAACGTCCATGTCGGGATGC |
| P1_gp003 | cre | GCCCCTGTTTCACTATCCAGGTTACGGATA | TTTCGTCCGCGAGTGACCAGGCGGTGATTGCACCAAAGCGGTGATTGCACCAAAGCCCCTGTTTCACTATCCAGGTTACGGATAAAGCGGTGATTGCACCAAAGCGGTGATTGCACCACAACGTCCATGTCGGGATGC |
| P1_gp003 | cre | GACCAGAGTCATCCTTAGCGCCGTAAATCA | TTTCGTCCGCGAGTGACCAGGCGGTGATTGCACCAAAGCGGTGATTGCACCAAAGACCAGAGTCATCCTTAGCGCCGTAAATCAAAGCGGTGATTGCACCAAAGCGGTGATTGCACCACAACGTCCATGTCGGGATGC |
| P1_gp003 | cre | CGGACAGAAGCATTTTCAGGTATGCTCAG | TTTCGTCCGCGAGTGACCAGGCGGTGATTGCACCAAAGCGGTGATTGCACCAAACGGACAGAAGCATTTTCAGGTATGCTCAGAAGCGGTGATTGCACCAAAGCGGTGATTGCACCACAACGTCCATGTCGGGATGC |
| P1_gp003 | cre | CATTCTCCCACCGTCAGTACGTGAGATATC | TTTCGTCCGCGAGTGACCAGGCGGTGATTGCACCAAAGCGGTGATTGCACCAAACATTCTCCCACCGTCAGTACGTGAGATATCAAGCGGTGATTGCACCAAAGCGGTGATTGCACCACAACGTCCATGTCGGGATGC |
| P1_gp003 | cre | CTGGCAATTTCGGCTATACGTAACAGGGTG | TTTCGTCCGCGAGTGACCAGGCGGTGATTGCACCAAAGCGGTGATTGCACCAAACTGGCAATTTCGGCTATACGTAACAGGGTGAAGCGGTGATTGCACCAAAGCGGTGATTGCACCACAACGTCCATGTCGGGATGC |
| P1_gp003 | cre | ATTGCTGTCACTTGGTCGTGGCAGCCCGGA | TTTCGTCCGCGAGTGACCAGGCGGTGATTGCACCAAAGCGGTGATTGCACCAAAATTGCTGTCACTTGGTCGTGGCAGCCCGGAAAGCGGTGATTGCACCAAAGCGGTGATTGCACCACAACGTCCATGTCGGGATGC |
| P1_gp003 | cre | CCGGTAATGCAGGCAAATTTTGGTGTACGG | TTTCGTCCGCGAGTGACCAGGCGGTGATTGCACCAAAGCGGTGATTGCACCAAACCGGTAATGCAGGCAAATTTTGGTGTACGGAAGCGGTGATTGCACCAAAGCGGTGATTGCACCACAACGTCCATGTCGGGATGC |
| P1_gp003 | cre | GGCACTGTGTCCAGACCAGGCCAGGTATCT | TTTCGTCCGCGAGTGACCAGGCGGTGATTGCACCAAAGCGGTGATTGCACCAAAGGCACTGTGTCCAGACCAGGCCAGGTATCTAAGCGGTGATTGCACCAAAGCGGTGATTGCACCACAACGTCCATGTCGGGATGC |
| P1_gp003 | cre | TTTCGGATCCGCCGCATAACCAGTGAAACA | TTTCGTCCGCGAGTGACCAGGCGGTGATTGCACCAAAGCGGTGATTGCACCAAATTTCGGATCCGCCGCATAACCAGTGAAACAAAGCGGTGATTGCACCAAAGCGGTGATTGCACCACAACGTCCATGTCGGGATGC |
| P1_gp003 | cre | GCAGCGATCGCTATTTTCCATGAGTGAACG | TTTCGTCCGCGAGTGACCAGGCGGTGATTGCACCAAAGCGGTGATTGCACCAAAGCAGCGATCGCTATTTTCCATGAGTGAACGAAGCGGTGATTGCACCAAAGCGGTGATTGCACCACAACGTCCATGTCGGGATGC |
| P1_gp003 | cre | AACGCCTGGCGATCCCTGAACATGTCCATC | TTTCGTCCGCGAGTGACCAGGCGGTGATTGCACCAAAGCGGTGATTGCACCAAAAACGCCTGGCGATCCCTGAACATGTCCATCAAGCGGTGATTGCACCAAAGCGGTGATTGCACCACAACGTCCATGTCGGGATGC |
| P1_gp003 | cre | GACCCGGCAAAACAGGTAGTTATTCGGATC | TTTCGTCCGCGAGTGACCAGGCGGTGATTGCACCAAAGCGGTGATTGCACCAAAGACCCGGCAAAACAGGTAGTTATTCGGATCAAGCGGTGATTGCACCAAAGCGGTGATTGCACCACAACGTCCATGTCGGGATGC |
| P1_gp003 | cre | AATCCCTTCCAGGGCGCGAGTTGATAGCTG | TTTCGTCCGCGAGTGACCAGGCGGTGATTGCACCAAAGCGGTGATTGCACCAAAAATCCCTTCCAGGGCGCGAGTTGATAGCTGAAGCGGTGATTGCACCAAAGCGGTGATTGCACCACAACGTCCATGTCGGGATGC |
| P1_gp003 | cre | GCATGTTTAGCTGGCCCAAATGTTGCTGGA | TTTCGTCCGCGAGTGACCAGGCGGTGATTGCACCAAAGCGGTGATTGCACCAAAGCATGTTTAGCTGGCCCAAATGTTGCTGGAAAGCGGTGATTGCACCAAAGCGGTGATTGCACCACAACGTCCATGTCGGGATGC |
| P1_gp003 | cre | TGGTCGAAATCAGTGCGTTCGAACGCTAGA | TTTCGTCCGCGAGTGACCAGGCGGTGATTGCACCAAAGCGGTGATTGCACCAAATGGTCGAAATCAGTGCGTTCGAACGCTAGAAAGCGGTGATTGCACCAAAGCGGTGATTGCACCACAACGTCCATGTCGGGATGC |
| PA14_00010 | dnaA | ATGGCGTTGAGCTGCAAGGCCTTCACCATG | TTTCGTCCGCGAGTGACCAGTGCTCCCGGATTACAAATGCTCCCGGATTACAAAATGGCGTTGAGCTGCAAGGCCTTCACCATGAATGCTCCCGGATTACAAATGCTCCCGGATTACACAACGTCCATGTCGGGATGC |
| PA14_00010 | dnaA | TGGTTGGTCAGCTCCTTGGAGAGCGCCATG | TTTCGTCCGCGAGTGACCAGTGCTCCCGGATTACAAATGCTCCCGGATTACAAATGGTTGGTCAGCTCCTTGGAGAGCGCCATGAATGCTCCCGGATTACAAATGCTCCCGGATTACACAACGTCCATGTCGGGATGC |
| PA14_00010 | dnaA | TTTCAGCCGCTCTTCCAGGCCTTCGATTTC | TTTCGTCCGCGAGTGACCAGTGCTCCCGGATTACAAATGCTCCCGGATTACAAATTTCAGCCGCTCTTCCAGGCCTTCGATTTCAATGCTCCCGGATTACAAATGCTCCCGGATTACACAACGTCCATGTCGGGATGC |
| PA14_00010 | dnaA | ACGACAGGCGTGCAACACCGTGGTGTGATC | TTTCGTCCGCGAGTGACCAGTGCTCCCGGATTACAAATGCTCCCGGATTACAAAACGACAGGCGTGCAACACCGTGGTGTGATCAATGCTCCCGGATTACAAATGCTCCCGGATTACACAACGTCCATGTCGGGATGC |
| PA14_00010 | dnaA | TCGGATAGCGGTCGCTGGTGAGGATCACCT | TTTCGTCCGCGAGTGACCAGTGCTCCCGGATTACAAATGCTCCCGGATTACAAATCGGATAGCGGTCGCTGGTGAGGATCACCTAATGCTCCCGGATTACAAATGCTCCCGGATTACACAACGTCCATGTCGGGATGC |
| PA14_00010 | dnaA | ACGAAGTTCTCGAAGGTGAAGGTACGGTTG | TTTCGTCCGCGAGTGACCAGTGCTCCCGGATTACAAATGCTCCCGGATTACAAAACGAAGTTCTCGAAGGTGAAGGTACGGTTGAATGCTCCCGGATTACAAATGCTCCCGGATTACACAACGTCCATGTCGGGATGC |
| PA14_00010 | dnaA | CGACCTGCAAGGGACGGATCCAGGTGTTGA | TTTCGTCCGCGAGTGACCAGTGCTCCCGGATTACAAATGCTCCCGGATTACAAACGACCTGCAAGGGACGGATCCAGGTGTTGAAATGCTCCCGGATTACAAATGCTCCCGGATTACACAACGTCCATGTCGGGATGC |
| PA14_00010 | dnaA | CGAGCAGTTCCAGAAGCCGACCGAGGTATT | TTTCGTCCGCGAGTGACCAGTGCTCCCGGATTACAAATGCTCCCGGATTACAAACGAGCAGTTCCAGAAGCCGACCGAGGTATTAATGCTCCCGGATTACAAATGCTCCCGGATTACACAACGTCCATGTCGGGATGC |
| PA14_00010 | dnaA | TGTTGGGACGGCAGCTCATCGCGGAGAAGA | TTTCGTCCGCGAGTGACCAGTGCTCCCGGATTACAAATGCTCCCGGATTACAAATGTTGGGACGGCAGCTCATCGCGGAGAAGAAATGCTCCCGGATTACAAATGCTCCCGGATTACACAACGTCCATGTCGGGATGC |
| PA14_00010 | dnaA | AAGAACTCCTCCTGGGAGCGCTCCTTACGG | TTTCGTCCGCGAGTGACCAGTGCTCCCGGATTACAAATGCTCCCGGATTACAAAAAGAACTCCTCCTGGGAGCGCTCCTTACGGAATGCTCCCGGATTACAAATGCTCCCGGATTACACAACGTCCATGTCGGGATGC |
| PA14_00010 | dnaA | CCACCGAGCGGTAGAAGCGCTTGAATTCGT | TTTCGTCCGCGAGTGACCAGTGCTCCCGGATTACAAATGCTCCCGGATTACAAACCACCGAGCGGTAGAAGCGCTTGAATTCGTAATGCTCCCGGATTACAAATGCTCCCGGATTACACAACGTCCATGTCGGGATGC |
| PA14_00010 | dnaA | CGCGACGAAACGTTCCGAATGCAGGTAGAC | TTTCGTCCGCGAGTGACCAGTGCTCCCGGATTACAAATGCTCCCGGATTACAAACGCGACGAAACGTTCCGAATGCAGGTAGACAATGCTCCCGGATTACAAATGCTCCCGGATTACACAACGTCCATGTCGGGATGC |
| PA14_00010 | dnaA | GCCGCTTGGACAACAGATCGGATATCTTGA | TTTCGTCCGCGAGTGACCAGTGCTCCCGGATTACAAATGCTCCCGGATTACAAAGCCGCTTGGACAACAGATCGGATATCTTGAAATGCTCCCGGATTACAAATGCTCCCGGATTACACAACGTCCATGTCGGGATGC |
| PA14_00010 | dnaA | TCCGCGTCGGATTCCCTAAGTTGAGCGATC | TTTCGTCCGCGAGTGACCAGTGCTCCCGGATTACAAATGCTCCCGGATTACAAATCCGCGTCGGATTCCCTAAGTTGAGCGATCAATGCTCCCGGATTACAAATGCTCCCGGATTACACAACGTCCATGTCGGGATGC |
| PA14_00010 | dnaA | CACGCACTGCTGCCAAAGTTCCACGGACAC | TTTCGTCCGCGAGTGACCAGTGCTCCCGGATTACAAATGCTCCCGGATTACAAACACGCACTGCTGCCAAAGTTCCACGGACACAATGCTCCCGGATTACAAATGCTCCCGGATTACACAACGTCCATGTCGGGATGC |
| PA14_09520 | mexI | TGACGAAGCCTTGCATGAGATCGGCGGAGG | TTTCGTCCGCGAGTGACCAGAATGACGCGACATAGAAAATGACGCGACATAGAATGACGAAGCCTTGCATGAGATCGGCGGAGGAAAATGACGCGACATAGAAAATGACGCGACATAGCAACGTCCATGTCGGGATGC |
| PA14_09520 | mexI | GTTTCCGGGATGCTGGTGTAGACCTGGTCG | TTTCGTCCGCGAGTGACCAGAATGACGCGACATAGAAAATGACGCGACATAGAAGTTTCCGGGATGCTGGTGTAGACCTGGTCGAAAATGACGCGACATAGAAAATGACGCGACATAGCAACGTCCATGTCGGGATGC |
| PA14_09520 | mexI | TGTTCGGTTCGACCTGAAGCTCCACCCGGA | TTTCGTCCGCGAGTGACCAGAATGACGCGACATAGAAAATGACGCGACATAGAATGTTCGGTTCGACCTGAAGCTCCACCCGGAAAAATGACGCGACATAGAAAATGACGCGACATAGCAACGTCCATGTCGGGATGC |
| PA14_09520 | mexI | TGCCATCGTTGCGGATGACCATCTCGCGGA | TTTCGTCCGCGAGTGACCAGAATGACGCGACATAGAAAATGACGCGACATAGAATGCCATCGTTGCGGATGACCATCTCGCGGAAAAATGACGCGACATAGAAAATGACGCGACATAGCAACGTCCATGTCGGGATGC |
| PA14_09520 | mexI | TCGTGGAGTTGCAGTTCGTTGGCGAACTCG | TTTCGTCCGCGAGTGACCAGAATGACGCGACATAGAAAATGACGCGACATAGAATCGTGGAGTTGCAGTTCGTTGGCGAACTCGAAAATGACGCGACATAGAAAATGACGCGACATAGCAACGTCCATGTCGGGATGC |
| PA14_09520 | mexI | TGAGCAGCATGCGGATGGTCACCACGCTAC | TTTCGTCCGCGAGTGACCAGAATGACGCGACATAGAAAATGACGCGACATAGAATGAGCAGCATGCGGATGGTCACCACGCTACAAAATGACGCGACATAGAAAATGACGCGACATAGCAACGTCCATGTCGGGATGC |
| PA14_09520 | mexI | ATGGTCGAGCTTTCCAGCAGCGGGTACTGG | TTTCGTCCGCGAGTGACCAGAATGACGCGACATAGAAAATGACGCGACATAGAAATGGTCGAGCTTTCCAGCAGCGGGTACTGGAAAATGACGCGACATAGAAAATGACGCGACATAGCAACGTCCATGTCGGGATGC |
| PA14_09520 | mexI | AAAGGTCTGGACCTTGGCCACGCCGTCGAT | TTTCGTCCGCGAGTGACCAGAATGACGCGACATAGAAAATGACGCGACATAGAAAAAGGTCTGGACCTTGGCCACGCCGTCGATAAAATGACGCGACATAGAAAATGACGCGACATAGCAACGTCCATGTCGGGATGC |
| PA14_09520 | mexI | TTCGCGCTTGGGCATCGAATACAGCAGCGG | TTTCGTCCGCGAGTGACCAGAATGACGCGACATAGAAAATGACGCGACATAGAATTCGCGCTTGGGCATCGAATACAGCAGCGGAAAATGACGCGACATAGAAAATGACGCGACATAGCAACGTCCATGTCGGGATGC |
| PA14_09520 | mexI | TGGACAGCACGTACTGCCCCTTCACCATCC | TTTCGTCCGCGAGTGACCAGAATGACGCGACATAGAAAATGACGCGACATAGAATGGACAGCACGTACTGCCCCTTCACCATCCAAAATGACGCGACATAGAAAATGACGCGACATAGCAACGTCCATGTCGGGATGC |
| PA14_09520 | mexI | CATGAGGATGCCGTGCTTGCTGATCAGGCC | TTTCGTCCGCGAGTGACCAGAATGACGCGACATAGAAAATGACGCGACATAGAACATGAGGATGCCGTGCTTGCTGATCAGGCCAAAATGACGCGACATAGAAAATGACGCGACATAGCAACGTCCATGTCGGGATGC |
| PA14_09520 | mexI | ATGGTCTCGGTCATCGCCTGGGTCGAATCG | TTTCGTCCGCGAGTGACCAGAATGACGCGACATAGAAAATGACGCGACATAGAAATGGTCTCGGTCATCGCCTGGGTCGAATCGAAAATGACGCGACATAGAAAATGACGCGACATAGCAACGTCCATGTCGGGATGC |
| PA14_09520 | mexI | CGTCGGTGCCGTTGATGATCCAGGTGCTCA | TTTCGTCCGCGAGTGACCAGAATGACGCGACATAGAAAATGACGCGACATAGAACGTCGGTGCCGTTGATGATCCAGGTGCTCAAAAATGACGCGACATAGAAAATGACGCGACATAGCAACGTCCATGTCGGGATGC |
| PA14_09520 | mexI | AAGGTAGTCGATGCCCTCCACCGACGACAC | TTTCGTCCGCGAGTGACCAGAATGACGCGACATAGAAAATGACGCGACATAGAAAAGGTAGTCGATGCCCTCCACCGACGACACAAAATGACGCGACATAGAAAATGACGCGACATAGCAACGTCCATGTCGGGATGC |
| PA14_09520 | mexI | CCGGCGGACGAACAGGTCGGTAAAGGTCAT | TTTCGTCCGCGAGTGACCAGAATGACGCGACATAGAAAATGACGCGACATAGAACCGGCGGACGAACAGGTCGGTAAAGGTCATAAAATGACGCGACATAGAAAATGACGCGACATAGCAACGTCCATGTCGGGATGC |
| PA14_25560 | rne | ACTTCCTTGATGTTGATCCGGCCTTCGGGG | TTTCGTCCGCGAGTGACCAGAGGGCCACGCACAAAAAAGGGCCACGCACAAAAAACTTCCTTGATGTTGATCCGGCCTTCGGGGAAAGGGCCACGCACAAAAAAGGGCCACGCACAAACAACGTCCATGTCGGGATGC |
| PA14_25560 | rne | AACAGGCGTTGGCCGTCGACCAGTGCAACA | TTTCGTCCGCGAGTGACCAGAGGGCCACGCACAAAAAAGGGCCACGCACAAAAAAACAGGCGTTGGCCGTCGACCAGTGCAACAAAAGGGCCACGCACAAAAAAGGGCCACGCACAAACAACGTCCATGTCGGGATGC |
| PA14_25560 | rne | TTTTCGCGTGCTCTGTGTCGGCGGCGTCTT | TTTCGTCCGCGAGTGACCAGAGGGCCACGCACAAAAAAGGGCCACGCACAAAAATTTTCGCGTGCTCTGTGTCGGCGGCGTCTTAAAGGGCCACGCACAAAAAAGGGCCACGCACAAACAACGTCCATGTCGGGATGC |
| PA14_25560 | rne | TTTCCTGGTAGATCAGGAAGGGCGCGCCAC | TTTCGTCCGCGAGTGACCAGAGGGCCACGCACAAAAAAGGGCCACGCACAAAAATTTCCTGGTAGATCAGGAAGGGCGCGCCACAAAGGGCCACGCACAAAAAAGGGCCACGCACAAACAACGTCCATGTCGGGATGC |
| PA14_25560 | rne | AGACGCTGGACTTCGAAATGCGGGGTTTCC | TTTCGTCCGCGAGTGACCAGAGGGCCACGCACAAAAAAGGGCCACGCACAAAAAAGACGCTGGACTTCGAAATGCGGGGTTTCCAAAGGGCCACGCACAAAAAAGGGCCACGCACAAACAACGTCCATGTCGGGATGC |
| PA14_25560 | rne | AGGTTGATCCTTGCCTGCGAACAGGCCGAC | TTTCGTCCGCGAGTGACCAGAGGGCCACGCACAAAAAAGGGCCACGCACAAAAAAGGTTGATCCTTGCCTGCGAACAGGCCGACAAAGGGCCACGCACAAAAAAGGGCCACGCACAAACAACGTCCATGTCGGGATGC |
| PA14_25560 | rne | GAAGTATTCGCGGGAGATTTCTTTGAGGGG | TTTCGTCCGCGAGTGACCAGAGGGCCACGCACAAAAAAGGGCCACGCACAAAAAGAAGTATTCGCGGGAGATTTCTTTGAGGGGAAAGGGCCACGCACAAAAAAGGGCCACGCACAAACAACGTCCATGTCGGGATGC |
| PA14_25560 | rne | AAGTTCAGGGCTTCTTCCTGGGCGTCGATG | TTTCGTCCGCGAGTGACCAGAGGGCCACGCACAAAAAAGGGCCACGCACAAAAAAAGTTCAGGGCTTCTTCCTGGGCGTCGATGAAAGGGCCACGCACAAAAAAGGGCCACGCACAAACAACGTCCATGTCGGGATGC |
| PA14_25560 | rne | GCGGTCTCGATCTGGCTCTCGATCTGGAAG | TTTCGTCCGCGAGTGACCAGAGGGCCACGCACAAAAAAGGGCCACGCACAAAAAGCGGTCTCGATCTGGCTCTCGATCTGGAAGAAAGGGCCACGCACAAAAAAGGGCCACGCACAAACAACGTCCATGTCGGGATGC |
| PA14_25560 | rne | AACTCCTCGGGTTGAGTCGCGTTGATCAGC | TTTCGTCCGCGAGTGACCAGAGGGCCACGCACAAAAAAGGGCCACGCACAAAAAAACTCCTCGGGTTGAGTCGCGTTGATCAGCAAAGGGCCACGCACAAAAAAGGGCCACGCACAAACAACGTCCATGTCGGGATGC |
| PA14_25560 | rne | CTTCTACGGCTTCGCTGGCGACAGCGATAC | TTTCGTCCGCGAGTGACCAGAGGGCCACGCACAAAAAAGGGCCACGCACAAAAACTTCTACGGCTTCGCTGGCGACAGCGATACAAAGGGCCACGCACAAAAAAGGGCCACGCACAAACAACGTCCATGTCGGGATGC |
| PA14_25560 | rne | TTCTTCAGCCTGGGGTGCAGCAGGTTCCTC | TTTCGTCCGCGAGTGACCAGAGGGCCACGCACAAAAAAGGGCCACGCACAAAAATTCTTCAGCCTGGGGTGCAGCAGGTTCCTCAAAGGGCCACGCACAAAAAAGGGCCACGCACAAACAACGTCCATGTCGGGATGC |
| PA14_25560 | rne | TTCATCGCTTGCCGTGGTTTCGCTGGCAGC | TTTCGTCCGCGAGTGACCAGAGGGCCACGCACAAAAAAGGGCCACGCACAAAAATTCATCGCTTGCCGTGGTTTCGCTGGCAGCAAAGGGCCACGCACAAAAAAGGGCCACGCACAAACAACGTCCATGTCGGGATGC |
| PA14_25560 | rne | CTTCCTTGATCGCGCTCCACAGTTGCAGCA | TTTCGTCCGCGAGTGACCAGAGGGCCACGCACAAAAAAGGGCCACGCACAAAAACTTCCTTGATCGCGCTCCACAGTTGCAGCAAAAGGGCCACGCACAAAAAAGGGCCACGCACAAACAACGTCCATGTCGGGATGC |
| PA14_25560 | rne | TAGATGTTGGCCTTCTTCTGCTCGCGGGCG | TTTCGTCCGCGAGTGACCAGAGGGCCACGCACAAAAAAGGGCCACGCACAAAAATAGATGTTGGCCTTCTTCTGCTCGCGGGCGAAAGGGCCACGCACAAAAAAGGGCCACGCACAAACAACGTCCATGTCGGGATGC |
| PA14_25560 | rne | TCACCTTGCTGGCGTACTGCGGCATCACCT | TTTCGTCCGCGAGTGACCAGAGGGCCACGCACAAAAAAGGGCCACGCACAAAAATCACCTTGCTGGCGTACTGCGGCATCACCTAAAGGGCCACGCACAAAAAAGGGCCACGCACAAACAACGTCCATGTCGGGATGC |
| PA14_25560 | rne | TGATGTCGATGGAAACCAGGGCCTCGGTCG | TTTCGTCCGCGAGTGACCAGAGGGCCACGCACAAAAAAGGGCCACGCACAAAAATGATGTCGATGGAAACCAGGGCCTCGGTCGAAAGGGCCACGCACAAAAAAGGGCCACGCACAAACAACGTCCATGTCGGGATGC |
| PA14_25560 | rne | TTGTTCGGCATCAGCACCAGGTAACGGCCG | TTTCGTCCGCGAGTGACCAGAGGGCCACGCACAAAAAAGGGCCACGCACAAAAATTGTTCGGCATCAGCACCAGGTAACGGCCGAAAGGGCCACGCACAAAAAAGGGCCACGCACAAACAACGTCCATGTCGGGATGC |
| PA14_25560 | rne | CCTGGACGATGACTTCCTGGCCTTCGCTCA | TTTCGTCCGCGAGTGACCAGAGGGCCACGCACAAAAAAGGGCCACGCACAAAAACCTGGACGATGACTTCCTGGCCTTCGCTCAAAAGGGCCACGCACAAAAAAGGGCCACGCACAAACAACGTCCATGTCGGGATGC |
| PA14_25560 | rne | TAGTCGAGGTCCCACTGCAGTTCCTCGGTG | TTTCGTCCGCGAGTGACCAGAGGGCCACGCACAAAAAAGGGCCACGCACAAAAATAGTCGAGGTCCCACTGCAGTTCCTCGGTGAAAGGGCCACGCACAAAAAAGGGCCACGCACAAACAACGTCCATGTCGGGATGC |
| PA14_30050 | aceA | CGCGCTCCAGCTGCTGCCGTTCTTCTCTTT | TTTCGTCCGCGAGTGACCAGCGCGGTGTTCAGGATAACGCGGTGTTCAGGATAACGCGCTCCAGCTGCTGCCGTTCTTCTCTTTAACGCGGTGTTCAGGATAACGCGGTGTTCAGGATCAACGTCCATGTCGGGATGC |
| PA14_30050 | aceA | CCAGTTGAAGGACGGGCTGTTGTTGTAGAC | TTTCGTCCGCGAGTGACCAGCGCGGTGTTCAGGATAACGCGGTGTTCAGGATAACCAGTTGAAGGACGGGCTGTTGTTGTAGACAACGCGGTGTTCAGGATAACGCGGTGTTCAGGATCAACGTCCATGTCGGGATGC |
| PA14_30050 | aceA | GTGGTCTTCAGGTGCTTCTTGATCGAGATC | TTTCGTCCGCGAGTGACCAGCGCGGTGTTCAGGATAACGCGGTGTTCAGGATAAGTGGTCTTCAGGTGCTTCTTGATCGAGATCAACGCGGTGTTCAGGATAACGCGGTGTTCAGGATCAACGTCCATGTCGGGATGC |
| PA14_30050 | aceA | ATACGCATGCGAGCTGCGTACTCGGGGTTG | TTTCGTCCGCGAGTGACCAGCGCGGTGTTCAGGATAACGCGGTGTTCAGGATAAATACGCATGCGAGCTGCGTACTCGGGGTTGAACGCGGTGTTCAGGATAACGCGGTGTTCAGGATCAACGTCCATGTCGGGATGC |
| PA14_30050 | aceA | AGCTCGGTGTCGTCGTACTCGACGCTCATC | TTTCGTCCGCGAGTGACCAGCGCGGTGTTCAGGATAACGCGGTGTTCAGGATAAAGCTCGGTGTCGTCGTACTCGACGCTCATCAACGCGGTGTTCAGGATAACGCGGTGTTCAGGATCAACGTCCATGTCGGGATGC |
| PA14_30050 | aceA | CGGTGTACTTGGCGATATCCAGGCCGGTCT | TTTCGTCCGCGAGTGACCAGCGCGGTGTTCAGGATAACGCGGTGTTCAGGATAACGGTGTACTTGGCGATATCCAGGCCGGTCTAACGCGGTGTTCAGGATAACGCGGTGTTCAGGATCAACGTCCATGTCGGGATGC |
| PA14_30050 | aceA | CTGCCTTGATCTCGTTCTGATATGCGGACA | TTTCGTCCGCGAGTGACCAGCGCGGTGTTCAGGATAACGCGGTGTTCAGGATAACTGCCTTGATCTCGTTCTGATATGCGGACAAACGCGGTGTTCAGGATAACGCGGTGTTCAGGATCAACGTCCATGTCGGGATGC |
| PA14_30050 | aceA | TTCTGCTGCCCGATGAAGCCGTGCCAGCAA | TTTCGTCCGCGAGTGACCAGCGCGGTGTTCAGGATAACGCGGTGTTCAGGATAATTCTGCTGCCCGATGAAGCCGTGCCAGCAAAACGCGGTGTTCAGGATAACGCGGTGTTCAGGATCAACGTCCATGTCGGGATGC |
| PA14_30050 | aceA | GGCGTAGCGTACGGCATTGATCTTGGCGAG | TTTCGTCCGCGAGTGACCAGCGCGGTGTTCAGGATAACGCGGTGTTCAGGATAAGGCGTAGCGTACGGCATTGATCTTGGCGAGAACGCGGTGTTCAGGATAACGCGGTGTTCAGGATCAACGTCCATGTCGGGATGC |
| PA14_30050 | aceA | TTTTCTCGTGCATGGACTGGTCCGGCAGCG | TTTCGTCCGCGAGTGACCAGCGCGGTGTTCAGGATAACGCGGTGTTCAGGATAATTTTCTCGTGCATGGACTGGTCCGGCAGCGAACGCGGTGTTCAGGATAACGCGGTGTTCAGGATCAACGTCCATGTCGGGATGC |
| PA14_30050 | aceA | TGATGCAGTCCAGGACGCAGCGATCTTCGC | TTTCGTCCGCGAGTGACCAGCGCGGTGTTCAGGATAACGCGGTGTTCAGGATAATGATGCAGTCCAGGACGCAGCGATCTTCGCAACGCGGTGTTCAGGATAACGCGGTGTTCAGGATCAACGTCCATGTCGGGATGC |
| PA14_30050 | aceA | AGCGACTGGGTGTAGACGGAAGAGTCCGCA | TTTCGTCCGCGAGTGACCAGCGCGGTGTTCAGGATAACGCGGTGTTCAGGATAAAGCGACTGGGTGTAGACGGAAGAGTCCGCAAACGCGGTGTTCAGGATAACGCGGTGTTCAGGATCAACGTCCATGTCGGGATGC |
| PA14_30050 | aceA | AAGGTGTACAGCTCTTCGATCAGGCCGGAA | TTTCGTCCGCGAGTGACCAGCGCGGTGTTCAGGATAACGCGGTGTTCAGGATAAAAGGTGTACAGCTCTTCGATCAGGCCGGAAAACGCGGTGTTCAGGATAACGCGGTGTTCAGGATCAACGTCCATGTCGGGATGC |
| PA14_30050 | aceA | TCCAACCGGACAGGTAGAGGTAGCGCTTGT | TTTCGTCCGCGAGTGACCAGCGCGGTGTTCAGGATAACGCGGTGTTCAGGATAATCCAACCGGACAGGTAGAGGTAGCGCTTGTAACGCGGTGTTCAGGATAACGCGGTGTTCAGGATCAACGTCCATGTCGGGATGC |
| PA14_30050 | aceA | GTATTCGGCCATGTCCTTGCGCATGATGGC | TTTCGTCCGCGAGTGACCAGCGCGGTGTTCAGGATAACGCGGTGTTCAGGATAAGTATTCGGCCATGTCCTTGCGCATGATGGCAACGCGGTGTTCAGGATAACGCGGTGTTCAGGATCAACGTCCATGTCGGGATGC |
| PA14_53470 | ackA | CAGGGTGCGCATGTCGTTGGACAGGTCGGA | TTTCGTCCGCGAGTGACCAGCATGAATTCTCGCCAAACATGAATTCTCGCCAAACAGGGTGCGCATGTCGTTGGACAGGTCGGAAACATGAATTCTCGCCAAACATGAATTCTCGCCACAACGTCCATGTCGGGATGC |
| PA14_53470 | ackA | TTGGGAATCATCAGGCTGTCGCTGTCGCCG | TTTCGTCCGCGAGTGACCAGCATGAATTCTCGCCAAACATGAATTCTCGCCAAATTGGGAATCATCAGGCTGTCGCTGTCGCCGAACATGAATTCTCGCCAAACATGAATTCTCGCCACAACGTCCATGTCGGGATGC |
| PA14_53470 | ackA | TCGATGCGCGTGGCATGGGTGAACAGCTCG | TTTCGTCCGCGAGTGACCAGCATGAATTCTCGCCAAACATGAATTCTCGCCAAATCGATGCGCGTGGCATGGGTGAACAGCTCGAACATGAATTCTCGCCAAACATGAATTCTCGCCACAACGTCCATGTCGGGATGC |
| PA14_53470 | ackA | TCTTCGTTGGTCGGGATCACCAGTACCCGC | TTTCGTCCGCGAGTGACCAGCATGAATTCTCGCCAAACATGAATTCTCGCCAAATCTTCGTTGGTCGGGATCACCAGTACCCGCAACATGAATTCTCGCCAAACATGAATTCTCGCCACAACGTCCATGTCGGGATGC |
| PA14_53470 | ackA | GCTTTCGTTGTTCAGCATCGAGTCGATGCG | TTTCGTCCGCGAGTGACCAGCATGAATTCTCGCCAAACATGAATTCTCGCCAAAGCTTTCGTTGTTCAGCATCGAGTCGATGCGAACATGAATTCTCGCCAAACATGAATTCTCGCCACAACGTCCATGTCGGGATGC |
| PA14_53470 | ackA | TTCTCGCCGATGCCACCGGTGAAGATCACC | TTTCGTCCGCGAGTGACCAGCATGAATTCTCGCCAAACATGAATTCTCGCCAAATTCTCGCCGATGCCACCGGTGAAGATCACCAACATGAATTCTCGCCAAACATGAATTCTCGCCACAACGTCCATGTCGGGATGC |
| PA14_53470 | ackA | TTGGCCAGGCGGTAGCAGAACACCTCGATC | TTTCGTCCGCGAGTGACCAGCATGAATTCTCGCCAAACATGAATTCTCGCCAAATTGGCCAGGCGGTAGCAGAACACCTCGATCAACATGAATTCTCGCCAAACATGAATTCTCGCCACAACGTCCATGTCGGGATGC |
| PA14_53470 | ackA | ACCAGGGCGAACTTGATCGACGAACTGCCG | TTTCGTCCGCGAGTGACCAGCATGAATTCTCGCCAAACATGAATTCTCGCCAAAACCAGGGCGAACTTGATCGACGAACTGCCGAACATGAATTCTCGCCAAACATGAATTCTCGCCACAACGTCCATGTCGGGATGC |
| PA14_53470 | ackA | GTTGATCACCAGTATGTTGCGTGAGGGCAT | TTTCGTCCGCGAGTGACCAGCATGAATTCTCGCCAAACATGAATTCTCGCCAAAGTTGATCACCAGTATGTTGCGTGAGGGCATAACATGAATTCTCGCCAAACATGAATTCTCGCCACAACGTCCATGTCGGGATGC |
| PA14_53470 | ackA | ATTGCCAAGGTGGGCGCTGAGCCAACTGCT | TTTCGTCCGCGAGTGACCAGCATGAATTCTCGCCAAACATGAATTCTCGCCAAAATTGCCAAGGTGGGCGCTGAGCCAACTGCTAACATGAATTCTCGCCAAACATGAATTCTCGCCACAACGTCCATGTCGGGATGC |
| PA14_53470 | ackA | ACACGGCGACGTGGGGCAGCTTTGGAAACA | TTTCGTCCGCGAGTGACCAGCATGAATTCTCGCCAAACATGAATTCTCGCCAAAACACGGCGACGTGGGGCAGCTTTGGAAACAAACATGAATTCTCGCCAAACATGAATTCTCGCCACAACGTCCATGTCGGGATGC |
| PA14_53470 | ackA | TGGTGTCGAGGCTCTGGCCGTTGACGATGG | TTTCGTCCGCGAGTGACCAGCATGAATTCTCGCCAAACATGAATTCTCGCCAAATGGTGTCGAGGCTCTGGCCGTTGACGATGGAACATGAATTCTCGCCAAACATGAATTCTCGCCACAACGTCCATGTCGGGATGC |

| | | | |
|---|---|---|---|
| PA14_53470 | ackA | TGCCCATTACCAGGCCTTCCAGCGGGGTCA | TTTCGTCCGCGAGTGACCAGCATGAATTCTCGCCAAACATGAATTCTCGCCAAATGCCCATTACCAGGCCTTCCAGCGGGGTCAAACATGAATTCTCGCCAAACATGAATTCTCGCCACAACGTCCATGTCGGGATGC |
| PA14_53470 | ackA | TCATCGCTGCCTCGATGCCTTGCAGGTTGG | TTTCGTCCGCGAGTGACCAGCATGAATTCTCGCCAAACATGAATTCTCGCCAAATCATCGCTGCCTCGATGCCTTGCAGGTTGGAACATGAATTCTCGCCAAACATGAATTCTCGCCACAACGTCCATGTCGGGATGC |
| PA14_01720 | ahpF | GGATTTGTCGCTGTCATCGAGGGACGCGAC | TTTCGTCCGCGAGTGACCAGTAGAGGTATGTCGGAAATAGAGGTATGTCGGAAAGGATTTGTCGCTGTCATCGAGGGACGCGACAATAGAGGTATGTCGGAAATAGAGGTATGTCGGACAACGTCCATGTCGGGATGC |
| PA14_01720 | ahpF | TGTATTCCTGCTCGCCGGGCACGCCCATTT | TTTCGTCCGCGAGTGACCAGTAGAGGTATGTCGGAAATAGAGGTATGTCGGAAATGTATTCCTGCTCGCCGGGCACGCCCATTTAATAGAGGTATGTCGGAAATAGAGGTATGTCGGACAACGTCCATGTCGGGATGC |
| PA14_01720 | ahpF | ACCTCGTAGTGACGCACGTGCTCTTCCAGC | TTTCGTCCGCGAGTGACCAGTAGAGGTATGTCGGAAATAGAGGTATGTCGGAAAACCTCGTAGTGACGCACGTGCTCTTCCAGCAATAGAGGTATGTCGGAAATAGAGGTATGTCGGACAACGTCCATGTCGGGATGC |
| PA14_01720 | ahpF | TCTTGGCGAGGATCTCTTCCACGCCCATGC | TTTCGTCCGCGAGTGACCAGTAGAGGTATGTCGGAAATAGAGGTATGTCGGAAATCTTGGCGAGGATCTCTTCCACGCCCATGCAATAGAGGTATGTCGGAAATAGAGGTATGTCGGACAACGTCCATGTCGGGATGC |
| PA14_01720 | ahpF | TTCGCTGGTCAGGGCACTGGTGATCACCTC | TTTCGTCCGCGAGTGACCAGTAGAGGTATGTCGGAAATAGAGGTATGTCGGAAATTCGCTGGTCAGGGCACTGGTGATCACCTCAATAGAGGTATGTCGGAAATAGAGGTATGTCGGACAACGTCCATGTCGGGATGC |
| PA14_01720 | ahpF | AATTCGTGGCCCATGGGGATACCGGCGAAG | TTTCGTCCGCGAGTGACCAGTAGAGGTATGTCGGAAATAGAGGTATGTCGGAAAAATTCGTGGCCCATGGGGATACCGGCGAAGAATAGAGGTATGTCGGAAATAGAGGTATGTCGGACAACGTCCATGTCGGGATGC |
| PA14_01720 | ahpF | CGTTGAGGTAGATGCTCGGCACCGACATGA | TTTCGTCCGCGAGTGACCAGTAGAGGTATGTCGGAAATAGAGGTATGTCGGAAACGTTGAGGTAGATGCTCGGCACCGACATGAAATAGAGGTATGTCGGAAATAGAGGTATGTCGGACAACGTCCATGTCGGGATGC |
| PA14_01720 | ahpF | CGGTCTTCAGGGTGATCTTGTCGGTCAGGC | TTTCGTCCGCGAGTGACCAGTAGAGGTATGTCGGAAATAGAGGTATGTCGGAAACGGTCTTCAGGGTGATCTTGTCGGTCAGGCAATAGAGGTATGTCGGAAATAGAGGTATGTCGGACAACGTCCATGTCGGGATGC |
| PA14_01720 | ahpF | TCCTTGTACACCAGGCCGGTGACCTTCTGG | TTTCGTCCGCGAGTGACCAGTAGAGGTATGTCGGAAATAGAGGTATGTCGGAAATCCTTGTACACCAGGCCGGTGACCTTCTGGAATAGAGGTATGTCGGAAATAGAGGTATGTCGGACAACGTCCATGTCGGGATGC |
| PA14_01720 | ahpF | TCTCGAACGGCTGGCTGACCTTTTCCAGGT | TTTCGTCCGCGAGTGACCAGTAGAGGTATGTCGGAAATAGAGGTATGTCGGAAATCTCGAACGGCTGGCTGACCTTTTCCAGGTAATAGAGGTATGTCGGAAATAGAGGTATGTCGGACAACGTCCATGTCGGGATGC |
| PA14_01720 | ahpF | ACGATCTGCTTGTACGGCACCGTGGTCACG | TTTCGTCCGCGAGTGACCAGTAGAGGTATGTCGGAAATAGAGGTATGTCGGAAAACGATCTGCTTGTACGGCACCGTGGTCACGAATAGAGGTATGTCGGAAATAGAGGTATGTCGGACAACGTCCATGTCGGGATGC |
| PA14_01720 | ahpF | ACGGATGTTCGGGTTGAGCACGGCCATCAG | TTTCGTCCGCGAGTGACCAGTAGAGGTATGTCGGAAATAGAGGTATGTCGGAAAACGGATGTTCGGGTTGAGCACGGCCATCAGAATAGAGGTATGTCGGAAATAGAGGTATGTCGGACAACGTCCATGTCGGGATGC |
| PA14_01720 | ahpF | GCCTTCAACTGGGTTTTAAGATTGGCGTCC | TTTCGTCCGCGAGTGACCAGTAGAGGTATGTCGGAAATAGAGGTATGTCGGAAAGCCTTCAACTGGGTTTTAAGATTGGCGTCCAATAGAGGTATGTCGGAAATAGAGGTATGTCGGACAACGTCCATGTCGGGATGC |
| PA14_01720 | ahpF | TTCGGCAGGCTGTAGAGCTTGCGTTGCAGC | TTTCGTCCGCGAGTGACCAGTAGAGGTATGTCGGAAATAGAGGTATGTCGGAAATTCGGCAGGCTGTAGAGCTTGCGTTGCAGCAATAGAGGTATGTCGGAAATAGAGGTATGTCGGACAACGTCCATGTCGGGATGC |
| PA14_01720 | ahpF | ATTCCAGCAGGGTCACATGGGCGACGATGC | TTTCGTCCGCGAGTGACCAGTAGAGGTATGTCGGAAATAGAGGTATGTCGGAAAATTCCAGCAGGGTCACATGGGCGACGATGCAATAGAGGTATGTCGGAAATAGAGGTATGTCGGACAACGTCCATGTCGGGATGC |
| PA14_01720 | ahpF | TTCAGCGAGAACGACGGTTTGCGCGCATCG | TTTCGTCCGCGAGTGACCAGTAGAGGTATGTCGGAAATAGAGGTATGTCGGAAATTCAGCGAGAACGACGGTTTGCGCGCATCGAATAGAGGTATGTCGGAAATAGAGGTATGTCGGACAACGTCCATGTCGGGATGC |
| PA14_01720 | ahpF | GCTCGATCGACTTGAACTCCTCGGAGTTGC | TTTCGTCCGCGAGTGACCAGTAGAGGTATGTCGGAAATAGAGGTATGTCGGAAAGCTCGATCGACTTGAACTCCTCGGAGTTGCAATAGAGGTATGTCGGAAATAGAGGTATGTCGGACAACGTCCATGTCGGGATGC |
| PA14_01720 | ahpF | GTTCTGGCAGGACAGCGAGAAGTAGGTTTC | TTTCGTCCGCGAGTGACCAGTAGAGGTATGTCGGAAATAGAGGTATGTCGGAAAGTTCTGGCAGGACAGCGAGAAGTAGGTTTCAATAGAGGTATGTCGGAAATAGAGGTATGTCGGACAACGTCCATGTCGGGATGC |
| PA14_01720 | ahpF | CGAGATGAAGTTCTCGATGGCCATGGTGTC | TTTCGTCCGCGAGTGACCAGTAGAGGTATGTCGGAAATAGAGGTATGTCGGAAACGAGATGAAGTTCTCGATGGCCATGGTGTCAATAGAGGTATGTCGGAAATAGAGGTATGTCGGACAACGTCCATGTCGGGATGC |
| PA14_01720 | ahpF | ACTCGAAGGTGCCCTCGATGCCCTTGACCT | TTTCGTCCGCGAGTGACCAGTAGAGGTATGTCGGAAATAGAGGTATGTCGGAAAACTCGAAGGTGCCCTCGATGCCCTTGACCTAATAGAGGTATGTCGGAAATAGAGGTATGTCGGACAACGTCCATGTCGGGATGC |
| PA14_54430 | algU | CTTCTCGCAACAAAGGCTGCAGAGCTTTGT | TTTCGTCCGCGAGTGACCAGGCAGGGTGAACCTACAAGCAGGGTGAACCTACAACTTCTCGCAACAAAGGCTGCAGAGCTTTGTAAGCAGGGTGAACCTACAAGCAGGGTGAACCTACCAACGTCCATGTCGGGATGC |
| PA14_54430 | algU | ATTGCTTCACGAGCGCGGAAGATCCGCGAC | TTTCGTCCGCGAGTGACCAGGCAGGGTGAACCTACAAGCAGGGTGAACCTACAAATTGCTTCACGAGCGCGGAAGATCCGCGACAAGCAGGGTGAACCTACAAGCAGGGTGAACCTACCAACGTCCATGTCGGGATGC |
| PA14_54430 | algU | GCACGATCAATCCCAGTATCTTGTGCTGGT | TTTCGTCCGCGAGTGACCAGGCAGGGTGAACCTACAAGCAGGGTGAACCTACAAGCACGATCAATCCCAGTATCTTGTGCTGGTAAGCAGGGTGAACCTACAAGCAGGGTGAACCTACCAACGTCCATGTCGGGATGC |
| PA14_54430 | algU | TCGATCTCATCCCGCAACATCGCCCGTTCC | TTTCGTCCGCGAGTGACCAGGCAGGGTGAACCTACAAGCAGGGTGAACCTACAATCGATCTCATCCCGCAACATCGCCCGTTCCAAGCAGGGTGAACCTACAAGCAGGGTGAACCTACCAACGTCCATGTCGGGATGC |
| PA14_54430 | algU | ATCCTCGGGCAACTGCTGGATGGTCTGGTG | TTTCGTCCGCGAGTGACCAGGCAGGGTGAACCTACAAGCAGGGTGAACCTACAAATCCTCGGGCAACTGCTGGATGGTCTGGTGAAGCAGGGTGAACCTACAAGCAGGGTGAACCTACCAACGTCCATGTCGGGATGC |
| PA14_54430 | algU | ATTCGATGTCCTTCAGGGCGTGGTCGCCCT | TTTCGTCCGCGAGTGACCAGGCAGGGTGAACCTACAAGCAGGGTGAACCTACAAATTCGATGTCCTTCAGGGCGTGGTCGCCCTAAGCAGGGTGAACCTACAAGCAGGGTGAACCTACCAACGTCCATGTCGGGATGC |
| PA14_54430 | algU | TGAAGGCTTCCTGCGCTACGTCCTGGGCTT | TTTCGTCCGCGAGTGACCAGGCAGGGTGAACCTACAAGCAGGGTGAACCTACAATGAAGGCTTCCTGCGCTACGTCCTGGGCTTAAGCAGGGTGAACCTACAAGCAGGGTGAACCTACCAACGTCCATGTCGGGATGC |
| PA14_54430 | algU | ATCGAAAGCCCGCTTGTCTCCGCGCTGTAC | TTTCGTCCGCGAGTGACCAGGCAGGGTGAACCTACAAGCAGGGTGAACCTACAAATCGAAAGCCCGCTTGTCTCCGCGCTGTACAAGCAGGGTGAACCTACAAGCAGGGTGAACCTACCAACGTCCATGTCGGGATGC |
| PA14_54430 | algU | CCACCGGACACTGCATCACGGTGGCGATAT | TTTCGTCCGCGAGTGACCAGGCAGGGTGAACCTACAAGCAGGGTGAACCTACAACCACCGGACACTGCATCACGGTGGCGATATAAGCAGGGTGAACCTACAAGCAGGGTGAACCTACCAACGTCCATGTCGGGATGC |
| PA14_54430 | algU | CAGTTGCTGATCCTGTTCCTGGGTTAGCAT | TTTCGTCCGCGAGTGACCAGGCAGGGTGAACCTACAAGCAGGGTGAACCTACAACAGTTGCTGATCCTGTTCCTGGGTTAGCATAAGCAGGGTGAACCTACAAGCAGGGTGAACCTACCAACGTCCATGTCGGGATGC |
| PA14_54430 | algU | TCGTAACTCAAACCTTCGAACTCGCGCAGG | TTTCGTCCGCGAGTGACCAGGCAGGGTGAACCTACAAGCAGGGTGAACCTACAATCGTAACTCAAACCTTCGAACTCGCGCAGGAAGCAGGGTGAACCTACAAGCAGGGTGAACCTACCAACGTCCATGTCGGGATGC |
| PA14_54430 | algU | AAGCACTATCGCCGCGGAAATTGCCGAGCG | TTTCGTCCGCGAGTGACCAGGCAGGGTGAACCTACAAGCAGGGTGAACCTACAAAAGCACTATCGCCGCGGAAATTGCCGAGCGAAGCAGGGTGAACCTACAAGCAGGGTGAACCTACCAACGTCCATGTCGGGATGC |
| PA14_54430 | algU | GTTGATGGCGATCCGATACAGCCAGGTATA | TTTCGTCCGCGAGTGACCAGGCAGGGTGAACCTACAAGCAGGGTGAACCTACAAGTTGATGGCGATCCGATACAGCCAGGTATAAAGCAGGGTGAACCTACAAGCAGGGTGAACCTACCAACGTCCATGTCGGGATGC |
| PA14_54430 | algU | ACTCCGCATCCTCTGCGGTCACATCGCTGT | TTTCGTCCGCGAGTGACCAGGCAGGGTGAACCTACAAGCAGGGTGAACCTACAAACTCCGCATCCTCTGCGGTCACATCGCTGTAAGCAGGGTGAACCTACAAGCAGGGTGAACCTACCAACGTCCATGTCGGGATGC |
| PA14_10790 | ampC | CGGCATTGGGATAGTTGCGGTTGGCCAGGA | TTTCGTCCGCGAGTGACCAGAATTGCATCTTCGCGAAAATTGCATCTTCGCGAACGGCATTGGGATAGTTGCGGTTGGCCAGGAAAAATTGCATCTTCGCGAAAATTGCATCTTCGCGCAACGTCCATGTCGGGATGC |
| PA14_10790 | ampC | CCGGATGCAGGTTGGCATCGACGAAGCGCA | TTTCGTCCGCGAGTGACCAGAATTGCATCTTCGCGAAAATTGCATCTTCGCGAACCGGATGCAGGTTGGCATCGACGAAGCGCAAAAATTGCATCTTCGCGAAAATTGCATCTTCGCGCAACGTCCATGTCGGGATGC |
| PA14_10790 | ampC | AGCCGTTGGTGGAGCCGGTCTTGTTCAGCA | TTTCGTCCGCGAGTGACCAGAATTGCATCTTCGCGAAAATTGCATCTTCGCGAAAGCCGTTGGTGGAGCCGGTCTTGTTCAGCAAAAATTGCATCTTCGCGAAAATTGCATCTTCGCGCAACGTCCATGTCGGGATGC |
| PA14_10790 | ampC | TTCAGGGAGATCGGCCAGTCGTAGGCTTCC | TTTCGTCCGCGAGTGACCAGAATTGCATCTTCGCGAAAATTGCATCTTCGCGAATTCAGGGAGATCGGCCAGTCGTAGGCTTCCAAAATTGCATCTTCGCGAAAATTGCATCTTCGCGCAACGTCCATGTCGGGATGC |
| PA14_10790 | ampC | TCCTTTCAGGCTGATGGCTACGGTCAGGCC | TTTCGTCCGCGAGTGACCAGAATTGCATCTTCGCGAAAATTGCATCTTCGCGAATCCTTTCAGGCTGATGGCTACGGTCAGGCCAAAATTGCATCTTCGCGAAAATTGCATCTTCGCGCAACGTCCATGTCGGGATGC |
| PA14_10790 | ampC | ATGCTCGGGTTGGAATAGAGGCGCTGGCTG | TTTCGTCCGCGAGTGACCAGAATTGCATCTTCGCGAAAATTGCATCTTCGCGAAATGCTCGGGTTGGAATAGAGGCGCTGGCTGAAAATTGCATCTTCGCGAAAATTGCATCTTCGCGCAACGTCCATGTCGGGATGC |
| PA14_10790 | ampC | ATTGGCCTTCATCACCGGTTGTACGGCGGC | TTTCGTCCGCGAGTGACCAGAATTGCATCTTCGCGAAAATTGCATCTTCGCGAAATTGGCCTTCATCACCGGTTGTACGGCGGCAAAATTGCATCTTCGCGAAAATTGCATCTTCGCGCAACGTCCATGTCGGGATGC |
| PA14_10790 | ampC | TGGTCTTCACCCCGTAGCCTTCGGCATCCA | TTTCGTCCGCGAGTGACCAGAATTGCATCTTCGCGAAAATTGCATCTTCGCGAATGGTCTTCACCCCGTAGCCTTCGGCATCCAAAAATTGCATCTTCGCGAAAATTGCATCTTCGCGCAACGTCCATGTCGGGATGC |
| PA14_10790 | ampC | TGCTCACCGAGCCGATCTCGAACAGGGTCT | TTTCGTCCGCGAGTGACCAGAATTGCATCTTCGCGAAAATTGCATCTTCGCGAATGCTCACCGAGCCGATCTCGAACAGGGTCTAAAATTGCATCTTCGCGAAAATTGCATCTTCGCGCAACGTCCATGTCGGGATGC |
| PA14_10790 | ampC | GCACAGGCAGGGGAATCTGGTATCGCGCAT | TTTCGTCCGCGAGTGACCAGAATTGCATCTTCGCGAAAATTGCATCTTCGCGAAGCACAGGCAGGGGAATCTGGTATCGCGCATAAAATTGCATCTTCGCGAAAATTGCATCTTCGCGCAACGTCCATGTCGGGATGC |
| PA14_10790 | ampC | TTGCTGCTCCATGAGCCGTTCGAACGGCTG | TTTCGTCCGCGAGTGACCAGAATTGCATCTTCGCGAAAATTGCATCTTCGCGAATTGCTGCTCCATGAGCCGTTCGAACGGCTGAAAATTGCATCTTCGCGAAAATTGCATCTTCGCGCAACGTCCATGTCGGGATGC |
| PA14_10790 | ampC | CTTGTAGTAACCGCGATGGGTGGCATCGAG | TTTCGTCCGCGAGTGACCAGAATTGCATCTTCGCGAAAATTGCATCTTCGCGAACTTGTAGTAACCGCGATGGGTGGCATCGAGAAAATTGCATCTTCGCGAAAATTGCATCTTCGCGCAACGTCCATGTCGGGATGC |
| PA14_10790 | ampC | CCGTCCTCTTTCGAGGCCAGCCCATAGCTG | TTTCGTCCGCGAGTGACCAGAATTGCATCTTCGCGAAAATTGCATCTTCGCGAACCGTCCTCTTTCGAGGCCAGCCCATAGCTGAAAATTGCATCTTCGCGAAAATTGCATCTTCGCGCAACGTCCATGTCGGGATGC |
| PA14_10790 | ampC | AGTAGTCGCGGATCTGTGCCCGGTCCTTCT | TTTCGTCCGCGAGTGACCAGAATTGCATCTTCGCGAAAATTGCATCTTCGCGAAAGTAGTCGCGGATCTGTGCCCGGTCCTTCTAAAATTGCATCTTCGCGAAAATTGCATCTTCGCGCAACGTCCATGTCGGGATGC |
| PA14_10790 | ampC | GTGGCGAACAGCAGTGTGGAAGCGGCGATG | TTTCGTCCGCGAGTGACCAGAATTGCATCTTCGCGAAAATTGCATCTTCGCGAAGTGGCGAACAGCAGTGTGGAAGCGGCGATGAAAATTGCATCTTCGCGAAAATTGCATCTTCGCGCAACGTCCATGTCGGGATGC |
| PA14_18580 | algD | CTTGATCGCGGTGCTCTCGCGGAGGAATTC | TTTCGTCCGCGAGTGACCAGAGCTCGGCCTGTGATAAAGCTCGGCCTGTGATAACTTGATCGCGGTGCTCTCGCGGAGGAATTCAAAGCTCGGCCTGTGATAAAGCTCGGCCTGTGATCAACGTCCATGTCGGGATGC |
| PA14_18580 | algD | GGTAGATTTCCTCGAGAAGGTCGCCGGTCT | TTTCGTCCGCGAGTGACCAGAGCTCGGCCTGTGATAAAGCTCGGCCTGTGATAAGGTAGATTTCCTCGAGAAGGTCGCCGGTCTAAAGCTCGGCCTGTGATAAAGCTCGGCCTGTGATCAACGTCCATGTCGGGATGC |
| PA14_18580 | algD | GACGCATGTAGTAGCGCGACAGGTTGAGCT | TTTCGTCCGCGAGTGACCAGAGCTCGGCCTGTGATAAAGCTCGGCCTGTGATAAGACGCATGTAGTAGCGCGACAGGTTGAGCTAAAGCTCGGCCTGTGATAAAGCTCGGCCTGTGATCAACGTCCATGTCGGGATGC |
| PA14_18580 | algD | TGCCGATCTCGTTGGCGAAGGTGACCTTGG | TTTCGTCCGCGAGTGACCAGAGCTCGGCCTGTGATAAAGCTCGGCCTGTGATAATGCCGATCTCGTTGGCGAAGGTGACCTTGGAAAGCTCGGCCTGTGATAAAGCTCGGCCTGTGATCAACGTCCATGTCGGGATGC |
| PA14_18580 | algD | GGTCGATCTTGGTGCTGGAGACATCCACAC | TTTCGTCCGCGAGTGACCAGAGCTCGGCCTGTGATAAAGCTCGGCCTGTGATAAGGTCGATCTTGGTGCTGGAGACATCCACACAAAGCTCGGCCTGTGATAAAGCTCGGCCTGTGATCAACGTCCATGTCGGGATGC |
| PA14_18580 | algD | ATGACTTCATGACCGCGTGCCGACAGGCAG | TTTCGTCCGCGAGTGACCAGAGCTCGGCCTGTGATAAAGCTCGGCCTGTGATAAATGACTTCATGACCGCGTGCCGACAGGCAGAAAGCTCGGCCTGTGATAAAGCTCGGCCTGTGATCAACGTCCATGTCGGGATGC |
| PA14_18580 | algD | AGGCCTTCTGCACCTGGTTGGAGTTGCTGC | TTTCGTCCGCGAGTGACCAGAGCTCGGCCTGTGATAAAGCTCGGCCTGTGATAAAGGCCTTCTGCACCTGGTTGGAGTTGCTGCAAAGCTCGGCCTGTGATAAAGCTCGGCCTGTGATCAACGTCCATGTCGGGATGC |
| PA14_18580 | algD | CACCGACATAGCCCAAACCAAAGATGCTGA | TTTCGTCCGCGAGTGACCAGAGCTCGGCCTGTGATAAAGCTCGGCCTGTGATAACACCGACATAGCCCAAACCAAAGATGCTGAAAAGCTCGGCCTGTGATAAAGCTCGGCCTGTGATCAACGTCCATGTCGGGATGC |
| PA14_18580 | algD | ATAGGTGAGGGCGCGTACATCCTTGGGCAG | TTTCGTCCGCGAGTGACCAGAGCTCGGCCTGTGATAAAGCTCGGCCTGTGATAAATAGGTGAGGGCGCGTACATCCTTGGGCAGAAAGCTCGGCCTGTGATAAAGCTCGGCCTGTGATCAACGTCCATGTCGGGATGC |
| PA14_18580 | algD | TGGTCCTGGCAGATCACGTCCATCACCTCG | TTTCGTCCGCGAGTGACCAGAGCTCGGCCTGTGATAAAGCTCGGCCTGTGATAATGGTCCTGGCAGATCACGTCCATCACCTCGAAAGCTCGGCCTGTGATAAAGCTCGGCCTGTGATCAACGTCCATGTCGGGATGC |
| PA14_18580 | algD | TCTGCCGGCCTTGCTGCAACAACGCTTCCA | TTTCGTCCGCGAGTGACCAGAGCTCGGCCTGTGATAAAGCTCGGCCTGTGATAATCTGCCGGCCTTGCTGCAACAACGCTTCCAAAAGCTCGGCCTGTGATAAAGCTCGGCCTGTGATCAACGTCCATGTCGGGATGC |
| PA14_18580 | algD | TTCGACGATGGGCGACTTGCCCTGGTTGAT | TTTCGTCCGCGAGTGACCAGAGCTCGGCCTGTGATAAAGCTCGGCCTGTGATAATTCGACGATGGGCGACTTGCCCTGGTTGATAAAGCTCGGCCTGTGATAAAGCTCGGCCTGTGATCAACGTCCATGTCGGGATGC |
| PA14_18580 | algD | AGACGGTCTCGATGTAGCCCAGGTCCAGGT | TTTCGTCCGCGAGTGACCAGAGCTCGGCCTGTGATAAAGCTCGGCCTGTGATAAAGACGGTCTCGATGTAGCCCAGGTCCAGGTAAAGCTCGGCCTGTGATAAAGCTCGGCCTGTGATCAACGTCCATGTCGGGATGC |
| PA14_18580 | algD | ACCAGCAGTGAGGAGACGTGCGGGATCTTC | TTTCGTCCGCGAGTGACCAGAGCTCGGCCTGTGATAAAGCTCGGCCTGTGATAAACCAGCAGTGAGGAGACGTGCGGGATCTTCAAAGCTCGGCCTGTGATAAAGCTCGGCCTGTGATCAACGTCCATGTCGGGATGC |
| PA14_18580 | algD | TTGTCCAGTTCGCCGATCACGGTCATCGGC | TTTCGTCCGCGAGTGACCAGAGCTCGGCCTGTGATAAAGCTCGGCCTGTGATAATTGTCCAGTTCGCCGATCACGGTCATCGGCAAAGCTCGGCCTGTGATAAAGCTCGGCCTGTGATCAACGTCCATGTCGGGATGC |
| PA14_18580 | algD | TCGATGTATTCCTTGTTGGCCCCGTGGACA | TTTCGTCCGCGAGTGACCAGAGCTCGGCCTGTGATAAAGCTCGGCCTGTGATAATCGATGTATTCCTTGTTGGCCCCGTGGACAAAAGCTCGGCCTGTGATAAAGCTCGGCCTGTGATCAACGTCCATGTCGGGATGC |
| PA14_18580 | algD | TCTTGAAATCGGTGGTGCCCGACAGCCGTC | TTTCGTCCGCGAGTGACCAGAGCTCGGCCTGTGATAAAGCTCGGCCTGTGATAATCTTGAAATCGGTGGTGCCCGACAGCCGTCAAAGCTCGGCCTGTGATAAAGCTCGGCCTGTGATCAACGTCCATGTCGGGATGC |
| PA14_18580 | algD | TCACCACGTTGTTGACGGTGCCCGGCAGTA | TTTCGTCCGCGAGTGACCAGAGCTCGGCCTGTGATAAAGCTCGGCCTGTGATAATCACCACGTTGTTGACGGTGCCCGGCAGTAAAAGCTCGGCCTGTGATAAAGCTCGGCCTGTGATCAACGTCCATGTCGGGATGC |
| PA14_18580 | algD | GATGAACGATACGTCGGAGTCCAGCACGGC | TTTCGTCCGCGAGTGACCAGAGCTCGGCCTGTGATAAAGCTCGGCCTGTGATAAGATGAACGATACGTCGGAGTCCAGCACGGCAAAGCTCGGCCTGTGATAAAGCTCGGCCTGTGATCAACGTCCATGTCGGGATGC |
| PA14_18580 | algD | TGATCATCTCGGCGACCTCGACGGTCTTGC | TTTCGTCCGCGAGTGACCAGAGCTCGGCCTGTGATAAAGCTCGGCCTGTGATAATGATCATCTCGGCGACCTCGACGGTCTTGCAAAGCTCGGCCTGTGATAAAGCTCGGCCTGTGATCAACGTCCATGTCGGGATGC |
| PA14_38410 | amrB | GTGGTGTTGATCTCGTCCATGCTCACGCCG | TTTCGTCCGCGAGTGACCAGGGGGTGGAATTTCGTAAAGGGTGGAATTTCGTAAAGTGGTGTTGATCTCGTCCATGCTCACGCCGAAGGGTGGAATTTCGTAAAGGGTGGAATTTCGTACAACGTCCATGTCGGGATGC |
| PA14_38410 | amrB | TGAGGGTGACGATCAGCTGGATGCTGTCCG | TTTCGTCCGCGAGTGACCAGGGGGTGGAATTTCGTAAAGGGTGGAATTTCGTAAATGAGGGTGACGATCAGCTGGATGCTGTCCGAAGGGTGGAATTTCGTAAAGGGTGGAATTTCGTACAACGTCCATGTCGGGATGC |
| PA14_38410 | amrB | AAGCCGTACTCGCTTTGGCCGAACTCGACG | TTTCGTCCGCGAGTGACCAGGGGGTGGAATTTCGTAAAGGGTGGAATTTCGTAAAAAGCCGTACTCGCTTTGGCCGAACTCGACGAAGGGTGGAATTTCGTAAAGGGTGGAATTTCGTACAACGTCCATGTCGGGATGC |
| PA14_38410 | amrB | TTTGGCGCATCGCCTTGACCGTGGTGTCGT | TTTCGTCCGCGAGTGACCAGGGGGTGGAATTTCGTAAAGGGTGGAATTTCGTAAATTTGGCGCATCGCCTTGACCGTGGTGTCGTAAGGGTGGAATTTCGTAAAGGGTGGAATTTCGTACAACGTCCATGTCGGGATGC |
| PA14_38410 | amrB | TGTTGTAGCTCACGCCCTCCGGGAAGTAGC | TTTCGTCCGCGAGTGACCAGGGGGTGGAATTTCGTAAAGGGTGGAATTTCGTAAATGTTGTAGCTCACGCCCTCCGGGAAGTAGCAAGGGTGGAATTTCGTAAAGGGTGGAATTTCGTACAACGTCCATGTCGGGATGC |
| PA14_38410 | amrB | AGATCTCGACGAACGCCGAGGTGTCGTAGG | TTTCGTCCGCGAGTGACCAGGGGGTGGAATTTCGTAAAGGGTGGAATTTCGTAAAAGATCTCGACGAACGCCGAGGTGTCGTAGGAAGGGTGGAATTTCGTAAAGGGTGGAATTTCGTACAACGTCCATGTCGGGATGC |
| PA14_38410 | amrB | GACCGGCCGGTCAATGAAGAAACGAGCCAT | TTTCGTCCGCGAGTGACCAGGGGGTGGAATTTCGTAAAGGGTGGAATTTCGTAAAGACCGGCCGGTCAATGAAGAAACGAGCCATAAGGGTGGAATTTCGTAAAGGGTGGAATTTCGTACAACGTCCATGTCGGGATGC |
| PA14_38410 | amrB | TAGATGTTGCCCACCGCGCCGCTGAAGAAC | TTTCGTCCGCGAGTGACCAGGGGGTGGAATTTCGTAAAGGGTGGAATTTCGTAAATAGATGTTGCCCACCGCGCCGCTGAAGAACAAGGGTGGAATTTCGTAAAGGGTGGAATTTCGTACAACGTCCATGTCGGGATGC |
| PA14_38410 | amrB | CTCGAGCAGGGTGCTGACCACCTTCCTGAT | TTTCGTCCGCGAGTGACCAGGGGGTGGAATTTCGTAAAGGGTGGAATTTCGTAAACTCGAGCAGGGTGCTGACCACCTTCCTGATAAGGGTGGAATTTCGTAAAGGGTGGAATTTCGTACAACGTCCATGTCGGGATGC |
| PA14_38410 | amrB | TTTCTCCACGTAGATACCGTCGCGCCGCAC | TTTCGTCCGCGAGTGACCAGGGGGTGGAATTTCGTAAAGGGTGGAATTTCGTAAATTTCTCCACGTAGATACCGTCGCGCCGCACAAGGGTGGAATTTCGTAAAGGGTGGAATTTCGTACAACGTCCATGTCGGGATGC |
| PA14_38410 | amrB | TGAGGTTCATCGAGGTCAGCTTGGCCGGGT | TTTCGTCCGCGAGTGACCAGGGGGTGGAATTTCGTAAAGGGTGGAATTTCGTAAATGAGGTTCATCGAGGTCAGCTTGGCCGGGTAAGGGTGGAATTTCGTAAAGGGTGGAATTTCGTACAACGTCCATGTCGGGATGC |
| PA14_38410 | amrB | TCATCGCATACACCGTACGGTTGGGCAGGC | TTTCGTCCGCGAGTGACCAGGGGGTGGAATTTCGTAAAGGGTGGAATTTCGTAAATCATCGCATACACCGTACGGTTGGGCAGGCAAGGGTGGAATTTCGTAAAGGGTGGAATTTCGTACAACGTCCATGTCGGGATGC |
| PA14_38410 | amrB | AATACCGAGACCAGCACCACGGTGATGCCG | TTTCGTCCGCGAGTGACCAGGGGGTGGAATTTCGTAAAGGGTGGAATTTCGTAAAAATACCGAGACCAGCACCACGGTGATGCCGAAGGGTGGAATTTCGTAAAGGGTGGAATTTCGTACAACGTCCATGTCGGGATGC |
| PA14_38410 | amrB | AGGCTCATGCCTTCCTGGTAATGGTCCTTG | TTTCGTCCGCGAGTGACCAGGGGGTGGAATTTCGTAAAGGGTGGAATTTCGTAAAAGGCTCATGCCTTCCTGGTAATGGTCCTTGAAGGGTGGAATTTCGTAAAGGGTGGAATTTCGTACAACGTCCATGTCGGGATGC |
| PA14_38410 | amrB | TCTACATCGGCGACGTTGGCCATGGTCTCC | TTTCGTCCGCGAGTGACCAGGGGGTGGAATTTCGTAAAGGGTGGAATTTCGTAAATCTACATCGGCGACGTTGGCCATGGTCTCCAAGGGTGGAATTTCGTAAAGGGTGGAATTTCGTACAACGTCCATGTCGGGATGC |
| PA14_44490 | anr | CACGGTTTCTACGGCCAGGCCGAGATAGTT | TTTCGTCCGCGAGTGACCAGGGCTTACTGAGTGAAAAGGCTTACTGAGTGAAAACACGGTTTCTACGGCCAGGCCGAGATAGTTAAGGCTTACTGAGTGAAAAGGCTTACTGAGTGAACAACGTCCATGTCGGGATGC |
| PA14_44490 | anr | TCATCAGGCGCATCAGTTGGCGGCGCAGTT | TTTCGTCCGCGAGTGACCAGGGCTTACTGAGTGAAAAGGCTTACTGAGTGAAAATCATCAGGCGCATCAGTTGGCGGCGCAGTTAAGGCTTACTGAGTGAAAAGGCTTACTGAGTGAACAACGTCCATGTCGGGATGC |
| PA14_44490 | anr | GCCAGACTGCAATCCTTGCAGTGTGCTTGG | TTTCGTCCGCGAGTGACCAGGGCTTACTGAGTGAAAAGGCTTACTGAGTGAAAAGCCAGACTGCAATCCTTGCAGTGTGCTTGGAAGGCTTACTGAGTGAAAAGGCTTACTGAGTGAACAACGTCCATGTCGGGATGC |
| PA14_44490 | anr | CAGTGCGCGCACCTTGATGGTTTCGGCCAT | TTTCGTCCGCGAGTGACCAGGGCTTACTGAGTGAAAAGGCTTACTGAGTGAAAACAGTGCGCGCACCTTGATGGTTTCGGCCATAAGGCTTACTGAGTGAAAAGGCTTACTGAGTGAACAACGTCCATGTCGGGATGC |
| PA14_44490 | anr | GAGGATGTGCACTTCCTTGCCTTCCGCGCT | TTTCGTCCGCGAGTGACCAGGGCTTACTGAGTGAAAAGGCTTACTGAGTGAAAAGAGGATGTGCACTTCCTTGCCTTCCGCGCTAAGGCTTACTGAGTGAAAAGGCTTACTGAGTGAACAACGTCCATGTCGGGATGC |
| PA14_44490 | anr | GATGCGCTCGTCGGCAGTCTTCTTCGACAG | TTTCGTCCGCGAGTGACCAGGGCTTACTGAGTGAAAAGGCTTACTGAGTGAAAAGATGCGCTCGTCGGCAGTCTTCTTCGACAGAAGGCTTACTGAGTGAAAAGGCTTACTGAGTGAACAACGTCCATGTCGGGATGC |
| PA14_44490 | anr | AAGGATCGCCCTGGCGGAACAGGAATTCGC | TTTCGTCCGCGAGTGACCAGGGCTTACTGAGTGAAAAGGCTTACTGAGTGAAAAAAGGATCGCCCTGGCGGAACAGGAATTCGCAAGGCTTACTGAGTGAAAAGGCTTACTGAGTGAACAACGTCCATGTCGGGATGC |
| PA14_44490 | anr | TGGGCAGGTGGAAGCCGGTGATCTGCTCTT | TTTCGTCCGCGAGTGACCAGGGCTTACTGAGTGAAAAGGCTTACTGAGTGAAAATGGGCAGGTGGAAGCCGGTGATCTGCTCTTAAGGCTTACTGAGTGAAAAGGCTTACTGAGTGAACAACGTCCATGTCGGGATGC |
| PA14_44490 | anr | TCAGGCCGTTCTGCTGGAAGCGGGTGAAGA | TTTCGTCCGCGAGTGACCAGGGCTTACTGAGTGAAAAGGCTTACTGAGTGAAAATCAGGCCGTTCTGCTGGAAGCGGGTGAAGAAAGGCTTACTGAGTGAAAAGGCTTACTGAGTGAACAACGTCCATGTCGGGATGC |
| PA14_44490 | anr | TCTTCAGGGGACGACCGCGCTTGACGATCT | TTTCGTCCGCGAGTGACCAGGGCTTACTGAGTGAAAAGGCTTACTGAGTGAAAATCTTCAGGGGACGACCGCGCTTGACGATCTAAGGCTTACTGAGTGAAAAGGCTTACTGAGTGAACAACGTCCATGTCGGGATGC |
| PA14_44490 | anr | CATCATCTGCTGGTCATCGCGGATTTCCCG | TTTCGTCCGCGAGTGACCAGGGCTTACTGAGTGAAAAGGCTTACTGAGTGAAAACATCATCTGCTGGTCATCGCGGATTTCCCGAAGGCTTACTGAGTGAAAAGGCTTACTGAGTGAACAACGTCCATGTCGGGATGC |
| PA14_44490 | anr | AATCTCGCAGACCGAGGTGGTTTCCAGGGC | TTTCGTCCGCGAGTGACCAGGGCTTACTGAGTGAAAAGGCTTACTGAGTGAAAAAATCTCGCAGACCGAGGTGGTTTCCAGGGCAAGGCTTACTGAGTGAAAAGGCTTACTGAGTGAACAACGTCCATGTCGGGATGC |
| PA14_44490 | anr | TGATGCTGAAGGTTTTCAGGGCGCCGGAGC | TTTCGTCCGCGAGTGACCAGGGCTTACTGAGTGAAAAGGCTTACTGAGTGAAAATGATGCTGAAGGTTTTCAGGGCGCCGGAGCAAGGCTTACTGAGTGAAAAGGCTTACTGAGTGAACAACGTCCATGTCGGGATGC |
| PA14_44490 | anr | ATCCATGTCTTCCACGGTCAGCGACAGGGG | TTTCGTCCGCGAGTGACCAGGGCTTACTGAGTGAAAAGGCTTACTGAGTGAAAAATCCATGTCTTCCACGGTCAGCGACAGGGGAAGGCTTACTGAGTGAAAAGGCTTACTGAGTGAACAACGTCCATGTCGGGATGC |
| PA14_44490 | anr | ATAGGTCTCGGTATCCATCCCGGACAGGCC | TTTCGTCCGCGAGTGACCAGGGCTTACTGAGTGAAAAGGCTTACTGAGTGAAAAATAGGTCTCGGTATCCATCCCGGACAGGCCAAGGCTTACTGAGTGAAAAGGCTTACTGAGTGAACAACGTCCATGTCGGGATGC |
| PA14_48060 | aprA | AACTGGACGAGGTGGCGCTGTAGAAGTCGC | TTTCGTCCGCGAGTGACCAGGGGGCCAGCGCAATAAAAGGGCCAGCGCAATAAAAAACTGGACGAGGTGGCGCTGTAGAAGTCGCAAGGGCCAGCGCAATAAAAGGGCCAGCGCAATAACAACGTCCATGTCGGGATGC |
| PA14_48060 | aprA | CCAGCGAGTAGATATCGCTGACATACTTCC | TTTCGTCCGCGAGTGACCAGGGGGCCAGCGCAATAAAAGGGCCAGCGCAATAAAACCAGCGAGTAGATATCGCTGACATACTTCCAAGGGCCAGCGCAATAAAAGGGCCAGCGCAATAACAACGTCCATGTCGGGATGC |
| PA14_48060 | aprA | TTGAGCGCATCCGGTACATCCGGCAGGAAG | TTTCGTCCGCGAGTGACCAGGGGGCCAGCGCAATAAAAGGGCCAGCGCAATAAAATTGAGCGCATCCGGTACATCCGGCAGGAAGAAGGGCCAGCGCAATAAAAGGGCCAGCGCAATAACAACGTCCATGTCGGGATGC |
| PA14_48060 | aprA | GTGCTTTCTGCCAAGACGCCTGTTCGCGGA | TTTCGTCCGCGAGTGACCAGGGGGCCAGCGCAATAAAAGGGCCAGCGCAATAAAAGTGCTTTCTGCCAAGACGCCTGTTCGCGGAAAGGGCCAGCGCAATAAAAGGGCCAGCGCAATAACAACGTCCATGTCGGGATGC |
| PA14_48060 | aprA | TTGGAGTTGAAGCCGTACACCGTGTCGCCG | TTTCGTCCGCGAGTGACCAGGGGGCCAGCGCAATAAAAGGGCCAGCGCAATAAAATTGGAGTTGAAGCCGTACACCGTGTCGCCGAAGGGCCAGCGCAATAAAAGGGCCAGCGCAATAACAACGTCCATGTCGGGATGC |
| PA14_48060 | aprA | CGTATTGAAGAAGTCGTTCGGCTTGGTCAG | TTTCGTCCGCGAGTGACCAGGGGGCCAGCGCAATAAAAGGGCCAGCGCAATAAAACGTATTGAAGAAGTCGTTCGGCTTGGTCAGAAGGGCCAGCGCAATAAAAGGGCCAGCGCAATAACAACGTCCATGTCGGGATGC |
| PA14_48060 | aprA | GAAGTCCTGGCCGGTGTTCTGCTCTTCCCA | TTTCGTCCGCGAGTGACCAGGGGGCCAGCGCAATAAAAGGGCCAGCGCAATAAAAGAAGTCCTGGCCGGTGTTCTGCTCTTCCCAAAGGGCCAGCGCAATAAAAGGGCCAGCGCAATAACAACGTCCATGTCGGGATGC |
| PA14_48060 | aprA | CCTGGGTATACGCATCACTACGACCTTTCA | TTTCGTCCGCGAGTGACCAGGGGGCCAGCGCAATAAAAGGGCCAGCGCAATAAAACCTGGGTATACGCATCACTACGACCTTTCAAAGGGCCAGCGCAATAAAAGGGCCAGCGCAATAACAACGTCCATGTCGGGATGC |
| PA14_48060 | aprA | ACGAATAGGACAGGGTCAGCACCGAGTCGC | TTTCGTCCGCGAGTGACCAGGGGGCCAGCGCAATAAAAGGGCCAGCGCAATAAAAACGAATAGGACAGGGTCAGCACCGAGTCGCAAGGGCCAGCGCAATAAAAGGGCCAGCGCAATAACAACGTCCATGTCGGGATGC |
| PA14_48060 | aprA | TAGTTGCCGTTGGCCGGATTGACGTTGGCG | TTTCGTCCGCGAGTGACCAGGGGGCCAGCGCAATAAAAGGGCCAGCGCAATAAAATAGTTGCCGTTGGCCGGATTGACGTTGGCGAAGGGCCAGCGCAATAAAAGGGCCAGCGCAATAACAACGTCCATGTCGGGATGC |
| PA14_48060 | aprA | GACACATTGCCCTTCAACCCGCCGACATCG | TTTCGTCCGCGAGTGACCAGGGGGCCAGCGCAATAAAAGGGCCAGCGCAATAAAAGACACATTGCCCTTCAACCCGCCGACATCGAAGGGCCAGCGCAATAAAAGGGCCAGCGCAATAACAACGTCCATGTCGGGATGC |
| PA14_48060 | aprA | AAGTTGCCGAAGGTCAGGTCGCCCTGATCG | TTTCGTCCGCGAGTGACCAGGGGGCCAGCGCAATAAAAGGGCCAGCGCAATAAAAAAGTTGCCGAAGGTCAGGTCGCCCTGATCGAAGGGCCAGCGCAATAAAAGGGCCAGCGCAATAACAACGTCCATGTCGGGATGC |
| PA14_48060 | aprA | ATCTGCTCCGCCGCCTGGTCTACGGTATAG | TTTCGTCCGCGAGTGACCAGGGGGCCAGCGCAATAAAAGGGCCAGCGCAATAAAAATCTGCTCCGCCGCCTGGTCTACGGTATAGAAGGGCCAGCGCAATAAAAGGGCCAGCGCAATAACAACGTCCATGTCGGGATGC |
| PA14_48060 | aprA | ATTGCAGCGACAACTTGGCCTGGGCCTGCT | TTTCGTCCGCGAGTGACCAGGGGGCCAGCGCAATAAAAGGGCCAGCGCAATAAAAATTGCAGCGACAACTTGGCCTGGGCCTGCTAAGGGCCAGCGCAATAAAAGGGCCAGCGCAATAACAACGTCCATGTCGGGATGC |
| PA14_48060 | aprA | AGCTGCTGTTGATCAGGTACCAGGATTGCC | TTTCGTCCGCGAGTGACCAGGGGGCCAGCGCAATAAAAGGGCCAGCGCAATAAAAAGCTGCTGTTGATCAGGTACCAGGATTGCCAAGGGCCAGCGCAATAAAAGGGCCAGCGCAATAACAACGTCCATGTCGGGATGC |
| PA14_48060 | aprA | ATGTCGTCCAGCAGCGGTGCCGAGGAATAG | TTTCGTCCGCGAGTGACCAGGGGGCCAGCGCAATAAAAGGGCCAGCGCAATAAAAATGTCGTCCAGCAGCGGTGCCGAGGAATAGAAGGGCCAGCGCAATAAAAGGGCCAGCGCAATAACAACGTCCATGTCGGGATGC |
| PA14_48060 | aprA | ATTGACCAATTCGTCCCCGCCCCGCGCATA | TTTCGTCCGCGAGTGACCAGGGGGCCAGCGCAATAAAAGGGCCAGCGCAATAAAAATTGACCAATTCGTCCCCGCCCCGCGCATAAAGGGCCAGCGCAATAAAAGGGCCAGCGCAATAACAACGTCCATGTCGGGATGC |
| PA14_48060 | aprA | GACGAAGTGGATATTGGTGACGTCCGCCCA | TTTCGTCCGCGAGTGACCAGGGGGCCAGCGCAATAAAAGGGCCAGCGCAATAAAAGACGAAGTGGATATTGGTGACGTCCGCCCAAAGGGCCAGCGCAATAAAAGGGCCAGCGCAATAACAACGTCCATGTCGGGATGC |
| PA14_48060 | aprA | TCAGGTTGGCCCCGTAGAGCTTCTGGATCG | TTTCGTCCGCGAGTGACCAGGGGGCCAGCGCAATAAAAGGGCCAGCGCAATAAAATCAGGTTGGCCCCGTAGAGCTTCTGGATCGAAGGGCCAGCGCAATAAAAGGGCCAGCGCAATAACAACGTCCATGTCGGGATGC |
| PA14_48060 | aprA | AGCTCATCACCGAATAGGCGCGGGTGTCCT | TTTCGTCCGCGAGTGACCAGGGGGCCAGCGCAATAAAAGGGCCAGCGCAATAAAAAGCTCATCACCGAATAGGCGCGGGTGTCCTAAGGGCCAGCGCAATAAAAGGGCCAGCGCAATAACAACGTCCATGTCGGGATGC |
| PA14_44370 | ccoN1 | AGCGGCCATTCCAGTTCGGCGTATTCCTTG | TTTCGTCCGCGAGTGACCAGCTGGTGATAACGCTAAACTGGTGATAACGCTAAAAGCGGCCATTCCAGTTCGGCGTATTCCTTGAACTGGTGATAACGCTAAACTGGTGATAACGCTACAACGTCCATGTCGGGATGC |
| PA14_44370 | ccoN1 | GGCGCCGGAGTAGGCGGAGTAGGACTTGAA | TTTCGTCCGCGAGTGACCAGCTGGTGATAACGCTAAACTGGTGATAACGCTAAAGGCGCCGGAGTAGGCGGAGTAGGACTTGAAAACTGGTGATAACGCTAAACTGGTGATAACGCTACAACGTCCATGTCGGGATGC |
| PA14_44370 | ccoN1 | TGCACCACGTAGTACGAGGTGGCGAAGAGG | TTTCGTCCGCGAGTGACCAGCTGGTGATAACGCTAAACTGGTGATAACGCTAAATGCACCACGTAGTACGAGGTGGCGAAGAGGAACTGGTGATAACGCTAAACTGGTGATAACGCTACAACGTCCATGTCGGGATGC |
| PA14_44370 | ccoN1 | GGCGAACTGGCGGACCACCTTATAGTTATA | TTTCGTCCGCGAGTGACCAGCTGGTGATAACGCTAAACTGGTGATAACGCTAAAGGCGAACTGGCGGACCACCTTATAGTTATAAACTGGTGATAACGCTAAACTGGTGATAACGCTACAACGTCCATGTCGGGATGC |
| PA14_44370 | ccoN1 | CAGGACACCGAGACCCATTCCAATGACCCC | TTTCGTCCGCGAGTGACCAGCTGGTGATAACGCTAAACTGGTGATAACGCTAAACAGGACACCGAGACCCATTCCAATGACCCCAACTGGTGATAACGCTAAACTGGTGATAACGCTACAACGTCCATGTCGGGATGC |
| PA14_44370 | ccoN1 | TGCGACCAGTGCTTCAACGAAGGAGTAGGT | TTTCGTCCGCGAGTGACCAGCTGGTGATAACGCTAAACTGGTGATAACGCTAAATGCGACCAGTGCTTCAACGAAGGAGTAGGTAACTGGTGATAACGCTAAACTGGTGATAACGCTACAACGTCCATGTCGGGATGC |
| PA14_44370 | ccoN1 | TCTGCGGGCGACCGAAGACTTTCGGGATCA | TTTCGTCCGCGAGTGACCAGCTGGTGATAACGCTAAACTGGTGATAACGCTAAATCTGCGGGCGACCGAAGACTTTCGGGATCAAACTGGTGATAACGCTAAACTGGTGATAACGCTACAACGTCCATGTCGGGATGC |
| PA14_44370 | ccoN1 | TTCAATTCGGGCCATACCAGTTGGGCGGCG | TTTCGTCCGCGAGTGACCAGCTGGTGATAACGCTAAACTGGTGATAACGCTAAATTCAATTCGGGCCATACCAGTTGGGCGGCGAACTGGTGATAACGCTAAACTGGTGATAACGCTACAACGTCCATGTCGGGATGC |

| | | | |
|---|---|---|---|
| PA14_44370 | ccoN1 | TGTCTGAAATCAGTCGGGCTTGCGAGGTAC | TTTCGTCCGCGAGTGACCAGCTGGTGATAACGCTAAACTGGTGATAACGCTAAATGTCTGAAATCAGTCGGGCTTGCGAGGTACAACTGGTGATAACGCTAAACTGGTGATAACGCTACAACGTCCATGTCGGGATGC |
| PA14_44370 | ccoN1 | CGATCGCGTAGGTGATCCAGACGATCGCCA | TTTCGTCCGCGAGTGACCAGCTGGTGATAACGCTAAACTGGTGATAACGCTAAACGATCGCGTAGGTGATCCAGACGATCGCCAAACTGGTGATAACGCTAAACTGGTGATAACGCTACAACGTCCATGTCGGGATGC |
| PA14_44370 | ccoN1 | CACCAGGATGAAGGCACCGTAGAACCAGTT | TTTCGTCCGCGAGTGACCAGCTGGTGATAACGCTAAACTGGTGATAACGCTAAACACCAGGATGAAGGCACCGTAGAACCAGTTAACTGGTGATAACGCTAAACTGGTGATAACGCTACAACGTCCATGTCGGGATGC |
| PA14_44370 | ccoN1 | AGGAAGAAACCACCGCCGATCATGCGCACG | TTTCGTCCGCGAGTGACCAGCTGGTGATAACGCTAAACTGGTGATAACGCTAAAGGAAGAAACCACCGCCGATCATGCGCACGAACTGGTGATAACGCTAAACTGGTGATAACGCTACAACGTCCATGTCGGGATGC |
| PA14_44370 | ccoN1 | GATCACGGCCTGCCAACCCCAGAAGGTGAA | TTTCGTCCGCGAGTGACCAGCTGGTGATAACGCTAAACTGGTGATAACGCTAAAGATCACGGCCTGCCAACCCCAGAAGGTGAAAACTGGTGATAACGCTAAACTGGTGATAACGCTACAACGTCCATGTCGGGATGC |
| PA14_68330 | arcA | CAGGTTGTGCATCTCGAGGACGTCGATGCC | TTTCGTCCGCGAGTGACCAGCCGCAAGTAATCTGTAACCGCAAGTAATCTGTAACAGGTTGTGCATCTCGAGGACGTCGATGCCAACCGCAAGTAATCTGTAACCGCAAGTAATCTGTCAACGTCCATGTCGGGATGC |
| PA14_68330 | arcA | TGTCGGCGGTGATCTTGCGATCGAGGATCC | TTTCGTCCGCGAGTGACCAGCCGCAAGTAATCTGTAACCGCAAGTAATCTGTAATGTCGGCGGTGATCTTGCGATCGAGGATCCAACCGCAAGTAATCTGTAACCGCAAGTAATCTGTCAACGTCCATGTCGGGATGC |
| PA14_68330 | arcA | GCTCGCGCATCTTGGTGACGAAGTCGAAGT | TTTCGTCCGCGAGTGACCAGCCGCAAGTAATCTGTAACCGCAAGTAATCTGTAAGCTCGCGCATCTTGGTGACGAAGTCGAAGTAACCGCAAGTAATCTGTAACCGCAAGTAATCTGTCAACGTCCATGTCGGGATGC |
| PA14_68330 | arcA | ATGTTCATGCCGTAGGGGCTGCTCGGATCG | TTTCGTCCGCGAGTGACCAGCCGCAAGTAATCTGTAACCGCAAGTAATCTGTAAATGTTCATGCCGTAGGGGCTGCTCGGATCGAACCGCAAGTAATCTGTAACCGCAAGTAATCTGTCAACGTCCATGTCGGGATGC |
| PA14_68330 | arcA | AAGTGGTGTCGCGGGTGAACTGGGTGTTCG | TTTCGTCCGCGAGTGACCAGCCGCAAGTAATCTGTAACCGCAAGTAATCTGTAAAAGTGGTGTCGCGGGTGAACTGGGTGTTCGAACCGCAAGTAATCTGTAACCGCAAGTAATCTGTCAACGTCCATGTCGGGATGC |
| PA14_68330 | arcA | GGAACTTGTAGATGGCGGTGGTCAGCAGGG | TTTCGTCCGCGAGTGACCAGCCGCAAGTAATCTGTAACCGCAAGTAATCTGTAAGGAACTTGTAGATGGCGGTGGTCAGCAGGGAACCGCAAGTAATCTGTAACCGCAAGTAATCTGTCAACGTCCATGTCGGGATGC |
| PA14_68330 | arcA | TAGGTGTTGCGGTCGTAGCCGACTACCACG | TTTCGTCCGCGAGTGACCAGCCGCAAGTAATCTGTAACCGCAAGTAATCTGTAATAGGTGTTGCGGTCGTAGCCGACTACCACGAACCGCAAGTAATCTGTAACCGCAAGTAATCTGTCAACGTCCATGTCGGGATGC |
| PA14_68330 | arcA | AGCAACTCGTCGCAGTTGCTAGGGGTCAGG | TTTCGTCCGCGAGTGACCAGCCGCAAGTAATCTGTAACCGCAAGTAATCTGTAAAGCAACTCGTCGCAGTTGCTAGGGGTCAGGAACCGCAAGTAATCTGTAACCGCAAGTAATCTGTCAACGTCCATGTCGGGATGC |
| PA14_68330 | arcA | ACTCGCGGTACATCTTGAGGATGTTGGCGC | TTTCGTCCGCGAGTGACCAGCCGCAAGTAATCTGTAACCGCAAGTAATCTGTAAACTCGCGGTACATCTTGAGGATGTTGGCGCAACCGCAAGTAATCTGTAACCGCAAGTAATCTGTCAACGTCCATGTCGGGATGC |
| PA14_68330 | arcA | TGGCCTGGTTCACCCAGATCACGTCGTCGA | TTTCGTCCGCGAGTGACCAGCCGCAAGTAATCTGTAACCGCAAGTAATCTGTAATGGCCTGGTTCACCCAGATCACGTCGTCGAAACCGCAAGTAATCTGTAACCGCAAGTAATCTGTCAACGTCCATGTCGGGATGC |
| PA14_68330 | arcA | GGCTTCGGAGTGGACGCCAAGTTTGGTTTT | TTTCGTCCGCGAGTGACCAGCCGCAAGTAATCTGTAACCGCAAGTAATCTGTAAGGCTTCGGAGTGGACGCCAAGTTTGGTTTTAACCGCAAGTAATCTGTAACCGCAAGTAATCTGTCAACGTCCATGTCGGGATGC |
| PA14_68330 | arcA | GGCGAGCAGACCATCACTTTGCGCAGTTTG | TTTCGTCCGCGAGTGACCAGCCGCAAGTAATCTGTAACCGCAAGTAATCTGTAAGGCGAGCAGACCATCACTTTGCGCAGTTTGAACCGCAAGTAATCTGTAACCGCAAGTAATCTGTCAACGTCCATGTCGGGATGC |
| PA14_68330 | arcA | TTGTTACCGTCGTCCCATTGCTCGCGCTCG | TTTCGTCCGCGAGTGACCAGCCGCAAGTAATCTGTAACCGCAAGTAATCTGTAATTGTTACCGTCGTCCCATTGCTCGCGCTCGAACCGCAAGTAATCTGTAACCGCAAGTAATCTGTCAACGTCCATGTCGGGATGC |
| PA14_68330 | arcA | TTCAGCGCTTCCGGGTTCTGGATGGTCTCG | TTTCGTCCGCGAGTGACCAGCCGCAAGTAATCTGTAACCGCAAGTAATCTGTAATTCAGCGCTTCCGGGTTCTGGATGGTCTCGAACCGCAAGTAATCTGTAACCGCAAGTAATCTGTCAACGTCCATGTCGGGATGC |
| PA14_68330 | arcA | TGACCACTTCCGGGAAGACCGTGACCAGGT | TTTCGTCCGCGAGTGACCAGCCGCAAGTAATCTGTAACCGCAAGTAATCTGTAATGACCACTTCCGGGAAGACCGTGACCAGGTAACCGCAAGTAATCTGTAACCGCAAGTAATCTGTCAACGTCCATGTCGGGATGC |
| PA14_73260 | atpA | TCACGGTACTGGGCCAGAGCGGTACGAATG | TTTCGTCCGCGAGTGACCAGTATCGCAACCGACCTAATATCGCAACCGACCTAATCACGGTACTGGGCCAGAGCGGTACGAATGAATATCGCAACCGACCTAATATCGCAACCGACCTCAACGTCCATGTCGGGATGC |
| PA14_73260 | atpA | TGGCCACGTAGACGCACTTAATGCCGCTGT | TTTCGTCCGCGAGTGACCAGTATCGCAACCGACCTAATATCGCAACCGACCTAATGGCCACGTAGACGCACTTAATGCCGCTGTAATATCGCAACCGACCTAATATCGCAACCGACCTCAACGTCCATGTCGGGATGC |
| PA14_73260 | atpA | TGCTCCAGGTTCAGCGCCATACCGTAGACG | TTTCGTCCGCGAGTGACCAGTATCGCAACCGACCTAATATCGCAACCGACCTAATGCTCCAGGTTCAGCGCCATACCGTAGACGAATATCGCAACCGACCTAATATCGCAACCGACCTCAACGTCCATGTCGGGATGC |
| PA14_73260 | atpA | TTCTTGATGATCTTGGTCTGGGCCGCGCCA | TTTCGTCCGCGAGTGACCAGTATCGCAACCGACCTAATATCGCAACCGACCTAATTCTTGATGATCTTGGTCTGGGCCGCGCCAAATATCGCAACCGACCTAATATCGCAACCGACCTCAACGTCCATGTCGGGATGC |
| PA14_73260 | atpA | CGGACGGATACCCGAGTTGAACATGGCGGA | TTTCGTCCGCGAGTGACCAGTATCGCAACCGACCTAATATCGCAACCGACCTAACGGACGGATACCCGAGTTGAACATGGCGGAAATATCGCAACCGACCTAATATCGCAACCGACCTCAACGTCCATGTCGGGATGC |
| PA14_73260 | atpA | ACGTTGGTCGGAACGAACGCGGAAACGTCG | TTTCGTCCGCGAGTGACCAGTATCGCAACCGACCTAATATCGCAACCGACCTAAACGTTGGTCGGAACGAACGCGGAAACGTCGAATATCGCAACCGACCTAATATCGCAACCGACCTCAACGTCCATGTCGGGATGC |
| PA14_73260 | atpA | AGGGACAGCGACATCTCGGCGATCGACATC | TTTCGTCCGCGAGTGACCAGTATCGCAACCGACCTAATATCGCAACCGACCTAAAGGGACAGCGACATCTCGGCGATCGACATCAATATCGCAACCGACCTAATATCGCAACCGACCTCAACGTCCATGTCGGGATGC |
| PA14_73260 | atpA | AGCAGCAGGGAGATCTGGCGATAAGCCACG | TTTCGTCCGCGAGTGACCAGTATCGCAACCGACCTAATATCGCAACCGACCTAAAGCAGCAGGGAGATCTGGCGATAAGCCACGAATATCGCAACCGACCTAATATCGCAACCGACCTCAACGTCCATGTCGGGATGC |
| PA14_73260 | atpA | TGCCTTCGTTTCGTGCTTGGGAGGCGACGT | TTTCGTCCGCGAGTGACCAGTATCGCAACCGACCTAATATCGCAACCGACCTAATGCCTTCGTTTCGTGCTTGGGAGGCGACGTAATATCGCAACCGACCTAATATCGCAACCGACCTCAACGTCCATGTCGGGATGC |
| PA14_73260 | atpA | TCATCTCGCCGTACATGACGTCGGCCAGAC | TTTCGTCCGCGAGTGACCAGTATCGCAACCGACCTAATATCGCAACCGACCTAATCATCTCGCCGTACATGACGTCGGCCAGACAATATCGCAACCGACCTAATATCGCAACCGACCTCAACGTCCATGTCGGGATGC |
| PA14_73260 | atpA | AGTTCCGGACCAACCGGAACCTCCAGGATG | TTTCGTCCGCGAGTGACCAGTATCGCAACCGACCTAATATCGCAACCGACCTAAAGTTCCGGACCAACCGGAACCTCCAGGATGAATATCGCAACCGACCTAATATCGCAACCGACCTCAACGTCCATGTCGGGATGC |
| PA14_73260 | atpA | AGATAGAACACGTCGCCCGGGTAGGCTTCG | TTTCGTCCGCGAGTGACCAGTATCGCAACCGACCTAATATCGCAACCGACCTAAAGATAGAACACGTCGCCCGGGTAGGCTTCGAATATCGCAACCGACCTAATATCGCAACCGACCTCAACGTCCATGTCGGGATGC |
| PA14_73260 | atpA | ATTCACCCAGGACCACGGCACCCACGGAAT | TTTCGTCCGCGAGTGACCAGTATCGCAACCGACCTAATATCGCAACCGACCTAAATTCACCCAGGACCACGGCACCCACGGAATAATATCGCAACCGACCTAATATCGCAACCGACCTCAACGTCCATGTCGGGATGC |
| PA14_73260 | atpA | TTCACCGCGATCACGGAAGTATTCGCCCAT | TTTCGTCCGCGAGTGACCAGTATCGCAACCGACCTAATATCGCAACCGACCTAATTCACCGCGATCACGGAAGTATTCGCCCATAATATCGCAACCGACCTAATATCGCAACCGACCTCAACGTCCATGTCGGGATGC |
| PA14_73260 | atpA | ATACTCTTCGGAAACGCGGGAAGCGCGCTC | TTTCGTCCGCGAGTGACCAGTATCGCAACCGACCTAATATCGCAACCGACCTAAATACTCTTCGGAAACGCGGGAAGCGCGCTCAATATCGCAACCGACCTAATATCGCAACCGACCTCAACGTCCATGTCGGGATGC |
| PA14_73260 | atpA | AGATGCGCACGATACCGTCGGACACGCTGA | TTTCGTCCGCGAGTGACCAGTATCGCAACCGACCTAATATCGCAACCGACCTAAAGATGCGCACGATACCGTCGGACACGCTGAAATATCGCAACCGACCTAATATCGCAACCGACCTCAACGTCCATGTCGGGATGC |
| PA14_73260 | atpA | TGGCGTCGACCGACTTGTAGCCGGTCTGTA | TTTCGTCCGCGAGTGACCAGTATCGCAACCGACCTAATATCGCAACCGACCTAATGGCGTCGACCGACTTGTAGCCGGTCTGTAAATATCGCAACCGACCTAATATCGCAACCGACCTCAACGTCCATGTCGGGATGC |
| PA14_73260 | atpA | CTTGATGCCCGCGTCGATTTCGTCGTTGAA | TTTCGTCCGCGAGTGACCAGTATCGCAACCGACCTAATATCGCAACCGACCTAACTTGATGCCCGCGTCGATTTCGTCGTTGAAAATATCGCAACCGACCTAATATCGCAACCGACCTCAACGTCCATGTCGGGATGC |
| PA14_73260 | atpA | GTTTCTCGATGCGCCCCTTGATGATTTCAC | TTTCGTCCGCGAGTGACCAGTATCGCAACCGACCTAATATCGCAACCGACCTAAGTTTCTCGATGCGCCCCTTGATGATTTCACAATATCGCAACCGACCTAATATCGCAACCGACCTCAACGTCCATGTCGGGATGC |
| PA14_73260 | atpA | GACGTTGGCGATAGTGGACTGCTTCTGGCC | TTTCGTCCGCGAGTGACCAGTATCGCAACCGACCTAATATCGCAACCGACCTAAGACGTTGGCGATAGTGGACTGCTTCTGGCCAATATCGCAACCGACCTAATATCGCAACCGACCTCAACGTCCATGTCGGGATGC |
| PA14_44340 | ccoN2 | TATCGATCGGCCACTCCAGCTCGGCGTACT | TTTCGTCCGCGAGTGACCAGATTCATCCGCAGGTCAAATTCATCCGCAGGTCAATATCGATCGGCCACTCCAGCTCGGCGTACTAAATTCATCCGCAGGTCAAATTCATCCGCAGGTCCAACGTCCATGTCGGGATGC |
| PA14_44340 | ccoN2 | AGCGAGCCGATGGAAACCATCGCGACCCAG | TTTCGTCCGCGAGTGACCAGATTCATCCGCAGGTCAAATTCATCCGCAGGTCAAAGCGAGCCGATGGAAACCATCGCGACCCAGAAATTCATCCGCAGGTCAAATTCATCCGCAGGTCCAACGTCCATGTCGGGATGC |
| PA14_44340 | ccoN2 | GAAGGTGAAGGAGGCGAGGGTGTCGGAGAA | TTTCGTCCGCGAGTGACCAGATTCATCCGCAGGTCAAATTCATCCGCAGGTCAAGAAGGTGAAGGAGGCGAGGGTGTCGGAGAAAAATTCATCCGCAGGTCAAATTCATCCGCAGGTCCAACGTCCATGTCGGGATGC |
| PA14_44340 | ccoN2 | CAGGAAGATGAAACCGCCGAGCATGCGCAC | TTTCGTCCGCGAGTGACCAGATTCATCCGCAGGTCAAATTCATCCGCAGGTCAACAGGAAGATGAAACCGCCGAGCATGCGCACAAATTCATCCGCAGGTCAAATTCATCCGCAGGTCCAACGTCCATGTCGGGATGC |
| PA14_44340 | ccoN2 | GATCAGGACACCGACGGTCATGCCGACGAT | TTTCGTCCGCGAGTGACCAGATTCATCCGCAGGTCAAATTCATCCGCAGGTCAAGATCAGGACACCGACGGTCATGCCGACGATAAATTCATCCGCAGGTCAAATTCATCCGCAGGTCCAACGTCCATGTCGGGATGC |
| PA14_44340 | ccoN2 | ATAGGCGGACTCTGGGTTAGTGGTGCTCAT | TTTCGTCCGCGAGTGACCAGATTCATCCGCAGGTCAAATTCATCCGCAGGTCAAATAGGCGGACTCTGGGTTAGTGGTGCTCATAAATTCATCCGCAGGTCAAATTCATCCGCAGGTCCAACGTCCATGTCGGGATGC |
| PA14_44340 | ccoN2 | TGAAGGTCACCGGCAGCTCGAGGTTGTTCA | TTTCGTCCGCGAGTGACCAGATTCATCCGCAGGTCAAATTCATCCGCAGGTCAATGAAGGTCACCGGCAGCTCGAGGTTGTTCAAAATTCATCCGCAGGTCAAATTCATCCGCAGGTCCAACGTCCATGTCGGGATGC |
| PA14_44340 | ccoN2 | ACGTGGAGCATGGCGACGGTGAGGATGAAC | TTTCGTCCGCGAGTGACCAGATTCATCCGCAGGTCAAATTCATCCGCAGGTCAAACGTGGAGCATGGCGACGGTGAGGATGAACAAATTCATCCGCAGGTCAAATTCATCCGCAGGTCCAACGTCCATGTCGGGATGC |
| PA14_44340 | ccoN2 | AGCCGAAGTTCAGGTCGGGCCATACCAGCT | TTTCGTCCGCGAGTGACCAGATTCATCCGCAGGTCAAATTCATCCGCAGGTCAAAGCCGAAGTTCAGGTCGGGCCATACCAGCTAAATTCATCCGCAGGTCAAATTCATCCGCAGGTCCAACGTCCATGTCGGGATGC |
| PA14_44340 | ccoN2 | GATGTAAAGCACGGTACCGATGGTGGCCAG | TTTCGTCCGCGAGTGACCAGATTCATCCGCAGGTCAAATTCATCCGCAGGTCAAGATGTAAAGCACGGTACCGATGGTGGCCAGAAATTCATCCGCAGGTCAAATTCATCCGCAGGTCCAACGTCCATGTCGGGATGC |
| PA14_44340 | ccoN2 | ATGATGGCGAATTGGCGGACCACCTTGTAG | TTTCGTCCGCGAGTGACCAGATTCATCCGCAGGTCAAATTCATCCGCAGGTCAAATGATGGCGAATTGGCGGACCACCTTGTAGAAATTCATCCGCAGGTCAAATTCATCCGCAGGTCCAACGTCCATGTCGGGATGC |
| PA14_44340 | ccoN2 | CCCAGATGTACACCGCGATCAGCGCCCAGA | TTTCGTCCGCGAGTGACCAGATTCATCCGCAGGTCAAATTCATCCGCAGGTCAACCCAGATGTACACCGCGATCAGCGCCCAGAAAATTCATCCGCAGGTCAAATTCATCCGCAGGTCCAACGTCCATGTCGGGATGC |
| PA14_44340 | ccoN2 | CATCTTCTCGCGGCCGAAGACCTTCGGGAT | TTTCGTCCGCGAGTGACCAGATTCATCCGCAGGTCAAATTCATCCGCAGGTCAACATCTTCTCGCGGCCGAAGACCTTCGGGATAAATTCATCCGCAGGTCAAATTCATCCGCAGGTCCAACGTCCATGTCGGGATGC |
| PA14_10500 | ccoN4 | AGTTCATCTGGGGCCAGACCAGTTGCGAGG | TTTCGTCCGCGAGTGACCAGACTCAAGGCCGGTATAAACTCAAGGCCGGTATAAAGTTCATCTGGGGCCAGACCAGTTGCGAGGAAACTCAAGGCCGGTATAAACTCAAGGCCGGTATCAACGTCCATGTCGGGATGC |
| PA14_10500 | ccoN4 | GTAGGCAGGTTCCGTCGTTGCTTTGTTCAT | TTTCGTCCGCGAGTGACCAGACTCAAGGCCGGTATAAACTCAAGGCCGGTATAAGTAGGCAGGTTCCGTCGTTGCTTTGTTCATAAACTCAAGGCCGGTATAAACTCAAGGCCGGTATCAACGTCCATGTCGGGATGC |
| PA14_10500 | ccoN4 | ATGATGGCGAACTGGCGAACCACCTTGTAG | TTTCGTCCGCGAGTGACCAGACTCAAGGCCGGTATAAACTCAAGGCCGGTATAAATGATGGCGAACTGGCGAACCACCTTGTAGAAACTCAAGGCCGGTATAAACTCAAGGCCGGTATCAACGTCCATGTCGGGATGC |
| PA14_10500 | ccoN4 | TGGACGATGGACAGCCGGTAGGAGTAGACC | TTTCGTCCGCGAGTGACCAGACTCAAGGCCGGTATAAACTCAAGGCCGGTATAATGGACGATGGACAGCCGGTAGGAGTAGACCAAACTCAAGGCCGGTATAAACTCAAGGCCGGTATCAACGTCCATGTCGGGATGC |
| PA14_10500 | ccoN4 | AGATCACCAGGTTGGTATGCAGCGGCCGCA | TTTCGTCCGCGAGTGACCAGACTCAAGGCCGGTATAAACTCAAGGCCGGTATAAAGATCACCAGGTTGGTATGCAGCGGCCGCAAAACTCAAGGCCGGTATAAACTCAAGGCCGGTATCAACGTCCATGTCGGGATGC |
| PA14_10500 | ccoN4 | TATTCCTTGGTGGTGGTGTTGCCCAGCGGC | TTTCGTCCGCGAGTGACCAGACTCAAGGCCGGTATAAACTCAAGGCCGGTATAATATTCCTTGGTGGTGGTGTTGCCCAGCGGCAAACTCAAGGCCGGTATAAACTCAAGGCCGGTATCAACGTCCATGTCGGGATGC |
| PA14_10500 | ccoN4 | AGGTACGCTGCACCGTGTAGTAGCTGGTGG | TTTCGTCCGCGAGTGACCAGACTCAAGGCCGGTATAAACTCAAGGCCGGTATAAAGGTACGCTGCACCGTGTAGTAGCTGGTGGAAACTCAAGGCCGGTATAAACTCAAGGCCGGTATCAACGTCCATGTCGGGATGC |
| PA14_10500 | ccoN4 | GGCGCCGGAGTACATCGAGTAGGACTTGAA | TTTCGTCCGCGAGTGACCAGACTCAAGGCCGGTATAAACTCAAGGCCGGTATAAGGCGCCGGAGTACATCGAGTAGGACTTGAAAAACTCAAGGCCGGTATAAACTCAAGGCCGGTATCAACGTCCATGTCGGGATGC |
| PA14_10500 | ccoN4 | TGGTCAGGATGAACGAGCCGAAGAACCAGT | TTTCGTCCGCGAGTGACCAGACTCAAGGCCGGTATAAACTCAAGGCCGGTATAATGGTCAGGATGAACGAGCCGAAGAACCAGTAAACTCAAGGCCGGTATAAACTCAAGGCCGGTATCAACGTCCATGTCGGGATGC |
| PA14_10500 | ccoN4 | CCCGTAGACCTTGGGAATCAGGTGGTAGAG | TTTCGTCCGCGAGTGACCAGACTCAAGGCCGGTATAAACTCAAGGCCGGTATAACCCGTAGACCTTGGGAATCAGGTGGTAGAGAAACTCAAGGCCGGTATAAACTCAAGGCCGGTATCAACGTCCATGTCGGGATGC |
| PA14_10500 | ccoN4 | ATCAGTACGCCCAACCCCATGCCGATCACG | TTTCGTCCGCGAGTGACCAGACTCAAGGCCGGTATAAACTCAAGGCCGGTATAAATCAGTACGCCCAACCCCATGCCGATCACGAAACTCAAGGCCGGTATAAACTCAAGGCCGGTATCAACGTCCATGTCGGGATGC |
| PA14_10500 | ccoN4 | GCCAGGTGTTGTAGCTCATCAGCAGCATGC | TTTCGTCCGCGAGTGACCAGACTCAAGGCCGGTATAAACTCAAGGCCGGTATAAGCCAGGTGTTGTAGCTCATCAGCAGCATGCAAACTCAAGGCCGGTATAAACTCAAGGCCGGTATCAACGTCCATGTCGGGATGC |
| PA14_40510 | ccoN3 | ACACGAAGCGCGCCAGGACGTCGGAGATCA | TTTCGTCCGCGAGTGACCAGGCCTTATACAGCGTAAAGCCTTATACAGCGTAAAACACGAAGCGCGCCAGGACGTCGGAGATCAAAGCCTTATACAGCGTAAAGCCTTATACAGCGTACAACGTCCATGTCGGGATGC |
| PA14_40510 | ccoN3 | GGACAGCGAGTTGACCGTCTTGATCGCCAT | TTTCGTCCGCGAGTGACCAGGCCTTATACAGCGTAAAGCCTTATACAGCGTAAAGGACAGCGAGTTGACCGTCTTGATCGCCATAAGCCTTATACAGCGTAAAGCCTTATACAGCGTACAACGTCCATGTCGGGATGC |
| PA14_40510 | ccoN3 | ACGTAGAAGGAGGTGGCGAACAACGCGCTG | TTTCGTCCGCGAGTGACCAGGCCTTATACAGCGTAAAGCCTTATACAGCGTAAAACGTAGAAGGAGGTGGCGAACAACGCGCTGAAGCCTTATACAGCGTAAAGCCTTATACAGCGTACAACGTCCATGTCGGGATGC |
| PA14_40510 | ccoN3 | GATCGCCCAGAAATGCACGATGGACAGGCG | TTTCGTCCGCGAGTGACCAGGCCTTATACAGCGTAAAGCCTTATACAGCGTAAAGATCGCCCAGAAATGCACGATGGACAGGCGAAGCCTTATACAGCGTAAAGCCTTATACAGCGTACAACGTCCATGTCGGGATGC |
| PA14_40510 | ccoN3 | CGTCCGGATCTTTTCTGCTATCCGTATCGT | TTTCGTCCGCGAGTGACCAGGCCTTATACAGCGTAAAGCCTTATACAGCGTAAACGTCCGGATCTTTTCTGCTATCCGTATCGTAAGCCTTATACAGCGTAAAGCCTTATACAGCGTACAACGTCCATGTCGGGATGC |
| PA14_40510 | ccoN3 | TGGTGCGGACCGGCCCAGATGTAGATCGAG | TTTCGTCCGCGAGTGACCAGGCCTTATACAGCGTAAAGCCTTATACAGCGTAAATGGTGCGGACCGGCCCAGATGTAGATCGAGAAGCCTTATACAGCGTAAAGCCTTATACAGCGTACAACGTCCATGTCGGGATGC |
| PA14_40510 | ccoN3 | ATAGGCCGAGTAGGACTTGCCCAGGCTCAC | TTTCGTCCGCGAGTGACCAGGCCTTATACAGCGTAAAGCCTTATACAGCGTAAAATAGGCCGAGTAGGACTTGCCCAGGCTCACAAGCCTTATACAGCGTAAAGCCTTATACAGCGTACAACGTCCATGTCGGGATGC |
| PA14_40510 | ccoN3 | AATGCGTGGTTCACGATATGGACCATCGCG | TTTCGTCCGCGAGTGACCAGGCCTTATACAGCGTAAAGCCTTATACAGCGTAAAAATGCGTGGTTCACGATATGGACCATCGCGAAGCCTTATACAGCGTAAAGCCTTATACAGCGTACAACGTCCATGTCGGGATGC |
| PA14_40510 | ccoN3 | TACCACCACTGGATCATCGCGTCGGTGGCG | TTTCGTCCGCGAGTGACCAGGCCTTATACAGCGTAAAGCCTTATACAGCGTAAATACCACCACTGGATCATCGCGTCGGTGGCGAAGCCTTATACAGCGTAAAGCCTTATACAGCGTACAACGTCCATGTCGGGATGC |
| PA14_40510 | ccoN3 | ATCCACAGCGCCAGTGGCCATTCCATCTCG | TTTCGTCCGCGAGTGACCAGGCCTTATACAGCGTAAAGCCTTATACAGCGTAAAATCCACAGCGCCAGTGGCCATTCCATCTCGAAGCCTTATACAGCGTAAAGCCTTATACAGCGTACAACGTCCATGTCGGGATGC |
| PA14_40510 | ccoN3 | CCGAAGAACAGGTAGGCATAGACCACCCAG | TTTCGTCCGCGAGTGACCAGGCCTTATACAGCGTAAAGCCTTATACAGCGTAAACCGAAGAACAGGTAGGCATAGACCACCCAGAAGCCTTATACAGCGTAAAGCCTTATACAGCGTACAACGTCCATGTCGGGATGC |
| PA14_40510 | ccoN3 | TAGATGTGCTTCACCTTGCGGCGGGCGATG | TTTCGTCCGCGAGTGACCAGGCCTTATACAGCGTAAAGCCTTATACAGCGTAAATAGATGTGCTTCACCTTGCGGCGGGCGATGAAGCCTTATACAGCGTAAAGCCTTATACAGCGTACAACGTCCATGTCGGGATGC |
| PA14_40510 | ccoN3 | AGCGGGTAGCTGACGATCATCCCCACGATC | TTTCGTCCGCGAGTGACCAGGCCTTATACAGCGTAAAGCCTTATACAGCGTAAAAGCGGGTAGCTGACGATCATCCCCACGATCAAGCCTTATACAGCGTAAAGCCTTATACAGCGTACAACGTCCATGTCGGGATGC |
| PA14_40510 | ccoN3 | GGTGAACTGGCGGACGACGTCGTAGTTGTA | TTTCGTCCGCGAGTGACCAGGCCTTATACAGCGTAAAGCCTTATACAGCGTAAAGGTGAACTGGCGGACGACGTCGTAGTTGTAAAGCCTTATACAGCGTAAAGCCTTATACAGCGTACAACGTCCATGTCGGGATGC |
| PA14_40510 | ccoN3 | TCATGTGGTAGAGCGTGCCGATGGAGATCA | TTTCGTCCGCGAGTGACCAGGCCTTATACAGCGTAAAGCCTTATACAGCGTAAATCATGTGGTAGAGCGTGCCGATGGAGATCAAAGCCTTATACAGCGTAAAGCCTTATACAGCGTACAACGTCCATGTCGGGATGC |
| PA14_40510 | ccoN3 | ATCACCAGGCTGGTGTGGATCGGGCGGATG | TTTCGTCCGCGAGTGACCAGGCCTTATACAGCGTAAAGCCTTATACAGCGTAAAATCACCAGGCTGGTGTGGATCGGGCGGATGAAGCCTTATACAGCGTAAAGCCTTATACAGCGTACAACGTCCATGTCGGGATGC |
| PA14_40510 | ccoN3 | GACGATGATGAAGGCGCCGTAGAACCAGTT | TTTCGTCCGCGAGTGACCAGGCCTTATACAGCGTAAAGCCTTATACAGCGTAAAGACGATGATGAAGGCGCCGTAGAACCAGTTAAGCCTTATACAGCGTAAAGCCTTATACAGCGTACAACGTCCATGTCGGGATGC |
| PA14_40510 | ccoN3 | GACGAACGAGTAGGTGAGCGTGCCGTCCTC | TTTCGTCCGCGAGTGACCAGGCCTTATACAGCGTAAAGCCTTATACAGCGTAAAGACGAACGAGTAGGTGAGCGTGCCGTCCTCAAGCCTTATACAGCGTAAAGCCTTATACAGCGTACAACGTCCATGTCGGGATGC |
| PA14_40510 | ccoN3 | AAGCCGACCGAGAGGATGAAGCCGACCACG | TTTCGTCCGCGAGTGACCAGGCCTTATACAGCGTAAAGCCTTATACAGCGTAAAAAGCCGACCGAGAGGATGAAGCCGACCACGAAGCCTTATACAGCGTAAAGCCTTATACAGCGTACAACGTCCATGTCGGGATGC |
| PA14_61200 | cdrA | ATCAATTGCGCCGGCGTGACGGTCAGCTTG | TTTCGTCCGCGAGTGACCAGTCCGTTATGGGCGTAAATCCGTTATGGGCGTAAAATCAATTGCGCCGGCGTGACGGTCAGCTTGAATCCGTTATGGGCGTAAATCCGTTATGGGCGTACAACGTCCATGTCGGGATGC |
| PA14_61200 | cdrA | TTCAGGCCACTGACCTGGTAGGTGAGGGTC | TTTCGTCCGCGAGTGACCAGTCCGTTATGGGCGTAAATCCGTTATGGGCGTAAATTCAGGCCACTGACCTGGTAGGTGAGGGTCAATCCGTTATGGGCGTAAATCCGTTATGGGCGTACAACGTCCATGTCGGGATGC |
| PA14_61200 | cdrA | AAGCCCTTCCAGTTGATGATCGCGCGCTGG | TTTCGTCCGCGAGTGACCAGTCCGTTATGGGCGTAAATCCGTTATGGGCGTAAAAAGCCCTTCCAGTTGATGATCGCGCGCTGGAATCCGTTATGGGCGTAAATCCGTTATGGGCGTACAACGTCCATGTCGGGATGC |
| PA14_61200 | cdrA | AACGCTGGCTGAAATACTCGCTGCCATTGG | TTTCGTCCGCGAGTGACCAGTCCGTTATGGGCGTAAATCCGTTATGGGCGTAAAAACGCTGGCTGAAATACTCGCTGCCATTGGAATCCGTTATGGGCGTAAATCCGTTATGGGCGTACAACGTCCATGTCGGGATGC |
| PA14_61200 | cdrA | ATCACCGTAGACGCGGGTAGCGTCATTGGC | TTTCGTCCGCGAGTGACCAGTCCGTTATGGGCGTAAATCCGTTATGGGCGTAAAATCACCGTAGACGCGGGTAGCGTCATTGGCAATCCGTTATGGGCGTAAATCCGTTATGGGCGTACAACGTCCATGTCGGGATGC |
| PA14_61200 | cdrA | ACGAGTCGTTGGTCGGGAAACTGGCGGAAC | TTTCGTCCGCGAGTGACCAGTCCGTTATGGGCGTAAATCCGTTATGGGCGTAAAACGAGTCGTTGGTCGGGAAACTGGCGGAACAATCCGTTATGGGCGTAAATCCGTTATGGGCGTACAACGTCCATGTCGGGATGC |
| PA14_61200 | cdrA | TGGTGATCAGGGAGCCCACGTCTACCTTGG | TTTCGTCCGCGAGTGACCAGTCCGTTATGGGCGTAAATCCGTTATGGGCGTAAATGGTGATCAGGGAGCCCACGTCTACCTTGGAATCCGTTATGGGCGTAAATCCGTTATGGGCGTACAACGTCCATGTCGGGATGC |
| PA14_61200 | cdrA | CTAGGTCATAGTTGCCACTGGTCAGCACCA | TTTCGTCCGCGAGTGACCAGTCCGTTATGGGCGTAAATCCGTTATGGGCGTAAACTAGGTCATAGTTGCCACTGGTCAGCACCAAATCCGTTATGGGCGTAAATCCGTTATGGGCGTACAACGTCCATGTCGGGATGC |
| PA14_61200 | cdrA | ATCGCTGACGTCCATACCCAGAGCGTAGGC | TTTCGTCCGCGAGTGACCAGTCCGTTATGGGCGTAAATCCGTTATGGGCGTAAAATCGCTGACGTCCATACCCAGAGCGTAGGCAATCCGTTATGGGCGTAAATCCGTTATGGGCGTACAACGTCCATGTCGGGATGC |
| PA14_61200 | cdrA | TCTGGATGTCCCCGGACTTGCTTACCAGCG | TTTCGTCCGCGAGTGACCAGTCCGTTATGGGCGTAAATCCGTTATGGGCGTAAATCTGGATGTCCCCGGACTTGCTTACCAGCGAATCCGTTATGGGCGTAAATCCGTTATGGGCGTACAACGTCCATGTCGGGATGC |
| PA14_61200 | cdrA | AGTATCCCGTAGTTGCCGACGTTCTCCCCG | TTTCGTCCGCGAGTGACCAGTCCGTTATGGGCGTAAATCCGTTATGGGCGTAAAAGTATCCCGTAGTTGCCGACGTTCTCCCCGAATCCGTTATGGGCGTAAATCCGTTATGGGCGTACAACGTCCATGTCGGGATGC |
| PA14_61200 | cdrA | TTCTTCGGACCGCTGACGTCGATCTTGCCG | TTTCGTCCGCGAGTGACCAGTCCGTTATGGGCGTAAATCCGTTATGGGCGTAAATTCTTCGGACCGCTGACGTCGATCTTGCCGAATCCGTTATGGGCGTAAATCCGTTATGGGCGTACAACGTCCATGTCGGGATGC |
| PA14_61200 | cdrA | ATTGCCCTTGCCGGTAGCCGAGACGTCGAT | TTTCGTCCGCGAGTGACCAGTCCGTTATGGGCGTAAATCCGTTATGGGCGTAAAATTGCCCTTGCCGGTAGCCGAGACGTCGATAATCCGTTATGGGCGTAAATCCGTTATGGGCGTACAACGTCCATGTCGGGATGC |
| PA14_61200 | cdrA | TTGATGCCGGACATCGTGGCCGTCAGAGTC | TTTCGTCCGCGAGTGACCAGTCCGTTATGGGCGTAAATCCGTTATGGGCGTAAATTGATGCCGGACATCGTGGCCGTCAGAGTCAATCCGTTATGGGCGTAAATCCGTTATGGGCGTACAACGTCCATGTCGGGATGC |
| PA14_61200 | cdrA | ACAGGATCGGCGCAGAGATATCCACCGCAC | TTTCGTCCGCGAGTGACCAGTCCGTTATGGGCGTAAATCCGTTATGGGCGTAAAACAGGATCGGCGCAGAGATATCCACCGCACAATCCGTTATGGGCGTAAATCCGTTATGGGCGTACAACGTCCATGTCGGGATGC |
| PA14_41230 | clpX | CCTCGTCGAGCTCATCAAGAGTCGCGATCA | TTTCGTCCGCGAGTGACCAGGTCCCGGCGTATCAAAAGTCCCGGCGTATCAAAACCTCGTCGAGCTCATCAAGAGTCGCGATCAAAGTCCCGGCGTATCAAAAGTCCCGGCGTATCAACAACGTCCATGTCGGGATGC |
| PA14_41230 | clpX | ATACCGCTACCGCCAGAACCTTCTTGGCCC | TTTCGTCCGCGAGTGACCAGGTCCCGGCGTATCAAAAGTCCCGGCGTATCAAAAATACCGCTACCGCCAGAACCTTCTTGGCCCAAGTCCCGGCGTATCAAAAGTCCCGGCGTATCAACAACGTCCATGTCGGGATGC |
| PA14_41230 | clpX | AACGCAAGCCACGTGCACCGGTCTTGCGTT | TTTCGTCCGCGAGTGACCAGGTCCCGGCGTATCAAAAGTCCCGGCGTATCAAAAAACGCAAGCCACGTGCACCGGTCTTGCGTTAAGTCCCGGCGTATCAAAAGTCCCGGCGTATCAACAACGTCCATGTCGGGATGC |
| PA14_41230 | clpX | TTGGCTGGCTTTTCGCTGTTCTCGTAGATC | TTTCGTCCGCGAGTGACCAGGTCCCGGCGTATCAAAAGTCCCGGCGTATCAAAATTGGCTGGCTTTTCGCTGTTCTCGTAGATCAAGTCCCGGCGTATCAAAAGTCCCGGCGTATCAACAACGTCCATGTCGGGATGC |
| PA14_41230 | clpX | CTCCACATCGTAATCGCACTTCTGCAAGAG | TTTCGTCCGCGAGTGACCAGGTCCCGGCGTATCAAAAGTCCCGGCGTATCAAAACTCCACATCGTAATCGCACTTCTGCAAGAGAAGTCCCGGCGTATCAAAAGTCCCGGCGTATCAACAACGTCCATGTCGGGATGC |
| PA14_41230 | clpX | GAACTTGACGAGATCTTCCGGCTCGACTTC | TTTCGTCCGCGAGTGACCAGGTCCCGGCGTATCAAAAGTCCCGGCGTATCAAAAGAACTTGACGAGATCTTCCGGCTCGACTTCAAGTCCCGGCGTATCAAAAGTCCCGGCGTATCAACAACGTCCATGTCGGGATGC |
| PA14_41230 | clpX | GTCACGCTGGTTGAGGCGCTTGTAGTGGTT | TTTCGTCCGCGAGTGACCAGGTCCCGGCGTATCAAAAGTCCCGGCGTATCAAAAGTCACGCTGGTTGAGGCGCTTGTAGTGGTTAAGTCCCGGCGTATCAAAAGTCCCGGCGTATCAACAACGTCCATGTCGGGATGC |
| PA14_41230 | clpX | TCGTCGATGTAGACGATGCCCATCTGGGCC | TTTCGTCCGCGAGTGACCAGGTCCCGGCGTATCAAAAGTCCCGGCGTATCAAAATCGTCGATGTAGACGATGCCCATCTGGGCCAAGTCCCGGCGTATCAAAAGTCCCGGCGTATCAACAACGTCCATGTCGGGATGC |
| PA14_41230 | clpX | ACACTCGTCGCAGATAAAGACCGAGGGGCC | TTTCGTCCGCGAGTGACCAGGTCCCGGCGTATCAAAAGTCCCGGCGTATCAAAAACACTCGTCGCAGATAAAGACCGAGGGGCCAAGTCCCGGCGTATCAAAAGTCCCGGCGTATCAACAACGTCCATGTCGGGATGC |
| PA14_41230 | clpX | CAGGGGTTGCGACGAGCCATCGATGACACT | TTTCGTCCGCGAGTGACCAGGTCCCGGCGTATCAAAAGTCCCGGCGTATCAAAACAGGGGTTGCGACGAGCCATCGATGACACTAAGTCCCGGCGTATCAAAAGTCCCGGCGTATCAACAACGTCCATGTCGGGATGC |
| PA14_41230 | clpX | GGTTGTCCGATTTCCGGGAAATCTTGTCGA | TTTCGTCCGCGAGTGACCAGGTCCCGGCGTATCAAAAGTCCCGGCGTATCAAAAGGTTGTCCGATTTCCGGGAAATCTTGTCGAAAGTCCCGGCGTATCAAAAGTCCCGGCGTATCAACAACGTCCATGTCGGGATGC |
| PA14_41230 | clpX | ATCGAGGATGGTGCGGATCTCTTTCGGCGC | TTTCGTCCGCGAGTGACCAGGTCCCGGCGTATCAAAAGTCCCGGCGTATCAAAAATCGAGGATGGTGCGGATCTCTTTCGGCGCAAGTCCCGGCGTATCAAAAGTCCCGGCGTATCAACAACGTCCATGTCGGGATGC |
| PA14_41230 | clpX | GAGGGAATCTCGTACATGGTATCGAGAAGG | TTTCGTCCGCGAGTGACCAGGTCCCGGCGTATCAAAAGTCCCGGCGTATCAAAAGAGGGAATCTCGTACATGGTATCGAGAAGGAAGTCCCGGCGTATCAAAAGTCCCGGCGTATCAACAACGTCCATGTCGGGATGC |
| PA14_41230 | clpX | GGCAAACGCCCTACAAACTCGGGGATAAGA | TTTCGTCCGCGAGTGACCAGGTCCCGGCGTATCAAAAGTCCCGGCGTATCAAAAGGCAAACGCCCTACAAACTCGGGGATAAGAAAGTCCCGGCGTATCAAAAGTCCCGGCGTATCAACAACGTCCATGTCGGGATGC |
| PA14_41230 | clpX | TACAGCAGCTTGCCGTTGTCCTCGCCGTTG | TTTCGTCCGCGAGTGACCAGGTCCCGGCGTATCAAAAGTCCCGGCGTATCAAAATACAGCAGCTTGCCGTTGTCCTCGCCGTTGAAGTCCCGGCGTATCAAAAGTCCCGGCGTATCAACAACGTCCATGTCGGGATGC |
| PA14_01300 | coxA | TGTGGTTGGTCGTCAGCACCCAGCGCATCA | TTTCGTCCGCGAGTGACCAGCACTCGTACAGCGTTAACACTCGTACAGCGTTAATGTGGTTGGTCGTCAGCACCCAGCGCATCAAACACTCGTACAGCGTTAACACTCGTACAGCGTTCAACGTCCATGTCGGGATGC |
| PA14_01300 | coxA | ATGAACAGGGTGCTGACCAGCAGGCCGAAG | TTTCGTCCGCGAGTGACCAGCACTCGTACAGCGTTAACACTCGTACAGCGTTAAATGAACAGGGTGCTGACCAGCAGGCCGAAGAACACTCGTACAGCGTTAACACTCGTACAGCGTTCAACGTCCATGTCGGGATGC |
| PA14_01300 | coxA | AGAAGCTGAAGTTGTTCATCCGCGGCAGGG | TTTCGTCCGCGAGTGACCAGCACTCGTACAGCGTTAACACTCGTACAGCGTTAAAGAAGCTGAAGTTGTTCATCCGCGGCAGGGAACACTCGTACAGCGTTAACACTCGTACAGCGTTCAACGTCCATGTCGGGATGC |
| PA14_01300 | coxA | ACGAACAGCGGCATCTTCATCAGGGTCATG | TTTCGTCCGCGAGTGACCAGCACTCGTACAGCGTTAACACTCGTACAGCGTTAAACGAACAGCGGCATCTTCATCAGGGTCATGAACACTCGTACAGCGTTAACACTCGTACAGCGTTCAACGTCCATGTCGGGATGC |
| PA14_01300 | coxA | GGATGTCCATCAGCATCATGGTCACCACGC | TTTCGTCCGCGAGTGACCAGCACTCGTACAGCGTTAACACTCGTACAGCGTTAAGGATGTCCATCAGCATCATGGTCACCACGCAACACTCGTACAGCGTTAACACTCGTACAGCGTTCAACGTCCATGTCGGGATGC |
| PA14_01300 | coxA | CATGATCGAGCTGATCCCGGCCAGGTGGAT | TTTCGTCCGCGAGTGACCAGCACTCGTACAGCGTTAACACTCGTACAGCGTTAACATGATCGAGCTGATCCCGGCCAGGTGGATAACACTCGTACAGCGTTAACACTCGTACAGCGTTCAACGTCCATGTCGGGATGC |
| PA14_01300 | coxA | GCTGAACCACAGGTAGAGGGTGCCGATGTC | TTTCGTCCGCGAGTGACCAGCACTCGTACAGCGTTAACACTCGTACAGCGTTAAGCTGAACCACAGGTAGAGGGTGCCGATGTCAACACTCGTACAGCGTTAACACTCGTACAGCGTTCAACGTCCATGTCGGGATGC |
| PA14_01300 | coxA | TTGAAGAACGCCGGCTCGACGATCTGCAGG | TTTCGTCCGCGAGTGACCAGCACTCGTACAGCGTTAACACTCGTACAGCGTTAATTGAAGAACGCCGGCTCGACGATCTGCAGGAACACTCGTACAGCGTTAACACTCGTACAGCGTTCAACGTCCATGTCGGGATGC |
| PA14_01300 | coxA | CGGATGCACTTGATGACGATGAACAGGAAC | TTTCGTCCGCGAGTGACCAGCACTCGTACAGCGTTAACACTCGTACAGCGTTAACGGATGCACTTGATGACGATGAACAGGAACAACACTCGTACAGCGTTAACACTCGTACAGCGTTCAACGTCCATGTCGGGATGC |
| PA14_01300 | coxA | TCTCGTCGTACATGTGGCCGGTCCACTTCG | TTTCGTCCGCGAGTGACCAGCACTCGTACAGCGTTAACACTCGTACAGCGTTAATCTCGTCGTACATGTGGCCGGTCCACTTCGAACACTCGTACAGCGTTAACACTCGTACAGCGTTCAACGTCCATGTCGGGATGC |
| PA14_01300 | coxA | GTGGTCGGGCGTATCGATCACAGCACTCAT | TTTCGTCCGCGAGTGACCAGCACTCGTACAGCGTTAACACTCGTACAGCGTTAAGTGGTCGGGCGTATCGATCACAGCACTCATAACACTCGTACAGCGTTAACACTCGTACAGCGTTCAACGTCCATGTCGGGATGC |
| PA14_01300 | coxA | ATGAAGAAGGTCACGCTGTGCGGGGCGAAG | TTTCGTCCGCGAGTGACCAGCACTCGTACAGCGTTAACACTCGTACAGCGTTAAATGAAGAAGGTCACGCTGTGCGGGGCGAAGAACACTCGTACAGCGTTAACACTCGTACAGCGTTCAACGTCCATGTCGGGATGC |
| PA14_01300 | coxA | TTGAGGATGGTGGCGATCACGTTGATCGCG | TTTCGTCCGCGAGTGACCAGCACTCGTACAGCGTTAACACTCGTACAGCGTTAATTGAGGATGGTGGCGATCACGTTGATCGCGAACACTCGTACAGCGTTAACACTCGTACAGCGTTCAACGTCCATGTCGGGATGC |
| PA14_01300 | coxA | CCATGATCAGGCCGTGCATGGTGGTCATCT | TTTCGTCCGCGAGTGACCAGCACTCGTACAGCGTTAACACTCGTACAGCGTTAACCATGATCAGGCCGTGCATGGTGGTCATCTAACACTCGTACAGCGTTAACACTCGTACAGCGTTCAACGTCCATGTCGGGATGC |
| PA14_01300 | coxA | AGTAGGTGTCGTGGTACTGGAAGTCCGCCG | TTTCGTCCGCGAGTGACCAGCACTCGTACAGCGTTAACACTCGTACAGCGTTAAAGTAGGTGTCGTGGTACTGGAAGTCCGCCGAACACTCGTACAGCGTTAACACTCGTACAGCGTTCAACGTCCATGTCGGGATGC |

| | | | |
|---|---|---|---|
| PA14_01300 | coxA | AGGCGGGCAGGATCATGATGTACACCTCGG | TTTCGTCCGCGAGTGACCAGCACTCGTACAGCGTTAACACTCGTACAGCGTTAAAGGCGGGCAGGATCATGATGTACACCTCGGAACACTCGTACAGCGTTAACACTCGTACAGCGTTCAACGTCCATGTCGGGATGC |
| PA14_01300 | coxA | ACCGCGATCAACAGGAACGCGGTGATCAGC | TTTCGTCCGCGAGTGACCAGCACTCGTACAGCGTTAACACTCGTACAGCGTTAAACCGCGATCAACAGGAACGCGGTGATCAGCAACACTCGTACAGCGTTAACACTCGTACAGCGTTCAACGTCCATGTCGGGATGC |
| PA14_01300 | coxA | TGGCGTAGACCATCGAGGTGTAGCCGAACA | TTTCGTCCGCGAGTGACCAGCACTCGTACAGCGTTAACACTCGTACAGCGTTAATGGCGTAGACCATCGAGGTGTAGCCGAACAAACACTCGTACAGCGTTAACACTCGTACAGCGTTCAACGTCCATGTCGGGATGC |
| PA14_01300 | coxA | ATCATCAGCGGGATCATCCAGTTGGCCAGG | TTTCGTCCGCGAGTGACCAGCACTCGTACAGCGTTAACACTCGTACAGCGTTAAATCATCAGCGGGATCATCCAGTTGGCCAGGAACACTCGTACAGCGTTAACACTCGTACAGCGTTCAACGTCCATGTCGGGATGC |
| PA14_01300 | coxA | AAGAACCAGAACACGTGCTGGAACAGCACC | TTTCGTCCGCGAGTGACCAGCACTCGTACAGCGTTAACACTCGTACAGCGTTAAAAGAACCAGAACACGTGCTGGAACAGCACCAACACTCGTACAGCGTTAACACTCGTACAGCGTTCAACGTCCATGTCGGGATGC |
| PA14_03650 | cysA | ACGTCCTCGCCATGGAACACGATGTTGCCG | TTTCGTCCGCGAGTGACCAGGACTGATTGGATGCAAAGACTGATTGGATGCAAAACGTCCTCGCCATGGAACACGATGTTGCCGAAGACTGATTGGATGCAAAGACTGATTGGATGCACAACGTCCATGTCGGGATGC |
| PA14_03650 | cysA | TCTGGATGTCGAGGTTGATGTCGTCCAGGG | TTTCGTCCGCGAGTGACCAGGACTGATTGGATGCAAAGACTGATTGGATGCAAATCTGGATGTCGAGGTTGATGTCGTCCAGGGAAGACTGATTGGATGCAAAGACTGATTGGATGCACAACGTCCATGTCGGGATGC |
| PA14_03650 | cysA | TTGCTCTGGATAGCGGTCGGCGAGCCAGTC | TTTCGTCCGCGAGTGACCAGGACTGATTGGATGCAAAGACTGATTGGATGCAAATTGCTCTGGATAGCGGTCGGCGAGCCAGTCAAGACTGATTGGATGCAAAGACTGATTGGATGCACAACGTCCATGTCGGGATGC |
| PA14_03650 | cysA | TGTCGAACACCGTCATGTGGCGGAACAGCG | TTTCGTCCGCGAGTGACCAGGACTGATTGGATGCAAAGACTGATTGGATGCAAATGTCGAACACCGTCATGTGGCGGAACAGCGAAGACTGATTGGATGCAAAGACTGATTGGATGCACAACGTCCATGTCGGGATGC |
| PA14_03650 | cysA | GGCGTTGAAGTTCTTGCTGACGTTACGGAT | TTTCGTCCGCGAGTGACCAGGACTGATTGGATGCAAAGACTGATTGGATGCAAAGGCGTTGAAGTTCTTGCTGACGTTACGGATAAGACTGATTGGATGCAAAGACTGATTGGATGCACAACGTCCATGTCGGGATGC |
| PA14_03650 | cysA | AGGGTTCGTCGAGGAGCAGGATCTTCGGCT | TTTCGTCCGCGAGTGACCAGGACTGATTGGATGCAAAGACTGATTGGATGCAAAAGGGTTCGTCGAGGAGCAGGATCTTCGGCTAAGACTGATTGGATGCAAAGACTGATTGGATGCACAACGTCCATGTCGGGATGC |
| PA14_03650 | cysA | GGCGGTTGGAGTCGCCGAGGAAGTGGTAGA | TTTCGTCCGCGAGTGACCAGGACTGATTGGATGCAAAGACTGATTGGATGCAAAGGCGGTTGGAGTCGCCGAGGAAGTGGTAGAAAGACTGATTGGATGCAAAGACTGATTGGATGCACAACGTCCATGTCGGGATGC |
| PA14_03650 | cysA | AGGCCTTCGGCTTGAAGTAGAGGGTCTCGC | TTTCGTCCGCGAGTGACCAGGACTGATTGGATGCAAAGACTGATTGGATGCAAAAGGCCTTCGGCTTGAAGTAGAGGGTCTCGCAAGACTGATTGGATGCAAAGACTGATTGGATGCACAACGTCCATGTCGGGATGC |
| PA14_03650 | cysA | TTCGTCCTGGCCGTCGACCTTCAGCGTTAC | TTTCGTCCGCGAGTGACCAGGACTGATTGGATGCAAAGACTGATTGGATGCAAATTCGTCCTGGCCGTCGACCTTCAGCGTTACAAGACTGATTGGATGCAAAGACTGATTGGATGCACAACGTCCATGTCGGGATGC |
| PA14_03650 | cysA | AAATCGCTGGCCGGGTTCTCGTAGACCTCG | TTTCGTCCGCGAGTGACCAGGACTGATTGGATGCAAAGACTGATTGGATGCAAAAAATCGCTGGCCGGGTTCTCGTAGACCTCGAAGACTGATTGGATGCAAAGACTGATTGGATGCACAACGTCCATGTCGGGATGC |
| PA14_03650 | cysA | TTGAGCAGCTCGTGGACCTTGGCCTTGATC | TTTCGTCCGCGAGTGACCAGGACTGATTGGATGCAAAGACTGATTGGATGCAAATTGAGCAGCTCGTGGACCTTGGCCTTGATCAAGACTGATTGGATGCAAAGACTGATTGGATGCACAACGTCCATGTCGGGATGC |
| PA14_03650 | cysA | TCGTGGTCCTTCACCACTTCGGCTTCGATC | TTTCGTCCGCGAGTGACCAGGACTGATTGGATGCAAAGACTGATTGGATGCAAATCGTGGTCCTTCACCACTTCGGCTTCGATCAAGACTGATTGGATGCAAAGACTGATTGGATGCACAACGTCCATGTCGGGATGC |
| PA14_03650 | cysA | ATCACCACGATGCGGTCGGCCACTTCCATC | TTTCGTCCGCGAGTGACCAGGACTGATTGGATGCAAAGACTGATTGGATGCAAAATCACCACGATGCGGTCGGCCACTTCCATCAAGACTGATTGGATGCAAAGACTGATTGGATGCACAACGTCCATGTCGGGATGC |
| PA14_03650 | cysA | AGTGCTGGAAGACGAAGCCGACGTTGCGGT | TTTCGTCCGCGAGTGACCAGGACTGATTGGATGCAAAGACTGATTGGATGCAAAAGTGCTGGAAGACGAAGCCGACGTTGCGGTAAGACTGATTGGATGCAAAGACTGATTGGATGCACAACGTCCATGTCGGGATGC |
| PA14_03650 | cysA | ACCGAGGTCAGGTTGATCTCCTCGTGCAGG | TTTCGTCCGCGAGTGACCAGGACTGATTGGATGCAAAGACTGATTGGATGCAAAACCGAGGTCAGGTTGATCTCCTCGTGCAGGAAGACTGATTGGATGCAAAGACTGATTGGATGCACAACGTCCATGTCGGGATGC |
| PA14_06870 | dnr | AGAATCTTCTCCTGGCCTTCCGGGGTCAGC | TTTCGTCCGCGAGTGACCAGGGCTAGCAGTCCGTCAAGGCTAGCAGTCCGTCAAAGAATCTTCTCCTGGCCTTCCGGGGTCAGCAAGGCTAGCAGTCCGTCAAGGCTAGCAGTCCGTCCAACGTCCATGTCGGGATGC |
| PA14_06870 | dnr | TTGTTGGAGAAACGGAACAGCTGGCTGGGC | TTTCGTCCGCGAGTGACCAGGGCTAGCAGTCCGTCAAGGCTAGCAGTCCGTCAATTGTTGGAGAAACGGAACAGCTGGCTGGGCAAGGCTAGCAGTCCGTCAAGGCTAGCAGTCCGTCCAACGTCCATGTCGGGATGC |
| PA14_06870 | dnr | AATCAGGTAGTAGAAGGCGTGCGCCGGCTC | TTTCGTCCGCGAGTGACCAGGGCTAGCAGTCCGTCAAGGCTAGCAGTCCGTCAAAATCAGGTAGTAGAAGGCGTGCGCCGGCTCAAGGCTAGCAGTCCGTCAAGGCTAGCAGTCCGTCCAACGTCCATGTCGGGATGC |
| PA14_06870 | dnr | AAGGTTTCCGGCTGGATCGACAGGTGACCG | TTTCGTCCGCGAGTGACCAGGGCTAGCAGTCCGTCAAGGCTAGCAGTCCGTCAAAAGGTTTCCGGCTGGATCGACAGGTGACCGAAGGCTAGCAGTCCGTCAAGGCTAGCAGTCCGTCCAACGTCCATGTCGGGATGC |
| PA14_06870 | dnr | ACACGTAGGCGCCCTTGTCGAGGTTGACCA | TTTCGTCCGCGAGTGACCAGGGCTAGCAGTCCGTCAAGGCTAGCAGTCCGTCAAACACGTAGGCGCCCTTGTCGAGGTTGACCAAAGGCTAGCAGTCCGTCAAGGCTAGCAGTCCGTCCAACGTCCATGTCGGGATGC |
| PA14_06870 | dnr | AAGCGTCTCGATCTCGTCGATGCGCTGGTG | TTTCGTCCGCGAGTGACCAGGGCTAGCAGTCCGTCAAGGCTAGCAGTCCGTCAAAAGCGTCTCGATCTCGTCGATGCGCTGGTGAAGGCTAGCAGTCCGTCAAGGCTAGCAGTCCGTCCAACGTCCATGTCGGGATGC |
| PA14_06870 | dnr | AGCAGTTCCTGGAGCTGGACGGGAGAAAGC | TTTCGTCCGCGAGTGACCAGGGCTAGCAGTCCGTCAAGGCTAGCAGTCCGTCAAAGCAGTTCCTGGAGCTGGACGGGAGAAAGCAAGGCTAGCAGTCCGTCAAGGCTAGCAGTCCGTCCAACGTCCATGTCGGGATGC |
| PA14_06870 | dnr | CGAACAGGTGGTGGCTTTGCAGCAACTGCT | TTTCGTCCGCGAGTGACCAGGGCTAGCAGTCCGTCAAGGCTAGCAGTCCGTCAACGAACAGGTGGTGGCTTTGCAGCAACTGCTAAGGCTAGCAGTCCGTCAAGGCTAGCAGTCCGTCCAACGTCCATGTCGGGATGC |
| PA14_06870 | dnr | CATGAACATCATCGCCTCGGCGAAGGTGTT | TTTCGTCCGCGAGTGACCAGGGCTAGCAGTCCGTCAAGGCTAGCAGTCCGTCAACATGAACATCATCGCCTCGGCGAAGGTGTTAAGGCTAGCAGTCCGTCAAGGCTAGCAGTCCGTCCAACGTCCATGTCGGGATGC |
| PA14_06870 | dnr | CAACGGGGTGTTGTCCTGCAACTGGCGCAG | TTTCGTCCGCGAGTGACCAGGGCTAGCAGTCCGTCAAGGCTAGCAGTCCGTCAACAACGGGGTGTTGTCCTGCAACTGGCGCAGAAGGCTAGCAGTCCGTCAAGGCTAGCAGTCCGTCCAACGTCCATGTCGGGATGC |
| PA14_06870 | dnr | AGGCGTTCGCGGTCGAGGATGCTGATTTCG | TTTCGTCCGCGAGTGACCAGGGCTAGCAGTCCGTCAAGGCTAGCAGTCCGTCAAAGGCGTTCGCGGTCGAGGATGCTGATTTCGAAGGCTAGCAGTCCGTCAAGGCTAGCAGTCCGTCCAACGTCCATGTCGGGATGC |
| PA14_06870 | dnr | ACCAGTTGCTTGGCCACCGGGATTTCCACC | TTTCGTCCGCGAGTGACCAGGGCTAGCAGTCCGTCAAGGCTAGCAGTCCGTCAAACCAGTTGCTTGGCCACCGGGATTTCCACCAAGGCTAGCAGTCCGTCAAGGCTAGCAGTCCGTCCAACGTCCATGTCGGGATGC |
| PA14_00560 | exoT | GCGTGTGATCCTTCGCCAGTCTCTCCTCTG | TTTCGTCCGCGAGTGACCAGGCGCAATAAACCCTAAAGCGCAATAAACCCTAAAGCGTGTGATCCTTCGCCAGTCTCTCCTCTGAAGCGCAATAAACCCTAAAGCGCAATAAACCCTACAACGTCCATGTCGGGATGC |
| PA14_00560 | exoT | AAAGTCGCCAGCTCGCTCGCCTTGCCAAGT | TTTCGTCCGCGAGTGACCAGGCGCAATAAACCCTAAAGCGCAATAAACCCTAAAAAAGTCGCCAGCTCGCTCGCCTTGCCAAGTAAGCGCAATAAACCCTAAAGCGCAATAAACCCTACAACGTCCATGTCGGGATGC |
| PA14_00560 | exoT | TCGATCGATATCTCGCTGACATCGATCCCG | TTTCGTCCGCGAGTGACCAGGCGCAATAAACCCTAAAGCGCAATAAACCCTAAATCGATCGATATCTCGCTGACATCGATCCCGAAGCGCAATAAACCCTAAAGCGCAATAAACCCTACAACGTCCATGTCGGGATGC |
| PA14_00560 | exoT | GAAGGTCTTCACGACCTGTTCAGCCGGTCC | TTTCGTCCGCGAGTGACCAGGCGCAATAAACCCTAAAGCGCAATAAACCCTAAAGAAGGTCTTCACGACCTGTTCAGCCGGTCCAAGCGCAATAAACCCTAAAGCGCAATAAACCCTACAACGTCCATGTCGGGATGC |
| PA14_00560 | exoT | TCTGCCGAACAGGGTGGTTATCGTGCCCTG | TTTCGTCCGCGAGTGACCAGGCGCAATAAACCCTAAAGCGCAATAAACCCTAAATCTGCCGAACAGGGTGGTTATCGTGCCCTGAAGCGCAATAAACCCTAAAGCGCAATAAACCCTACAACGTCCATGTCGGGATGC |
| PA14_00560 | exoT | ATCTGCTTGATCTCGGCGGCCGAGAGACTG | TTTCGTCCGCGAGTGACCAGGCGCAATAAACCCTAAAGCGCAATAAACCCTAAAATCTGCTTGATCTCGGCGGCCGAGAGACTGAAGCGCAATAAACCCTAAAGCGCAATAAACCCTACAACGTCCATGTCGGGATGC |
| PA14_00560 | exoT | CCAAGGTCAGGGGCAGTGCCTTTTGCAGCA | TTTCGTCCGCGAGTGACCAGGCGCAATAAACCCTAAAGCGCAATAAACCCTAAACCAAGGTCAGGGGCAGTGCCTTTTGCAGCAAAGCGCAATAAACCCTAAAGCGCAATAAACCCTACAACGTCCATGTCGGGATGC |
| PA14_00560 | exoT | TTGCTCCGGGCTTGCGCTGAGTACATGCTG | TTTCGTCCGCGAGTGACCAGGCGCAATAAACCCTAAAGCGCAATAAACCCTAAATTGCTCCGGGCTTGCGCTGAGTACATGCTGAAGCGCAATAAACCCTAAAGCGCAATAAACCCTACAACGTCCATGTCGGGATGC |
| PA14_00560 | exoT | GACGCAGGGAACGATTCAGGTGCTGGTACT | TTTCGTCCGCGAGTGACCAGGCGCAATAAACCCTAAAGCGCAATAAACCCTAAAGACGCAGGGAACGATTCAGGTGCTGGTACTAAGCGCAATAAACCCTAAAGCGCAATAAACCCTACAACGTCCATGTCGGGATGC |
| PA14_00560 | exoT | TCTTTTCGAACGCGGCAGACATGCCCCGGT | TTTCGTCCGCGAGTGACCAGGCGCAATAAACCCTAAAGCGCAATAAACCCTAAATCTTTTCGAACGCGGCAGACATGCCCCGGTAAGCGCAATAAACCCTAAAGCGCAATAAACCCTACAACGTCCATGTCGGGATGC |
| PA14_00560 | exoT | TAGGGCTTGCCTGAACGATCCGTCCCGGTT | TTTCGTCCGCGAGTGACCAGGCGCAATAAACCCTAAAGCGCAATAAACCCTAAATAGGGCTTGCCTGAACGATCCGTCCCGGTTAAGCGCAATAAACCCTAAAGCGCAATAAACCCTACAACGTCCATGTCGGGATGC |
| PA14_00560 | exoT | TTGCCCAGCCACTCGATGATCGCCACGAAG | TTTCGTCCGCGAGTGACCAGGCGCAATAAACCCTAAAGCGCAATAAACCCTAAATTGCCCAGCCACTCGATGATCGCCACGAAGAAGCGCAATAAACCCTAAAGCGCAATAAACCCTACAACGTCCATGTCGGGATGC |
| PA14_00560 | exoT | ATCCTTGGCGCTGAGCAGTACGCGCATGTC | TTTCGTCCGCGAGTGACCAGGCGCAATAAACCCTAAAGCGCAATAAACCCTAAAATCCTTGGCGCTGAGCAGTACGCGCATGTCAAGCGCAATAAACCCTAAAGCGCAATAAACCCTACAACGTCCATGTCGGGATGC |
| PA14_00560 | exoT | CACGAAAGACGGGTTCTGCTGAGATGATTG | TTTCGTCCGCGAGTGACCAGGCGCAATAAACCCTAAAGCGCAATAAACCCTAAACACGAAAGACGGGTTCTGCTGAGATGATTGAAGCGCAATAAACCCTAAAGCGCAATAAACCCTACAACGTCCATGTCGGGATGC |
| PA14_00560 | exoT | TTTCACCGCCTCGAAGGCATCCCTGCCCTG | TTTCGTCCGCGAGTGACCAGGCGCAATAAACCCTAAAGCGCAATAAACCCTAAATTTCACCGCCTCGAAGGCATCCCTGCCCTGAAGCGCAATAAACCCTAAAGCGCAATAAACCCTACAACGTCCATGTCGGGATGC |
| PA14_00560 | exoT | ATAGAGACCGAGCGCCATCACCTCGGCATC | TTTCGTCCGCGAGTGACCAGGCGCAATAAACCCTAAAGCGCAATAAACCCTAAAATAGAGACCGAGCGCCATCACCTCGGCATCAAGCGCAATAAACCCTAAAGCGCAATAAACCCTACAACGTCCATGTCGGGATGC |
| PA14_00560 | exoT | CCTTGTCGTAGAGGATCTCCTGCTCATCGC | TTTCGTCCGCGAGTGACCAGGCGCAATAAACCCTAAAGCGCAATAAACCCTAAACCTTGTCGTAGAGGATCTCCTGCTCATCGCAAGCGCAATAAACCCTAAAGCGCAATAAACCCTACAACGTCCATGTCGGGATGC |
| PA14_00560 | exoT | TTACCTCGCTCTCTACCGCGTGGCGCAACA | TTTCGTCCGCGAGTGACCAGGCGCAATAAACCCTAAAGCGCAATAAACCCTAAATTACCTCGCTCTCTACCGCGTGGCGCAACAAAGCGCAATAAACCCTAAAGCGCAATAAACCCTACAACGTCCATGTCGGGATGC |
| PA14_00560 | exoT | TACCGCATGCAGTTGGACGGCGATTTCGCG | TTTCGTCCGCGAGTGACCAGGCGCAATAAACCCTAAAGCGCAATAAACCCTAAATACCGCATGCAGTTGGACGGCGATTTCGCGAAGCGCAATAAACCCTAAAGCGCAATAAACCCTACAACGTCCATGTCGGGATGC |
| PA14_36345 | exoY | GCTTTGCCAACGACCTTCGCGCATCAGACC | TTTCGTCCGCGAGTGACCAGGTCCTAGAGGTAGTTAAGTCCTAGAGGTAGTTAAGCTTTGCCAACGACCTTCGCGCATCAGACCAAGTCCTAGAGGTAGTTAAGTCCTAGAGGTAGTTCAACGTCCATGTCGGGATGC |
| PA14_36345 | exoY | GTTTCCCCTGGCCATATCCTCCCTGCCATA | TTTCGTCCGCGAGTGACCAGGTCCTAGAGGTAGTTAAGTCCTAGAGGTAGTTAAGTTTCCCCTGGCCATATCCTCCCTGCCATAAAGTCCTAGAGGTAGTTAAGTCCTAGAGGTAGTTCAACGTCCATGTCGGGATGC |
| PA14_36345 | exoY | ATACGCAGGTCGGTCTGGGTATAGGCTCCG | TTTCGTCCGCGAGTGACCAGGTCCTAGAGGTAGTTAAGTCCTAGAGGTAGTTAAATACGCAGGTCGGTCTGGGTATAGGCTCCGAAGTCCTAGAGGTAGTTAAGTCCTAGAGGTAGTTCAACGTCCATGTCGGGATGC |
| PA14_36345 | exoY | TATTCCGGCAAGCGGAAGGCCTTGGACAGG | TTTCGTCCGCGAGTGACCAGGTCCTAGAGGTAGTTAAGTCCTAGAGGTAGTTAATATTCCGGCAAGCGGAAGGCCTTGGACAGGAAGTCCTAGAGGTAGTTAAGTCCTAGAGGTAGTTCAACGTCCATGTCGGGATGC |
| PA14_36345 | exoY | AAGCCCTTGGTCGGGAAACCCTCTTCGATC | TTTCGTCCGCGAGTGACCAGGTCCTAGAGGTAGTTAAGTCCTAGAGGTAGTTAAAAGCCCTTGGTCGGGAAACCCTCTTCGATCAAGTCCTAGAGGTAGTTAAGTCCTAGAGGTAGTTCAACGTCCATGTCGGGATGC |
| PA14_36345 | exoY | CGTAGTGACGATTCTTTCAACCGGGCGAAT | TTTCGTCCGCGAGTGACCAGGTCCTAGAGGTAGTTAAGTCCTAGAGGTAGTTAACGTAGTGACGATTCTTTCAACCGGGCGAATAAGTCCTAGAGGTAGTTAAGTCCTAGAGGTAGTTCAACGTCCATGTCGGGATGC |
| PA14_36345 | exoY | ATCCGTCCTCGCTCAGGGGATCCTTTGCAC | TTTCGTCCGCGAGTGACCAGGTCCTAGAGGTAGTTAAGTCCTAGAGGTAGTTAAATCCGTCCTCGCTCAGGGGATCCTTTGCACAAGTCCTAGAGGTAGTTAAGTCCTAGAGGTAGTTCAACGTCCATGTCGGGATGC |
| PA14_36345 | exoY | GCTGTGGTGAAACATCTCCCTGTGCTCCCC | TTTCGTCCGCGAGTGACCAGGTCCTAGAGGTAGTTAAGTCCTAGAGGTAGTTAAGCTGTGGTGAAACATCTCCCTGTGCTCCCCAAGTCCTAGAGGTAGTTAAGTCCTAGAGGTAGTTCAACGTCCATGTCGGGATGC |
| PA14_36345 | exoY | GACCCCAGTTCGAGCTTTTCCCCTTCACCG | TTTCGTCCGCGAGTGACCAGGTCCTAGAGGTAGTTAAGTCCTAGAGGTAGTTAAGACCCCAGTTCGAGCTTTTCCCCTTCACCGAAGTCCTAGAGGTAGTTAAGTCCTAGAGGTAGTTCAACGTCCATGTCGGGATGC |
| PA14_36345 | exoY | AAAAGCTCTTCCGATCCGCCACCACGCAGA | TTTCGTCCGCGAGTGACCAGGTCCTAGAGGTAGTTAAGTCCTAGAGGTAGTTAAAAAAGCTCTTCCGATCCGCCACCACGCAGAAAGTCCTAGAGGTAGTTAAGTCCTAGAGGTAGTTCAACGTCCATGTCGGGATGC |
| PA14_36345 | exoY | ATTTCCGCCGTGTCGCGGTTTTCCCGCTTG | TTTCGTCCGCGAGTGACCAGGTCCTAGAGGTAGTTAAGTCCTAGAGGTAGTTAAATTTCCGCCGTGTCGCGGTTTTCCCGCTTGAAGTCCTAGAGGTAGTTAAGTCCTAGAGGTAGTTCAACGTCCATGTCGGGATGC |
| PA14_36345 | exoY | CAAAGACGGTATTCGTCTGCCTGGCCACCG | TTTCGTCCGCGAGTGACCAGGTCCTAGAGGTAGTTAAGTCCTAGAGGTAGTTAACAAAGACGGTATTCGTCTGCCTGGCCACCGAAGTCCTAGAGGTAGTTAAGTCCTAGAGGTAGTTCAACGTCCATGTCGGGATGC |
| PA14_36345 | exoY | ACGTTCCAGTCGGGATGGGCGGTGAAGTGA | TTTCGTCCGCGAGTGACCAGGTCCTAGAGGTAGTTAAGTCCTAGAGGTAGTTAAACGTTCCAGTCGGGATGGGCGGTGAAGTGAAAGTCCTAGAGGTAGTTAAGTCCTAGAGGTAGTTCAACGTCCATGTCGGGATGC |
| PA14_36345 | exoY | TTCCCTTCGCCACAGCCAGGTTCAGCTTGC | TTTCGTCCGCGAGTGACCAGGTCCTAGAGGTAGTTAAGTCCTAGAGGTAGTTAATTCCCTTCGCCACAGCCAGGTTCAGCTTGCAAGTCCTAGAGGTAGTTAAGTCCTAGAGGTAGTTCAACGTCCATGTCGGGATGC |
| PA14_36345 | exoY | TCATCGAAGCGAACGGACTCTTCTCCGAGC | TTTCGTCCGCGAGTGACCAGGTCCTAGAGGTAGTTAAGTCCTAGAGGTAGTTAATCATCGAAGCGAACGGACTCTTCTCCGAGCAAGTCCTAGAGGTAGTTAAGTCCTAGAGGTAGTTCAACGTCCATGTCGGGATGC |
| PA14_25080 | fadB | ACGCCCTTGACCGATGCATCGGCCTTGATC | TTTCGTCCGCGAGTGACCAGCCCTCGTACGCCTATAACCCTCGTACGCCTATAAACGCCCTTGACCGATGCATCGGCCTTGATCAACCCTCGTACGCCTATAACCCTCGTACGCCTATCAACGTCCATGTCGGGATGC |
| PA14_25080 | fadB | ACCAACGATGAACACGTCCTTTCCGCTGGT | TTTCGTCCGCGAGTGACCAGCCCTCGTACGCCTATAACCCTCGTACGCCTATAAACCAACGATGAACACGTCCTTTCCGCTGGTAACCCTCGTACGCCTATAACCCTCGTACGCCTATCAACGTCCATGTCGGGATGC |
| PA14_25080 | fadB | TCTTCTCGCCACGGATCACTTCCACCAGCG | TTTCGTCCGCGAGTGACCAGCCCTCGTACGCCTATAACCCTCGTACGCCTATAATCTTCTCGCCACGGATCACTTCCACCAGCGAACCCTCGTACGCCTATAACCCTCGTACGCCTATCAACGTCCATGTCGGGATGC |
| PA14_25080 | fadB | CGATCGAATCGATGTAGCGCAGAGCACCAC | TTTCGTCCGCGAGTGACCAGCCCTCGTACGCCTATAACCCTCGTACGCCTATAACGATCGAATCGATGTAGCGCAGAGCACCACAACCCTCGTACGCCTATAACCCTCGTACGCCTATCAACGTCCATGTCGGGATGC |
| PA14_25080 | fadB | CGGTTGAACTTGTTGACGGACTCGCCCTTG | TTTCGTCCGCGAGTGACCAGCCCTCGTACGCCTATAACCCTCGTACGCCTATAACGGTTGAACTTGTTGACGGACTCGCCCTTGAACCCTCGTACGCCTATAACCCTCGTACGCCTATCAACGTCCATGTCGGGATGC |
| PA14_25080 | fadB | TTTCACGCAGCTTCGCGGTCGGGTGGTACA | TTTCGTCCGCGAGTGACCAGCCCTCGTACGCCTATAACCCTCGTACGCCTATAATTTCACGCAGCTTCGCGGTCGGGTGGTACAAACCCTCGTACGCCTATAACCCTCGTACGCCTATCAACGTCCATGTCGGGATGC |
| PA14_25080 | fadB | CCTTGGCGACTTCGTCGTATTTCTTGGCTT | TTTCGTCCGCGAGTGACCAGCCCTCGTACGCCTATAACCCTCGTACGCCTATAACCTTGGCGACTTCGTCGTATTTCTTGGCTTAACCCTCGTACGCCTATAACCCTCGTACGCCTATCAACGTCCATGTCGGGATGC |
| PA14_25080 | fadB | CGAAGGCCATCATCTGCTCGATTGCATTCA | TTTCGTCCGCGAGTGACCAGCCCTCGTACGCCTATAACCCTCGTACGCCTATAACGAAGGCCATCATCTGCTCGATTGCATTCAAACCCTCGTACGCCTATAACCCTCGTACGCCTATCAACGTCCATGTCGGGATGC |
| PA14_25080 | fadB | TCAGGCCCATCTGGATACCCTCTTCGCGGA | TTTCGTCCGCGAGTGACCAGCCCTCGTACGCCTATAACCCTCGTACGCCTATAATCAGGCCCATCTGGATACCCTCTTCGCGGAAACCCTCGTACGCCTATAACCCTCGTACGCCTATCAACGTCCATGTCGGGATGC |
| PA14_25080 | fadB | ATCGAAATTCAACTCGACGATGCCGCCCTC | TTTCGTCCGCGAGTGACCAGCCCTCGTACGCCTATAACCCTCGTACGCCTATAAATCGAAATTCAACTCGACGATGCCGCCCTCAACCCTCGTACGCCTATAACCCTCGTACGCCTATCAACGTCCATGTCGGGATGC |
| PA14_25080 | fadB | AGCCGAACTTCTCCATCACCTTGTCGATGC | TTTCGTCCGCGAGTGACCAGCCCTCGTACGCCTATAACCCTCGTACGCCTATAAAGCCGAACTTCTCCATCACCTTGTCGATGCAACCCTCGTACGCCTATAACCCTCGTACGCCTATCAACGTCCATGTCGGGATGC |
| PA14_25080 | fadB | GAGGCTTAACCGTGATGGCTTTACCTTGGT | TTTCGTCCGCGAGTGACCAGCCCTCGTACGCCTATAACCCTCGTACGCCTATAAGAGGCTTAACCGTGATGGCTTTACCTTGGTAACCCTCGTACGCCTATAACCCTCGTACGCCTATCAACGTCCATGTCGGGATGC |
| PA14_25080 | fadB | TCCTTCATCAGGATCGGGGTGCCCTTGAGC | TTTCGTCCGCGAGTGACCAGCCCTCGTACGCCTATAACCCTCGTACGCCTATAATCCTTCATCAGGATCGGGGTGCCCTTGAGCAACCCTCGTACGCCTATAACCCTCGTACGCCTATCAACGTCCATGTCGGGATGC |
| PA14_25080 | fadB | ATCGACTGCCGCACGCAACTCACTGAGGGT | TTTCGTCCGCGAGTGACCAGCCCTCGTACGCCTATAACCCTCGTACGCCTATAAATCGACTGCCGCACGCAACTCACTGAGGGTAACCCTCGTACGCCTATAACCCTCGTACGCCTATCAACGTCCATGTCGGGATGC |
| PA14_25080 | fadB | TGAGCACTTCGTAGGCCTGCGGATCGGTGA | TTTCGTCCGCGAGTGACCAGCCCTCGTACGCCTATAACCCTCGTACGCCTATAATGAGCACTTCGTAGGCCTGCGGATCGGTGAAACCCTCGTACGCCTATAACCCTCGTACGCCTATCAACGTCCATGTCGGGATGC |
| PA14_50360 | flgK | ATCCAGGACGGTATCGCGATACGCCAGCAG | TTTCGTCCGCGAGTGACCAGAATCCAGACGGGATGAAAATCCAGACGGGATGAAATCCAGGACGGTATCGCGATACGCCAGCAGAAAATCCAGACGGGATGAAAATCCAGACGGGATGCAACGTCCATGTCGGGATGC |
| PA14_50360 | flgK | TGTTCACCTGCGAGGTCAGGGCACCCAACT | TTTCGTCCGCGAGTGACCAGAATCCAGACGGGATGAAAATCCAGACGGGATGAATGTTCACCTGCGAGGTCAGGGCACCCAACTAAAATCCAGACGGGATGAAAATCCAGACGGGATGCAACGTCCATGTCGGGATGC |
| PA14_50360 | flgK | AGCTTGCTGGTATCGGTGATGTTCAGGTTG | TTTCGTCCGCGAGTGACCAGAATCCAGACGGGATGAAAATCCAGACGGGATGAAAGCTTGCTGGTATCGGTGATGTTCAGGTTGAAAATCCAGACGGGATGAAAATCCAGACGGGATGCAACGTCCATGTCGGGATGC |
| PA14_50360 | flgK | AGCGCAAGTTGGCCGAGCTTGTTGTAGGAG | TTTCGTCCGCGAGTGACCAGAATCCAGACGGGATGAAAATCCAGACGGGATGAAAGCGCAAGTTGGCCGAGCTTGTTGTAGGAGAAAATCCAGACGGGATGAAAATCCAGACGGGATGCAACGTCCATGTCGGGATGC |
| PA14_50360 | flgK | TTATCCGCCAGCAGCGAGTTGAGCTGGTCG | TTTCGTCCGCGAGTGACCAGAATCCAGACGGGATGAAAATCCAGACGGGATGAATTATCCGCCAGCAGCGAGTTGAGCTGGTCGAAAATCCAGACGGGATGAAAATCCAGACGGGATGCAACGTCCATGTCGGGATGC |
| PA14_50360 | flgK | CAGGCTGTTCTGCTTGTCCAGTTGGTCGTA | TTTCGTCCGCGAGTGACCAGAATCCAGACGGGATGAAAATCCAGACGGGATGAACAGGCTGTTCTGCTTGTCCAGTTGGTCGTAAAAATCCAGACGGGATGAAAATCCAGACGGGATGCAACGTCCATGTCGGGATGC |
| PA14_50360 | flgK | CCAGTTGCTTGTTGACGGTATCGGCGAAGG | TTTCGTCCGCGAGTGACCAGAATCCAGACGGGATGAAAATCCAGACGGGATGAACCAGTTGCTTGTTGACGGTATCGGCGAAGGAAAATCCAGACGGGATGAAAATCCAGACGGGATGCAACGTCCATGTCGGGATGC |
| PA14_50360 | flgK | ATCGTTGTACTCGGCCACGCTCTGGCTGAG | TTTCGTCCGCGAGTGACCAGAATCCAGACGGGATGAAAATCCAGACGGGATGAAATCGTTGTACTCGGCCACGCTCTGGCTGAGAAAATCCAGACGGGATGAAAATCCAGACGGGATGCAACGTCCATGTCGGGATGC |
| PA14_50360 | flgK | AAGGCGGAAAAGAAACGCTGCATCGCCGGG | TTTCGTCCGCGAGTGACCAGAATCCAGACGGGATGAAAATCCAGACGGGATGAAAAGGCGGAAAAGAAACGCTGCATCGCCGGGAAAATCCAGACGGGATGAAAATCCAGACGGGATGCAACGTCCATGTCGGGATGC |
| PA14_50360 | flgK | AAGGCGTTCTGCAGAACCGCCGGATCGATC | TTTCGTCCGCGAGTGACCAGAATCCAGACGGGATGAAAATCCAGACGGGATGAAAAGGCGTTCTGCAGAACCGCCGGATCGATCAAAATCCAGACGGGATGAAAATCCAGACGGGATGCAACGTCCATGTCGGGATGC |
| PA14_50360 | flgK | TTTGCGTGGAGTCGCCAAGCGTGAGCTGCA | TTTCGTCCGCGAGTGACCAGAATCCAGACGGGATGAAAATCCAGACGGGATGAATTTGCGTGGAGTCGCCAAGCGTGAGCTGCAAAAATCCAGACGGGATGAAAATCCAGACGGGATGCAACGTCCATGTCGGGATGC |
| PA14_50360 | flgK | CTGGCCCAGGAAAGCGCTCAACTCGCTGTT | TTTCGTCCGCGAGTGACCAGAATCCAGACGGGATGAAAATCCAGACGGGATGAACTGGCCCAGGAAAGCGCTCAACTCGCTGTTAAAATCCAGACGGGATGAAAATCCAGACGGGATGCAACGTCCATGTCGGGATGC |
| PA14_50360 | flgK | GGGTATTGAAGGTCTTGCTCAGGCCCTGCG | TTTCGTCCGCGAGTGACCAGAATCCAGACGGGATGAAAATCCAGACGGGATGAAGGGTATTGAAGGTCTTGCTCAGGCCCTGCGAAAATCCAGACGGGATGAAAATCCAGACGGGATGCAACGTCCATGTCGGGATGC |
| PA14_50360 | flgK | CAGCGTCGAAACGGTATTGCCCACCACCAG | TTTCGTCCGCGAGTGACCAGAATCCAGACGGGATGAAAATCCAGACGGGATGAACAGCGTCGAAACGGTATTGCCCACCACCAGAAAATCCAGACGGGATGAAAATCCAGACGGGATGCAACGTCCATGTCGGGATGC |
| PA14_50360 | flgK | TGTTGTGCCCGGTGATGGTCAGCCAGGTCT | TTTCGTCCGCGAGTGACCAGAATCCAGACGGGATGAAAATCCAGACGGGATGAATGTTGTGCCCGGTGATGGTCAGCCAGGTCTAAAATCCAGACGGGATGAAAATCCAGACGGGATGCAACGTCCATGTCGGGATGC |
| PA14_50360 | flgK | TCGTTGTAGGTAGACCCCGTGCTGCCGGTA | TTTCGTCCGCGAGTGACCAGAATCCAGACGGGATGAAAATCCAGACGGGATGAATCGTTGTAGGTAGACCCCGTGCTGCCGGTAAAAATCCAGACGGGATGAAAATCCAGACGGGATGCAACGTCCATGTCGGGATGC |
| PA14_50360 | flgK | ACAGGCTGACGCTGTTGTCGTCCTGGGTGA | TTTCGTCCGCGAGTGACCAGAATCCAGACGGGATGAAAATCCAGACGGGATGAAACAGGCTGACGCTGTTGTCGTCCTGGGTGAAAAATCCAGACGGGATGAAAATCCAGACGGGATGCAACGTCCATGTCGGGATGC |
| PA14_50360 | flgK | GTCCTTGTCGACATAACTCCAACCGGCCGG | TTTCGTCCGCGAGTGACCAGAATCCAGACGGGATGAAAATCCAGACGGGATGAAGTCCTTGTCGACATAACTCCAACCGGCCGGAAAATCCAGACGGGATGAAAATCCAGACGGGATGCAACGTCCATGTCGGGATGC |
| PA14_50360 | flgK | TTGCGCAACTGGCCGGTGAGGAAGTCACTG | TTTCGTCCGCGAGTGACCAGAATCCAGACGGGATGAAAATCCAGACGGGATGAATTGCGCAACTGGCCGGTGAGGAAGTCACTGAAAATCCAGACGGGATGAAAATCCAGACGGGATGCAACGTCCATGTCGGGATGC |
| PA14_50360 | flgK | GGACAGGCCTATCGAGAGTAGGTCGGACAT | TTTCGTCCGCGAGTGACCAGAATCCAGACGGGATGAAAATCCAGACGGGATGAAGGACAGGCCTATCGAGAGTAGGTCGGACATAAAATCCAGACGGGATGAAAATCCAGACGGGATGCAACGTCCATGTCGGGATGC |
| PA14_45630 | fliA | ATGCGGATGCCGGCATAGGTCTCGAAGCTG | TTTCGTCCGCGAGTGACCAGGCACGAGATTCACATAAGCACGAGATTCACATAAATGCGGATGCCGGCATAGGTCTCGAAGCTGAAGCACGAGATTCACATAAGCACGAGATTCACATCAACGTCCATGTCGGGATGC |
| PA14_45630 | fliA | TTGCTCCTGGGAGTTCTGCGCTTGCGCCTT | TTTCGTCCGCGAGTGACCAGGCACGAGATTCACATAAGCACGAGATTCACATAATTGCTCCTGGGAGTTCTGCGCTTGCGCCTTAAGCACGAGATTCACATAAGCACGAGATTCACATCAACGTCCATGTCGGGATGC |
| PA14_45630 | fliA | ATACATACGCACTCCAGAGGCCGCTGTCAT | TTTCGTCCGCGAGTGACCAGGCACGAGATTCACATAAGCACGAGATTCACATAAATACATACGCACTCCAGAGGCCGCTGTCATAAGCACGAGATTCACATAAGCACGAGATTCACATCAACGTCCATGTCGGGATGC |
| PA14_45630 | fliA | TGTCTTCCGGCAATCCATGTTCGCCGTCCT | TTTCGTCCGCGAGTGACCAGGCACGAGATTCACATAAGCACGAGATTCACATAATGTCTTCCGGCAATCCATGTTCGCCGTCCTAAGCACGAGATTCACATAAGCACGAGATTCACATCAACGTCCATGTCGGGATGC |
| PA14_45630 | fliA | GCCGTGGATCGGTTCGTTGTGACTGAGGCT | TTTCGTCCGCGAGTGACCAGGCACGAGATTCACATAAGCACGAGATTCACATAAGCCGTGGATCGGTTCGTTGTGACTGAGGCTAAGCACGAGATTCACATAAGCACGAGATTCACATCAACGTCCATGTCGGGATGC |
| PA14_45630 | fliA | TCGTAATACAACGCCAGCACCAGGCGCTCG | TTTCGTCCGCGAGTGACCAGGCACGAGATTCACATAAGCACGAGATTCACATAATCGTAATACAACGCCAGCACCAGGCGCTCGAAGCACGAGATTCACATAAGCACGAGATTCACATCAACGTCCATGTCGGGATGC |
| PA14_45630 | fliA | ACTGGCTGTGCAACTGGCTGACCCGCGATT | TTTCGTCCGCGAGTGACCAGGCACGAGATTCACATAAGCACGAGATTCACATAAACTGGCTGTGCAACTGGCTGACCCGCGATTAAGCACGAGATTCACATAAGCACGAGATTCACATCAACGTCCATGTCGGGATGC |
| PA14_45630 | fliA | GTCGTATTTCTTCGCGGCCTCGAGCAGGCC | TTTCGTCCGCGAGTGACCAGGCACGAGATTCACATAAGCACGAGATTCACATAAGTCGTATTTCTTCGCGGCCTCGAGCAGGCCAAGCACGAGATTCACATAAGCACGAGATTCACATCAACGTCCATGTCGGGATGC |
| PA14_45630 | fliA | CTCAATTGGAGTTCGGCAGCAACTTCGTGA | TTTCGTCCGCGAGTGACCAGGCACGAGATTCACATAAGCACGAGATTCACATAACTCAATTGGAGTTCGGCAGCAACTTCGTGAAAGCACGAGATTCACATAAGCACGAGATTCACATCAACGTCCATGTCGGGATGC |
| PA14_45630 | fliA | TCATGCCGGCCTGCATCAGGTCCTCCACCT | TTTCGTCCGCGAGTGACCAGGCACGAGATTCACATAAGCACGAGATTCACATAATCATGCCGGCCTGCATCAGGTCCTCCACCTAAGCACGAGATTCACATAAGCACGAGATTCACATCAACGTCCATGTCGGGATGC |
| PA14_45630 | fliA | TGACGCCCAGTACCTCGCCGATCTCCTTGA | TTTCGTCCGCGAGTGACCAGGCACGAGATTCACATAAGCACGAGATTCACATAATGACGCCCAGTACCTCGCCGATCTCCTTGAAAGCACGAGATTCACATAAGCACGAGATTCACATCAACGTCCATGTCGGGATGC |
| PA14_45630 | fliA | AACAGGTCGTCGAAGCTGTACAGGCGGCTG | TTTCGTCCGCGAGTGACCAGGCACGAGATTCACATAAGCACGAGATTCACATAAAACAGGTCGTCGAAGCTGTACAGGCGGCTGAAGCACGAGATTCACATAAGCACGAGATTCACATCAACGTCCATGTCGGGATGC |
| PA14_45630 | fliA | TGTCGCTGAGGATGCCGTAGTAATCTTCGA | TTTCGTCCGCGAGTGACCAGGCACGAGATTCACATAAGCACGAGATTCACATAATGTCGCTGAGGATGCCGTAGTAATCTTCGAAAGCACGAGATTCACATAAGCACGAGATTCACATCAACGTCCATGTCGGGATGC |
| PA14_45630 | fliA | CGCTTGACCAGTGGTGCGTAACGCTGGATC | TTTCGTCCGCGAGTGACCAGGCACGAGATTCACATAAGCACGAGATTCACATAACGCTTGACCAGTGGTGCGTAACGCTGGATCAAGCACGAGATTCACATAAGCACGAGATTCACATCAACGTCCATGTCGGGATGC |
| PA14_45630 | fliA | TTTAGCGTCGCGTCCGGTTCTCGCCTCGAT | TTTCGTCCGCGAGTGACCAGGCACGAGATTCACATAAGCACGAGATTCACATAATTTAGCGTCGCGTCCGGTTCTCGCCTCGATAAGCACGAGATTCACATAAGCACGAGATTCACATCAACGTCCATGTCGGGATGC |
| PA14_61060 | fnr-2 | TGAACTGCAGCTTGTGGGCATGCTCGGCCA | TTTCGTCCGCGAGTGACCAGTCCGCCTGGATCCAAAATCCGCCTGGATCCAAAATGAACTGCAGCTTGTGGGCATGCTCGGCCAAATCCGCCTGGATCCAAAATCCGCCTGGATCCAACAACGTCCATGTCGGGATGC |
| PA14_61060 | fnr-2 | TACGTCCGTTCAGGCAACCGGGTACCTGCT | TTTCGTCCGCGAGTGACCAGTCCGCCTGGATCCAAAATCCGCCTGGATCCAAAATACGTCCGTTCAGGCAACCGGGTACCTGCTAATCCGCCTGGATCCAAAATCCGCCTGGATCCAACAACGTCCATGTCGGGATGC |
| PA14_61060 | fnr-2 | TCATGAGGCGCCGAAACCATCGAGTAGGCG | TTTCGTCCGCGAGTGACCAGTCCGCCTGGATCCAAAATCCGCCTGGATCCAAAATCATGAGGCGCCGAAACCATCGAGTAGGCGAATCCGCCTGGATCCAAAATCCGCCTGGATCCAACAACGTCCATGTCGGGATGC |
| PA14_61060 | fnr-2 | TTCGAAGCGCTCCCAGACCTCGAAGTCCTG | TTTCGTCCGCGAGTGACCAGTCCGCCTGGATCCAAAATCCGCCTGGATCCAAAATTCGAAGCGCTCCCAGACCTCGAAGTCCTGAATCCGCCTGGATCCAAAATCCGCCTGGATCCAACAACGTCCATGTCGGGATGC |
| PA14_61060 | fnr-2 | GGGACGACGATGGAAAAGAAGTCGAGGAAT | TTTCGTCCGCGAGTGACCAGTCCGCCTGGATCCAAAATCCGCCTGGATCCAAAAGGGACGACGATGGAAAAGAAGTCGAGGAATAATCCGCCTGGATCCAAAATCCGCCTGGATCCAACAACGTCCATGTCGGGATGC |
| PA14_61060 | fnr-2 | CTCTCGCGCACGCTGTAGACCAGCTTGATG | TTTCGTCCGCGAGTGACCAGTCCGCCTGGATCCAAAATCCGCCTGGATCCAAAACTCTCGCGCACGCTGTAGACCAGCTTGATGAATCCGCCTGGATCCAAAATCCGCCTGGATCCAACAACGTCCATGTCGGGATGC |
| PA14_61060 | fnr-2 | ACAGGTTGGGAGTCAGAGGCTGCACGTCGA | TTTCGTCCGCGAGTGACCAGTCCGCCTGGATCCAAAATCCGCCTGGATCCAAAAACAGGTTGGGAGTCAGAGGCTGCACGTCGAAATCCGCCTGGATCCAAAATCCGCCTGGATCCAACAACGTCCATGTCGGGATGC |
| PA14_61060 | fnr-2 | TACCAGTAGTTCTCCACCGCCACCTGCCCC | TTTCGTCCGCGAGTGACCAGTCCGCCTGGATCCAAAATCCGCCTGGATCCAAAATACCAGTAGTTCTCCACCGCCACCTGCCCCAATCCGCCTGGATCCAAAATCCGCCTGGATCCAACAACGTCCATGTCGGGATGC |
| PA14_61060 | fnr-2 | ACCATCCACGAAGCGGTCGAGGGTGAGGAA | TTTCGTCCGCGAGTGACCAGTCCGCCTGGATCCAAAATCCGCCTGGATCCAAAAACCATCCACGAAGCGGTCGAGGGTGAGGAAAATCCGCCTGGATCCAAAATCCGCCTGGATCCAACAACGTCCATGTCGGGATGC |
| PA14_61060 | fnr-2 | ATCATCTGCGGGTTGCCGCAGAGCATCACC | TTTCGTCCGCGAGTGACCAGTCCGCCTGGATCCAAAATCCGCCTGGATCCAAAAATCATCTGCGGGTTGCCGCAGAGCATCACCAATCCGCCTGGATCCAAAATCCGCCTGGATCCAACAACGTCCATGTCGGGATGC |
| PA14_61060 | fnr-2 | GCGGGTGAATTTCTCTTCGCTGGCGGTCAT | TTTCGTCCGCGAGTGACCAGTCCGCCTGGATCCAAAATCCGCCTGGATCCAAAAGCGGGTGAATTTCTCTTCGCTGGCGGTCATAATCCGCCTGGATCCAAAATCCGCCTGGATCCAACAACGTCCATGTCGGGATGC |
| PA14_61060 | fnr-2 | ACTGAGGTTCATGCCGCGAGCCTTCAGCAC | TTTCGTCCGCGAGTGACCAGTCCGCCTGGATCCAAAATCCGCCTGGATCCAAAAACTGAGGTTCATGCCGCGAGCCTTCAGCACAATCCGCCTGGATCCAAAATCCGCCTGGATCCAACAACGTCCATGTCGGGATGC |
| PA14_61060 | fnr-2 | AGACGATACTGCCGCTGGGCTTGTACACGC | TTTCGTCCGCGAGTGACCAGTCCGCCTGGATCCAAAATCCGCCTGGATCCAAAAAGACGATACTGCCGCTGGGCTTGTACACGCAATCCGCCTGGATCCAAAATCCGCCTGGATCCAACAACGTCCATGTCGGGATGC |

| | | | |
|---|---|---|---|
| PA14_50290 | fliC | AGGCATTCTGCAGGCTGATGTCAATGGTCT | TTTCGTCCGCGAGTGACCAGAAGCGGACGTCAGTTAAAAGCGGACGTCAGTTAAAGGCATTCTGCAGGCTGATGTCAATGGTCTAAAAGCGGACGTCAGTTAAAAGCGGACGTCAGTTCAACGTCCATGTCGGGATGC |
| PA14_50290 | fliC | TGGTCAGTACACCCTTGTCGTTGATGCTGG | TTTCGTCCGCGAGTGACCAGAAGCGGACGTCAGTTAAAAGCGGACGTCAGTTAATGGTCAGTACACCCTTGTCGTTGATGCTGGAAAAGCGGACGTCAGTTAAAAGCGGACGTCAGTTCAACGTCCATGTCGGGATGC |
| PA14_50290 | fliC | ATTGGTGGACTGCTGCAGGGCACCTTCAGC | TTTCGTCCGCGAGTGACCAGAAGCGGACGTCAGTTAAAAGCGGACGTCAGTTAAATTGGTGGACTGCTGCAGGGCACCTTCAGCAAAAGCGGACGTCAGTTAAAAGCGGACGTCAGTTCAACGTCCATGTCGGGATGC |
| PA14_50290 | fliC | TCAGACGCTGCAACGAGGTGTTGAGGTCGT | TTTCGTCCGCGAGTGACCAGAAGCGGACGTCAGTTAAAAGCGGACGTCAGTTAATCAGACGCTGCAACGAGGTGTTGAGGTCGTAAAAGCGGACGTCAGTTAAAAGCGGACGTCAGTTCAACGTCCATGTCGGGATGC |
| PA14_50290 | fliC | TCGGAACCCTGGACCTTCACTGCGACCTGA | TTTCGTCCGCGAGTGACCAGAAGCGGACGTCAGTTAAAAGCGGACGTCAGTTAATCGGAACCCTGGACCTTCACTGCGACCTGAAAAAGCGGACGTCAGTTAAAAGCGGACGTCAGTTCAACGTCCATGTCGGGATGC |
| PA14_50290 | fliC | ACCACGTTCTTGGCGGCCGCTTCGAACTTG | TTTCGTCCGCGAGTGACCAGAAGCGGACGTCAGTTAAAAGCGGACGTCAGTTAAACCACGTTCTTGGCGGCCGCTTCGAACTTGAAAAGCGGACGTCAGTTAAAAGCGGACGTCAGTTCAACGTCCATGTCGGGATGC |
| PA14_50290 | fliC | ATCGTCCTTGGCGCTGTTGATGCGGTAGCC | TTTCGTCCGCGAGTGACCAGAAGCGGACGTCAGTTAAAAGCGGACGTCAGTTAAATCGTCCTTGGCGCTGTTGATGCGGTAGCCAAAAGCGGACGTCAGTTAAAAGCGGACGTCAGTTCAACGTCCATGTCGGGATGC |
| PA14_50290 | fliC | AGCAACTTCTTTCTGCAGGGCGGCACGGTC | TTTCGTCCGCGAGTGACCAGAAGCGGACGTCAGTTAAAAGCGGACGTCAGTTAAAGCAACTTCTTTCTGCAGGGCGGCACGGTCAAAAGCGGACGTCAGTTAAAAGCGGACGTCAGTTCAACGTCCATGTCGGGATGC |
| PA14_50290 | fliC | AGGTTCGGGATGGCACCGTCCATCTTCTCG | TTTCGTCCGCGAGTGACCAGAAGCGGACGTCAGTTAAAAGCGGACGTCAGTTAAAGGTTCGGGATGGCACCGTCCATCTTCTCGAAAAGCGGACGTCAGTTAAAAGCGGACGTCAGTTCAACGTCCATGTCGGGATGC |
| PA14_50290 | fliC | TCGGAGATACGGGTCAGTTCGGCCTGTTGC | TTTCGTCCGCGAGTGACCAGAAGCGGACGTCAGTTAAAAGCGGACGTCAGTTAATCGGAGATACGGGTCAGTTCGGCCTGTTGCAAAAGCGGACGTCAGTTAAAAGCGGACGTCAGTTCAACGTCCATGTCGGGATGC |
| PA14_50290 | fliC | CCTTGATGCGGCTACGAGCGTTGGTAGCGT | TTTCGTCCGCGAGTGACCAGAAGCGGACGTCAGTTAAAAGCGGACGTCAGTTAACCTTGATGCGGCTACGAGCGTTGGTAGCGTAAAAGCGGACGTCAGTTAAAAGCGGACGTCAGTTCAACGTCCATGTCGGGATGC |
| PA14_50290 | fliC | GAAGCAATGTTCGTGTTGACTGTAAGGGCC | TTTCGTCCGCGAGTGACCAGAAGCGGACGTCAGTTAAAAGCGGACGTCAGTTAAGAAGCAATGTTCGTGTTGACTGTAAGGGCCAAAAGCGGACGTCAGTTAAAAGCGGACGTCAGTTCAACGTCCATGTCGGGATGC |
| PA14_50290 | fliC | TTATCGACTACCGCGATGGCGTTCTGGGCG | TTTCGTCCGCGAGTGACCAGAAGCGGACGTCAGTTAAAAGCGGACGTCAGTTAATTATCGACTACCGCGATGGCGTTCTGGGCGAAAAGCGGACGTCAGTTAAAAGCGGACGTCAGTTCAACGTCCATGTCGGGATGC |
| PA14_50290 | fliC | CGTTCAGACCGCTGATCTGGTTGGACAGGC | TTTCGTCCGCGAGTGACCAGAAGCGGACGTCAGTTAAAAGCGGACGTCAGTTAACGTTCAGACCGCTGATCTGGTTGGACAGGCAAAAGCGGACGTCAGTTAAAAGCGGACGTCAGTTCAACGTCCATGTCGGGATGC |
| PA14_50290 | fliC | TAGGCGTTGGAACCGACCTGGAAGCTGGTG | TTTCGTCCGCGAGTGACCAGAAGCGGACGTCAGTTAAAAGCGGACGTCAGTTAATAGGCGTTGGAACCGACCTGGAAGCTGGTGAAAAGCGGACGTCAGTTAAAAGCGGACGTCAGTTCAACGTCCATGTCGGGATGC |
| PA14_50290 | fliC | ATGTCGACGCTGGCAACGCTGCTCTTCTGC | TTTCGTCCGCGAGTGACCAGAAGCGGACGTCAGTTAAAAGCGGACGTCAGTTAAATGTCGACGCTGGCAACGCTGCTCTTCTGCAAAAGCGGACGTCAGTTAAAAGCGGACGTCAGTTCAACGTCCATGTCGGGATGC |
| PA14_50290 | fliC | ACATCAGCGGTGAACACGGTACGGGCACGA | TTTCGTCCGCGAGTGACCAGAAGCGGACGTCAGTTAAAAGCGGACGTCAGTTAAACATCAGCGGTGAACACGGTACGGGCACGAAAAAGCGGACGTCAGTTAAAAGCGGACGTCAGTTCAACGTCCATGTCGGGATGC |
| PA14_50290 | fliC | GTGATGCCCAGCTTCGACGAGTTGGAGTTC | TTTCGTCCGCGAGTGACCAGAAGCGGACGTCAGTTAAAAGCGGACGTCAGTTAAGTGATGCCCAGCTTCGACGAGTTGGAGTTCAAAAGCGGACGTCAGTTAAAAGCGGACGTCAGTTCAACGTCCATGTCGGGATGC |
| PA14_50290 | fliC | TTGCTGCCGACCTGGTAAGAACCGATGGCG | TTTCGTCCGCGAGTGACCAGAAGCGGACGTCAGTTAAAAGCGGACGTCAGTTAATTGCTGCCGACCTGGTAAGAACCGATGGCGAAAAGCGGACGTCAGTTAAAAGCGGACGTCAGTTCAACGTCCATGTCGGGATGC |
| PA14_50290 | fliC | GAAGAAGCATTCAGGTTGCGCTGAGTGTTC | TTTCGTCCGCGAGTGACCAGAAGCGGACGTCAGTTAAAAGCGGACGTCAGTTAAGAAGAAGCATTCAGGTTGCGCTGAGTGTTCAAAAGCGGACGTCAGTTAAAAGCGGACGTCAGTTCAACGTCCATGTCGGGATGC |
| PA14_62860 | ftsH | CCGTCGCTACGCTTGCCGGTGATGACATAG | TTTCGTCCGCGAGTGACCAGACGGAAGCTCTAATCAAACGGAAGCTCTAATCAACCGTCGCTACGCTTGCCGGTGATGACATAGAAACGGAAGCTCTAATCAAACGGAAGCTCTAATCCAACGTCCATGTCGGGATGC |
| PA14_62860 | ftsH | GCTCGAGAAATTGTTCATCACGGTCACCAG | TTTCGTCCGCGAGTGACCAGACGGAAGCTCTAATCAAACGGAAGCTCTAATCAAGCTCGAGAAATTGTTCATCACGGTCACCAGAAACGGAAGCTCTAATCAAACGGAAGCTCTAATCCAACGTCCATGTCGGGATGC |
| PA14_62860 | ftsH | GAACCGGAAATGGTGAAGAACGGCACCTTG | TTTCGTCCGCGAGTGACCAGACGGAAGCTCTAATCAAACGGAAGCTCTAATCAAGAACCGGAAATGGTGAAGAACGGCACCTTGAAACGGAAGCTCTAATCAAACGGAAGCTCTAATCCAACGTCCATGTCGGGATGC |
| PA14_62860 | ftsH | ACTTGCCCGGATCGCGAAGGAACTCGACCA | TTTCGTCCGCGAGTGACCAGACGGAAGCTCTAATCAAACGGAAGCTCTAATCAAACTTGCCCGGATCGCGAAGGAACTCGACCAAAACGGAAGCTCTAATCAAACGGAAGCTCTAATCCAACGTCCATGTCGGGATGC |
| PA14_62860 | ftsH | ACACCTTGTAGACCGGGTCATGCTCCGGCA | TTTCGTCCGCGAGTGACCAGACGGAAGCTCTAATCAAACGGAAGCTCTAATCAAACACCTTGTAGACCGGGTCATGCTCCGGCAAAACGGAAGCTCTAATCAAACGGAAGCTCTAATCCAACGTCCATGTCGGGATGC |
| PA14_62860 | ftsH | CCCTCGACCACGACGTTGTTGTTGACCAGG | TTTCGTCCGCGAGTGACCAGACGGAAGCTCTAATCAAACGGAAGCTCTAATCAACCCTCGACCACGACGTTGTTGTTGACCAGGAAACGGAAGCTCTAATCAAACGGAAGCTCTAATCCAACGTCCATGTCGGGATGC |
| PA14_62860 | ftsH | AGGTGGTCTTGACCTGGTCCTCGGACAGCA | TTTCGTCCGCGAGTGACCAGACGGAAGCTCTAATCAAACGGAAGCTCTAATCAAAGGTGGTCTTGACCTGGTCCTCGGACAGCAAAACGGAAGCTCTAATCAAACGGAAGCTCTAATCCAACGTCCATGTCGGGATGC |
| PA14_62860 | ftsH | TGGCGGTACCGTAGCAGTCGTCGATGATGC | TTTCGTCCGCGAGTGACCAGACGGAAGCTCTAATCAAACGGAAGCTCTAATCAATGGCGGTACCGTAGCAGTCGTCGATGATGCAAACGGAAGCTCTAATCAAACGGAAGCTCTAATCCAACGTCCATGTCGGGATGC |
| PA14_62860 | ftsH | CATCTGGCGCATGAAGAACATGAAGACGGC | TTTCGTCCGCGAGTGACCAGACGGAAGCTCTAATCAAACGGAAGCTCTAATCAACATCTGGCGCATGAAGAACATGAAGACGGCAAACGGAAGCTCTAATCAAACGGAAGCTCTAATCCAACGTCCATGTCGGGATGC |
| PA14_62860 | ftsH | ACTGGGTCCAGATGCTCTGCTGTTCGGGCT | TTTCGTCCGCGAGTGACCAGACGGAAGCTCTAATCAAACGGAAGCTCTAATCAAACTGGGTCCAGATGCTCTGCTGTTCGGGCTAAACGGAAGCTCTAATCAAACGGAAGCTCTAATCCAACGTCCATGTCGGGATGC |
| PA14_62860 | ftsH | TGATCACCAGGATCGGGAAGCTGGCGACCA | TTTCGTCCGCGAGTGACCAGACGGAAGCTCTAATCAAACGGAAGCTCTAATCAATGATCACCAGGATCGGGAAGCTGGCGACCAAAACGGAAGCTCTAATCAAACGGAAGCTCTAATCCAACGTCCATGTCGGGATGC |
| PA14_62860 | ftsH | TCCGAGTAGTTCAGCGTCTGTGGCTCGCTC | TTTCGTCCGCGAGTGACCAGACGGAAGCTCTAATCAAACGGAAGCTCTAATCAATCCGAGTAGTTCAGCGTCTGTGGCTCGCTCAAACGGAAGCTCTAATCAAACGGAAGCTCTAATCCAACGTCCATGTCGGGATGC |
| PA14_62860 | ftsH | ATCCACGGTCACTCGCTCGACCTTGCCATC | TTTCGTCCGCGAGTGACCAGACGGAAGCTCTAATCAAACGGAAGCTCTAATCAAATCCACGGTCACTCGCTCGACCTTGCCATCAAACGGAAGCTCTAATCAAACGGAAGCTCTAATCCAACGTCCATGTCGGGATGC |
| PA14_62860 | ftsH | CCGCGATGATCAGCCACAGAATCAGATTCT | TTTCGTCCGCGAGTGACCAGACGGAAGCTCTAATCAAACGGAAGCTCTAATCAACCGCGATGATCAGCCACAGAATCAGATTCTAAACGGAAGCTCTAATCAAACGGAAGCTCTAATCCAACGTCCATGTCGGGATGC |
| PA14_62860 | ftsH | GCATGATGTCGTTGGAAGCGCCAGTGGTGA | TTTCGTCCGCGAGTGACCAGACGGAAGCTCTAATCAAACGGAAGCTCTAATCAAGCATGATGTCGTTGGAAGCGCCAGTGGTGAAAACGGAAGCTCTAATCAAACGGAAGCTCTAATCCAACGTCCATGTCGGGATGC |
| PA14_08150 | PA14_08150 | TCAGCTTGAGCAGGCGGTCATTCAGCCGCT | TTTCGTCCGCGAGTGACCAGTAGAATTCGCGGCCAAATAGAATTCGCGGCCAAATCAGCTTGAGCAGGCGGTCATTCAGCCGCTAATAGAATTCGCGGCCAAATAGAATTCGCGGCCACAACGTCCATGTCGGGATGC |
| PA14_08150 | PA14_08150 | TCCACGGTAAAGCGCCCCATGCGCATCAGT | TTTCGTCCGCGAGTGACCAGTAGAATTCGCGGCCAAATAGAATTCGCGGCCAAATCCACGGTAAAGCGCCCCATGCGCATCAGTAATAGAATTCGCGGCCAAATAGAATTCGCGGCCACAACGTCCATGTCGGGATGC |
| PA14_08150 | PA14_08150 | ATTCCCGCCTTGAAGCCTTGGAGCAGCAGG | TTTCGTCCGCGAGTGACCAGTAGAATTCGCGGCCAAATAGAATTCGCGGCCAAAATTCCCGCCTTGAAGCCTTGGAGCAGCAGGAATAGAATTCGCGGCCAAATAGAATTCGCGGCCACAACGTCCATGTCGGGATGC |
| PA14_08150 | PA14_08150 | AACACCAAAGGTTCGATCCGACGCCCCGTG | TTTCGTCCGCGAGTGACCAGTAGAATTCGCGGCCAAATAGAATTCGCGGCCAAAAACACCAAAGGTTCGATCCGACGCCCCGTGAATAGAATTCGCGGCCAAATAGAATTCGCGGCCACAACGTCCATGTCGGGATGC |
| PA14_08150 | PA14_08150 | TGGGTAAGGCCACCTGTCCATCGCGATCGT | TTTCGTCCGCGAGTGACCAGTAGAATTCGCGGCCAAATAGAATTCGCGGCCAAATGGGTAAGGCCACCTGTCCATCGCGATCGTAATAGAATTCGCGGCCAAATAGAATTCGCGGCCACAACGTCCATGTCGGGATGC |
| PA14_08150 | PA14_08150 | CACGACAGTGGTCCATCCGCTGGAGCTGTA | TTTCGTCCGCGAGTGACCAGTAGAATTCGCGGCCAAATAGAATTCGCGGCCAAACACGACAGTGGTCCATCCGCTGGAGCTGTAAATAGAATTCGCGGCCAAATAGAATTCGCGGCCACAACGTCCATGTCGGGATGC |
| PA14_08150 | PA14_08150 | TGTGCTCCACCGAGTCGATCAGGAACTCGC | TTTCGTCCGCGAGTGACCAGTAGAATTCGCGGCCAAATAGAATTCGCGGCCAAATGTGCTCCACCGAGTCGATCAGGAACTCGCAATAGAATTCGCGGCCAAATAGAATTCGCGGCCACAACGTCCATGTCGGGATGC |
| PA14_08150 | PA14_08150 | AGCCTCAAGGTCAGGCTGTCGGACTCCATG | TTTCGTCCGCGAGTGACCAGTAGAATTCGCGGCCAAATAGAATTCGCGGCCAAAAGCCTCAAGGTCAGGCTGTCGGACTCCATGAATAGAATTCGCGGCCAAATAGAATTCGCGGCCACAACGTCCATGTCGGGATGC |
| PA14_08150 | PA14_08150 | AGTCTTCACCGTTCTCATCAGGCTGCGGTC | TTTCGTCCGCGAGTGACCAGTAGAATTCGCGGCCAAATAGAATTCGCGGCCAAAAGTCTTCACCGTTCTCATCAGGCTGCGGTCAATAGAATTCGCGGCCAAATAGAATTCGCGGCCACAACGTCCATGTCGGGATGC |
| PA14_08150 | PA14_08150 | ATAGCCGAGATGAACCTCCAGCACCGCACC | TTTCGTCCGCGAGTGACCAGTAGAATTCGCGGCCAAATAGAATTCGCGGCCAAAATAGCCGAGATGAACCTCCAGCACCGCACCAATAGAATTCGCGGCCAAATAGAATTCGCGGCCACAACGTCCATGTCGGGATGC |
| PA14_08150 | PA14_08150 | TAACGTCGAAGCTGCCGACGTCAGCACGTC | TTTCGTCCGCGAGTGACCAGTAGAATTCGCGGCCAAATAGAATTCGCGGCCAAATAACGTCGAAGCTGCCGACGTCAGCACGTCAATAGAATTCGCGGCCAAATAGAATTCGCGGCCACAACGTCCATGTCGGGATGC |
| PA14_08150 | PA14_08150 | GTCGGCAACGATACGGAAACTCGGTTGCAC | TTTCGTCCGCGAGTGACCAGTAGAATTCGCGGCCAAATAGAATTCGCGGCCAAAGTCGGCAACGATACGGAAACTCGGTTGCACAATAGAATTCGCGGCCAAATAGAATTCGCGGCCACAACGTCCATGTCGGGATGC |
| PA14_08150 | PA14_08150 | TGCGTATCGTCTTGCCACTGCCGCGCATGT | TTTCGTCCGCGAGTGACCAGTAGAATTCGCGGCCAAATAGAATTCGCGGCCAAATGCGTATCGTCTTGCCACTGCCGCGCATGTAATAGAATTCGCGGCCAAATAGAATTCGCGGCCACAACGTCCATGTCGGGATGC |
| PA14_08150 | PA14_08150 | AACATCGCGGCAGACCTGAGCCAGGGTAGT | TTTCGTCCGCGAGTGACCAGTAGAATTCGCGGCCAAATAGAATTCGCGGCCAAAAACATCGCGGCAGACCTGAGCCAGGGTAGTAATAGAATTCGCGGCCAAATAGAATTCGCGGCCACAACGTCCATGTCGGGATGC |
| PA14_08150 | PA14_08150 | TAGTCGGACTCATTGACCTGGTCGACCCGG | TTTCGTCCGCGAGTGACCAGTAGAATTCGCGGCCAAATAGAATTCGCGGCCAAATAGTCGGACTCATTGACCTGGTCGACCCGGAATAGAATTCGCGGCCAAATAGAATTCGCGGCCACAACGTCCATGTCGGGATGC |
| PA14_22980 | gltB | TGAAGTCCTTCTGGAAGGACGGATCGAGCA | TTTCGTCCGCGAGTGACCAGTGATACGCGAACTGTAATGATACGCGAACTGTAATGAAGTCCTTCTGGAAGGACGGATCGAGCAAATGATACGCGAACTGTAATGATACGCGAACTGTCAACGTCCATGTCGGGATGC |
| PA14_22980 | gltB | TCGCCTTGTCCTGTTCGACGAAGGCCTTGC | TTTCGTCCGCGAGTGACCAGTGATACGCGAACTGTAATGATACGCGAACTGTAATCGCCTTGTCCTGTTCGACGAAGGCCTTGCAATGATACGCGAACTGTAATGATACGCGAACTGTCAACGTCCATGTCGGGATGC |
| PA14_22980 | gltB | TCGGCATCGACGTAGCCGCGGAGCTTCTTC | TTTCGTCCGCGAGTGACCAGTGATACGCGAACTGTAATGATACGCGAACTGTAATCGGCATCGACGTAGCCGCGGAGCTTCTTCAATGATACGCGAACTGTAATGATACGCGAACTGTCAACGTCCATGTCGGGATGC |
| PA14_22980 | gltB | ACCTTGCCGGCGGCGGTGAATTCGCTCTTC | TTTCGTCCGCGAGTGACCAGTGATACGCGAACTGTAATGATACGCGAACTGTAAACCTTGCCGGCGGCGGTGAATTCGCTCTTCAATGATACGCGAACTGTAATGATACGCGAACTGTCAACGTCCATGTCGGGATGC |
| PA14_22980 | gltB | TGAGCTTGTCGGCGGCGACGAACAGTTCGT | TTTCGTCCGCGAGTGACCAGTGATACGCGAACTGTAATGATACGCGAACTGTAATGAGCTTGTCGGCGGCGACGAACAGTTCGTAATGATACGCGAACTGTAATGATACGCGAACTGTCAACGTCCATGTCGGGATGC |
| PA14_22980 | gltB | CCGGGTTGATGTAGAGCCAGTTGACCCGGT | TTTCGTCCGCGAGTGACCAGTGATACGCGAACTGTAATGATACGCGAACTGTAACCGGGTTGATGTAGAGCCAGTTGACCCGGTAATGATACGCGAACTGTAATGATACGCGAACTGTCAACGTCCATGTCGGGATGC |
| PA14_22980 | gltB | GACGGCAGAGAGGCGACGGATCGCATTCAT | TTTCGTCCGCGAGTGACCAGTGATACGCGAACTGTAATGATACGCGAACTGTAAGACGGCAGAGAGGCGACGGATCGCATTCATAATGATACGCGAACTGTAATGATACGCGAACTGTCAACGTCCATGTCGGGATGC |
| PA14_22980 | gltB | CCATGATCTGCATCCCGGCCTTGCCGTTGA | TTTCGTCCGCGAGTGACCAGTGATACGCGAACTGTAATGATACGCGAACTGTAACCATGATCTGCATCCCGGCCTTGCCGTTGAAATGATACGCGAACTGTAATGATACGCGAACTGTCAACGTCCATGTCGGGATGC |
| PA14_22980 | gltB | GAACACCTCGACCATCTGCGCTCCGGTGAG | TTTCGTCCGCGAGTGACCAGTGATACGCGAACTGTAATGATACGCGAACTGTAAGAACACCTCGACCATCTGCGCTCCGGTGAGAATGATACGCGAACTGTAATGATACGCGAACTGTCAACGTCCATGTCGGGATGC |
| PA14_22980 | gltB | GAAGTTGGTCACCACGTCGAAGATCGCCCC | TTTCGTCCGCGAGTGACCAGTGATACGCGAACTGTAATGATACGCGAACTGTAAGAAGTTGGTCACCACGTCGAAGATCGCCCCAATGATACGCGAACTGTAATGATACGCGAACTGTCAACGTCCATGTCGGGATGC |
| PA14_22980 | gltB | GTACTTCATGATCTGCGCCACCTGCTTCGG | TTTCGTCCGCGAGTGACCAGTGATACGCGAACTGTAATGATACGCGAACTGTAAGTACTTCATGATCTGCGCCACCTGCTTCGGAATGATACGCGAACTGTAATGATACGCGAACTGTCAACGTCCATGTCGGGATGC |
| PA14_22980 | gltB | AGCGGGGAGAGACAGAGGGAGGAAAGACAG | TTTCGTCCGCGAGTGACCAGTGATACGCGAACTGTAATGATACGCGAACTGTAAAGCGGGGAGAGACAGAGGGAGGAAAGACAGAATGATACGCGAACTGTAATGATACGCGAACTGTCAACGTCCATGTCGGGATGC |
| PA14_22980 | gltB | TGTTGTAGTCGAAGGCCTTCTGCGTGCCGG | TTTCGTCCGCGAGTGACCAGTGATACGCGAACTGTAATGATACGCGAACTGTAATGTTGTAGTCGAAGGCCTTCTGCGTGCCGGAATGATACGCGAACTGTAATGATACGCGAACTGTCAACGTCCATGTCGGGATGC |
| PA14_22980 | gltB | ACTCCTGGATATCCGGCCCCTTGATCTGCG | TTTCGTCCGCGAGTGACCAGTGATACGCGAACTGTAATGATACGCGAACTGTAAACTCCTGGATATCCGGCCCCTTGATCTGCGAATGATACGCGAACTGTAATGATACGCGAACTGTCAACGTCCATGTCGGGATGC |
| PA14_22980 | gltB | TGGCCCTTGGCTTCGACCAGCTTCTTCAGG | TTTCGTCCGCGAGTGACCAGTGATACGCGAACTGTAATGATACGCGAACTGTAATGGCCCTTGGCTTCGACCAGCTTCTTCAGGAATGATACGCGAACTGTAATGATACGCGAACTGTCAACGTCCATGTCGGGATGC |
| PA14_22980 | gltB | AGGTCCACCAGTGCAGCACTTCGACTTCGC | TTTCGTCCGCGAGTGACCAGTGATACGCGAACTGTAATGATACGCGAACTGTAAAGGTCCACCAGTGCAGCACTTCGACTTCGCAATGATACGCGAACTGTAATGATACGCGAACTGTCAACGTCCATGTCGGGATGC |
| PA14_22980 | gltB | ACGTTGATCGGCACTGCTACGTAGTCGCCG | TTTCGTCCGCGAGTGACCAGTGATACGCGAACTGTAATGATACGCGAACTGTAAACGTTGATCGGCACTGCTACGTAGTCGCCGAATGATACGCGAACTGTAATGATACGCGAACTGTCAACGTCCATGTCGGGATGC |
| PA14_22980 | gltB | CTTGAAGTCCTTCATCGACTGCTGGGCGCA | TTTCGTCCGCGAGTGACCAGTGATACGCGAACTGTAATGATACGCGAACTGTAACTTGAAGTCCTTCATCGACTGCTGGGCGCAAATGATACGCGAACTGTAATGATACGCGAACTGTCAACGTCCATGTCGGGATGC |
| PA14_22980 | gltB | CGGACCCATCTTGCTCAGCACCAGGTTCTC | TTTCGTCCGCGAGTGACCAGTGATACGCGAACTGTAATGATACGCGAACTGTAACGGACCCATCTTGCTCAGCACCAGGTTCTCAATGATACGCGAACTGTAATGATACGCGAACTGTCAACGTCCATGTCGGGATGC |
| PA14_22980 | gltB | GTTGCTCAGCTTGAACATCACCAGCGAGTC | TTTCGTCCGCGAGTGACCAGTGATACGCGAACTGTAATGATACGCGAACTGTAAGTTGCTCAGCTTGAACATCACCAGCGAGTCAATGATACGCGAACTGTAATGATACGCGAACTGTCAACGTCCATGTCGGGATGC |
| PA14_05310 | gshB | AACTTCTCGTTGCAGTCGCGCAGGCTCTGC | TTTCGTCCGCGAGTGACCAGAATCAGGATACGGCGAAAATCAGGATACGGCGAAAACTTCTCGTTGCAGTCGCGCAGGCTCTGCAAAATCAGGATACGGCGAAAATCAGGATACGGCGCAACGTCCATGTCGGGATGC |
| PA14_05310 | gshB | AGCGGGAGGCATCGTTGAAGACCTTCAGCG | TTTCGTCCGCGAGTGACCAGAATCAGGATACGGCGAAAATCAGGATACGGCGAAAGCGGGAGGCATCGTTGAAGACCTTCAGCGAAAATCAGGATACGGCGAAAATCAGGATACGGCGCAACGTCCATGTCGGGATGC |
| PA14_05310 | gshB | ACGAACCGCCCATTCCGTCCAGGGGTTTGA | TTTCGTCCGCGAGTGACCAGAATCAGGATACGGCGAAAATCAGGATACGGCGAAACGAACCGCCCATTCCGTCCAGGGGTTTGAAAAATCAGGATACGGCGAAAATCAGGATACGGCGCAACGTCCATGTCGGGATGC |
| PA14_05310 | gshB | TTCGCCGTCGATCATCAGGATGCGTTTGTC | TTTCGTCCGCGAGTGACCAGAATCAGGATACGGCGAAAATCAGGATACGGCGAATTCGCCGTCGATCATCAGGATGCGTTTGTCAAAATCAGGATACGGCGAAAATCAGGATACGGCGCAACGTCCATGTCGGGATGC |
| PA14_05310 | gshB | CTTGCGCATCAGGATCACGTCCAGTTCGTG | TTTCGTCCGCGAGTGACCAGAATCAGGATACGGCGAAAATCAGGATACGGCGAACTTGCGCATCAGGATCACGTCCAGTTCGTGAAAATCAGGATACGGCGAAAATCAGGATACGGCGCAACGTCCATGTCGGGATGC |
| PA14_05310 | gshB | GTCCATGACGATCCCGAGGCGTACGCTCAT | TTTCGTCCGCGAGTGACCAGAATCAGGATACGGCGAAAATCAGGATACGGCGAAGTCCATGACGATCCCGAGGCGTACGCTCATAAAATCAGGATACGGCGAAAATCAGGATACGGCGCAACGTCCATGTCGGGATGC |
| PA14_05310 | gshB | ATGATCTGCTGGCTGCCATGCTGGGTGAGG | TTTCGTCCGCGAGTGACCAGAATCAGGATACGGCGAAAATCAGGATACGGCGAAATGATCTGCTGGCTGCCATGCTGGGTGAGGAAAATCAGGATACGGCGAAAATCAGGATACGGCGCAACGTCCATGTCGGGATGC |
| PA14_05310 | gshB | GCGAGCGAGCTGTCCTTCTTGAAGTTGATG | TTTCGTCCGCGAGTGACCAGAATCAGGATACGGCGAAAATCAGGATACGGCGAAGCGAGCGAGCTGTCCTTCTTGAAGTTGATGAAAATCAGGATACGGCGAAAATCAGGATACGGCGCAACGTCCATGTCGGGATGC |
| PA14_05310 | gshB | AGATAATCGCCGATCACGTCGAGGCCGACG | TTTCGTCCGCGAGTGACCAGAATCAGGATACGGCGAAAATCAGGATACGGCGAAAGATAATCGCCGATCACGTCGAGGCCGACGAAAATCAGGATACGGCGAAAATCAGGATACGGCGCAACGTCCATGTCGGGATGC |
| PA14_05310 | gshB | CGTCCTTGATTTCCGGCAGATAGCGCTGCG | TTTCGTCCGCGAGTGACCAGAATCAGGATACGGCGAAAATCAGGATACGGCGAACGTCCTTGATTTCCGGCAGATAGCGCTGCGAAAATCAGGATACGGCGAAAATCAGGATACGGCGCAACGTCCATGTCGGGATGC |
| PA14_05310 | gshB | GAATAGACGAATTCGTTATCGAAGGGCGGG | TTTCGTCCGCGAGTGACCAGAATCAGGATACGGCGAAAATCAGGATACGGCGAAGAATAGACGAATTCGTTATCGAAGGGCGGGAAAATCAGGATACGGCGAAAATCAGGATACGGCGCAACGTCCATGTCGGGATGC |
| PA14_05310 | gshB | GCGGTTGGTCGCTTTCCGCTTCCAGTTCGA | TTTCGTCCGCGAGTGACCAGAATCAGGATACGGCGAAAATCAGGATACGGCGAAGCGGTTGGTCGCTTTCCGCTTCCAGTTCGAAAAATCAGGATACGGCGAAAATCAGGATACGGCGCAACGTCCATGTCGGGATGC |
| PA14_05310 | gshB | AATATCGGATCGCCGGCTGACCATCGTCGG | TTTCGTCCGCGAGTGACCAGAATCAGGATACGGCGAAAATCAGGATACGGCGAAAATATCGGATCGCCGGCTGACCATCGTCGGAAAATCAGGATACGGCGAAAATCAGGATACGGCGCAACGTCCATGTCGGGATGC |
| PA14_05310 | gshB | TGTTCCATGTAGAACAGCGACCAGCCGCGG | TTTCGTCCGCGAGTGACCAGAATCAGGATACGGCGAAAATCAGGATACGGCGAATGTTCCATGTAGAACAGCGACCAGCCGCGGAAAATCAGGATACGGCGAAAATCAGGATACGGCGCAACGTCCATGTCGGGATGC |
| PA14_05310 | gshB | TGATGTCACGATGCTCGGCGGCAAACTCGC | TTTCGTCCGCGAGTGACCAGAATCAGGATACGGCGAAAATCAGGATACGGCGAATGATGTCACGATGCTCGGCGGCAAACTCGCAAAATCAGGATACGGCGAAAATCAGGATACGGCGCAACGTCCATGTCGGGATGC |
| PA14_36310 | hcnC | CCGGCAATCACGATGTCATAGGTTCTGTTC | TTTCGTCCGCGAGTGACCAGGGTCTCGGATATGCAAAGGTCTCGGATATGCAAACCGGCAATCACGATGTCATAGGTTCTGTTCAAGGTCTCGGATATGCAAAGGTCTCGGATATGCACAACGTCCATGTCGGGATGC |
| PA14_36310 | hcnC | TCGTCCTGGATCACGTATTTCAGGCCGGTG | TTTCGTCCGCGAGTGACCAGGGTCTCGGATATGCAAAGGTCTCGGATATGCAAATCGTCCTGGATCACGTATTTCAGGCCGGTGAAGGTCTCGGATATGCAAAGGTCTCGGATATGCACAACGTCCATGTCGGGATGC |
| PA14_36310 | hcnC | CCACGATGTGCTCGGCGTACTGGCGATCCT | TTTCGTCCGCGAGTGACCAGGGTCTCGGATATGCAAAGGTCTCGGATATGCAAACCACGATGTGCTCGGCGTACTGGCGATCCTAAGGTCTCGGATATGCAAAGGTCTCGGATATGCACAACGTCCATGTCGGGATGC |
| PA14_36310 | hcnC | TTGCTGACGTCGAAGCCCTTGTCCTCGGTG | TTTCGTCCGCGAGTGACCAGGGTCTCGGATATGCAAAGGTCTCGGATATGCAAATTGCTGACGTCGAAGCCCTTGTCCTCGGTGAAGGTCTCGGATATGCAAAGGTCTCGGATATGCACAACGTCCATGTCGGGATGC |
| PA14_36310 | hcnC | GGGAGGACATCATGCGGAAGAAGATCACCC | TTTCGTCCGCGAGTGACCAGGGTCTCGGATATGCAAAGGTCTCGGATATGCAAAGGGAGGACATCATGCGGAAGAAGATCACCCAAGGTCTCGGATATGCAAAGGTCTCGGATATGCACAACGTCCATGTCGGGATGC |
| PA14_36310 | hcnC | TCAGGCAACCGTTCAGCAGTCGCGGCATAC | TTTCGTCCGCGAGTGACCAGGGTCTCGGATATGCAAAGGTCTCGGATATGCAAATCAGGCAACCGTTCAGCAGTCGCGGCATACAAGGTCTCGGATATGCAAAGGTCTCGGATATGCACAACGTCCATGTCGGGATGC |
| PA14_36310 | hcnC | TCAGTTCTCGGTGCAGTTCCGGGTACAGCG | TTTCGTCCGCGAGTGACCAGGGTCTCGGATATGCAAAGGTCTCGGATATGCAAATCAGTTCTCGGTGCAGTTCCGGGTACAGCGAAGGTCTCGGATATGCAAAGGTCTCGGATATGCACAACGTCCATGTCGGGATGC |
| PA14_36310 | hcnC | CTCGAACTTGAAGTCCATCCCGTGGCGTTC | TTTCGTCCGCGAGTGACCAGGGTCTCGGATATGCAAAGGTCTCGGATATGCAAACTCGAACTTGAAGTCCATCCCGTGGCGTTCAAGGTCTCGGATATGCAAAGGTCTCGGATATGCACAACGTCCATGTCGGGATGC |
| PA14_36310 | hcnC | TTGGATTGCAGGGCGAGGTCGAAGAACGCC | TTTCGTCCGCGAGTGACCAGGGTCTCGGATATGCAAAGGTCTCGGATATGCAAATTGGATTGCAGGGCGAGGTCGAAGAACGCCAAGGTCTCGGATATGCAAAGGTCTCGGATATGCACAACGTCCATGTCGGGATGC |
| PA14_36310 | hcnC | AGGTTGACCTGTTGCAGCTCCGGCACGCAA | TTTCGTCCGCGAGTGACCAGGGTCTCGGATATGCAAAGGTCTCGGATATGCAAAAGGTTGACCTGTTGCAGCTCCGGCACGCAAAAGGTCTCGGATATGCAAAGGTCTCGGATATGCACAACGTCCATGTCGGGATGC |
| PA14_36310 | hcnC | TGACCTGGTGGTCGCAGAGGAATTCCAGCG | TTTCGTCCGCGAGTGACCAGGGTCTCGGATATGCAAAGGTCTCGGATATGCAAATGACCTGGTGGTCGCAGAGGAATTCCAGCGAAGGTCTCGGATATGCAAAGGTCTCGGATATGCACAACGTCCATGTCGGGATGC |
| PA14_36310 | hcnC | GTCCTTCTGCGCCATGTAGCAGTCGCTGGT | TTTCGTCCGCGAGTGACCAGGGTCTCGGATATGCAAAGGTCTCGGATATGCAAAGTCCTTCTGCGCCATGTAGCAGTCGCTGGTAAGGTCTCGGATATGCAAAGGTCTCGGATATGCACAACGTCCATGTCGGGATGC |
| PA14_36310 | hcnC | AATGCAGTACCCCGGCATTGTCCGTGCGCA | TTTCGTCCGCGAGTGACCAGGGTCTCGGATATGCAAAGGTCTCGGATATGCAAAAATGCAGTACCCCGGCATTGTCCGTGCGCAAAGGTCTCGGATATGCAAAGGTCTCGGATATGCACAACGTCCATGTCGGGATGC |
| PA14_36310 | hcnC | ATCTGGCCCTTCACCGGTTTCACCGGGATG | TTTCGTCCGCGAGTGACCAGGGTCTCGGATATGCAAAGGTCTCGGATATGCAAAATCTGGCCCTTCACCGGTTTCACCGGGATGAAGGTCTCGGATATGCAAAGGTCTCGGATATGCACAACGTCCATGTCGGGATGC |
| PA14_36310 | hcnC | TGGTTCCCAGCAACAGCTCGACGCCGTTCT | TTTCGTCCGCGAGTGACCAGGGTCTCGGATATGCAAAGGTCTCGGATATGCAAATGGTTCCCAGCAACAGCTCGACGCCGTTCTAAGGTCTCGGATATGCAAAGGTCTCGGATATGCACAACGTCCATGTCGGGATGC |
| PA14_62160 | ilvI | GAGTTGATCCAGCGATTGGGCTTGTTGAAC | TTTCGTCCGCGAGTGACCAGTGCGTACGACAATGTAATGCGTACGACAATGTAAGAGTTGATCCAGCGATTGGGCTTGTTGAACAATGCGTACGACAATGTAATGCGTACGACAATGTCAACGTCCATGTCGGGATGC |
| PA14_62160 | ilvI | GACCATTTCAGCGCCTGATAAAAGCTCCAC | TTTCGTCCGCGAGTGACCAGTGCGTACGACAATGTAATGCGTACGACAATGTAAGACCATTTCAGCGCCTGATAAAAGCTCCACAATGCGTACGACAATGTAATGCGTACGACAATGTCAACGTCCATGTCGGGATGC |
| PA14_62160 | ilvI | TACTTAACGCCTTCGTCGCGCAACGAGCGG | TTTCGTCCGCGAGTGACCAGTGCGTACGACAATGTAATGCGTACGACAATGTAATACTTAACGCCTTCGTCGCGCAACGAGCGGAATGCGTACGACAATGTAATGCGTACGACAATGTCAACGTCCATGTCGGGATGC |
| PA14_62160 | ilvI | TTCATCTGGATACTGCCCTCGCCGGTCACG | TTTCGTCCGCGAGTGACCAGTGCGTACGACAATGTAATGCGTACGACAATGTAATTCATCTGGATACTGCCCTCGCCGGTCACGAATGCGTACGACAATGTAATGCGTACGACAATGTCAACGTCCATGTCGGGATGC |
| PA14_62160 | ilvI | ATCGCCATGGGTAACCTCGTAGAGCGTCTC | TTTCGTCCGCGAGTGACCAGTGCGTACGACAATGTAATGCGTACGACAATGTAAATCGCCATGGGTAACCTCGTAGAGCGTCTCAATGCGTACGACAATGTAATGCGTACGACAATGTCAACGTCCATGTCGGGATGC |
| PA14_62160 | ilvI | GTAGGCTTCGGCAAGCTTGACGAAGTCCGG | TTTCGTCCGCGAGTGACCAGTGCGTACGACAATGTAATGCGTACGACAATGTAAGTAGGCTTCGGCAAGCTTGACGAAGTCCGGAATGCGTACGACAATGTAATGCGTACGACAATGTCAACGTCCATGTCGGGATGC |
| PA14_62160 | ilvI | ACCTCGGGAATTTCCGAGGGATGCTTGATG | TTTCGTCCGCGAGTGACCAGTGCGTACGACAATGTAATGCGTACGACAATGTAAACCTCGGGAATTTCCGAGGGATGCTTGATGAATGCGTACGACAATGTAATGCGTACGACAATGTCAACGTCCATGTCGGGATGC |
| PA14_62160 | ilvI | CAGGCGATTCTTCATGGCGAACGCCTCCTC | TTTCGTCCGCGAGTGACCAGTGCGTACGACAATGTAATGCGTACGACAATGTAACAGGCGATTCTTCATGGCGAACGCCTCCTCAATGCGTACGACAATGTAATGCGTACGACAATGTCAACGTCCATGTCGGGATGC |
| PA14_62160 | ilvI | TGTCCTTCGGAATATCCACGACCACCGGCC | TTTCGTCCGCGAGTGACCAGTGCGTACGACAATGTAATGCGTACGACAATGTAATGTCCTTCGGAATATCCACGACCACCGGCCAATGCGTACGACAATGTAATGCGTACGACAATGTCAACGTCCATGTCGGGATGC |
| PA14_62160 | ilvI | CCGACTGCGCCAGGTAAAAGGCCTTCTTGA | TTTCGTCCGCGAGTGACCAGTGCGTACGACAATGTAATGCGTACGACAATGTAACCGACTGCGCCAGGTAAAAGGCCTTCTTGAAATGCGTACGACAATGTAATGCGTACGACAATGTCAACGTCCATGTCGGGATGC |
| PA14_62160 | ilvI | GGGATTCCATGTAGGAATGCGAATAACGGC | TTTCGTCCGCGAGTGACCAGTGCGTACGACAATGTAATGCGTACGACAATGTAAGGGATTCCATGTAGGAATGCGAATAACGGCAATGCGTACGACAATGTAATGCGTACGACAATGTCAACGTCCATGTCGGGATGC |
| PA14_62160 | ilvI | ACCATTTCGGTGAGAACGCTGTCCACCGGG | TTTCGTCCGCGAGTGACCAGTGCGTACGACAATGTAATGCGTACGACAATGTAAACCATTTCGGTGAGAACGCTGTCCACCGGGAATGCGTACGACAATGTAATGCGTACGACAATGTCAACGTCCATGTCGGGATGC |
| PA14_62160 | ilvI | GAAGCTGTGCTTCACGATCGGCCGGGAGAT | TTTCGTCCGCGAGTGACCAGTGCGTACGACAATGTAATGCGTACGACAATGTAAGAAGCTGTGCTTCACGATCGGCCGGGAGATAATGCGTACGACAATGTAATGCGTACGACAATGTCAACGTCCATGTCGGGATGC |
| PA14_62160 | ilvI | TGTGGGTCACGTCGTGCTCCTTGAAGAGCG | TTTCGTCCGCGAGTGACCAGTGCGTACGACAATGTAATGCGTACGACAATGTAATGTGGGTCACGTCGTGCTCCTTGAAGAGCGAATGCGTACGACAATGTAATGCGTACGACAATGTCAACGTCCATGTCGGGATGC |
| PA14_62160 | ilvI | TAGATATGCAGGAGGGCACCACCCGGGTAC | TTTCGTCCGCGAGTGACCAGTGCGTACGACAATGTAATGCGTACGACAATGTAATAGATATGCAGGAGGGCACCACCCGGGTACAATGCGTACGACAATGTAATGCGTACGACAATGTCAACGTCCATGTCGGGATGC |
| PA14_62160 | ilvI | CGTACTGCAGGCAGGTAGACAGTTCCTGGA | TTTCGTCCGCGAGTGACCAGTGCGTACGACAATGTAATGCGTACGACAATGTAACGTACTGCAGGCAGGTAGACAGTTCCTGGAAATGCGTACGACAATGTAATGCGTACGACAATGTCAACGTCCATGTCGGGATGC |
| PA14_62160 | ilvI | GGAATATTCGAACTTCTGCGTCGGGTCGCC | TTTCGTCCGCGAGTGACCAGTGCGTACGACAATGTAATGCGTACGACAATGTAAGGAATATTCGAACTTCTGCGTCGGGTCGCCAATGCGTACGACAATGTAATGCGTACGACAATGTCAACGTCCATGTCGGGATGC |
| PA14_62160 | ilvI | CTGTACGAACGCAACTTGACCTTCTTCGGA | TTTCGTCCGCGAGTGACCAGTGCGTACGACAATGTAATGCGTACGACAATGTAACTGTACGAACGCAACTTGACCTTCTTCGGAAATGCGTACGACAATGTAATGCGTACGACAATGTCAACGTCCATGTCGGGATGC |
| PA14_62160 | ilvI | TCCGGGAAGTTGAGCTTGATGCCCATCGCC | TTTCGTCCGCGAGTGACCAGTGCGTACGACAATGTAATGCGTACGACAATGTAATCCGGGAAGTTGAGCTTGATGCCCATCGCCAATGCGTACGACAATGTAATGCGTACGACAATGTCAACGTCCATGTCGGGATGC |
| PA14_62160 | ilvI | CAGGGCACCATTGTTCAGGTTGACGATCTT | TTTCGTCCGCGAGTGACCAGTGCGTACGACAATGTAATGCGTACGACAATGTAACAGGGCACCATTGTTCAGGTTGACGATCTTAATGCGTACGACAATGTAATGCGTACGACAATGTCAACGTCCATGTCGGGATGC |
| PA14_27480 | htpX | TGAGGCCGCCATTGATGCCGAAGGCGTTCA | TTTCGTCCGCGAGTGACCAGACAACGTGCGCCTGAAAACAACGTGCGCCTGAAATGAGGCCGCCATTGATGCCGAAGGCGTTCAAAAACAACGTGCGCCTGAAAACAACGTGCGCCTGACAACGTCCATGTCGGGATGC |
| PA14_27480 | htpX | AACCGGCGAAACCGAACACGGCGCAGAAGA | TTTCGTCCGCGAGTGACCAGACAACGTGCGCCTGAAAACAACGTGCGCCTGAAAAACCGGCGAAACCGAACACGGCGCAGAAGAAAAACAACGTGCGCCTGAAAACAACGTGCGCCTGACAACGTCCATGTCGGGATGC |
| PA14_27480 | htpX | AGGTGTTCACCACGCCCTGGATCAACGCCA | TTTCGTCCGCGAGTGACCAGACAACGTGCGCCTGAAAACAACGTGCGCCTGAAAAGGTGTTCACCACGCCCTGGATCAACGCCAAAAACAACGTGCGCCTGAAAACAACGTGCGCCTGACAACGTCCATGTCGGGATGC |
| PA14_27480 | htpX | GCGCGAGAACCACATGACGATGATGCTGGC | TTTCGTCCGCGAGTGACCAGACAACGTGCGCCTGAAAACAACGTGCGCCTGAAAGCGCGAGAACCACATGACGATGATGCTGGCAAACAACGTGCGCCTGAAAACAACGTGCGCCTGACAACGTCCATGTCGGGATGC |
| PA14_27480 | htpX | CTGATGACTTCGGTACCGGTGCTCATCTTC | TTTCGTCCGCGAGTGACCAGACAACGTGCGCCTGAAAACAACGTGCGCCTGAAACTGATGACTTCGGTACCGGTGCTCATCTTCAAACAACGTGCGCCTGAAAACAACGTGCGCCTGACAACGTCCATGTCGGGATGC |
| PA14_27480 | htpX | TCGACGGTCTGCAGCAGCCACTGTTCGTGA | TTTCGTCCGCGAGTGACCAGACAACGTGCGCCTGAAAACAACGTGCGCCTGAAATCGACGGTCTGCAGCAGCCACTGTTCGTGAAAACAACGTGCGCCTGAAAACAACGTGCGCCTGACAACGTCCATGTCGGGATGC |
| PA14_27480 | htpX | AGCAGGCTGCCGTAATTCTGGCCGGTGAAG | TTTCGTCCGCGAGTGACCAGACAACGTGCGCCTGAAAACAACGTGCGCCTGAAAAGCAGGCTGCCGTAATTCTGGCCGGTGAAGAAACAACGTGCGCCTGAAAACAACGTGCGCCTGACAACGTCCATGTCGGGATGC |
| PA14_27480 | htpX | AAGATGGTCGCGACGAAGTAGCCGATGCCC | TTTCGTCCGCGAGTGACCAGACAACGTGCGCCTGAAAACAACGTGCGCCTGAAAAAGATGGTCGCGACGAAGTAGCCGATGCCCAAACAACGTGCGCCTGAAAACAACGTGCGCCTGACAACGTCCATGTCGGGATGC |
| PA14_27480 | htpX | TCACCTCATCGGGCGAGAAACGTTCGAGCA | TTTCGTCCGCGAGTGACCAGACAACGTGCGCCTGAAAACAACGTGCGCCTGAAATCACCTCATCGGGCGAGAAACGTTCGAGCAAAACAACGTGCGCCTGAAAACAACGTGCGCCTGACAACGTCCATGTCGGGATGC |

| | | | |
|---|---|---|---|
| PA14_27480 | htpX | GTTGCCGAAGATTCGCGCGAAGAACATCAC | TTTCGTCCGCGAGTGACCAGACAACGTGCGCCTGAAAACAACGTGCGCCTGAAAGTTGCCGAAGATTCGCGCGAAGAACATCACAAACAACGTGCGCCTGAAAACAACGTGCGCCTGACAACGTCCATGTCGGGATGC |
| PA14_27480 | htpX | GTTTCAGGGTGATGCTGGCAATCACCAGGA | TTTCGTCCGCGAGTGACCAGACAACGTGCGCCTGAAAACAACGTGCGCCTGAAAGTTTCAGGGTGATGCTGGCAATCACCAGGAAAACAACGTGCGCCTGAAAACAACGTGCGCCTGACAACGTCCATGTCGGGATGC |
| PA14_27480 | htpX | CATCCACTTGGAGATGAACAGCGAGACCAG | TTTCGTCCGCGAGTGACCAGACAACGTGCGCCTGAAAACAACGTGCGCCTGAAACATCCACTTGGAGATGAACAGCGAGACCAGAAACAACGTGCGCCTGAAAACAACGTGCGCCTGACAACGTCCATGTCGGGATGC |
| PA14_27480 | htpX | GCCAGGTTGGTGGCCAGGAACAACAGGATG | TTTCGTCCGCGAGTGACCAGACAACGTGCGCCTGAAAACAACGTGCGCCTGAAAGCCAGGTTGGTGGCCAGGAACAACAGGATGAAACAACGTGCGCCTGAAAACAACGTGCGCCTGACAACGTCCATGTCGGGATGC |
| PA14_27480 | htpX | TCCTCGTTCTTCAGGATCGCCTTGTCGACG | TTTCGTCCGCGAGTGACCAGACAACGTGCGCCTGAAAACAACGTGCGCCTGAAATCCTCGTTCTTCAGGATCGCCTTGTCGACGAAACAACGTGCGCCTGAAAACAACGTGCGCCTGACAACGTCCATGTCGGGATGC |
| PA14_27480 | htpX | TGGCGCGCAGCGCTTCGATACGATCTTCCA | TTTCGTCCGCGAGTGACCAGACAACGTGCGCCTGAAAACAACGTGCGCCTGAAATGGCGCGCAGCGCTTCGATACGATCTTCCAAAACAACGTGCGCCTGAAAACAACGTGCGCCTGACAACGTCCATGTCGGGATGC |
| PA14_34050 | impC | TGTAGAGCCTGGCCTTGTTGGTGGTCTGGC | TTTCGTCCGCGAGTGACCAGTAAGATGTCGCCGTGAATAAGATGTCGCCGTGAATGTAGAGCCTGGCCTTGTTGGTGGTCTGGCAATAAGATGTCGCCGTGAATAAGATGTCGCCGTGCAACGTCCATGTCGGGATGC |
| PA14_34050 | impC | CTTCCAGCTTCTGCAACTCGGGGTGGTGGA | TTTCGTCCGCGAGTGACCAGTAAGATGTCGCCGTGAATAAGATGTCGCCGTGAACTTCCAGCTTCTGCAACTCGGGGTGGTGGAAATAAGATGTCGCCGTGAATAAGATGTCGCCGTGCAACGTCCATGTCGGGATGC |
| PA14_34050 | impC | GGATTGTTCGGCGGCGGATGACTTGGGCAT | TTTCGTCCGCGAGTGACCAGTAAGATGTCGCCGTGAATAAGATGTCGCCGTGAAGGATTGTTCGGCGGCGGATGACTTGGGCATAATAAGATGTCGCCGTGAATAAGATGTCGCCGTGCAACGTCCATGTCGGGATGC |
| PA14_34050 | impC | AGCTGTTCATGTCGAACAGGCGTGGGCTGG | TTTCGTCCGCGAGTGACCAGTAAGATGTCGCCGTGAATAAGATGTCGCCGTGAAAGCTGTTCATGTCGAACAGGCGTGGGCTGGAATAAGATGTCGCCGTGAATAAGATGTCGCCGTGCAACGTCCATGTCGGGATGC |
| PA14_34050 | impC | TCGACAGCTTCTCCAGCAGGCCGATGTCCT | TTTCGTCCGCGAGTGACCAGTAAGATGTCGCCGTGAATAAGATGTCGCCGTGAATCGACAGCTTCTCCAGCAGGCCGATGTCCTAATAAGATGTCGCCGTGAATAAGATGTCGCCGTGCAACGTCCATGTCGGGATGC |
| PA14_34050 | impC | GCTGTCCGCCGAAGGTGCCGTATTCCTCTT | TTTCGTCCGCGAGTGACCAGTAAGATGTCGCCGTGAATAAGATGTCGCCGTGAAGCTGTCCGCCGAAGGTGCCGTATTCCTCTTAATAAGATGTCGCCGTGAATAAGATGTCGCCGTGCAACGTCCATGTCGGGATGC |
| PA14_34050 | impC | TGCAGTTCGTTCTTGCCGACGTTCAGCAGG | TTTCGTCCGCGAGTGACCAGTAAGATGTCGCCGTGAATAAGATGTCGCCGTGAATGCAGTTCGTTCTTGCCGACGTTCAGCAGGAATAAGATGTCGCCGTGAATAAGATGTCGCCGTGCAACGTCCATGTCGGGATGC |
| PA14_34050 | impC | ATAGTCCTGCTGGCTGTCGTCGTGGGCCAT | TTTCGTCCGCGAGTGACCAGTAAGATGTCGCCGTGAATAAGATGTCGCCGTGAAATAGTCCTGCTGGCTGTCGTCGTGGGCCATAATAAGATGTCGCCGTGAATAAGATGTCGCCGTGCAACGTCCATGTCGGGATGC |
| PA14_34050 | impC | ATCTGGTTGAGCTGGTCGCTGATCAGCGCA | TTTCGTCCGCGAGTGACCAGTAAGATGTCGCCGTGAATAAGATGTCGCCGTGAAATCTGGTTGAGCTGGTCGCTGATCAGCGCAAATAAGATGTCGCCGTGAATAAGATGTCGCCGTGCAACGTCCATGTCGGGATGC |
| PA14_34050 | impC | TCTCGGTGTTGCTCACCAGTTGGTGCAGGC | TTTCGTCCGCGAGTGACCAGTAAGATGTCGCCGTGAATAAGATGTCGCCGTGAATCTCGGTGTTGCTCACCAGTTGGTGCAGGCAATAAGATGTCGCCGTGAATAAGATGTCGCCGTGCAACGTCCATGTCGGGATGC |
| PA14_34050 | impC | TTGATTTCCTGCGGTGCGTTGTCGTTGATC | TTTCGTCCGCGAGTGACCAGTAAGATGTCGCCGTGAATAAGATGTCGCCGTGAATTGATTTCCTGCGGTGCGTTGTCGTTGATCAATAAGATGTCGCCGTGAATAAGATGTCGCCGTGCAACGTCCATGTCGGGATGC |
| PA14_34050 | impC | ATGGCGACCGAAGGTGAAGTCGCCGATCAG | TTTCGTCCGCGAGTGACCAGTAAGATGTCGCCGTGAATAAGATGTCGCCGTGAAATGGCGACCGAAGGTGAAGTCGCCGATCAGAATAAGATGTCGCCGTGAATAAGATGTCGCCGTGCAACGTCCATGTCGGGATGC |
| PA14_34050 | impC | GATCTGGCTGATGCGGTCGTTGATCATCGC | TTTCGTCCGCGAGTGACCAGTAAGATGTCGCCGTGAATAAGATGTCGCCGTGAAGATCTGGCTGATGCGGTCGTTGATCATCGCAATAAGATGTCGCCGTGAATAAGATGTCGCCGTGCAACGTCCATGTCGGGATGC |
| PA14_34050 | impC | TCTTGAACAGTGCGCTCTGGTCGAACTCGA | TTTCGTCCGCGAGTGACCAGTAAGATGTCGCCGTGAATAAGATGTCGCCGTGAATCTTGAACAGTGCGCTCTGGTCGAACTCGAAATAAGATGTCGCCGTGAATAAGATGTCGCCGTGCAACGTCCATGTCGGGATGC |
| PA14_34050 | impC | ATGATCTCGTCGAGCAGGGACAGGGTCTGG | TTTCGTCCGCGAGTGACCAGTAAGATGTCGCCGTGAATAAGATGTCGCCGTGAAATGATCTCGTCGAGCAGGGACAGGGTCTGGAATAAGATGTCGCCGTGAATAAGATGTCGCCGTGCAACGTCCATGTCGGGATGC |
| PA14_09150 | katA | GCTTCTCGAGGAACCACACGTCCTGCAGCA | TTTCGTCCGCGAGTGACCAGTATCATACGACGTGGAATATCATACGACGTGGAAGCTTCTCGAGGAACCACACGTCCTGCAGCAAATATCATACGACGTGGAATATCATACGACGTGGCAACGTCCATGTCGGGATGC |
| PA14_09150 | katA | TCATGGGTGACGGTGAAGGTGCCATAGGCC | TTTCGTCCGCGAGTGACCAGTATCATACGACGTGGAATATCATACGACGTGGAATCATGGGTGACGGTGAAGGTGCCATAGGCCAATATCATACGACGTGGAATATCATACGACGTGGCAACGTCCATGTCGGGATGC |
| PA14_09150 | katA | TCTCGTACAAGTCGCGCTGCGAACTCTCGC | TTTCGTCCGCGAGTGACCAGTATCATACGACGTGGAATATCATACGACGTGGAATCTCGTACAAGTCGCGCTGCGAACTCTCGCAATATCATACGACGTGGAATATCATACGACGTGGCAACGTCCATGTCGGGATGC |
| PA14_09150 | katA | GATGCAGCGACTCGGGGAGGTGAGAGAAAA | TTTCGTCCGCGAGTGACCAGTATCATACGACGTGGAATATCATACGACGTGGAAGATGCAGCGACTCGGGGAGGTGAGAGAAAAAATATCATACGACGTGGAATATCATACGACGTGGCAACGTCCATGTCGGGATGC |
| PA14_09150 | katA | TTCGATCAACGGGTAGTCACCGTGGGGCCA | TTTCGTCCGCGAGTGACCAGTATCATACGACGTGGAATATCATACGACGTGGAATTCGATCAACGGGTAGTCACCGTGGGGCCAAATATCATACGACGTGGAATATCATACGACGTGGCAACGTCCATGTCGGGATGC |
| PA14_09150 | katA | TGGTTGAGATCGGGGAACTTGAGCGGGTCG | TTTCGTCCGCGAGTGACCAGTATCATACGACGTGGAATATCATACGACGTGGAATGGTTGAGATCGGGGAACTTGAGCGGGTCGAATATCATACGACGTGGAATATCATACGACGTGGCAACGTCCATGTCGGGATGC |
| PA14_09150 | katA | TCCGGGTTACGGTTCAGCTCGAAGAAGCCC | TTTCGTCCGCGAGTGACCAGTATCATACGACGTGGAATATCATACGACGTGGAATCCGGGTTACGGTTCAGCTCGAAGAAGCCCAATATCATACGACGTGGAATATCATACGACGTGGCAACGTCCATGTCGGGATGC |
| PA14_09150 | katA | ATAGCTCTTCGGCAGACCGCGGTCGCTGAA | TTTCGTCCGCGAGTGACCAGTATCATACGACGTGGAATATCATACGACGTGGAAATAGCTCTTCGGCAGACCGCGGTCGCTGAAAATATCATACGACGTGGAATATCATACGACGTGGCAACGTCCATGTCGGGATGC |
| PA14_09150 | katA | AAGGTATGGCTGCCGAAACCATGGATGTGC | TTTCGTCCGCGAGTGACCAGTATCATACGACGTGGAATATCATACGACGTGGAAAAGGTATGGCTGCCGAAACCATGGATGTGCAATATCATACGACGTGGAATATCATACGACGTGGCAACGTCCATGTCGGGATGC |
| PA14_09150 | katA | AGATAGAACACCGGGGTGTTGTTGCCGACC | TTTCGTCCGCGAGTGACCAGTATCATACGACGTGGAATATCATACGACGTGGAAAGATAGAACACCGGGGTGTTGTTGCCGACCAATATCATACGACGTGGAATATCATACGACGTGGCAACGTCCATGTCGGGATGC |
| PA14_09150 | katA | AGAAGCGCTCGTTGTTCGCATTGATGAAGC | TTTCGTCCGCGAGTGACCAGTATCATACGACGTGGAATATCATACGACGTGGAAAGAAGCGCTCGTTGTTCGCATTGATGAAGCAATATCATACGACGTGGAATATCATACGACGTGGCAACGTCCATGTCGGGATGC |
| PA14_09150 | katA | ACCTGGGAGAAGATCTTCGCCCGGGTGTAG | TTTCGTCCGCGAGTGACCAGTATCATACGACGTGGAATATCATACGACGTGGAAACCTGGGAGAAGATCTTCGCCCGGGTGTAGAATATCATACGACGTGGAATATCATACGACGTGGCAACGTCCATGTCGGGATGC |
| PA14_09150 | katA | CTTCCAGCGCGGGAAGTCGCCTTTCTCGAT | TTTCGTCCGCGAGTGACCAGTATCATACGACGTGGAATATCATACGACGTGGAACTTCCAGCGCGGGAAGTCGCCTTTCTCGATAATATCATACGACGTGGAATATCATACGACGTGGCAACGTCCATGTCGGGATGC |
| PA14_09150 | katA | TCTTGTCCGGCGAGAAACCGATACCGGGCA | TTTCGTCCGCGAGTGACCAGTATCATACGACGTGGAATATCATACGACGTGGAATCTTGTCCGGCGAGAAACCGATACCGGGCAAATATCATACGACGTGGAATATCATACGACGTGGCAACGTCCATGTCGGGATGC |
| PA14_09150 | katA | ACGTTCTGGTTATCGACCACCGGTGCGCCA | TTTCGTCCGCGAGTGACCAGTATCATACGACGTGGAATATCATACGACGTGGAAACGTTCTGGTTATCGACCACCGGTGCGCCAAATATCATACGACGTGGAATATCATACGACGTGGCAACGTCCATGTCGGGATGC |
| PA14_09150 | katA | TGGAGAAGCCGCGGATGTCTCGCTCTGCAT | TTTCGTCCGCGAGTGACCAGTATCATACGACGTGGAATATCATACGACGTGGAATGGAGAAGCCGCGGATGTCTCGCTCTGCATAATATCATACGACGTGGAATATCATACGACGTGGCAACGTCCATGTCGGGATGC |
| PA14_09150 | katA | TTGAAGGTCGCATTGCGCAGGTTGGTGCGC | TTTCGTCCGCGAGTGACCAGTATCATACGACGTGGAATATCATACGACGTGGAATTGAAGGTCGCATTGCGCAGGTTGGTGCGCAATATCATACGACGTGGAATATCATACGACGTGGCAACGTCCATGTCGGGATGC |
| PA14_09150 | katA | AGAAGCGCAGGAACATATCGGTCTTCTTGC | TTTCGTCCGCGAGTGACCAGTATCATACGACGTGGAATATCATACGACGTGGAAAGAAGCGCAGGAACATATCGGTCTTCTTGCAATATCATACGACGTGGAATATCATACGACGTGGCAACGTCCATGTCGGGATGC |
| PA14_09150 | katA | GTCCCAGTTGCCCTGCTCGGTGTAGAAACG | TTTCGTCCGCGAGTGACCAGTATCATACGACGTGGAATATCATACGACGTGGAAGTCCCAGTTGCCCTGCTCGGTGTAGAAACGAATATCATACGACGTGGAATATCATACGACGTGGCAACGTCCATGTCGGGATGC |
| PA14_09150 | katA | GCCCTGTTGGGTCTTGAAGTGGAACTTGAC | TTTCGTCCGCGAGTGACCAGTATCATACGACGTGGAATATCATACGACGTGGAAGCCCTGTTGGGTCTTGAAGTGGAACTTGACAATATCATACGACGTGGAATATCATACGACGTGGCAACGTCCATGTCGGGATGC |
| PA14_61040 | katB | ATACCGGGGTCTTCTCGCCCTTGCGGAAGA | TTTCGTCCGCGAGTGACCAGTGGATCATTCTACGGAATGGATCATTCTACGGAAATACCGGGGTCTTCTCGCCCTTGCGGAAGAAATGGATCATTCTACGGAATGGATCATTCTACGGCAACGTCCATGTCGGGATGC |
| PA14_61040 | katB | TTGGAATACAGCAGGGTCAGGGTGCGGGTC | TTTCGTCCGCGAGTGACCAGTGGATCATTCTACGGAATGGATCATTCTACGGAATTGGAATACAGCAGGGTCAGGGTGCGGGTCAATGGATCATTCTACGGAATGGATCATTCTACGGCAACGTCCATGTCGGGATGC |
| PA14_61040 | katB | AGAAGGTCGGGAAGTTGTTGCCCACCAGGT | TTTCGTCCGCGAGTGACCAGTGGATCATTCTACGGAATGGATCATTCTACGGAAAGAAGGTCGGGAAGTTGTTGCCCACCAGGTAATGGATCATTCTACGGAATGGATCATTCTACGGCAACGTCCATGTCGGGATGC |
| PA14_61040 | katB | TCCTGGTTGACGGTATTGACCTCGCTGCGC | TTTCGTCCGCGAGTGACCAGTGGATCATTCTACGGAATGGATCATTCTACGGAATCCTGGTTGACGGTATTGACCTCGCTGCGCAATGGATCATTCTACGGAATGGATCATTCTACGGCAACGTCCATGTCGGGATGC |
| PA14_61040 | katB | TCGCGGTCGAAACGTTGCAACTTCTGCAGC | TTTCGTCCGCGAGTGACCAGTGGATCATTCTACGGAATGGATCATTCTACGGAATCGCGGTCGAAACGTTGCAACTTCTGCAGCAATGGATCATTCTACGGAATGGATCATTCTACGGCAACGTCCATGTCGGGATGC |
| PA14_61040 | katB | GGGACGGAAGGCGTTCAGGGAAGGGTTCAT | TTTCGTCCGCGAGTGACCAGTGGATCATTCTACGGAATGGATCATTCTACGGAAGGGACGGAAGGCGTTCAGGGAAGGGTTCATAATGGATCATTCTACGGAATGGATCATTCTACGGCAACGTCCATGTCGGGATGC |
| PA14_61040 | katB | TTGCACGTCCTGCAGCAGCACCGAACCGTT | TTTCGTCCGCGAGTGACCAGTGGATCATTCTACGGAATGGATCATTCTACGGAATTGCACGTCCTGCAGCAGCACCGAACCGTTAATGGATCATTCTACGGAATGGATCATTCTACGGCAACGTCCATGTCGGGATGC |
| PA14_61040 | katB | GTAGTCGCTGTCAGCCTTGTAGAAGTAGGA | TTTCGTCCGCGAGTGACCAGTGGATCATTCTACGGAATGGATCATTCTACGGAAGTAGTCGCTGTCAGCCTTGTAGAAGTAGGAAATGGATCATTCTACGGAATGGATCATTCTACGGCAACGTCCATGTCGGGATGC |
| PA14_61040 | katB | TAGGCATGCACGCTGTTGCCGTCCATTTCC | TTTCGTCCGCGAGTGACCAGTGGATCATTCTACGGAATGGATCATTCTACGGAATAGGCATGCACGCTGTTGCCGTCCATTTCCAATGGATCATTCTACGGAATGGATCATTCTACGGCAACGTCCATGTCGGGATGC |
| PA14_61040 | katB | ATCGAAGCGTCGCGAGTCGTCGTCGAGGTT | TTTCGTCCGCGAGTGACCAGTGGATCATTCTACGGAATGGATCATTCTACGGAAATCGAAGCGTCGCGAGTCGTCGTCGAGGTTAATGGATCATTCTACGGAATGGATCATTCTACGGCAACGTCCATGTCGGGATGC |
| PA14_61040 | katB | CGGTATAGAACTTGGTGGCGAAACCGCGCG | TTTCGTCCGCGAGTGACCAGTGGATCATTCTACGGAATGGATCATTCTACGGAACGGTATAGAACTTGGTGGCGAAACCGCGCGAATGGATCATTCTACGGAATGGATCATTCTACGGCAACGTCCATGTCGGGATGC |
| PA14_61040 | katB | TTGGCCATGCTCAGGTCGCTGATATCGGCG | TTTCGTCCGCGAGTGACCAGTGGATCATTCTACGGAATGGATCATTCTACGGAATTGGCCATGCTCAGGTCGCTGATATCGGCGAATGGATCATTCTACGGAATGGATCATTCTACGGCAACGTCCATGTCGGGATGC |
| PA14_61040 | katB | ACCATCTGGCCGATCTTGCGCTCGGGAATG | TTTCGTCCGCGAGTGACCAGTGGATCATTCTACGGAATGGATCATTCTACGGAAACCATCTGGCCGATCTTGCGCTCGGGAATGAATGGATCATTCTACGGAATGGATCATTCTACGGCAACGTCCATGTCGGGATGC |
| PA14_61040 | katB | GGCATGGACCATGTCCGGGAACTTGATCGC | TTTCGTCCGCGAGTGACCAGTGGATCATTCTACGGAATGGATCATTCTACGGAAGGCATGGACCATGTCCGGGAACTTGATCGCAATGGATCATTCTACGGAATGGATCATTCTACGGCAACGTCCATGTCGGGATGC |
| PA14_61040 | katB | TGGTAGTTCACCCCGCTGGTACTGTGTCCG | TTTCGTCCGCGAGTGACCAGTGGATCATTCTACGGAATGGATCATTCTACGGAATGGTAGTTCACCCCGCTGGTACTGTGTCCGAATGGATCATTCTACGGAATGGATCATTCTACGGCAACGTCCATGTCGGGATGC |
| PA14_72960 | kgtP | AGACGCGGAAATGGCGTTGGCGCTTTCCAT | TTTCGTCCGCGAGTGACCAGTCTAGTCTCACGCAAAATCTAGTCTCACGCAAAAAGACGCGGAAATGGCGTTGGCGCTTTCCATAATCTAGTCTCACGCAAAATCTAGTCTCACGCAACAACGTCCATGTCGGGATGC |
| PA14_72960 | kgtP | TGGGGAACAGTTCGGCCTTCACTACCGCGT | TTTCGTCCGCGAGTGACCAGTCTAGTCTCACGCAAAATCTAGTCTCACGCAAAATGGGGAACAGTTCGGCCTTCACTACCGCGTAATCTAGTCTCACGCAAAATCTAGTCTCACGCAACAACGTCCATGTCGGGATGC |
| PA14_72960 | kgtP | AGCCAAAGCACATCAGCAGCACCGAGGCCA | TTTCGTCCGCGAGTGACCAGTCTAGTCTCACGCAAAATCTAGTCTCACGCAAAAAGCCAAAGCACATCAGCAGCACCGAGGCCAAATCTAGTCTCACGCAAAATCTAGTCTCACGCAACAACGTCCATGTCGGGATGC |
| PA14_72960 | kgtP | TGTTCAGCAGTTGGGCGGTCATGTCGCCCT | TTTCGTCCGCGAGTGACCAGTCTAGTCTCACGCAAAATCTAGTCTCACGCAAAATGTTCAGCAGTTGGGCGGTCATGTCGCCCTAATCTAGTCTCACGCAAAATCTAGTCTCACGCAACAACGTCCATGTCGGGATGC |
| PA14_72960 | kgtP | TTGTTCGCCATCTCGCTCAGGTAGGTGGCG | TTTCGTCCGCGAGTGACCAGTCTAGTCTCACGCAAAATCTAGTCTCACGCAAAATTGTTCGCCATCTCGCTCAGGTAGGTGGCGAATCTAGTCTCACGCAAAATCTAGTCTCACGCAACAACGTCCATGTCGGGATGC |
| PA14_72960 | kgtP | AAGGCGGCGTAGACGTACCAGTCGTACCAC | TTTCGTCCGCGAGTGACCAGTCTAGTCTCACGCAAAATCTAGTCTCACGCAAAAAAGGCGGCGTAGACGTACCAGTCGTACCACAATCTAGTCTCACGCAAAATCTAGTCTCACGCAACAACGTCCATGTCGGGATGC |
| PA14_72960 | kgtP | TAGGTGAAGACGGTGCCGAGTACGCCGAAG | TTTCGTCCGCGAGTGACCAGTCTAGTCTCACGCAAAATCTAGTCTCACGCAAAATAGGTGAAGACGGTGCCGAGTACGCCGAAGAATCTAGTCTCACGCAAAATCTAGTCTCACGCAACAACGTCCATGTCGGGATGC |
| PA14_72960 | kgtP | GCGCCAGCCCCAGCTTTCCAGTTGTTCTAC | TTTCGTCCGCGAGTGACCAGTCTAGTCTCACGCAAAATCTAGTCTCACGCAAAAGCGCCAGCCCCAGCTTTCCAGTTGTTCTACAATCTAGTCTCACGCAAAATCTAGTCTCACGCAACAACGTCCATGTCGGGATGC |
| PA14_72960 | kgtP | CGGTCGTGATCTTCGAGTGCTTCTGGGTGT | TTTCGTCCGCGAGTGACCAGTCTAGTCTCACGCAAAATCTAGTCTCACGCAAAACGGTCGTGATCTTCGAGTGCTTCTGGGTGTAATCTAGTCTCACGCAAAATCTAGTCTCACGCAACAACGTCCATGTCGGGATGC |
| PA14_72960 | kgtP | TAGATACCCATCAGCCAGCCACCGATCGGG | TTTCGTCCGCGAGTGACCAGTCTAGTCTCACGCAAAATCTAGTCTCACGCAAAATAGATACCCATCAGCCAGCCACCGATCGGGAATCTAGTCTCACGCAAAATCTAGTCTCACGCAACAACGTCCATGTCGGGATGC |
| PA14_72960 | kgtP | ACTTGATGCGTTGCGAAGTGGTGCGGGGCT | TTTCGTCCGCGAGTGACCAGTCTAGTCTCACGCAAAATCTAGTCTCACGCAAAAACTTGATGCGTTGCGAAGTGGTGCGGGGCTAATCTAGTCTCACGCAAAATCTAGTCTCACGCAACAACGTCCATGTCGGGATGC |
| PA14_72960 | kgtP | CATGCCTACCGTGTTCACCAGGTACTTCTG | TTTCGTCCGCGAGTGACCAGTCTAGTCTCACGCAAAATCTAGTCTCACGCAAAACATGCCTACCGTGTTCACCAGGTACTTCTGAATCTAGTCTCACGCAAAATCTAGTCTCACGCAACAACGTCCATGTCGGGATGC |
| PA14_72960 | kgtP | GACCAGGTTGCCGACGGAACCGCTGAAGAT | TTTCGTCCGCGAGTGACCAGTCTAGTCTCACGCAAAATCTAGTCTCACGCAAAAGACCAGGTTGCCGACGGAACCGCTGAAGATAATCTAGTCTCACGCAAAATCTAGTCTCACGCAACAACGTCCATGTCGGGATGC |
| PA14_72960 | kgtP | GCAGCCGAGATCATGGTCGAATCGGTCTTG | TTTCGTCCGCGAGTGACCAGTCTAGTCTCACGCAAAATCTAGTCTCACGCAAAAGCAGCCGAGATCATGGTCGAATCGGTCTTGAATCTAGTCTCACGCAAAATCTAGTCTCACGCAACAACGTCCATGTCGGGATGC |
| PA14_72960 | kgtP | AACCACAAGGCAACGTATTCGGCGGTGCCG | TTTCGTCCGCGAGTGACCAGTCTAGTCTCACGCAAAATCTAGTCTCACGCAAAAAACCACAAGGCAACGTATTCGGCGGTGCCGAATCTAGTCTCACGCAAAATCTAGTCTCACGCAACAACGTCCATGTCGGGATGC |
| PA14_40290 | lasA | TTGCTGGCATAGACGCCGAGCTTGGTGTCG | TTTCGTCCGCGAGTGACCAGAGATTGGCGCGACATAAAGATTGGCGCGACATAATTGCTGGCATAGACGCCGAGCTTGGTGTCGAAAGATTGGCGCGACATAAAGATTGGCGCGACATCAACGTCCATGTCGGGATGC |
| PA14_40290 | lasA | AACAGTTCGTCATTCAGCGGCAGGGCCACG | TTTCGTCCGCGAGTGACCAGAGATTGGCGCGACATAAAGATTGGCGCGACATAAAACAGTTCGTCATTCAGCGGCAGGGCCACGAAAGATTGGCGCGACATAAAGATTGGCGCGACATCAACGTCCATGTCGGGATGC |
| PA14_40290 | lasA | TGCAACACGTCGCGTACCTGGGCATCGAAG | TTTCGTCCGCGAGTGACCAGAGATTGGCGCGACATAAAGATTGGCGCGACATAATGCAACACGTCGCGTACCTGGGCATCGAAGAAAGATTGGCGCGACATAAAGATTGGCGCGACATCAACGTCCATGTCGGGATGC |
| PA14_40290 | lasA | TGACCTGCTGGCCGTTGCTCACCTGGATCT | TTTCGTCCGCGAGTGACCAGAGATTGGCGCGACATAAAGATTGGCGCGACATAATGACCTGCTGGCCGTTGCTCACCTGGATCTAAAGATTGGCGCGACATAAAGATTGGCGCGACATCAACGTCCATGTCGGGATGC |
| PA14_40290 | lasA | AAGGTCTGCGCGTAGGCTCCGAGCGGATTG | TTTCGTCCGCGAGTGACCAGAGATTGGCGCGACATAAAGATTGGCGCGACATAAAAGGTCTGCGCGTAGGCTCCGAGCGGATTGAAAGATTGGCGCGACATAAAGATTGGCGCGACATCAACGTCCATGTCGGGATGC |
| PA14_40290 | lasA | GGGTCAACAACACTTTCGGGTTGATGCTGT | TTTCGTCCGCGAGTGACCAGAGATTGGCGCGACATAAAGATTGGCGCGACATAAGGGTCAACAACACTTTCGGGTTGATGCTGTAAAGATTGGCGCGACATAAAGATTGGCGCGACATCAACGTCCATGTCGGGATGC |
| PA14_40290 | lasA | CCTCGAAACCGTAGTAGCGTCGCGACAACT | TTTCGTCCGCGAGTGACCAGAGATTGGCGCGACATAAAGATTGGCGCGACATAACCTCGAAACCGTAGTAGCGTCGCGACAACTAAAGATTGGCGCGACATAAAGATTGGCGCGACATCAACGTCCATGTCGGGATGC |
| PA14_40290 | lasA | ATCGAAGGACGAGTACGGATAGCCCGAGCC | TTTCGTCCGCGAGTGACCAGAGATTGGCGCGACATAAAGATTGGCGCGACATAAATCGAAGGACGAGTACGGATAGCCCGAGCCAAAGATTGGCGCGACATAAAGATTGGCGCGACATCAACGTCCATGTCGGGATGC |
| PA14_40290 | lasA | TAGAGCAGCGAGAAATGCAGGTGCGGTCCG | TTTCGTCCGCGAGTGACCAGAGATTGGCGCGACATAAAGATTGGCGCGACATAATAGAGCAGCGAGAAATGCAGGTGCGGTCCGAAAGATTGGCGCGACATAAAGATTGGCGCGACATCAACGTCCATGTCGGGATGC |
| PA14_40290 | lasA | GCCGCTCCAGTGCTCCAGGTATTCGCTCTT | TTTCGTCCGCGAGTGACCAGAGATTGGCGCGACATAAAGATTGGCGCGACATAAGCCGCTCCAGTGCTCCAGGTATTCGCTCTTAAAGATTGGCGCGACATAAAGATTGGCGCGACATCAACGTCCATGTCGGGATGC |
| PA14_40290 | lasA | AATGCGCTCCGTTGGGCTGCCAGGAATAGC | TTTCGTCCGCGAGTGACCAGAGATTGGCGCGACATAAAGATTGGCGCGACATAAAATGCGCTCCGTTGGGCTGCCAGGAATAGCAAAGATTGGCGCGACATAAAGATTGGCGCGACATCAACGTCCATGTCGGGATGC |
| PA14_40290 | lasA | AATTGCATCAGGTTGGATGGCGGCGCCAGC | TTTCGTCCGCGAGTGACCAGAGATTGGCGCGACATAAAGATTGGCGCGACATAAAATTGCATCAGGTTGGATGGCGGCGCCAGCAAAGATTGGCGCGACATAAAGATTGGCGCGACATCAACGTCCATGTCGGGATGC |
| PA14_40290 | lasA | TAGTTGCTGGTGCCGACGTTGATCCGGTAC | TTTCGTCCGCGAGTGACCAGAGATTGGCGCGACATAAAGATTGGCGCGACATAATAGTTGCTGGTGCCGACGTTGATCCGGTACAAAGATTGGCGCGACATAAAGATTGGCGCGACATCAACGTCCATGTCGGGATGC |
| PA14_40290 | lasA | ATGTGGTAGTAGTTGGTCGCCCAGCCGCTG | TTTCGTCCGCGAGTGACCAGAGATTGGCGCGACATAAAGATTGGCGCGACATAAATGTGGTAGTAGTTGGTCGCCCAGCCGCTGAAAGATTGGCGCGACATAAAGATTGGCGCGACATCAACGTCCATGTCGGGATGC |
| PA14_40290 | lasA | ACCGAAGCTGGCTCCCTGCAGGGAAACGAA | TTTCGTCCGCGAGTGACCAGAGATTGGCGCGACATAAAGATTGGCGCGACATAAACCGAAGCTGGCTCCCTGCAGGGAAACGAAAAAGATTGGCGCGACATAAAGATTGGCGCGACATCAACGTCCATGTCGGGATGC |
| PA14_40290 | lasA | TGCAACTGGCCGAGTAGCTCCGCCGAATAG | TTTCGTCCGCGAGTGACCAGAGATTGGCGCGACATAAAGATTGGCGCGACATAATGCAACTGGCCGAGTAGCTCCGCCGAATAGAAAGATTGGCGCGACATAAAGATTGGCGCGACATCAACGTCCATGTCGGGATGC |
| PA14_40290 | lasA | TACCTGGCAGCGCGACAACACCCGTACCGT | TTTCGTCCGCGAGTGACCAGAGATTGGCGCGACATAAAGATTGGCGCGACATAATACCTGGCAGCGCGACAACACCCGTACCGTAAAGATTGGCGCGACATAAAGATTGGCGCGACATCAACGTCCATGTCGGGATGC |
| PA14_40290 | lasA | GGTTGTAGAAATAGTAGCGGCGACAGTCGT | TTTCGTCCGCGAGTGACCAGAGATTGGCGCGACATAAAGATTGGCGCGACATAAGGTTGTAGAAATAGTAGCGGCGACAGTCGTAAAGATTGGCGCGACATAAAGATTGGCGCGACATCAACGTCCATGTCGGGATGC |
| PA14_40290 | lasA | CGCCAGGGCGCGGGATCTTTTGTGCTGCAT | TTTCGTCCGCGAGTGACCAGAGATTGGCGCGACATAAAGATTGGCGCGACATAACGCCAGGGCGCGGGATCTTTTGTGCTGCATAAAGATTGGCGCGACATAAAGATTGGCGCGACATCAACGTCCATGTCGGGATGC |
| PA14_40290 | lasA | TACAACGGGCGGAAAGCGCAATGGGTGGTG | TTTCGTCCGCGAGTGACCAGAGATTGGCGCGACATAAAGATTGGCGCGACATAATACAACGGGCGGAAAGCGCAATGGGTGGTGAAAGATTGGCGCGACATAAAGATTGGCGCGACATCAACGTCCATGTCGGGATGC |
| PA14_45940 | lasI | CGCAACTTGTGCATCTCGCCCAGCAGTTTT | TTTCGTCCGCGAGTGACCAGAACGCGTAGGCTAGAAAAACGCGTAGGCTAGAAACGCAACTTGTGCATCTCGCCCAGCAGTTTTAAAACGCGTAGGCTAGAAAAACGCGTAGGCTAGACAACGTCCATGTCGGGATGC |
| PA14_45940 | lasI | AGCAAGGCGCTTCCTTGCCGTGCAGAAGCT | TTTCGTCCGCGAGTGACCAGAACGCGTAGGCTAGAAAAACGCGTAGGCTAGAAAAGCAAGGCGCTTCCTTGCCGTGCAGAAGCTAAAACGCGTAGGCTAGAAAAACGCGTAGGCTAGACAACGTCCATGTCGGGATGC |
| PA14_45940 | lasI | CTCTTCGCGCCGACCAATTTGTACGATCAT | TTTCGTCCGCGAGTGACCAGAACGCGTAGGCTAGAAAAACGCGTAGGCTAGAAACTCTTCGCGCCGACCAATTTGTACGATCATAAAACGCGTAGGCTAGAAAAACGCGTAGGCTAGACAACGTCCATGTCGGGATGC |
| PA14_45940 | lasI | CGATACGTCCAGGCCGGCACGGATCATCAT | TTTCGTCCGCGAGTGACCAGAACGCGTAGGCTAGAAAAACGCGTAGGCTAGAAACGATACGTCCAGGCCGGCACGGATCATCATAAAACGCGTAGGCTAGAAAAACGCGTAGGCTAGACAACGTCCATGTCGGGATGC |
| PA14_45940 | lasI | CCCAGCCTTTGCGCTCCTTGAACACTTGAG | TTTCGTCCGCGAGTGACCAGAACGCGTAGGCTAGAAAAACGCGTAGGCTAGAAACCCAGCCTTTGCGCTCCTTGAACACTTGAGAAAACGCGTAGGCTAGAAAAACGCGTAGGCTAGACAACGTCCATGTCGGGATGC |
| PA14_45940 | lasI | GGGAAGGTGTTCTTCAGCATGTAGGGGCCA | TTTCGTCCGCGAGTGACCAGAACGCGTAGGCTAGAAAAACGCGTAGGCTAGAAAGGGAAGGTGTTCTTCAGCATGTAGGGGCCAAAAACGCGTAGGCTAGAAAAACGCGTAGGCTAGACAACGTCCATGTCGGGATGC |
| PA14_45940 | lasI | TGGCGAAACGGCTGAGTTCCCAGATGTGCG | TTTCGTCCGCGAGTGACCAGAACGCGTAGGCTAGAAAAACGCGTAGGCTAGAAATGGCGAAACGGCTGAGTTCCCAGATGTGCGAAAACGCGTAGGCTAGAAAAACGCGTAGGCTAGACAACGTCCATGTCGGGATGC |
| PA14_45940 | lasI | AAACCGCCAGTCGCTGTTCCACCAGCACTC | TTTCGTCCGCGAGTGACCAGAACGCGTAGGCTAGAAAAACGCGTAGGCTAGAAAAAACCGCCAGTCGCTGTTCCACCAGCACTCAAAACGCGTAGGCTAGAAAAACGCGTAGGCTAGACAACGTCCATGTCGGGATGC |
| PA14_45940 | lasI | TCTCCACCCCTACGGTGGTTACCGTCACCA | TTTCGTCCGCGAGTGACCAGAACGCGTAGGCTAGAAAAACGCGTAGGCTAGAAATCTCCACCCCTACGGTGGTTACCGTCACCAAAAACGCGTAGGCTAGAAAAACGCGTAGGCTAGACAACGTCCATGTCGGGATGC |
| PA14_45940 | lasI | GGGCTTCAGGAGTATCTTCCTGGATCAACA | TTTCGTCCGCGAGTGACCAGAACGCGTAGGCTAGAAAAACGCGTAGGCTAGAAAGGGCTTCAGGAGTATCTTCCTGGATCAACAAAAACGCGTAGGCTAGAAAAACGCGTAGGCTAGACAACGTCCATGTCGGGATGC |
| PA14_45940 | lasI | TCTGGATGTCGTTCTGCAGGCTGTAGCGGG | TTTCGTCCGCGAGTGACCAGAACGCGTAGGCTAGAAAAACGCGTAGGCTAGAAATCTGGATGTCGTTCTGCAGGCTGTAGCGGGAAAACGCGTAGGCTAGAAAAACGCGTAGGCTAGACAACGTCCATGTCGGGATGC |
| PA14_45940 | lasI | AAAGCCCAGCGAGCCTTTCTGTCCAGAGTT | TTTCGTCCGCGAGTGACCAGAACGCGTAGGCTAGAAAAACGCGTAGGCTAGAAAAAAGCCCAGCGAGCCTTTCTGTCCAGAGTTAAAACGCGTAGGCTAGAAAAACGCGTAGGCTAGACAACGTCCATGTCGGGATGC |
| PA14_45940 | lasI | GCGTCATAACCATCGATTTCCATCTCGTCG | TTTCGTCCGCGAGTGACCAGAACGCGTAGGCTAGAAAAACGCGTAGGCTAGAAAGCGTCATAACCATCGATTTCCATCTCGTCGAAAACGCGTAGGCTAGAAAAACGCGTAGGCTAGACAACGTCCATGTCGGGATGC |
| PA14_45940 | lasI | TGGCATTGAGTTCGATGCGCAAGGCCACCG | TTTCGTCCGCGAGTGACCAGAACGCGTAGGCTAGAAAAACGCGTAGGCTAGAAATGGCATTGAGTTCGATGCGCAAGGCCACCGAAAACGCGTAGGCTAGAAAAACGCGTAGGCTAGACAACGTCCATGTCGGGATGC |
| PA14_45940 | lasI | GGTATCGAGAATTCGCCAGCAACCGAAAAC | TTTCGTCCGCGAGTGACCAGAACGCGTAGGCTAGAAAAACGCGTAGGCTAGAAAGGTATCGAGAATTCGCCAGCAACCGAAAACAAAACGCGTAGGCTAGAAAAACGCGTAGGCTAGACAACGTCCATGTCGGGATGC |
| PA14_45960 | lasR | ATCTCCCAACTGGTCTTGCCGATGGCGCAC | TTTCGTCCGCGAGTGACCAGTTGACGAAGCCTGACAATTGACGAAGCCTGACAAATCTCCCAACTGGTCTTGCCGATGGCGCACAATTGACGAAGCCTGACAATTGACGAAGCCTGACCAACGTCCATGTCGGGATGC |
| PA14_45960 | lasR | CCGAACTTCCGCCGAATATTTCCCATATGG | TTTCGTCCGCGAGTGACCAGTTGACGAAGCCTGACAATTGACGAAGCCTGACAACCGAACTTCCGCCGAATATTTCCCATATGGAATTGACGAAGCCTGACAATTGACGAAGCCTGACCAACGTCCATGTCGGGATGC |
| PA14_45960 | lasR | CCTCGAAGAACTCGTGCTGCTTTCGCGTCT | TTTCGTCCGCGAGTGACCAGTTGACGAAGCCTGACAATTGACGAAGCCTGACAACCTCGAAGAACTCGTGCTGCTTTCGCGTCTAATTGACGAAGCCTGACAATTGACGAAGCCTGACCAACGTCCATGTCGGGATGC |
| PA14_45960 | lasR | TGGGTACAGTGACTGACCGTTGGGTCGACC | TTTCGTCCGCGAGTGACCAGTTGACGAAGCCTGACAATTGACGAAGCCTGACAATGGGTACAGTGACTGACCGTTGGGTCGACCAATTGACGAAGCCTGACAATTGACGAAGCCTGACCAACGTCCATGTCGGGATGC |
| PA14_45960 | lasR | CATTGGCTTCCGAGCAGTTGCAGATAACCG | TTTCGTCCGCGAGTGACCAGTTGACGAAGCCTGACAATTGACGAAGCCTGACAACATTGGCTTCCGAGCAGTTGCAGATAACCGAATTGACGAAGCCTGACAATTGACGAAGCCTGACCAACGTCCATGTCGGGATGC |
| PA14_45960 | lasR | ATTAACGGCCATAATGGCCGCTACGCGGCG | TTTCGTCCGCGAGTGACCAGTTGACGAAGCCTGACAATTGACGAAGCCTGACAAATTAACGGCCATAATGGCCGCTACGCGGCGAATTGACGAAGCCTGACAATTGACGAAGCCTGACCAACGTCCATGTCGGGATGC |
| PA14_45960 | lasR | TGGACGGTTCCCAGAAAATCGGCAGTACGC | TTTCGTCCGCGAGTGACCAGTTGACGAAGCCTGACAATTGACGAAGCCTGACAATGGACGGTTCCCAGAAAATCGGCAGTACGCAATTGACGAAGCCTGACAATTGACGAAGCCTGACCAACGTCCATGTCGGGATGC |
| PA14_45960 | lasR | GAATCCAAGGTCGCTCGCCATCTTCTGCAG | TTTCGTCCGCGAGTGACCAGTTGACGAAGCCTGACAATTGACGAAGCCTGACAAGAATCCAAGGTCGCTCGCCATCTTCTGCAGAATTGACGAAGCCTGACAATTGACGAAGCCTGACCAACGTCCATGTCGGGATGC |
| PA14_45960 | lasR | TTGCTGACCGGATGTTCGAAGGCCAGTCCG | TTTCGTCCGCGAGTGACCAGTTGACGAAGCCTGACAATTGACGAAGCCTGACAATTGCTGACCGGATGTTCGAAGGCCAGTCCGAATTGACGAAGCCTGACAATTGACGAAGCCTGACCAACGTCCATGTCGGGATGC |
| PA14_45960 | lasR | GCGCTCCACTCCAATTTTCCACTTGAGCGT | TTTCGTCCGCGAGTGACCAGTTGACGAAGCCTGACAATTGACGAAGCCTGACAAGCGCTCCACTCCAATTTTCCACTTGAGCGTAATTGACGAAGCCTGACAATTGACGAAGCCTGACCAACGTCCATGTCGGGATGC |
| PA14_45960 | lasR | CAGCTCAAGAAAACCGTCAACCAAGGCCAT | TTTCGTCCGCGAGTGACCAGTTGACGAAGCCTGACAATTGACGAAGCCTGACAACAGCTCAAGAAAACCGTCAACCAAGGCCATAATTGACGAAGCCTGACAATTGACGAAGCCTGACCAACGTCCATGTCGGGATGC |
| PA14_45960 | lasR | ACTTCCTTCTCCCGGCTGGTCAGAACCACC | TTTCGTCCGCGAGTGACCAGTTGACGAAGCCTGACAATTGACGAAGCCTGACAAACTTCCTTCTCCCGGCTGGTCAGAACCACCAATTGACGAAGCCTGACAATTGACGAAGCCTGACCAACGTCCATGTCGGGATGC |
| PA14_45960 | lasR | AGTCCTTGAGCATCCACAGGGTCGGCAGGA | TTTCGTCCGCGAGTGACCAGTTGACGAAGCCTGACAATTGACGAAGCCTGACAAAGTCCTTGAGCATCCACAGGGTCGGCAGGAAATTGACGAAGCCTGACAATTGACGAAGCCTGACCAACGTCCATGTCGGGATGC |
| PA14_45960 | lasR | TCCTTAGGCAACAGGCCGAAGAGGATCTTC | TTTCGTCCGCGAGTGACCAGTTGACGAAGCCTGACAATTGACGAAGCCTGACAATCCTTAGGCAACAGGCCGAAGAGGATCTTCAATTGACGAAGCCTGACAATTGACGAAGCCTGACCAACGTCCATGTCGGGATGC |
| PA14_45960 | lasR | ATGAAACGGTTGGCCTCGGCCCGGTTTTCC | TTTCGTCCGCGAGTGACCAGTTGACGAAGCCTGACAATTGACGAAGCCTGACAAATGAAACGGTTGGCCTCGGCCCGGTTTTCCAATTGACGAAGCCTGACAATTGACGAAGCCTGACCAACGTCCATGTCGGGATGC |
| PA14_41220 | lon | CCTGGACCTGCTTCGGAATCAGGTACTTGG | TTTCGTCCGCGAGTGACCAGACTACGCGCGGATATAAACTACGCGCGGATATAACCTGGACCTGCTTCGGAATCAGGTACTTGGAAACTACGCGCGGATATAAACTACGCGCGGATATCAACGTCCATGTCGGGATGC |
| PA14_41220 | lon | CAGCCAGTCTATGTAGGAACGCACCACGGT | TTTCGTCCGCGAGTGACCAGACTACGCGCGGATATAAACTACGCGCGGATATAACAGCCAGTCTATGTAGGAACGCACCACGGTAAACTACGCGCGGATATAAACTACGCGCGGATATCAACGTCCATGTCGGGATGC |
| PA14_41220 | lon | TCTTCAGCGCCATGTGCGCGGCCATGGTAT | TTTCGTCCGCGAGTGACCAGACTACGCGCGGATATAAACTACGCGCGGATATAATCTTCAGCGCCATGTGCGCGGCCATGGTATAAACTACGCGCGGATATAAACTACGCGCGGATATCAACGTCCATGTCGGGATGC |
| PA14_41220 | lon | TTCGGCGAGGCCATAACGGAACTTGCGCAC | TTTCGTCCGCGAGTGACCAGACTACGCGCGGATATAAACTACGCGCGGATATAATTCGGCGAGGCCATAACGGAACTTGCGCACAAACTACGCGCGGATATAAACTACGCGCGGATATCAACGTCCATGTCGGGATGC |
| PA14_41220 | lon | AGGCGGATGACTTCCATCCGATCCAGCAGG | TTTCGTCCGCGAGTGACCAGACTACGCGCGGATATAAACTACGCGCGGATATAAAGGCGGATGACTTCCATCCGATCCAGCAGGAAACTACGCGCGGATATAAACTACGCGCGGATATCAACGTCCATGTCGGGATGC |
| PA14_41220 | lon | GACTTTTCCCGACCGACGAACAACGGGATG | TTTCGTCCGCGAGTGACCAGACTACGCGCGGATATAAACTACGCGCGGATATAAGACTTTTCCCGACCGACGAACAACGGGATGAAACTACGCGCGGATATAAACTACGCGCGGATATCAACGTCCATGTCGGGATGC |
| PA14_41220 | lon | TCCTTTGCAACCATCTCCGGAGCCGCATCG | TTTCGTCCGCGAGTGACCAGACTACGCGCGGATATAAACTACGCGCGGATATAATCCTTTGCAACCATCTCCGGAGCCGCATCGAAACTACGCGCGGATATAAACTACGCGCGGATATCAACGTCCATGTCGGGATGC |
| PA14_41220 | lon | ATCCGGCAGCTTGAGCAACTGCAGCACGGT | TTTCGTCCGCGAGTGACCAGACTACGCGCGGATATAAACTACGCGCGGATATAAATCCGGCAGCTTGAGCAACTGCAGCACGGTAAACTACGCGCGGATATAAACTACGCGCGGATATCAACGTCCATGTCGGGATGC |

| | | | |
|---|---|---|---|
| PA14_41220 | lon | TTGTTCAGCTCGGCAGTGGCCTTGGTGTGC | TTTCGTCCGCGAGTGACCAGACTACGCGCGGATATAAACTACGCGCGGATATAATTGTTCAGCTCGGCAGTGGCCTTGGTGTGCAAACTACGCGCGGATATAAACTACGCGCGGATATCAACGTCCATGTCGGGATGC |
| PA14_41220 | lon | CTGGGATGTTCATCGAGTTTGCGGTGCAGA | TTTCGTCCGCGAGTGACCAGACTACGCGCGGATATAAACTACGCGCGGATATAACTGGGATGTTCATCGAGTTTGCGGTGCAGAAAACTACGCGCGGATATAAACTACGCGCGGATATCAACGTCCATGTCGGGATGC |
| PA14_41220 | lon | TATAGGTACGACGGTGACCGCGGATCTCGG | TTTCGTCCGCGAGTGACCAGACTACGCGCGGATATAAACTACGCGCGGATATAATATAGGTACGACGGTGACCGCGGATCTCGGAAACTACGCGCGGATATAAACTACGCGCGGATATCAACGTCCATGTCGGGATGC |
| PA14_41220 | lon | ACCTTGGTCATCTTCTGGATCAGGCGGCCC | TTTCGTCCGCGAGTGACCAGACTACGCGCGGATATAAACTACGCGCGGATATAAACCTTGGTCATCTTCTGGATCAGGCGGCCCAAACTACGCGCGGATATAAACTACGCGCGGATATCAACGTCCATGTCGGGATGC |
| PA14_41220 | lon | ATCATGGCGCACCTTGCTCTCGGCCTTCCA | TTTCGTCCGCGAGTGACCAGACTACGCGCGGATATAAACTACGCGCGGATATAAATCATGGCGCACCTTGCTCTCGGCCTTCCAAAACTACGCGCGGATATAAACTACGCGCGGATATCAACGTCCATGTCGGGATGC |
| PA14_41220 | lon | GCAGCAAGGGCAATTCGACGAGTGTTTTCA | TTTCGTCCGCGAGTGACCAGACTACGCGCGGATATAAACTACGCGCGGATATAAGCAGCAAGGGCAATTCGACGAGTGTTTTCAAAACTACGCGCGGATATAAACTACGCGCGGATATCAACGTCCATGTCGGGATGC |
| PA14_41220 | lon | TTCTCGTGGAAGTCCGCCGCGATTCCTAGG | TTTCGTCCGCGAGTGACCAGACTACGCGCGGATATAAACTACGCGCGGATATAATTCTCGTGGAAGTCCGCCGCGATTCCTAGGAAACTACGCGCGGATATAAACTACGCGCGGATATCAACGTCCATGTCGGGATGC |
| PA14_41220 | lon | CGCCTTCCGGAACGTGGATATGGATGTCAC | TTTCGTCCGCGAGTGACCAGACTACGCGCGGATATAAACTACGCGCGGATATAACGCCTTCCGGAACGTGGATATGGATGTCACAAACTACGCGCGGATATAAACTACGCGCGGATATCAACGTCCATGTCGGGATGC |
| PA14_41220 | lon | GTCCGGAATTTCCTTCAGGTCGCGAACATT | TTTCGTCCGCGAGTGACCAGACTACGCGCGGATATAAACTACGCGCGGATATAAGTCCGGAATTTCCTTCAGGTCGCGAACATTAAACTACGCGCGGATATAAACTACGCGCGGATATCAACGTCCATGTCGGGATGC |
| PA14_41220 | lon | AGATACTCGAGAATGCGCTCCTTGACCTCT | TTTCGTCCGCGAGTGACCAGACTACGCGCGGATATAAACTACGCGCGGATATAAAGATACTCGAGAATGCGCTCCTTGACCTCTAAACTACGCGCGGATATAAACTACGCGCGGATATCAACGTCCATGTCGGGATGC |
| PA14_41220 | lon | ATTCGCTCCTTGGAATCAGGCTCGCGTTTT | TTTCGTCCGCGAGTGACCAGACTACGCGCGGATATAAACTACGCGCGGATATAAATTCGCTCCTTGGAATCAGGCTCGCGTTTTAAACTACGCGCGGATATAAACTACGCGCGGATATCAACGTCCATGTCGGGATGC |
| PA14_41220 | lon | CATGTGCGGGTACACCACCACGTCACGTAG | TTTCGTCCGCGAGTGACCAGACTACGCGCGGATATAAACTACGCGCGGATATAACATGTGCGGGTACACCACCACGTCACGTAGAAACTACGCGCGGATATAAACTACGCGCGGATATCAACGTCCATGTCGGGATGC |
| PA14_52270 | ldhA | AAGCCGGTCAGCCCATGCAGCGAGAAATCG | TTTCGTCCGCGAGTGACCAGTTCCACTGCGTAGTGAATTCCACTGCGTAGTGAAAAGCCGGTCAGCCCATGCAGCGAGAAATCGAATTCCACTGCGTAGTGAATTCCACTGCGTAGTGCAACGTCCATGTCGGGATGC |
| PA14_52270 | ldhA | GGTCGGAGCGATCCTCGAAGAAGATATCCG | TTTCGTCCGCGAGTGACCAGTTCCACTGCGTAGTGAATTCCACTGCGTAGTGAAGGTCGGAGCGATCCTCGAAGAAGATATCCGAATTCCACTGCGTAGTGAATTCCACTGCGTAGTGCAACGTCCATGTCGGGATGC |
| PA14_52270 | ldhA | CGATCACTCCGACGCGTTTGCCGTGGAGGT | TTTCGTCCGCGAGTGACCAGTTCCACTGCGTAGTGAATTCCACTGCGTAGTGAACGATCACTCCGACGCGTTTGCCGTGGAGGTAATTCCACTGCGTAGTGAATTCCACTGCGTAGTGCAACGTCCATGTCGGGATGC |
| PA14_52270 | ldhA | TGATCAGCATCGCGCCGGGTTTCATCGTCG | TTTCGTCCGCGAGTGACCAGTTCCACTGCGTAGTGAATTCCACTGCGTAGTGAATGATCAGCATCGCGCCGGGTTTCATCGTCGAATTCCACTGCGTAGTGAATTCCACTGCGTAGTGCAACGTCCATGTCGGGATGC |
| PA14_52270 | ldhA | ACCACCACGTTGGGGAAGCTCAGCAGGCGT | TTTCGTCCGCGAGTGACCAGTTCCACTGCGTAGTGAATTCCACTGCGTAGTGAAACCACCACGTTGGGGAAGCTCAGCAGGCGTAATTCCACTGCGTAGTGAATTCCACTGCGTAGTGCAACGTCCATGTCGGGATGC |
| PA14_52270 | ldhA | TTGGGATAGGGATCGTAGGCCAGCAGCTCG | TTTCGTCCGCGAGTGACCAGTTCCACTGCGTAGTGAATTCCACTGCGTAGTGAATTGGGATAGGGATCGTAGGCCAGCAGCTCGAATTCCACTGCGTAGTGAATTCCACTGCGTAGTGCAACGTCCATGTCGGGATGC |
| PA14_52270 | ldhA | TTGCTGGCCTGGAAGCTCTCGCTGTCGTAG | TTTCGTCCGCGAGTGACCAGTTCCACTGCGTAGTGAATTCCACTGCGTAGTGAATTGCTGGCCTGGAAGCTCTCGCTGTCGTAGAATTCCACTGCGTAGTGAATTCCACTGCGTAGTGCAACGTCCATGTCGGGATGC |
| PA14_52270 | ldhA | ACGAAGGCGCAGACTACTTCGAAGCCCTGG | TTTCGTCCGCGAGTGACCAGTTCCACTGCGTAGTGAATTCCACTGCGTAGTGAAACGAAGGCGCAGACTACTTCGAAGCCCTGGAATTCCACTGCGTAGTGAATTCCACTGCGTAGTGCAACGTCCATGTCGGGATGC |
| PA14_52270 | ldhA | TCCTCTTCATAGACGTCCAGGCCGAGGTAG | TTTCGTCCGCGAGTGACCAGTTCCACTGCGTAGTGAATTCCACTGCGTAGTGAATCCTCTTCATAGACGTCCAGGCCGAGGTAGAATTCCACTGCGTAGTGAATTCCACTGCGTAGTGCAACGTCCATGTCGGGATGC |
| PA14_52270 | ldhA | AGGTGGGCCTGCTGGAAGTGCAGTTCGAAG | TTTCGTCCGCGAGTGACCAGTTCCACTGCGTAGTGAATTCCACTGCGTAGTGAAAGGTGGGCCTGCTGGAAGTGCAGTTCGAAGAATTCCACTGCGTAGTGAATTCCACTGCGTAGTGCAACGTCCATGTCGGGATGC |
| PA14_52270 | ldhA | CGATGTTGTCCAGGGTGGTGTCGGCGATCG | TTTCGTCCGCGAGTGACCAGTTCCACTGCGTAGTGAATTCCACTGCGTAGTGAACGATGTTGTCCAGGGTGGTGTCGGCGATCGAATTCCACTGCGTAGTGAATTCCACTGCGTAGTGCAACGTCCATGTCGGGATGC |
| PA14_52270 | ldhA | AATGCAGGCTGACGATGTCGGACTCGGCGA | TTTCGTCCGCGAGTGACCAGTTCCACTGCGTAGTGAATTCCACTGCGTAGTGAAAATGCAGGCTGACGATGTCGGACTCGGCGAAATTCCACTGCGTAGTGAATTCCACTGCGTAGTGCAACGTCCATGTCGGGATGC |
| PA14_52580 | lysC | TACGCACCTGGTTGCCGGTATAGGAGACCG | TTTCGTCCGCGAGTGACCAGATTGTGCGAGGTCCAAAATTGTGCGAGGTCCAAATACGCACCTGGTTGCCGGTATAGGAGACCGAAATTGTGCGAGGTCCAAAATTGTGCGAGGTCCACAACGTCCATGTCGGGATGC |
| PA14_52580 | lysC | GGCTGTTCCATGGATTCCTCTTCATCATCG | TTTCGTCCGCGAGTGACCAGATTGTGCGAGGTCCAAAATTGTGCGAGGTCCAAAGGCTGTTCCATGGATTCCTCTTCATCATCGAAATTGTGCGAGGTCCAAAATTGTGCGAGGTCCACAACGTCCATGTCGGGATGC |
| PA14_52580 | lysC | CGCCAGTTCCAGGTACTTCTCTTCGATCAC | TTTCGTCCGCGAGTGACCAGATTGTGCGAGGTCCAAAATTGTGCGAGGTCCAAACGCCAGTTCCAGGTACTTCTCTTCGATCACAAATTGTGCGAGGTCCAAAATTGTGCGAGGTCCACAACGTCCATGTCGGGATGC |
| PA14_52580 | lysC | TTGATCAGAGCCATGCTGAGCAGGGCGATG | TTTCGTCCGCGAGTGACCAGATTGTGCGAGGTCCAAAATTGTGCGAGGTCCAAATTGATCAGAGCCATGCTGAGCAGGGCGATGAAATTGTGCGAGGTCCAAAATTGTGCGAGGTCCACAACGTCCATGTCGGGATGC |
| PA14_52580 | lysC | GTTCAGGTAATCGTTGCGATGCACCGTGAA | TTTCGTCCGCGAGTGACCAGATTGTGCGAGGTCCAAAATTGTGCGAGGTCCAAAGTTCAGGTAATCGTTGCGATGCACCGTGAAAAATTGTGCGAGGTCCAAAATTGTGCGAGGTCCACAACGTCCATGTCGGGATGC |
| PA14_52580 | lysC | TCACCTTCTCGGCCACCTGCTCGATACGCT | TTTCGTCCGCGAGTGACCAGATTGTGCGAGGTCCAAAATTGTGCGAGGTCCAAATCACCTTCTCGGCCACCTGCTCGATACGCTAAATTGTGCGAGGTCCAAAATTGTGCGAGGTCCACAACGTCCATGTCGGGATGC |
| PA14_52580 | lysC | TGAAGTCGGTGGTGTTATCGTGCGCCACGT | TTTCGTCCGCGAGTGACCAGATTGTGCGAGGTCCAAAATTGTGCGAGGTCCAAATGAAGTCGGTGGTGTTATCGTGCGCCACGTAAATTGTGCGAGGTCCAAAATTGTGCGAGGTCCACAACGTCCATGTCGGGATGC |
| PA14_52580 | lysC | ATGTTGCCGTTGCCGTCGACGCCCTGGAAA | TTTCGTCCGCGAGTGACCAGATTGTGCGAGGTCCAAAATTGTGCGAGGTCCAAAATGTTGCCGTTGCCGTCGACGCCCTGGAAAAAATTGTGCGAGGTCCAAAATTGTGCGAGGTCCACAACGTCCATGTCGGGATGC |
| PA14_52580 | lysC | AGTAATGAGGGTGCCCGGACCCTCCTGGAA | TTTCGTCCGCGAGTGACCAGATTGTGCGAGGTCCAAAATTGTGCGAGGTCCAAAAGTAATGAGGGTGCCCGGACCCTCCTGGAAAAATTGTGCGAGGTCCAAAATTGTGCGAGGTCCACAACGTCCATGTCGGGATGC |
| PA14_52580 | lysC | TGCTCCATGATCTGGTTGGCCAGGCCGATC | TTTCGTCCGCGAGTGACCAGATTGTGCGAGGTCCAAAATTGTGCGAGGTCCAAATGCTCCATGATCTGGTTGGCCAGGCCGATCAAATTGTGCGAGGTCCAAAATTGTGCGAGGTCCACAACGTCCATGTCGGGATGC |
| PA14_52580 | lysC | GGTATCGGGTACGCCACGGATGGTCAGCTT | TTTCGTCCGCGAGTGACCAGATTGTGCGAGGTCCAAAATTGTGCGAGGTCCAAAGGTATCGGGTACGCCACGGATGGTCAGCTTAAATTGTGCGAGGTCCAAAATTGTGCGAGGTCCACAACGTCCATGTCGGGATGC |
| PA14_52580 | lysC | GTCCCTCCAAACTTCTGTACGATCAAAGCC | TTTCGTCCGCGAGTGACCAGATTGTGCGAGGTCCAAAATTGTGCGAGGTCCAAAGTCCCTCCAAACTTCTGTACGATCAAAGCCAAATTGTGCGAGGTCCAAAATTGTGCGAGGTCCACAACGTCCATGTCGGGATGC |
| PA14_52580 | lysC | TCGAGACCATGACGTCCAGTTCGCGCGGAA | TTTCGTCCGCGAGTGACCAGATTGTGCGAGGTCCAAAATTGTGCGAGGTCCAAATCGAGACCATGACGTCCAGTTCGCGCGGAAAAATTGTGCGAGGTCCAAAATTGTGCGAGGTCCACAACGTCCATGTCGGGATGC |
| PA14_52580 | lysC | ATACCACCACGACGACGTCATCGCCCGCTT | TTTCGTCCGCGAGTGACCAGATTGTGCGAGGTCCAAAATTGTGCGAGGTCCAAAATACCACCACGACGACGTCATCGCCCGCTTAAATTGTGCGAGGTCCAAAATTGTGCGAGGTCCACAACGTCCATGTCGGGATGC |
| PA14_52580 | lysC | TCGATGTGCAGGATGCGCGCCTTGGTGTGT | TTTCGTCCGCGAGTGACCAGATTGTGCGAGGTCCAAAATTGTGCGAGGTCCAAATCGATGTGCAGGATGCGCGCCTTGGTGTGTAAATTGTGCGAGGTCCAAAATTGTGCGAGGTCCACAACGTCCATGTCGGGATGC |
| PA14_05540 | mexB | CTGGGAACCGTCGGGATTGACCTTGAGCAG | TTTCGTCCGCGAGTGACCAGTCAATTGGGCGCTCAAATCAATTGGGCGCTCAAACTGGGAACCGTCGGGATTGACCTTGAGCAGAATCAATTGGGCGCTCAAATCAATTGGGCGCTCACAACGTCCATGTCGGGATGC |
| PA14_05540 | mexB | GTCGTTGACGTAGCTGGAACCCCAGGCGAT | TTTCGTCCGCGAGTGACCAGTCAATTGGGCGCTCAAATCAATTGGGCGCTCAAAGTCGTTGACGTAGCTGGAACCCCAGGCGATAATCAATTGGGCGCTCAAATCAATTGGGCGCTCACAACGTCCATGTCGGGATGC |
| PA14_05540 | mexB | CGAGCTGACCGAAGAGCTTTCCTTCTCCAG | TTTCGTCCGCGAGTGACCAGTCAATTGGGCGCTCAAATCAATTGGGCGCTCAAACGAGCTGACCGAAGAGCTTTCCTTCTCCAGAATCAATTGGGCGCTCAAATCAATTGGGCGCTCACAACGTCCATGTCGGGATGC |
| PA14_05540 | mexB | CACCCACGCGAAAATGGGCCTATCAATGAA | TTTCGTCCGCGAGTGACCAGTCAATTGGGCGCTCAAATCAATTGGGCGCTCAAACACCCACGCGAAAATGGGCCTATCAATGAAAATCAATTGGGCGCTCAAATCAATTGGGCGCTCACAACGTCCATGTCGGGATGC |
| PA14_05540 | mexB | TTTCGACGGACGCTTGCTGCTTGGACGCCT | TTTCGTCCGCGAGTGACCAGTCAATTGGGCGCTCAAATCAATTGGGCGCTCAAATTTCGACGGACGCTTGCTGCTTGGACGCCTAATCAATTGGGCGCTCAAATCAATTGGGCGCTCACAACGTCCATGTCGGGATGC |
| PA14_05540 | mexB | GAACTCCACGATGAGAATGGCGTTCTTCGC | TTTCGTCCGCGAGTGACCAGTCAATTGGGCGCTCAAATCAATTGGGCGCTCAAAGAACTCCACGATGAGAATGGCGTTCTTCGCAATCAATTGGGCGCTCAAATCAATTGGGCGCTCACAACGTCCATGTCGGGATGC |
| PA14_05540 | mexB | AGATCAGCAGGTACGGCGCGCGATGCTTGA | TTTCGTCCGCGAGTGACCAGTCAATTGGGCGCTCAAATCAATTGGGCGCTCAAAAGATCAGCAGGTACGGCGCGCGATGCTTGAAATCAATTGGGCGCTCAAATCAATTGGGCGCTCACAACGTCCATGTCGGGATGC |
| PA14_05540 | mexB | TGGTTGACCGGCAGACTGAGGATCGACAGG | TTTCGTCCGCGAGTGACCAGTCAATTGGGCGCTCAAATCAATTGGGCGCTCAAATGGTTGACCGGCAGACTGAGGATCGACAGGAATCAATTGGGCGCTCAAATCAATTGGGCGCTCACAACGTCCATGTCGGGATGC |
| PA14_05540 | mexB | ATGGAGAACTGCCGGTAGATCACCCCGGTG | TTTCGTCCGCGAGTGACCAGTCAATTGGGCGCTCAAATCAATTGGGCGCTCAAAATGGAGAACTGCCGGTAGATCACCCCGGTGAATCAATTGGGCGCTCAAATCAATTGGGCGCTCACAACGTCCATGTCGGGATGC |
| PA14_05540 | mexB | ACACGCTGTTCTCACCACCGGGACGCTCTT | TTTCGTCCGCGAGTGACCAGTCAATTGGGCGCTCAAATCAATTGGGCGCTCAAAACACGCTGTTCTCACCACCGGGACGCTCTTAATCAATTGGGCGCTCAAATCAATTGGGCGCTCACAACGTCCATGTCGGGATGC |
| PA14_05540 | mexB | TTCGGCGAACCGTATTCCCACTTGCCGGTG | TTTCGTCCGCGAGTGACCAGTCAATTGGGCGCTCAAATCAATTGGGCGCTCAAATTCGGCGAACCGTATTCCCACTTGCCGGTGAATCAATTGGGCGCTCAAATCAATTGGGCGCTCACAACGTCCATGTCGGGATGC |
| PA14_05540 | mexB | AAGATCGCCAGGACGGTCGCAGTGACCATG | TTTCGTCCGCGAGTGACCAGTCAATTGGGCGCTCAAATCAATTGGGCGCTCAAAAAGATCGCCAGGACGGTCGCAGTGACCATGAATCAATTGGGCGCTCAAATCAATTGGGCGCTCACAACGTCCATGTCGGGATGC |
| PA14_05540 | mexB | GCGAGAGCGGGTCCTGGATGTTGGAAACGA | TTTCGTCCGCGAGTGACCAGTCAATTGGGCGCTCAAATCAATTGGGCGCTCAAAGCGAGAGCGGGTCCTGGATGTTGGAAACGAAATCAATTGGGCGCTCAAATCAATTGGGCGCTCACAACGTCCATGTCGGGATGC |
| PA14_05540 | mexB | TGTTCTCGAATTGCTCCGCGGTCTGCAGGC | TTTCGTCCGCGAGTGACCAGTCAATTGGGCGCTCAAATCAATTGGGCGCTCAAATGTTCTCGAATTGCTCCGCGGTCTGCAGGCAATCAATTGGGCGCTCAAATCAATTGGGCGCTCACAACGTCCATGTCGGGATGC |
| PA14_05540 | mexB | TGTACTGCGGTTCGTCGCTCATGCCGTTGG | TTTCGTCCGCGAGTGACCAGTCAATTGGGCGCTCAAATCAATTGGGCGCTCAAATGTACTGCGGTTCGTCGCTCATGCCGTTGGAATCAATTGGGCGCTCAAATCAATTGGGCGCTCACAACGTCCATGTCGGGATGC |
| PA14_05540 | mexB | AACACGTCGTTAGACAGGCCGCGCATGGAC | TTTCGTCCGCGAGTGACCAGTCAATTGGGCGCTCAAATCAATTGGGCGCTCAAAAACACGTCGTTAGACAGGCCGCGCATGGACAATCAATTGGGCGCTCAAATCAATTGGGCGCTCACAACGTCCATGTCGGGATGC |
| PA14_05540 | mexB | TCGAGCCAGATGCGCATCGAGTACTGCGAG | TTTCGTCCGCGAGTGACCAGTCAATTGGGCGCTCAAATCAATTGGGCGCTCAAATCGAGCCAGATGCGCATCGAGTACTGCGAGAATCAATTGGGCGCTCAAATCAATTGGGCGCTCACAACGTCCATGTCGGGATGC |
| PA14_05540 | mexB | TGAAGAAGTGCATCTGCGCGCGCTTGGCCA | TTTCGTCCGCGAGTGACCAGTCAATTGGGCGCTCAAATCAATTGGGCGCTCAAATGAAGAAGTGCATCTGCGCGCGCTTGGCCAAATCAATTGGGCGCTCAAATCAATTGGGCGCTCACAACGTCCATGTCGGGATGC |
| PA14_05540 | mexB | GGAGATGTAGCGCAGATTGTCGATCCCGTT | TTTCGTCCGCGAGTGACCAGTCAATTGGGCGCTCAAATCAATTGGGCGCTCAAAGGAGATGTAGCGCAGATTGTCGATCCCGTTAATCAATTGGGCGCTCAAATCAATTGGGCGCTCACAACGTCCATGTCGGGATGC |
| PA14_05540 | mexB | CATCCAGATCATCCCGGCCACGATCACCAC | TTTCGTCCGCGAGTGACCAGTCAATTGGGCGCTCAAATCAATTGGGCGCTCAAACATCCAGATCATCCCGGCCACGATCACCACAATCAATTGGGCGCTCAAATCAATTGGGCGCTCACAACGTCCATGTCGGGATGC |
| PA14_32390 | mexF | GGTTGAAGTCGTTCGCGTACAGCGAGCCGA | TTTCGTCCGCGAGTGACCAGCTCTAGGATCGTACGAACTCTAGGATCGTACGAAGGTTGAAGTCGTTCGCGTACAGCGAGCCGAAACTCTAGGATCGTACGAACTCTAGGATCGTACGCAACGTCCATGTCGGGATGC |
| PA14_32390 | mexF | TGATCGGTAGCTGGAACAGGGAGATGGCGC | TTTCGTCCGCGAGTGACCAGCTCTAGGATCGTACGAACTCTAGGATCGTACGAATGATCGGTAGCTGGAACAGGGAGATGGCGCAACTCTAGGATCGTACGAACTCTAGGATCGTACGCAACGTCCATGTCGGGATGC |
| PA14_32390 | mexF | ATGACTTTCGGGTTGGCGCCGGGGAAGTTG | TTTCGTCCGCGAGTGACCAGCTCTAGGATCGTACGAACTCTAGGATCGTACGAAATGACTTTCGGGTTGGCGCCGGGGAAGTTGAACTCTAGGATCGTACGAACTCTAGGATCGTACGCAACGTCCATGTCGGGATGC |
| PA14_32390 | mexF | TGAGGACGTAGAACACCGGCGTCAGCAGCA | TTTCGTCCGCGAGTGACCAGCTCTAGGATCGTACGAACTCTAGGATCGTACGAATGAGGACGTAGAACACCGGCGTCAGCAGCAAACTCTAGGATCGTACGAACTCTAGGATCGTACGCAACGTCCATGTCGGGATGC |
| PA14_32390 | mexF | GACATGTAGAGCATGTTCTCCACCCCGGTG | TTTCGTCCGCGAGTGACCAGCTCTAGGATCGTACGAACTCTAGGATCGTACGAAGACATGTAGAGCATGTTCTCCACCCCGGTGAACTCTAGGATCGTACGAACTCTAGGATCGTACGCAACGTCCATGTCGGGATGC |
| PA14_32390 | mexF | GGCCGCTGGATGAAGAATTGGGAGAAATTC | TTTCGTCCGCGAGTGACCAGCTCTAGGATCGTACGAACTCTAGGATCGTACGAAGGCCGCTGGATGAAGAATTGGGAGAAATTCAACTCTAGGATCGTACGAACTCTAGGATCGTACGCAACGTCCATGTCGGGATGC |
| PA14_32390 | mexF | AGATTTCGATGGCGTTCGAGCCGGGACGCT | TTTCGTCCGCGAGTGACCAGCTCTAGGATCGTACGAACTCTAGGATCGTACGAAAGATTTCGATGGCGTTCGAGCCGGGACGCTAACTCTAGGATCGTACGAACTCTAGGATCGTACGCAACGTCCATGTCGGGATGC |
| PA14_32390 | mexF | AGCTCGGTCCACTCGTAGGTCATGCCGTTG | TTTCGTCCGCGAGTGACCAGCTCTAGGATCGTACGAACTCTAGGATCGTACGAAAGCTCGGTCCACTCGTAGGTCATGCCGTTGAACTCTAGGATCGTACGAACTCTAGGATCGTACGCAACGTCCATGTCGGGATGC |
| PA14_32390 | mexF | TTGTAGTGCATCACACGGTCCGGACCGGAG | TTTCGTCCGCGAGTGACCAGCTCTAGGATCGTACGAACTCTAGGATCGTACGAATTGTAGTGCATCACACGGTCCGGACCGGAGAACTCTAGGATCGTACGAACTCTAGGATCGTACGCAACGTCCATGTCGGGATGC |
| PA14_32390 | mexF | GACGATGGAGTAGTCCATGCCCTGCGGGAA | TTTCGTCCGCGAGTGACCAGCTCTAGGATCGTACGAACTCTAGGATCGTACGAAGACGATGGAGTAGTCCATGCCCTGCGGGAAAACTCTAGGATCGTACGAACTCTAGGATCGTACGCAACGTCCATGTCGGGATGC |
| PA14_32390 | mexF | TCCTGGATGTCGGCGTACTTGGCGTTCAGC | TTTCGTCCGCGAGTGACCAGCTCTAGGATCGTACGAACTCTAGGATCGTACGAATCCTGGATGTCGGCGTACTTGGCGTTCAGCAACTCTAGGATCGTACGAACTCTAGGATCGTACGCAACGTCCATGTCGGGATGC |
| PA14_32390 | mexF | TAGAACTGCCCGGTGAGGCCGGAGATGAAC | TTTCGTCCGCGAGTGACCAGCTCTAGGATCGTACGAACTCTAGGATCGTACGAATAGAACTGCCCGGTGAGGCCGGAGATGAACAACTCTAGGATCGTACGAACTCTAGGATCGTACGCAACGTCCATGTCGGGATGC |
| PA14_32390 | mexF | GATGAAGGCGATGGACGTCATCAGGATCGG | TTTCGTCCGCGAGTGACCAGCTCTAGGATCGTACGAACTCTAGGATCGTACGAAGATGAAGGCGATGGACGTCATCAGGATCGGAACTCTAGGATCGTACGAACTCTAGGATCGTACGCAACGTCCATGTCGGGATGC |
| PA14_32390 | mexF | TGAACAGCTCCTCGTAGCCCTGGTTGCCAC | TTTCGTCCGCGAGTGACCAGCTCTAGGATCGTACGAACTCTAGGATCGTACGAATGAACAGCTCCTCGTAGCCCTGGTTGCCACAACTCTAGGATCGTACGAACTCTAGGATCGTACGCAACGTCCATGTCGGGATGC |
| PA14_32390 | mexF | TGACCTTGATGAAGGACGCCAGCGGGACCA | TTTCGTCCGCGAGTGACCAGCTCTAGGATCGTACGAACTCTAGGATCGTACGAATGACCTTGATGAAGGACGCCAGCGGGACCAAACTCTAGGATCGTACGAACTCTAGGATCGTACGCAACGTCCATGTCGGGATGC |
| PA14_51340 | mvfR | TTGAAGCGGGAGATCTTCAGTTCCTCGTCG | TTTCGTCCGCGAGTGACCAGTGACTCGCACTGGGAAATGACTCGCACTGGGAAATTGAAGCGGGAGATCTTCAGTTCCTCGTCGAATGACTCGCACTGGGAAATGACTCGCACTGGGACAACGTCCATGTCGGGATGC |
| PA14_51340 | mvfR | AAATAATGCGGCGCGATGCCCCATCCGACG | TTTCGTCCGCGAGTGACCAGTGACTCGCACTGGGAAATGACTCGCACTGGGAAAAAATAATGCGGCGCGATGCCCCATCCGACGAATGACTCGCACTGGGAAATGACTCGCACTGGGACAACGTCCATGTCGGGATGC |
| PA14_51340 | mvfR | TACGCTGCTCACCGTATCGCAGAACGACGG | TTTCGTCCGCGAGTGACCAGTGACTCGCACTGGGAAATGACTCGCACTGGGAAATACGCTGCTCACCGTATCGCAGAACGACGGAATGACTCGCACTGGGAAATGACTCGCACTGGGACAACGTCCATGTCGGGATGC |
| PA14_51340 | mvfR | CCTGGAGGAACATGTTCACGTGATTCAGGT | TTTCGTCCGCGAGTGACCAGTGACTCGCACTGGGAAATGACTCGCACTGGGAAACCTGGAGGAACATGTTCACGTGATTCAGGTAATGACTCGCACTGGGAAATGACTCGCACTGGGACAACGTCCATGTCGGGATGC |
| PA14_51340 | mvfR | TGTGCAGGGAGGCATTGCACAACGGGTGCT | TTTCGTCCGCGAGTGACCAGTGACTCGCACTGGGAAATGACTCGCACTGGGAAATGTGCAGGGAGGCATTGCACAACGGGTGCTAATGACTCGCACTGGGAAATGACTCGCACTGGGACAACGTCCATGTCGGGATGC |
| PA14_51340 | mvfR | ATCAGGCTGACCATGTTGAAATCGTCGAGC | TTTCGTCCGCGAGTGACCAGTGACTCGCACTGGGAAATGACTCGCACTGGGAAAATCAGGCTGACCATGTTGAAATCGTCGAGCAATGACTCGCACTGGGAAATGACTCGCACTGGGACAACGTCCATGTCGGGATGC |
| PA14_51340 | mvfR | GATCTGCCGGTAATTGGCCAGGCTCGCGAT | TTTCGTCCGCGAGTGACCAGTGACTCGCACTGGGAAATGACTCGCACTGGGAAAGATCTGCCGGTAATTGGCCAGGCTCGCGATAATGACTCGCACTGGGAAATGACTCGCACTGGGACAACGTCCATGTCGGGATGC |
| PA14_51340 | mvfR | CGATTTCCAGGTTGCTGACCGCCGAGCTGA | TTTCGTCCGCGAGTGACCAGTGACTCGCACTGGGAAATGACTCGCACTGGGAAACGATTTCCAGGTTGCTGACCGCCGAGCTGAAATGACTCGCACTGGGAAATGACTCGCACTGGGACAACGTCCATGTCGGGATGC |
| PA14_51340 | mvfR | GTCGAAGTTTTCCACGAAGAGCACCTTGTC | TTTCGTCCGCGAGTGACCAGTGACTCGCACTGGGAAATGACTCGCACTGGGAAAGTCGAAGTTTTCCACGAAGAGCACCTTGTCAATGACTCGCACTGGGAAATGACTCGCACTGGGACAACGTCCATGTCGGGATGC |
| PA14_51340 | mvfR | AGTTCGCTGAGGACTGCCAGGGTACCGTTG | TTTCGTCCGCGAGTGACCAGTGACTCGCACTGGGAAATGACTCGCACTGGGAAAAGTTCGCTGAGGACTGCCAGGGTACCGTTGAATGACTCGCACTGGGAAATGACTCGCACTGGGACAACGTCCATGTCGGGATGC |
| PA14_51340 | mvfR | TTCAGCAGGCTGCGCATGTAAGGGATCAGG | TTTCGTCCGCGAGTGACCAGTGACTCGCACTGGGAAATGACTCGCACTGGGAAATTCAGCAGGCTGCGCATGTAAGGGATCAGGAATGACTCGCACTGGGAAATGACTCGCACTGGGACAACGTCCATGTCGGGATGC |
| PA14_51340 | mvfR | AAGCGCCTGCTCGGTGGGTTCGACCTTGTA | TTTCGTCCGCGAGTGACCAGTGACTCGCACTGGGAAATGACTCGCACTGGGAAAAAGCGCCTGCTCGGTGGGTTCGACCTTGTAAATGACTCGCACTGGGAAATGACTCGCACTGGGACAACGTCCATGTCGGGATGC |
| PA14_51340 | mvfR | TGCACCGTTTCGACGATGCTCGGTTGCCAG | TTTCGTCCGCGAGTGACCAGTGACTCGCACTGGGAAATGACTCGCACTGGGAAATGCACCGTTTCGACGATGCTCGGTTGCCAGAATGACTCGCACTGGGAAATGACTCGCACTGGGACAACGTCCATGTCGGGATGC |
| PA14_51340 | mvfR | TGAAGGCGATGTCGCCGATCAGCTGCTGGT | TTTCGTCCGCGAGTGACCAGTGACTCGCACTGGGAAATGACTCGCACTGGGAAATGAAGGCGATGTCGCCGATCAGCTGCTGGTAATGACTCGCACTGGGAAATGACTCGCACTGGGACAACGTCCATGTCGGGATGC |
| PA14_51340 | mvfR | TCGATTTCCGCGTTGTCCTGCTTGATCGTC | TTTCGTCCGCGAGTGACCAGTGACTCGCACTGGGAAATGACTCGCACTGGGAAATCGATTTCCGCGTTGTCCTGCTTGATCGTCAATGACTCGCACTGGGAAATGACTCGCACTGGGACAACGTCCATGTCGGGATGC |
| PA14_49250 | napA | ACTTCTCGGCCATGATGTCGAAGGCCTGTT | TTTCGTCCGCGAGTGACCAGTTATTACGGAGGCTGAATTATTACGGAGGCTGAAACTTCTCGGCCATGATGTCGAAGGCCTGTTAATTATTACGGAGGCTGAATTATTACGGAGGCTGCAACGTCCATGTCGGGATGC |
| PA14_49250 | napA | TTCCACCAGTTGCCAGAGGTCGGACTTCGC | TTTCGTCCGCGAGTGACCAGTTATTACGGAGGCTGAATTATTACGGAGGCTGAATTCCACCAGTTGCCAGAGGTCGGACTTCGCAATTATTACGGAGGCTGAATTATTACGGAGGCTGCAACGTCCATGTCGGGATGC |
| PA14_49250 | napA | AACTCGGCGAAGGCCTTGAAGTCGATGTCG | TTTCGTCCGCGAGTGACCAGTTATTACGGAGGCTGAATTATTACGGAGGCTGAAAACTCGGCGAAGGCCTTGAAGTCGATGTCGAATTATTACGGAGGCTGAATTATTACGGAGGCTGCAACGTCCATGTCGGGATGC |
| PA14_49250 | napA | TGATCCCGCGGTTGACCTCGGCCTTGATGT | TTTCGTCCGCGAGTGACCAGTTATTACGGAGGCTGAATTATTACGGAGGCTGAATGATCCCGCGGTTGACCTCGGCCTTGATGTAATTATTACGGAGGCTGAATTATTACGGAGGCTGCAACGTCCATGTCGGGATGC |
| PA14_49250 | napA | AAGGGGCTGTTGCCCGGTTCGCTGATCTTC | TTTCGTCCGCGAGTGACCAGTTATTACGGAGGCTGAATTATTACGGAGGCTGAAAAGGGGCTGTTGCCCGGTTCGCTGATCTTCAATTATTACGGAGGCTGAATTATTACGGAGGCTGCAACGTCCATGTCGGGATGC |
| PA14_49250 | napA | TTCACCAACTGGTGCCAGAACTGCGTGCGG | TTTCGTCCGCGAGTGACCAGTTATTACGGAGGCTGAATTATTACGGAGGCTGAATTCACCAACTGGTGCCAGAACTGCGTGCGGAATTATTACGGAGGCTGAATTATTACGGAGGCTGCAACGTCCATGTCGGGATGC |
| PA14_49250 | napA | TTCGATGCGAACGGCGCACTTCTTGTAGTC | TTTCGTCCGCGAGTGACCAGTTATTACGGAGGCTGAATTATTACGGAGGCTGAATTCGATGCGAACGGCGCACTTCTTGTAGTCAATTATTACGGAGGCTGAATTATTACGGAGGCTGCAACGTCCATGTCGGGATGC |
| PA14_49250 | napA | TCAGGTCGGTCTGCGGCTTGAACACCATGG | TTTCGTCCGCGAGTGACCAGTTATTACGGAGGCTGAATTATTACGGAGGCTGAATCAGGTCGGTCTGCGGCTTGAACACCATGGAATTATTACGGAGGCTGAATTATTACGGAGGCTGCAACGTCCATGTCGGGATGC |
| PA14_49250 | napA | CGATGATGTGGTTGGCGATGTAGTTGAGGA | TTTCGTCCGCGAGTGACCAGTTATTACGGAGGCTGAATTATTACGGAGGCTGAACGATGATGTGGTTGGCGATGTAGTTGAGGAAATTATTACGGAGGCTGAATTATTACGGAGGCTGCAACGTCCATGTCGGGATGC |
| PA14_49250 | napA | GATCTTCGACAGGAAGTAGCCCTTCACGCA | TTTCGTCCGCGAGTGACCAGTTATTACGGAGGCTGAATTATTACGGAGGCTGAAGATCTTCGACAGGAAGTAGCCCTTCACGCAAATTATTACGGAGGCTGAATTATTACGGAGGCTGCAACGTCCATGTCGGGATGC |
| PA14_49250 | napA | ATCGGCTGGAACTCGCCCTGCTTGTCGAAC | TTTCGTCCGCGAGTGACCAGTTATTACGGAGGCTGAATTATTACGGAGGCTGAAATCGGCTGGAACTCGCCCTGCTTGTCGAACAATTATTACGGAGGCTGAATTATTACGGAGGCTGCAACGTCCATGTCGGGATGC |
| PA14_49250 | napA | GGCCTTGGCGAATTCACGACGGGTGAGGTT | TTTCGTCCGCGAGTGACCAGTTATTACGGAGGCTGAATTATTACGGAGGCTGAAGGCCTTGGCGAATTCACGACGGGTGAGGTTAATTATTACGGAGGCTGAATTATTACGGAGGCTGCAACGTCCATGTCGGGATGC |
| PA14_49250 | napA | ATGCCGAAGCTGCGCATGAAGCCCATCACC | TTTCGTCCGCGAGTGACCAGTTATTACGGAGGCTGAATTATTACGGAGGCTGAAATGCCGAAGCTGCGCATGAAGCCCATCACCAATTATTACGGAGGCTGAATTATTACGGAGGCTGCAACGTCCATGTCGGGATGC |
| PA14_49250 | napA | CATCACGTTGGGACCGGCCTGCATGTTGTT | TTTCGTCCGCGAGTGACCAGTTATTACGGAGGCTGAATTATTACGGAGGCTGAACATCACGTTGGGACCGGCCTGCATGTTGTTAATTATTACGGAGGCTGAATTATTACGGAGGCTGCAACGTCCATGTCGGGATGC |
| PA14_49250 | napA | TGGTCCAGAACGACACCACCTTGCGCTTGG | TTTCGTCCGCGAGTGACCAGTTATTACGGAGGCTGAATTATTACGGAGGCTGAATGGTCCAGAACGACACCACCTTGCGCTTGGAATTATTACGGAGGCTGAATTATTACGGAGGCTGCAACGTCCATGTCGGGATGC |
| PA14_49250 | napA | TTGTTCGCCGCATAGCCTTCCCAGACCGTC | TTTCGTCCGCGAGTGACCAGTTATTACGGAGGCTGAATTATTACGGAGGCTGAATTGTTCGCCGCATAGCCTTCCCAGACCGTCAATTATTACGGAGGCTGAATTATTACGGAGGCTGCAACGTCCATGTCGGGATGC |
| PA14_49250 | napA | TCATGCTCCCGGTGTGCCAGTGTTCGAGCA | TTTCGTCCGCGAGTGACCAGTTATTACGGAGGCTGAATTATTACGGAGGCTGAATCATGCTCCCGGTGTGCCAGTGTTCGAGCAAATTATTACGGAGGCTGAATTATTACGGAGGCTGCAACGTCCATGTCGGGATGC |
| PA14_49250 | napA | TGACCTGGGTCCAGTAGACGTTGAGCACGC | TTTCGTCCGCGAGTGACCAGTTATTACGGAGGCTGAATTATTACGGAGGCTGAATGACCTGGGTCCAGTAGACGTTGAGCACGCAATTATTACGGAGGCTGAATTATTACGGAGGCTGCAACGTCCATGTCGGGATGC |
| PA14_49250 | napA | TGCCTTGTTCCAGACCAGGCTGGTGACGTC | TTTCGTCCGCGAGTGACCAGTTATTACGGAGGCTGAATTATTACGGAGGCTGAATGCCTTGTTCCAGACCAGGCTGGTGACGTCAATTATTACGGAGGCTGAATTATTACGGAGGCTGCAACGTCCATGTCGGGATGC |
| PA14_49250 | napA | TTGGCCTTGTCGGCGTTCTTCGCCTTCTTC | TTTCGTCCGCGAGTGACCAGTTATTACGGAGGCTGAATTATTACGGAGGCTGAATTGGCCTTGTCGGCGTTCTTCGCCTTCTTCAATTATTACGGAGGCTGAATTATTACGGAGGCTGCAACGTCCATGTCGGGATGC |
| PA14_41510 | nasA | AGAACATGAACCACCAGCTGTCGCGGTCGC | TTTCGTCCGCGAGTGACCAGTAGCTCGACAGGGTTAATAGCTCGACAGGGTTAAAGAACATGAACCACCAGCTGTCGCGGTCGCAATAGCTCGACAGGGTTAATAGCTCGACAGGGTTCAACGTCCATGTCGGGATGC |
| PA14_41510 | nasA | GAGCACGTACCAGACCATGAAGCTCAGGTC | TTTCGTCCGCGAGTGACCAGTAGCTCGACAGGGTTAATAGCTCGACAGGGTTAAGAGCACGTACCAGACCATGAAGCTCAGGTCAATAGCTCGACAGGGTTAATAGCTCGACAGGGTTCAACGTCCATGTCGGGATGC |
| PA14_41510 | nasA | ACCGAGACTGAGCATGGCGACCACGAACAG | TTTCGTCCGCGAGTGACCAGTAGCTCGACAGGGTTAATAGCTCGACAGGGTTAAACCGAGACTGAGCATGGCGACCACGAACAGAATAGCTCGACAGGGTTAATAGCTCGACAGGGTTCAACGTCCATGTCGGGATGC |
| PA14_41510 | nasA | TGTTGCGGCGGGTACCATTGCGATGCGAGC | TTTCGTCCGCGAGTGACCAGTAGCTCGACAGGGTTAATAGCTCGACAGGGTTAATGTTGCGGCGGGTACCATTGCGATGCGAGCAATAGCTCGACAGGGTTAATAGCTCGACAGGGTTCAACGTCCATGTCGGGATGC |
| PA14_41510 | nasA | TGCGGTACCAACTGGAACACCGCGCCATTG | TTTCGTCCGCGAGTGACCAGTAGCTCGACAGGGTTAATAGCTCGACAGGGTTAATGCGGTACCAACTGGAACACCGCGCCATTGAATAGCTCGACAGGGTTAATAGCTCGACAGGGTTCAACGTCCATGTCGGGATGC |
| PA14_41510 | nasA | AAGCCGAACACGTTGTTCCAGCCGAACGCG | TTTCGTCCGCGAGTGACCAGTAGCTCGACAGGGTTAATAGCTCGACAGGGTTAAAAGCCGAACACGTTGTTCCAGCCGAACGCGAATAGCTCGACAGGGTTAATAGCTCGACAGGGTTCAACGTCCATGTCGGGATGC |
| PA14_41510 | nasA | GGCTTTCAGGTAATCGCCCATGGCCTTGGG | TTTCGTCCGCGAGTGACCAGTAGCTCGACAGGGTTAATAGCTCGACAGGGTTAAGGCTTTCAGGTAATCGCCCATGGCCTTGGGAATAGCTCGACAGGGTTAATAGCTCGACAGGGTTCAACGTCCATGTCGGGATGC |
| PA14_41510 | nasA | AGATGGCGGCCAGGGTGTACACCATCAGCA | TTTCGTCCGCGAGTGACCAGTAGCTCGACAGGGTTAATAGCTCGACAGGGTTAAAGATGGCGGCCAGGGTGTACACCATCAGCAAATAGCTCGACAGGGTTAATAGCTCGACAGGGTTCAACGTCCATGTCGGGATGC |
| PA14_41510 | nasA | AATAGAGAAACGCCGCGAACAGGGTCGGCG | TTTCGTCCGCGAGTGACCAGTAGCTCGACAGGGTTAATAGCTCGACAGGGTTAAAATAGAGAAACGCCGCGAACAGGGTCGGCGAATAGCTCGACAGGGTTAATAGCTCGACAGGGTTCAACGTCCATGTCGGGATGC |
| PA14_41510 | nasA | ATGGCGAACACCAGGAGGGTCAGGACGAGC | TTTCGTCCGCGAGTGACCAGTAGCTCGACAGGGTTAATAGCTCGACAGGGTTAAATGGCGAACACCAGGAGGGTCAGGACGAGCAATAGCTCGACAGGGTTAATAGCTCGACAGGGTTCAACGTCCATGTCGGGATGC |
| PA14_41510 | nasA | TGGTAGTCGCCGGTGCTTTGCTTGATCGCG | TTTCGTCCGCGAGTGACCAGTAGCTCGACAGGGTTAATAGCTCGACAGGGTTAATGGTAGTCGCCGGTGCTTTGCTTGATCGCGAATAGCTCGACAGGGTTAATAGCTCGACAGGGTTCAACGTCCATGTCGGGATGC |
| PA14_41510 | nasA | TCATCGCGCGCTGTTGGGTGCTCAGTTGCA | TTTCGTCCGCGAGTGACCAGTAGCTCGACAGGGTTAATAGCTCGACAGGGTTAATCATCGCGCGCTGTTGGGTGCTCAGTTGCAAATAGCTCGACAGGGTTAATAGCTCGACAGGGTTCAACGTCCATGTCGGGATGC |
| PA14_41510 | nasA | GTTCAGGCCGTACTGGTCGTGGAAATAGCC | TTTCGTCCGCGAGTGACCAGTAGCTCGACAGGGTTAATAGCTCGACAGGGTTAAGTTCAGGCCGTACTGGTCGTGGAAATAGCCAATAGCTCGACAGGGTTAATAGCTCGACAGGGTTCAACGTCCATGTCGGGATGC |
| PA14_41510 | nasA | AGCAGCAGTGCCTGCGCATAGCTGTTCACG | TTTCGTCCGCGAGTGACCAGTAGCTCGACAGGGTTAATAGCTCGACAGGGTTAAAGCAGCAGTGCCTGCGCATAGCTGTTCACGAATAGCTCGACAGGGTTAATAGCTCGACAGGGTTCAACGTCCATGTCGGGATGC |
| PA14_13780 | narG | TTGCGCACCTTCGGGTACTTCAGGCGGTTG | TTTCGTCCGCGAGTGACCAGTGCGGACCTCTTATAAATGCGGACCTCTTATAAATTGCGCACCTTCGGGTACTTCAGGCGGTTGAATGCGGACCTCTTATAAATGCGGACCTCTTATACAACGTCCATGTCGGGATGC |
| PA14_13780 | narG | GGTTGGAGCGCCAGATGAACATGTTGCGCG | TTTCGTCCGCGAGTGACCAGTGCGGACCTCTTATAAATGCGGACCTCTTATAAAGGTTGGAGCGCCAGATGAACATGTTGCGCGAATGCGGACCTCTTATAAATGCGGACCTCTTATACAACGTCCATGTCGGGATGC |
| PA14_13780 | narG | TAGCTGGAGTTGTACCAGTCGGCCGACTCC | TTTCGTCCGCGAGTGACCAGTGCGGACCTCTTATAAATGCGGACCTCTTATAAATAGCTGGAGTTGTACCAGTCGGCCGACTCCAATGCGGACCTCTTATAAATGCGGACCTCTTATACAACGTCCATGTCGGGATGC |
| PA14_13780 | narG | ACAGTTTCTCGGTGGTCCGGGTGTTGACCG | TTTCGTCCGCGAGTGACCAGTGCGGACCTCTTATAAATGCGGACCTCTTATAAAACAGTTTCTCGGTGGTCCGGGTGTTGACCGAATGCGGACCTCTTATAAATGCGGACCTCTTATACAACGTCCATGTCGGGATGC |
| PA14_13780 | narG | TCAGCATGTTGATGACCGCGCGGTAGTTCA | TTTCGTCCGCGAGTGACCAGTGCGGACCTCTTATAAATGCGGACCTCTTATAAATCAGCATGTTGATGACCGCGCGGTAGTTCAAATGCGGACCTCTTATAAATGCGGACCTCTTATACAACGTCCATGTCGGGATGC |
| PA14_13780 | narG | GTGGATGCCCCATTTCTGGTGCGGGGTGAT | TTTCGTCCGCGAGTGACCAGTGCGGACCTCTTATAAATGCGGACCTCTTATAAAGTGGATGCCCCATTTCTGGTGCGGGGTGATAATGCGGACCTCTTATAAATGCGGACCTCTTATACAACGTCCATGTCGGGATGC |

| | | | |
|---|---|---|---|
| PA14_13780 | narG | ACCGTAGGTCTTGGCGGTGTAGACGTTGGC | TTTCGTCCGCGAGTGACCAGTGCGGACCTCTTATAAATGCGGACCTCTTATAAAACCGTAGGTCTTGGCGGTGTAGACGTTGGCAATGCGGACCTCTTATAAATGCGGACCTCTTATACAACGTCCATGTCGGGATGC |
| PA14_13780 | narG | ACCCAGTCGTTATCGACGATCCCGGCCTTG | TTTCGTCCGCGAGTGACCAGTGCGGACCTCTTATAAATGCGGACCTCTTATAAAACCCAGTCGTTATCGACGATCCCGGCCTTGAATGCGGACCTCTTATAAATGCGGACCTCTTATACAACGTCCATGTCGGGATGC |
| PA14_13780 | narG | CGTGCATCGACAGCTTCTCGTGGCGCCATT | TTTCGTCCGCGAGTGACCAGTGCGGACCTCTTATAAATGCGGACCTCTTATAAACGTGCATCGACAGCTTCTCGTGGCGCCATTAATGCGGACCTCTTATAAATGCGGACCTCTTATACAACGTCCATGTCGGGATGC |
| PA14_13780 | narG | GTTCTTCACGTAGATCTTCCAGGAGCAGGA | TTTCGTCCGCGAGTGACCAGTGCGGACCTCTTATAAATGCGGACCTCTTATAAAGTTCTTCACGTAGATCTTCCAGGAGCAGGAAATGCGGACCTCTTATAAATGCGGACCTCTTATACAACGTCCATGTCGGGATGC |
| PA14_13780 | narG | AGATCTCCCAGTCGCTCTTGGCTTCCCAGG | TTTCGTCCGCGAGTGACCAGTGCGGACCTCTTATAAATGCGGACCTCTTATAAAAGATCTCCCAGTCGCTCTTGGCTTCCCAGGAATGCGGACCTCTTATAAATGCGGACCTCTTATACAACGTCCATGTCGGGATGC |
| PA14_13780 | narG | TCGGTCTGCTGGGTTTCCCAGGTGATCAGG | TTTCGTCCGCGAGTGACCAGTGCGGACCTCTTATAAATGCGGACCTCTTATAAATCGGTCTGCTGGGTTTCCCAGGTGATCAGGAATGCGGACCTCTTATAAATGCGGACCTCTTATACAACGTCCATGTCGGGATGC |
| PA14_13780 | narG | CCGACAGCGGATGGATGAAGGGGTGCATGT | TTTCGTCCGCGAGTGACCAGTGCGGACCTCTTATAAATGCGGACCTCTTATAAACCGACAGCGGATGGATGAAGGGGTGCATGTAATGCGGACCTCTTATAAATGCGGACCTCTTATACAACGTCCATGTCGGGATGC |
| PA14_13780 | narG | ACTTGCGGTAGACGTTCGGGTAGTCGCGCT | TTTCGTCCGCGAGTGACCAGTGCGGACCTCTTATAAATGCGGACCTCTTATAAAACTTGCGGTAGACGTTCGGGTAGTCGCGCTAATGCGGACCTCTTATAAATGCGGACCTCTTATACAACGTCCATGTCGGGATGC |
| PA14_13780 | narG | CTGCTTCTTCTTGAAGAACTGCAGGCGGTC | TTTCGTCCGCGAGTGACCAGTGCGGACCTCTTATAAATGCGGACCTCTTATAAACTGCTTCTTCTTGAAGAACTGCAGGCGGTCAATGCGGACCTCTTATAAATGCGGACCTCTTATACAACGTCCATGTCGGGATGC |
| PA14_06750 | nirS | CACGGCAACCTTGTTGGAGTTGTTGGCGGC | TTTCGTCCGCGAGTGACCAGGCTAGGCACCTGGATAAGCTAGGCACCTGGATAACACGGCAACCTTGTTGGAGTTGTTGGCGGCAAGCTAGGCACCTGGATAAGCTAGGCACCTGGATCAACGTCCATGTCGGGATGC |
| PA14_06750 | nirS | TCCACACCGAGAACCAGACTTCATCGCCGC | TTTCGTCCGCGAGTGACCAGGCTAGGCACCTGGATAAGCTAGGCACCTGGATAATCCACACCGAGAACCAGACTTCATCGCCGCAAGCTAGGCACCTGGATAAGCTAGGCACCTGGATCAACGTCCATGTCGGGATGC |
| PA14_06750 | nirS | TGGTAGGTCTGGGTGTCTACGGTCATGCCG | TTTCGTCCGCGAGTGACCAGGCTAGGCACCTGGATAAGCTAGGCACCTGGATAATGGTAGGTCTGGGTGTCTACGGTCATGCCGAAGCTAGGCACCTGGATAAGCTAGGCACCTGGATCAACGTCCATGTCGGGATGC |
| PA14_06750 | nirS | TGAACTTGGAGCTTTCCACCGAGCGCGCCT | TTTCGTCCGCGAGTGACCAGGCTAGGCACCTGGATAAGCTAGGCACCTGGATAATGAACTTGGAGCTTTCCACCGAGCGCGCCTAAGCTAGGCACCTGGATAAGCTAGGCACCTGGATCAACGTCCATGTCGGGATGC |
| PA14_06750 | nirS | TATTGCTGGCCGCGTTGCTGGGTGATGTCC | TTTCGTCCGCGAGTGACCAGGCTAGGCACCTGGATAAGCTAGGCACCTGGATAATATTGCTGGCCGCGTTGCTGGGTGATGTCCAAGCTAGGCACCTGGATAAGCTAGGCACCTGGATCAACGTCCATGTCGGGATGC |
| PA14_06750 | nirS | TTGACGATGAACTCGGGGTGCTCGTGGGAG | TTTCGTCCGCGAGTGACCAGGCTAGGCACCTGGATAAGCTAGGCACCTGGATAATTGACGATGAACTCGGGGTGCTCGTGGGAGAAGCTAGGCACCTGGATAAGCTAGGCACCTGGATCAACGTCCATGTCGGGATGC |
| PA14_06750 | nirS | TGGAGACGATCTGCTTCGGCTCCAGGGTCT | TTTCGTCCGCGAGTGACCAGGCTAGGCACCTGGATAAGCTAGGCACCTGGATAATGGAGACGATCTGCTTCGGCTCCAGGGTCTAAGCTAGGCACCTGGATAAGCTAGGCACCTGGATCAACGTCCATGTCGGGATGC |
| PA14_06750 | nirS | TTGTTGTACTCGGGCTGCACCACCCGCTTG | TTTCGTCCGCGAGTGACCAGGCTAGGCACCTGGATAAGCTAGGCACCTGGATAATTGTTGTACTCGGGCTGCACCACCCGCTTGAAGCTAGGCACCTGGATAAGCTAGGCACCTGGATCAACGTCCATGTCGGGATGC |
| PA14_06750 | nirS | TATGCACGGCATAGCCGGTATCGATGACCT | TTTCGTCCGCGAGTGACCAGGCTAGGCACCTGGATAAGCTAGGCACCTGGATAATATGCACGGCATAGCCGGTATCGATGACCTAAGCTAGGCACCTGGATAAGCTAGGCACCTGGATCAACGTCCATGTCGGGATGC |
| PA14_06750 | nirS | ACCTTCCACGATTCGCGCATCTCCGGCATG | TTTCGTCCGCGAGTGACCAGGCTAGGCACCTGGATAAGCTAGGCACCTGGATAAACCTTCCACGATTCGCGCATCTCCGGCATGAAGCTAGGCACCTGGATAAGCTAGGCACCTGGATCAACGTCCATGTCGGGATGC |
| PA14_06750 | nirS | ACGATCTTCTTGCTGTCGCCGTCGACCAGG | TTTCGTCCGCGAGTGACCAGGCTAGGCACCTGGATAAGCTAGGCACCTGGATAAACGATCTTCTTGCTGTCGCCGTCGACCAGGAAGCTAGGCACCTGGATAAGCTAGGCACCTGGATCAACGTCCATGTCGGGATGC |
| PA14_06750 | nirS | GGTGCCCACCAGTGGCTTGCCAAATGGCAT | TTTCGTCCGCGAGTGACCAGGCTAGGCACCTGGATAAGCTAGGCACCTGGATAAGGTGCCCACCAGTGGCTTGCCAAATGGCATAAGCTAGGCACCTGGATAAGCTAGGCACCTGGATCAACGTCCATGTCGGGATGC |
| PA14_06750 | nirS | TCGTTGAGCTGTTTCTTCGGCCGGTCCTCC | TTTCGTCCGCGAGTGACCAGGCTAGGCACCTGGATAAGCTAGGCACCTGGATAATCGTTGAGCTGTTTCTTCGGCCGGTCCTCCAAGCTAGGCACCTGGATAAGCTAGGCACCTGGATCAACGTCCATGTCGGGATGC |
| PA14_06750 | nirS | TTTCATGTCGTCCTTGGCGTGAGCGGTGGC | TTTCGTCCGCGAGTGACCAGGCTAGGCACCTGGATAAGCTAGGCACCTGGATAATTTCATGTCGTCCTTGGCGTGAGCGGTGGCAAGCTAGGCACCTGGATAAGCTAGGCACCTGGATCAACGTCCATGTCGGGATGC |
| PA14_06750 | nirS | CGGCGCAGCGTTGGAAGTAGATCTGCTTGG | TTTCGTCCGCGAGTGACCAGGCTAGGCACCTGGATAAGCTAGGCACCTGGATAACGGCGCAGCGTTGGAAGTAGATCTGCTTGGAAGCTAGGCACCTGGATAAGCTAGGCACCTGGATCAACGTCCATGTCGGGATGC |
| PA14_06750 | nirS | ACCGGGCCGTACTTGGGATGCACGAAGTTG | TTTCGTCCGCGAGTGACCAGGCTAGGCACCTGGATAAGCTAGGCACCTGGATAAACCGGGCCGTACTTGGGATGCACGAAGTTGAAGCTAGGCACCTGGATAAGCTAGGCACCTGGATCAACGTCCATGTCGGGATGC |
| PA14_06750 | nirS | TTGGCCCACAGGTCGATCATGTCGATCCGC | TTTCGTCCGCGAGTGACCAGGCTAGGCACCTGGATAAGCTAGGCACCTGGATAATTGGCCCACAGGTCGATCATGTCGATCCGCAAGCTAGGCACCTGGATAAGCTAGGCACCTGGATCAACGTCCATGTCGGGATGC |
| PA14_06750 | nirS | ATGTACTTGGCCATCAGGGTGATCTGGTCC | TTTCGTCCGCGAGTGACCAGGCTAGGCACCTGGATAAGCTAGGCACCTGGATAAATGTACTTGGCCATCAGGGTGATCTGGTCCAAGCTAGGCACCTGGATAAGCTAGGCACCTGGATCAACGTCCATGTCGGGATGC |
| PA14_06750 | nirS | TCATGAAGTAGCGGTGGCTGCTGTCCCAGC | TTTCGTCCGCGAGTGACCAGGCTAGGCACCTGGATAAGCTAGGCACCTGGATAATCATGAAGTAGCGGTGGCTGCTGTCCCAGCAAGCTAGGCACCTGGATAAGCTAGGCACCTGGATCAACGTCCATGTCGGGATGC |
| PA14_06750 | nirS | TCGTTGAACTCGCTTTCACTCATGTCGGGG | TTTCGTCCGCGAGTGACCAGGCTAGGCACCTGGATAAGCTAGGCACCTGGATAATCGTTGAACTCGCTTTCACTCATGTCGGGGAAGCTAGGCACCTGGATAAGCTAGGCACCTGGATCAACGTCCATGTCGGGATGC |
| PA14_70560 | OxyR | CAGCACATAGAAAGGCTCGTCGAACAGCGG | TTTCGTCCGCGAGTGACCAGGCGGCATTGACTAAGAAGCGGCATTGACTAAGAACAGCACATAGAAAGGCTCGTCGAACAGCGGAAGCGGCATTGACTAAGAAGCGGCATTGACTAAGCAACGTCCATGTCGGGATGC |
| PA14_70560 | OxyR | ACAGGTAGGGACCGATGGTGTAGATGGCGC | TTTCGTCCGCGAGTGACCAGGCGGCATTGACTAAGAAGCGGCATTGACTAAGAAACAGGTAGGGACCGATGGTGTAGATGGCGCAAGCGGCATTGACTAAGAAGCGGCATTGACTAAGCAACGTCCATGTCGGGATGC |
| PA14_70560 | OxyR | GACCATGTGGCGGATGGTCTCCAGCGACGA | TTTCGTCCGCGAGTGACCAGGCGGCATTGACTAAGAAGCGGCATTGACTAAGAAGACCATGTGGCGGATGGTCTCCAGCGACGAAAGCGGCATTGACTAAGAAGCGGCATTGACTAAGCAACGTCCATGTCGGGATGC |
| PA14_70560 | OxyR | TGAGCAGCTCGGAATCGATGCTGGCCTTGG | TTTCGTCCGCGAGTGACCAGGCGGCATTGACTAAGAAGCGGCATTGACTAAGAATGAGCAGCTCGGAATCGATGCTGGCCTTGGAAGCGGCATTGACTAAGAAGCGGCATTGACTAAGCAACGTCCATGTCGGGATGC |
| PA14_70560 | OxyR | GCTCTTGCTGCGCTCGAAGATCAGCACCCC | TTTCGTCCGCGAGTGACCAGGCGGCATTGACTAAGAAGCGGCATTGACTAAGAAGCTCTTGCTGCGCTCGAAGATCAGCACCCCAAGCGGCATTGACTAAGAAGCGGCATTGACTAAGCAACGTCCATGTCGGGATGC |
| PA14_70560 | OxyR | AGAGGGTCGGTTGGCTGACGTGGCAACGCT | TTTCGTCCGCGAGTGACCAGGCGGCATTGACTAAGAAGCGGCATTGACTAAGAAAGAGGGTCGGTTGGCTGACGTGGCAACGCTAAGCGGCATTGACTAAGAAGCGGCATTGACTAAGCAACGTCCATGTCGGGATGC |
| PA14_70560 | OxyR | ATCGAGTCGGCCAGCACTTCGATGGCGCGT | TTTCGTCCGCGAGTGACCAGGCGGCATTGACTAAGAAGCGGCATTGACTAAGAAATCGAGTCGGCCAGCACTTCGATGGCGCGTAAGCGGCATTGACTAAGAAGCGGCATTGACTAAGCAACGTCCATGTCGGGATGC |
| PA14_70560 | OxyR | TGCGCAGCTTGTCGCGGAGGATATGGGTGA | TTTCGTCCGCGAGTGACCAGGCGGCATTGACTAAGAAGCGGCATTGACTAAGAATGCGCAGCTTGTCGCGGAGGATATGGGTGAAAGCGGCATTGACTAAGAAGCGGCATTGACTAAGCAACGTCCATGTCGGGATGC |
| PA14_70560 | OxyR | GACGATGTAGCGCAGTTCGGTGAGGGTCAT | TTTCGTCCGCGAGTGACCAGGCGGCATTGACTAAGAAGCGGCATTGACTAAGAAGACGATGTAGCGCAGTTCGGTGAGGGTCATAAGCGGCATTGACTAAGAAGCGGCATTGACTAAGCAACGTCCATGTCGGGATGC |
| PA14_70560 | OxyR | ACTCCACCGTGGTGTGCTTGTTCTCGTCGC | TTTCGTCCGCGAGTGACCAGGCGGCATTGACTAAGAAGCGGCATTGACTAAGAAACTCCACCGTGGTGTGCTTGTTCTCGTCGCAAGCGGCATTGACTAAGAAGCGGCATTGACTAAGCAACGTCCATGTCGGGATGC |
| PA14_70560 | OxyR | TGGTCAGGACGTCGGCTTCCTGGAACGGCA | TTTCGTCCGCGAGTGACCAGGCGGCATTGACTAAGAAGCGGCATTGACTAAGAATGGTCAGGACGTCGGCTTCCTGGAACGGCAAAGCGGCATTGACTAAGAAGCGGCATTGACTAAGCAACGTCCATGTCGGGATGC |
| PA14_70560 | OxyR | TTCTCCTCGATGTACAGCGGCATCTGCGGC | TTTCGTCCGCGAGTGACCAGGCGGCATTGACTAAGAAGCGGCATTGACTAAGAATTCTCCTCGATGTACAGCGGCATCTGCGGCAAGCGGCATTGACTAAGAAGCGGCATTGACTAAGCAACGTCCATGTCGGGATGC |
| PA14_70560 | OxyR | ATGCCCTGGGCCTGTTCGAGGACCTTCTGC | TTTCGTCCGCGAGTGACCAGGCGGCATTGACTAAGAAGCGGCATTGACTAAGAAATGCCCTGGGCCTGTTCGAGGACCTTCTGCAAGCGGCATTGACTAAGAAGCGGCATTGACTAAGCAACGTCCATGTCGGGATGC |
| PA14_70560 | OxyR | ATTTGCGGTTGTTCCTGGGTTTGCGGGCGG | TTTCGTCCGCGAGTGACCAGGCGGCATTGACTAAGAAGCGGCATTGACTAAGAAATTTGCGGTTGTTCCTGGGTTTGCGGGCGGAAGCGGCATTGACTAAGAAGCGGCATTGACTAAGCAACGTCCATGTCGGGATGC |
| PA14_06830 | norB | GAACAGGGTGATGAACACCATCGCCACGGT | TTTCGTCCGCGAGTGACCAGTCGGGCGGCTATAGAAATCGGGCGGCTATAGAAAGAACAGGGTGATGAACACCATCGCCACGGTAATCGGGCGGCTATAGAAATCGGGCGGCTATAGACAACGTCCATGTCGGGATGC |
| PA14_06830 | norB | ACAGGATCCAGGCCAGCTTCGGGCTGTACA | TTTCGTCCGCGAGTGACCAGTCGGGCGGCTATAGAAATCGGGCGGCTATAGAAAACAGGATCCAGGCCAGCTTCGGGCTGTACAAATCGGGCGGCTATAGAAATCGGGCGGCTATAGACAACGTCCATGTCGGGATGC |
| PA14_06830 | norB | GAATTTCAGGGAGCCATTGGGCGACATCAT | TTTCGTCCGCGAGTGACCAGTCGGGCGGCTATAGAAATCGGGCGGCTATAGAAAGAATTTCAGGGAGCCATTGGGCGACATCATAATCGGGCGGCTATAGAAATCGGGCGGCTATAGACAACGTCCATGTCGGGATGC |
| PA14_06830 | norB | AGTTGACCGGCGCCAGGGTATGCATGAAGC | TTTCGTCCGCGAGTGACCAGTCGGGCGGCTATAGAAATCGGGCGGCTATAGAAAAGTTGACCGGCGCCAGGGTATGCATGAAGCAATCGGGCGGCTATAGAAATCGGGCGGCTATAGACAACGTCCATGTCGGGATGC |
| PA14_06830 | norB | ACGATCAGCAGGTTGGTGTGCACCATGCGC | TTTCGTCCGCGAGTGACCAGTCGGGCGGCTATAGAAATCGGGCGGCTATAGAAAACGATCAGCAGGTTGGTGTGCACCATGCGCAATCGGGCGGCTATAGAAATCGGGCGGCTATAGACAACGTCCATGTCGGGATGC |
| PA14_06830 | norB | AGCTGGTCCTGGGTGGCCATGAAGGTCATC | TTTCGTCCGCGAGTGACCAGTCGGGCGGCTATAGAAATCGGGCGGCTATAGAAAAGCTGGTCCTGGGTGGCCATGAAGGTCATCAATCGGGCGGCTATAGAAATCGGGCGGCTATAGACAACGTCCATGTCGGGATGC |
| PA14_06830 | norB | CTTGCTGATGGTCGGCTGTTCGAGGAACTC | TTTCGTCCGCGAGTGACCAGTCGGGCGGCTATAGAAATCGGGCGGCTATAGAAACTTGCTGATGGTCGGCTGTTCGAGGAACTCAATCGGGCGGCTATAGAAATCGGGCGGCTATAGACAACGTCCATGTCGGGATGC |
| PA14_06830 | norB | GCACCACCCACCACCAGTAGAACTTGTCGC | TTTCGTCCGCGAGTGACCAGTCGGGCGGCTATAGAAATCGGGCGGCTATAGAAAGCACCACCCACCACCAGTAGAACTTGTCGCAATCGGGCGGCTATAGAAATCGGGCGGCTATAGACAACGTCCATGTCGGGATGC |
| PA14_06830 | norB | GCATGGCGTAGGAGATGATGGTCATCACGA | TTTCGTCCGCGAGTGACCAGTCGGGCGGCTATAGAAATCGGGCGGCTATAGAAAGCATGGCGTAGGAGATGATGGTCATCACGAAATCGGGCGGCTATAGAAATCGGGCGGCTATAGACAACGTCCATGTCGGGATGC |
| PA14_06830 | norB | TTGAAGGGGATCGCCGGGAACAGGAAGTCG | TTTCGTCCGCGAGTGACCAGTCGGGCGGCTATAGAAATCGGGCGGCTATAGAAATTGAAGGGGATCGCCGGGAACAGGAAGTCGAATCGGGCGGCTATAGAAATCGGGCGGCTATAGACAACGTCCATGTCGGGATGC |
| PA14_06830 | norB | GATCAGTTCCCACACGCCTTCCACCCACAG | TTTCGTCCGCGAGTGACCAGTCGGGCGGCTATAGAAATCGGGCGGCTATAGAAAGATCAGTTCCCACACGCCTTCCACCCACAGAATCGGGCGGCTATAGAAATCGGGCGGCTATAGACAACGTCCATGTCGGGATGC |
| PA14_06830 | norB | TCAGCCAGAAGCCCCACATCTCCAGGACCT | TTTCGTCCGCGAGTGACCAGTCGGGCGGCTATAGAAATCGGGCGGCTATAGAAATCAGCCAGAAGCCCCACATCTCCAGGACCTAATCGGGCGGCTATAGAAATCGGGCGGCTATAGACAACGTCCATGTCGGGATGC |
| PA14_06830 | norB | AGAATACCGAGCCGAGCCACAGCCAGTAGC | TTTCGTCCGCGAGTGACCAGTCGGGCGGCTATAGAAATCGGGCGGCTATAGAAAAGAATACCGAGCCGAGCCACAGCCAGTAGCAATCGGGCGGCTATAGAAATCGGGCGGCTATAGACAACGTCCATGTCGGGATGC |
| PA14_06830 | norB | ATCGCGTAGGCGCCATAGAAGGCCATGTGG | TTTCGTCCGCGAGTGACCAGTCGGGCGGCTATAGAAATCGGGCGGCTATAGAAAATCGCGTAGGCGCCATAGAAGGCCATGTGGAATCGGGCGGCTATAGAAATCGGGCGGCTATAGACAACGTCCATGTCGGGATGC |
| PA14_06830 | norB | CGAACAGGATCTGCCCGACGAAGAGGATCA | TTTCGTCCGCGAGTGACCAGTCGGGCGGCTATAGAAATCGGGCGGCTATAGAAACGAACAGGATCTGCCCGACGAAGAGGATCAAATCGGGCGGCTATAGAAATCGGGCGGCTATAGACAACGTCCATGTCGGGATGC |
| PA14_20200 | nosZ | AACCCGTAGTACTCATCCAGTTCGCCGGGG | TTTCGTCCGCGAGTGACCAGTGTTCGCCAGGATTGAATGTTCGCCAGGATTGAAAACCCGTAGTACTCATCCAGTTCGCCGGGGAATGTTCGCCAGGATTGAATGTTCGCCAGGATTGCAACGTCCATGTCGGGATGC |
| PA14_20200 | nosZ | ACGGCCTTCTCCGAGTTGTAGCAGGTGGAG | TTTCGTCCGCGAGTGACCAGTGTTCGCCAGGATTGAATGTTCGCCAGGATTGAAACGGCCTTCTCCGAGTTGTAGCAGGTGGAGAATGTTCGCCAGGATTGAATGTTCGCCAGGATTGCAACGTCCATGTCGGGATGC |
| PA14_20200 | nosZ | TGACGAAGCCGTGGGTCACGTCCTCGATCT | TTTCGTCCGCGAGTGACCAGTGTTCGCCAGGATTGAATGTTCGCCAGGATTGAATGACGAAGCCGTGGGTCACGTCCTCGATCTAATGTTCGCCAGGATTGAATGTTCGCCAGGATTGCAACGTCCATGTCGGGATGC |
| PA14_20200 | nosZ | AGTAGTACCAGTGCACCCCGGGCTTATCGG | TTTCGTCCGCGAGTGACCAGTGTTCGCCAGGATTGAATGTTCGCCAGGATTGAAAGTAGTACCAGTGCACCCCGGGCTTATCGGAATGTTCGCCAGGATTGAATGTTCGCCAGGATTGCAACGTCCATGTCGGGATGC |
| PA14_20200 | nosZ | CTGGACGTTGGGAATGGTGACGATGCGGTC | TTTCGTCCGCGAGTGACCAGTGTTCGCCAGGATTGAATGTTCGCCAGGATTGAACTGGACGTTGGGAATGGTGACGATGCGGTCAATGTTCGCCAGGATTGAATGTTCGCCAGGATTGCAACGTCCATGTCGGGATGC |
| PA14_20200 | nosZ | ACAGGTCGAAGGTCGAGCCGTCGTTGGGAT | TTTCGTCCGCGAGTGACCAGTGTTCGCCAGGATTGAATGTTCGCCAGGATTGAAACAGGTCGAAGGTCGAGCCGTCGTTGGGATAATGTTCGCCAGGATTGAATGTTCGCCAGGATTGCAACGTCCATGTCGGGATGC |
| PA14_20200 | nosZ | CCATGTCGGTGTTGTCGAGGTTGCCGTCGA | TTTCGTCCGCGAGTGACCAGTGTTCGCCAGGATTGAATGTTCGCCAGGATTGAACCATGTCGGTGTTGTCGAGGTTGCCGTCGAAATGTTCGCCAGGATTGAATGTTCGCCAGGATTGCAACGTCCATGTCGGGATGC |
| PA14_20200 | nosZ | GCTGATCTCCATGCAGACGCCGTGGTTGAC | TTTCGTCCGCGAGTGACCAGTGTTCGCCAGGATTGAATGTTCGCCAGGATTGAAGCTGATCTCCATGCAGACGCCGTGGTTGACAATGTTCGCCAGGATTGAATGTTCGCCAGGATTGCAACGTCCATGTCGGGATGC |
| PA14_20200 | nosZ | GAACACCGGTATGCGCATCAGCTCGCGCAT | TTTCGTCCGCGAGTGACCAGTGTTCGCCAGGATTGAATGTTCGCCAGGATTGAAGAACACCGGTATGCGCATCAGCTCGCGCATAATGTTCGCCAGGATTGAATGTTCGCCAGGATTGCAACGTCCATGTCGGGATGC |
| PA14_20200 | nosZ | AAGTACTTGCCGTCCGGCGAGGTGTTCAGC | TTTCGTCCGCGAGTGACCAGTGTTCGCCAGGATTGAATGTTCGCCAGGATTGAAAAGTACTTGCCGTCCGGCGAGGTGTTCAGCAATGTTCGCCAGGATTGAATGTTCGCCAGGATTGCAACGTCCATGTCGGGATGC |
| PA14_20200 | nosZ | ATGAACAGGTACTTGCCGTCGTACTTGCCG | TTTCGTCCGCGAGTGACCAGTGTTCGCCAGGATTGAATGTTCGCCAGGATTGAAATGAACAGGTACTTGCCGTCGTACTTGCCGAATGTTCGCCAGGATTGAATGTTCGCCAGGATTGCAACGTCCATGTCGGGATGC |
| PA14_20200 | nosZ | ACCACCCAGTCGCGCTCGTTACGCATCATG | TTTCGTCCGCGAGTGACCAGTGTTCGCCAGGATTGAATGTTCGCCAGGATTGAAACCACCCAGTCGCGCTCGTTACGCATCATGAATGTTCGCCAGGATTGAATGTTCGCCAGGATTGCAACGTCCATGTCGGGATGC |
| PA14_20200 | nosZ | ATCACCTGCCAGGCCACTTCCATGGTCTCG | TTTCGTCCGCGAGTGACCAGTGTTCGCCAGGATTGAATGTTCGCCAGGATTGAAATCACCTGCCAGGCCACTTCCATGGTCTCGAATGTTCGCCAGGATTGAATGTTCGCCAGGATTGCAACGTCCATGTCGGGATGC |
| PA14_20200 | nosZ | ACGTAGCGGGTATGCGGCACCTTTTGCAGG | TTTCGTCCGCGAGTGACCAGTGTTCGCCAGGATTGAATGTTCGCCAGGATTGAAACGTAGCGGGTATGCGGCACCTTTTGCAGGAATGTTCGCCAGGATTGAATGTTCGCCAGGATTGCAACGTCCATGTCGGGATGC |
| PA14_20200 | nosZ | AGATGTGCGGATGGTGGCAGTCGCCGTTGA | TTTCGTCCGCGAGTGACCAGTGTTCGCCAGGATTGAATGTTCGCCAGGATTGAAAGATGTGCGGATGGTGGCAGTCGCCGTTGAAATGTTCGCCAGGATTGAATGTTCGCCAGGATTGCAACGTCCATGTCGGGATGC |
| PA14_20200 | nosZ | AAGCGCTTCGCCTTGATCTCGGCCTCGATG | TTTCGTCCGCGAGTGACCAGTGTTCGCCAGGATTGAATGTTCGCCAGGATTGAAAAGCGCTTCGCCTTGATCTCGGCCTCGATGAATGTTCGCCAGGATTGAATGTTCGCCAGGATTGCAACGTCCATGTCGGGATGC |
| PA14_20200 | nosZ | TCACCTTGAACTCGGTGAGGCCGAAGGAGG | TTTCGTCCGCGAGTGACCAGTGTTCGCCAGGATTGAATGTTCGCCAGGATTGAATCACCTTGAACTCGGTGAGGCCGAAGGAGGAATGTTCGCCAGGATTGAATGTTCGCCAGGATTGCAACGTCCATGTCGGGATGC |
| PA14_20200 | nosZ | ATTTCATGACGTCCAGGCGGATGCGCGCGA | TTTCGTCCGCGAGTGACCAGTGTTCGCCAGGATTGAATGTTCGCCAGGATTGAAATTTCATGACGTCCAGGCGGATGCGCGCGAAATGTTCGCCAGGATTGAATGTTCGCCAGGATTGCAACGTCCATGTCGGGATGC |
| PA14_20200 | nosZ | TGGCGTTGTACAGGGTGAAGGCGTTGTCGC | TTTCGTCCGCGAGTGACCAGTGTTCGCCAGGATTGAATGTTCGCCAGGATTGAATGGCGTTGTACAGGGTGAAGGCGTTGTCGCAATGTTCGCCAGGATTGAATGTTCGCCAGGATTGCAACGTCCATGTCGGGATGC |
| PA14_20200 | nosZ | GGACCAGCTTCATTTCCTCGCCGGAAATGT | TTTCGTCCGCGAGTGACCAGTGTTCGCCAGGATTGAATGTTCGCCAGGATTGAAGGACCAGCTTCATTTCCTCGCCGGAAATGTAATGTTCGCCAGGATTGAATGTTCGCCAGGATTGCAACGTCCATGTCGGGATGC |
| PA14_48930 | PA14_48930 | GGCAGGTACTGGCAACCATTCTTGCAGGCA | TTTCGTCCGCGAGTGACCAGTGGTCGAGGTATTCGAATGGTCGAGGTATTCGAAGGCAGGTACTGGCAACCATTCTTGCAGGCAAATGGTCGAGGTATTCGAATGGTCGAGGTATTCGCAACGTCCATGTCGGGATGC |
| PA14_48930 | PA14_48930 | TATAGCCGGGAGGACAGCCAGGGTCATTCG | TTTCGTCCGCGAGTGACCAGTGGTCGAGGTATTCGAATGGTCGAGGTATTCGAATATAGCCGGGAGGACAGCCAGGGTCATTCGAATGGTCGAGGTATTCGAATGGTCGAGGTATTCGCAACGTCCATGTCGGGATGC |
| PA14_48930 | PA14_48930 | GGGCGGCGAGCTATTCACGAAGTTTCTGCC | TTTCGTCCGCGAGTGACCAGTGGTCGAGGTATTCGAATGGTCGAGGTATTCGAAGGGCGGCGAGCTATTCACGAAGTTTCTGCCAATGGTCGAGGTATTCGAATGGTCGAGGTATTCGCAACGTCCATGTCGGGATGC |
| PA14_48930 | PA14_48930 | TTCAACGAGGGACCGGTCATGCCGGGATTG | TTTCGTCCGCGAGTGACCAGTGGTCGAGGTATTCGAATGGTCGAGGTATTCGAATTCAACGAGGGACCGGTCATGCCGGGATTGAATGGTCGAGGTATTCGAATGGTCGAGGTATTCGCAACGTCCATGTCGGGATGC |
| PA14_48930 | PA14_48930 | AATGGTCATGGACTCGCAGGGAAGCTGGCC | TTTCGTCCGCGAGTGACCAGTGGTCGAGGTATTCGAATGGTCGAGGTATTCGAAAATGGTCATGGACTCGCAGGGAAGCTGGCCAATGGTCGAGGTATTCGAATGGTCGAGGTATTCGCAACGTCCATGTCGGGATGC |
| PA14_48930 | PA14_48930 | AGTCGGATCCGTGGGCGTCTTGACGCAAGT | TTTCGTCCGCGAGTGACCAGTGGTCGAGGTATTCGAATGGTCGAGGTATTCGAAAGTCGGATCCGTGGGCGTCTTGACGCAAGTAATGGTCGAGGTATTCGAATGGTCGAGGTATTCGCAACGTCCATGTCGGGATGC |
| PA14_48930 | PA14_48930 | CGCGAATTGATCACCATTTCAACCGTGGGA | TTTCGTCCGCGAGTGACCAGTGGTCGAGGTATTCGAATGGTCGAGGTATTCGAACGCGAATTGATCACCATTTCAACCGTGGGAAATGGTCGAGGTATTCGAATGGTCGAGGTATTCGCAACGTCCATGTCGGGATGC |
| PA14_48930 | PA14_48930 | AACAGATCCGTCAGCCCTTTGATGCAGTTG | TTTCGTCCGCGAGTGACCAGTGGTCGAGGTATTCGAATGGTCGAGGTATTCGAAAACAGATCCGTCAGCCCTTTGATGCAGTTGAATGGTCGAGGTATTCGAATGGTCGAGGTATTCGCAACGTCCATGTCGGGATGC |
| PA14_48930 | PA14_48930 | GTCGAACGCGCCCTTCATCTGATCGACGTT | TTTCGTCCGCGAGTGACCAGTGGTCGAGGTATTCGAATGGTCGAGGTATTCGAAGTCGAACGCGCCCTTCATCTGATCGACGTTAATGGTCGAGGTATTCGAATGGTCGAGGTATTCGCAACGTCCATGTCGGGATGC |
| PA14_48930 | PA14_48930 | TGCACACATAGTTGGTCGAGCCCCTCATCT | TTTCGTCCGCGAGTGACCAGTGGTCGAGGTATTCGAATGGTCGAGGTATTCGAATGCACACATAGTTGGTCGAGCCCCTCATCTAATGGTCGAGGTATTCGAATGGTCGAGGTATTCGCAACGTCCATGTCGGGATGC |
| PA14_48930 | PA14_48930 | TATACGTGGAGCCCTCAGCACAACCGGTGC | TTTCGTCCGCGAGTGACCAGTGGTCGAGGTATTCGAATGGTCGAGGTATTCGAATATACGTGGAGCCCTCAGCACAACCGGTGCAATGGTCGAGGTATTCGAATGGTCGAGGTATTCGCAACGTCCATGTCGGGATGC |
| PA14_48930 | PA14_48930 | GCGCATGACGGAAATAGTGAAATTCGGCTC | TTTCGTCCGCGAGTGACCAGTGGTCGAGGTATTCGAATGGTCGAGGTATTCGAAGCGCATGACGGAAATAGTGAAATTCGGCTCAATGGTCGAGGTATTCGAATGGTCGAGGTATTCGCAACGTCCATGTCGGGATGC |
| PA14_48930 | PA14_48930 | CAGTAATAAGGCTCCGCGTAGGCACTCCAG | TTTCGTCCGCGAGTGACCAGTGGTCGAGGTATTCGAATGGTCGAGGTATTCGAACAGTAATAAGGCTCCGCGTAGGCACTCCAGAATGGTCGAGGTATTCGAATGGTCGAGGTATTCGCAACGTCCATGTCGGGATGC |
| PA14_48930 | PA14_48930 | ACCGGTCAGGAGCAGTCGGGCCGTTTAATT | TTTCGTCCGCGAGTGACCAGTGGTCGAGGTATTCGAATGGTCGAGGTATTCGAAACCGGTCAGGAGCAGTCGGGCCGTTTAATTAATGGTCGAGGTATTCGAATGGTCGAGGTATTCGCAACGTCCATGTCGGGATGC |
| PA14_48930 | PA14_48930 | ATTCCCAGCCCGACCTCACAAGCCGCACTA | TTTCGTCCGCGAGTGACCAGTGGTCGAGGTATTCGAATGGTCGAGGTATTCGAAATTCCCAGCCCGACCTCACAAGCCGCACTAAATGGTCGAGGTATTCGAATGGTCGAGGTATTCGCAACGTCCATGTCGGGATGC |
| PA14_09280 | pchF | TACCGTCAGGTTGAGACAGAACGCCGGTGC | TTTCGTCCGCGAGTGACCAGACATTCGTTATCGCGAAACATTCGTTATCGCGAATACCGTCAGGTTGAGACAGAACGCCGGTGCAAACATTCGTTATCGCGAAACATTCGTTATCGCGCAACGTCCATGTCGGGATGC |
| PA14_09280 | pchF | TGCCACTGGTGAACACCACCGGCATCAGGT | TTTCGTCCGCGAGTGACCAGACATTCGTTATCGCGAAACATTCGTTATCGCGAATGCCACTGGTGAACACCACCGGCATCAGGTAAACATTCGTTATCGCGAAACATTCGTTATCGCGCAACGTCCATGTCGGGATGC |
| PA14_09280 | pchF | GAGTGACAGAACAGCCGGTAGAGGGTGTCG | TTTCGTCCGCGAGTGACCAGACATTCGTTATCGCGAAACATTCGTTATCGCGAAGAGTGACAGAACAGCCGGTAGAGGGTGTCGAAACATTCGTTATCGCGAAACATTCGTTATCGCGCAACGTCCATGTCGGGATGC |
| PA14_09280 | pchF | TAGCTGGAGACGATATCCAGCTGGCGCACG | TTTCGTCCGCGAGTGACCAGACATTCGTTATCGCGAAACATTCGTTATCGCGAATAGCTGGAGACGATATCCAGCTGGCGCACGAAACATTCGTTATCGCGAAACATTCGTTATCGCGCAACGTCCATGTCGGGATGC |
| PA14_09280 | pchF | CAATTGCGCGGCGCTTTTCGGCAGCATCAC | TTTCGTCCGCGAGTGACCAGACATTCGTTATCGCGAAACATTCGTTATCGCGAACAATTGCGCGGCGCTTTTCGGCAGCATCACAAACATTCGTTATCGCGAAACATTCGTTATCGCGCAACGTCCATGTCGGGATGC |
| PA14_09280 | pchF | ATTCGCCAAGCAGCAATTGCAGGCTGGCGT | TTTCGTCCGCGAGTGACCAGACATTCGTTATCGCGAAACATTCGTTATCGCGAAATTCGCCAAGCAGCAATTGCAGGCTGGCGTAAACATTCGTTATCGCGAAACATTCGTTATCGCGCAACGTCCATGTCGGGATGC |
| PA14_09280 | pchF | TACGCAGTCATCCCGGCCGATACTCAGCTC | TTTCGTCCGCGAGTGACCAGACATTCGTTATCGCGAAACATTCGTTATCGCGAATACGCAGTCATCCCGGCCGATACTCAGCTCAAACATTCGTTATCGCGAAACATTCGTTATCGCGCAACGTCCATGTCGGGATGC |
| PA14_09280 | pchF | TCGTTGTGCTCGCGCGTCAGTTCCTGGATC | TTTCGTCCGCGAGTGACCAGACATTCGTTATCGCGAAACATTCGTTATCGCGAATCGTTGTGCTCGCGCGTCAGTTCCTGGATCAAACATTCGTTATCGCGAAACATTCGTTATCGCGCAACGTCCATGTCGGGATGC |
| PA14_09280 | pchF | ACGAAGTCGTCGAACATGGCTTCCGCCTGC | TTTCGTCCGCGAGTGACCAGACATTCGTTATCGCGAAACATTCGTTATCGCGAAACGAAGTCGTCGAACATGGCTTCCGCCTGCAAACATTCGTTATCGCGAAACATTCGTTATCGCGCAACGTCCATGTCGGGATGC |
| PA14_09280 | pchF | GATACGGTGGCCGCGAATCTTCACCTGGTC | TTTCGTCCGCGAGTGACCAGACATTCGTTATCGCGAAACATTCGTTATCGCGAAGATACGGTGGCCGCGAATCTTCACCTGGTCAAACATTCGTTATCGCGAAACATTCGTTATCGCGCAACGTCCATGTCGGGATGC |
| PA14_09280 | pchF | TCGTACTCGGCGTACAGCTGGCAGGCGTTA | TTTCGTCCGCGAGTGACCAGACATTCGTTATCGCGAAACATTCGTTATCGCGAATCGTACTCGGCGTACAGCTGGCAGGCGTTAAAACATTCGTTATCGCGAAACATTCGTTATCGCGCAACGTCCATGTCGGGATGC |
| PA14_09280 | pchF | AGCAGTTCCGCCCAATGCGAAGGATCGCTG | TTTCGTCCGCGAGTGACCAGACATTCGTTATCGCGAAACATTCGTTATCGCGAAAGCAGTTCCGCCCAATGCGAAGGATCGCTGAAACATTCGTTATCGCGAAACATTCGTTATCGCGCAACGTCCATGTCGGGATGC |
| PA14_09280 | pchF | TCGACCAGGGTGAAATCCACCGAGCAGTGC | TTTCGTCCGCGAGTGACCAGACATTCGTTATCGCGAAACATTCGTTATCGCGAATCGACCAGGGTGAAATCCACCGAGCAGTGCAAACATTCGTTATCGCGAAACATTCGTTATCGCGCAACGTCCATGTCGGGATGC |
| PA14_09280 | pchF | AGAGTGCCAGCAACGCACTCTCCAGCTCGT | TTTCGTCCGCGAGTGACCAGACATTCGTTATCGCGAAACATTCGTTATCGCGAAAGAGTGCCAGCAACGCACTCTCCAGCTCGTAAACATTCGTTATCGCGAAACATTCGTTATCGCGCAACGTCCATGTCGGGATGC |
| PA14_09280 | pchF | GGTTCCGCCGTCCATTGCGGATCGAGATAG | TTTCGTCCGCGAGTGACCAGACATTCGTTATCGCGAAACATTCGTTATCGCGAAGGTTCCGCCGTCCATTGCGGATCGAGATAGAAACATTCGTTATCGCGAAACATTCGTTATCGCGCAACGTCCATGTCGGGATGC |
| PA14_42500 | pcrD | CAACGGCGAGCCGATGTACACCGACATCAT | TTTCGTCCGCGAGTGACCAGATATTCAGGCTAGCGAAATATTCAGGCTAGCGAACAACGGCGAGCCGATGTACACCGACATCATAAATATTCAGGCTAGCGAAATATTCAGGCTAGCGCAACGTCCATGTCGGGATGC |
| PA14_42500 | pcrD | CAACACGATCCTGCCAAGCGGCTGGATGTT | TTTCGTCCGCGAGTGACCAGATATTCAGGCTAGCGAAATATTCAGGCTAGCGAACAACACGATCCTGCCAAGCGGCTGGATGTTAAATATTCAGGCTAGCGAAATATTCAGGCTAGCGCAACGTCCATGTCGGGATGC |
| PA14_42500 | pcrD | TCACCAGCTCGCCGTAACCCTGCTCCATCT | TTTCGTCCGCGAGTGACCAGATATTCAGGCTAGCGAAATATTCAGGCTAGCGAATCACCAGCTCGCCGTAACCCTGCTCCATCTAAATATTCAGGCTAGCGAAATATTCAGGCTAGCGCAACGTCCATGTCGGGATGC |
| PA14_42500 | pcrD | ACGGCACGGTGAGGGCGAATTCTTCCTGCT | TTTCGTCCGCGAGTGACCAGATATTCAGGCTAGCGAAATATTCAGGCTAGCGAAACGGCACGGTGAGGGCGAATTCTTCCTGCTAAATATTCAGGCTAGCGAAATATTCAGGCTAGCGCAACGTCCATGTCGGGATGC |
| PA14_42500 | pcrD | GAAGATGATGATGCCCACCACCAGGTTGCC | TTTCGTCCGCGAGTGACCAGATATTCAGGCTAGCGAAATATTCAGGCTAGCGAAGAAGATGATGATGCCCACCACCAGGTTGCCAAATATTCAGGCTAGCGAAATATTCAGGCTAGCGCAACGTCCATGTCGGGATGC |
| PA14_42500 | pcrD | ATCAGCAGCACTGCCGGGAACACCGAGAAC | TTTCGTCCGCGAGTGACCAGATATTCAGGCTAGCGAAATATTCAGGCTAGCGAAATCAGCAGCACTGCCGGGAACACCGAGAACAAATATTCAGGCTAGCGAAATATTCAGGCTAGCGCAACGTCCATGTCGGGATGC |
| PA14_42500 | pcrD | CATCTGGCTTTCCTTTTCGATCACCGCGCG | TTTCGTCCGCGAGTGACCAGATATTCAGGCTAGCGAAATATTCAGGCTAGCGAACATCTGGCTTTCCTTTTCGATCACCGCGCGAAATATTCAGGCTAGCGAAATATTCAGGCTAGCGCAACGTCCATGTCGGGATGC |
| PA14_42500 | pcrD | GATGTTGTTGCCGCTGGAGTACTTGTAGCA | TTTCGTCCGCGAGTGACCAGATATTCAGGCTAGCGAAATATTCAGGCTAGCGAAGATGTTGTTGCCGCTGGAGTACTTGTAGCAAAATATTCAGGCTAGCGAAATATTCAGGCTAGCGCAACGTCCATGTCGGGATGC |
| PA14_42500 | pcrD | TTGACATCGATCACGCCGGCACGCATGTCG | TTTCGTCCGCGAGTGACCAGATATTCAGGCTAGCGAAATATTCAGGCTAGCGAATTGACATCGATCACGCCGGCACGCATGTCGAAATATTCAGGCTAGCGAAATATTCAGGCTAGCGCAACGTCCATGTCGGGATGC |
| PA14_42500 | pcrD | ATCTCGGTCATCCGCTGCAGGGGAACGATC | TTTCGTCCGCGAGTGACCAGATATTCAGGCTAGCGAAATATTCAGGCTAGCGAAATCTCGGTCATCCGCTGCAGGGGAACGATCAAATATTCAGGCTAGCGAAATATTCAGGCTAGCGCAACGTCCATGTCGGGATGC |
| PA14_42500 | pcrD | ATCATGAACACCACCGCCAGCAGCAACACC | TTTCGTCCGCGAGTGACCAGATATTCAGGCTAGCGAAATATTCAGGCTAGCGAAATCATGAACACCACCGCCAGCAGCAACACCAAATATTCAGGCTAGCGAAATATTCAGGCTAGCGCAACGTCCATGTCGGGATGC |
| PA14_42500 | pcrD | ATGAGGATGTCCAGCAGCAGCGGTGGCAAC | TTTCGTCCGCGAGTGACCAGATATTCAGGCTAGCGAAATATTCAGGCTAGCGAAATGAGGATGTCCAGCAGCAGCGGTGGCAACAAATATTCAGGCTAGCGAAATATTCAGGCTAGCGCAACGTCCATGTCGGGATGC |
| PA14_42500 | pcrD | ATGATCCCGGCGGTGATGGCGATCAGCAAC | TTTCGTCCGCGAGTGACCAGATATTCAGGCTAGCGAAATATTCAGGCTAGCGAAATGATCCCGGCGGTGATGGCGATCAGCAACAAATATTCAGGCTAGCGAAATATTCAGGCTAGCGCAACGTCCATGTCGGGATGC |
| PA14_42500 | pcrD | ACGAATTTCATCGCGCCGTCCATAGAGCCG | TTTCGTCCGCGAGTGACCAGATATTCAGGCTAGCGAAATATTCAGGCTAGCGAAACGAATTTCATCGCGCCGTCCATAGAGCCGAAATATTCAGGCTAGCGAAATATTCAGGCTAGCGCAACGTCCATGTCGGGATGC |
| PA14_42500 | pcrD | TGCTCATCTGCTTGCCCGGCATGGCATCCA | TTTCGTCCGCGAGTGACCAGATATTCAGGCTAGCGAAATATTCAGGCTAGCGAATGCTCATCTGCTTGCCCGGCATGGCATCCAAAATATTCAGGCTAGCGAAATATTCAGGCTAGCGCAACGTCCATGTCGGGATGC |
| PA14_42500 | pcrD | TTGGTGATCACCAGGAACTGCACGATGGTG | TTTCGTCCGCGAGTGACCAGATATTCAGGCTAGCGAAATATTCAGGCTAGCGAATTGGTGATCACCAGGAACTGCACGATGGTGAAATATTCAGGCTAGCGAAATATTCAGGCTAGCGCAACGTCCATGTCGGGATGC |
| PA14_42500 | pcrD | GCAGGACCACGGAGATGGTGATGTTGACGG | TTTCGTCCGCGAGTGACCAGATATTCAGGCTAGCGAAATATTCAGGCTAGCGAAGCAGGACCACGGAGATGGTGATGTTGACGGAAATATTCAGGCTAGCGAAATATTCAGGCTAGCGCAACGTCCATGTCGGGATGC |
| PA14_42500 | pcrD | TCGGGAAACCGGGAATCAGGCCGAACAGCA | TTTCGTCCGCGAGTGACCAGATATTCAGGCTAGCGAAATATTCAGGCTAGCGAATCGGGAAACCGGGAATCAGGCCGAACAGCAAAATATTCAGGCTAGCGAAATATTCAGGCTAGCGCAACGTCCATGTCGGGATGC |
| PA14_42500 | pcrD | TGAGCTGCACCACGTCCTTCTCCTTCTGCC | TTTCGTCCGCGAGTGACCAGATATTCAGGCTAGCGAAATATTCAGGCTAGCGAATGAGCTGCACCACGTCCTTCTCCTTCTGCCAAATATTCAGGCTAGCGAAATATTCAGGCTAGCGCAACGTCCATGTCGGGATGC |
| PA14_42500 | pcrD | TTGACGACGATAATGATCAGCCCGGCGATG | TTTCGTCCGCGAGTGACCAGATATTCAGGCTAGCGAAATATTCAGGCTAGCGAATTGACGACGATAATGATCAGCCCGGCGATGAAATATTCAGGCTAGCGAAATATTCAGGCTAGCGCAACGTCCATGTCGGGATGC |
| PA14_24480 | pelA | CCTGTAGCTGGAAGCTGTCCAGGGTATCGA | TTTCGTCCGCGAGTGACCAGACGCTAGTAACGTCTAAACGCTAGTAACGTCTAACCTGTAGCTGGAAGCTGTCCAGGGTATCGAAAACGCTAGTAACGTCTAAACGCTAGTAACGTCTCAACGTCCATGTCGGGATGC |

| | | | |
|---|---|---|---|
| PA14_24480 | pelA | ATCGGCAACTCGAACGTCCACAGGCCGTTG | TTTCGTCCGCGAGTGACCAGACGCTAGTAACGTCTAAACGCTAGTAACGTCTAAATCGGCAACTCGAACGTCCACAGGCCGTTGAAACGCTAGTAACGTCTAAACGCTAGTAACGTCTCAACGTCCATGTCGGGATGC |
| PA14_24480 | pelA | CACGGCGATTCCTTTCTTGCTGAACCGCAT | TTTCGTCCGCGAGTGACCAGACGCTAGTAACGTCTAAACGCTAGTAACGTCTAACACGGCGATTCCTTTCTTGCTGAACCGCATAAACGCTAGTAACGTCTAAACGCTAGTAACGTCTCAACGTCCATGTCGGGATGC |
| PA14_24480 | pelA | ATTCGCCGCACTCAGGTGTACATAGCGCCC | TTTCGTCCGCGAGTGACCAGACGCTAGTAACGTCTAAACGCTAGTAACGTCTAAATTCGCCGCACTCAGGTGTACATAGCGCCCAAACGCTAGTAACGTCTAAACGCTAGTAACGTCTCAACGTCCATGTCGGGATGC |
| PA14_24480 | pelA | TGACCCTTCAGCCAATCGCGATCGTCCTGG | TTTCGTCCGCGAGTGACCAGACGCTAGTAACGTCTAAACGCTAGTAACGTCTAATGACCCTTCAGCCAATCGCGATCGTCCTGGAAACGCTAGTAACGTCTAAACGCTAGTAACGTCTCAACGTCCATGTCGGGATGC |
| PA14_24480 | pelA | CGACCCGATAACCGAGGTATTCGAGCAGGC | TTTCGTCCGCGAGTGACCAGACGCTAGTAACGTCTAAACGCTAGTAACGTCTAACGACCCGATAACCGAGGTATTCGAGCAGGCAAACGCTAGTAACGTCTAAACGCTAGTAACGTCTCAACGTCCATGTCGGGATGC |
| PA14_24480 | pelA | TACCTGGCTGTTCCAGGCTTGGTTGCGGAC | TTTCGTCCGCGAGTGACCAGACGCTAGTAACGTCTAAACGCTAGTAACGTCTAATACCTGGCTGTTCCAGGCTTGGTTGCGGACAAACGCTAGTAACGTCTAAACGCTAGTAACGTCTCAACGTCCATGTCGGGATGC |
| PA14_24480 | pelA | AGTTTCTGCGACAGTCGGCGAATCCCCAGG | TTTCGTCCGCGAGTGACCAGACGCTAGTAACGTCTAAACGCTAGTAACGTCTAAAGTTTCTGCGACAGTCGGCGAATCCCCAGGAAACGCTAGTAACGTCTAAACGCTAGTAACGTCTCAACGTCCATGTCGGGATGC |
| PA14_24480 | pelA | TCTTTCAGATACCCGACATCGGCCGGCTTG | TTTCGTCCGCGAGTGACCAGACGCTAGTAACGTCTAAACGCTAGTAACGTCTAATCTTTCAGATACCCGACATCGGCCGGCTTGAAACGCTAGTAACGTCTAAACGCTAGTAACGTCTCAACGTCCATGTCGGGATGC |
| PA14_24480 | pelA | AAAACAGCTTGAGGCCCGGCTCCTGGCGAT | TTTCGTCCGCGAGTGACCAGACGCTAGTAACGTCTAAACGCTAGTAACGTCTAAAAAACAGCTTGAGGCCCGGCTCCTGGCGATAAACGCTAGTAACGTCTAAACGCTAGTAACGTCTCAACGTCCATGTCGGGATGC |
| PA14_24480 | pelA | TCGATGGGTCGAAGGGTGTTACGGCAGGAC | TTTCGTCCGCGAGTGACCAGACGCTAGTAACGTCTAAACGCTAGTAACGTCTAATCGATGGGTCGAAGGGTGTTACGGCAGGACAAACGCTAGTAACGTCTAAACGCTAGTAACGTCTCAACGTCCATGTCGGGATGC |
| PA14_24480 | pelA | TTGTCGTAGCCCGGGATGGCCATCTTGTAG | TTTCGTCCGCGAGTGACCAGACGCTAGTAACGTCTAAACGCTAGTAACGTCTAATTGTCGTAGCCCGGGATGGCCATCTTGTAGAAACGCTAGTAACGTCTAAACGCTAGTAACGTCTCAACGTCCATGTCGGGATGC |
| PA14_24480 | pelA | TCATCCACAGCGACAACGGATGCTCCGCCT | TTTCGTCCGCGAGTGACCAGACGCTAGTAACGTCTAAACGCTAGTAACGTCTAATCATCCACAGCGACAACGGATGCTCCGCCTAAACGCTAGTAACGTCTAAACGCTAGTAACGTCTCAACGTCCATGTCGGGATGC |
| PA14_24480 | pelA | TGCAACAGACCGTCGTTCTCGGTAGGCAGG | TTTCGTCCGCGAGTGACCAGACGCTAGTAACGTCTAAACGCTAGTAACGTCTAATGCAACAGACCGTCGTTCTCGGTAGGCAGGAAACGCTAGTAACGTCTAAACGCTAGTAACGTCTCAACGTCCATGTCGGGATGC |
| PA14_24480 | pelA | TGACGAACGGCACGTAGCCTTCGCTACGCA | TTTCGTCCGCGAGTGACCAGACGCTAGTAACGTCTAAACGCTAGTAACGTCTAATGACGAACGGCACGTAGCCTTCGCTACGCAAAACGCTAGTAACGTCTAAACGCTAGTAACGTCTCAACGTCCATGTCGGGATGC |
| PA14_09440 | phzE1 | TCCGCGCCATCATCTTCAGCTCCTCGTCCA | TTTCGTCCGCGAGTGACCAGCCTAGGCATATTCGAAACCTAGGCATATTCGAAATCCGCGCCATCATCTTCAGCTCCTCGTCCAAACCTAGGCATATTCGAAACCTAGGCATATTCGACAACGTCCATGTCGGGATGC |
| PA14_09440 | phzE1 | ACGATGAAGGTCCAGTAGGCGCCCTTCTCG | TTTCGTCCGCGAGTGACCAGCCTAGGCATATTCGAAACCTAGGCATATTCGAAAACGATGAAGGTCCAGTAGGCGCCCTTCTCGAACCTAGGCATATTCGAAACCTAGGCATATTCGACAACGTCCATGTCGGGATGC |
| PA14_09440 | phzE1 | AGCGGTAGGTGCCGCTGATCGGGTTCATCA | TTTCGTCCGCGAGTGACCAGCCTAGGCATATTCGAAACCTAGGCATATTCGAAAAGCGGTAGGTGCCGCTGATCGGGTTCATCAAACCTAGGCATATTCGAAACCTAGGCATATTCGACAACGTCCATGTCGGGATGC |
| PA14_09440 | phzE1 | TCTGTTTCTGCACGCCCTGGTTGGGCTCCT | TTTCGTCCGCGAGTGACCAGCCTAGGCATATTCGAAACCTAGGCATATTCGAAATCTGTTTCTGCACGCCCTGGTTGGGCTCCTAACCTAGGCATATTCGAAACCTAGGCATATTCGACAACGTCCATGTCGGGATGC |
| PA14_09440 | phzE1 | TTCCTGCTCCAGAACCTTCAGCGCCAGCAG | TTTCGTCCGCGAGTGACCAGCCTAGGCATATTCGAAACCTAGGCATATTCGAAATTCCTGCTCCAGAACCTTCAGCGCCAGCAGAACCTAGGCATATTCGAAACCTAGGCATATTCGACAACGTCCATGTCGGGATGC |
| PA14_09440 | phzE1 | GATCATCGAGGTGAAGGTGTCCTCGGCGTC | TTTCGTCCGCGAGTGACCAGCCTAGGCATATTCGAAACCTAGGCATATTCGAAAGATCATCGAGGTGAAGGTGTCCTCGGCGTCAACCTAGGCATATTCGAAACCTAGGCATATTCGACAACGTCCATGTCGGGATGC |
| PA14_09440 | phzE1 | TCGATGAAGTACTCGGTGTGCGCCAGGTGC | TTTCGTCCGCGAGTGACCAGCCTAGGCATATTCGAAACCTAGGCATATTCGAAATCGATGAAGTACTCGGTGTGCGCCAGGTGCAACCTAGGCATATTCGAAACCTAGGCATATTCGACAACGTCCATGTCGGGATGC |
| PA14_09440 | phzE1 | TTCGGGGTGGAATTGCATGGAGGCGAAGCG | TTTCGTCCGCGAGTGACCAGCCTAGGCATATTCGAAACCTAGGCATATTCGAAATTCGGGGTGGAATTGCATGGAGGCGAAGCGAACCTAGGCATATTCGAAACCTAGGCATATTCGACAACGTCCATGTCGGGATGC |
| PA14_09440 | phzE1 | CTGCAACAGCGATGTGGGCAGGGCGTTCAT | TTTCGTCCGCGAGTGACCAGCCTAGGCATATTCGAAACCTAGGCATATTCGAAACTGCAACAGCGATGTGGGCAGGGCGTTCATAACCTAGGCATATTCGAAACCTAGGCATATTCGACAACGTCCATGTCGGGATGC |
| PA14_09440 | phzE1 | AGCAACTGGCGGAAGAAACTCAGCGCGCTG | TTTCGTCCGCGAGTGACCAGCCTAGGCATATTCGAAACCTAGGCATATTCGAAAAGCAACTGGCGGAAGAAACTCAGCGCGCTGAACCTAGGCATATTCGAAACCTAGGCATATTCGACAACGTCCATGTCGGGATGC |
| PA14_09440 | phzE1 | TGGAAGCGCCGCTTGATCACGAAGTTGGCG | TTTCGTCCGCGAGTGACCAGCCTAGGCATATTCGAAACCTAGGCATATTCGAAATGGAAGCGCCGCTTGATCACGAAGTTGGCGAACCTAGGCATATTCGAAACCTAGGCATATTCGACAACGTCCATGTCGGGATGC |
| PA14_09440 | phzE1 | AGGAATTCCATGACCTCGGCCAGGTTCGGC | TTTCGTCCGCGAGTGACCAGCCTAGGCATATTCGAAACCTAGGCATATTCGAAAAGGAATTCCATGACCTCGGCCAGGTTCGGCAACCTAGGCATATTCGAAACCTAGGCATATTCGACAACGTCCATGTCGGGATGC |
| PA14_09440 | phzE1 | AAAGCTTCCTCGCTCAGTTCCAGCGCCTGG | TTTCGTCCGCGAGTGACCAGCCTAGGCATATTCGAAACCTAGGCATATTCGAAAAAAGCTTCCTCGCTCAGTTCCAGCGCCTGGAACCTAGGCATATTCGAAACCTAGGCATATTCGACAACGTCCATGTCGGGATGC |
| PA14_09440 | phzE1 | ACCATGTACAGCTCGTCGGCTTCCTTGCGG | TTTCGTCCGCGAGTGACCAGCCTAGGCATATTCGAAACCTAGGCATATTCGAAAACCATGTACAGCTCGTCGGCTTCCTTGCGGAACCTAGGCATATTCGAAACCTAGGCATATTCGACAACGTCCATGTCGGGATGC |
| PA14_09440 | phzE1 | TTGCTTTCCGGGCGGTAGAGCAAGGCGAAC | TTTCGTCCGCGAGTGACCAGCCTAGGCATATTCGAAACCTAGGCATATTCGAAATTGCTTTCCGGGCGGTAGAGCAAGGCGAACAACCTAGGCATATTCGAAACCTAGGCATATTCGACAACGTCCATGTCGGGATGC |
| PA14_00640 | phzH | AGCGAGGTATCCATGTCGCCGAAGGTGAAA | TTTCGTCCGCGAGTGACCAGAGTACTCTTACGCGAAAGTACTCTTACGCGAAAAGCGAGGTATCCATGTCGCCGAAGGTGAAAAAAGTACTCTTACGCGAAAGTACTCTTACGCGACAACGTCCATGTCGGGATGC |
| PA14_00640 | phzH | TGAAGTACCCCTCCGGATGCTCCAGTTCGG | TTTCGTCCGCGAGTGACCAGAGTACTCTTACGCGAAAGTACTCTTACGCGAAATGAAGTACCCCTCCGGATGCTCCAGTTCGGAAAGTACTCTTACGCGAAAGTACTCTTACGCGACAACGTCCATGTCGGGATGC |
| PA14_00640 | phzH | TCGGATAAGGGCTCTTGCGGCGCTTGAGCA | TTTCGTCCGCGAGTGACCAGAGTACTCTTACGCGAAAGTACTCTTACGCGAAATCGGATAAGGGCTCTTGCGGCGCTTGAGCAAAAGTACTCTTACGCGAAAGTACTCTTACGCGACAACGTCCATGTCGGGATGC |
| PA14_00640 | phzH | AGAGCCCGTACAACCTGAGCCAGCCTTCCA | TTTCGTCCGCGAGTGACCAGAGTACTCTTACGCGAAAGTACTCTTACGCGAAAAGAGCCCGTACAACCTGAGCCAGCCTTCCAAAAGTACTCTTACGCGAAAGTACTCTTACGCGACAACGTCCATGTCGGGATGC |
| PA14_00640 | phzH | ACTTCGCCTGGGAATTGGGACGCCAGGACA | TTTCGTCCGCGAGTGACCAGAGTACTCTTACGCGAAAGTACTCTTACGCGAAAACTTCGCCTGGGAATTGGGACGCCAGGACAAAAGTACTCTTACGCGAAAGTACTCTTACGCGACAACGTCCATGTCGGGATGC |
| PA14_00640 | phzH | ACTCCACCAGCTCATGGTCGGTGTAGGGCA | TTTCGTCCGCGAGTGACCAGAGTACTCTTACGCGAAAGTACTCTTACGCGAAAACTCCACCAGCTCATGGTCGGTGTAGGGCAAAAGTACTCTTACGCGAAAGTACTCTTACGCGACAACGTCCATGTCGGGATGC |
| PA14_00640 | phzH | TACGTCCTTGGCCCGGAATACCTCTTCGCG | TTTCGTCCGCGAGTGACCAGAGTACTCTTACGCGAAAGTACTCTTACGCGAAATACGTCCTTGGCCCGGAATACCTCTTCGCGAAAGTACTCTTACGCGAAAGTACTCTTACGCGACAACGTCCATGTCGGGATGC |
| PA14_00640 | phzH | TCGGTATCGCTGCGGGTGCGGAACTCATGT | TTTCGTCCGCGAGTGACCAGAGTACTCTTACGCGAAAGTACTCTTACGCGAAATCGGTATCGCTGCGGGTGCGGAACTCATGTAAAGTACTCTTACGCGAAAGTACTCTTACGCGACAACGTCCATGTCGGGATGC |
| PA14_00640 | phzH | TTGATCTCCGAGCCGAACAGCAGTCCCTCG | TTTCGTCCGCGAGTGACCAGAGTACTCTTACGCGAAAGTACTCTTACGCGAAATTGATCTCCGAGCCGAACAGCAGTCCCTCGAAAGTACTCTTACGCGAAAGTACTCTTACGCGACAACGTCCATGTCGGGATGC |
| PA14_00640 | phzH | AATTCCTGGACCTCGCGGAATGGCGTCTGC | TTTCGTCCGCGAGTGACCAGAGTACTCTTACGCGAAAGTACTCTTACGCGAAAAATTCCTGGACCTCGCGGAATGGCGTCTGCAAAGTACTCTTACGCGAAAGTACTCTTACGCGACAACGTCCATGTCGGGATGC |
| PA14_00640 | phzH | AGAACGAATTGATGTCGCCGCCGTGCTCCG | TTTCGTCCGCGAGTGACCAGAGTACTCTTACGCGAAAGTACTCTTACGCGAAAAGAACGAATTGATGTCGCCGCCGTGCTCCGAAAGTACTCTTACGCGAAAGTACTCTTACGCGACAACGTCCATGTCGGGATGC |
| PA14_00640 | phzH | GTACTCGCAACAACGCTCGCCCCATTGCAG | TTTCGTCCGCGAGTGACCAGAGTACTCTTACGCGAAAGTACTCTTACGCGAAAGTACTCGCAACAACGCTCGCCCCATTGCAGAAAGTACTCTTACGCGAAAGTACTCTTACGCGACAACGTCCATGTCGGGATGC |
| PA14_00640 | phzH | GTAATCCACCCAACCCGCGAGACCGCACAT | TTTCGTCCGCGAGTGACCAGAGTACTCTTACGCGAAAGTACTCTTACGCGAAAGTAATCCACCCAACCCGCGAGACCGCACATAAAGTACTCTTACGCGAAAGTACTCTTACGCGACAACGTCCATGTCGGGATGC |
| PA14_00640 | phzH | ACTGGAGGAGATCGCAGCGGCGGTTGAAAC | TTTCGTCCGCGAGTGACCAGAGTACTCTTACGCGAAAGTACTCTTACGCGAAAACTGGAGGAGATCGCAGCGGCGGTTGAAACAAAGTACTCTTACGCGAAAGTACTCTTACGCGACAACGTCCATGTCGGGATGC |
| PA14_00640 | phzH | TGGGAAAGCGATAGGACATCGGCTGCACGC | TTTCGTCCGCGAGTGACCAGAGTACTCTTACGCGAAAGTACTCTTACGCGAAATGGGAAAGCGATAGGACATCGGCTGCACGCAAAGTACTCTTACGCGAAAGTACTCTTACGCGACAACGTCCATGTCGGGATGC |
| PA14_00640 | phzH | TCAGATGCAGGTGGCTGAACTCGCGCATCC | TTTCGTCCGCGAGTGACCAGAGTACTCTTACGCGAAAGTACTCTTACGCGAAATCAGATGCAGGTGGCTGAACTCGCGCATCCAAAGTACTCTTACGCGAAAGTACTCTTACGCGACAACGTCCATGTCGGGATGC |
| PA14_00640 | phzH | ACACACCGGAACGTCGGCGTGCAATTGCGA | TTTCGTCCGCGAGTGACCAGAGTACTCTTACGCGAAAGTACTCTTACGCGAAAACACACCGGAACGTCGGCGTGCAATTGCGAAAAGTACTCTTACGCGAAAGTACTCTTACGCGACAACGTCCATGTCGGGATGC |
| PA14_00640 | phzH | ATTGTCGATGAGCACGGTGCGGTGACGGCT | TTTCGTCCGCGAGTGACCAGAGTACTCTTACGCGAAAGTACTCTTACGCGAAAATTGTCGATGAGCACGGTGCGGTGACGGCTAAAGTACTCTTACGCGAAAGTACTCTTACGCGACAACGTCCATGTCGGGATGC |
| PA14_00640 | phzH | ACACCTCGCCGGTATAGACGAGGGTGACCT | TTTCGTCCGCGAGTGACCAGAGTACTCTTACGCGAAAGTACTCTTACGCGAAAACACCTCGCCGGTATAGACGAGGGTGACCTAAAGTACTCTTACGCGAAAGTACTCTTACGCGACAACGTCCATGTCGGGATGC |
| PA14_00640 | phzH | GATCGCCGGAAATTCGTCGTCGAGCTTGCG | TTTCGTCCGCGAGTGACCAGAGTACTCTTACGCGAAAGTACTCTTACGCGAAAGATCGCCGGAAATTCGTCGTCGAGCTTGCGAAAGTACTCTTACGCGAAAGTACTCTTACGCGACAACGTCCATGTCGGGATGC |
| PA14_09490 | phzM | ATGGCCTTGGTCAATTCGCCGGAACCGCCA | TTTCGTCCGCGAGTGACCAGTAACGAGTGCTAACGAATAACGAGTGCTAACGAAATGGCCTTGGTCAATTCGCCGGAACCGCCAAATAACGAGTGCTAACGAATAACGAGTGCTAACGCAACGTCCATGTCGGGATGC |
| PA14_09490 | phzM | TGGCCGAGATGGTCCGCTCGATCACCACCA | TTTCGTCCGCGAGTGACCAGTAACGAGTGCTAACGAATAACGAGTGCTAACGAATGGCCGAGATGGTCCGCTCGATCACCACCAAATAACGAGTGCTAACGAATAACGAGTGCTAACGCAACGTCCATGTCGGGATGC |
| PA14_09490 | phzM | TATTAGCGTAGCCGTCGCGGGTATCGCCCT | TTTCGTCCGCGAGTGACCAGTAACGAGTGCTAACGAATAACGAGTGCTAACGAATATTAGCGTAGCCGTCGCGGGTATCGCCCTAATAACGAGTGCTAACGAATAACGAGTGCTAACGCAACGTCCATGTCGGGATGC |
| PA14_09490 | phzM | TCGTGGAAGGCCAGGTTGCTCGCCTTCATC | TTTCGTCCGCGAGTGACCAGTAACGAGTGCTAACGAATAACGAGTGCTAACGAATCGTGGAAGGCCAGGTTGCTCGCCTTCATCAATAACGAGTGCTAACGAATAACGAGTGCTAACGCAACGTCCATGTCGGGATGC |
| PA14_09490 | phzM | AGTCTTCGCCGAACGCCAGTTCGAAGCCTG | TTTCGTCCGCGAGTGACCAGTAACGAGTGCTAACGAATAACGAGTGCTAACGAAAGTCTTCGCCGAACGCCAGTTCGAAGCCTGAATAACGAGTGCTAACGAATAACGAGTGCTAACGCAACGTCCATGTCGGGATGC |
| PA14_09490 | phzM | AATTGCCGAGCAACCTCAGGCTGGCGGCTT | TTTCGTCCGCGAGTGACCAGTAACGAGTGCTAACGAATAACGAGTGCTAACGAAAATTGCCGAGCAACCTCAGGCTGGCGGCTTAATAACGAGTGCTAACGAATAACGAGTGCTAACGCAACGTCCATGTCGGGATGC |
| PA14_09490 | phzM | AAGCTACGCCCGCGGAAATCCAGGAGCCTG | TTTCGTCCGCGAGTGACCAGTAACGAGTGCTAACGAATAACGAGTGCTAACGAAAAGCTACGCCCGCGGAAATCCAGGAGCCTGAATAACGAGTGCTAACGAATAACGAGTGCTAACGCAACGTCCATGTCGGGATGC |
| PA14_09490 | phzM | TGTCGATCCCGCTCTCGATCAGATCGGCCA | TTTCGTCCGCGAGTGACCAGTAACGAGTGCTAACGAATAACGAGTGCTAACGAATGTCGATCCCGCTCTCGATCAGATCGGCCAAATAACGAGTGCTAACGAATAACGAGTGCTAACGCAACGTCCATGTCGGGATGC |
| PA14_09490 | phzM | AGCGACGATCATGCGGGTTTCCATCGGCAG | TTTCGTCCGCGAGTGACCAGTAACGAGTGCTAACGAATAACGAGTGCTAACGAAAGCGACGATCATGCGGGTTTCCATCGGCAGAATAACGAGTGCTAACGAATAACGAGTGCTAACGCAACGTCCATGTCGGGATGC |
| PA14_09490 | phzM | AACAGGTGCACGTCCCAGAGCACCGACATC | TTTCGTCCGCGAGTGACCAGTAACGAGTGCTAACGAATAACGAGTGCTAACGAAAACAGGTGCACGTCCCAGAGCACCGACATCAATAACGAGTGCTAACGAATAACGAGTGCTAACGCAACGTCCATGTCGGGATGC |
| PA14_09490 | phzM | GGGACTTCCATTCCCCGGTAACAACTTGTA | TTTCGTCCGCGAGTGACCAGTAACGAGTGCTAACGAATAACGAGTGCTAACGAAGGGACTTCCATTCCCCGGTAACAACTTGTAAATAACGAGTGCTAACGAATAACGAGTGCTAACGCAACGTCCATGTCGGGATGC |
| PA14_09490 | phzM | AGAACAGCACCATGTCGCGGAAGGAGCCCT | TTTCGTCCGCGAGTGACCAGTAACGAGTGCTAACGAATAACGAGTGCTAACGAAAGAACAGCACCATGTCGCGGAAGGAGCCCTAATAACGAGTGCTAACGAATAACGAGTGCTAACGCAACGTCCATGTCGGGATGC |
| PA14_09490 | phzM | AGCAGGCGCATCAGTCGATGGATGCGCTCG | TTTCGTCCGCGAGTGACCAGTAACGAGTGCTAACGAATAACGAGTGCTAACGAAAGCAGGCGCATCAGTCGATGGATGCGCTCGAATAACGAGTGCTAACGAATAACGAGTGCTAACGCAACGTCCATGTCGGGATGC |
| PA14_09490 | phzM | ATCGCCGTTGGACGGCACCTCTTGCAGCAT | TTTCGTCCGCGAGTGACCAGTAACGAGTGCTAACGAATAACGAGTGCTAACGAAATCGCCGTTGGACGGCACCTCTTGCAGCATAATAACGAGTGCTAACGAATAACGAGTGCTAACGCAACGTCCATGTCGGGATGC |
| PA14_09490 | phzM | TCCCCTGCCAACAGGCTGGAGAGGTTGTCG | TTTCGTCCGCGAGTGACCAGTAACGAGTGCTAACGAATAACGAGTGCTAACGAATCCCCTGCCAACAGGCTGGAGAGGTTGTCGAATAACGAGTGCTAACGAATAACGAGTGCTAACGCAACGTCCATGTCGGGATGC |
| PA14_09400 | phzS | AGTACTGCGGATAGGCGTTGCCGGCTTCCA | TTTCGTCCGCGAGTGACCAGTATTCGTGAATCGCGAATATTCGTGAATCGCGAAAGTACTGCGGATAGGCGTTGCCGGCTTCCAAATATTCGTGAATCGCGAATATTCGTGAATCGCGCAACGTCCATGTCGGGATGC |
| PA14_09400 | phzS | CAGGATCATCTGCAGTTCGCCGCGATGGAT | TTTCGTCCGCGAGTGACCAGTATTCGTGAATCGCGAATATTCGTGAATCGCGAACAGGATCATCTGCAGTTCGCCGCGATGGATAATATTCGTGAATCGCGAATATTCGTGAATCGCGCAACGTCCATGTCGGGATGC |
| PA14_09400 | phzS | AACACGTCTTCCAGGCGTCCGTTGCGGTTC | TTTCGTCCGCGAGTGACCAGTATTCGTGAATCGCGAATATTCGTGAATCGCGAAAACACGTCTTCCAGGCGTCCGTTGCGGTTCAATATTCGTGAATCGCGAATATTCGTGAATCGCGCAACGTCCATGTCGGGATGC |
| PA14_09400 | phzS | TCTTCTCAGTCTTCGGTCGCGAAGCCGCGG | TTTCGTCCGCGAGTGACCAGTATTCGTGAATCGCGAATATTCGTGAATCGCGAATCTTCTCAGTCTTCGGTCGCGAAGCCGCGGAATATTCGTGAATCGCGAATATTCGTGAATCGCGCAACGTCCATGTCGGGATGC |
| PA14_09400 | phzS | TGCTTTCCAGCAGCGTGACCTTGCCGATGC | TTTCGTCCGCGAGTGACCAGTATTCGTGAATCGCGAATATTCGTGAATCGCGAATGCTTTCCAGCAGCGTGACCTTGCCGATGCAATATTCGTGAATCGCGAATATTCGTGAATCGCGCAACGTCCATGTCGGGATGC |
| PA14_09400 | phzS | TAGCGTGGCCGTTCCACCTGGTTGCGGTAG | TTTCGTCCGCGAGTGACCAGTATTCGTGAATCGCGAATATTCGTGAATCGCGAATAGCGTGGCCGTTCCACCTGGTTGCGGTAGAATATTCGTGAATCGCGAATATTCGTGAATCGCGCAACGTCCATGTCGGGATGC |
| PA14_09400 | phzS | TGGTCTTGCCGTCGAGGAAGCGGTCGAACT | TTTCGTCCGCGAGTGACCAGTATTCGTGAATCGCGAATATTCGTGAATCGCGAATGGTCTTGCCGTCGAGGAAGCGGTCGAACTAATATTCGTGAATCGCGAATATTCGTGAATCGCGCAACGTCCATGTCGGGATGC |
| PA14_09400 | phzS | TTGGCCAGGATGATCTTGTTGGCGGTCGGC | TTTCGTCCGCGAGTGACCAGTATTCGTGAATCGCGAATATTCGTGAATCGCGAATTGGCCAGGATGATCTTGTTGGCGGTCGGCAATATTCGTGAATCGCGAATATTCGTGAATCGCGCAACGTCCATGTCGGGATGC |
| PA14_09400 | phzS | CGGCTGGATATTGATGCCGACGCCAAGGGG | TTTCGTCCGCGAGTGACCAGTATTCGTGAATCGCGAATATTCGTGAATCGCGAACGGCTGGATATTGATGCCGACGCCAAGGGGAATATTCGTGAATCGCGAATATTCGTGAATCGCGCAACGTCCATGTCGGGATGC |
| PA14_09400 | phzS | GGCAATGAGGATATCGATGGGTTCGCTCAT | TTTCGTCCGCGAGTGACCAGTATTCGTGAATCGCGAATATTCGTGAATCGCGAAGGCAATGAGGATATCGATGGGTTCGCTCATAATATTCGTGAATCGCGAATATTCGTGAATCGCGCAACGTCCATGTCGGGATGC |
| PA14_09400 | phzS | ACCATCGGGTACTGCAGGATCAACTGGTTG | TTTCGTCCGCGAGTGACCAGTATTCGTGAATCGCGAATATTCGTGAATCGCGAAACCATCGGGTACTGCAGGATCAACTGGTTGAATATTCGTGAATCGCGAATATTCGTGAATCGCGCAACGTCCATGTCGGGATGC |
| PA14_09400 | phzS | TCGATGTAGCGCAGCTCATGGGTGGGGATG | TTTCGTCCGCGAGTGACCAGTATTCGTGAATCGCGAATATTCGTGAATCGCGAATCGATGTAGCGCAGCTCATGGGTGGGGATGAATATTCGTGAATCGCGAATATTCGTGAATCGCGCAACGTCCATGTCGGGATGC |
| PA14_09400 | phzS | TTCTTCGTATTCGCGCAGGGCTGCGGCCAC | TTTCGTCCGCGAGTGACCAGTATTCGTGAATCGCGAATATTCGTGAATCGCGAATTCTTCGTATTCGCGCAGGGCTGCGGCCACAATATTCGTGAATCGCGAATATTCGTGAATCGCGCAACGTCCATGTCGGGATGC |
| PA14_09400 | phzS | ACACCCAGTTCACCAGCGACTTGCCTTCGG | TTTCGTCCGCGAGTGACCAGTATTCGTGAATCGCGAATATTCGTGAATCGCGAAACACCCAGTTCACCAGCGACTTGCCTTCGGAATATTCGTGAATCGCGAATATTCGTGAATCGCGCAACGTCCATGTCGGGATGC |
| PA14_09400 | phzS | TTGGCGCCCATCGGATACATCAGGTGGGCG | TTTCGTCCGCGAGTGACCAGTATTCGTGAATCGCGAATATTCGTGAATCGCGAATTGGCGCCCATCGGATACATCAGGTGGGCGAATATTCGTGAATCGCGAATATTCGTGAATCGCGCAACGTCCATGTCGGGATGC |
| PA14_60310 | pilY1 | ACGTCCGCCAGTATATGGATTGAGGCCGTA | TTTCGTCCGCGAGTGACCAGACGCGCGTCGTTATAAAACGCGCGTCGTTATAAAACGTCCGCCAGTATATGGATTGAGGCCGTAAAACGCGCGTCGTTATAAAACGCGCGTCGTTATACAACGTCCATGTCGGGATGC |
| PA14_60310 | pilY1 | ATTGTTGTCGGCGCGAGCACGGAAAAGCGG | TTTCGTCCGCGAGTGACCAGACGCGCGTCGTTATAAAACGCGCGTCGTTATAAAATTGTTGTCGGCGCGAGCACGGAAAAGCGGAAACGCGCGTCGTTATAAAACGCGCGTCGTTATACAACGTCCATGTCGGGATGC |
| PA14_60310 | pilY1 | TGTTGCGCTCGGAAATCCGTGTCAGGATCT | TTTCGTCCGCGAGTGACCAGACGCGCGTCGTTATAAAACGCGCGTCGTTATAAATGTTGCGCTCGGAAATCCGTGTCAGGATCTAAACGCGCGTCGTTATAAAACGCGCGTCGTTATACAACGTCCATGTCGGGATGC |
| PA14_60310 | pilY1 | CATCGCTGTTGTTATCAGCCAGGCGAGGCG | TTTCGTCCGCGAGTGACCAGACGCGCGTCGTTATAAAACGCGCGTCGTTATAAACATCGCTGTTGTTATCAGCCAGGCGAGGCGAAACGCGCGTCGTTATAAAACGCGCGTCGTTATACAACGTCCATGTCGGGATGC |
| PA14_60310 | pilY1 | TTACATCGAGTGCGAACAGGCCGCGACCTC | TTTCGTCCGCGAGTGACCAGACGCGCGTCGTTATAAAACGCGCGTCGTTATAAATTACATCGAGTGCGAACAGGCCGCGACCTCAAACGCGCGTCGTTATAAAACGCGCGTCGTTATACAACGTCCATGTCGGGATGC |
| PA14_60310 | pilY1 | GGCTTCAGGTCATTGTCGATGCTGGGACGT | TTTCGTCCGCGAGTGACCAGACGCGCGTCGTTATAAAACGCGCGTCGTTATAAAGGCTTCAGGTCATTGTCGATGCTGGGACGTAAACGCGCGTCGTTATAAAACGCGCGTCGTTATACAACGTCCATGTCGGGATGC |
| PA14_60310 | pilY1 | ACGGATGAAGTCGACCCGCTGTGCTCCTTT | TTTCGTCCGCGAGTGACCAGACGCGCGTCGTTATAAAACGCGCGTCGTTATAAAACGGATGAAGTCGACCCGCTGTGCTCCTTTAAACGCGCGTCGTTATAAAACGCGCGTCGTTATACAACGTCCATGTCGGGATGC |
| PA14_60310 | pilY1 | CAGATCGCCAGCATAGGCGTAGTCAGCAAT | TTTCGTCCGCGAGTGACCAGACGCGCGTCGTTATAAAACGCGCGTCGTTATAAACAGATCGCCAGCATAGGCGTAGTCAGCAATAAACGCGCGTCGTTATAAAACGCGCGTCGTTATACAACGTCCATGTCGGGATGC |
| PA14_60310 | pilY1 | ACGAGTGCACGATGTCCCCTAACACCGCCT | TTTCGTCCGCGAGTGACCAGACGCGCGTCGTTATAAAACGCGCGTCGTTATAAAACGAGTGCACGATGTCCCCTAACACCGCCTAAACGCGCGTCGTTATAAAACGCGCGTCGTTATACAACGTCCATGTCGGGATGC |
| PA14_60310 | pilY1 | CCTACGATGACGATGTAGCCCTTACGGCTA | TTTCGTCCGCGAGTGACCAGACGCGCGTCGTTATAAAACGCGCGTCGTTATAAACCTACGATGACGATGTAGCCCTTACGGCTAAAACGCGCGTCGTTATAAAACGCGCGTCGTTATACAACGTCCATGTCGGGATGC |
| PA14_60310 | pilY1 | GTCTGGTTCTGGTTGTCGAGATTGCCGGGG | TTTCGTCCGCGAGTGACCAGACGCGCGTCGTTATAAAACGCGCGTCGTTATAAAGTCTGGTTCTGGTTGTCGAGATTGCCGGGGAAACGCGCGTCGTTATAAAACGCGCGTCGTTATACAACGTCCATGTCGGGATGC |
| PA14_60310 | pilY1 | CAGCCAAGTGGCTAACTGACTTCCCTCAAT | TTTCGTCCGCGAGTGACCAGACGCGCGTCGTTATAAAACGCGCGTCGTTATAAACAGCCAAGTGGCTAACTGACTTCCCTCAATAAACGCGCGTCGTTATAAAACGCGCGTCGTTATACAACGTCCATGTCGGGATGC |
| PA14_60310 | pilY1 | GCCGCTGAACGATACTCTGAAAGCAGAAGG | TTTCGTCCGCGAGTGACCAGACGCGCGTCGTTATAAAACGCGCGTCGTTATAAAGCCGCTGAACGATACTCTGAAAGCAGAAGGAAACGCGCGTCGTTATAAAACGCGCGTCGTTATACAACGTCCATGTCGGGATGC |
| PA14_60310 | pilY1 | ATAGAGCGTCATGGCTCGGCTGGTATCGGC | TTTCGTCCGCGAGTGACCAGACGCGCGTCGTTATAAAACGCGCGTCGTTATAAAATAGAGCGTCATGGCTCGGCTGGTATCGGCAAACGCGCGTCGTTATAAAACGCGCGTCGTTATACAACGTCCATGTCGGGATGC |
| PA14_60310 | pilY1 | CCAAGTTGGTGCAGAGTTCAATGTGAGCGA | TTTCGTCCGCGAGTGACCAGACGCGCGTCGTTATAAAACGCGCGTCGTTATAAACCAAGTTGGTGCAGAGTTCAATGTGAGCGAAAACGCGCGTCGTTATAAAACGCGCGTCGTTATACAACGTCCATGTCGGGATGC |
| PA14_60310 | pilY1 | CGTTACTCTTGTCGCTGGGGTTGAAGATGA | TTTCGTCCGCGAGTGACCAGACGCGCGTCGTTATAAAACGCGCGTCGTTATAAACGTTACTCTTGTCGCTGGGGTTGAAGATGAAAACGCGCGTCGTTATAAAACGCGCGTCGTTATACAACGTCCATGTCGGGATGC |
| PA14_60310 | pilY1 | CGCGGAAGTCATTGCTGGTTTGGAGGAGGA | TTTCGTCCGCGAGTGACCAGACGCGCGTCGTTATAAAACGCGCGTCGTTATAAACGCGGAAGTCATTGCTGGTTTGGAGGAGGAAAACGCGCGTCGTTATAAAACGCGCGTCGTTATACAACGTCCATGTCGGGATGC |
| PA14_60310 | pilY1 | GAAACGATGATCTGATCGCCTGACAGTGTC | TTTCGTCCGCGAGTGACCAGACGCGCGTCGTTATAAAACGCGCGTCGTTATAAAGAAACGATGATCTGATCGCCTGACAGTGTCAAACGCGCGTCGTTATAAAACGCGCGTCGTTATACAACGTCCATGTCGGGATGC |
| PA14_60310 | pilY1 | TAGCCACATCCGAGTTCGCGGTACCGGTTG | TTTCGTCCGCGAGTGACCAGACGCGCGTCGTTATAAAACGCGCGTCGTTATAAATAGCCACATCCGAGTTCGCGGTACCGGTTGAAACGCGCGTCGTTATAAAACGCGCGTCGTTATACAACGTCCATGTCGGGATGC |
| PA14_60310 | pilY1 | TTCAGGGCCGTGGCGGCATAACTGTCAGAG | TTTCGTCCGCGAGTGACCAGACGCGCGTCGTTATAAAACGCGCGTCGTTATAAATTCAGGGCCGTGGCGGCATAACTGTCAGAGAAACGCGCGTCGTTATAAAACGCGCGTCGTTATACAACGTCCATGTCGGGATGC |
| PA14_51410 | pqsC | GAAGTCGAGCAGGTTGGAGATGTACTCGCT | TTTCGTCCGCGAGTGACCAGTATTGGCTACGGAGCAATATTGGCTACGGAGCAAGAAGTCGAGCAGGTTGGAGATGTACTCGCTAATATTGGCTACGGAGCAATATTGGCTACGGAGCCAACGTCCATGTCGGGATGC |
| PA14_51410 | pqsC | GTACGGGATGGAAACGGAGATCATCACGCC | TTTCGTCCGCGAGTGACCAGTATTGGCTACGGAGCAATATTGGCTACGGAGCAAGTACGGGATGGAAACGGAGATCATCACGCCAATATTGGCTACGGAGCAATATTGGCTACGGAGCCAACGTCCATGTCGGGATGC |
| PA14_51410 | pqsC | TATCGCCCATCGTCAGCTGGTACTGCTCCG | TTTCGTCCGCGAGTGACCAGTATTGGCTACGGAGCAATATTGGCTACGGAGCAATATCGCCCATCGTCAGCTGGTACTGCTCCGAATATTGGCTACGGAGCAATATTGGCTACGGAGCCAACGTCCATGTCGGGATGC |
| PA14_51410 | pqsC | ACTCCATCTGCGAATCCAATGGCAGCGCCC | TTTCGTCCGCGAGTGACCAGTATTGGCTACGGAGCAATATTGGCTACGGAGCAAACTCCATCTGCGAATCCAATGGCAGCGCCCAATATTGGCTACGGAGCAATATTGGCTACGGAGCCAACGTCCATGTCGGGATGC |
| PA14_51410 | pqsC | TCGTTTTCGTAGCTGCGAGCCGGAAGTTCA | TTTCGTCCGCGAGTGACCAGTATTGGCTACGGAGCAATATTGGCTACGGAGCAATCGTTTTCGTAGCTGCGAGCCGGAAGTTCAAATATTGGCTACGGAGCAATATTGGCTACGGAGCCAACGTCCATGTCGGGATGC |
| PA14_51410 | pqsC | ATGTCCACGCTATCCGGCGCAGTATCGCTG | TTTCGTCCGCGAGTGACCAGTATTGGCTACGGAGCAATATTGGCTACGGAGCAAATGTCCACGCTATCCGGCGCAGTATCGCTGAATATTGGCTACGGAGCAATATTGGCTACGGAGCCAACGTCCATGTCGGGATGC |
| PA14_51410 | pqsC | TTGCCACCAAGACTCGCTGAGGTCCGGTAC | TTTCGTCCGCGAGTGACCAGTATTGGCTACGGAGCAATATTGGCTACGGAGCAATTGCCACCAAGACTCGCTGAGGTCCGGTACAATATTGGCTACGGAGCAATATTGGCTACGGAGCCAACGTCCATGTCGGGATGC |
| PA14_51410 | pqsC | ACAGCACGTTGGCCATGCGCGGATAGAGAC | TTTCGTCCGCGAGTGACCAGTATTGGCTACGGAGCAATATTGGCTACGGAGCAAACAGCACGTTGGCCATGCGCGGATAGAGACAATATTGGCTACGGAGCAATATTGGCTACGGAGCCAACGTCCATGTCGGGATGC |
| PA14_51410 | pqsC | ACACCACCAGCACTTTCTCGGCCTTGCCCT | TTTCGTCCGCGAGTGACCAGTATTGGCTACGGAGCAATATTGGCTACGGAGCAAACACCACCAGCACTTTCTCGGCCTTGCCCTAATATTGGCTACGGAGCAATATTGGCTACGGAGCCAACGTCCATGTCGGGATGC |
| PA14_51410 | pqsC | ACGAAGCTGGCCATCTTGTTCTGGCCGTCG | TTTCGTCCGCGAGTGACCAGTATTGGCTACGGAGCAATATTGGCTACGGAGCAAACGAAGCTGGCCATCTTGTTCTGGCCGTCGAATATTGGCTACGGAGCAATATTGGCTACGGAGCCAACGTCCATGTCGGGATGC |
| PA14_51410 | pqsC | TAGAACGTGGCGTCGCTGTGTTCGGCCGAA | TTTCGTCCGCGAGTGACCAGTATTGGCTACGGAGCAATATTGGCTACGGAGCAATAGAACGTGGCGTCGCTGTGTTCGGCCGAAAATATTGGCTACGGAGCAATATTGGCTACGGAGCCAACGTCCATGTCGGGATGC |
| PA14_51410 | pqsC | ACCACCAGGCTGAACTCGTTCTCGCCGTTC | TTTCGTCCGCGAGTGACCAGTATTGGCTACGGAGCAATATTGGCTACGGAGCAAACCACCAGGCTGAACTCGTTCTCGCCGTTCAATATTGGCTACGGAGCAATATTGGCTACGGAGCCAACGTCCATGTCGGGATGC |
| PA14_51410 | pqsC | AACGAGAAATACAGCCGCGGCTTGGCCTCG | TTTCGTCCGCGAGTGACCAGTATTGGCTACGGAGCAATATTGGCTACGGAGCAAAACGAGAAATACAGCCGCGGCTTGGCCTCGAATATTGGCTACGGAGCAATATTGGCTACGGAGCCAACGTCCATGTCGGGATGC |
| PA14_51410 | pqsC | GGTGATCGCTGCCAGTTTGACCTTATGCAT | TTTCGTCCGCGAGTGACCAGTATTGGCTACGGAGCAATATTGGCTACGGAGCAAGGTGATCGCTGCCAGTTTGACCTTATGCATAATATTGGCTACGGAGCAATATTGGCTACGGAGCCAACGTCCATGTCGGGATGC |
| PA14_51410 | pqsC | GCTGGCTGGTGGAAGACGAAATAATCGATG | TTTCGTCCGCGAGTGACCAGTATTGGCTACGGAGCAATATTGGCTACGGAGCAAGCTGGCTGGTGGAAGACGAAATAATCGATGAATATTGGCTACGGAGCAATATTGGCTACGGAGCCAACGTCCATGTCGGGATGC |
| PA14_30630 | pqsH | CCCACCCAGACTGGAAGGCATCGACATCAG | TTTCGTCCGCGAGTGACCAGATACGGACAATTCGGAAATACGGACAATTCGGAACCCACCCAGACTGGAAGGCATCGACATCAGAAATACGGACAATTCGGAAATACGGACAATTCGGCAACGTCCATGTCGGGATGC |
| PA14_30630 | pqsH | TCGTGGGCGTATACGTTGATGCCGGCCAAC | TTTCGTCCGCGAGTGACCAGATACGGACAATTCGGAAATACGGACAATTCGGAATCGTGGGCGTATACGTTGATGCCGGCCAACAAATACGGACAATTCGGAAATACGGACAATTCGGCAACGTCCATGTCGGGATGC |
| PA14_30630 | pqsH | ACCGACCACCAGCGAACAGTCGTGGACAGT | TTTCGTCCGCGAGTGACCAGATACGGACAATTCGGAAATACGGACAATTCGGAAACCGACCACCAGCGAACAGTCGTGGACAGTAAATACGGACAATTCGGAAATACGGACAATTCGGCAACGTCCATGTCGGGATGC |
| PA14_30630 | pqsH | ACCAAGGCGGCCGACGATCCAGGATTGCTT | TTTCGTCCGCGAGTGACCAGATACGGACAATTCGGAAATACGGACAATTCGGAAACCAAGGCGGCCGACGATCCAGGATTGCTTAAATACGGACAATTCGGAAATACGGACAATTCGGCAACGTCCATGTCGGGATGC |
| PA14_30630 | pqsH | TAACGTCGCACGCTCGAACGAATGCCATCC | TTTCGTCCGCGAGTGACCAGATACGGACAATTCGGAAATACGGACAATTCGGAATAACGTCGCACGCTCGAACGAATGCCATCCAAATACGGACAATTCGGAAATACGGACAATTCGGCAACGTCCATGTCGGGATGC |
| PA14_30630 | pqsH | TCAGGGTCGCGTAGACATACATCTCGCGCG | TTTCGTCCGCGAGTGACCAGATACGGACAATTCGGAAATACGGACAATTCGGAATCAGGGTCGCGTAGACATACATCTCGCGCGAAATACGGACAATTCGGAAATACGGACAATTCGGCAACGTCCATGTCGGGATGC |
| PA14_30630 | pqsH | ATACCGGTACCGATGGGCCTGATCTCGCTG | TTTCGTCCGCGAGTGACCAGATACGGACAATTCGGAAATACGGACAATTCGGAAATACCGGTACCGATGGGCCTGATCTCGCTGAAATACGGACAATTCGGAAATACGGACAATTCGGCAACGTCCATGTCGGGATGC |
| PA14_30630 | pqsH | ACCGACACCGGTGAAAAGACGCTGGTGCAG | TTTCGTCCGCGAGTGACCAGATACGGACAATTCGGAAATACGGACAATTCGGAAACCGACACCGGTGAAAAGACGCTGGTGCAGAAATACGGACAATTCGGAAATACGGACAATTCGGCAACGTCCATGTCGGGATGC |
| PA14_30630 | pqsH | CAGCCAGTCGATGCCTGCCTTGGTGAATTC | TTTCGTCCGCGAGTGACCAGATACGGACAATTCGGAAATACGGACAATTCGGAACAGCCAGTCGATGCCTGCCTTGGTGAATTCAAATACGGACAATTCGGAAATACGGACAATTCGGCAACGTCCATGTCGGGATGC |
| PA14_30630 | pqsH | AATGCGCGACTCATCCAGCCCCTCCAGTAG | TTTCGTCCGCGAGTGACCAGATACGGACAATTCGGAAATACGGACAATTCGGAAAATGCGCGACTCATCCAGCCCCTCCAGTAGAAATACGGACAATTCGGAAATACGGACAATTCGGCAACGTCCATGTCGGGATGC |
| PA14_30630 | pqsH | TGATCTGGATGAAGCCGAGGCGCTTGCCGT | TTTCGTCCGCGAGTGACCAGATACGGACAATTCGGAAATACGGACAATTCGGAATGATCTGGATGAAGCCGAGGCGCTTGCCGTAAATACGGACAATTCGGAAATACGGACAATTCGGCAACGTCCATGTCGGGATGC |
| PA14_30630 | pqsH | TGGACAACGCCGTCAACGCATTGCTCGCCA | TTTCGTCCGCGAGTGACCAGATACGGACAATTCGGAAATACGGACAATTCGGAATGGACAACGCCGTCAACGCATTGCTCGCCAAAATACGGACAATTCGGAAATACGGACAATTCGGCAACGTCCATGTCGGGATGC |
| PA14_30630 | pqsH | TTCGTGGTCGAGTCCGTCGAGGATCTGCAC | TTTCGTCCGCGAGTGACCAGATACGGACAATTCGGAAATACGGACAATTCGGAATTCGTGGTCGAGTCCGTCGAGGATCTGCACAAATACGGACAATTCGGAAATACGGACAATTCGGCAACGTCCATGTCGGGATGC |
| PA14_30630 | pqsH | AACTCCTCGAGGTCGTTGTGCACCAGCGTG | TTTCGTCCGCGAGTGACCAGATACGGACAATTCGGAAATACGGACAATTCGGAAAACTCCTCGAGGTCGTTGTGCACCAGCGTGAAATACGGACAATTCGGAAATACGGACAATTCGGCAACGTCCATGTCGGGATGC |
| PA14_30630 | pqsH | AGGTTTCGCCGGAATGACGCAAGGTCGCCT | TTTCGTCCGCGAGTGACCAGATACGGACAATTCGGAAATACGGACAATTCGGAAAGGTTTCGCCGGAATGACGCAAGGTCGCCTAAATACGGACAATTCGGAAATACGGACAATTCGGCAACGTCCATGTCGGGATGC |

| | | | |
|---|---|---|---|
| PA14_62710 | pnp | ACGCTGACGTCGTCCATGGTGACCAGTACG | TTTCGTCCGCGAGTGACCAGGTCGTAATAGCACGCAAGTCGTAATAGCACGCAAACGCTGACGTCGTCCATGGTGACCAGTACGAAGTCGTAATAGCACGCAAGTCGTAATAGCACGCCAACGTCCATGTCGGGATGC |
| PA14_62710 | pnp | TCTGCAGTTGCTCGTAGGTGGGGTTGAGGA | TTTCGTCCGCGAGTGACCAGGTCGTAATAGCACGCAAGTCGTAATAGCACGCAATCTGCAGTTGCTCGTAGGTGGGGTTGAGGAAAGTCGTAATAGCACGCAAGTCGTAATAGCACGCCAACGTCCATGTCGGGATGC |
| PA14_62710 | pnp | AGGTCAGGGTTTCCTTCTCGGAAGGACGTC | TTTCGTCCGCGAGTGACCAGGTCGTAATAGCACGCAAGTCGTAATAGCACGCAAAGGTCAGGGTTTCCTTCTCGGAAGGACGTCAAGTCGTAATAGCACGCAAGTCGTAATAGCACGCCAACGTCCATGTCGGGATGC |
| PA14_62710 | pnp | ACTCCACCATCAGCACGGCGTCTTCGGTAC | TTTCGTCCGCGAGTGACCAGGTCGTAATAGCACGCAAGTCGTAATAGCACGCAAACTCCACCATCAGCACGGCGTCTTCGGTACAAGTCGTAATAGCACGCAAGTCGTAATAGCACGCCAACGTCCATGTCGGGATGC |
| PA14_62710 | pnp | CGATGACGTCGCGGATCTTGTCCGAGTCGA | TTTCGTCCGCGAGTGACCAGGTCGTAATAGCACGCAAGTCGTAATAGCACGCAACGATGACGTCGCGGATCTTGTCCGAGTCGAAAGTCGTAATAGCACGCAAGTCGTAATAGCACGCCAACGTCCATGTCGGGATGC |
| PA14_62710 | pnp | TGACCTCGGAAGCCGGGAACTTGCCCTCTT | TTTCGTCCGCGAGTGACCAGGTCGTAATAGCACGCAAGTCGTAATAGCACGCAATGACCTCGGAAGCCGGGAACTTGCCCTCTTAAGTCGTAATAGCACGCAAGTCGTAATAGCACGCCAACGTCCATGTCGGGATGC |
| PA14_62710 | pnp | CGATCTCCATGATCTCTTCGGTGATACCGT | TTTCGTCCGCGAGTGACCAGGTCGTAATAGCACGCAAGTCGTAATAGCACGCAACGATCTCCATGATCTCTTCGGTGATACCGTAAGTCGTAATAGCACGCAAGTCGTAATAGCACGCCAACGTCCATGTCGGGATGC |
| PA14_62710 | pnp | CCGTTGGACTCGGTGATCTCGGAAACCACG | TTTCGTCCGCGAGTGACCAGGTCGTAATAGCACGCAAGTCGTAATAGCACGCAACCGTTGGACTCGGTGATCTCGGAAACCACGAAGTCGTAATAGCACGCAAGTCGTAATAGCACGCCAACGTCCATGTCGGGATGC |
| PA14_62710 | pnp | TAGCCGATTTCCGGATGGAAACCGACGCGG | TTTCGTCCGCGAGTGACCAGGTCGTAATAGCACGCAAGTCGTAATAGCACGCAATAGCCGATTTCCGGATGGAAACCGACGCGGAAGTCGTAATAGCACGCAAGTCGTAATAGCACGCCAACGTCCATGTCGGGATGC |
| PA14_62710 | pnp | GAACTGGAATTGCTTGGTTACCGGGTTCAC | TTTCGTCCGCGAGTGACCAGGTCGTAATAGCACGCAAGTCGTAATAGCACGCAAGAACTGGAATTGCTTGGTTACCGGGTTCACAAGTCGTAATAGCACGCAAGTCGTAATAGCACGCCAACGTCCATGTCGGGATGC |
| PA14_62710 | pnp | GCGATGTCCGGATCGGACTTCTTGTTGGTG | TTTCGTCCGCGAGTGACCAGGTCGTAATAGCACGCAAGTCGTAATAGCACGCAAGCGATGTCCGGATCGGACTTCTTGTTGGTGAAGTCGTAATAGCACGCAAGTCGTAATAGCACGCCAACGTCCATGTCGGGATGC |
| PA14_62710 | pnp | TCTCTTCGCAGATACCACGGATGGTCGCGC | TTTCGTCCGCGAGTGACCAGGTCGTAATAGCACGCAAGTCGTAATAGCACGCAATCTCTTCGCAGATACCACGGATGGTCGCGCAAGTCGTAATAGCACGCAAGTCGTAATAGCACGCCAACGTCCATGTCGGGATGC |
| PA14_62710 | pnp | TGGTAGTGCACGGACAGCGGGAAGAAGTCG | TTTCGTCCGCGAGTGACCAGGTCGTAATAGCACGCAAGTCGTAATAGCACGCAATGGTAGTGCACGGACAGCGGGAAGAAGTCGAAGTCGTAATAGCACGCAAGTCGTAATAGCACGCCAACGTCCATGTCGGGATGC |
| PA14_62710 | pnp | GCTTCCTTGGTTTCGCCGTAGATCTTCACG | TTTCGTCCGCGAGTGACCAGGTCGTAATAGCACGCAAGTCGTAATAGCACGCAAGCTTCCTTGGTTTCGCCGTAGATCTTCACGAAGTCGTAATAGCACGCAAGTCGTAATAGCACGCCAACGTCCATGTCGGGATGC |
| PA14_62710 | pnp | TCACGCTTGAAGAAACCGCCGGGGATACGG | TTTCGTCCGCGAGTGACCAGGTCGTAATAGCACGCAAGTCGTAATAGCACGCAATCACGCTTGAAGAAACCGCCGGGGATACGGAAGTCGTAATAGCACGCAAGTCGTAATAGCACGCCAACGTCCATGTCGGGATGC |
| PA14_62710 | pnp | AAACCTTCGGGGAACAGCGGACGGATCGGA | TTTCGTCCGCGAGTGACCAGGTCGTAATAGCACGCAAGTCGTAATAGCACGCAAAAACCTTCGGGGAACAGCGGACGGATCGGAAAGTCGTAATAGCACGCAAGTCGTAATAGCACGCCAACGTCCATGTCGGGATGC |
| PA14_62710 | pnp | AACGACGGTGCAGACGACCTGCACTTCATT | TTTCGTCCGCGAGTGACCAGGTCGTAATAGCACGCAAGTCGTAATAGCACGCAAAACGACGGTGCAGACGACCTGCACTTCATTAAGTCGTAATAGCACGCAAGTCGTAATAGCACGCCAACGTCCATGTCGGGATGC |
| PA14_62710 | pnp | AGCATCTGGTCTTCGGTCAGTTCGTCGGCC | TTTCGTCCGCGAGTGACCAGGTCGTAATAGCACGCAAGTCGTAATAGCACGCAAAGCATCTGGTCTTCGGTCAGTTCGTCGGCCAAGTCGTAATAGCACGCAAGTCGTAATAGCACGCCAACGTCCATGTCGGGATGC |
| PA14_62710 | pnp | TGTTGACGAAGGCCCCGAAGTCGACGATGC | TTTCGTCCGCGAGTGACCAGGTCGTAATAGCACGCAAGTCGTAATAGCACGCAATGTTGACGAAGGCCCCGAAGTCGACGATGCAAGTCGTAATAGCACGCAAGTCGTAATAGCACGCCAACGTCCATGTCGGGATGC |
| PA14_62710 | pnp | TGATCGACAGCTTGATGCGGCCACGGTTGT | TTTCGTCCGCGAGTGACCAGGTCGTAATAGCACGCAAGTCGTAATAGCACGCAATGATCGACAGCTTGATGCGGCCACGGTTGTAAGTCGTAATAGCACGCAAGTCGTAATAGCACGCCAACGTCCATGTCGGGATGC |
| PA14_42350 | pscC | TCGACTTGTTGTTGCCCAGGCGAATCCCGG | TTTCGTCCGCGAGTGACCAGTCTCGTGATACGGAAAATCTCGTGATACGGAAAATCGACTTGTTGTTGCCCAGGCGAATCCCGGAATCTCGTGATACGGAAAATCTCGTGATACGGAACAACGTCCATGTCGGGATGC |
| PA14_42350 | pscC | ATACATGGGCAACCGGTCCTTGTGGTCGCG | TTTCGTCCGCGAGTGACCAGTCTCGTGATACGGAAAATCTCGTGATACGGAAAAATACATGGGCAACCGGTCCTTGTGGTCGCGAATCTCGTGATACGGAAAATCTCGTGATACGGAACAACGTCCATGTCGGGATGC |
| PA14_42350 | pscC | GTGGTCTTGAATACGTACAGCACGGTGCCG | TTTCGTCCGCGAGTGACCAGTCTCGTGATACGGAAAATCTCGTGATACGGAAAAGTGGTCTTGAATACGTACAGCACGGTGCCGAATCTCGTGATACGGAAAATCTCGTGATACGGAACAACGTCCATGTCGGGATGC |
| PA14_42350 | pscC | TCAACTGCAGGAAGGCCTGCGGGCTTTCCA | TTTCGTCCGCGAGTGACCAGTCTCGTGATACGGAAAATCTCGTGATACGGAAAATCAACTGCAGGAAGGCCTGCGGGCTTTCCAAATCTCGTGATACGGAAAATCTCGTGATACGGAACAACGTCCATGTCGGGATGC |
| PA14_42350 | pscC | AGGCATCGTAGTTGGCGCCGAAGTTGGCCA | TTTCGTCCGCGAGTGACCAGTCTCGTGATACGGAAAATCTCGTGATACGGAAAAAGGCATCGTAGTTGGCGCCGAAGTTGGCCAAATCTCGTGATACGGAAAATCTCGTGATACGGAACAACGTCCATGTCGGGATGC |
| PA14_42350 | pscC | TCTCGTCGTCGCGGTACTCGATCTTGCGGT | TTTCGTCCGCGAGTGACCAGTCTCGTGATACGGAAAATCTCGTGATACGGAAAATCTCGTCGTCGCGGTACTCGATCTTGCGGTAATCTCGTGATACGGAAAATCTCGTGATACGGAACAACGTCCATGTCGGGATGC |
| PA14_42350 | pscC | TGTGCAGGCTGAGGCTGATCTCCGGCGTAT | TTTCGTCCGCGAGTGACCAGTCTCGTGATACGGAAAATCTCGTGATACGGAAAATGTGCAGGCTGAGGCTGATCTCCGGCGTATAATCTCGTGATACGGAAAATCTCGTGATACGGAACAACGTCCATGTCGGGATGC |
| PA14_42350 | pscC | TGCGCAGGGTGTATTGCTGCTCCAGCACCT | TTTCGTCCGCGAGTGACCAGTCTCGTGATACGGAAAATCTCGTGATACGGAAAATGCGCAGGGTGTATTGCTGCTCCAGCACCTAATCTCGTGATACGGAAAATCTCGTGATACGGAACAACGTCCATGTCGGGATGC |
| PA14_42350 | pscC | ATAGCGCAGCGGGAAGATCTCCACGCTGAG | TTTCGTCCGCGAGTGACCAGTCTCGTGATACGGAAAATCTCGTGATACGGAAAAATAGCGCAGCGGGAAGATCTCCACGCTGAGAATCTCGTGATACGGAAAATCTCGTGATACGGAACAACGTCCATGTCGGGATGC |
| PA14_42350 | pscC | ATAGTCGTAAGGCAGGCTGGGCCAGTCCAG | TTTCGTCCGCGAGTGACCAGTCTCGTGATACGGAAAATCTCGTGATACGGAAAAATAGTCGTAAGGCAGGCTGGGCCAGTCCAGAATCTCGTGATACGGAAAATCTCGTGATACGGAACAACGTCCATGTCGGGATGC |
| PA14_42350 | pscC | TCGATGATCGACAAGCCGACCTCGATCCGC | TTTCGTCCGCGAGTGACCAGTCTCGTGATACGGAAAATCTCGTGATACGGAAAATCGATGATCGACAAGCCGACCTCGATCCGCAATCTCGTGATACGGAAAATCTCGTGATACGGAACAACGTCCATGTCGGGATGC |
| PA14_42350 | pscC | TTTTCAGCATGGTGCCGTAGGTGATCGCCT | TTTCGTCCGCGAGTGACCAGTCTCGTGATACGGAAAATCTCGTGATACGGAAAATTTTCAGCATGGTGCCGTAGGTGATCGCCTAATCTCGTGATACGGAAAATCTCGTGATACGGAACAACGTCCATGTCGGGATGC |
| PA14_42350 | pscC | GTAGTACCAGCCGAGGTTGTAGAGGGAGGC | TTTCGTCCGCGAGTGACCAGTCTCGTGATACGGAAAATCTCGTGATACGGAAAAGTAGTACCAGCCGAGGTTGTAGAGGGAGGCAATCTCGTGATACGGAAAATCTCGTGATACGGAACAACGTCCATGTCGGGATGC |
| PA14_42350 | pscC | CGCACATAGTAGGTTTCGCTCTGGTCGAGC | TTTCGTCCGCGAGTGACCAGTCTCGTGATACGGAAAATCTCGTGATACGGAAAACGCACATAGTAGGTTTCGCTCTGGTCGAGCAATCTCGTGATACGGAAAATCTCGTGATACGGAACAACGTCCATGTCGGGATGC |
| PA14_42350 | pscC | CTGGTCGTTGACCTTGTCGCTGACGATCAC | TTTCGTCCGCGAGTGACCAGTCTCGTGATACGGAAAATCTCGTGATACGGAAAACTGGTCGTTGACCTTGTCGCTGACGATCACAATCTCGTGATACGGAAAATCTCGTGATACGGAACAACGTCCATGTCGGGATGC |
| PA14_35670 | pslG | TGACCCCGAGGAAGTCTTTCCAGACCACCG | TTTCGTCCGCGAGTGACCAGCTGCGAGACATTGCTAACTGCGAGACATTGCTAATGACCCCGAGGAAGTCTTTCCAGACCACCGAACTGCGAGACATTGCTAACTGCGAGACATTGCTCAACGTCCATGTCGGGATGC |
| PA14_35670 | pslG | TGTAGCGCTCCGGGCTGAACCAGAGGAACT | TTTCGTCCGCGAGTGACCAGCTGCGAGACATTGCTAACTGCGAGACATTGCTAATGTAGCGCTCCGGGCTGAACCAGAGGAACTAACTGCGAGACATTGCTAACTGCGAGACATTGCTCAACGTCCATGTCGGGATGC |
| PA14_35670 | pslG | TGTCCTGCAACTCCTTCGGCCCCTTGTAGG | TTTCGTCCGCGAGTGACCAGCTGCGAGACATTGCTAACTGCGAGACATTGCTAATGTCCTGCAACTCCTTCGGCCCCTTGTAGGAACTGCGAGACATTGCTAACTGCGAGACATTGCTCAACGTCCATGTCGGGATGC |
| PA14_35670 | pslG | ACCAGGTAGAACACCGACTTCAGCTGGCGC | TTTCGTCCGCGAGTGACCAGCTGCGAGACATTGCTAACTGCGAGACATTGCTAAACCAGGTAGAACACCGACTTCAGCTGGCGCAACTGCGAGACATTGCTAACTGCGAGACATTGCTCAACGTCCATGTCGGGATGC |
| PA14_35670 | pslG | AGTCCAGGTTCCACGGGTAGTTGCCTTCCG | TTTCGTCCGCGAGTGACCAGCTGCGAGACATTGCTAACTGCGAGACATTGCTAAAGTCCAGGTTCCACGGGTAGTTGCCTTCCGAACTGCGAGACATTGCTAACTGCGAGACATTGCTCAACGTCCATGTCGGGATGC |
| PA14_35670 | pslG | TCTCGCTGAAGAAGGCCATGCCGGCGGAAA | TTTCGTCCGCGAGTGACCAGCTGCGAGACATTGCTAACTGCGAGACATTGCTAATCTCGCTGAAGAAGGCCATGCCGGCGGAAAAACTGCGAGACATTGCTAACTGCGAGACATTGCTCAACGTCCATGTCGGGATGC |
| PA14_35670 | pslG | TCAGGTTGGGCTCGTTCCATACCTGCCAGG | TTTCGTCCGCGAGTGACCAGCTGCGAGACATTGCTAACTGCGAGACATTGCTAATCAGGTTGGGCTCGTTCCATACCTGCCAGGAACTGCGAGACATTGCTAACTGCGAGACATTGCTCAACGTCCATGTCGGGATGC |
| PA14_35670 | pslG | ACTGGTCCTGGAAGGGCGAATAGAACGGCG | TTTCGTCCGCGAGTGACCAGCTGCGAGACATTGCTAACTGCGAGACATTGCTAAACTGGTCCTGGAAGGGCGAATAGAACGGCGAACTGCGAGACATTGCTAACTGCGAGACATTGCTCAACGTCCATGTCGGGATGC |
| PA14_35670 | pslG | CTGGGTATAGGGGTGGTAGGTGGCGATGGT | TTTCGTCCGCGAGTGACCAGCTGCGAGACATTGCTAACTGCGAGACATTGCTAACTGGGTATAGGGGTGGTAGGTGGCGATGGTAACTGCGAGACATTGCTAACTGCGAGACATTGCTCAACGTCCATGTCGGGATGC |
| PA14_35670 | pslG | ATCCGGTCGTAGTCCAGCGCACTCATCAGC | TTTCGTCCGCGAGTGACCAGCTGCGAGACATTGCTAACTGCGAGACATTGCTAAATCCGGTCGTAGTCCAGCGCACTCATCAGCAACTGCGAGACATTGCTAACTGCGAGACATTGCTCAACGTCCATGTCGGGATGC |
| PA14_35670 | pslG | AGCTGGTACTGGTCTTCGGCGGTTTCCAGG | TTTCGTCCGCGAGTGACCAGCTGCGAGACATTGCTAACTGCGAGACATTGCTAAAGCTGGTACTGGTCTTCGGCGGTTTCCAGGAACTGCGAGACATTGCTAACTGCGAGACATTGCTCAACGTCCATGTCGGGATGC |
| PA14_35670 | pslG | GCCGCCCAGATAGAGTCCCTTACGTGCCAT | TTTCGTCCGCGAGTGACCAGCTGCGAGACATTGCTAACTGCGAGACATTGCTAAGCCGCCCAGATAGAGTCCCTTACGTGCCATAACTGCGAGACATTGCTAACTGCGAGACATTGCTCAACGTCCATGTCGGGATGC |
| PA14_35670 | pslG | TCTGCAGGCTGGACTTCACCGGGACTTCCA | TTTCGTCCGCGAGTGACCAGCTGCGAGACATTGCTAACTGCGAGACATTGCTAATCTGCAGGCTGGACTTCACCGGGACTTCCAAACTGCGAGACATTGCTAACTGCGAGACATTGCTCAACGTCCATGTCGGGATGC |
| PA14_35670 | pslG | CGAGATCCTTGACCAACTGGTCGAGGGAGG | TTTCGTCCGCGAGTGACCAGCTGCGAGACATTGCTAACTGCGAGACATTGCTAACGAGATCCTTGACCAACTGGTCGAGGGAGGAACTGCGAGACATTGCTAACTGCGAGACATTGCTCAACGTCCATGTCGGGATGC |
| PA14_35670 | pslG | AGAACACCAGCAGCAGTACCACGGCGAGCA | TTTCGTCCGCGAGTGACCAGCTGCGAGACATTGCTAACTGCGAGACATTGCTAAAGAACACCAGCAGCAGTACCACGGCGAGCAAACTGCGAGACATTGCTAACTGCGAGACATTGCTCAACGTCCATGTCGGGATGC |
| PA14_70860 | pstS | TCCTCGACGAAGGCAGCGCCTTCCTTCTTG | TTTCGTCCGCGAGTGACCAGCGCTAATTGCCGAGTAACGCTAATTGCCGAGTAATCCTCGACGAAGGCAGCGCCTTCCTTCTTGAACGCTAATTGCCGAGTAACGCTAATTGCCGAGTCAACGTCCATGTCGGGATGC |
| PA14_70860 | pstS | CGGGTTGTCCTTGTGCACGAAGATCGCCAG | TTTCGTCCGCGAGTGACCAGCGCTAATTGCCGAGTAACGCTAATTGCCGAGTAACGGGTTGTCCTTGTGCACGAAGATCGCCAGAACGCTAATTGCCGAGTAACGCTAATTGCCGAGTCAACGTCCATGTCGGGATGC |
| PA14_70860 | pstS | GATGTAGCCGTCCTTGACCACGACCTGCTG | TTTCGTCCGCGAGTGACCAGCGCTAATTGCCGAGTAACGCTAATTGCCGAGTAAGATGTAGCCGTCCTTGACCACGACCTGCTGAACGCTAATTGCCGAGTAACGCTAATTGCCGAGTCAACGTCCATGTCGGGATGC |
| PA14_70860 | pstS | TGGCTTTCTGATATTCCGGCAGCGCCGGGT | TTTCGTCCGCGAGTGACCAGCGCTAATTGCCGAGTAACGCTAATTGCCGAGTAATGGCTTTCTGATATTCCGGCAGCGCCGGGTAACGCTAATTGCCGAGTAACGCTAATTGCCGAGTCAACGTCCATGTCGGGATGC |
| PA14_70860 | pstS | TAGACGTAGAGGAAGCGCGACAGCGGGTAG | TTTCGTCCGCGAGTGACCAGCGCTAATTGCCGAGTAACGCTAATTGCCGAGTAATAGACGTAGAGGAAGCGCGACAGCGGGTAGAACGCTAATTGCCGAGTAACGCTAATTGCCGAGTCAACGTCCATGTCGGGATGC |
| PA14_70860 | pstS | TCATCAGGTTGGCCAGAGTGTCGGAGCCGA | TTTCGTCCGCGAGTGACCAGCGCTAATTGCCGAGTAACGCTAATTGCCGAGTAATCATCAGGTTGGCCAGAGTGTCGGAGCCGAAACGCTAATTGCCGAGTAACGCTAATTGCCGAGTCAACGTCCATGTCGGGATGC |
| PA14_70860 | pstS | CTTCACGCTGGCGGTCTTGTAGCCGATACC | TTTCGTCCGCGAGTGACCAGCGCTAATTGCCGAGTAACGCTAATTGCCGAGTAACTTCACGCTGGCGGTCTTGTAGCCGATACCAACGCTAATTGCCGAGTAACGCTAATTGCCGAGTCAACGTCCATGTCGGGATGC |
| PA14_70860 | pstS | CCGGCAGCCTGGATCTGGATGTTCACGTTC | TTTCGTCCGCGAGTGACCAGCGCTAATTGCCGAGTAACGCTAATTGCCGAGTAACCGGCAGCCTGGATCTGGATGTTCACGTTCAACGCTAATTGCCGAGTAACGCTAATTGCCGAGTCAACGTCCATGTCGGGATGC |
| PA14_70860 | pstS | CGGTCTTGGACAGGACCAGCTTGAGGAACT | TTTCGTCCGCGAGTGACCAGCGCTAATTGCCGAGTAACGCTAATTGCCGAGTAACGGTCTTGGACAGGACCAGCTTGAGGAACTAACGCTAATTGCCGAGTAACGCTAATTGCCGAGTCAACGTCCATGTCGGGATGC |
| PA14_70860 | pstS | ACAGTTGCACCGGCTTCTTCGCCCAGTCAC | TTTCGTCCGCGAGTGACCAGCGCTAATTGCCGAGTAACGCTAATTGCCGAGTAAACAGTTGCACCGGCTTCTTCGCCCAGTCACAACGCTAATTGCCGAGTAACGCTAATTGCCGAGTCAACGTCCATGTCGGGATGC |
| PA14_70860 | pstS | AGTAGCCGATGCCGTTCAGCGACTGGCTGA | TTTCGTCCGCGAGTGACCAGCGCTAATTGCCGAGTAACGCTAATTGCCGAGTAAAGTAGCCGATGCCGTTCAGCGACTGGCTGAAACGCTAATTGCCGAGTAACGCTAATTGCCGAGTCAACGTCCATGTCGGGATGC |
| PA14_70860 | pstS | GAAGTCGCCTTTGCACAGGGCTTCTTCCTT | TTTCGTCCGCGAGTGACCAGCGCTAATTGCCGAGTAACGCTAATTGCCGAGTAAGAAGTCGCCTTTGCACAGGGCTTCTTCCTTAACGCTAATTGCCGAGTAACGCTAATTGCCGAGTCAACGTCCATGTCGGGATGC |
| PA14_70860 | pstS | CAGGGCCGCCATCAAACGCTTGAGTTTCAT | TTTCGTCCGCGAGTGACCAGCGCTAATTGCCGAGTAACGCTAATTGCCGAGTAACAGGGCCGCCATCAAACGCTTGAGTTTCATAACGCTAATTGCCGAGTAACGCTAATTGCCGAGTCAACGTCCATGTCGGGATGC |
| PA14_70860 | pstS | ATAGAGGCGCTTGTACTCCTCGGCCCACAT | TTTCGTCCGCGAGTGACCAGCGCTAATTGCCGAGTAACGCTAATTGCCGAGTAAATAGAGGCGCTTGTACTCCTCGGCCCACATAACGCTAATTGCCGAGTAACGCTAATTGCCGAGTCAACGTCCATGTCGGGATGC |
| PA14_70860 | pstS | AGGTCTTCACGTCCTGCTTGCTGCCGCACA | TTTCGTCCGCGAGTGACCAGCGCTAATTGCCGAGTAACGCTAATTGCCGAGTAAAGGTCTTCACGTCCTGCTTGCTGCCGCACAAACGCTAATTGCCGAGTAACGCTAATTGCCGAGTCAACGTCCATGTCGGGATGC |
| PA14_70860 | pstS | GATCCAGCGGCTTGTTCGGTGCCTTGTTCA | TTTCGTCCGCGAGTGACCAGCGCTAATTGCCGAGTAACGCTAATTGCCGAGTAAGATCCAGCGGCTTGTTCGGTGCCTTGTTCAAACGCTAATTGCCGAGTAACGCTAATTGCCGAGTCAACGTCCATGTCGGGATGC |
| PA14_70860 | pstS | ACTTCTGCTCGAAAGCCTGCAGCTCGACGT | TTTCGTCCGCGAGTGACCAGCGCTAATTGCCGAGTAACGCTAATTGCCGAGTAAACTTCTGCTCGAAAGCCTGCAGCTCGACGTAACGCTAATTGCCGAGTAACGCTAATTGCCGAGTCAACGTCCATGTCGGGATGC |
| PA14_70860 | pstS | TTCATCTTGCGGCTCATCGGGCCCAGGTTG | TTTCGTCCGCGAGTGACCAGCGCTAATTGCCGAGTAACGCTAATTGCCGAGTAATTCATCTTGCGGCTCATCGGGCCCAGGTTGAACGCTAATTGCCGAGTAACGCTAATTGCCGAGTCAACGTCCATGTCGGGATGC |
| PA14_70860 | pstS | AGTAGCCGTAGGTGCCGGAAACCGAGTTGC | TTTCGTCCGCGAGTGACCAGCGCTAATTGCCGAGTAACGCTAATTGCCGAGTAAAGTAGCCGTAGGTGCCGGAAACCGAGTTGCAACGCTAATTGCCGAGTAACGCTAATTGCCGAGTCAACGTCCATGTCGGGATGC |
| PA14_70860 | pstS | GCATCGACCTGCTGCATGGTCAGGCCTTTG | TTTCGTCCGCGAGTGACCAGCGCTAATTGCCGAGTAACGCTAATTGCCGAGTAAGCATCGACCTGCTGCATGGTCAGGCCTTTGAACGCTAATTGCCGAGTAACGCTAATTGCCGAGTCAACGTCCATGTCGGGATGC |
| PA14_04410 | ptsP | CTGTTCCTTCTCGCTGGGGAAGCGGTCGTT | TTTCGTCCGCGAGTGACCAGTGCCTTGTCACGCAAAATGCCTTGTCACGCAAAACTGTTCCTTCTCGCTGGGGAAGCGGTCGTTAATGCCTTGTCACGCAAAATGCCTTGTCACGCAACAACGTCCATGTCGGGATGC |
| PA14_04410 | ptsP | CCATGCTGACCTTGCCGATGGAACGCTTGT | TTTCGTCCGCGAGTGACCAGTGCCTTGTCACGCAAAATGCCTTGTCACGCAAAACCATGCTGACCTTGCCGATGGAACGCTTGTAATGCCTTGTCACGCAAAATGCCTTGTCACGCAACAACGTCCATGTCGGGATGC |
| PA14_04410 | ptsP | TACCGTCGACCTTGGAGTACGGCAGGTCGA | TTTCGTCCGCGAGTGACCAGTGCCTTGTCACGCAAAATGCCTTGTCACGCAAAATACCGTCGACCTTGGAGTACGGCAGGTCGAAATGCCTTGTCACGCAAAATGCCTTGTCACGCAACAACGTCCATGTCGGGATGC |
| PA14_04410 | ptsP | GCCGCCTTCAGATCCTTGGCGGAGTTCACT | TTTCGTCCGCGAGTGACCAGTGCCTTGTCACGCAAAATGCCTTGTCACGCAAAAGCCGCCTTCAGATCCTTGGCGGAGTTCACTAATGCCTTGTCACGCAAAATGCCTTGTCACGCAACAACGTCCATGTCGGGATGC |
| PA14_04410 | ptsP | TGGAAGGCACTGAGCTGCTCGCGGTAGATC | TTTCGTCCGCGAGTGACCAGTGCCTTGTCACGCAAAATGCCTTGTCACGCAAAATGGAAGGCACTGAGCTGCTCGCGGTAGATCAATGCCTTGTCACGCAAAATGCCTTGTCACGCAACAACGTCCATGTCGGGATGC |
| PA14_04410 | ptsP | TCATGGTGACGAGGAAGGCCTCCTCGCCTT | TTTCGTCCGCGAGTGACCAGTGCCTTGTCACGCAAAATGCCTTGTCACGCAAAATCATGGTGACGAGGAAGGCCTCCTCGCCTTAATGCCTTGTCACGCAAAATGCCTTGTCACGCAACAACGTCCATGTCGGGATGC |
| PA14_04410 | ptsP | TTGGCTTCCTGGATGCCCTTGCCGAGCTTG | TTTCGTCCGCGAGTGACCAGTGCCTTGTCACGCAAAATGCCTTGTCACGCAAAATTGGCTTCCTGGATGCCCTTGCCGAGCTTGAATGCCTTGTCACGCAAAATGCCTTGTCACGCAACAACGTCCATGTCGGGATGC |
| PA14_04410 | ptsP | CTGGACGATCTTGCGCAGCGTGTTGAGCAT | TTTCGTCCGCGAGTGACCAGTGCCTTGTCACGCAAAATGCCTTGTCACGCAAAACTGGACGATCTTGCGCAGCGTGTTGAGCATAATGCCTTGTCACGCAAAATGCCTTGTCACGCAACAACGTCCATGTCGGGATGC |
| PA14_04410 | ptsP | CGAGCATCATCAGGTAGACGTCGAACAGCG | TTTCGTCCGCGAGTGACCAGTGCCTTGTCACGCAAAATGCCTTGTCACGCAAAACGAGCATCATCAGGTAGACGTCGAACAGCGAATGCCTTGTCACGCAAAATGCCTTGTCACGCAACAACGTCCATGTCGGGATGC |
| PA14_04410 | ptsP | TCAGCTCGAAGCGCTGCACGTGCTCCATCA | TTTCGTCCGCGAGTGACCAGTGCCTTGTCACGCAAAATGCCTTGTCACGCAAAATCAGCTCGAAGCGCTGCACGTGCTCCATCAAATGCCTTGTCACGCAAAATGCCTTGTCACGCAACAACGTCCATGTCGGGATGC |
| PA14_04410 | ptsP | TCAGGACGAAACGCTGGGTCTCGGTGTCGA | TTTCGTCCGCGAGTGACCAGTGCCTTGTCACGCAAAATGCCTTGTCACGCAAAATCAGGACGAAACGCTGGGTCTCGGTGTCGAAATGCCTTGTCACGCAAAATGCCTTGTCACGCAACAACGTCCATGTCGGGATGC |
| PA14_04410 | ptsP | TTCCTTGCGCAACTGGCTGGCGAGCTTGCT | TTTCGTCCGCGAGTGACCAGTGCCTTGTCACGCAAAATGCCTTGTCACGCAAAATTCCTTGCGCAACTGGCTGGCGAGCTTGCTAATGCCTTGTCACGCAAAATGCCTTGTCACGCAACAACGTCCATGTCGGGATGC |
| PA14_04410 | ptsP | TGTCGTCGACCTGCTTGTCCGGCACCACTT | TTTCGTCCGCGAGTGACCAGTGCCTTGTCACGCAAAATGCCTTGTCACGCAAAATGTCGTCGACCTGCTTGTCCGGCACCACTTAATGCCTTGTCACGCAAAATGCCTTGTCACGCAACAACGTCCATGTCGGGATGC |
| PA14_04410 | ptsP | TGCGCTTGACCTCGTTGCCGATGGAGGCAT | TTTCGTCCGCGAGTGACCAGTGCCTTGTCACGCAAAATGCCTTGTCACGCAAAATGCGCTTGACCTCGTTGCCGATGGAGGCATAATGCCTTGTCACGCAAAATGCCTTGTCACGCAACAACGTCCATGTCGGGATGC |
| PA14_04410 | ptsP | AGGTACACCGAGCAGACCTGGGTACCCATG | TTTCGTCCGCGAGTGACCAGTGCCTTGTCACGCAAAATGCCTTGTCACGCAAAAAGGTACACCGAGCAGACCTGGGTACCCATGAATGCCTTGTCACGCAAAATGCCTTGTCACGCAACAACGTCCATGTCGGGATGC |
| PA14_33700 | pvdF | CACTTCGCCGGAATCGATGCCATTGTCCAC | TTTCGTCCGCGAGTGACCAGAGCACCTTTACATCGAAAGCACCTTTACATCGAACACTTCGCCGGAATCGATGCCATTGTCCACAAAGCACCTTTACATCGAAAGCACCTTTACATCGCAACGTCCATGTCGGGATGC |
| PA14_33700 | pvdF | AGTGGAACGATGCTCCGGTCCAGTACAGCG | TTTCGTCCGCGAGTGACCAGAGCACCTTTACATCGAAAGCACCTTTACATCGAAAGTGGAACGATGCTCCGGTCCAGTACAGCGAAAGCACCTTTACATCGAAAGCACCTTTACATCGCAACGTCCATGTCGGGATGC |
| PA14_33700 | pvdF | TCATGCAGGCGTCGCTCGAAATCGCTCTTG | TTTCGTCCGCGAGTGACCAGAGCACCTTTACATCGAAAGCACCTTTACATCGAATCATGCAGGCGTCGCTCGAAATCGCTCTTGAAAGCACCTTTACATCGAAAGCACCTTTACATCGCAACGTCCATGTCGGGATGC |
| PA14_33700 | pvdF | TAGATCACGCCCACGAGCTCGTAGGCATCG | TTTCGTCCGCGAGTGACCAGAGCACCTTTACATCGAAAGCACCTTTACATCGAATAGATCACGCCCACGAGCTCGTAGGCATCGAAAGCACCTTTACATCGAAAGCACCTTTACATCGCAACGTCCATGTCGGGATGC |
| PA14_33700 | pvdF | ACCGCTGTTCGCCCTTGTACGGCACGTACT | TTTCGTCCGCGAGTGACCAGAGCACCTTTACATCGAAAGCACCTTTACATCGAAACCGCTGTTCGCCCTTGTACGGCACGTACTAAAGCACCTTTACATCGAAAGCACCTTTACATCGCAACGTCCATGTCGGGATGC |
| PA14_33700 | pvdF | GTCGGGTGAAATCTCCGTTTTCAGCACATC | TTTCGTCCGCGAGTGACCAGAGCACCTTTACATCGAAAGCACCTTTACATCGAAGTCGGGTGAAATCTCCGTTTTCAGCACATCAAAGCACCTTTACATCGAAAGCACCTTTACATCGCAACGTCCATGTCGGGATGC |
| PA14_33700 | pvdF | GGGACCAGATATAGGCCAGTTTCCTTTTCG | TTTCGTCCGCGAGTGACCAGAGCACCTTTACATCGAAAGCACCTTTACATCGAAGGGACCAGATATAGGCCAGTTTCCTTTTCGAAAGCACCTTTACATCGAAAGCACCTTTACATCGCAACGTCCATGTCGGGATGC |
| PA14_33700 | pvdF | ACCATTGCTGCCCGGGACGATAGGCGAAAC | TTTCGTCCGCGAGTGACCAGAGCACCTTTACATCGAAAGCACCTTTACATCGAAACCATTGCTGCCCGGGACGATAGGCGAAACAAAGCACCTTTACATCGAAAGCACCTTTACATCGCAACGTCCATGTCGGGATGC |
| PA14_33700 | pvdF | CTTGCCTTGCACCTGCAGGTCTGCCGGATA | TTTCGTCCGCGAGTGACCAGAGCACCTTTACATCGAAAGCACCTTTACATCGAACTTGCCTTGCACCTGCAGGTCTGCCGGATAAAAGCACCTTTACATCGAAAGCACCTTTACATCGCAACGTCCATGTCGGGATGC |
| PA14_33700 | pvdF | ACCAGGAGCCCGTCCAATACCACCACATCG | TTTCGTCCGCGAGTGACCAGAGCACCTTTACATCGAAAGCACCTTTACATCGAAACCAGGAGCCCGTCCAATACCACCACATCGAAAGCACCTTTACATCGAAAGCACCTTTACATCGCAACGTCCATGTCGGGATGC |
| PA14_33700 | pvdF | TGGACGGCACGCTGAGCAAGAGGTCGTTCA | TTTCGTCCGCGAGTGACCAGAGCACCTTTACATCGAAAGCACCTTTACATCGAATGGACGGCACGCTGAGCAAGAGGTCGTTCAAAAGCACCTTTACATCGAAAGCACCTTTACATCGCAACGTCCATGTCGGGATGC |
| PA14_33700 | pvdF | TCCATACAACGCGTCCAGGGTCGCATAGGC | TTTCGTCCGCGAGTGACCAGAGCACCTTTACATCGAAAGCACCTTTACATCGAATCCATACAACGCGTCCAGGGTCGCATAGGCAAAGCACCTTTACATCGAAAGCACCTTTACATCGCAACGTCCATGTCGGGATGC |
| PA14_33700 | pvdF | TCCATGGTCGCCCAATCCACCACCTTCTCG | TTTCGTCCGCGAGTGACCAGAGCACCTTTACATCGAAAGCACCTTTACATCGAATCCATGGTCGCCCAATCCACCACCTTCTCGAAAGCACCTTTACATCGAAAGCACCTTTACATCGCAACGTCCATGTCGGGATGC |
| PA14_33700 | pvdF | CTCCACCAAGGACTCCAGTACCGACTTCAT | TTTCGTCCGCGAGTGACCAGAGCACCTTTACATCGAAAGCACCTTTACATCGAACTCCACCAAGGACTCCAGTACCGACTTCATAAAGCACCTTTACATCGAAAGCACCTTTACATCGCAACGTCCATGTCGGGATGC |
| PA14_33700 | pvdF | TATTCATGATCCGCCGTGCGAACGGAGCGC | TTTCGTCCGCGAGTGACCAGAGCACCTTTACATCGAAAGCACCTTTACATCGAATATTCATGATCCGCCGTGCGAACGGAGCGCAAAGCACCTTTACATCGAAAGCACCTTTACATCGCAACGTCCATGTCGGGATGC |
| PA14_23920 | purF | ATTGGGGTCGCGGAAACCGACGATGCCATG | TTTCGTCCGCGAGTGACCAGCCGGTTAGCAATGCTAACCGGTTAGCAATGCTAAATTGGGGTCGCGGAAACCGACGATGCCATGAACCGGTTAGCAATGCTAACCGGTTAGCAATGCTCAACGTCCATGTCGGGATGC |
| PA14_23920 | purF | GCGATCATGTATTCGACGCCGTTCTCGGTA | TTTCGTCCGCGAGTGACCAGCCGGTTAGCAATGCTAACCGGTTAGCAATGCTAAGCGATCATGTATTCGACGCCGTTCTCGGTAAACCGGTTAGCAATGCTAACCGGTTAGCAATGCTCAACGTCCATGTCGGGATGC |
| PA14_23920 | purF | GATCGGGATGACCACGTCGATGTCGTGGTC | TTTCGTCCGCGAGTGACCAGCCGGTTAGCAATGCTAACCGGTTAGCAATGCTAAGATCGGGATGACCACGTCGATGTCGTGGTCAACCGGTTAGCAATGCTAACCGGTTAGCAATGCTCAACGTCCATGTCGGGATGC |
| PA14_23920 | purF | ACGTTCTTGCCGCGGAACTCCAGCTCGATG | TTTCGTCCGCGAGTGACCAGCCGGTTAGCAATGCTAACCGGTTAGCAATGCTAAACGTTCTTGCCGCGGAACTCCAGCTCGATGAACCGGTTAGCAATGCTAACCGGTTAGCAATGCTCAACGTCCATGTCGGGATGC |
| PA14_23920 | purF | AGATGCCGTCCATGATCGAATCCGGACGCG | TTTCGTCCGCGAGTGACCAGCCGGTTAGCAATGCTAACCGGTTAGCAATGCTAAAGATGCCGTCCATGATCGAATCCGGACGCGAACCGGTTAGCAATGCTAACCGGTTAGCAATGCTCAACGTCCATGTCGGGATGC |
| PA14_23920 | purF | ACGTGCTCGAAGATGCACGGCGCGTACTTC | TTTCGTCCGCGAGTGACCAGCCGGTTAGCAATGCTAACCGGTTAGCAATGCTAAACGTGCTCGAAGATGCACGGCGCGTACTTCAACCGGTTAGCAATGCTAACCGGTTAGCAATGCTCAACGTCCATGTCGGGATGC |
| PA14_23920 | purF | TTGCTGACGTCCTCGGTGCTGCGATTGTGC | TTTCGTCCGCGAGTGACCAGCCGGTTAGCAATGCTAACCGGTTAGCAATGCTAATTGCTGACGTCCTCGGTGCTGCGATTGTGCAACCGGTTAGCAATGCTAACCGGTTAGCAATGCTCAACGTCCATGTCGGGATGC |
| PA14_23920 | purF | ATCGGAGTTGGTGTTCACGTGGCGCAGGTC | TTTCGTCCGCGAGTGACCAGCCGGTTAGCAATGCTAACCGGTTAGCAATGCTAAATCGGAGTTGGTGTTCACGTGGCGCAGGTCAACCGGTTAGCAATGCTAACCGGTTAGCAATGCTCAACGTCCATGTCGGGATGC |
| PA14_23920 | purF | ATCGTCATGGCAGGTGACGATACCGGCAGC | TTTCGTCCGCGAGTGACCAGCCGGTTAGCAATGCTAACCGGTTAGCAATGCTAAATCGTCATGGCAGGTGACGATACCGGCAGCAACCGGTTAGCAATGCTAACCGGTTAGCAATGCTCAACGTCCATGTCGGGATGC |
| PA14_23920 | purF | AGCGCGTCATACAGCGCCTGATTGACGTTC | TTTCGTCCGCGAGTGACCAGCCGGTTAGCAATGCTAACCGGTTAGCAATGCTAAAGCGCGTCATACAGCGCCTGATTGACGTTCAACCGGTTAGCAATGCTAACCGGTTAGCAATGCTCAACGTCCATGTCGGGATGC |
| PA14_23920 | purF | TTGCCTTCCTCGGTGATGTACACCGCCTCG | TTTCGTCCGCGAGTGACCAGCCGGTTAGCAATGCTAACCGGTTAGCAATGCTAATTGCCTTCCTCGGTGATGTACACCGCCTCGAACCGGTTAGCAATGCTAACCGGTTAGCAATGCTCAACGTCCATGTCGGGATGC |
| PA14_23920 | purF | TCGTGGGCGAACACGTTGAGCAGCACTTCC | TTTCGTCCGCGAGTGACCAGCCGGTTAGCAATGCTAACCGGTTAGCAATGCTAATCGTGGGCGAACACGTTGAGCAGCACTTCCAACCGGTTAGCAATGCTAACCGGTTAGCAATGCTCAACGTCCATGTCGGGATGC |
| PA14_23920 | purF | CGATGTAGCGGTTCTTGACGAAGCCCTCGC | TTTCGTCCGCGAGTGACCAGCCGGTTAGCAATGCTAACCGGTTAGCAATGCTAACGATGTAGCGGTTCTTGACGAAGCCCTCGCAACCGGTTAGCAATGCTAACCGGTTAGCAATGCTCAACGTCCATGTCGGGATGC |
| PA14_23920 | purF | TTGAGCTTCTGCCGCACGGACTTCTTCCGC | TTTCGTCCGCGAGTGACCAGCCGGTTAGCAATGCTAACCGGTTAGCAATGCTAATTGAGCTTCTGCCGCACGGACTTCTTCCGCAACCGGTTAGCAATGCTAACCGGTTAGCAATGCTCAACGTCCATGTCGGGATGC |
| PA14_23920 | purF | TGCCGTAGGGCGAGTTGACGTAGAACGGCT | TTTCGTCCGCGAGTGACCAGCCGGTTAGCAATGCTAACCGGTTAGCAATGCTAATGCCGTAGGGCGAGTTGACGTAGAACGGCTAACCGGTTAGCAATGCTAACCGGTTAGCAATGCTCAACGTCCATGTCGGGATGC |
| PA14_23920 | purF | ATGTCCTCCTCGGTGGGCTGCAGCTTGTTG | TTTCGTCCGCGAGTGACCAGCCGGTTAGCAATGCTAACCGGTTAGCAATGCTAAATGTCCTCCTCGGTGGGCTGCAGCTTGTTGAACCGGTTAGCAATGCTAACCGGTTAGCAATGCTCAACGTCCATGTCGGGATGC |
| PA14_23920 | purF | TCTTGTCGGCAAGCTTCTCGCCCATGCGCA | TTTCGTCCGCGAGTGACCAGCCGGTTAGCAATGCTAACCGGTTAGCAATGCTAATCTTGTCGGCAAGCTTCTCGCCCATGCGCAAACCGGTTAGCAATGCTAACCGGTTAGCAATGCTCAACGTCCATGTCGGGATGC |
| PA14_23920 | purF | CGTAGATTTCCTTGGCCAACTGCTCGACAT | TTTCGTCCGCGAGTGACCAGCCGGTTAGCAATGCTAACCGGTTAGCAATGCTAACGTAGATTTCCTTGGCCAACTGCTCGACATAACCGGTTAGCAATGCTAACCGGTTAGCAATGCTCAACGTCCATGTCGGGATGC |
| PA14_23920 | purF | GTTGTACAGATCGATGATCGCGCTGACCGC | TTTCGTCCGCGAGTGACCAGCCGGTTAGCAATGCTAACCGGTTAGCAATGCTAAGTTGTACAGATCGATGATCGCGCTGACCGCAACCGGTTAGCAATGCTAACCGGTTAGCAATGCTCAACGTCCATGTCGGGATGC |
| PA14_23920 | purF | CTTGCCCACGATACCGACGATGCCACACAT | TTTCGTCCGCGAGTGACCAGCCGGTTAGCAATGCTAACCGGTTAGCAATGCTAACTTGCCCACGATACCGACGATGCCACACATAACCGGTTAGCAATGCTAACCGGTTAGCAATGCTCAACGTCCATGTCGGGATGC |
| PA14_17290 | pyrG | ACCACCCGTGACGAAGATGTAGCGCGTCAT | TTTCGTCCGCGAGTGACCAGCCACGATCGCTGATTAACCACGATCGCTGATTAAACCACCCGTGACGAAGATGTAGCGCGTCATAACCACGATCGCTGATTAACCACGATCGCTGATTCAACGTCCATGTCGGGATGC |
| PA14_17290 | pyrG | CATCACGCGGTGTGGAGGTGAATTCCGGGT | TTTCGTCCGCGAGTGACCAGCCACGATCGCTGATTAACCACGATCGCTGATTAACATCACGCGGTGTGGAGGTGAATTCCGGGTAACCACGATCGCTGATTAACCACGATCGCTGATTCAACGTCCATGTCGGGATGC |
| PA14_17290 | pyrG | AACGAAGCGCTCGTAATGGCCCAGGTCCAG | TTTCGTCCGCGAGTGACCAGCCACGATCGCTGATTAACCACGATCGCTGATTAAAACGAAGCGCTCGTAATGGCCCAGGTCCAGAACCACGATCGCTGATTAACCACGATCGCTGATTCAACGTCCATGTCGGGATGC |
| PA14_17290 | pyrG | CCACGCGCTTCCAAGATCGCAGCCAGAGAA | TTTCGTCCGCGAGTGACCAGCCACGATCGCTGATTAACCACGATCGCTGATTAACCACGCGCTTCCAAGATCGCAGCCAGAGAAAACCACGATCGCTGATTAACCACGATCGCTGATTCAACGTCCATGTCGGGATGC |
| PA14_17290 | pyrG | TCCTTGCGCAACACGTCCATGTAGACGCGG | TTTCGTCCGCGAGTGACCAGCCACGATCGCTGATTAACCACGATCGCTGATTAATCCTTGCGCAACACGTCCATGTAGACGCGGAACCACGATCGCTGATTAACCACGATCGCTGATTCAACGTCCATGTCGGGATGC |
| PA14_17290 | pyrG | CTTGATGATGCGCCGCTTGATCTCGTCGGT | TTTCGTCCGCGAGTGACCAGCCACGATCGCTGATTAACCACGATCGCTGATTAACTTGATGATGCGCCGCTTGATCTCGTCGGTAACCACGATCGCTGATTAACCACGATCGCTGATTCAACGTCCATGTCGGGATGC |
| PA14_17290 | pyrG | AAGACCTCACCGTGCTGGAACGGGCTCATG | TTTCGTCCGCGAGTGACCAGCCACGATCGCTGATTAACCACGATCGCTGATTAAAAGACCTCACCGTGCTGGAACGGGCTCATGAACCACGATCGCTGATTAACCACGATCGCTGATTCAACGTCCATGTCGGGATGC |
| PA14_17290 | pyrG | CGAAGCGATGCCTTTCCCCAATGAAGAAAC | TTTCGTCCGCGAGTGACCAGCCACGATCGCTGATTAACCACGATCGCTGATTAACGAAGCGATGCCTTTCCCCAATGAAGAAACAACCACGATCGCTGATTAACCACGATCGCTGATTCAACGTCCATGTCGGGATGC |
| PA14_17290 | pyrG | ATGCCCAGGTAGGGGATCTTGTTCTCGCGG | TTTCGTCCGCGAGTGACCAGCCACGATCGCTGATTAACCACGATCGCTGATTAAATGCCCAGGTAGGGGATCTTGTTCTCGCGGAACCACGATCGCTGATTAACCACGATCGCTGATTCAACGTCCATGTCGGGATGC |

| | | | |
|---|---|---|---|
| PA14_17290 | pyrG | TGTTGTTCACTTCGTAGCGATGGCGATGGC | TTTCGTCCGCGAGTGACCAGCCACGATCGCTGATTAACCACGATCGCTGATTAATGTTGTTCACTTCGTAGCGATGGCGATGGCAACCACGATCGCTGATTAACCACGATCGCTGATTCAACGTCCATGTCGGGATGC |
| PA14_17290 | pyrG | AACTGGCAGGCAACGAACCACGGATGCTCC | TTTCGTCCGCGAGTGACCAGCCACGATCGCTGATTAACCACGATCGCTGATTAAAACTGGCAGGCAACGAACCACGGATGCTCCAACCACGATCGCTGATTAACCACGATCGCTGATTCAACGTCCATGTCGGGATGC |
| PA14_17290 | pyrG | AAGGGTCCAGCTTGAGCATCGTGATCTTCA | TTTCGTCCGCGAGTGACCAGCCACGATCGCTGATTAACCACGATCGCTGATTAAAAGGGTCCAGCTTGAGCATCGTGATCTTCAAACCACGATCGCTGATTAACCACGATCGCTGATTCAACGTCCATGTCGGGATGC |
| PA14_17290 | pyrG | AGCCGAGCACGTTGCGGGCGTATTCGATGA | TTTCGTCCGCGAGTGACCAGCCACGATCGCTGATTAACCACGATCGCTGATTAAAGCCGAGCACGTTGCGGGCGTATTCGATGAAACCACGATCGCTGATTAACCACGATCGCTGATTCAACGTCCATGTCGGGATGC |
| PA14_17290 | pyrG | GGTGAAGTTGTTGTTCTGGGTCATGGTGGT | TTTCGTCCGCGAGTGACCAGCCACGATCGCTGATTAACCACGATCGCTGATTAAGGTGAAGTTGTTGTTCTGGGTCATGGTGGTAACCACGATCGCTGATTAACCACGATCGCTGATTCAACGTCCATGTCGGGATGC |
| PA14_17290 | pyrG | TACGGGACCAGGGTCAGGTGCATCAGCATC | TTTCGTCCGCGAGTGACCAGCCACGATCGCTGATTAACCACGATCGCTGATTAATACGGGACCAGGGTCAGGTGCATCAGCATCAACCACGATCGCTGATTAACCACGATCGCTGATTCAACGTCCATGTCGGGATGC |
| PA14_39330 | rbsA | ACCTCGAACGACACCTCATGCACCTTGCCG | TTTCGTCCGCGAGTGACCAGTGGCCCATACTCGTTAATGGCCCATACTCGTTAAACCTCGAACGACACCTCATGCACCTTGCCGAATGGCCCATACTCGTTAATGGCCCATACTCGTTCAACGTCCATGTCGGGATGC |
| PA14_39330 | rbsA | ATGCGCTGCGAAATCCGCGCCAGTTCCTCC | TTTCGTCCGCGAGTGACCAGTGGCCCATACTCGTTAATGGCCCATACTCGTTAAATGCGCTGCGAAATCCGCGCCAGTTCCTCCAATGGCCCATACTCGTTAATGGCCCATACTCGTTCAACGTCCATGTCGGGATGC |
| PA14_39330 | rbsA | TCGATGGGAGATGTAGACGATCGCCACGCC | TTTCGTCCGCGAGTGACCAGTGGCCCATACTCGTTAATGGCCCATACTCGTTAATCGATGGGAGATGTAGACGATCGCCACGCCAATGGCCCATACTCGTTAATGGCCCATACTCGTTCAACGTCCATGTCGGGATGC |
| PA14_39330 | rbsA | ATAGATGTCGAACTTGGCGCCGACGTCGAT | TTTCGTCCGCGAGTGACCAGTGGCCCATACTCGTTAATGGCCCATACTCGTTAAATAGATGTCGAACTTGGCGCCGACGTCGATAATGGCCCATACTCGTTAATGGCCCATACTCGTTCAACGTCCATGTCGGGATGC |
| PA14_39330 | rbsA | ATCAGGGTCACCAGTTGGTCGGCGTCGTAG | TTTCGTCCGCGAGTGACCAGTGGCCCATACTCGTTAATGGCCCATACTCGTTAAATCAGGGTCACCAGTTGGTCGGCGTCGTAGAATGGCCCATACTCGTTAATGGCCCATACTCGTTCAACGTCCATGTCGGGATGC |
| PA14_39330 | rbsA | AGCGCGTCGCGTTTCTTGTAACCGGCGAAG | TTTCGTCCGCGAGTGACCAGTGGCCCATACTCGTTAATGGCCCATACTCGTTAAAGCGCGTCGCGTTTCTTGTAACCGGCGAAGAATGGCCCATACTCGTTAATGGCCCATACTCGTTCAACGTCCATGTCGGGATGC |
| PA14_39330 | rbsA | TTGGCGCTGATCGACTGGTCCAGCAGCAGG | TTTCGTCCGCGAGTGACCAGTGGCCCATACTCGTTAATGGCCCATACTCGTTAATTGGCGCTGATCGACTGGTCCAGCAGCAGGAATGGCCCATACTCGTTAATGGCCCATACTCGTTCAACGTCCATGTCGGGATGC |
| PA14_39330 | rbsA | CGCCGACGATCTTCGACAGGGTGCTCTTGC | TTTCGTCCGCGAGTGACCAGTGGCCCATACTCGTTAATGGCCCATACTCGTTAACGCCGACGATCTTCGACAGGGTGCTCTTGCAATGGCCCATACTCGTTAATGGCCCATACTCGTTCAACGTCCATGTCGGGATGC |
| PA14_39330 | rbsA | GATCAGTCCGGAAATGCCGAAGATCTCCCC | TTTCGTCCGCGAGTGACCAGTGGCCCATACTCGTTAATGGCCCATACTCGTTAAGATCAGTCCGGAAATGCCGAAGATCTCCCCAATGGCCCATACTCGTTAATGGCCCATACTCGTTCAACGTCCATGTCGGGATGC |
| PA14_39330 | rbsA | TAGATCAGGCGCAACAATTCGGTGCGCCCG | TTTCGTCCGCGAGTGACCAGTGGCCCATACTCGTTAATGGCCCATACTCGTTAATAGATCAGGCGCAACAATTCGGTGCGCCCGAATGGCCCATACTCGTTAATGGCCCATACTCGTTCAACGTCCATGTCGGGATGC |
| PA14_39330 | rbsA | GATCAGCAAGCGGCAGTCGCCGATCAGGTT | TTTCGTCCGCGAGTGACCAGTGGCCCATACTCGTTAATGGCCCATACTCGTTAAGATCAGCAAGCGGCAGTCGCCGATCAGGTTAATGGCCCATACTCGTTAATGGCCCATACTCGTTCAACGTCCATGTCGGGATGC |
| PA14_39330 | rbsA | TCAGTTCCTGCATCACCATGCGCACGCCCA | TTTCGTCCGCGAGTGACCAGTGGCCCATACTCGTTAATGGCCCATACTCGTTAATCAGTTCCTGCATCACCATGCGCACGCCCAAATGGCCCATACTCGTTAATGGCCCATACTCGTTCAACGTCCATGTCGGGATGC |
| PA14_39330 | rbsA | AGGAACAGGTTCTCCGCCACCGACAGGGTC | TTTCGTCCGCGAGTGACCAGTGGCCCATACTCGTTAATGGCCCATACTCGTTAAAGGAACAGGTTCTCCGCCACCGACAGGGTCAATGGCCCATACTCGTTAATGGCCCATACTCGTTCAACGTCCATGTCGGGATGC |
| PA14_39330 | rbsA | AGCATGTCCACTTCGCGCGCGGTAAGCATC | TTTCGTCCGCGAGTGACCAGTGGCCCATACTCGTTAATGGCCCATACTCGTTAAAGCATGTCCACTTCGCGCGCGGTAAGCATCAATGGCCCATACTCGTTAATGGCCCATACTCGTTCAACGTCCATGTCGGGATGC |
| PA14_39330 | rbsA | ATCGCAGATCAGCATCAGCTCGCGCAGGTC | TTTCGTCCGCGAGTGACCAGTGGCCCATACTCGTTAATGGCCCATACTCGTTAAATCGCAGATCAGCATCAGCTCGCGCAGGTCAATGGCCCATACTCGTTAATGGCCCATACTCGTTCAACGTCCATGTCGGGATGC |
| PA14_17530 | recA | TGCCGAATTGGCGTTCGATCTGTCCCAGGG | TTTCGTCCGCGAGTGACCAGCTTCAGCGATAGGCAAACTTCAGCGATAGGCAAATGCCGAATTGGCGTTCGATCTGTCCCAGGGAACTTCAGCGATAGGCAAACTTCAGCGATAGGCACAACGTCCATGTCGGGATGC |
| PA14_17530 | recA | AGCATGTCGGTGATTTCCAGGGCCTGCTCG | TTTCGTCCGCGAGTGACCAGCTTCAGCGATAGGCAAACTTCAGCGATAGGCAAAAGCATGTCGGTGATTTCCAGGGCCTGCTCGAACTTCAGCGATAGGCAAACTTCAGCGATAGGCACAACGTCCATGTCGGGATGC |
| PA14_17530 | recA | GGGGAAACCTTGTTCTTCACCACCTTGACG | TTTCGTCCGCGAGTGACCAGCTTCAGCGATAGGCAAACTTCAGCGATAGGCAAAGGGGAAACCTTGTTCTTCACCACCTTGACGAACTTCAGCGATAGGCAAACTTCAGCGATAGGCACAACGTCCATGTCGGGATGC |
| PA14_17530 | recA | GATGGTCTTCTCCAGTACCGAACCGATTTC | TTTCGTCCGCGAGTGACCAGCTTCAGCGATAGGCAAACTTCAGCGATAGGCAAAGATGGTCTTCTCCAGTACCGAACCGATTTCAACTTCAGCGATAGGCAAACTTCAGCGATAGGCACAACGTCCATGTCGGGATGC |
| PA14_17530 | recA | AGCAGGTCGTCGACGTTGACGCCCAGCTTG | TTTCGTCCGCGAGTGACCAGCTTCAGCGATAGGCAAACTTCAGCGATAGGCAAAAGCAGGTCGTCGACGTTGACGCCCAGCTTGAACTTCAGCGATAGGCAAACTTCAGCGATAGGCACAACGTCCATGTCGGGATGC |
| PA14_17530 | recA | TTTCGCTACCCACCACCTCGTCGCCTTCCT | TTTCGTCCGCGAGTGACCAGCTTCAGCGATAGGCAAACTTCAGCGATAGGCAAATTTCGCTACCCACCACCTCGTCGCCTTCCTAACTTCAGCGATAGGCAAACTTCAGCGATAGGCACAACGTCCATGTCGGGATGC |
| PA14_17530 | recA | ATCACCGAGAGGGTCAGGGTGGTCTTGCCC | TTTCGTCCGCGAGTGACCAGCTTCAGCGATAGGCAAACTTCAGCGATAGGCAAAATCACCGAGAGGGTCAGGGTGGTCTTGCCCAACTTCAGCGATAGGCAAACTTCAGCGATAGGCACAACGTCCATGTCGGGATGC |
| PA14_17530 | recA | TTCAGTGCGTTACCGCCGGTGGTGGTTTCC | TTTCGTCCGCGAGTGACCAGCTTCAGCGATAGGCAAACTTCAGCGATAGGCAAATTCAGTGCGTTACCGCCGGTGGTGGTTTCCAACTTCAGCGATAGGCAAACTTCAGCGATAGGCACAACGTCCATGTCGGGATGC |
| PA14_17530 | recA | ATTCCGGACCGTAGATCTCGACGATCCGGC | TTTCGTCCGCGAGTGACCAGCTTCAGCGATAGGCAAACTTCAGCGATAGGCAAAATTCCGGACCGTAGATCTCGACGATCCGGCAACTTCAGCGATAGGCAAACTTCAGCGATAGGCACAACGTCCATGTCGGGATGC |
| PA14_17530 | recA | TGATCTCGCCGGTACGGTAGATGCCCTTGC | TTTCGTCCGCGAGTGACCAGCTTCAGCGATAGGCAAACTTCAGCGATAGGCAAATGATCTCGCCGGTACGGTAGATGCCCTTGCAACTTCAGCGATAGGCAAACTTCAGCGATAGGCACAACGTCCATGTCGGGATGC |
| PA14_17530 | recA | AATCGGCTTCGGCGTCAGCCACTTCTTCGG | TTTCGTCCGCGAGTGACCAGCTTCAGCGATAGGCAAACTTCAGCGATAGGCAAAAATCGGCTTCGGCGTCAGCCACTTCTTCGGAACTTCAGCGATAGGCAAACTTCAGCGATAGGCACAACGTCCATGTCGGGATGC |
| PA14_17530 | recA | ATCTTGCTGCCCTGGTAGCTGTACCAGGCG | TTTCGTCCGCGAGTGACCAGCTTCAGCGATAGGCAAACTTCAGCGATAGGCAAAATCTTGCTGCCCTGGTAGCTGTACCAGGCGAACTTCAGCGATAGGCAAACTTCAGCGATAGGCACAACGTCCATGTCGGGATGC |
| PA14_17530 | recA | TGATGAAGATGACCAGGCAGTTGGCGTTCT | TTTCGTCCGCGAGTGACCAGCTTCAGCGATAGGCAAACTTCAGCGATAGGCAAATGATGAAGATGACCAGGCAGTTGGCGTTCTAACTTCAGCGATAGGCAAACTTCAGCGATAGGCACAACGTCCATGTCGGGATGC |
| PA14_17530 | recA | ATATTGCCGGTGATCTTGCGCAGCGCCTGG | TTTCGTCCGCGAGTGACCAGCTTCAGCGATAGGCAAACTTCAGCGATAGGCAAAATATTGCCGGTGATCTTGCGCAGCGCCTGGAACTTCAGCGATAGGCAAACTTCAGCGATAGGCACAACGTCCATGTCGGGATGC |
| PA14_17530 | recA | ACTTCTCGACCAGGCCCAATTGCACGCCCA | TTTCGTCCGCGAGTGACCAGCTTCAGCGATAGGCAAACTTCAGCGATAGGCAAAACTTCTCGACCAGGCCCAATTGCACGCCCAAACTTCAGCGATAGGCAAACTTCAGCGATAGGCACAACGTCCATGTCGGGATGC |
| PA14_17530 | recA | TAGAGGATCTGGAACTCGGCCTGGCGGAAC | TTTCGTCCGCGAGTGACCAGCTTCAGCGATAGGCAAACTTCAGCGATAGGCAAATAGAGGATCTGGAACTCGGCCTGGCGGAACAACTTCAGCGATAGGCAAACTTCAGCGATAGGCACAACGTCCATGTCGGGATGC |
| PA14_17530 | recA | TCTTCCAGGTACTTGGCGGCGTTCGCCTTG | TTTCGTCCGCGAGTGACCAGCTTCAGCGATAGGCAAACTTCAGCGATAGGCAAATCTTCCAGGTACTTGGCGGCGTTCGCCTTGAACTTCAGCGATAGGCAAACTTCAGCGATAGGCACAACGTCCATGTCGGGATGC |
| PA14_17530 | recA | GAACATGACGCCGATCTTCATGCGGATCTG | TTTCGTCCGCGAGTGACCAGCTTCAGCGATAGGCAAACTTCAGCGATAGGCAAAGAACATGACGCCGATCTTCATGCGGATCTGAACTTCAGCGATAGGCAAACTTCAGCGATAGGCACAACGTCCATGTCGGGATGC |
| PA14_17530 | recA | TCGACGATGATCACGTCGACCGCGTTGGAG | TTTCGTCCGCGAGTGACCAGCTTCAGCGATAGGCAAACTTCAGCGATAGGCAAATCGACGATGATCACGTCGACCGCGTTGGAGAACTTCAGCGATAGGCAAACTTCAGCGATAGGCACAACGTCCATGTCGGGATGC |
| PA14_17530 | recA | GGCCAGGGCGCGCTTCTTGTTCTCGTCCAT | TTTCGTCCGCGAGTGACCAGCTTCAGCGATAGGCAAACTTCAGCGATAGGCAAAGGCCAGGGCGCGCTTCTTGTTCTCGTCCATAACTTCAGCGATAGGCAAACTTCAGCGATAGGCACAACGTCCATGTCGGGATGC |
| PA14_19130 | rhlI | AAGACTTGATGCCGGTAGCGTCCCAGCTCG | TTTCGTCCGCGAGTGACCAGGTAAGCTATAAGCGGAAGTAAGCTATAAGCGGAAAAGACTTGATGCCGGTAGCGTCCCAGCTCGAAGTAAGCTATAAGCGGAAGTAAGCTATAAGCGGCAACGTCCATGTCGGGATGC |
| PA14_19130 | rhlI | CCTTCCAGCGATTCAGAGAGCAATTCGATC | TTTCGTCCGCGAGTGACCAGGTAAGCTATAAGCGGAAGTAAGCTATAAGCGGAACCTTCCAGCGATTCAGAGAGCAATTCGATCAAGTAAGCTATAAGCGGAAGTAAGCTATAAGCGGCAACGTCCATGTCGGGATGC |
| PA14_19130 | rhlI | TTCCGGGTGGTAGCGCAACAGCATCTCCAG | TTTCGTCCGCGAGTGACCAGGTAAGCTATAAGCGGAAGTAAGCTATAAGCGGAATTCCGGGTGGTAGCGCAACAGCATCTCCAGAAGTAAGCTATAAGCGGAAGTAAGCTATAAGCGGCAACGTCCATGTCGGGATGC |
| PA14_19130 | rhlI | CAGGCTGGACCAGAATATCTTCATCGCCAG | TTTCGTCCGCGAGTGACCAGGTAAGCTATAAGCGGAAGTAAGCTATAAGCGGAACAGGCTGGACCAGAATATCTTCATCGCCAGAAGTAAGCTATAAGCGGAAGTAAGCTATAAGCGGCAACGTCCATGTCGGGATGC |
| PA14_19130 | rhlI | CTGCACAGGTAGGCGAAGACGTCCTTGAGC | TTTCGTCCGCGAGTGACCAGGTAAGCTATAAGCGGAAGTAAGCTATAAGCGGAACTGCACAGGTAGGCGAAGACGTCCTTGAGCAAGTAAGCTATAAGCGGAAGTAAGCTATAAGCGGCAACGTCCATGTCGGGATGC |
| PA14_19130 | rhlI | ACACCGCCATCGACAGCGGTACACCCTGCA | TTTCGTCCGCGAGTGACCAGGTAAGCTATAAGCGGAAGTAAGCTATAAGCGGAAACACCGCCATCGACAGCGGTACACCCTGCAAAGTAAGCTATAAGCGGAAGTAAGCTATAAGCGGCAACGTCCATGTCGGGATGC |
| PA14_19130 | rhlI | AGCTGATCGCGACCAGCGTCTCGCCCTTGA | TTTCGTCCGCGAGTGACCAGGTAAGCTATAAGCGGAAGTAAGCTATAAGCGGAAAGCTGATCGCGACCAGCGTCTCGCCCTTGAAAGTAAGCTATAAGCGGAAGTAAGCTATAAGCGGCAACGTCCATGTCGGGATGC |
| PA14_19130 | rhlI | GAAATAGCGCTCCATGGCCGTGGTGGTCAC | TTTCGTCCGCGAGTGACCAGGTAAGCTATAAGCGGAAGTAAGCTATAAGCGGAAGAAATAGCGCTCCATGGCCGTGGTGGTCACAAGTAAGCTATAAGCGGAAGTAAGCTATAAGCGGCAACGTCCATGTCGGGATGC |
| PA14_19130 | rhlI | AACTGGTCGAATTCCTGGTCGCGGACCCTG | TTTCGTCCGCGAGTGACCAGGTAAGCTATAAGCGGAAGTAAGCTATAAGCGGAAAACTGGTCGAATTCCTGGTCGCGGACCCTGAAGTAAGCTATAAGCGGAAGTAAGCTATAAGCGGCAACGTCCATGTCGGGATGC |
| PA14_19130 | rhlI | TGGAGACCACGTCCCAGCCCAGCTTCTCGA | TTTCGTCCGCGAGTGACCAGGTAAGCTATAAGCGGAAGTAAGCTATAAGCGGAATGGAGACCACGTCCCAGCCCAGCTTCTCGAAAGTAAGCTATAAGCGGAAGTAAGCTATAAGCGGCAACGTCCATGTCGGGATGC |
| PA14_19130 | rhlI | TAGCGCGAAAGCTCCCAGACCGACGGATCG | TTTCGTCCGCGAGTGACCAGGTAAGCTATAAGCGGAAGTAAGCTATAAGCGGAATAGCGCGAAAGCTCCCAGACCGACGGATCGAAGTAAGCTATAAGCGGAAGTAAGCTATAAGCGGCAACGTCCATGTCGGGATGC |
| PA14_19130 | rhlI | TGGCGACGATGTAGCGGGTTTGCGGATGGT | TTTCGTCCGCGAGTGACCAGGTAAGCTATAAGCGGAAGTAAGCTATAAGCGGAATGGCGACGATGTAGCGGGTTTGCGGATGGTAAGTAAGCTATAAGCGGAAGTAAGCTATAAGCGGCAACGTCCATGTCGGGATGC |
| PA14_19100 | rhlA | CGAATGCCATCACCACCGAGCTGCGGATAC | TTTCGTCCGCGAGTGACCAGATTGCGAGGCCACTAAAATTGCGAGGCCACTAAACGAATGCCATCACCACCGAGCTGCGGATACAAATTGCGAGGCCACTAAAATTGCGAGGCCACTACAACGTCCATGTCGGGATGC |
| PA14_19100 | rhlA | AAATGCACGTGGCTCTGGATCCGCTCCAGG | TTTCGTCCGCGAGTGACCAGATTGCGAGGCCACTAAAATTGCGAGGCCACTAAAAAATGCACGTGGCTCTGGATCCGCTCCAGGAAATTGCGAGGCCACTAAAATTGCGAGGCCACTACAACGTCCATGTCGGGATGC |
| PA14_19100 | rhlA | ATGTGAAAGCGCGCCTGCTCGTATTCGCCG | TTTCGTCCGCGAGTGACCAGATTGCGAGGCCACTAAAATTGCGAGGCCACTAAAATGTGAAAGCGCGCCTGCTCGTATTCGCCGAAATTGCGAGGCCACTAAAATTGCGAGGCCACTACAACGTCCATGTCGGGATGC |
| PA14_19100 | rhlA | AGGAGGATTTCCACCTCGTCGTCCTTGGTG | TTTCGTCCGCGAGTGACCAGATTGCGAGGCCACTAAAATTGCGAGGCCACTAAAAGGAGGATTTCCACCTCGTCGTCCTTGGTGAAATTGCGAGGCCACTAAAATTGCGAGGCCACTACAACGTCCATGTCGGGATGC |
| PA14_19100 | rhlA | AGCTTGGACTCCAGGTCGAGGAAATGCCCG | TTTCGTCCGCGAGTGACCAGATTGCGAGGCCACTAAAATTGCGAGGCCACTAAAAGCTTGGACTCCAGGTCGAGGAAATGCCCGAAATTGCGAGGCCACTAAAATTGCGAGGCCACTACAACGTCCATGTCGGGATGC |
| PA14_19100 | rhlA | TATTTGCCGACGGTCTCGTTGAGCAGATGG | TTTCGTCCGCGAGTGACCAGATTGCGAGGCCACTAAAATTGCGAGGCCACTAAATATTTGCCGACGGTCTCGTTGAGCAGATGGAAATTGCGAGGCCACTAAAATTGCGAGGCCACTACAACGTCCATGTCGGGATGC |
| PA14_19100 | rhlA | AGGCGTAGCCGATGGCCATCTCGTGGAATC | TTTCGTCCGCGAGTGACCAGATTGCGAGGCCACTAAAATTGCGAGGCCACTAAAAGGCGTAGCCGATGGCCATCTCGTGGAATCAAATTGCGAGGCCACTAAAATTGCGAGGCCACTACAACGTCCATGTCGGGATGC |
| PA14_19100 | rhlA | AACCGATACCAACAGACTTTCGCGCCGCAT | TTTCGTCCGCGAGTGACCAGATTGCGAGGCCACTAAAATTGCGAGGCCACTAAAAACCGATACCAACAGACTTTCGCGCCGCATAAATTGCGAGGCCACTAAAATTGCGAGGCCACTACAACGTCCATGTCGGGATGC |
| PA14_19100 | rhlA | GAGACCAGGTGATTGACCTCGAAGCGCTCG | TTTCGTCCGCGAGTGACCAGATTGCGAGGCCACTAAAATTGCGAGGCCACTAAAGAGACCAGGTGATTGACCTCGAAGCGCTCGAAATTGCGAGGCCACTAAAATTGCGAGGCCACTACAACGTCCATGTCGGGATGC |
| PA14_19100 | rhlA | AACGCGCTCGACATGTACCCGCAGGCCCTT | TTTCGTCCGCGAGTGACCAGATTGCGAGGCCACTAAAATTGCGAGGCCACTAAAAACGCGCTCGACATGTACCCGCAGGCCCTTAAATTGCGAGGCCACTAAAATTGCGAGGCCACTACAACGTCCATGTCGGGATGC |
| PA14_19100 | rhlA | AACTGCAGTGCGGCAGGTAGTCGCGGAACT | TTTCGTCCGCGAGTGACCAGATTGCGAGGCCACTAAAATTGCGAGGCCACTAAAAACTGCAGTGCGGCAGGTAGTCGCGGAACTAAATTGCGAGGCCACTAAAATTGCGAGGCCACTACAACGTCCATGTCGGGATGC |
| PA14_19100 | rhlA | GATCGCCGACTTGTCGTCCAGCTCGATCAG | TTTCGTCCGCGAGTGACCAGATTGCGAGGCCACTAAAATTGCGAGGCCACTAAAGATCGCCGACTTGTCGTCCAGCTCGATCAGAAATTGCGAGGCCACTAAAATTGCGAGGCCACTACAACGTCCATGTCGGGATGC |
| PA14_19100 | rhlA | ACCACGTTGAAATGTTCGGCCAGGCACTTG | TTTCGTCCGCGAGTGACCAGATTGCGAGGCCACTAAAATTGCGAGGCCACTAAAACCACGTTGAAATGTTCGGCCAGGCACTTGAAATTGCGAGGCCACTAAAATTGCGAGGCCACTACAACGTCCATGTCGGGATGC |
| PA14_19100 | rhlA | CGACGTAGTCGAGCATCGCCTGGTTCAGTC | TTTCGTCCGCGAGTGACCAGATTGCGAGGCCACTAAAATTGCGAGGCCACTAAACGACGTAGTCGAGCATCGCCTGGTTCAGTCAAATTGCGAGGCCACTAAAATTGCGAGGCCACTACAACGTCCATGTCGGGATGC |
| PA14_19100 | rhlA | TGTGCTGATGGTTGCTGGCTTTCAGGCGCG | TTTCGTCCGCGAGTGACCAGATTGCGAGGCCACTAAAATTGCGAGGCCACTAAATGTGCTGATGGTTGCTGGCTTTCAGGCGCGAAATTGCGAGGCCACTAAAATTGCGAGGCCACTACAACGTCCATGTCGGGATGC |
| PA14_52180 | relA | TGATGTAGCCGACGATGGGATCTCCCGGCA | TTTCGTCCGCGAGTGACCAGACTTCAACGAGTGACAAACTTCAACGAGTGACAATGATGTAGCCGACGATGGGATCTCCCGGCAAAACTTCAACGAGTGACAAACTTCAACGAGTGACCAACGTCCATGTCGGGATGC |
| PA14_52180 | relA | AGCACCATGGAATGGCGCGGATTGAGGCTG | TTTCGTCCGCGAGTGACCAGACTTCAACGAGTGACAAACTTCAACGAGTGACAAAGCACCATGGAATGGCGCGGATTGAGGCTGAAACTTCAACGAGTGACAAACTTCAACGAGTGACCAACGTCCATGTCGGGATGC |
| PA14_52180 | relA | TGGCCGCGAGAGCTTCCTTCAACTGGCTCA | TTTCGTCCGCGAGTGACCAGACTTCAACGAGTGACAAACTTCAACGAGTGACAATGGCCGCGAGAGCTTCCTTCAACTGGCTCAAAACTTCAACGAGTGACAAACTTCAACGAGTGACCAACGTCCATGTCGGGATGC |
| PA14_52180 | relA | TTCAGTGTGGACCCGATAGGCGAAATCCAG | TTTCGTCCGCGAGTGACCAGACTTCAACGAGTGACAAACTTCAACGAGTGACAATTCAGTGTGGACCCGATAGGCGAAATCCAGAAACTTCAACGAGTGACAAACTTCAACGAGTGACCAACGTCCATGTCGGGATGC |
| PA14_52180 | relA | TGTTGACCGGCTGGTGCGCTCTCACCTGTA | TTTCGTCCGCGAGTGACCAGACTTCAACGAGTGACAAACTTCAACGAGTGACAATGTTGACCGGCTGGTGCGCTCTCACCTGTAAAACTTCAACGAGTGACAAACTTCAACGAGTGACCAACGTCCATGTCGGGATGC |
| PA14_52180 | relA | AATGTGCCGCCAGAGGGTATGCACGATGCC | TTTCGTCCGCGAGTGACCAGACTTCAACGAGTGACAAACTTCAACGAGTGACAAAATGTGCCGCCAGAGGGTATGCACGATGCCAAACTTCAACGAGTGACAAACTTCAACGAGTGACCAACGTCCATGTCGGGATGC |
| PA14_52180 | relA | ACCAGGGACTCCTGATCCAGCTTGAGGTCC | TTTCGTCCGCGAGTGACCAGACTTCAACGAGTGACAAACTTCAACGAGTGACAAACCAGGGACTCCTGATCCAGCTTGAGGTCCAAACTTCAACGAGTGACAAACTTCAACGAGTGACCAACGTCCATGTCGGGATGC |
| PA14_52180 | relA | CGAAATGCTTGTTGACCGCTTCCAGGGTGA | TTTCGTCCGCGAGTGACCAGACTTCAACGAGTGACAAACTTCAACGAGTGACAACGAAATGCTTGTTGACCGCTTCCAGGGTGAAAACTTCAACGAGTGACAAACTTCAACGAGTGACCAACGTCCATGTCGGGATGC |
| PA14_52180 | relA | TGGACGTGCCTTCTGCCCAGGAGTTCTGTG | TTTCGTCCGCGAGTGACCAGACTTCAACGAGTGACAAACTTCAACGAGTGACAATGGACGTGCCTTCTGCCCAGGAGTTCTGTGAAACTTCAACGAGTGACAAACTTCAACGAGTGACCAACGTCCATGTCGGGATGC |
| PA14_52180 | relA | AACTGACCTGGATCATCCGCTCCGGTTCGC | TTTCGTCCGCGAGTGACCAGACTTCAACGAGTGACAAACTTCAACGAGTGACAAAACTGACCTGGATCATCCGCTCCGGTTCGCAAACTTCAACGAGTGACAAACTTCAACGAGTGACCAACGTCCATGTCGGGATGC |
| PA14_52180 | relA | TGTAGCAGTCGCGCATCTCCGGAACGAGCA | TTTCGTCCGCGAGTGACCAGACTTCAACGAGTGACAAACTTCAACGAGTGACAATGTAGCAGTCGCGCATCTCCGGAACGAGCAAAACTTCAACGAGTGACAAACTTCAACGAGTGACCAACGTCCATGTCGGGATGC |
| PA14_52180 | relA | TAGATGTGTTTCGCCCGCCCGCTGAGATCG | TTTCGTCCGCGAGTGACCAGACTTCAACGAGTGACAAACTTCAACGAGTGACAATAGATGTGTTTCGCCCGCCCGCTGAGATCGAAACTTCAACGAGTGACAAACTTCAACGAGTGACCAACGTCCATGTCGGGATGC |
| PA14_52180 | relA | GTTCGAACGGGTATTCACCGCCAGAACGTT | TTTCGTCCGCGAGTGACCAGACTTCAACGAGTGACAAACTTCAACGAGTGACAAGTTCGAACGGGTATTCACCGCCAGAACGTTAAACTTCAACGAGTGACAAACTTCAACGAGTGACCAACGTCCATGTCGGGATGC |
| PA14_52180 | relA | GTCCAAGAGCGTCCAGACCGGGAATCTCGA | TTTCGTCCGCGAGTGACCAGACTTCAACGAGTGACAAACTTCAACGAGTGACAAGTCCAAGAGCGTCCAGACCGGGAATCTCGAAAACTTCAACGAGTGACAAACTTCAACGAGTGACCAACGTCCATGTCGGGATGC |
| PA14_52180 | relA | TTGCCTTCCCGCACGCCTCGATAGATGACC | TTTCGTCCGCGAGTGACCAGACTTCAACGAGTGACAAACTTCAACGAGTGACAATTGCCTTCCCGCACGCCTCGATAGATGACCAAACTTCAACGAGTGACAAACTTCAACGAGTGACCAACGTCCATGTCGGGATGC |
| PA14_52180 | relA | AATGGTTGGAGCTGGCCTTGACGTCGGTCC | TTTCGTCCGCGAGTGACCAGACTTCAACGAGTGACAAACTTCAACGAGTGACAAAATGGTTGGAGCTGGCCTTGACGTCGGTCCAAACTTCAACGAGTGACAAACTTCAACGAGTGACCAACGTCCATGTCGGGATGC |
| PA14_52180 | relA | AGCTCCGCCTCTTCGTGCATGGAGTGGGTA | TTTCGTCCGCGAGTGACCAGACTTCAACGAGTGACAAACTTCAACGAGTGACAAAGCTCCGCCTCTTCGTGCATGGAGTGGGTAAAACTTCAACGAGTGACAAACTTCAACGAGTGACCAACGTCCATGTCGGGATGC |
| PA14_52180 | relA | TGATGTTGGGCAGTTGCGAAACGCGCGCCA | TTTCGTCCGCGAGTGACCAGACTTCAACGAGTGACAAACTTCAACGAGTGACAATGATGTTGGGCAGTTGCGAAACGCGCGCCAAAACTTCAACGAGTGACAAACTTCAACGAGTGACCAACGTCCATGTCGGGATGC |
| PA14_52180 | relA | TCCGCTCATTGAGCAGCACCTGGGACACGT | TTTCGTCCGCGAGTGACCAGACTTCAACGAGTGACAAACTTCAACGAGTGACAATCCGCTCATTGAGCAGCACCTGGGACACGTAAACTTCAACGAGTGACAAACTTCAACGAGTGACCAACGTCCATGTCGGGATGC |
| PA14_52180 | relA | GTCAGTTGCATGGTCGCGGTGTTGTCTTCC | TTTCGTCCGCGAGTGACCAGACTTCAACGAGTGACAAACTTCAACGAGTGACAAGTCAGTTGCATGGTCGCGGTGTTGTCTTCCAAACTTCAACGAGTGACAAACTTCAACGAGTGACCAACGTCCATGTCGGGATGC |
| PA14_19120 | rhlR | AGTTCACCGTGCTCTCGGAGATGCTCAGGA | TTTCGTCCGCGAGTGACCAGAGTCGTATACGCGTGAAAGTCGTATACGCGTGAAAGTTCACCGTGCTCTCGGAGATGCTCAGGAAAAGTCGTATACGCGTGAAAGTCGTATACGCGTGCAACGTCCATGTCGGGATGC |
| PA14_19120 | rhlR | CAGGTCGGTCAGCTTCTGGGTCAGCAACTC | TTTCGTCCGCGAGTGACCAGAGTCGTATACGCGTGAAAGTCGTATACGCGTGAACAGGTCGGTCAGCTTCTGGGTCAGCAACTCAAAGTCGTATACGCGTGAAAGTCGTATACGCGTGCAACGTCCATGTCGGGATGC |
| PA14_19120 | rhlR | TCATGCAACGCAGCCGCAGGCGGATTTCCT | TTTCGTCCGCGAGTGACCAGAGTCGTATACGCGTGAAAGTCGTATACGCGTGAATCATGCAACGCAGCCGCAGGCGGATTTCCTAAAGTCGTATACGCGTGAAAGTCGTATACGCGTGCAACGTCCATGTCGGGATGC |
| PA14_19120 | rhlR | GTTCTGCATCTGGTATCGCTCCAGCCAGGC | TTTCGTCCGCGAGTGACCAGAGTCGTATACGCGTGAAAGTCGTATACGCGTGAAGTTCTGCATCTGGTATCGCTCCAGCCAGGCAAAGTCGTATACGCGTGAAAGTCGTATACGCGTGCAACGTCCATGTCGGGATGC |
| PA14_19120 | rhlR | TTCCTTTTCCAGGACGGCGAACACGCCCTG | TTTCGTCCGCGAGTGACCAGAGTCGTATACGCGTGAAAGTCGTATACGCGTGAATTCCTTTTCCAGGACGGCGAACACGCCCTGAAAGTCGTATACGCGTGAAAGTCGTATACGCGTGCAACGTCCATGTCGGGATGC |
| PA14_19120 | rhlR | CGCTACGCAAACCGTCCCACCACAGCAAAA | TTTCGTCCGCGAGTGACCAGAGTCGTATACGCGTGAAAGTCGTATACGCGTGAACGCTACGCAAACCGTCCCACCACAGCAAAAAAAGTCGTATACGCGTGAAAGTCGTATACGCGTGCAACGTCCATGTCGGGATGC |
| PA14_19120 | rhlR | TGGGATAGGTGCCATGGACCTCGGTCTTCG | TTTCGTCCGCGAGTGACCAGAGTCGTATACGCGTGAAAGTCGTATACGCGTGAATGGGATAGGTGCCATGGACCTCGGTCTTCGAAAGTCGTATACGCGTGAAAGTCGTATACGCGTGCAACGTCCATGTCGGGATGC |
| PA14_19120 | rhlR | ATTGCAGGATCTCGCGTTCGCGATGGCTCA | TTTCGTCCGCGAGTGACCAGAGTCGTATACGCGTGAAAGTCGTATACGCGTGAAATTGCAGGATCTCGCGTTCGCGATGGCTCAAAAGTCGTATACGCGTGAAAGTCGTATACGCGTGCAACGTCCATGTCGGGATGC |
| PA14_19120 | rhlR | CAGACCGGGTTGGACATCAGCATCGGATGC | TTTCGTCCGCGAGTGACCAGAGTCGTATACGCGTGAAAGTCGTATACGCGTGAACAGACCGGGTTGGACATCAGCATCGGATGCAAAGTCGTATACGCGTGAAAGTCGTATACGCGTGCAACGTCCATGTCGGGATGC |
| PA14_19120 | rhlR | TGTCGCTCCAGACCACCATTTCCGAGGAGC | TTTCGTCCGCGAGTGACCAGAGTCGTATACGCGTGAAAGTCGTATACGCGTGAATGTCGCTCCAGACCACCATTTCCGAGGAGCAAAGTCGTATACGCGTGAAAGTCGTATACGCGTGCAACGTCCATGTCGGGATGC |
| PA14_19120 | rhlR | GAAAGCACGCTGAGCAAATTGTTCGGCGCG | TTTCGTCCGCGAGTGACCAGAGTCGTATACGCGTGAAAGTCGTATACGCGTGAAGAAAGCACGCTGAGCAAATTGTTCGGCGCGAAAGTCGTATACGCGTGAAAGTCGTATACGCGTGCAACGTCCATGTCGGGATGC |
| PA14_19120 | rhlR | TCGAAGCTGGAGATGTTCTGCTGGTCGCGC | TTTCGTCCGCGAGTGACCAGAGTCGTATACGCGTGAAAGTCGTATACGCGTGAATCGAAGCTGGAGATGTTCTGCTGGTCGCGCAAAGTCGTATACGCGTGAAAGTCGTATACGCGTGCAACGTCCATGTCGGGATGC |
| PA14_19120 | rhlR | ATGGCGATTTCCCCGGAACTCTTGCCGTCG | TTTCGTCCGCGAGTGACCAGAGTCGTATACGCGTGAAAGTCGTATACGCGTGAAATGGCGATTTCCCCGGAACTCTTGCCGTCGAAAGTCGTATACGCGTGAAAGTCGTATACGCGTGCAACGTCCATGTCGGGATGC |
| PA14_19120 | rhlR | GTTCCAGAGCATCCGGCTCTGGTCGAACAG | TTTCGTCCGCGAGTGACCAGAGTCGTATACGCGTGAAAGTCGTATACGCGTGAAGTTCCAGAGCATCCGGCTCTGGTCGAACAGAAAGTCGTATACGCGTGAAAGTCGTATACGCGTGCAACGTCCATGTCGGGATGC |
| PA14_19120 | rhlR | ATAGGCGTAGTAATCGAAGCCCAGGCGCCG | TTTCGTCCGCGAGTGACCAGAGTCGTATACGCGTGAAAGTCGTATACGCGTGAAATAGGCGTAGTAATCGAAGCCCAGGCGCCGAAAGTCGTATACGCGTGAAAGTCGTATACGCGTGCAACGTCCATGTCGGGATGC |
| PA14_69190 | rho | TTGCCGGAGCTGGGGATCACGGTGTTGTAG | TTTCGTCCGCGAGTGACCAGCACTTAACTAGACGCAACACTTAACTAGACGCAATTGCCGGAGCTGGGGATCACGGTGTTGTAGAACACTTAACTAGACGCAACACTTAACTAGACGCCAACGTCCATGTCGGGATGC |
| PA14_69190 | rho | TACGTGACGGGTCGGCGGTTCGTCAAAGGT | TTTCGTCCGCGAGTGACCAGCACTTAACTAGACGCAACACTTAACTAGACGCAATACGTGACGGGTCGGCGGTTCGTCAAAGGTAACACTTAACTAGACGCAACACTTAACTAGACGCCAACGTCCATGTCGGGATGC |
| PA14_69190 | rho | GCAATCGGCTTTTGCTTGAGTTCGGTCAGA | TTTCGTCCGCGAGTGACCAGCACTTAACTAGACGCAACACTTAACTAGACGCAAGCAATCGGCTTTTGCTTGAGTTCGGTCAGAAACACTTAACTAGACGCAACACTTAACTAGACGCCAACGTCCATGTCGGGATGC |
| PA14_69190 | rho | GGCCCATGGCATCGGACATTTCCAGCAGTT | TTTCGTCCGCGAGTGACCAGCACTTAACTAGACGCAACACTTAACTAGACGCAAGGCCCATGGCATCGGACATTTCCAGCAGTTAACACTTAACTAGACGCAACACTTAACTAGACGCCAACGTCCATGTCGGGATGC |
| PA14_69190 | rho | GAGAATCTTGTTCTTCGCATTCTCCGGACG | TTTCGTCCGCGAGTGACCAGCACTTAACTAGACGCAACACTTAACTAGACGCAAGAGAATCTTGTTCTTCGCATTCTCCGGACGAACACTTAACTAGACGCAACACTTAACTAGACGCCAACGTCCATGTCGGGATGC |
| PA14_69190 | rho | TCTTGCGATCCAATGGGAGTTCCATGTTGC | TTTCGTCCGCGAGTGACCAGCACTTAACTAGACGCAACACTTAACTAGACGCAATCTTGCGATCCAATGGGAGTTCCATGTTGCAACACTTAACTAGACGCAACACTTAACTAGACGCCAACGTCCATGTCGGGATGC |
| PA14_69190 | rho | TTCGATCGCGGAGATCTCGTCCATCGGGTG | TTTCGTCCGCGAGTGACCAGCACTTAACTAGACGCAACACTTAACTAGACGCAATTCGATCGCGGAGATCTCGTCCATCGGGTGAACACTTAACTAGACGCAACACTTAACTAGACGCCAACGTCCATGTCGGGATGC |
| PA14_69190 | rho | GTCCTGCTTGCGCGAACGGGCCATGTTTTC | TTTCGTCCGCGAGTGACCAGCACTTAACTAGACGCAACACTTAACTAGACGCAAGTCCTGCTTGCGCGAACGGGCCATGTTTTCAACACTTAACTAGACGCAACACTTAACTAGACGCCAACGTCCATGTCGGGATGC |
| PA14_69190 | rho | TGACCACGTCCTTCTTGTGCTCGACCAGGC | TTTCGTCCGCGAGTGACCAGCACTTAACTAGACGCAACACTTAACTAGACGCAATGACCACGTCCTTCTTGTGCTCGACCAGGCAACACTTAACTAGACGCAACACTTAACTAGACGCCAACGTCCATGTCGGGATGC |
| PA14_69190 | rho | TCTTTCGGCGGCCGGATCTTGCCGATGATG | TTTCGTCCGCGAGTGACCAGCACTTAACTAGACGCAACACTTAACTAGACGCAATCTTTCGGCGGCCGGATCTTGCCGATGATGAACACTTAACTAGACGCAACACTTAACTAGACGCCAACGTCCATGTCGGGATGC |
| PA14_69190 | rho | GCCGTCCTGAAGAATCTCCAGCACGCCATC | TTTCGTCCGCGAGTGACCAGCACTTAACTAGACGCAACACTTAACTAGACGCAAGCCGTCCTGAAGAATCTCCAGCACGCCATCAACACTTAACTAGACGCAACACTTAACTAGACGCCAACGTCCATGTCGGGATGC |
| PA14_69190 | rho | GTACCCTTGAATTCCTCGTAGATCACCTCG | TTTCGTCCGCGAGTGACCAGCACTTAACTAGACGCAACACTTAACTAGACGCAAGTACCCTTGAATTCCTCGTAGATCACCTCGAACACTTAACTAGACGCAACACTTAACTAGACGCCAACGTCCATGTCGGGATGC |
| PA14_69190 | rho | AAGAGAACCGCCCTCTTCGATGTTCCGCGC | TTTCGTCCGCGAGTGACCAGCACTTAACTAGACGCAACACTTAACTAGACGCAAAAGAGAACCGCCCTCTTCGATGTTCCGCGCAACACTTAACTAGACGCAACACTTAACTAGACGCCAACGTCCATGTCGGGATGC |
| PA14_69190 | rho | TGGCATTCGGGGTTGTTGCGGGTGATGTTG | TTTCGTCCGCGAGTGACCAGCACTTAACTAGACGCAACACTTAACTAGACGCAATGGCATTCGGGGTTGTTGCGGGTGATGTTGAACACTTAACTAGACGCAACACTTAACTAGACGCCAACGTCCATGTCGGGATGC |
| PA14_69190 | rho | TTGGCCTTCTCGATCACCATCTCGGCGACC | TTTCGTCCGCGAGTGACCAGCACTTAACTAGACGCAACACTTAACTAGACGCAATTGGCCTTCTCGATCACCATCTCGGCGACCAACACTTAACTAGACGCAACACTTAACTAGACGCCAACGTCCATGTCGGGATGC |
| PA14_09115 | rpoA | ACCACGCTCGAGAGGCTCGAGCGTGATCTT | TTTCGTCCGCGAGTGACCAGCACAGACGCATATTCAACACAGACGCATATTCAAACCACGCTCGAGAGGCTCGAGCGTGATCTTAACACAGACGCATATTCAACACAGACGCATATTCCAACGTCCATGTCGGGATGC |
| PA14_09115 | rpoA | CAGCAGGATCTCGATTACATCTTCCTGCAC | TTTCGTCCGCGAGTGACCAGCACAGACGCATATTCAACACAGACGCATATTCAACAGCAGGATCTCGATTACATCTTCCTGCACAACACAGACGCATATTCAACACAGACGCATATTCCAACGTCCATGTCGGGATGC |
| PA14_09115 | rpoA | CGTTGTCTGCCAGGTTGGCGATAACGTGGT | TTTCGTCCGCGAGTGACCAGCACAGACGCATATTCAACACAGACGCATATTCAACGTTGTCTGCCAGGTTGGCGATAACGTGGTAACACAGACGCATATTCAACACAGACGCATATTCCAACGTCCATGTCGGGATGC |
| PA14_09115 | rpoA | TCTTAGCCAGGGTCAGCGTCACTTCATCAC | TTTCGTCCGCGAGTGACCAGCACAGACGCATATTCAACACAGACGCATATTCAATCTTAGCCAGGGTCAGCGTCACTTCATCACAACACAGACGCATATTCAACACAGACGCATATTCCAACGTCCATGTCGGGATGC |
| PA14_09115 | rpoA | ACGAGCTACCTTCAGCTTCATGTTCAGCGC | TTTCGTCCGCGAGTGACCAGCACAGACGCATATTCAACACAGACGCATATTCAAACGAGCTACCTTCAGCTTCATGTTCAGCGCAACACAGACGCATATTCAACACAGACGCATATTCCAACGTCCATGTCGGGATGC |
| PA14_09115 | rpoA | AATATCGGCAGCAGTCACAACACCCGAGCC | TTTCGTCCGCGAGTGACCAGCACAGACGCATATTCAACACAGACGCATATTCAAAATATCGGCAGCAGTCACAACACCCGAGCCAACACAGACGCATATTCAACACAGACGCATATTCCAACGTCCATGTCGGGATGC |
| PA14_09115 | rpoA | ACAGCTCCACTTCGGTGCGCTGGATCAGGT | TTTCGTCCGCGAGTGACCAGCACAGACGCATATTCAACACAGACGCATATTCAAACAGCTCCACTTCGGTGCGCTGGATCAGGTAACACAGACGCATATTCAACACAGACGCATATTCCAACGTCCATGTCGGGATGC |
| PA14_09115 | rpoA | TGTTGCAGGATGGTAGCGGCGCGACGGATA | TTTCGTCCGCGAGTGACCAGCACAGACGCATATTCAACACAGACGCATATTCAATGTTGCAGGATGGTAGCGGCGCGACGGATAAACACAGACGCATATTCAACACAGACGCATATTCCAACGTCCATGTCGGGATGC |
| PA14_09115 | rpoA | TTCAACGACGGGTTCGCTGTCGCCCTTGAG | TTTCGTCCGCGAGTGACCAGCACAGACGCATATTCAACACAGACGCATATTCAATTCAACGACGGGTTCGCTGTCGCCCTTGAGAACACAGACGCATATTCAACACAGACGCATATTCCAACGTCCATGTCGGGATGC |
| PA14_09115 | rpoA | AGGCATGGAGGACAACAGGATGCGACGCAG | TTTCGTCCGCGAGTGACCAGCACAGACGCATATTCAACACAGACGCATATTCAAAGGCATGGAGGACAACAGGATGCGACGCAGAACACAGACGCATATTCAACACAGACGCATATTCCAACGTCCATGTCGGGATGC |
| PA14_09115 | rpoA | TAGGAGACACGACGGACCGGGCTGAACGAT | TTTCGTCCGCGAGTGACCAGCACAGACGCATATTCAACACAGACGCATATTCAATAGGAGACACGACGGACCGGGCTGAACGATAACACAGACGCATATTCAACACAGACGCATATTCCAACGTCCATGTCGGGATGC |

| | | | |
|---|---|---|---|
| PA14_09115 | rpoA | TTGACTGACCACCTGCACATCGATGTGGCG | TTTCGTCCGCGAGTGACCAGCACAGACGCATATTCAACACAGACGCATATTCAATTGACTGACCACCTGCACATCGATGTGGCGAACACAGACGCATATTCAACACAGACGCATATTCCAACGTCCATGTCGGGATGC |
| PA14_09115 | rpoA | TTTGTCCAGGTTGGTGCGCTGCTCGACACG | TTTCGTCCGCGAGTGACCAGCACAGACGCATATTCAACACAGACGCATATTCAATTTGTCCAGGTTGGTGCGCTGCTCGACACGAACACAGACGCATATTCAACACAGACGCATATTCCAACGTCCATGTCGGGATGC |
| PA14_09115 | rpoA | GGGTCAGGAACTCATTTACCGAACTCTGCA | TTTCGTCCGCGAGTGACCAGCACAGACGCATATTCAACACAGACGCATATTCAAGGGTCAGGAACTCATTTACCGAACTCTGCAAACACAGACGCATATTCAACACAGACGCATATTCCAACGTCCATGTCGGGATGC |
| PA14_09115 | rpoA | AGAGTGCCGTTGGTTTCCAGGTCCAGGACC | TTTCGTCCGCGAGTGACCAGCACAGACGCATATTCAACACAGACGCATATTCAAAGAGTGCCGTTGGTTTCCAGGTCCAGGACCAACACAGACGCATATTCAACACAGACGCATATTCCAACGTCCATGTCGGGATGC |
| PA14_09115 | rpoA | CCGTGCAGCTTGATGGCCAGACCTTTCAGG | TTTCGTCCGCGAGTGACCAGCACAGACGCATATTCAACACAGACGCATATTCAACCGTGCAGCTTGATGGCCAGACCTTTCAGGAACACAGACGCATATTCAACACAGACGCATATTCCAACGTCCATGTCGGGATGC |
| PA14_09115 | rpoA | AGGCGCATACCGAGGGACAAACCACGGGAA | TTTCGTCCGCGAGTGACCAGCACAGACGCATATTCAACACAGACGCATATTCAAAGGCGCATACCGAGGGACAAACCACGGGAAAACACAGACGCATATTCAACACAGACGCATATTCCAACGTCCATGTCGGGATGC |
| PA14_09115 | rpoA | CCGTTGATGATCTCAACATCGTGATCCAGC | TTTCGTCCGCGAGTGACCAGCACAGACGCATATTCAACACAGACGCATATTCAACCGTTGATGATCTCAACATCGTGATCCAGCAACACAGACGCATATTCAACACAGACGCATATTCCAACGTCCATGTCGGGATGC |
| PA14_09115 | rpoA | TTCGATCGCCGAGTACTCGTGGAGTACGCC | TTTCGTCCGCGAGTGACCAGCACAGACGCATATTCAACACAGACGCATATTCAATTCGATCGCCGAGTACTCGTGGAGTACGCCAACACAGACGCATATTCAACACAGACGCATATTCCAACGTCCATGTCGGGATGC |
| PA14_09115 | rpoA | TCGTCTTCGTCGCTCTGACGTGCATCGGCA | TTTCGTCCGCGAGTGACCAGCACAGACGCATATTCAACACAGACGCATATTCAATCGTCTTCGTCGCTCTGACGTGCATCGGCAAACACAGACGCATATTCAACACAGACGCATATTCCAACGTCCATGTCGGGATGC |
| PA14_04930 | rpoH | TCTTCGAGGTAATGCGCCGGCGACTGGTAG | TTTCGTCCGCGAGTGACCAGCGCGTCACGTTCAATAACGCGTCACGTTCAATAATCTTCGAGGTAATGCGCCGGCGACTGGTAGAACGCGTCACGTTCAATAACGCGTCACGTTCAATCAACGTCCATGTCGGGATGC |
| PA14_04930 | rpoH | CCGTACCCCGAATAACTCTTGGCGATATGA | TTTCGTCCGCGAGTGACCAGCGCGTCACGTTCAATAACGCGTCACGTTCAATAACCGTACCCCGAATAACTCTTGGCGATATGAAACGCGTCACGTTCAATAACGCGTCACGTTCAATCAACGTCCATGTCGGGATGC |
| PA14_04930 | rpoH | TTGATCCAGTGCACGGCGAAGGACACCAGG | TTTCGTCCGCGAGTGACCAGCGCGTCACGTTCAATAACGCGTCACGTTCAATAATTGATCCAGTGCACGGCGAAGGACACCAGGAACGCGTCACGTTCAATAACGCGTCACGTTCAATCAACGTCCATGTCGGGATGC |
| PA14_04930 | rpoH | GTTGAAACGCTTCACGGCCTTCATCAGGCC | TTTCGTCCGCGAGTGACCAGCGCGTCACGTTCAATAACGCGTCACGTTCAATAAGTTGAAACGCTTCACGGCCTTCATCAGGCCAACGCGTCACGTTCAATAACGCGTCACGTTCAATCAACGTCCATGTCGGGATGC |
| PA14_04930 | rpoH | AGAAGAGGCGTTCGGCCAGTTCACGCTCCT | TTTCGTCCGCGAGTGACCAGCGCGTCACGTTCAATAACGCGTCACGTTCAATAAAGAAGAGGCGTTCGGCCAGTTCACGCTCCTAACGCGTCACGTTCAATAACGCGTCACGTTCAATCAACGTCCATGTCGGGATGC |
| PA14_04930 | rpoH | TCGTTGTTCAACCAGGCCAGGCGCTTCTTC | TTTCGTCCGCGAGTGACCAGCGCGTCACGTTCAATAACGCGTCACGTTCAATAATCGTTGTTCAACCAGGCCAGGCGCTTCTTCAACGCGTCACGTTCAATAACGCGTCACGTTCAATCAACGTCCATGTCGGGATGC |
| PA14_04930 | rpoH | TGTACTTTTCCGCCAGGTCGTGCAGCGTGG | TTTCGTCCGCGAGTGACCAGCGCGTCACGTTCAATAACGCGTCACGTTCAATAATGTACTTTTCCGCCAGGTCGTGCAGCGTGGAACGCGTCACGTTCAATAACGCGTCACGTTCAATCAACGTCCATGTCGGGATGC |
| PA14_04930 | rpoH | TCCGGCGACAACAGCGGAATGCTGTTCACC | TTTCGTCCGCGAGTGACCAGCGCGTCACGTTCAATAACGCGTCACGTTCAATAATCCGGCGACAACAGCGGAATGCTGTTCACCAACGCGTCACGTTCAATAACGCGTCACGTTCAATCAACGTCCATGTCGGGATGC |
| PA14_04930 | rpoH | TGACGATCCGCCAGTTGCGCAGGATGAACT | TTTCGTCCGCGAGTGACCAGCGCGTCACGTTCAATAACGCGTCACGTTCAATAATGACGATCCGCCAGTTGCGCAGGATGAACTAACGCGTCACGTTCAATAACGCGTCACGTTCAATCAACGTCCATGTCGGGATGC |
| PA14_04930 | rpoH | CCAAGGCATGTACAGGTTGCAAAGAAGTGG | TTTCGTCCGCGAGTGACCAGCGCGTCACGTTCAATAACGCGTCACGTTCAATAACCAAGGCATGTACAGGTTGCAAAGAAGTGGAACGCGTCACGTTCAATAACGCGTCACGTTCAATCAACGTCCATGTCGGGATGC |
| PA14_04930 | rpoH | AGCTGTCGCTCCAGTCGGCGTCTTCCAACT | TTTCGTCCGCGAGTGACCAGCGCGTCACGTTCAATAACGCGTCACGTTCAATAAAGCTGTCGCTCCAGTCGGCGTCTTCCAACTAACGCGTCACGTTCAATAACGCGTCACGTTCAATCAACGTCCATGTCGGGATGC |
| PA14_04930 | rpoH | AAGAACAGCTTGCGCTGCGCCTTGGTGGTG | TTTCGTCCGCGAGTGACCAGCGCGTCACGTTCAATAACGCGTCACGTTCAATAAAAGAACAGCTTGCGCTGCGCCTTGGTGGTGAACGCGTCACGTTCAATAACGCGTCACGTTCAATCAACGTCCATGTCGGGATGC |
| PA14_04930 | rpoH | CAGCTTGCTCATGGCGTTCTTTTCCAGCTG | TTTCGTCCGCGAGTGACCAGCGCGTCACGTTCAATAACGCGTCACGTTCAATAACAGCTTGCTCATGGCGTTCTTTTCCAGCTGAACGCGTCACGTTCAATAACGCGTCACGTTCAATCAACGTCCATGTCGGGATGC |
| PA14_04930 | rpoH | AACGAAGCGCAGGTGCGCCAACACCATCTG | TTTCGTCCGCGAGTGACCAGCGCGTCACGTTCAATAACGCGTCACGTTCAATAAAACGAAGCGCAGGTGCGCCAACACCATCTGAACGCGTCACGTTCAATAACGCGTCACGTTCAATCAACGTCCATGTCGGGATGC |
| PA14_04930 | rpoH | GTGCACATAGGCTTCCAGGTTTGCGCCTGG | TTTCGTCCGCGAGTGACCAGCGCGTCACGTTCAATAACGCGTCACGTTCAATAAGTGCACATAGGCTTCCAGGTTTGCGCCTGGAACGCGTCACGTTCAATAACGCGTCACGTTCAATCAACGTCCATGTCGGGATGC |
| PA14_07520 | rpoD | TTCGGCTTGCTCTTCAGGACGCTGTCGACC | TTTCGTCCGCGAGTGACCAGCTGATAAGCTTGGACAACTGATAAGCTTGGACAATTCGGCTTGCTCTTCAGGACGCTGTCGACCAACTGATAAGCTTGGACAACTGATAAGCTTGGACCAACGTCCATGTCGGGATGC |
| PA14_07520 | rpoD | CGTCGTCCTTCTCTTTCGAGTCGGCGGAAT | TTTCGTCCGCGAGTGACCAGCTGATAAGCTTGGACAACTGATAAGCTTGGACAACGTCGTCCTTCTCTTTCGAGTCGGCGGAATAACTGATAAGCTTGGACAACTGATAAGCTTGGACCAACGTCCATGTCGGGATGC |
| PA14_07520 | rpoD | TGCTCACGGCCTCGGGCGATCAACTCTTTC | TTTCGTCCGCGAGTGACCAGCTGATAAGCTTGGACAACTGATAAGCTTGGACAATGCTCACGGCCTCGGGCGATCAACTCTTTCAACTGATAAGCTTGGACAACTGATAAGCTTGGACCAACGTCCATGTCGGGATGC |
| PA14_07520 | rpoD | TTCTCGTCGGTCTCGTGGTTCGGGAACAGG | TTTCGTCCGCGAGTGACCAGCTGATAAGCTTGGACAACTGATAAGCTTGGACAATTCTCGTCGGTCTCGTGGTTCGGGAACAGGAACTGATAAGCTTGGACAACTGATAAGCTTGGACCAACGTCCATGTCGGGATGC |
| PA14_07520 | rpoD | TGGGCTCGCGACCCATTTCCTGGAGCATCT | TTTCGTCCGCGAGTGACCAGCTGATAAGCTTGGACAACTGATAAGCTTGGACAATGGGCTCGCGACCCATTTCCTGGAGCATCTAACTGATAAGCTTGGACAACTGATAAGCTTGGACCAACGTCCATGTCGGGATGC |
| PA14_07520 | rpoD | ACGCAGAGTTGCATGATGGCGCGTTCCTGG | TTTCGTCCGCGAGTGACCAGCTGATAAGCTTGGACAACTGATAAGCTTGGACAAACGCAGAGTTGCATGATGGCGCGTTCCTGGAACTGATAAGCTTGGACAACTGATAAGCTTGGACCAACGTCCATGTCGGGATGC |
| PA14_07520 | rpoD | CGATCGGGGTTTCCATGGAGATCGGCTCTT | TTTCGTCCGCGAGTGACCAGCTGATAAGCTTGGACAACTGATAAGCTTGGACAACGATCGGGGTTTCCATGGAGATCGGCTCTTAACTGATAAGCTTGGACAACTGATAAGCTTGGACCAACGTCCATGTCGGGATGC |
| PA14_07520 | rpoD | ATATAGCCGCTGAGGACGTCGGAGAGGCGA | TTTCGTCCGCGAGTGACCAGCTGATAAGCTTGGACAACTGATAAGCTTGGACAAATATAGCCGCTGAGGACGTCGGAGAGGCGAAACTGATAAGCTTGGACAACTGATAAGCTTGGACCAACGTCCATGTCGGGATGC |
| PA14_07520 | rpoD | GGTCGTTGACCTCCGCGTAAGTCAGGTATC | TTTCGTCCGCGAGTGACCAGCTGATAAGCTTGGACAACTGATAAGCTTGGACAAGGTCGTTGACCTCCGCGTAAGTCAGGTATCAACTGATAAGCTTGGACAACTGATAAGCTTGGACCAACGTCCATGTCGGGATGC |
| PA14_07520 | rpoD | ACGAGATTGCTGTTGCGCTTTTCCGGACAT | TTTCGTCCGCGAGTGACCAGCTGATAAGCTTGGACAACTGATAAGCTTGGACAAACGAGATTGCTGTTGCGCTTTTCCGGACATAACTGATAAGCTTGGACAACTGATAAGCTTGGACCAACGTCCATGTCGGGATGC |
| PA14_07520 | rpoD | GCGATCTTCAGTACCTTGCGGATCTTGTCC | TTTCGTCCGCGAGTGACCAGCTGATAAGCTTGGACAACTGATAAGCTTGGACAAGCGATCTTCAGTACCTTGCGGATCTTGTCCAACTGATAAGCTTGGACAACTGATAAGCTTGGACCAACGTCCATGTCGGGATGC |
| PA14_07520 | rpoD | CTGTTCCGGATCGGAAATATCCTCCGGCAG | TTTCGTCCGCGAGTGACCAGCTGATAAGCTTGGACAACTGATAAGCTTGGACAACTGTTCCGGATCGGAAATATCCTCCGGCAGAACTGATAAGCTTGGACAACTGATAAGCTTGGACCAACGTCCATGTCGGGATGC |
| PA14_07520 | rpoD | GTGGCGTAGGTAGAGAATTTGTAGCCACGA | TTTCGTCCGCGAGTGACCAGCTGATAAGCTTGGACAACTGATAAGCTTGGACAAGTGGCGTAGGTAGAGAATTTGTAGCCACGAAACTGATAAGCTTGGACAACTGATAAGCTTGGACCAACGTCCATGTCGGGATGC |
| PA14_07520 | rpoD | CCGCATCCGGGGCTGTCTCGAATACGTTGA | TTTCGTCCGCGAGTGACCAGCTGATAAGCTTGGACAACTGATAAGCTTGGACAACCGCATCCGGGGCTGTCTCGAATACGTTGAAACTGATAAGCTTGGACAACTGATAAGCTTGGACCAACGTCCATGTCGGGATGC |
| PA14_07520 | rpoD | GGAGATGCGGTTGAGCTTGTTGATCGTCTC | TTTCGTCCGCGAGTGACCAGCTGATAAGCTTGGACAACTGATAAGCTTGGACAAGGAGATGCGGTTGAGCTTGTTGATCGTCTCAACTGATAAGCTTGGACAACTGATAAGCTTGGACCAACGTCCATGTCGGGATGC |
| PA14_57940 | rpoN | ACATCGGTGCCAGGTTGACCTGCCAGAGCA | TTTCGTCCGCGAGTGACCAGATGCACGACTTCGGAAAATGCACGACTTCGGAAAACATCGGTGCCAGGTTGACCTGCCAGAGCAAAATGCACGACTTCGGAAAATGCACGACTTCGGACAACGTCCATGTCGGGATGC |
| PA14_57940 | rpoN | TCGTCGAGTTCGACGTCCAGTTCCGGATCG | TTTCGTCCGCGAGTGACCAGATGCACGACTTCGGAAAATGCACGACTTCGGAAATCGTCGAGTTCGACGTCCAGTTCCGGATCGAAATGCACGACTTCGGAAAATGCACGACTTCGGACAACGTCCATGTCGGGATGC |
| PA14_57940 | rpoN | CCTTCAGCGGTACTGACGTGACTTGAAAAG | TTTCGTCCGCGAGTGACCAGATGCACGACTTCGGAAAATGCACGACTTCGGAAACCTTCAGCGGTACTGACGTGACTTGAAAAGAAATGCACGACTTCGGAAAATGCACGACTTCGGACAACGTCCATGTCGGGATGC |
| PA14_57940 | rpoN | TCCTGGATTTCCTGCTGGAGGTCCAGGGTG | TTTCGTCCGCGAGTGACCAGATGCACGACTTCGGAAAATGCACGACTTCGGAAATCCTGGATTTCCTGCTGGAGGTCCAGGGTGAAATGCACGACTTCGGAAAATGCACGACTTCGGACAACGTCCATGTCGGGATGC |
| PA14_57940 | rpoN | TTCAGCTCGAAAATGCCGCGCGGCGTATGC | TTTCGTCCGCGAGTGACCAGATGCACGACTTCGGAAAATGCACGACTTCGGAAATTCAGCTCGAAAATGCCGCGCGGCGTATGCAAATGCACGACTTCGGAAAATGCACGACTTCGGACAACGTCCATGTCGGGATGC |
| PA14_57940 | rpoN | GTCGCTGTTGTCGAAATCGTCGCCGTCTTC | TTTCGTCCGCGAGTGACCAGATGCACGACTTCGGAAAATGCACGACTTCGGAAAGTCGCTGTTGTCGAAATCGTCGCCGTCTTCAAATGCACGACTTCGGAAAATGCACGACTTCGGACAACGTCCATGTCGGGATGC |
| PA14_57940 | rpoN | AGATGTCTTCCCAGGCGGTATCCACCGGCA | TTTCGTCCGCGAGTGACCAGATGCACGACTTCGGAAAATGCACGACTTCGGAAAAGATGTCTTCCCAGGCGGTATCCACCGGCAAAATGCACGACTTCGGAAAATGCACGACTTCGGACAACGTCCATGTCGGGATGC |
| PA14_57940 | rpoN | TAGCTGGAGGAGGCGTATGGCCTGTTGCAG | TTTCGTCCGCGAGTGACCAGATGCACGACTTCGGAAAATGCACGACTTCGGAAATAGCTGGAGGAGGCGTATGGCCTGTTGCAGAAATGCACGACTTCGGAAAATGCACGACTTCGGACAACGTCCATGTCGGGATGC |
| PA14_57940 | rpoN | GGCGTTCGAGCATGGGATTGGATTCCAGCG | TTTCGTCCGCGAGTGACCAGATGCACGACTTCGGAAAATGCACGACTTCGGAAAGGCGTTCGAGCATGGGATTGGATTCCAGCGAAATGCACGACTTCGGAAAATGCACGACTTCGGACAACGTCCATGTCGGGATGC |
| PA14_57940 | rpoN | ATGCGCCGCATCAATTGGCTGTAGTCGCGG | TTTCGTCCGCGAGTGACCAGATGCACGACTTCGGAAAATGCACGACTTCGGAAAATGCGCCGCATCAATTGGCTGTAGTCGCGGAAATGCACGACTTCGGAAAATGCACGACTTCGGACAACGTCCATGTCGGGATGC |
| PA14_57940 | rpoN | CAGCGATCTTGCTGTCGCTCAATGGCTTTT | TTTCGTCCGCGAGTGACCAGATGCACGACTTCGGAAAATGCACGACTTCGGAAACAGCGATCTTGCTGTCGCTCAATGGCTTTTAAATGCACGACTTCGGAAAATGCACGACTTCGGACAACGTCCATGTCGGGATGC |
| PA14_57940 | rpoN | GGGACTCTTCGAGATAGCCATCGTTGTTGA | TTTCGTCCGCGAGTGACCAGATGCACGACTTCGGAAAATGCACGACTTCGGAAAGGGACTCTTCGAGATAGCCATCGTTGTTGAAAATGCACGACTTCGGAAAATGCACGACTTCGGACAACGTCCATGTCGGGATGC |
| PA14_57940 | rpoN | AGTCGGATACCAGGCGCAGCGCTTCGTTCA | TTTCGTCCGCGAGTGACCAGATGCACGACTTCGGAAAATGCACGACTTCGGAAAAGTCGGATACCAGGCGCAGCGCTTCGTTCAAAATGCACGACTTCGGAAAATGCACGACTTCGGACAACGTCCATGTCGGGATGC |
| PA14_57940 | rpoN | AGCCACTTGAATGCCCTGTGCCTCCAGTAA | TTTCGTCCGCGAGTGACCAGATGCACGACTTCGGAAAATGCACGACTTCGGAAAAGCCACTTGAATGCCCTGTGCCTCCAGTAAAAATGCACGACTTCGGAAAATGCACGACTTCGGACAACGTCCATGTCGGGATGC |
| PA14_57940 | rpoN | TCGGCTTCGCTCGATTCGATCTGCGAACCG | TTTCGTCCGCGAGTGACCAGATGCACGACTTCGGAAAATGCACGACTTCGGAAATCGGCTTCGCTCGATTCGATCTGCGAACCGAAATGCACGACTTCGGAAAATGCACGACTTCGGACAACGTCCATGTCGGGATGC |
| PA14_57940 | rpoN | GCCCATCTTGAGGACTAGCGATGGTTTCAT | TTTCGTCCGCGAGTGACCAGATGCACGACTTCGGAAAATGCACGACTTCGGAAAGCCCATCTTGAGGACTAGCGATGGTTTCATAAATGCACGACTTCGGAAAATGCACGACTTCGGACAACGTCCATGTCGGGATGC |
| PA14_57940 | rpoN | TCAGCGTTTCGTTGCGGCTCTGCAGGCTCT | TTTCGTCCGCGAGTGACCAGATGCACGACTTCGGAAAATGCACGACTTCGGAAATCAGCGTTTCGTTGCGGCTCTGCAGGCTCTAAATGCACGACTTCGGAAAATGCACGACTTCGGACAACGTCCATGTCGGGATGC |
| PA14_57940 | rpoN | GATGGTCACCGCGATCATCCGATCGGTGTC | TTTCGTCCGCGAGTGACCAGATGCACGACTTCGGAAAATGCACGACTTCGGAAAGATGGTCACCGCGATCATCCGATCGGTGTCAAATGCACGACTTCGGAAAATGCACGACTTCGGACAACGTCCATGTCGGGATGC |
| PA14_57940 | rpoN | TTCGCTCCACTGGTCGTCGTCGAGGCTTTC | TTTCGTCCGCGAGTGACCAGATGCACGACTTCGGAAAATGCACGACTTCGGAAATTCGCTCCACTGGTCGTCGTCGAGGCTTTCAAATGCACGACTTCGGAAAATGCACGACTTCGGACAACGTCCATGTCGGGATGC |
| PA14_57940 | rpoN | TTCGTCATCGTCGTTGCTGGGCAGGCTGCT | TTTCGTCCGCGAGTGACCAGATGCACGACTTCGGAAAATGCACGACTTCGGAAATTCGTCATCGTCGTTGCTGGGCAGGCTGCTAAATGCACGACTTCGGAAAATGCACGACTTCGGACAACGTCCATGTCGGGATGC |
| PA14_41575 | sigX | ACATGGACGAGCCAACGATCCAATCCGCCC | TTTCGTCCGCGAGTGACCAGGCTGTCGCCTATAAGAAGCTGTCGCCTATAAGAAACATGGACGAGCCAACGATCCAATCCGCCCAAGCTGTCGCCTATAAGAAGCTGTCGCCTATAAGCAACGTCCATGTCGGGATGC |
| PA14_41575 | sigX | AAGCTGGCGTGGGTCATACCGCTGTGACAG | TTTCGTCCGCGAGTGACCAGGCTGTCGCCTATAAGAAGCTGTCGCCTATAAGAAAAGCTGGCGTGGGTCATACCGCTGTGACAGAAGCTGTCGCCTATAAGAAGCTGTCGCCTATAAGCAACGTCCATGTCGGGATGC |
| PA14_41575 | sigX | CTATCTCCTGAAACTCCAGTTCCGCTACGA | TTTCGTCCGCGAGTGACCAGGCTGTCGCCTATAAGAAGCTGTCGCCTATAAGAACTATCTCCTGAAACTCCAGTTCCGCTACGAAAGCTGTCGCCTATAAGAAGCTGTCGCCTATAAGCAACGTCCATGTCGGGATGC |
| PA14_41575 | sigX | AACCTTCGGCGACTTCTCCTCGGAAGCCTC | TTTCGTCCGCGAGTGACCAGGCTGTCGCCTATAAGAAGCTGTCGCCTATAAGAAAACCTTCGGCGACTTCTCCTCGGAAGCCTCAAGCTGTCGCCTATAAGAAGCTGTCGCCTATAAGCAACGTCCATGTCGGGATGC |
| PA14_41575 | sigX | AACAGCGTGCGCTGGTAACGCCGCATCAAT | TTTCGTCCGCGAGTGACCAGGCTGTCGCCTATAAGAAGCTGTCGCCTATAAGAAAACAGCGTGCGCTGGTAACGCCGCATCAATAAGCTGTCGCCTATAAGAAGCTGTCGCCTATAAGCAACGTCCATGTCGGGATGC |
| PA14_41575 | sigX | AGGCTGAGCGCATCCATCAATCGGCGCTTG | TTTCGTCCGCGAGTGACCAGGCTGTCGCCTATAAGAAGCTGTCGCCTATAAGAAAGGCTGAGCGCATCCATCAATCGGCGCTTGAAGCTGTCGCCTATAAGAAGCTGTCGCCTATAAGCAACGTCCATGTCGGGATGC |
| PA14_41575 | sigX | AAAACTTTTCGCGCAGGCGGTCCAGTGCCC | TTTCGTCCGCGAGTGACCAGGCTGTCGCCTATAAGAAGCTGTCGCCTATAAGAAAAAACTTTTCGCGCAGGCGGTCCAGTGCCCAAGCTGTCGCCTATAAGAAGCTGTCGCCTATAAGCAACGTCCATGTCGGGATGC |
| PA14_41575 | sigX | CGAAGAACCAGAATCTCCCGATCTATCGGA | TTTCGTCCGCGAGTGACCAGGCTGTCGCCTATAAGAAGCTGTCGCCTATAAGAACGAAGAACCAGAATCTCCCGATCTATCGGAAAGCTGTCGCCTATAAGAAGCTGTCGCCTATAAGCAACGTCCATGTCGGGATGC |
| PA14_41575 | sigX | CGTCGCACTCAGACCCATGTGCATGATATC | TTTCGTCCGCGAGTGACCAGGCTGTCGCCTATAAGAAGCTGTCGCCTATAAGAACGTCGCACTCAGACCCATGTGCATGATATCAAGCTGTCGCCTATAAGAAGCTGTCGCCTATAAGCAACGTCCATGTCGGGATGC |
| PA14_41575 | sigX | GCGTGATGCACTCGTTGTACGTGATGCTAT | TTTCGTCCGCGAGTGACCAGGCTGTCGCCTATAAGAAGCTGTCGCCTATAAGAAGCGTGATGCACTCGTTGTACGTGATGCTATAAGCTGTCGCCTATAAGAAGCTGTCGCCTATAAGCAACGTCCATGTCGGGATGC |
| PA14_41575 | sigX | TTCATAGGCGCGCGTCACGTGGAACAGCTC | TTTCGTCCGCGAGTGACCAGGCTGTCGCCTATAAGAAGCTGTCGCCTATAAGAATTCATAGGCGCGCGTCACGTGGAACAGCTCAAGCTGTCGCCTATAAGAAGCTGTCGCCTATAAGCAACGTCCATGTCGGGATGC |
| PA14_41575 | sigX | CCGTACAAAACTTTCAGCATCACCTCCTGG | TTTCGTCCGCGAGTGACCAGGCTGTCGCCTATAAGAAGCTGTCGCCTATAAGAACCGTACAAAACTTTCAGCATCACCTCCTGGAAGCTGTCGCCTATAAGAAGCTGTCGCCTATAAGCAACGTCCATGTCGGGATGC |
| PA14_41575 | sigX | GCCATGTTTTGAACTTCGACTTGCCCTCAA | TTTCGTCCGCGAGTGACCAGGCTGTCGCCTATAAGAAGCTGTCGCCTATAAGAAGCCATGTTTTGAACTTCGACTTGCCCTCAAAAGCTGTCGCCTATAAGAAGCTGTCGCCTATAAGCAACGTCCATGTCGGGATGC |
| PA14_08910 | rpsC | TGATGTGCACGGGCACACCCATTTGCTTGG | TTTCGTCCGCGAGTGACCAGTTAATCGGCCGAACGAATTAATCGGCCGAACGAATGATGTGCACGGGCACACCCATTTGCTTGGAATTAATCGGCCGAACGAATTAATCGGCCGAACGCAACGTCCATGTCGGGATGC |
| PA14_08910 | rpsC | CGAATGCCATTGGGATGTACTTTCTGACCC | TTTCGTCCGCGAGTGACCAGTTAATCGGCCGAACGAATTAATCGGCCGAACGAACGAATGCCATTGGGATGTACTTTCTGACCCAATTAATCGGCCGAACGAATTAATCGGCCGAACGCAACGTCCATGTCGGGATGC |
| PA14_08910 | rpsC | GTAATCCGCATAGTTCTTGCGGTCTGCGTA | TTTCGTCCGCGAGTGACCAGTTAATCGGCCGAACGAATTAATCGGCCGAACGAAGTAATCCGCATAGTTCTTGCGGTCTGCGTAAATTAATCGGCCGAACGAATTAATCGGCCGAACGCAACGTCCATGTCGGGATGC |
| PA14_08910 | rpsC | AGGTACTCACGCACCTTCAGGTCGGCAAAC | TTTCGTCCGCGAGTGACCAGTTAATCGGCCGAACGAATTAATCGGCCGAACGAAAGGTACTCACGCACCTTCAGGTCGGCAAACAATTAATCGGCCGAACGAATTAATCGGCCGAACGCAACGTCCATGTCGGGATGC |
| PA14_08910 | rpsC | TTTCTTGCCGATCACGATGCCGGGACGAGC | TTTCGTCCGCGAGTGACCAGTTAATCGGCCGAACGAATTAATCGGCCGAACGAATTTCTTGCCGATCACGATGCCGGGACGAGCAATTAATCGGCCGAACGAATTAATCGGCCGAACGCAACGTCCATGTCGGGATGC |
| PA14_08910 | rpsC | CATCGAGTTCCGGCTTGCGGATTTCCTCGA | TTTCGTCCGCGAGTGACCAGTTAATCGGCCGAACGAATTAATCGGCCGAACGAACATCGAGTTCCGGCTTGCGGATTTCCTCGAAATTAATCGGCCGAACGAATTAATCGGCCGAACGCAACGTCCATGTCGGGATGC |
| PA14_08910 | rpsC | CACCGATCACACCGTAAGTGGTGTGCGCTT | TTTCGTCCGCGAGTGACCAGTTAATCGGCCGAACGAATTAATCGGCCGAACGAACACCGATCACACCGTAAGTGGTGTGCGCTTAATTAATCGGCCGAACGAATTAATCGGCCGAACGCAACGTCCATGTCGGGATGC |
| PA14_08910 | rpsC | GATACCATTCGGTACGGGCGATTTCAGCCC | TTTCGTCCGCGAGTGACCAGTTAATCGGCCGAACGAATTAATCGGCCGAACGAAGATACCATTCGGTACGGGCGATTTCAGCCCAATTAATCGGCCGAACGAATTAATCGGCCGAACGCAACGTCCATGTCGGGATGC |
| PA14_08910 | rpsC | CCCAGACGACCGCTGACCTGAATCTTGATG | TTTCGTCCGCGAGTGACCAGTTAATCGGCCGAACGAATTAATCGGCCGAACGAACCCAGACGACCGCTGACCTGAATCTTGATGAATTAATCGGCCGAACGAATTAATCGGCCGAACGCAACGTCCATGTCGGGATGC |
| PA14_08910 | rpsC | GAATATCGATACGGCTTACGGACGCGCTTT | TTTCGTCCGCGAGTGACCAGTTAATCGGCCGAACGAATTAATCGGCCGAACGAAGAATATCGATACGGCTTACGGACGCGCTTTAATTAATCGGCCGAACGAATTAATCGGCCGAACGCAACGTCCATGTCGGGATGC |
| PA14_08910 | rpsC | AGGTCCTGGCGCAGTTTCTCAACATCTTCA | TTTCGTCCGCGAGTGACCAGTTAATCGGCCGAACGAATTAATCGGCCGAACGAAAGGTCCTGGCGCAGTTTCTCAACATCTTCAAATTAATCGGCCGAACGAATTAATCGGCCGAACGCAACGTCCATGTCGGGATGC |
| PA14_08910 | rpsC | ACATTACGCGACGCTCCAGCTGCTGGGCTA | TTTCGTCCGCGAGTGACCAGTTAATCGGCCGAACGAATTAATCGGCCGAACGAAACATTACGCGACGCTCCAGCTGCTGGGCTAAATTAATCGGCCGAACGAATTAATCGGCCGAACGCAACGTCCATGTCGGGATGC |
| PA14_08910 | rpsC | TGGATGGTGATGCGTGCGGTTTGAGCCGGA | TTTCGTCCGCGAGTGACCAGTTAATCGGCCGAACGAATTAATCGGCCGAACGAATGGATGGTGATGCGTGCGGTTTGAGCCGGAAATTAATCGGCCGAACGAATTAATCGGCCGAACGCAACGTCCATGTCGGGATGC |
| PA14_08910 | rpsC | TTTACGAGGAGCGGGTGCAACCGGCTTCAG | TTTCGTCCGCGAGTGACCAGTTAATCGGCCGAACGAATTAATCGGCCGAACGAATTTACGAGGAGCGGGTGCAACCGGCTTCAGAATTAATCGGCCGAACGAATTAATCGGCCGAACGCAACGTCCATGTCGGGATGC |
| PA14_08910 | rpsC | AAACGGAGGTGTGCTCCTTGACGATACCCA | TTTCGTCCGCGAGTGACCAGTTAATCGGCCGAACGAATTAATCGGCCGAACGAAAAACGGAGGTGTGCTCCTTGACGATACCCAAATTAATCGGCCGAACGAATTAATCGGCCGAACGCAACGTCCATGTCGGGATGC |
| PA14_17480 | rpoS | AATCCGCGCTCCGGATCGAACTTCTCCACG | TTTCGTCCGCGAGTGACCAGAATTCGACGGTCTAGAAAATTCGACGGTCTAGAAAATCCGCGCTCCGGATCGAACTTCTCCACGAAAATTCGACGGTCTAGAAAATTCGACGGTCTAGCAACGTCCATGTCGGGATGC |
| PA14_17480 | rpoS | ATGGATCGGCAAGCGAATGGTCCGGGTCTG | TTTCGTCCGCGAGTGACCAGAATTCGACGGTCTAGAAAATTCGACGGTCTAGAAATGGATCGGCAAGCGAATGGTCCGGGTCTGAAAATTCGACGGTCTAGAAAATTCGACGGTCTAGCAACGTCCATGTCGGGATGC |
| PA14_17480 | rpoS | GAAGTCACCCGTTCGTTCAGGCCGAGCATG | TTTCGTCCGCGAGTGACCAGAATTCGACGGTCTAGAAAATTCGACGGTCTAGAAGAAGTCACCCGTTCGTTCAGGCCGAGCATGAAAATTCGACGGTCTAGAAAATTCGACGGTCTAGCAACGTCCATGTCGGGATGC |
| PA14_17480 | rpoS | TCTCGATCATCCGCTTCCGACCAGCGGGAT | TTTCGTCCGCGAGTGACCAGAATTCGACGGTCTAGAAAATTCGACGGTCTAGAATCTCGATCATCCGCTTCCGACCAGCGGGATAAAATTCGACGGTCTAGAAAATTCGACGGTCTAGCAACGTCCATGTCGGGATGC |
| PA14_17480 | rpoS | TCGGGCGTCAACAGGGGCGAGAAACCGATT | TTTCGTCCGCGAGTGACCAGAATTCGACGGTCTAGAAAATTCGACGGTCTAGAATCGGGCGTCAACAGGGGCGAGAAACCGATTAAAATTCGACGGTCTAGAAAATTCGACGGTCTAGCAACGTCCATGTCGGGATGC |
| PA14_17480 | rpoS | ACATAGCGCCGGGCGATCTTCACCACCAAC | TTTCGTCCGCGAGTGACCAGAATTCGACGGTCTAGAAAATTCGACGGTCTAGAAACATAGCGCCGGGCGATCTTCACCACCAACAAAATTCGACGGTCTAGAAAATTCGACGGTCTAGCAACGTCCATGTCGGGATGC |
| PA14_17480 | rpoS | TGTCCGAGTCCGGACCAAGAGAGACGTCTA | TTTCGTCCGCGAGTGACCAGAATTCGACGGTCTAGAAAATTCGACGGTCTAGAATGTCCGAGTCCGGACCAAGAGAGACGTCTAAAAATTCGACGGTCTAGAAAATTCGACGGTCTAGCAACGTCCATGTCGGGATGC |
| PA14_17480 | rpoS | TCGTCGGCAGACGACTCGTCCAGCATGATG | TTTCGTCCGCGAGTGACCAGAATTCGACGGTCTAGAAAATTCGACGGTCTAGAATCGTCGGCAGACGACTCGTCCAGCATGATGAAAATTCGACGGTCTAGAAAATTCGACGGTCTAGCAACGTCCATGTCGGGATGC |
| PA14_17480 | rpoS | GTGAAGGTTCGTGGTCGAGCTTGTGGGTCA | TTTCGTCCGCGAGTGACCAGAATTCGACGGTCTAGAAAATTCGACGGTCTAGAAGTGAAGGTTCGTGGTCGAGCTTGTGGGTCAAAAATTCGACGGTCTAGAAAATTCGACGGTCTAGCAACGTCCATGTCGGGATGC |
| PA14_17480 | rpoS | TCATCGTGGTCAAACTCCGGCCCTTCTTTT | TTTCGTCCGCGAGTGACCAGAATTCGACGGTCTAGAAAATTCGACGGTCTAGAATCATCGTGGTCAAACTCCGGCCCTTCTTTTAAAATTCGACGGTCTAGAAAATTCGACGGTCTAGCAACGTCCATGTCGGGATGC |
| PA14_17480 | rpoS | ATCGTCATCCTGCAGCAGCTCGCACGGATC | TTTCGTCCGCGAGTGACCAGAATTCGACGGTCTAGAAAATTCGACGGTCTAGAAATCGTCATCCTGCAGCAGCTCGCACGGATCAAAATTCGACGGTCTAGAAAATTCGACGGTCTAGCAACGTCCATGTCGGGATGC |
| PA14_17480 | rpoS | TGGCTTTTGGAGTTGCCCGGGGAGAAGGCT | TTTCGTCCGCGAGTGACCAGAATTCGACGGTCTAGAAAATTCGACGGTCTAGAATGGCTTTTGGAGTTGCCCGGGGAGAAGGCTAAAATTCGACGGTCTAGAAAATTCGACGGTCTAGCAACGTCCATGTCGGGATGC |
| PA14_17480 | rpoS | TCTTCTCCAGGATCTCCCGCAGGCGCTTCA | TTTCGTCCGCGAGTGACCAGAATTCGACGGTCTAGAAAATTCGACGGTCTAGAATCTTCTCCAGGATCTCCCGCAGGCGCTTCAAAAATTCGACGGTCTAGAAAATTCGACGGTCTAGCAACGTCCATGTCGGGATGC |
| PA14_17480 | rpoS | TGTCGGTGAGTTCCGTCAGCCACTGGTCGA | TTTCGTCCGCGAGTGACCAGAATTCGACGGTCTAGAAAATTCGACGGTCTAGAATGTCGGTGAGTTCCGTCAGCCACTGGTCGAAAAATTCGACGGTCTAGAAAATTCGACGGTCTAGCAACGTCCATGTCGGGATGC |
| PA14_17480 | rpoS | TAGGTTGCCTTCCTCGATCAGGTCGAGCAG | TTTCGTCCGCGAGTGACCAGAATTCGACGGTCTAGAAAATTCGACGGTCTAGAATAGGTTGCCTTCCTCGATCAGGTCGAGCAGAAAATTCGACGGTCTAGAAAATTCGACGGTCTAGCAACGTCCATGTCGGGATGC |
| PA14_17480 | rpoS | GCACGCAGGTAGACGTTGAGCTCCTTGACC | TTTCGTCCGCGAGTGACCAGAATTCGACGGTCTAGAAAATTCGACGGTCTAGAAGCACGCAGGTAGACGTTGAGCTCCTTGACCAAAATTCGACGGTCTAGAAAATTCGACGGTCTAGCAACGTCCATGTCGGGATGC |
| PA14_17480 | rpoS | GTTGAGATACAGCTGCGTTGCGTCCAACGC | TTTCGTCCGCGAGTGACCAGAATTCGACGGTCTAGAAAATTCGACGGTCTAGAAGTTGAGATACAGCTGCGTTGCGTCCAACGCAAAATTCGACGGTCTAGAAAATTCGACGGTCTAGCAACGTCCATGTCGGGATGC |
| PA14_17480 | rpoS | ATGATGGCCCGTTCGATGGTCTGGCGGATC | TTTCGTCCGCGAGTGACCAGAATTCGACGGTCTAGAAAATTCGACGGTCTAGAAATGATGGCCCGTTCGATGGTCTGGCGGATCAAAATTCGACGGTCTAGAAAATTCGACGGTCTAGCAACGTCCATGTCGGGATGC |
| PA14_17480 | rpoS | GCGTGTAGTCGATGTGCTTGTGTTGTTTGG | TTTCGTCCGCGAGTGACCAGAATTCGACGGTCTAGAAAATTCGACGGTCTAGAAGCGTGTAGTCGATGTGCTTGTGTTGTTTGGAAAATTCGACGGTCTAGAAAATTCGACGGTCTAGCAACGTCCATGTCGGGATGC |
| PA14_17480 | rpoS | TCTGCGCCAGACGAGCGAAGTGGACTTCCT | TTTCGTCCGCGAGTGACCAGAATTCGACGGTCTAGAAAATTCGACGGTCTAGAATCTGCGCCAGACGAGCGAAGTGGACTTCCTAAAATTCGACGGTCTAGAAAATTCGACGGTCTAGCAACGTCCATGTCGGGATGC |
| PA14_58000 | sodM | GTTCAGGTTGTTCACGTAGGTCTGGTGGTG | TTTCGTCCGCGAGTGACCAGGCGACTGGAACACTTAAGCGACTGGAACACTTAAGTTCAGGTTGTTCACGTAGGTCTGGTGGTGAAGCGACTGGAACACTTAAGCGACTGGAACACTTCAACGTCCATGTCGGGATGC |
| PA14_58000 | sodM | TCCAGAACAGCGAGTGGTTGGCATGCCCAC | TTTCGTCCGCGAGTGACCAGGCGACTGGAACACTTAAGCGACTGGAACACTTAATCCAGAACAGCGAGTGGTTGGCATGCCCACAAGCGACTGGAACACTTAAGCGACTGGAACACTTCAACGTCCATGTCGGGATGC |
| PA14_58000 | sodM | AATAGGCGTGCTCCCATACGTCCAGGCCGA | TTTCGTCCGCGAGTGACCAGGCGACTGGAACACTTAAGCGACTGGAACACTTAAAATAGGCGTGCTCCCATACGTCCAGGCCGAAAGCGACTGGAACACTTAAGCGACTGGAACACTTCAACGTCCATGTCGGGATGC |
| PA14_58000 | sodM | GTATTCCGGGCGGCGGTTCTGGTACTTCAG | TTTCGTCCGCGAGTGACCAGGCGACTGGAACACTTAAGCGACTGGAACACTTAAGTATTCCGGGCGGCGGTTCTGGTACTTCAGAAGCGACTGGAACACTTAAGCGACTGGAACACTTCAACGTCCATGTCGGGATGC |
| PA14_58000 | sodM | CGCCAGTCGATCACGTTGTAGAAAGCGCCG | TTTCGTCCGCGAGTGACCAGGCGACTGGAACACTTAAGCGACTGGAACACTTAACGCCAGTCGATCACGTTGTAGAAAGCGCCGAAGCGACTGGAACACTTAAGCGACTGGAACACTTCAACGTCCATGTCGGGATGC |
| PA14_58000 | sodM | TGAACGCATCCTTGAACGCCTCGAAGCCCC | TTTCGTCCGCGAGTGACCAGGCGACTGGAACACTTAAGCGACTGGAACACTTAATGAACGCATCCTTGAACGCCTCGAAGCCCCAAGCGACTGGAACACTTAAGCGACTGGAACACTTCAACGTCCATGTCGGGATGC |
| PA14_58000 | sodM | TCGGCGTATTGCCGTTCATCAGCGGGCTGT | TTTCGTCCGCGAGTGACCAGGCGACTGGAACACTTAAGCGACTGGAACACTTAATCGGCGTATTGCCGTTCATCAGCGGGCTGTAAGCGACTGGAACACTTAAGCGACTGGAACACTTCAACGTCCATGTCGGGATGC |
| PA14_58000 | sodM | TTGTTGACAACCGGGGTGCGCAGCTTCTCC | TTTCGTCCGCGAGTGACCAGGCGACTGGAACACTTAAGCGACTGGAACACTTAATTGTTGACAACCGGGGTGCGCAGCTTCTCCAAGCGACTGGAACACTTAAGCGACTGGAACACTTCAACGTCCATGTCGGGATGC |
| PA14_58000 | sodM | ATGTGCGGTTCCAGGGCATCGTAGGCGTAG | TTTCGTCCGCGAGTGACCAGGCGACTGGAACACTTAAGCGACTGGAACACTTAAATGTGCGGTTCCAGGGCATCGTAGGCGTAGAAGCGACTGGAACACTTAAGCGACTGGAACACTTCAACGTCCATGTCGGGATGC |
| PA14_58000 | sodM | TGGTTACCGCTGCTCTCCACCAGCAAGCTG | TTTCGTCCGCGAGTGACCAGGCGACTGGAACACTTAAGCGACTGGAACACTTAATGGTTACCGCTGCTCTCCACCAGCAAGCTGAAGCGACTGGAACACTTAAGCGACTGGAACACTTCAACGTCCATGTCGGGATGC |
| PA14_58000 | sodM | TGCTGTGGTGGATCTCCATGGTCAGCGCAT | TTTCGTCCGCGAGTGACCAGGCGACTGGAACACTTAAGCGACTGGAACACTTAATGCTGTGGTGGATCTCCATGGTCAGCGCATAAGCGACTGGAACACTTAAGCGACTGGAACACTTCAACGTCCATGTCGGGATGC |
| PA14_58000 | sodM | AGCAGGCTTTCCACCGGCTGCTCGGCATAG | TTTCGTCCGCGAGTGACCAGGCGACTGGAACACTTAAGCGACTGGAACACTTAAAGCAGGCTTTCCACCGGCTGCTCGGCATAGAAGCGACTGGAACACTTAAGCGACTGGAACACTTCAACGTCCATGTCGGGATGC |
| PA14_03920 | spuD | CTTGTCGAGCTTCTGGTAGACCCCGGCCTT | TTTCGTCCGCGAGTGACCAGTGATACGCTTGGCATAATGATACGCTTGGCATAACTTGTCGAGCTTCTGGTAGACCCCGGCCTTAATGATACGCTTGGCATAATGATACGCTTGGCATCAACGTCCATGTCGGGATGC |
| PA14_03920 | spuD | CGCCGCACTGCTTCAGCTTCTGGATGTTCT | TTTCGTCCGCGAGTGACCAGTGATACGCTTGGCATAATGATACGCTTGGCATAACGCCGCACTGCTTCAGCTTCTGGATGTTCTAATGATACGCTTGGCATAATGATACGCTTGGCATCAACGTCCATGTCGGGATGC |
| PA14_03920 | spuD | TACCAGGTCCCAGGAATCCACCGGCGCATT | TTTCGTCCGCGAGTGACCAGTGATACGCTTGGCATAATGATACGCTTGGCATAATACCAGGTCCCAGGAATCCACCGGCGCATTAATGATACGCTTGGCATAATGATACGCTTGGCATCAACGTCCATGTCGGGATGC |
| PA14_03920 | spuD | TGTAGGGAATCGCATGCTCGTTGCCCGGGT | TTTCGTCCGCGAGTGACCAGTGATACGCTTGGCATAATGATACGCTTGGCATAATGTAGGGAATCGCATGCTCGTTGCCCGGGTAATGATACGCTTGGCATAATGATACGCTTGGCATCAACGTCCATGTCGGGATGC |
| PA14_03920 | spuD | CGAGCAGGGTCTTACCGAAACGTTTCATCA | TTTCGTCCGCGAGTGACCAGTGATACGCTTGGCATAATGATACGCTTGGCATAACGAGCAGGGTCTTACCGAAACGTTTCATCAAATGATACGCTTGGCATAATGATACGCTTGGCATCAACGTCCATGTCGGGATGC |
| PA14_03920 | spuD | GCGATGTAGTCGGACCAGTTGTACACGTGC | TTTCGTCCGCGAGTGACCAGTGATACGCTTGGCATAATGATACGCTTGGCATAAGCGATGTAGTCGGACCAGTTGTACACGTGCAATGATACGCTTGGCATAATGATACGCTTGGCATCAACGTCCATGTCGGGATGC |
| PA14_03920 | spuD | AAGGCCGCCTTGACCTTGTCCGGGTTGTAG | TTTCGTCCGCGAGTGACCAGTGATACGCTTGGCATAATGATACGCTTGGCATAAAAGGCCGCCTTGACCTTGTCCGGGTTGTAGAATGATACGCTTGGCATAATGATACGCTTGGCATCAACGTCCATGTCGGGATGC |
| PA14_03920 | spuD | TACCTTGTTGTCCGCAGCCTGCGCCATACC | TTTCGTCCGCGAGTGACCAGTGATACGCTTGGCATAATGATACGCTTGGCATAATACCTTGTTGTCCGCAGCCTGCGCCATACCAATGATACGCTTGGCATAATGATACGCTTGGCATCAACGTCCATGTCGGGATGC |
| PA14_03920 | spuD | GGTCGGAAATGTACTTCGACGAGTGGAAGT | TTTCGTCCGCGAGTGACCAGTGATACGCTTGGCATAATGATACGCTTGGCATAAGGTCGGAAATGTACTTCGACGAGTGGAAGTAATGATACGCTTGGCATAATGATACGCTTGGCATCAACGTCCATGTCGGGATGC |
| PA14_03920 | spuD | GCATCAGGTCCTTGTTCAGGTTCTTCCAGT | TTTCGTCCGCGAGTGACCAGTGATACGCTTGGCATAATGATACGCTTGGCATAAGCATCAGGTCCTTGTTCAGGTTCTTCCAGTAATGATACGCTTGGCATAATGATACGCTTGGCATCAACGTCCATGTCGGGATGC |

| | | | |
|---|---|---|---|
| PA14_03920 | spuD | TTTCCTTGGTGAACTTCTCCAGGGTGTCCG | TTTCGTCCGCGAGTGACCAGTGATACGCTTGGCATAATGATACGCTTGGCATAATTTCCTTGGTGAACTTCTCCAGGGTGTCCGAATGATACGCTTGGCATAATGATACGCTTGGCATCAACGTCCATGTCGGGATGC |
| PA14_03920 | spuD | TCTGCTTGGCGAGGAAGGAGTTGGACGGCA | TTTCGTCCGCGAGTGACCAGTGATACGCTTGGCATAATGATACGCTTGGCATAATCTGCTTGGCGAGGAAGGAGTTGGACGGCAAATGATACGCTTGGCATAATGATACGCTTGGCATCAACGTCCATGTCGGGATGC |
| PA14_03920 | spuD | GTCGTAGACGTCGTAGACGACCTTGATCCC | TTTCGTCCGCGAGTGACCAGTGATACGCTTGGCATAATGATACGCTTGGCATAAGTCGTAGACGTCGTAGACGACCTTGATCCCAATGATACGCTTGGCATAATGATACGCTTGGCATCAACGTCCATGTCGGGATGC |
| PA14_03920 | spuD | TCATGACCTCTTCGCTCGGGTAGATGCCCG | TTTCGTCCGCGAGTGACCAGTGATACGCTTGGCATAATGATACGCTTGGCATAATCATGACCTCTTCGCTCGGGTAGATGCCCGAATGATACGCTTGGCATAATGATACGCTTGGCATCAACGTCCATGTCGGGATGC |
| PA14_03920 | spuD | GCCAGCAACTTGGCTTCCAGCACTTCGTTG | TTTCGTCCGCGAGTGACCAGTGATACGCTTGGCATAATGATACGCTTGGCATAAGCCAGCAACTTGGCTTCCAGCACTTCGTTGAATGATACGCTTGGCATAATGATACGCTTGGCATCAACGTCCATGTCGGGATGC |
| PA14_43950 | sucC | ACCTTGAGGATCTTCTCGGGAGTGTCGTGG | TTTCGTCCGCGAGTGACCAGTGGCAGATCGCTAATAATGGCAGATCGCTAATAAACCTTGAGGATCTTCTCGGGAGTGTCGTGGAATGGCAGATCGCTAATAATGGCAGATCGCTAATCAACGTCCATGTCGGGATGC |
| PA14_43950 | sucC | AAGCCATGAACACGATGCGGCGGCTGGAAC | TTTCGTCCGCGAGTGACCAGTGGCAGATCGCTAATAATGGCAGATCGCTAATAAAAGCCATGAACACGATGCGGCGGCTGGAACAATGGCAGATCGCTAATAATGGCAGATCGCTAATCAACGTCCATGTCGGGATGC |
| PA14_43950 | sucC | GTTCACCAGGACAGCCTTCACGTTGCTGTC | TTTCGTCCGCGAGTGACCAGTGGCAGATCGCTAATAATGGCAGATCGCTAATAAGTTCACCAGGACAGCCTTCACGTTGCTGTCAATGGCAGATCGCTAATAATGGCAGATCGCTAATCAACGTCCATGTCGGGATGC |
| PA14_43950 | sucC | TTCACCAGCTTCACACCGCCCGCTTTACCG | TTTCGTCCGCGAGTGACCAGTGGCAGATCGCTAATAATGGCAGATCGCTAATAATTCACCAGCTTCACACCGCCCGCTTTACCGAATGGCAGATCGCTAATAATGGCAGATCGCTAATCAACGTCCATGTCGGGATGC |
| PA14_43950 | sucC | TGCACCTGAGCTTTGACGACCCACTCGCTG | TTTCGTCCGCGAGTGACCAGTGGCAGATCGCTAATAATGGCAGATCGCTAATAATGCACCTGAGCTTTGACGACCCACTCGCTGAATGGCAGATCGCTAATAATGGCAGATCGCTAATCAACGTCCATGTCGGGATGC |
| PA14_43950 | sucC | AGGACTCCACCAGGATCTTGCTGACCGGCT | TTTCGTCCGCGAGTGACCAGTGGCAGATCGCTAATAATGGCAGATCGCTAATAAAGGACTCCACCAGGATCTTGCTGACCGGCTAATGGCAGATCGCTAATAATGGCAGATCGCTAATCAACGTCCATGTCGGGATGC |
| PA14_43950 | sucC | GTCGGTCTGGTAGGTCACCAGGTTCTTGCC | TTTCGTCCGCGAGTGACCAGTGGCAGATCGCTAATAATGGCAGATCGCTAATAAGTCGGTCTGGTAGGTCACCAGGTTCTTGCCAATGGCAGATCGCTAATAATGGCAGATCGCTAATCAACGTCCATGTCGGGATGC |
| PA14_43950 | sucC | ACTTACCCTCCGCGGCCTTGACGACTTGCT | TTTCGTCCGCGAGTGACCAGTGGCAGATCGCTAATAATGGCAGATCGCTAATAAACTTACCCTCCGCGGCCTTGACGACTTGCTAATGGCAGATCGCTAATAATGGCAGATCGCTAATCAACGTCCATGTCGGGATGC |
| PA14_43950 | sucC | GAACAGCTGCTTACCCTGATATTCGTGGAG | TTTCGTCCGCGAGTGACCAGTGGCAGATCGCTAATAATGGCAGATCGCTAATAAGAACAGCTGCTTACCCTGATATTCGTGGAGAATGGCAGATCGCTAATAATGGCAGATCGCTAATCAACGTCCATGTCGGGATGC |
| PA14_43950 | sucC | AAGCCCTTGGATACAGGCAGGCCGTATTCA | TTTCGTCCGCGAGTGACCAGTGGCAGATCGCTAATAATGGCAGATCGCTAATAAAAGCCCTTGGATACAGGCAGGCCGTATTCAAATGGCAGATCGCTAATAATGGCAGATCGCTAATCAACGTCCATGTCGGGATGC |
| PA14_43950 | sucC | TTCCAGGCGGACGACAACCGGAACCTTCAC | TTTCGTCCGCGAGTGACCAGTGGCAGATCGCTAATAATGGCAGATCGCTAATAATTCCAGGCGGACGACAACCGGAACCTTCACAATGGCAGATCGCTAATAATGGCAGATCGCTAATCAACGTCCATGTCGGGATGC |
| PA14_43950 | sucC | ATCTTGTCGCAGGCTTCTGCGGCCTCTTCG | TTTCGTCCGCGAGTGACCAGTGGCAGATCGCTAATAATGGCAGATCGCTAATAAATCTTGTCGCAGGCTTCTGCGGCCTCTTCGAATGGCAGATCGCTAATAATGGCAGATCGCTAATCAACGTCCATGTCGGGATGC |
| PA14_43950 | sucC | GACCTTCTCGATGTCCACGCCACCTTCGGT | TTTCGTCCGCGAGTGACCAGTGGCAGATCGCTAATAATGGCAGATCGCTAATAAGACCTTCTCGATGTCCACGCCACCTTCGGTAATGGCAGATCGCTAATAATGGCAGATCGCTAATCAACGTCCATGTCGGGATGC |
| PA14_43950 | sucC | GCCGAGGTACAGCTCCTTGTCGATGTCGGT | TTTCGTCCGCGAGTGACCAGTGGCAGATCGCTAATAATGGCAGATCGCTAATAAGCCGAGGTACAGCTCCTTGTCGATGTCGGTAATGGCAGATCGCTAATAATGGCAGATCGCTAATCAACGTCCATGTCGGGATGC |
| PA14_43950 | sucC | TCAGCTCCCACTTCTGCGCATGGGCTTCAC | TTTCGTCCGCGAGTGACCAGTGGCAGATCGCTAATAATGGCAGATCGCTAATAATCAGCTCCCACTTCTGCGCATGGGCTTCACAATGGCAGATCGCTAATAATGGCAGATCGCTAATCAACGTCCATGTCGGGATGC |
| PA14_43950 | sucC | GATATGGGTGAACTGCTTGATCTGGTCGCC | TTTCGTCCGCGAGTGACCAGTGGCAGATCGCTAATAATGGCAGATCGCTAATAAGATATGGGTGAACTGCTTGATCTGGTCGCCAATGGCAGATCGCTAATAATGGCAGATCGCTAATCAACGTCCATGTCGGGATGC |
| PA14_43950 | sucC | ATCATGTCGCAACGCACGATGCCGCCGAAG | TTTCGTCCGCGAGTGACCAGTGGCAGATCGCTAATAATGGCAGATCGCTAATAAATCATGTCGCAACGCACGATGCCGCCGAAGAATGGCAGATCGCTAATAATGGCAGATCGCTAATCAACGTCCATGTCGGGATGC |
| PA14_43950 | sucC | GATCTTGGCATCCAGGCAGTGCAGGTTGCC | TTTCGTCCGCGAGTGACCAGTGGCAGATCGCTAATAATGGCAGATCGCTAATAAGATCTTGGCATCCAGGCAGTGCAGGTTGCCAATGGCAGATCGCTAATAATGGCAGATCGCTAATCAACGTCCATGTCGGGATGC |
| PA14_43950 | sucC | TGACGATGTCCATGGTACCCATGGCCAGGC | TTTCGTCCGCGAGTGACCAGTGGCAGATCGCTAATAATGGCAGATCGCTAATAATGACGATGTCCATGGTACCCATGGCCAGGCAATGGCAGATCGCTAATAATGGCAGATCGCTAATCAACGTCCATGTCGGGATGC |
| PA14_43950 | sucC | AGGGCCAGGTCGTAGTCCTGGAACAGCTTC | TTTCGTCCGCGAGTGACCAGTGGCAGATCGCTAATAATGGCAGATCGCTAATAAAGGGCCAGGTCGTAGTCCTGGAACAGCTTCAATGGCAGATCGCTAATAATGGCAGATCGCTAATCAACGTCCATGTCGGGATGC |
| PA14_14680 | suhB | CGGTGAATTCTTCCTGGCGCACCGGGTCGA | TTTCGTCCGCGAGTGACCAGCGCGCAGACGTTAATAACGCGCAGACGTTAATAACGGTGAATTCTTCCTGGCGCACCGGGTCGAAACGCGCAGACGTTAATAACGCGCAGACGTTAATCAACGTCCATGTCGGGATGC |
| PA14_14680 | suhB | ACCCTTCTCGAGGAATTCGTGGCTGCCGGT | TTTCGTCCGCGAGTGACCAGCGCGCAGACGTTAATAACGCGCAGACGTTAATAAACCCTTCTCGAGGAATTCGTGGCTGCCGGTAACGCGCAGACGTTAATAACGCGCAGACGTTAATCAACGTCCATGTCGGGATGC |
| PA14_14680 | suhB | CATGGATGAAGTTGGTGGTGCCGTCCAGCG | TTTCGTCCGCGAGTGACCAGCGCGCAGACGTTAATAACGCGCAGACGTTAATAACATGGATGAAGTTGGTGGTGCCGTCCAGCGAACGCGCAGACGTTAATAACGCGCAGACGTTAATCAACGTCCATGTCGGGATGC |
| PA14_14680 | suhB | AGGCCGAATTCCCAGAAGGCGTCGTAGCGT | TTTCGTCCGCGAGTGACCAGCGCGCAGACGTTAATAACGCGCAGACGTTAATAAAGGCCGAATTCCCAGAAGGCGTCGTAGCGTAACGCGCAGACGTTAATAACGCGCAGACGTTAATCAACGTCCATGTCGGGATGC |
| PA14_14680 | suhB | TCGATCACCCAGAGGTAATCGGCACCCTCG | TTTCGTCCGCGAGTGACCAGCGCGCAGACGTTAATAACGCGCAGACGTTAATAATCGATCACCCAGAGGTAATCGGCACCCTCGAACGCGCAGACGTTAATAACGCGCAGACGTTAATCAACGTCCATGTCGGGATGC |
| PA14_14680 | suhB | ACTTCGGTCACGTAGTCCTTGGCATCCTTC | TTTCGTCCGCGAGTGACCAGCGCGCAGACGTTAATAACGCGCAGACGTTAATAAACTTCGGTCACGTAGTCCTTGGCATCCTTCAACGCGCAGACGTTAATAACGCGCAGACGTTAATCAACGTCCATGTCGGGATGC |
| PA14_14680 | suhB | GTTGACCGAGATGACATCCAGGCGTTCGAT | TTTCGTCCGCGAGTGACCAGCGCGCAGACGTTAATAACGCGCAGACGTTAATAAGTTGACCGAGATGACATCCAGGCGTTCGATAACGCGCAGACGTTAATAACGCGCAGACGTTAATCAACGTCCATGTCGGGATGC |
| PA14_14680 | suhB | TGTCGATCTGGTTGTCGCGGAACGGGAACC | TTTCGTCCGCGAGTGACCAGCGCGCAGACGTTAATAACGCGCAGACGTTAATAATGTCGATCTGGTTGTCGCGGAACGGGAACCAACGCGCAGACGTTAATAACGCGCAGACGTTAATCAACGTCCATGTCGGGATGC |
| PA14_14680 | suhB | TGCAGGCGATGCTCACGGCAAAGTGCGGTA | TTTCGTCCGCGAGTGACCAGCGCGCAGACGTTAATAACGCGCAGACGTTAATAATGCAGGCGATGCTCACGGCAAAGTGCGGTAAACGCGCAGACGTTAATAACGCGCAGACGTTAATCAACGTCCATGTCGGGATGC |
| PA14_14680 | suhB | TGATCGCGTGAGTCGGATAGGCCTTGCGCA | TTTCGTCCGCGAGTGACCAGCGCGCAGACGTTAATAACGCGCAGACGTTAATAATGATCGCGTGAGTCGGATAGGCCTTGCGCAAACGCGCAGACGTTAATAACGCGCAGACGTTAATCAACGTCCATGTCGGGATGC |
| PA14_14680 | suhB | GCTGCGGAACATGTTCAGGTAGTTGTCGAG | TTTCGTCCGCGAGTGACCAGCGCGCAGACGTTAATAACGCGCAGACGTTAATAAGCTGCGGAACATGTTCAGGTAGTTGTCGAGAACGCGCAGACGTTAATAACGCGCAGACGTTAATCAACGTCCATGTCGGGATGC |
| PA14_14680 | suhB | AGCGGAAAATCAGTTCACCGGCGCTGCGAG | TTTCGTCCGCGAGTGACCAGCGCGCAGACGTTAATAACGCGCAGACGTTAATAAAGCGGAAAATCAGTTCACCGGCGCTGCGAGAACGCGCAGACGTTAATAACGCGCAGACGTTAATCAACGTCCATGTCGGGATGC |
| PA14_14680 | suhB | TTGAAGCATTTGGTGTTGCCGGCGACGATG | TTTCGTCCGCGAGTGACCAGCGCGCAGACGTTAATAACGCGCAGACGTTAATAATTGAAGCATTTGGTGTTGCCGGCGACGATGAACGCGCAGACGTTAATAACGCGCAGACGTTAATCAACGTCCATGTCGGGATGC |
| PA14_62730 | truB | ACAGGGCGGCTCGAAGGAGAGCAAATCCAG | TTTCGTCCGCGAGTGACCAGCATGATACAATCCGGAACATGATACAATCCGGAAACAGGGCGGCTCGAAGGAGAGCAAATCCAGAACATGATACAATCCGGAACATGATACAATCCGGCAACGTCCATGTCGGGATGC |
| PA14_62730 | truB | TCACTTCACCGATACCGATGAAACGACCTG | TTTCGTCCGCGAGTGACCAGCATGATACAATCCGGAACATGATACAATCCGGAATCACTTCACCGATACCGATGAAACGACCTGAACATGATACAATCCGGAACATGATACAATCCGGCAACGTCCATGTCGGGATGC |
| PA14_62730 | truB | TCCGTCTTTCTTCAGCGCCGAGTACATCGG | TTTCGTCCGCGAGTGACCAGCATGATACAATCCGGAACATGATACAATCCGGAATCCGTCTTTCTTCAGCGCCGAGTACATCGGAACATGATACAATCCGGAACATGATACAATCCGGCAACGTCCATGTCGGGATGC |
| PA14_62730 | truB | CCGTCTCGTAACCCTTGTCGGCATCGAGAA | TTTCGTCCGCGAGTGACCAGCATGATACAATCCGGAACATGATACAATCCGGAACCGTCTCGTAACCCTTGTCGGCATCGAGAAAACATGATACAATCCGGAACATGATACAATCCGGCAACGTCCATGTCGGGATGC |
| PA14_62730 | truB | ATGACCGAATCAGTCGACGCGGGGCGATAC | TTTCGTCCGCGAGTGACCAGCATGATACAATCCGGAACATGATACAATCCGGAAATGACCGAATCAGTCGACGCGGGGCGATACAACATGATACAATCCGGAACATGATACAATCCGGCAACGTCCATGTCGGGATGC |
| PA14_62730 | truB | CCTGGCCAAGATCTTCCACCAGCGTACGCA | TTTCGTCCGCGAGTGACCAGCATGATACAATCCGGAACATGATACAATCCGGAACCTGGCCAAGATCTTCCACCAGCGTACGCAAACATGATACAATCCGGAACATGATACAATCCGGCAACGTCCATGTCGGGATGC |
| PA14_62730 | truB | TGATCCTGCACCCGCAACCAGCCGAACTTC | TTTCGTCCGCGAGTGACCAGCATGATACAATCCGGAACATGATACAATCCGGAATGATCCTGCACCCGCAACCAGCCGAACTTCAACATGATACAATCCGGAACATGATACAATCCGGCAACGTCCATGTCGGGATGC |
| PA14_62730 | truB | TACCTCTCCTGCACGGGCGAGCTTGTACAG | TTTCGTCCGCGAGTGACCAGCATGATACAATCCGGAACATGATACAATCCGGAATACCTCTCCTGCACGGGCGAGCTTGTACAGAACATGATACAATCCGGAACATGATACAATCCGGCAACGTCCATGTCGGGATGC |
| PA14_62730 | truB | ATGCACGCGTTCGAGCGTCTCGAGGGTGAT | TTTCGTCCGCGAGTGACCAGCATGATACAATCCGGAACATGATACAATCCGGAAATGCACGCGTTCGAGCGTCTCGAGGGTGATAACATGATACAATCCGGAACATGATACAATCCGGCAACGTCCATGTCGGGATGC |
| PA14_62730 | truB | TAGGTGCCTTTGCTGCAGGAAACGGCCAGG | TTTCGTCCGCGAGTGACCAGCATGATACAATCCGGAACATGATACAATCCGGAATAGGTGCCTTTGCTGCAGGAAACGGCCAGGAACATGATACAATCCGGAACATGATACAATCCGGCAACGTCCATGTCGGGATGC |
| PA14_62730 | truB | TTGTCGAGCACCAGGATGCCGCTGATGGAG | TTTCGTCCGCGAGTGACCAGCATGATACAATCCGGAACATGATACAATCCGGAATTGTCGAGCACCAGGATGCCGCTGATGGAGAACATGATACAATCCGGAACATGATACAATCCGGCAACGTCCATGTCGGGATGC |
| PA14_62730 | truB | GTAATAGTAACAGAACGCGCCTCGCGCTCC | TTTCGTCCGCGAGTGACCAGCATGATACAATCCGGAACATGATACAATCCGGAAGTAATAGTAACAGAACGCGCCTCGCGCTCCAACATGATACAATCCGGAACATGATACAATCCGGCAACGTCCATGTCGGGATGC |
| PA14_62730 | truB | TGCAGTGCCTGGTTGGAACTCATGCCGCGA | TTTCGTCCGCGAGTGACCAGCATGATACAATCCGGAACATGATACAATCCGGAATGCAGTGCCTGGTTGGAACTCATGCCGCGAAACATGATACAATCCGGAACATGATACAATCCGGCAACGTCCATGTCGGGATGC |
| PA14_62730 | truB | CCAGTAGTACGCACTGTGCTCGGACAACTG | TTTCGTCCGCGAGTGACCAGCATGATACAATCCGGAACATGATACAATCCGGAACCAGTAGTACGCACTGTGCTCGGACAACTGAACATGATACAATCCGGAACATGATACAATCCGGCAACGTCCATGTCGGGATGC |
| PA14_62730 | truB | TACTGGGAGAACTTGGTCGCCTCGCCGAAG | TTTCGTCCGCGAGTGACCAGCATGATACAATCCGGAACATGATACAATCCGGAATACTGGGAGAACTTGGTCGCCTCGCCGAAGAACATGATACAATCCGGAACATGATACAATCCGGCAACGTCCATGTCGGGATGC |
| PA14_01970 | TriC | CGGTGTATTCCATGAACACGAAGACCATCG | TTTCGTCCGCGAGTGACCAGGGCGTATCGAACATTAAGGCGTATCGAACATTAACGGTGTATTCCATGAACACGAAGACCATCGAAGGCGTATCGAACATTAAGGCGTATCGAACATTCAACGTCCATGTCGGGATGC |
| PA14_01970 | TriC | GTCGGAAGACGGGAAGAACTGGTGCTGGAC | TTTCGTCCGCGAGTGACCAGGGCGTATCGAACATTAAGGCGTATCGAACATTAAGTCGGAAGACGGGAAGAACTGGTGCTGGACAAGGCGTATCGAACATTAAGGCGTATCGAACATTCAACGTCCATGTCGGGATGC |
| PA14_01970 | TriC | GTAGATGACCTCGTTCTGCGCACCGATCAT | TTTCGTCCGCGAGTGACCAGGGCGTATCGAACATTAAGGCGTATCGAACATTAAGTAGATGACCTCGTTCTGCGCACCGATCATAAGGCGTATCGAACATTAAGGCGTATCGAACATTCAACGTCCATGTCGGGATGC |
| PA14_01970 | TriC | ACCGTACCGCCCACTTCGATCCGGTAGTTC | TTTCGTCCGCGAGTGACCAGGGCGTATCGAACATTAAGGCGTATCGAACATTAAACCGTACCGCCCACTTCGATCCGGTAGTTCAAGGCGTATCGAACATTAAGGCGTATCGAACATTCAACGTCCATGTCGGGATGC |
| PA14_01970 | TriC | TCTTGATGGCGAACGAGGGATCCTCCTCGC | TTTCGTCCGCGAGTGACCAGGGCGTATCGAACATTAAGGCGTATCGAACATTAATCTTGATGGCGAACGAGGGATCCTCCTCGCAAGGCGTATCGAACATTAAGGCGTATCGAACATTCAACGTCCATGTCGGGATGC |
| PA14_01970 | TriC | TTTCCGAACGGAAGTTGCCGGAGGTGCGCA | TTTCGTCCGCGAGTGACCAGGGCGTATCGAACATTAAGGCGTATCGAACATTAATTTCCGAACGGAAGTTGCCGGAGGTGCGCAAAGGCGTATCGAACATTAAGGCGTATCGAACATTCAACGTCCATGTCGGGATGC |
| PA14_01970 | TriC | AGGATATTGCCGCCCTCCTTCATCGCCACC | TTTCGTCCGCGAGTGACCAGGGCGTATCGAACATTAAGGCGTATCGAACATTAAAGGATATTGCCGCCCTCCTTCATCGCCACCAAGGCGTATCGAACATTAAGGCGTATCGAACATTCAACGTCCATGTCGGGATGC |
| PA14_01970 | TriC | GTACCAGATATCCGGGATGTCGCCGGCCTT | TTTCGTCCGCGAGTGACCAGGGCGTATCGAACATTAAGGCGTATCGAACATTAAGTACCAGATATCCGGGATGTCGCCGGCCTTAAGGCGTATCGAACATTAAGGCGTATCGAACATTCAACGTCCATGTCGGGATGC |
| PA14_01970 | TriC | TTTCACCGAGCGGATGTCCAGGCGCACCTT | TTTCGTCCGCGAGTGACCAGGGCGTATCGAACATTAAGGCGTATCGAACATTAATTTCACCGAGCGGATGTCCAGGCGCACCTTAAGGCGTATCGAACATTAAGGCGTATCGAACATTCAACGTCCATGTCGGGATGC |
| PA14_01970 | TriC | ATCTCCAGCGGCTGCACGTAGGTGCTGATG | TTTCGTCCGCGAGTGACCAGGGCGTATCGAACATTAAGGCGTATCGAACATTAAATCTCCAGCGGCTGCACGTAGGTGCTGATGAAGGCGTATCGAACATTAAGGCGTATCGAACATTCAACGTCCATGTCGGGATGC |
| PA14_01970 | TriC | TGGTGTCCTTGAGGTAGACGAAGACCGTCG | TTTCGTCCGCGAGTGACCAGGGCGTATCGAACATTAAGGCGTATCGAACATTAATGGTGTCCTTGAGGTAGACGAAGACCGTCGAAGGCGTATCGAACATTAAGGCGTATCGAACATTCAACGTCCATGTCGGGATGC |
| PA14_01970 | TriC | ACATAGTCCTTGCGCAAGCGATCGCGCAGG | TTTCGTCCGCGAGTGACCAGGGCGTATCGAACATTAAGGCGTATCGAACATTAAACATAGTCCTTGCGCAAGCGATCGCGCAGGAAGGCGTATCGAACATTAAGGCGTATCGAACATTCAACGTCCATGTCGGGATGC |
| PA14_01970 | TriC | CATCTTCCCGGGCTCGTTCCAGTCGTAGAC | TTTCGTCCGCGAGTGACCAGGGCGTATCGAACATTAAGGCGTATCGAACATTAACATCTTCCCGGGCTCGTTCCAGTCGTAGACAAGGCGTATCGAACATTAAGGCGTATCGAACATTCAACGTCCATGTCGGGATGC |
| PA14_01970 | TriC | TTGAGGTAGGAGAACACGCCCATCACCAGC | TTTCGTCCGCGAGTGACCAGGGCGTATCGAACATTAAGGCGTATCGAACATTAATTGAGGTAGGAGAACACGCCCATCACCAGCAAGGCGTATCGAACATTAAGGCGTATCGAACATTCAACGTCCATGTCGGGATGC |
| PA14_01970 | TriC | CAGCGAGTCCAGCTCCTCGAGCTTCTTCTC | TTTCGTCCGCGAGTGACCAGGGCGTATCGAACATTAAGGCGTATCGAACATTAACAGCGAGTCCAGCTCCTCGAGCTTCTTCTCAAGGCGTATCGAACATTAAGGCGTATCGAACATTCAACGTCCATGTCGGGATGC |
| PA14_01970 | TriC | TCGCCGGGTCGCGTGTAGCTCTTCACGTAA | TTTCGTCCGCGAGTGACCAGGGCGTATCGAACATTAAGGCGTATCGAACATTAATCGCCGGGTCGCGTGTAGCTCTTCACGTAAAAGGCGTATCGAACATTAAGGCGTATCGAACATTCAACGTCCATGTCGGGATGC |
| PA14_01970 | TriC | AGTCTTCGGCAGGATGTGCACGGCGATCAC | TTTCGTCCGCGAGTGACCAGGGCGTATCGAACATTAAGGCGTATCGAACATTAAAGTCTTCGGCAGGATGTGCACGGCGATCACAAGGCGTATCGAACATTAAGGCGTATCGAACATTCAACGTCCATGTCGGGATGC |
| PA14_01970 | TriC | TCCACGGTGATCATGGCATCGTCCACCAGC | TTTCGTCCGCGAGTGACCAGGGCGTATCGAACATTAAGGCGTATCGAACATTAATCCACGGTGATCATGGCATCGTCCACCAGCAAGGCGTATCGAACATTAAGGCGTATCGAACATTCAACGTCCATGTCGGGATGC |
| PA14_01970 | TriC | TACCGCCATCAGGTACCAGACCAGGGACTG | TTTCGTCCGCGAGTGACCAGGGCGTATCGAACATTAAGGCGTATCGAACATTAATACCGCCATCAGGTACCAGACCAGGGACTGAAGGCGTATCGAACATTAAGGCGTATCGAACATTCAACGTCCATGTCGGGATGC |
| PA14_01970 | TriC | CCAGTCCGACAGGTTGAAACCGCCCTTCAT | TTTCGTCCGCGAGTGACCAGGGCGTATCGAACATTAAGGCGTATCGAACATTAACCAGTCCGACAGGTTGAAACCGCCCTTCATAAGGCGTATCGAACATTAAGGCGTATCGAACATTCAACGTCCATGTCGGGATGC |
| PA14_41440 | uspL | TTCCAGGCCTGGTTGGTCGAGACGCTGTAG | TTTCGTCCGCGAGTGACCAGATTGCACAACCGATGAAATTGCACAACCGATGAATTCCAGGCCTGGTTGGTCGAGACGCTGTAGAAATTGCACAACCGATGAAATTGCACAACCGATGCAACGTCCATGTCGGGATGC |
| PA14_41440 | uspL | AGCACCACTTCCGCGGTATTGCCGATCAGC | TTTCGTCCGCGAGTGACCAGATTGCACAACCGATGAAATTGCACAACCGATGAAAGCACCACTTCCGCGGTATTGCCGATCAGCAAATTGCACAACCGATGAAATTGCACAACCGATGCAACGTCCATGTCGGGATGC |
| PA14_41440 | uspL | TCAACACCAGCACGTCGCTTTCCAGGGTAT | TTTCGTCCGCGAGTGACCAGATTGCACAACCGATGAAATTGCACAACCGATGAATCAACACCAGCACGTCGCTTTCCAGGGTATAAATTGCACAACCGATGAAATTGCACAACCGATGCAACGTCCATGTCGGGATGC |
| PA14_41440 | uspL | TGTCGTAGGCATGGCTGATGATCCCGGCGT | TTTCGTCCGCGAGTGACCAGATTGCACAACCGATGAAATTGCACAACCGATGAATGTCGTAGGCATGGCTGATGATCCCGGCGTAAATTGCACAACCGATGAAATTGCACAACCGATGCAACGTCCATGTCGGGATGC |
| PA14_41440 | uspL | TGCTGCTTGATGATCAGGCCGCAGCCTTCG | TTTCGTCCGCGAGTGACCAGATTGCACAACCGATGAAATTGCACAACCGATGAATGCTGCTTGATGATCAGGCCGCAGCCTTCGAAATTGCACAACCGATGAAATTGCACAACCGATGCAACGTCCATGTCGGGATGC |
| PA14_41440 | uspL | TCCAGGGCCGGGCGGTCTTGGTCATCAGTA | TTTCGTCCGCGAGTGACCAGATTGCACAACCGATGAAATTGCACAACCGATGAATCCAGGGCCGGGCGGTCTTGGTCATCAGTAAAATTGCACAACCGATGAAATTGCACAACCGATGCAACGTCCATGTCGGGATGC |
| PA14_41440 | uspL | CACCACGAGGATGCTGCGAATGGCTTGCAT | TTTCGTCCGCGAGTGACCAGATTGCACAACCGATGAAATTGCACAACCGATGAACACCACGAGGATGCTGCGAATGGCTTGCATAAATTGCACAACCGATGAAATTGCACAACCGATGCAACGTCCATGTCGGGATGC |
| PA14_41440 | uspL | TTCCTCACGCAGTTCCTGGGCCAGGTCGTT | TTTCGTCCGCGAGTGACCAGATTGCACAACCGATGAAATTGCACAACCGATGAATTCCTCACGCAGTTCCTGGGCCAGGTCGTTAAATTGCACAACCGATGAAATTGCACAACCGATGCAACGTCCATGTCGGGATGC |
| PA14_41440 | uspL | TTGCTCGGCGATGATGGTCTGGTGCAGGGA | TTTCGTCCGCGAGTGACCAGATTGCACAACCGATGAAATTGCACAACCGATGAATTGCTCGGCGATGATGGTCTGGTGCAGGGAAAATTGCACAACCGATGAAATTGCACAACCGATGCAACGTCCATGTCGGGATGC |
| PA14_41440 | uspL | TTGTGCCCGTTTCAGGGCGAGACCTTCCAG | TTTCGTCCGCGAGTGACCAGATTGCACAACCGATGAAATTGCACAACCGATGAATTGTGCCCGTTTCAGGGCGAGACCTTCCAGAAATTGCACAACCGATGAAATTGCACAACCGATGCAACGTCCATGTCGGGATGC |
| PA14_41440 | uspL | TCGCTGAAACCGTACTCGGCCTGGAAGGTG | TTTCGTCCGCGAGTGACCAGATTGCACAACCGATGAAATTGCACAACCGATGAATCGCTGAAACCGTACTCGGCCTGGAAGGTGAAATTGCACAACCGATGAAATTGCACAACCGATGCAACGTCCATGTCGGGATGC |
| PA14_41440 | uspL | CAGCAGTTTCCAGTCGTCCGGGGTGAGGAT | TTTCGTCCGCGAGTGACCAGATTGCACAACCGATGAAATTGCACAACCGATGAACAGCAGTTTCCAGTCGTCCGGGGTGAGGATAAATTGCACAACCGATGAAATTGCACAACCGATGCAACGTCCATGTCGGGATGC |
| PA14_41440 | uspL | AGCTCCTCCAGGTGGGCGATGATGTCGTCC | TTTCGTCCGCGAGTGACCAGATTGCACAACCGATGAAATTGCACAACCGATGAAAGCTCCTCCAGGTGGGCGATGATGTCGTCCAAATTGCACAACCGATGAAATTGCACAACCGATGCAACGTCCATGTCGGGATGC |
| PA14_41440 | uspL | CGGCCGGACCTTCCTCGATGTGCAATTGTT | TTTCGTCCGCGAGTGACCAGATTGCACAACCGATGAAATTGCACAACCGATGAACGGCCGGACCTTCCTCGATGTGCAATTGTTAAATTGCACAACCGATGAAATTGCACAACCGATGCAACGTCCATGTCGGGATGC |
| PA14_41440 | uspL | ATGGTTTCGCTGAGCTGGAAGGTCGGGTCG | TTTCGTCCGCGAGTGACCAGATTGCACAACCGATGAAATTGCACAACCGATGAAATGGTTTCGCTGAGCTGGAAGGTCGGGTCGAAATTGCACAACCGATGAAATTGCACAACCGATGCAACGTCCATGTCGGGATGC |
| PA14_56220 | uspM | TGGCGGATCAATTGCCAGTCGCTGTTGCCC | TTTCGTCCGCGAGTGACCAGCGTACTGCAGATCGAAACGTACTGCAGATCGAAATGGCGGATCAATTGCCAGTCGCTGTTGCCCAACGTACTGCAGATCGAAACGTACTGCAGATCGACAACGTCCATGTCGGGATGC |
| PA14_56220 | uspM | TTGCAGGACCATACGGTCCAGCGGCTTGCC | TTTCGTCCGCGAGTGACCAGCGTACTGCAGATCGAAACGTACTGCAGATCGAAATTGCAGGACCATACGGTCCAGCGGCTTGCCAACGTACTGCAGATCGAAACGTACTGCAGATCGACAACGTCCATGTCGGGATGC |
| PA14_56220 | uspM | CATTTGTCGGTTTGAGCACCAGCAGGTCGC | TTTCGTCCGCGAGTGACCAGCGTACTGCAGATCGAAACGTACTGCAGATCGAAACATTTGTCGGTTTGAGCACCAGCAGGTCGCAACGTACTGCAGATCGAAACGTACTGCAGATCGACAACGTCCATGTCGGGATGC |
| PA14_56220 | uspM | AGGTGACCCGTTCGTCCAGGTAAAGGTCGC | TTTCGTCCGCGAGTGACCAGCGTACTGCAGATCGAAACGTACTGCAGATCGAAAAGGTGACCCGTTCGTCCAGGTAAAGGTCGCAACGTACTGCAGATCGAAACGTACTGCAGATCGACAACGTCCATGTCGGGATGC |
| PA14_56220 | uspM | ACGAAACGCTCGTAGCCGGCGAGCATTTCC | TTTCGTCCGCGAGTGACCAGCGTACTGCAGATCGAAACGTACTGCAGATCGAAAACGAAACGCTCGTAGCCGGCGAGCATTTCCAACGTACTGCAGATCGAAACGTACTGCAGATCGACAACGTCCATGTCGGGATGC |
| PA14_56220 | uspM | TGCAGCGAGTGACTGTCGAAGAACAGCCCG | TTTCGTCCGCGAGTGACCAGCGTACTGCAGATCGAAACGTACTGCAGATCGAAATGCAGCGAGTGACTGTCGAAGAACAGCCCGAACGTACTGCAGATCGAAACGTACTGCAGATCGACAACGTCCATGTCGGGATGC |
| PA14_56220 | uspM | ACAGGTCGATGTCGTTGGCCCGGACGAAGT | TTTCGTCCGCGAGTGACCAGCGTACTGCAGATCGAAACGTACTGCAGATCGAAAACAGGTCGATGTCGTTGGCCCGGACGAAGTAACGTACTGCAGATCGAAACGTACTGCAGATCGACAACGTCCATGTCGGGATGC |
| PA14_56220 | uspM | TGAAGTCGCACAGCAGCAACTCGACGCTGG | TTTCGTCCGCGAGTGACCAGCGTACTGCAGATCGAAACGTACTGCAGATCGAAATGAAGTCGCACAGCAGCAACTCGACGCTGGAACGTACTGCAGATCGAAACGTACTGCAGATCGACAACGTCCATGTCGGGATGC |
| PA14_56220 | uspM | TGTGCTTGCGGGTGCTCTTGAGGACCAGGT | TTTCGTCCGCGAGTGACCAGCGTACTGCAGATCGAAACGTACTGCAGATCGAAATGTGCTTGCGGGTGCTCTTGAGGACCAGGTAACGTACTGCAGATCGAAACGTACTGCAGATCGACAACGTCCATGTCGGGATGC |
| PA14_56220 | uspM | GACGACCACCAGCAGGTTATGCAGGTTCAT | TTTCGTCCGCGAGTGACCAGCGTACTGCAGATCGAAACGTACTGCAGATCGAAAGACGACCACCAGCAGGTTATGCAGGTTCATAACGTACTGCAGATCGAAACGTACTGCAGATCGACAACGTCCATGTCGGGATGC |
| PA14_56220 | uspM | CTGCTGCAGCGGCATGGAAAGCTGCTCCAG | TTTCGTCCGCGAGTGACCAGCGTACTGCAGATCGAAACGTACTGCAGATCGAAACTGCTGCAGCGGCATGGAAAGCTGCTCCAGAACGTACTGCAGATCGAAACGTACTGCAGATCGACAACGTCCATGTCGGGATGC |
| PA14_56220 | uspM | AGGCCTGGCGGTGACGTTCCTCATGTTGCA | TTTCGTCCGCGAGTGACCAGCGTACTGCAGATCGAAACGTACTGCAGATCGAAAAGGCCTGGCGGTGACGTTCCTCATGTTGCAAACGTACTGCAGATCGAAACGTACTGCAGATCGACAACGTCCATGTCGGGATGC |
| PA14_56220 | uspM | TGCGTTCGGCGATCCACTGGGCACGGGTAA | TTTCGTCCGCGAGTGACCAGCGTACTGCAGATCGAAACGTACTGCAGATCGAAATGCGTTCGGCGATCCACTGGGCACGGGTAAAACGTACTGCAGATCGAAACGTACTGCAGATCGACAACGTCCATGTCGGGATGC |
| PA14_01160 | vgrG | CTGGCTCTGTACGTATTCGCCAGGGTAGTC | TTTCGTCCGCGAGTGACCAGTACGACTCGGTCATTAATACGACTCGGTCATTAACTGGCTCTGTACGTATTCGCCAGGGTAGTCAATACGACTCGGTCATTAATACGACTCGGTCATTCAACGTCCATGTCGGGATGC |
| PA14_01160 | vgrG | GGAGGATCTTGCGTCCCTTCACCACCACGT | TTTCGTCCGCGAGTGACCAGTACGACTCGGTCATTAATACGACTCGGTCATTAAGGAGGATCTTGCGTCCCTTCACCACCACGTAATACGACTCGGTCATTAATACGACTCGGTCATTCAACGTCCATGTCGGGATGC |
| PA14_01160 | vgrG | AGAACCAGGATGTGCCGATTCCGCTCGTGG | TTTCGTCCGCGAGTGACCAGTACGACTCGGTCATTAATACGACTCGGTCATTAAAGAACCAGGATGTGCCGATTCCGCTCGTGGAATACGACTCGGTCATTAATACGACTCGGTCATTCAACGTCCATGTCGGGATGC |
| PA14_01160 | vgrG | AGTCGCCGGCATTGAGCGTGAAGCTCTTGC | TTTCGTCCGCGAGTGACCAGTACGACTCGGTCATTAATACGACTCGGTCATTAAAGTCGCCGGCATTGAGCGTGAAGCTCTTGCAATACGACTCGGTCATTAATACGACTCGGTCATTCAACGTCCATGTCGGGATGC |
| PA14_01160 | vgrG | AGTCGTAGAAATGGTCGCGCTCGCGCTGTT | TTTCGTCCGCGAGTGACCAGTACGACTCGGTCATTAATACGACTCGGTCATTAAAGTCGTAGAAATGGTCGCGCTCGCGCTGTTAATACGACTCGGTCATTAATACGACTCGGTCATTCAACGTCCATGTCGGGATGC |
| PA14_01160 | vgrG | GACCAGGCGGGTCTGTTGCGCAAGTGCCAT | TTTCGTCCGCGAGTGACCAGTACGACTCGGTCATTAATACGACTCGGTCATTAAGACCAGGCGGGTCTGTTGCGCAAGTGCCATAATACGACTCGGTCATTAATACGACTCGGTCATTCAACGTCCATGTCGGGATGC |
| PA14_01160 | vgrG | ACCCTCCATTCGCTGCAATTGCAGGACCTC | TTTCGTCCGCGAGTGACCAGTACGACTCGGTCATTAATACGACTCGGTCATTAAACCCTCCATTCGCTGCAATTGCAGGACCTCAATACGACTCGGTCATTAATACGACTCGGTCATTCAACGTCCATGTCGGGATGC |
| PA14_01160 | vgrG | TAGCCGCTGAGCTTGAACAGGTGGCCGGAA | TTTCGTCCGCGAGTGACCAGTACGACTCGGTCATTAATACGACTCGGTCATTAATAGCCGCTGAGCTTGAACAGGTGGCCGGAAAATACGACTCGGTCATTAATACGACTCGGTCATTCAACGTCCATGTCGGGATGC |
| PA14_01160 | vgrG | TGATCGTCTCGTTGTTGCCCACCGACTCGC | TTTCGTCCGCGAGTGACCAGTACGACTCGGTCATTAATACGACTCGGTCATTAATGATCGTCTCGTTGTTGCCCACCGACTCGCAATACGACTCGGTCATTAATACGACTCGGTCATTCAACGTCCATGTCGGGATGC |
| PA14_01160 | vgrG | CCAGGCTATCGTCCTTGCCCACCGAGGTAT | TTTCGTCCGCGAGTGACCAGTACGACTCGGTCATTAATACGACTCGGTCATTAACCAGGCTATCGTCCTTGCCCACCGAGGTATAATACGACTCGGTCATTAATACGACTCGGTCATTCAACGTCCATGTCGGGATGC |
| PA14_01160 | vgrG | TATGTCGATGGTCTCGTTGGCGCCCACCGA | TTTCGTCCGCGAGTGACCAGTACGACTCGGTCATTAATACGACTCGGTCATTAATATGTCGATGGTCTCGTTGGCGCCCACCGAAATACGACTCGGTCATTAATACGACTCGGTCATTCAACGTCCATGTCGGGATGC |
| PA14_01160 | vgrG | TCGATGTCCTGGTTCTTCTCGGCATGGATG | TTTCGTCCGCGAGTGACCAGTACGACTCGGTCATTAATACGACTCGGTCATTAATCGATGTCCTGGTTCTTCTCGGCATGGATGAATACGACTCGGTCATTAATACGACTCGGTCATTCAACGTCCATGTCGGGATGC |
| PA14_01160 | vgrG | TGCTGCCGTCCTTCTTCATGCGGATGCTGG | TTTCGTCCGCGAGTGACCAGTACGACTCGGTCATTAATACGACTCGGTCATTAATGCTGCCGTCCTTCTTCATGCGGATGCTGGAATACGACTCGGTCATTAATACGACTCGGTCATTCAACGTCCATGTCGGGATGC |
| PA14_01160 | vgrG | CGTCTCGTCGTAGTCGATGGTCTTGGTCCG | TTTCGTCCGCGAGTGACCAGTACGACTCGGTCATTAATACGACTCGGTCATTAACGTCTCGTCGTAGTCGATGGTCTTGGTCCGAATACGACTCGGTCATTAATACGACTCGGTCATTCAACGTCCATGTCGGGATGC |
| PA14_01160 | vgrG | GTCGATGGTGATGTTCTTGCCGCTGATGAC | TTTCGTCCGCGAGTGACCAGTACGACTCGGTCATTAATACGACTCGGTCATTAAGTCGATGGTGATGTTCTTGCCGCTGATGACAATACGACTCGGTCATTAATACGACTCGGTCATTCAACGTCCATGTCGGGATGC |
| PA14_01160 | vgrG | TTGTCGGCCTTGATGTTGATCGCCCCGGAT | TTTCGTCCGCGAGTGACCAGTACGACTCGGTCATTAATACGACTCGGTCATTAATTGTCGGCCTTGATGTTGATCGCCCCGGATAATACGACTCGGTCATTAATACGACTCGGTCATTCAACGTCCATGTCGGGATGC |
| PA14_01160 | vgrG | GATCTTCTCGTTGTTACCGACCTTCTCGGT | TTTCGTCCGCGAGTGACCAGTACGACTCGGTCATTAATACGACTCGGTCATTAAGATCTTCTCGTTGTTACCGACCTTCTCGGTAATACGACTCGGTCATTAATACGACTCGGTCATTCAACGTCCATGTCGGGATGC |
| PA14_01160 | vgrG | GCTGACGAAGTCGAAACTGGTCTCGCGGTA | TTTCGTCCGCGAGTGACCAGTACGACTCGGTCATTAATACGACTCGGTCATTAAGCTGACGAAGTCGAAACTGGTCTCGCGGTAAATACGACTCGGTCATTAATACGACTCGGTCATTCAACGTCCATGTCGGGATGC |
| PA14_01160 | vgrG | GTCGACGCCGATGGTTATGGTCTCGTTGTT | TTTCGTCCGCGAGTGACCAGTACGACTCGGTCATTAATACGACTCGGTCATTAAGTCGACGCCGATGGTTATGGTCTCGTTGTTAATACGACTCGGTCATTAATACGACTCGGTCATTCAACGTCCATGTCGGGATGC |
| PA14_01160 | vgrG | GTTTTCCGAGATCAGTTCCAGCTCGTAGGC | TTTCGTCCGCGAGTGACCAGTACGACTCGGTCATTAATACGACTCGGTCATTAAGTTTTCCGAGATCAGTTCCAGCTCGTAGGCAATACGACTCGGTCATTAATACGACTCGGTCATTCAACGTCCATGTCGGGATGC |
| PA14_08370 | vfr | CGACTTCGCATTCCACCTTGGCACGAACCC | TTTCGTCCGCGAGTGACCAGTCGTAGAGCCTAATCAATCGTAGAGCCTAATCAACGACTTCGCATTCCACCTTGGCACGAACCCAATCGTAGAGCCTAATCAATCGTAGAGCCTAATCCAACGTCCATGTCGGGATGC |
| PA14_08370 | vfr | TCCGGTTGCTGGCACAGGTCCAGCAAGGTG | TTTCGTCCGCGAGTGACCAGTCGTAGAGCCTAATCAATCGTAGAGCCTAATCAATCCGGTTGCTGGCACAGGTCCAGCAAGGTGAATCGTAGAGCCTAATCAATCGTAGAGCCTAATCCAACGTCCATGTCGGGATGC |
| PA14_08370 | vfr | GTCTAGGTGTTTGAGTTTGGGTGTGTGGGT | TTTCGTCCGCGAGTGACCAGTCGTAGAGCCTAATCAATCGTAGAGCCTAATCAAGTCTAGGTGTTTGAGTTTGGGTGTGTGGGTAATCGTAGAGCCTAATCAATCGTAGAGCCTAATCCAACGTCCATGTCGGGATGC |
| PA14_08370 | vfr | AGGCTCTTCAGCACGCGTCCGACCATTTCC | TTTCGTCCGCGAGTGACCAGTCGTAGAGCCTAATCAATCGTAGAGCCTAATCAAAGGCTCTTCAGCACGCGTCCGACCATTTCCAATCGTAGAGCCTAATCAATCGTAGAGCCTAATCCAACGTCCATGTCGGGATGC |
| PA14_08370 | vfr | AGGGTGTAGAGGATCTCCGAGTCCTGCTGC | TTTCGTCCGCGAGTGACCAGTCGTAGAGCCTAATCAATCGTAGAGCCTAATCAAAGGGTGTAGAGGATCTCCGAGTCCTGCTGCAATCGTAGAGCCTAATCAATCGTAGAGCCTAATCCAACGTCCATGTCGGGATGC |
| PA14_08370 | vfr | GAGGTAGCCGATGATCATTTCGCGGCCGTC | TTTCGTCCGCGAGTGACCAGTCGTAGAGCCTAATCAATCGTAGAGCCTAATCAAGAGGTAGCCGATGATCATTTCGCGGCCGTCAATCGTAGAGCCTAATCAATCGTAGAGCCTAATCCAACGTCCATGTCGGGATGC |
| PA14_08370 | vfr | GATGAGGATGGTGACCGAACCCTTGATGAT | TTTCGTCCGCGAGTGACCAGTCGTAGAGCCTAATCAATCGTAGAGCCTAATCAAGATGAGGATGGTGACCGAACCCTTGATGATAATCGTAGAGCCTAATCAATCGTAGAGCCTAATCCAACGTCCATGTCGGGATGC |
| PA14_08370 | vfr | AGAACAGCGTTTCGCAGCGATCGCCGGCAT | TTTCGTCCGCGAGTGACCAGTCGTAGAGCCTAATCAATCGTAGAGCCTAATCAAAGAACAGCGTTTCGCAGCGATCGCCGGCATAATCGTAGAGCCTAATCAATCGTAGAGCCTAATCCAACGTCCATGTCGGGATGC |

| | | | |
|---|---|---|---|
| PA14_08370 | vfr | TTCCTTTCACATGCACCAGGCCCTGCTCCT | TTTCGTCCGCGAGTGACCAGTCGTAGAGCCTAATCAATCGTAGAGCCTAATCAATTCCTTTCACATGCACCAGGCCCTGCTCCTAATCGTAGAGCCTAATCAATCGTAGAGCCTAATCCAACGTCCATGTCGGGATGC |
| PA14_08370 | vfr | GAACAATCCCAGCTCGCCGAAGAAATCACC | TTTCGTCCGCGAGTGACCAGTCGTAGAGCCTAATCAATCGTAGAGCCTAATCAAGAACAATCCCAGCTCGCCGAAGAAATCACCAATCGTAGAGCCTAATCAATCGTAGAGCCTAATCCAACGTCCATGTCGGGATGC |
| PA14_08370 | vfr | GCGCTGCGTTCCTGTTCGCTGCCTTCCTTT | TTTCGTCCGCGAGTGACCAGTCGTAGAGCCTAATCAATCGTAGAGCCTAATCAAGCGCTGCGTTCCTGTTCGCTGCCTTCCTTTAATCGTAGAGCCTAATCAATCGTAGAGCCTAATCCAACGTCCATGTCGGGATGC |
| PA14_08370 | vfr | CAGTTCGCGGAACTTCGCATAGCTGATCTC | TTTCGTCCGCGAGTGACCAGTCGTAGAGCCTAATCAATCGTAGAGCCTAATCAACAGTTCGCGGAACTTCGCATAGCTGATCTCAATCGTAGAGCCTAATCAATCGTAGAGCCTAATCCAACGTCCATGTCGGGATGC |
| PA14_08370 | vfr | ATGATGGTGCTCTTTGCGGTGTAGCGGCGG | TTTCGTCCGCGAGTGACCAGTCGTAGAGCCTAATCAATCGTAGAGCCTAATCAAATGATGGTGCTCTTTGCGGTGTAGCGGCGGAATCGTAGAGCCTAATCAATCGTAGAGCCTAATCCAACGTCCATGTCGGGATGC |
| PA14_08370 | vfr | TTGATCTGCATGCCGTCCGGGTGGGTCATG | TTTCGTCCGCGAGTGACCAGTCGTAGAGCCTAATCAATCGTAGAGCCTAATCAATTGATCTGCATGCCGTCCGGGTGGGTCATGAATCGTAGAGCCTAATCAATCGTAGAGCCTAATCCAACGTCCATGTCGGGATGC |
| PA14_16500 | wspR | ATGTACGAACGCGAGTGGTAGCGGATCCGC | TTTCGTCCGCGAGTGACCAGGGCGCACAACGAAATAAGGCGCACAACGAAATAAATGTACGAACGCGAGTGGTAGCGGATCCGCAAGGCGCACAACGAAATAAGGCGCACAACGAAATCAACGTCCATGTCGGGATGC |
| PA14_16500 | wspR | TCTCGATGAGAACCCGGAAGGTCTGTCCGC | TTTCGTCCGCGAGTGACCAGGGCGCACAACGAAATAAGGCGCACAACGAAATAATCTCGATGAGAACCCGGAAGGTCTGTCCGCAAGGCGCACAACGAAATAAGGCGCACAACGAAATCAACGTCCATGTCGGGATGC |
| PA14_16500 | wspR | TCTCCACCGTACGCCGGACCTTTTCCGCCA | TTTCGTCCGCGAGTGACCAGGGCGCACAACGAAATAAGGCGCACAACGAAATAATCTCCACCGTACGCCGGACCTTTTCCGCCAAAGGCGCACAACGAAATAAGGCGCACAACGAAATCAACGTCCATGTCGGGATGC |
| PA14_16500 | wspR | GAGGATCACCGTCGGCTTGATCTGGTTGGC | TTTCGTCCGCGAGTGACCAGGGCGCACAACGAAATAAGGCGCACAACGAAATAAGAGGATCACCGTCGGCTTGATCTGGTTGGCAAGGCGCACAACGAAATAAGGCGCACAACGAAATCAACGTCCATGTCGGGATGC |
| PA14_16500 | wspR | GTCGGTCTTGCTCTCATGAGGGTTGTGCAT | TTTCGTCCGCGAGTGACCAGGGCGCACAACGAAATAAGGCGCACAACGAAATAAGTCGGTCTTGCTCTCATGAGGGTTGTGCATAAGGCGCACAACGAAATAAGGCGCACAACGAAATCAACGTCCATGTCGGGATGC |
| PA14_16500 | wspR | GACGTGGCCGAAGGTGTCGTTGTAGCTCTT | TTTCGTCCGCGAGTGACCAGGGCGCACAACGAAATAAGGCGCACAACGAAATAAGACGTGGCCGAAGGTGTCGTTGTAGCTCTTAAGGCGCACAACGAAATAAGGCGCACAACGAAATCAACGTCCATGTCGGGATGC |
| PA14_16500 | wspR | TTGCGGCCATTGTTCTTGGCCTGGTAGAGC | TTTCGTCCGCGAGTGACCAGGGCGCACAACGAAATAAGGCGCACAACGAAATAATTGCGGCCATTGTTCTTGGCCTGGTAGAGCAAGGCGCACAACGAAATAAGGCGCACAACGAAATCAACGTCCATGTCGGGATGC |
| PA14_16500 | wspR | GAACAGAAGTGGAAGTCGATGCCCGCCTCG | TTTCGTCCGCGAGTGACCAGGGCGCACAACGAAATAAGGCGCACAACGAAATAAGAACAGAAGTGGAAGTCGATGCCCGCCTCGAAGGCGCACAACGAAATAAGGCGCACAACGAAATCAACGTCCATGTCGGGATGC |
| PA14_16500 | wspR | AATAGTCCACGTCGATCATCAGCAGCGACA | TTTCGTCCGCGAGTGACCAGGGCGCACAACGAAATAAGGCGCACAACGAAATAAAATAGTCCACGTCGATCATCAGCAGCGACAAAGGCGCACAACGAAATAAGGCGCACAACGAAATCAACGTCCATGTCGGGATGC |
| PA14_16500 | wspR | TTCGTCGAAGTGACGACGGTTGGAGAGCCC | TTTCGTCCGCGAGTGACCAGGGCGCACAACGAAATAAGGCGCACAACGAAATAATTCGTCGAAGTGACGACGGTTGGAGAGCCCAAGGCGCACAACGAAATAAGGCGCACAACGAAATCAACGTCCATGTCGGGATGC |
| PA14_16500 | wspR | GTCCGGCAGCTTGACCAGGTAGTCGTTGGC | TTTCGTCCGCGAGTGACCAGGGCGCACAACGAAATAAGGCGCACAACGAAATAAGTCCGGCAGCTTGACCAGGTAGTCGTTGGCAAGGCGCACAACGAAATAAGGCGCACAACGAAATCAACGTCCATGTCGGGATGC |
| PA14_16500 | wspR | TGGTCGACAGGACGATGATCGGGATGTCGC | TTTCGTCCGCGAGTGACCAGGGCGCACAACGAAATAAGGCGCACAACGAAATAATGGTCGACAGGACGATGATCGGGATGTCGCAAGGCGCACAACGAAATAAGGCGCACAACGAAATCAACGTCCATGTCGGGATGC |
| PA14_16500 | wspR | GCACCAGGTTGGTCTCCAGCAACTGCTGCT | TTTCGTCCGCGAGTGACCAGGGCGCACAACGAAATAAGGCGCACAACGAAATAAGCACCAGGTTGGTCTCCAGCAACTGCTGCTAAGGCGCACAACGAAATAAGGCGCACAACGAAATCAACGTCCATGTCGGGATGC |
| PA14_16500 | wspR | ATCGACAAGCAGTACCATGACCGCGCCGTC | TTTCGTCCGCGAGTGACCAGGGCGCACAACGAAATAAGGCGCACAACGAAATAAATCGACAAGCAGTACCATGACCGCGCCGTCAAGGCGCACAACGAAATAAGGCGCACAACGAAATCAACGTCCATGTCGGGATGC |
| PA14_16500 | wspR | CGACGCACAGCCTCTCCGATCATGGCCTGA | TTTCGTCCGCGAGTGACCAGGGCGCACAACGAAATAAGGCGCACAACGAAATAACGACGCACAGCCTCTCCGATCATGGCCTGAAAGGCGCACAACGAAATAAGGCGCACAACGAAATCAACGTCCATGTCGGGATGC |
| PA14_70470 | spoT | TGTGCATGTCGGCGAGGATGTTGGAGACCG | TTTCGTCCGCGAGTGACCAGTCGATGAATCGTCGAAATCGATGAATCGTCGAAATGTGCATGTCGGCGAGGATGTTGGAGACCGAATCGATGAATCGTCGAAATCGATGAATCGTCGACAACGTCCATGTCGGGATGC |
| PA14_70470 | spoT | TCCTCGAACTCCACGCGCATGGTGTGCATG | TTTCGTCCGCGAGTGACCAGTCGATGAATCGTCGAAATCGATGAATCGTCGAAATCCTCGAACTCCACGCGCATGGTGTGCATGAATCGATGAATCGTCGAAATCGATGAATCGTCGACAACGTCCATGTCGGGATGC |
| PA14_70470 | spoT | GAAGTTCATCTGGGTCAGCTTGCTGACCCC | TTTCGTCCGCGAGTGACCAGTCGATGAATCGTCGAAATCGATGAATCGTCGAAAGAAGTTCATCTGGGTCAGCTTGCTGACCCCAATCGATGAATCGTCGAAATCGATGAATCGTCGACAACGTCCATGTCGGGATGC |
| PA14_70470 | spoT | ATCTTCTGGAAGTTCTCGGCCTGGGCCTCG | TTTCGTCCGCGAGTGACCAGTCGATGAATCGTCGAAATCGATGAATCGTCGAAAATCTTCTGGAAGTTCTCGGCCTGGGCCTCGAATCGATGAATCGTCGAAATCGATGAATCGTCGACAACGTCCATGTCGGGATGC |
| PA14_70470 | spoT | CTTACGCTTGCCGCGCATCTTCTGGTAGAT | TTTCGTCCGCGAGTGACCAGTCGATGAATCGTCGAAATCGATGAATCGTCGAAACTTACGCTTGCCGCGCATCTTCTGGTAGATAATCGATGAATCGTCGAAATCGATGAATCGTCGACAACGTCCATGTCGGGATGC |
| PA14_70470 | spoT | ATGCAACTGTTGCCGACGTCGGTGTGCACC | TTTCGTCCGCGAGTGACCAGTCGATGAATCGTCGAAATCGATGAATCGTCGAAAATGCAACTGTTGCCGACGTCGGTGTGCACCAATCGATGAATCGTCGAAATCGATGAATCGTCGACAACGTCCATGTCGGGATGC |
| PA14_70470 | spoT | ACCCTTGGGCAGTTCCATGATGCGGCCCTT | TTTCGTCCGCGAGTGACCAGTCGATGAATCGTCGAAATCGATGAATCGTCGAAAACCCTTGGGCAGTTCCATGATGCGGCCCTTAATCGATGAATCGTCGAAATCGATGAATCGTCGACAACGTCCATGTCGGGATGC |
| PA14_70470 | spoT | AGCGACTCCTGGATCTTGCCGACGATCTCG | TTTCGTCCGCGAGTGACCAGTCGATGAATCGTCGAAATCGATGAATCGTCGAAAAGCGACTCCTGGATCTTGCCGACGATCTCGAATCGATGAATCGTCGAAATCGATGAATCGTCGACAACGTCCATGTCGGGATGC |
| PA14_70470 | spoT | TCGACAGTCTGTCGGCGAAGGCGTCTATGC | TTTCGTCCGCGAGTGACCAGTCGATGAATCGTCGAAATCGATGAATCGTCGAAATCGACAGTCTGTCGGCGAAGGCGTCTATGCAATCGATGAATCGTCGAAATCGATGAATCGTCGACAACGTCCATGTCGGGATGC |
| PA14_70470 | spoT | AGCCGTTGGCCTTGGGAATCGCGATGTAGT | TTTCGTCCGCGAGTGACCAGTCGATGAATCGTCGAAATCGATGAATCGTCGAAAAGCCGTTGGCCTTGGGAATCGCGATGTAGTAATCGATGAATCGTCGAAATCGATGAATCGTCGACAACGTCCATGTCGGGATGC |
| PA14_70470 | spoT | ATTCGCCGGTGACATCCTTGGCCCAGCTGA | TTTCGTCCGCGAGTGACCAGTCGATGAATCGTCGAAATCGATGAATCGTCGAAAATTCGCCGGTGACATCCTTGGCCCAGCTGAAATCGATGAATCGTCGAAATCGATGAATCGTCGACAACGTCCATGTCGGGATGC |
| PA14_70470 | spoT | TGCGTTCCTGGGGAATCTTCTCGAGGTGGC | TTTCGTCCGCGAGTGACCAGTCGATGAATCGTCGAAATCGATGAATCGTCGAAATGCGTTCCTGGGGAATCTTCTCGAGGTGGCAATCGATGAATCGTCGAAATCGATGAATCGTCGACAACGTCCATGTCGGGATGC |
| PA14_70470 | spoT | GTTGTGGCGTACTTCGCTGATGTTCTTGCA | TTTCGTCCGCGAGTGACCAGTCGATGAATCGTCGAAATCGATGAATCGTCGAAAGTTGTGGCGTACTTCGCTGATGTTCTTGCAAATCGATGAATCGTCGAAATCGATGAATCGTCGACAACGTCCATGTCGGGATGC |
| PA14_70470 | spoT | CAGCTTGACCAGGATCACGCGGATATCGCG | TTTCGTCCGCGAGTGACCAGTCGATGAATCGTCGAAATCGATGAATCGTCGAAACAGCTTGACCAGGATCACGCGGATATCGCGAATCGATGAATCGTCGAAATCGATGAATCGTCGACAACGTCCATGTCGGGATGC |
| PA14_70470 | spoT | ATTTCCAGGGTTTCCTTGGCGATGCGCCGG | TTTCGTCCGCGAGTGACCAGTCGATGAATCGTCGAAATCGATGAATCGTCGAAAATTTCCAGGGTTTCCTTGGCGATGCGCCGGAATCGATGAATCGTCGAAATCGATGAATCGTCGACAACGTCCATGTCGGGATGC |
| PA14_70470 | spoT | CGACCAGTTCCGAAACCGTCTCGCCGAACT | TTTCGTCCGCGAGTGACCAGTCGATGAATCGTCGAAATCGATGAATCGTCGAAACGACCAGTTCCGAAACCGTCTCGCCGAACTAATCGATGAATCGTCGAAATCGATGAATCGTCGACAACGTCCATGTCGGGATGC |
| PA14_70470 | spoT | ACAGCACCTCGAGGGTGCGCATGTTGTGCA | TTTCGTCCGCGAGTGACCAGTCGATGAATCGTCGAAATCGATGAATCGTCGAAAACAGCACCTCGAGGGTGCGCATGTTGTGCAAATCGATGAATCGTCGAAATCGATGAATCGTCGACAACGTCCATGTCGGGATGC |
| PA14_70470 | spoT | AGCACTTGGCGTAGCTCAGGACCAGGCCTT | TTTCGTCCGCGAGTGACCAGTCGATGAATCGTCGAAATCGATGAATCGTCGAAAAGCACTTGGCGTAGCTCAGGACCAGGCCTTAATCGATGAATCGTCGAAATCGATGAATCGTCGACAACGTCCATGTCGGGATGC |
| PA14_70470 | spoT | TGATCGACTCGGAGCGGCGTTGCAGCTTGA | TTTCGTCCGCGAGTGACCAGTCGATGAATCGTCGAAATCGATGAATCGTCGAAATGATCGACTCGGAGCGGCGTTGCAGCTTGAAATCGATGAATCGTCGAAATCGATGAATCGTCGACAACGTCCATGTCGGGATGC |
| PA14_70470 | spoT | TTGAAGCGTCCGGGGAACGGCTTGTACAGG | TTTCGTCCGCGAGTGACCAGTCGATGAATCGTCGAAATCGATGAATCGTCGAAATTGAAGCGTCCGGGGAACGGCTTGTACAGGAATCGATGAATCGTCGAAATCGATGAATCGTCGACAACGTCCATGTCGGGATGC |
| PA14_56780 | sodB | GTACTTCGGACGCAGGTTGCGGTAGTCGAT | TTTCGTCCGCGAGTGACCAGTGTGCGAACACTCGTAATGTGCGAACACTCGTAAGTACTTCGGACGCAGGTTGCGGTAGTCGATAATGTGCGAACACTCGTAATGTGCGAACACTCGTCAACGTCCATGTCGGGATGC |
| PA14_56780 | sodB | ATCAGGTTGTTCAGGTTCACCACGTAGGTG | TTTCGTCCGCGAGTGACCAGTGTGCGAACACTCGTAATGTGCGAACACTCGTAAATCAGGTTGTTCAGGTTCACCACGTAGGTGAATGTGCGAACACTCGTAATGTGCGAACACTCGTCAACGTCCATGTCGGGATGC |
| PA14_56780 | sodB | AGTAGAAGGTGTGGTTCCACACCTGGGCGG | TTTCGTCCGCGAGTGACCAGTGTGCGAACACTCGTAATGTGCGAACACTCGTAAAGTAGAAGGTGTGGTTCCACACCTGGGCGGAATGTGCGAACACTCGTAATGTGCGAACACTCGTCAACGTCCATGTCGGGATGC |
| PA14_56780 | sodB | TCGAGGCTCTTGCCTTCGAACTCGGTACCC | TTTCGTCCGCGAGTGACCAGTGTGCGAACACTCGTAATGTGCGAACACTCGTAATCGAGGCTCTTGCCTTCGAACTCGGTACCCAATGTGCGAACACTCGTAATGTGCGAACACTCGTCAACGTCCATGTCGGGATGC |
| PA14_56780 | sodB | CTGCGGAAATGTGCGGCTCAAGGGCGTTCT | TTTCGTCCGCGAGTGACCAGTGTGCGAACACTCGTAATGTGCGAACACTCGTAACTGCGGAAATGTGCGGCTCAAGGGCGTTCTAATGTGCGAACACTCGTAATGTGCGAACACTCGTCAACGTCCATGTCGGGATGC |
| PA14_56780 | sodB | AGATGCCGCCGGAGGAGCTCTTGACGATCT | TTTCGTCCGCGAGTGACCAGTGTGCGAACACTCGTAATGTGCGAACACTCGTAAAGATGCCGCCGGAGGAGCTCTTGACGATCTAATGTGCGAACACTCGTAATGTGCGAACACTCGTCAACGTCCATGTCGGGATGC |
| PA14_56780 | sodB | AGTAGGCGTGTTCCCAGACGTCGCAGGTCA | TTTCGTCCGCGAGTGACCAGTGTGCGAACACTCGTAATGTGCGAACACTCGTAAAGTAGGCGTGTTCCCAGACGTCGCAGGTCAAATGTGCGAACACTCGTAATGTGCGAACACTCGTCAACGTCCATGTCGGGATGC |
| PA14_56780 | sodB | GTGGTGCTTGTCGTGGTGGTATTCCAGGGT | TTTCGTCCGCGAGTGACCAGTGTGCGAACACTCGTAATGTGCGAACACTCGTAAGTGGTGCTTGTCGTGGTGGTATTCCAGGGTAATGTGCGAACACTCGTAATGTGCGAACACTCGTCAACGTCCATGTCGGGATGC |
| PA14_56780 | sodB | CGACGGAAGTCTTGGTGAACTCTTCCTTGA | TTTCGTCCGCGAGTGACCAGTGTGCGAACACTCGTAATGTGCGAACACTCGTAACGACGGAAGTCTTGGTGAACTCTTCCTTGAAATGTGCGAACACTCGTAATGTGCGAACACTCGTCAACGTCCATGTCGGGATGC |
| PA14_56780 | sodB | TTGTCGAAGGAGCCGAAGGCGGCGTTGATG | TTTCGTCCGCGAGTGACCAGTGTGCGAACACTCGTAATGTGCGAACACTCGTAATTGTCGAAGGAGCCGAAGGCGGCGTTGATGAATGTGCGAACACTCGTAATGTGCGAACACTCGTCAACGTCCATGTCGGGATGC |
| PA14_56780 | sodB | TCCCAGTTGACCAGGTTCCAGAACGCCTCG | TTTCGTCCGCGAGTGACCAGTGTGCGAACACTCGTAATGTGCGAACACTCGTAATCCCAGTTGACCAGGTTCCAGAACGCCTCGAATGTGCGAACACTCGTAATGTGCGAACACTCGTCAACGTCCATGTCGGGATGC |
| PA14_56780 | sodB | TCGTAAGGCAGCGGCGGCAATTCGAAAGCC | TTTCGTCCGCGAGTGACCAGTGTGCGAACACTCGTAATGTGCGAACACTCGTAATCGTAAGGCAGCGGCGGCAATTCGAAAGCCAATGTGCGAACACTCGTAATGTGCGAACACTCGTCAACGTCCATGTCGGGATGC |
| PA14_58730 | pilA | GGCTTGAGCATTCCAGACCCTTGCCGAAGC | TTTCGTCCGCGAGTGACCAGAACGATCGCGGTGATAAAACGATCGCGGTGATAAGGCTTGAGCATTCCAGACCCTTGCCGAAGCAAAACGATCGCGGTGATAAAACGATCGCGGTGATCAACGTCCATGTCGGGATGC |
| PA14_58730 | pilA | TTAGCGGCATTCGCTCGGAGCGAAACGAGC | TTTCGTCCGCGAGTGACCAGAACGATCGCGGTGATAAAACGATCGCGGTGATAATTAGCGGCATTCGCTCGGAGCGAAACGAGCAAAACGATCGCGGTGATAAAACGATCGCGGTGATCAACGTCCATGTCGGGATGC |
| PA14_58730 | pilA | CCGGCAGCATTCTGCACATAGGGAGTAAAT | TTTCGTCCGCGAGTGACCAGAACGATCGCGGTGATAAAACGATCGCGGTGATAACCGGCAGCATTCTGCACATAGGGAGTAAATAAAACGATCGCGGTGATAAAACGATCGCGGTGATCAACGTCCATGTCGGGATGC |
| PA14_58730 | pilA | GCCGGAATATTACCCACGTTTGTGGCATTG | TTTCGTCCGCGAGTGACCAGAACGATCGCGGTGATAAAACGATCGCGGTGATAAGCCGGAATATTACCCACGTTTGTGGCATTGAAAACGATCGCGGTGATAAAACGATCGCGGTGATCAACGTCCATGTCGGGATGC |
| PA14_58730 | pilA | ATATGCCGGCAAAGCGACGGCAGCAAGAAT | TTTCGTCCGCGAGTGACCAGAACGATCGCGGTGATAAAACGATCGCGGTGATAAATATGCCGGCAAAGCGACGGCAGCAAGAATAAAACGATCGCGGTGATAAAACGATCGCGGTGATCAACGTCCATGTCGGGATGC |
| PA14_58730 | pilA | GGACCACTGGAAACCGCATTGGATTCCGAT | TTTCGTCCGCGAGTGACCAGAACGATCGCGGTGATAAAACGATCGCGGTGATAAGGACCACTGGAAACCGCATTGGATTCCGATAAAACGATCGCGGTGATAAAACGATCGCGGTGATCAACGTCCATGTCGGGATGC |
| PA14_58730 | pilA | ACAACCCCAGTCAATAGAGCCAGTCACACC | TTTCGTCCGCGAGTGACCAGAACGATCGCGGTGATAAAACGATCGCGGTGATAAACAACCCCAGTCAATAGAGCCAGTCACACCAAAACGATCGCGGTGATAAAACGATCGCGGTGATCAACGTCCATGTCGGGATGC |
| PA14_58730 | pilA | TTTCACCAGTCGCCTCTGCAATTCGCACAG | TTTCGTCCGCGAGTGACCAGAACGATCGCGGTGATAAAACGATCGCGGTGATAATTTCACCAGTCGCCTCTGCAATTCGCACAGAAAACGATCGCGGTGATAAAACGATCGCGGTGATCAACGTCCATGTCGGGATGC |
| PA14_58730 | pilA | TCTGTCACGCGAGCACGAATGGTGTAATCC | TTTCGTCCGCGAGTGACCAGAACGATCGCGGTGATAAAACGATCGCGGTGATAATCTGTCACGCGAGCACGAATGGTGTAATCCAAAACGATCGCGGTGATAAAACGATCGCGGTGATCAACGTCCATGTCGGGATGC |
| PA14_58730 | pilA | TAGCTCACCGGCAGTGGCAGAGCTATCGCC | TTTCGTCCGCGAGTGACCAGAACGATCGCGGTGATAAAACGATCGCGGTGATAATAGCTCACCGGCAGTGGCAGAGCTATCGCCAAAACGATCGCGGTGATAAAACGATCGCGGTGATCAACGTCCATGTCGGGATGC |
| PA14_58730 | pilA | GTCACATACTTACTGGTAGCACCGGCGTTA | TTTCGTCCGCGAGTGACCAGAACGATCGCGGTGATAAAACGATCGCGGTGATAAGTCACATACTTACTGGTAGCACCGGCGTTAAAAACGATCGCGGTGATAAAACGATCGCGGTGATCAACGTCCATGTCGGGATGC |
| PA14_58730 | pilA | TACCTGCAGTCAGGGCAGGCATATTGCGGT | TTTCGTCCGCGAGTGACCAGAACGATCGCGGTGATAAAACGATCGCGGTGATAATACCTGCAGTCAGGGCAGGCATATTGCGGTAAAACGATCGCGGTGATAAAACGATCGCGGTGATCAACGTCCATGTCGGGATGC |
| PA14_58730 | pilA | CGATGATCGCAACCACGATCATCAGTTCGA | TTTCGTCCGCGAGTGACCAGAACGATCGCGGTGATAAAACGATCGCGGTGATAACGATGATCGCAACCACGATCATCAGTTCGAAAAACGATCGCGGTGATAAAACGATCGCGGTGATCAACGTCCATGTCGGGATGC |
| PA14_58730 | pilA | CAGCATAACTGGCACCCAATTGAGTCGGGG | TTTCGTCCGCGAGTGACCAGAACGATCGCGGTGATAAAACGATCGCGGTGATAACAGCATAACTGGCACCCAATTGAGTCGGGGAAAACGATCGCGGTGATAAAACGATCGCGGTGATCAACGTCCATGTCGGGATGC |
| PA14_58730 | pilA | TAAGCGTCTTGGCGCTGGCAGCCAGACCAA | TTTCGTCCGCGAGTGACCAGAACGATCGCGGTGATAAAACGATCGCGGTGATAATAAGCGTCTTGGCGCTGGCAGCCAGACCAAAAAACGATCGCGGTGATAAAACGATCGCGGTGATCAACGTCCATGTCGGGATGC |
| PA14_16250 | lasB | TACTTGCCGATCTTCTGGTTGCCACCGGGG | TTTCGTCCGCGAGTGACCAGATAGAATATCGGCCGAAATAGAATATCGGCCGAATACTTGCCGATCTTCTGGTTGCCACCGGGGAAATAGAATATCGGCCGAAATAGAATATCGGCCGCAACGTCCATGTCGGGATGC |
| PA14_16250 | lasB | TACACGCCGCTGGAGTGGTGCACGTCGATA | TTTCGTCCGCGAGTGACCAGATAGAATATCGGCCGAAATAGAATATCGGCCGAATACACGCCGCTGGAGTGGTGCACGTCGATAAAATAGAATATCGGCCGAAATAGAATATCGGCCGCAACGTCCATGTCGGGATGC |
| PA14_16250 | lasB | TTGGGCGATGTTGTTCTCGCCCAGGCGGAT | TTTCGTCCGCGAGTGACCAGATAGAATATCGGCCGAAATAGAATATCGGCCGAATTGGGCGATGTTGTTCTCGCCCAGGCGGATAAATAGAATATCGGCCGAAATAGAATATCGGCCGCAACGTCCATGTCGGGATGC |
| PA14_16250 | lasB | AGCTAATCACTCCACAGGCACCGCTGTTGT | TTTCGTCCGCGAGTGACCAGATAGAATATCGGCCGAAATAGAATATCGGCCGAAAGCTAATCACTCCACAGGCACCGCTGTTGTAAATAGAATATCGGCCGAAATAGAATATCGGCCGCAACGTCCATGTCGGGATGC |
| PA14_16250 | lasB | TGCTGCGGATCGCTTTCAGTTCGTCGGCAC | TTTCGTCCGCGAGTGACCAGATAGAATATCGGCCGAAATAGAATATCGGCCGAATGCTGCGGATCGCTTTCAGTTCGTCGGCACAAATAGAATATCGGCCGAAATAGAATATCGGCCGCAACGTCCATGTCGGGATGC |
| PA14_16250 | lasB | TTGACTTCGGTGATGGCTTCGCCGACCACC | TTTCGTCCGCGAGTGACCAGATAGAATATCGGCCGAAATAGAATATCGGCCGAATTGACTTCGGTGATGGCTTCGCCGACCACCAAATAGAATATCGGCCGAAATAGAATATCGGCCGCAACGTCCATGTCGGGATGC |
| PA14_16250 | lasB | CGTCGTTCAGTGGCGAATAAGCGCCGTTGA | TTTCGTCCGCGAGTGACCAGATAGAATATCGGCCGAAATAGAATATCGGCCGAACGTCGTTCAGTGGCGAATAAGCGCCGTTGAAAATAGAATATCGGCCGAAATAGAATATCGGCCGCAACGTCCATGTCGGGATGC |
| PA14_16250 | lasB | TGACGTCAGCCGCCGAGTAGTTGCGGTTCT | TTTCGTCCGCGAGTGACCAGATAGAATATCGGCCGAAATAGAATATCGGCCGAATGACGTCAGCCGCCGAGTAGTTGCGGTTCTAAATAGAATATCGGCCGAAATAGAATATCGGCCGCAACGTCCATGTCGGGATGC |
| PA14_16250 | lasB | TGCTTGTAGGTGTTGGTCGGGCAGGCGAAG | TTTCGTCCGCGAGTGACCAGATAGAATATCGGCCGAAATAGAATATCGGCCGAATGCTTGTAGGTGTTGGTCGGGCAGGCGAAGAAATAGAATATCGGCCGAAATAGAATATCGGCCGCAACGTCCATGTCGGGATGC |
| PA14_16250 | lasB | TTTGTCATTCTCGGTCTTGCGGCCCTGGGC | TTTCGTCCGCGAGTGACCAGATAGAATATCGGCCGAAATAGAATATCGGCCGAATTTGTCATTCTCGGTCTTGCGGCCCTGGGCAAATAGAATATCGGCCGAAATAGAATATCGGCCGCAACGTCCATGTCGGGATGC |
| PA14_16250 | lasB | ACACCAGCGGATAGAACATGGTGGCGCCGT | TTTCGTCCGCGAGTGACCAGATAGAATATCGGCCGAAATAGAATATCGGCCGAAACACCAGCGGATAGAACATGGTGGCGCCGTAAATAGAATATCGGCCGAAATAGAATATCGGCCGCAACGTCCATGTCGGGATGC |
| PA14_16250 | lasB | ACCGGTCTTGGCGTCGATGACGAAATGCGG | TTTCGTCCGCGAGTGACCAGATAGAATATCGGCCGAAATAGAATATCGGCCGAAACCGGTCTTGGCGTCGATGACGAAATGCGGAAATAGAATATCGGCCGAAATAGAATATCGGCCGCAACGTCCATGTCGGGATGC |
| PA14_16250 | lasB | TTGCTCGTAGCGGGTGACCTGCTTGCCGTT | TTTCGTCCGCGAGTGACCAGATAGAATATCGGCCGAAATAGAATATCGGCCGAATTGCTCGTAGCGGGTGACCTGCTTGCCGTTAAATAGAATATCGGCCGAAATAGAATATCGGCCGCAACGTCCATGTCGGGATGC |
| PA14_16250 | lasB | ACCAGTCCCGGTACAGGTTGAACACCACGC | TTTCGTCCGCGAGTGACCAGATAGAATATCGGCCGAAATAGAATATCGGCCGAAACCAGTCCCGGTACAGGTTGAACACCACGCAAATAGAATATCGGCCGAAATAGAATATCGGCCGCAACGTCCATGTCGGGATGC |
| PA14_16250 | lasB | AGTTCTGCTCGGTGAAGCCGTGGCTGACCT | TTTCGTCCGCGAGTGACCAGATAGAATATCGGCCGAAATAGAATATCGGCCGAAAGTTCTGCTCGGTGAAGCCGTGGCTGACCTAAATAGAATATCGGCCGAAATAGAATATCGGCCGCAACGTCCATGTCGGGATGC |
| PA14_16250 | lasB | TCATTCCGCCGGATTGCCCGCGGTAGATCA | TTTCGTCCGCGAGTGACCAGATAGAATATCGGCCGAAATAGAATATCGGCCGAATCATTCCGCCGGATTGCCCGCGGTAGATCAAAATAGAATATCGGCCGAAATAGAATATCGGCCGCAACGTCCATGTCGGGATGC |
| PA14_16250 | lasB | GCAACGAACAACAGGTCAAGCGTAGAAACC | TTTCGTCCGCGAGTGACCAGATAGAATATCGGCCGAAATAGAATATCGGCCGAAGCAACGAACAACAGGTCAAGCGTAGAAACCAAATAGAATATCGGCCGAAATAGAATATCGGCCGCAACGTCCATGTCGGGATGC |
| PA14_16250 | lasB | TTGCTGGTGGCGGTCCAGTAGTAGCGGTTG | TTTCGTCCGCGAGTGACCAGATAGAATATCGGCCGAAATAGAATATCGGCCGAATTGCTGGTGGCGGTCCAGTAGTAGCGGTTGAAATAGAATATCGGCCGAAATAGAATATCGGCCGCAACGTCCATGTCGGGATGC |
| PA14_16250 | lasB | CCGGGAATCAGGTAGGAGACGTTGTAGACC | TTTCGTCCGCGAGTGACCAGATAGAATATCGGCCGAAATAGAATATCGGCCGAACCGGGAATCAGGTAGGAGACGTTGTAGACCAAATAGAATATCGGCCGAAATAGAATATCGGCCGCAACGTCCATGTCGGGATGC |
| PA14_16250 | lasB | TGGACACGTCGATCAGGTCGGCGGCAAAAG | TTTCGTCCGCGAGTGACCAGATAGAATATCGGCCGAAATAGAATATCGGCCGAATGGACACGTCGATCAGGTCGGCGGCAAAAGAAATAGAATATCGGCCGAAATAGAATATCGGCCGCAACGTCCATGTCGGGATGC |
