## Supplemental Table 3 for "In situ single-cell activities of microbial populations revealed by spatial transcriptomics"

**Table S3: 16S rRNA probes**

| Probe_id | 16S pos | 16S seq | Full probe sequence |
|---|---|---|---|
| Ribo-Tag_1 | 771-799 | GCGTGGACTACCAGGGTATCTAATCCTG | CGATTAGTCGTCACTAAGCGTGGACTACCAGGGTATCTAATCCTGAAACTCCGAATGCTACG |
| Ribo-Tag_2 | 771-799 | GCGTGGACTACCAGGGTATCTAATCCTG | CGATTAGTCGTCACTAAGCGTGGACTACCAGGGTATCTAATCCTGAAGGTTACACGCGACTA |
| Ribo-Tag_3 | 771-799 | GCGTGGACTACCAGGGTATCTAATCCTG | ACTCCGAATGCTACGAAGCGTGGACTACCAGGGTATCTAATCCTGAAGGTTACACGCGACTA |
| Ribo-Tag_4 | 771-799 | GCGTGGACTACCAGGGTATCTAATCCTG | ATGTAACCAAGCGTCAAGCGTGGACTACCAGGGTATCTAATCCTGAATCAGTTACCGGTGTA |
| Ribo-Tag_5 | 771-799 | GCGTGGACTACCAGGGTATCTAATCCTG | ATGTAACCAAGCGTCAAGCGTGGACTACCAGGGTATCTAATCCTGAATCCAGCTTACGTTCG |
| Ribo-Tag_6 | 771-799 | GCGTGGACTACCAGGGTATCTAATCCTG | TCAGTTACCGGTGTAAAGCGTGGACTACCAGGGTATCTAATCCTGAATCCAGCTTACGTTCG |
| Ribo-Tag_7 | 771-799 | GCGTGGACTACCAGGGTATCTAATCCTG | CGATTAGTCGTCACTAAGCGTGGACTACCAGGGTATCTAATCCTGAATCAGTTACCGGTGTA |
| Ribo-Tag_8 | 771-799 | GCGTGGACTACCAGGGTATCTAATCCTG | CGATTAGTCGTCACTAAGCGTGGACTACCAGGGTATCTAATCCTGAATCCAGCTTACGTTCG |
| Ribo-Tag_9 | 771-799 | GCGTGGACTACCAGGGTATCTAATCCTG | ATGTAACCAAGCGTCAAGCGTGGACTACCAGGGTATCTAATCCTGAAACTCCGAATGCTACG |
| Ribo-Tag_10 | 771-799 | GCGTGGACTACCAGGGTATCTAATCCTG | ATGTAACCAAGCGTCAAGCGTGGACTACCAGGGTATCTAATCCTGAAGGTTACACGCGACTA |
| Ribo-Tag_11 | 771-799 | GCGTGGACTACCAGGGTATCTAATCCTG | ACTCCGAATGCTACGAAGCGTGGACTACCAGGGTATCTAATCCTGAATCCAGCTTACGTTCG |
| Ribo-Tag_12 | 771-799 | GCGTGGACTACCAGGGTATCTAATCCTG | TCAGTTACCGGTGTAAAGCGTGGACTACCAGGGTATCTAATCCTGAAGGTTACACGCGACTA |
| Reference (all samples) | 714-743 | CAGTGTCAGTATCAGTCCAGGTGGTCGCCT | TATGTGACTACGCACAACGTCGTATCCAGTATAACAGTGTCAGTATCAGTCCAGGTGGTCGCCTAACTATTATCGCCGAGA |
