## Supplemental Table 4 for "In situ single-cell activities of microbial populations revealed by spatial transcriptomics"

**Table S4 - UMAP cluster frequency (%) per LB growth condition**

| Cluster | OD = 0.03 (lag_0.5h) | OD = 0.03 (lag_1h) | OD = 0.06 | OD_0.2 | OD_0.45 | OD_0.85 | OD_1.2 | OD_1.5 | OD_1.8 | OD_2.1 | OD_3.2 |
|---|---|---|---|---|---|---|---|---|---|---|---|
| 1 | 6.71% | 29.94% | 45.89% | 56.18% | 74.52% | 2.31% | 0.05% | 0.00% | 0.02% | 0.00% | 0.00% |
| 2 | 0.04% | 0.02% | 0.00% | 0.09% | 0.03% | 0.27% | 88.59% | 17.43% | 0.82% | 0.87% | 0.75% |
| 3 | 53.44% | 47.55% | 12.96% | 3.52% | 6.33% | 1.93% | 0.02% | 0.02% | 0.21% | 0.19% | 0.04% |
| 4 | 0.00% | 0.00% | 0.00% | 0.00% | 0.07% | 0.15% | 2.38% | 31.72% | 19.97% | 15.96% | 7.10% |
| 5 | 0.22% | 0.13% | 0.03% | 0.00% | 0.00% | 0.08% | 0.86% | 8.23% | 17.19% | 20.77% | 21.23% |
| 6 | 0.04% | 0.02% | 0.09% | 0.05% | 2.80% | 92.22% | 0.66% | 0.44% | 0.15% | 0.05% | 0.04% |
| 7 | 0.13% | 0.04% | 0.00% | 0.00% | 0.17% | 0.50% | 1.45% | 9.83% | 12.33% | 13.91% | 9.46% |
| 8 | 0.61% | 8.05% | 30.30% | 28.34% | 9.33% | 0.31% | 0.00% | 0.02% | 0.02% | 0.03% | 0.00% |
| 9 | 0.17% | 0.11% | 0.03% | 0.00% | 0.00% | 0.12% | 0.52% | 4.76% | 9.74% | 9.32% | 9.94% |
| 10 | 0.04% | 0.16% | 0.03% | 0.05% | 0.17% | 0.27% | 0.70% | 4.78% | 8.37% | 10.13% | 7.47% |
| 11 | 0.35% | 0.40% | 0.18% | 0.27% | 0.31% | 0.69% | 1.31% | 4.49% | 7.71% | 8.61% | 7.08% |
| 12 | 0.09% | 0.09% | 0.00% | 0.14% | 0.35% | 0.08% | 0.38% | 0.46% | 1.35% | 1.50% | 23.30% |
| 13 | 34.56% | 5.03% | 1.80% | 1.85% | 1.73% | 0.12% | 0.18% | 0.53% | 0.71% | 0.65% | 0.42% |
| 14 | 0.04% | 0.04% | 0.03% | 0.00% | 0.03% | 0.23% | 0.88% | 5.10% | 6.82% | 6.53% | 3.78% |
| 15 | 0.13% | 0.04% | 0.00% | 0.00% | 0.03% | 0.27% | 1.04% | 5.80% | 6.55% | 4.67% | 3.82% |
| 16 | 2.90% | 8.09% | 8.19% | 8.94% | 2.73% | 0.12% | 0.00% | 0.00% | 0.00% | 0.02% | 0.00% |
| 17 | 0.17% | 0.04% | 0.12% | 0.00% | 0.07% | 0.04% | 0.23% | 2.04% | 3.16% | 2.73% | 3.76% |
| 18 | 0.04% | 0.02% | 0.03% | 0.00% | 0.07% | 0.08% | 0.29% | 2.77% | 2.74% | 2.15% | 0.39% |
| 19 | 0.09% | 0.02% | 0.00% | 0.00% | 0.00% | 0.12% | 0.29% | 0.73% | 1.42% | 1.47% | 0.62% |
| 20 | 0.22% | 0.18% | 0.31% | 0.59% | 1.24% | 0.12% | 0.16% | 0.85% | 0.74% | 0.44% | 0.81% |
